## Supplementary Figures and Tables for "TROP2 represents a negative prognostic factor in colorectal adenocarcinoma and its expression is associated with features of epithelial-mesenchymal transition and invasiveness"

#### Supplementary Information

##### Supplementary Figure and Table Legends

###### Supplementary Figure S1

Tissue microarray (TMA) analysis of TROP2 expression in patients with CRC. (A) Left, an example of a TMA with 20 2-mm-diameter cylindrical samples from 10 cases; each case is represented by two samples taken from random locations on the paraffin block. Right, representative images of TROP2 immunostaining; score 0: TROP2 negative; score 4: weak TROP2 positivity, score 9: strong "patchy" positivity; score 12: strong uniform positivity in tumor tissue. Scale bar: 50 µm. (B) Categorization of TROP2 staining by total immunostaining score (TIS) and binarization into low/medium and high expression groups.

###### Supplementary Figure S2

Kaplan-Meier survival data for groups with low (score 0-4), intermediate (score 6-8), and high (score 9-12) TROP2 expression, censored at 5 years of follow-up, document a significant survival difference between patients with low/intermediate and high TROP2 expression ($p = 0.00073$).

###### Supplementary Figure S3

Bar graphs of univariate logistic regression show a positive significant correlation of high TROP2 expression (score 9-12) with high grading ($p<0.001$), mucinous and signet ring cell morphology ($p=0.025$), lymph node metastasis ($p=0.012$), advanced stage according to Union for International Cancer Control staging (UICC stage) III+IV ($p=0.047$), localization of primary CRC in the right colon ($p=0.026$), positive PD-L1≥1% ($p=0.018$), CK7 staining ≥10% ($p=0.027$). High TROP2 expression was inversely correlated with SATB2 staining >40% ($p<0.001$).

###### Supplementary Figure S4

(A) Principal component analysis (PCA) based on the 500 most variably expressed genes between $TROP2^{high}$ and $TROP2^{low}$ human colon tumor cells. Red dots represent $TROP2^{high}$ cells, blue dots represent $TROP2^{low}$ cells. (B) The graph shows the normalized number of *TROP2* mRNA reads detected in $TROP2^{high}$ and $TROP2^{low}$ human CRC cells. The boxes correspond to the first and third quartiles; the median of the relative expression values is shown as a blue line. The range of values is indicated by "whiskers" above and below each box; S12, S4, S5, and S7 refer to individual patients; colored lines connect $TROP2^{high}$ and $TROP2^{low}$ samples from the same patient.

###### Supplementary Figure S5

Trop2 protein and mRNA are increased in small intestine and colon tumors of $Apc^{Min/+}$ mice. (A) Immunohistochemical detection of Trop2 protein in adenomas that developed in the small intestine and colon; the framed areas are shown at higher magnification in the images below. Membranous and cytoplasmic Trop2 localization is clearly visible throughout tumor tissue. Scale bar: 100 µm (top images); 20 µm (bottom images). (B) Quantitative RT-PCR analysis of *Trop2* expression along the rostro-caudal axis of the intestine. Total RNA was isolated from adenomas that had developed in different regions of the small or large intestine as indicated; healthy tissue was isolated from the same intestinal regions of Apc WT littermates. Two samples, i.e. biological replicates, were obtained for each type of tissue specimen. Trop2 expression was assessed in technical triplicates for each sample and normalized to the *Gapdh* expression levels. The graph shows the expression of *Trop* relative to its average expression in all samples from healthy intestine. The individual dots then represent the average value in the triplicates analyzed. Black bars represent the mean of two biological replicates.

**Supplementary Figure S6**

TROP2-deficient SW480 cells show increased migration. (A) Western blot analysis of whole cell lysates prepared from sorted single cell clones of DLD1 (left) and SW480 (right) human CRC cells edited with the TROP2-specific CRISPR/Cas9 system; edited clones are labeled "KO". Clones labeled "WT" were treated in the same way as KO clones except that they were transfected with a construct that did not produce TROP2-specific sgRNA, but the vector used was "empty." Results for three KO and three WT clones for each cell line are shown along with immunoblotting of the lysate of parental cells. CTBP, loading control. (B) Left, cell viability assay showed no difference in viability/proliferation between DLD1 KO/WT (n=6/5) and SW480 KO/WT (n=5/7) clones. Viability is indicated by the relative fluorescence intensity of the metabolic product of alamarBlue; the intensity at day 0 was set to 1. The experiment was performed in technical triplicates for each cell clone; mean values of relative fluorescence intensity are plotted; error bars indicate standard deviations (SD). Right, tumor formation after xenotransplantation of SW480 KO (n=3) and SW480 WT (n=3) cell clones. Immunodeficient mice of the NSG strain were injected subcutaneously with $10^6$ cells into the lumbar region. The mice were sacrificed 28 days later, and the growing tumors were resected and weighed. The experiment was performed in triplicate for each cell clone. The graph shows the weight of each tumor; the mean is indicated by the black line; no significance level (ns) was reached for the difference in weight between tumors derived from WT and KO cells. (C) Wound healing assay showed increased migration of SW480 KO cells compared to WT clones. The graph shows the percentage of "healed" area 24 hours after wound formation. The experiment was performed with five KO and five WT cell clones for each cell line; six positions were photographed for each clone and time point. The median is indicated by the black line. The significance level ($p < 0.05$) was not reached for the percentage of healed area for DLD1 cells; ****, $p < 0.0001$.

**Supplementary Table S1**

Patients and histopathological characteristics of clinical samples. Hyp, hyperplastic polyps; LGD, adenomas with low-grade dysplasia; HGD, adenomas with high-grade dysplasia; CRC, invasive carcinoma.

**Supplementary Table S2**

Oligonucleotides and primers used in the study

**Supplementary Table S3**

Patient characteristics and correlation of TROP2 expression scores with clinicopathological parameters in CRC patients. Gene abbreviations: CK7, cytokeratin 7; CK20, cytokeratin 20; PD-L1, programmed death ligand; SATB2, special AT-rich sequence binding protein 2. Other abbreviations: CI, confidence interval; NOS, not otherwise specified; SD, standard deviation; UICC, Union for International Cancer Control.

**Supplementary Table S4**

Differentially expressed genes in Trop2$^{high}$ compared to Trop2$^{low}$ tumor cells. Genes with significantly altered expression in hyperplastic epithelium 7 days and in adenomas 6 weeks after Apc inactivation are listed on the appropriate sheet according to whether they were upregulated or downregulated. Genes with an adjusted p-value < 0.05 in both experiments were considered significant; pseudogenes were removed from the lists.

**Supplementary Table S5**

Differentially expressed genes (DEGs) in two populations of RFP-positive, i.e. proliferating cells sorted for Trop2 expression, and in a population of RFP-negative, i.e. differentiated cells sorted from hyperplastic epithelium 7 days after Apc inactivation. Genes with an adjusted p-value < 0.05 and $|\log_2 FC| \geq 1$ were considered significant. DEGs between Trop2$^+$RFP$^+$ and Trop2$^-$RFP$^+$ cells and between the two RFP$^+$ cell-derived gene expression profiles and the RFP$^-$ cell dataset are shown on the corresponding sheet.

**Supplementary Table S6**

Differentially expressed genes (DEGs) in two populations of tdTomato-positive tumor cells sorted for Trop2 expression and in a population of tdTomato-negative (differentiated) cells obtained from microroadenomas 6 weeks after Apc inactivation. DEGs between Trop2$^+$ tdTomato$^+$ and Trop2$^-$ tdTomato$^+$ cells and between the two tdTomato-positive cell groups and tdTomato-negative cells are shown on the corresponding sheets. Genes with an adjusted p-value < 0.05 and $|\log_2 FC| \geq 1$ were considered significant. Wnt-responsive genes are highlighted in red.

**Supplementary Table S7**

Overlap of differentially expressed genes in Trop2$^+$RFP/tdTomato$^+$ vs Trop2$^-$RFP/tdTomato$^+$ neoplastic cells obtained in hyperplastic epithelium and adenomas 7 days and 6 weeks after Apc inactivation, respectively. Genes with adjusted p-value < 0.05 and $|\log_2 FC| \geq 1$ were used for the comparison.

**Supplementary Figure S1**

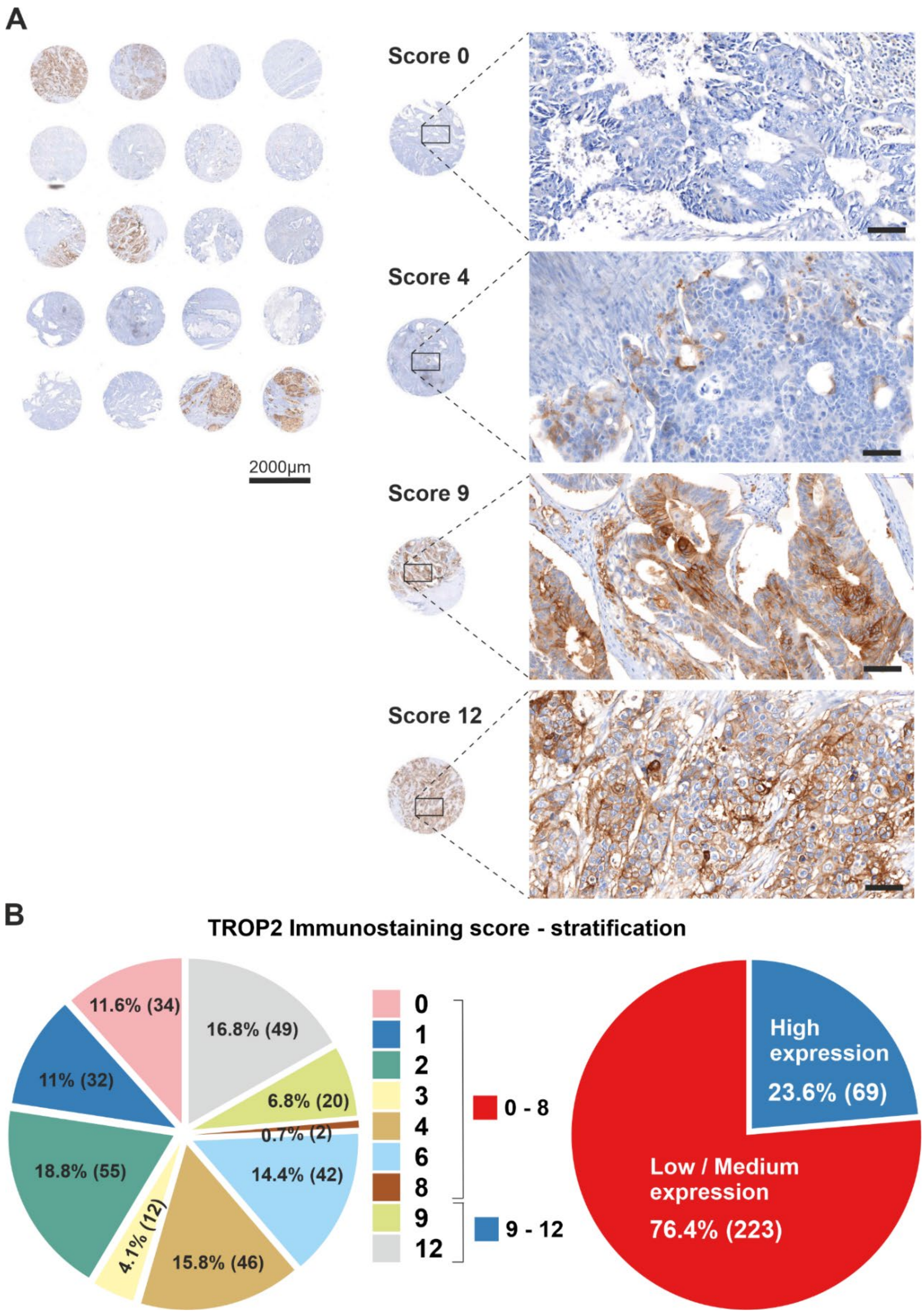

#### Supplementary Figure S2

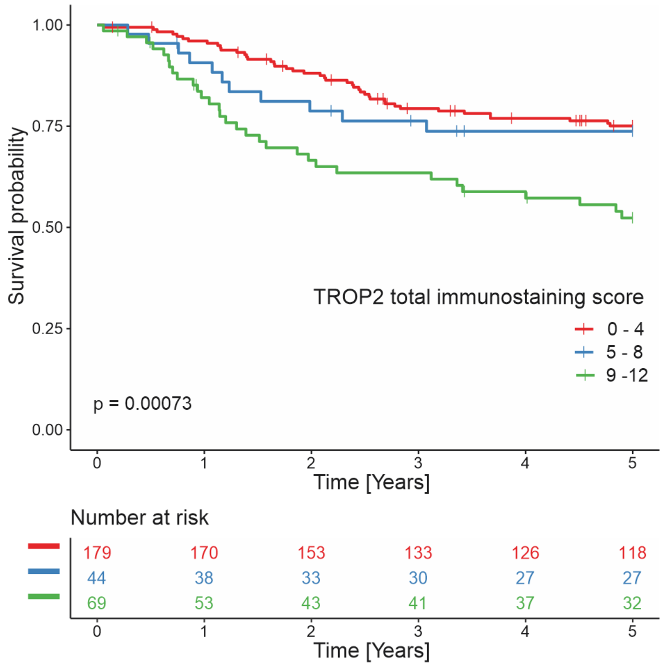

Supplementary Figure S3

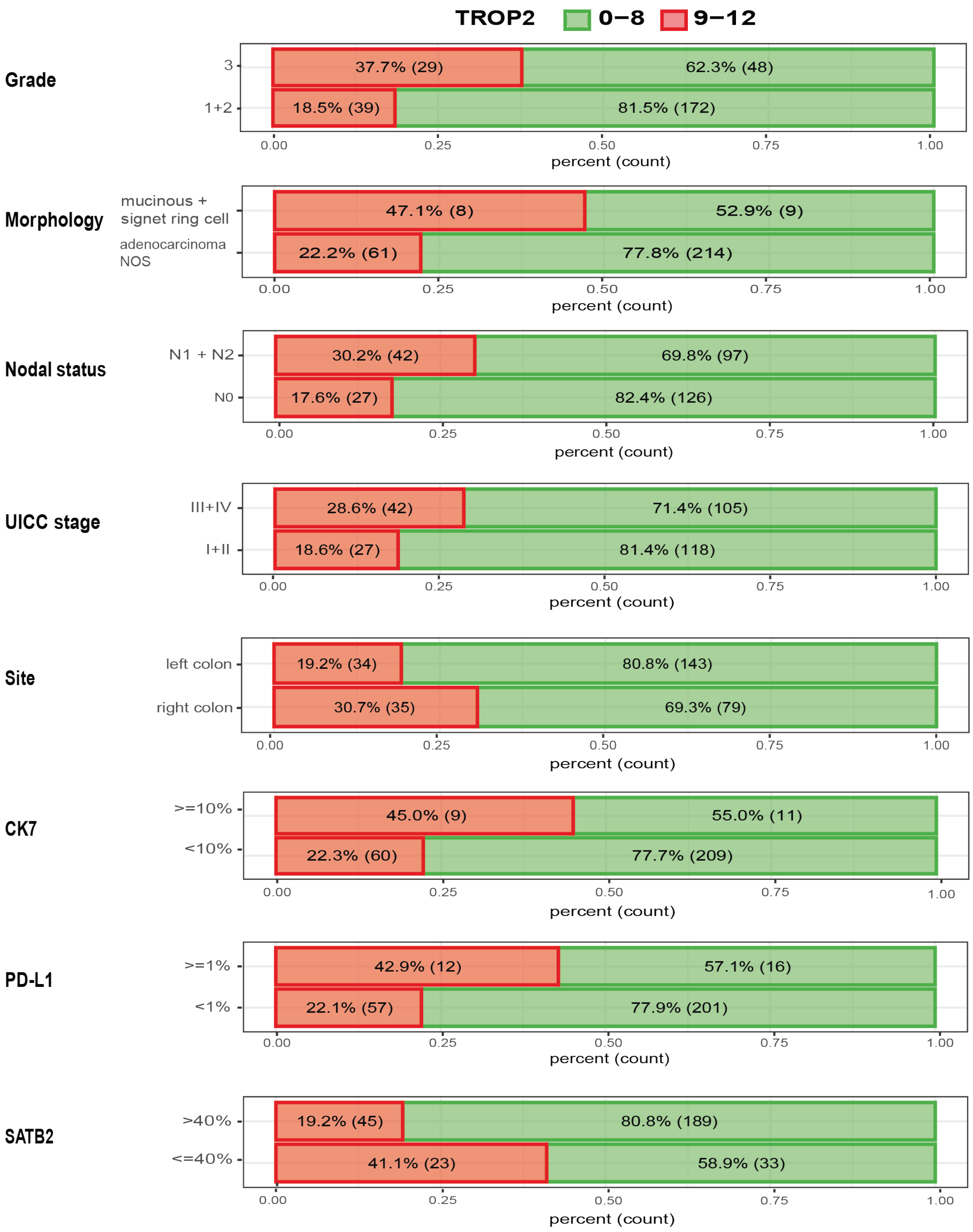

**Supplementary Figure S4**

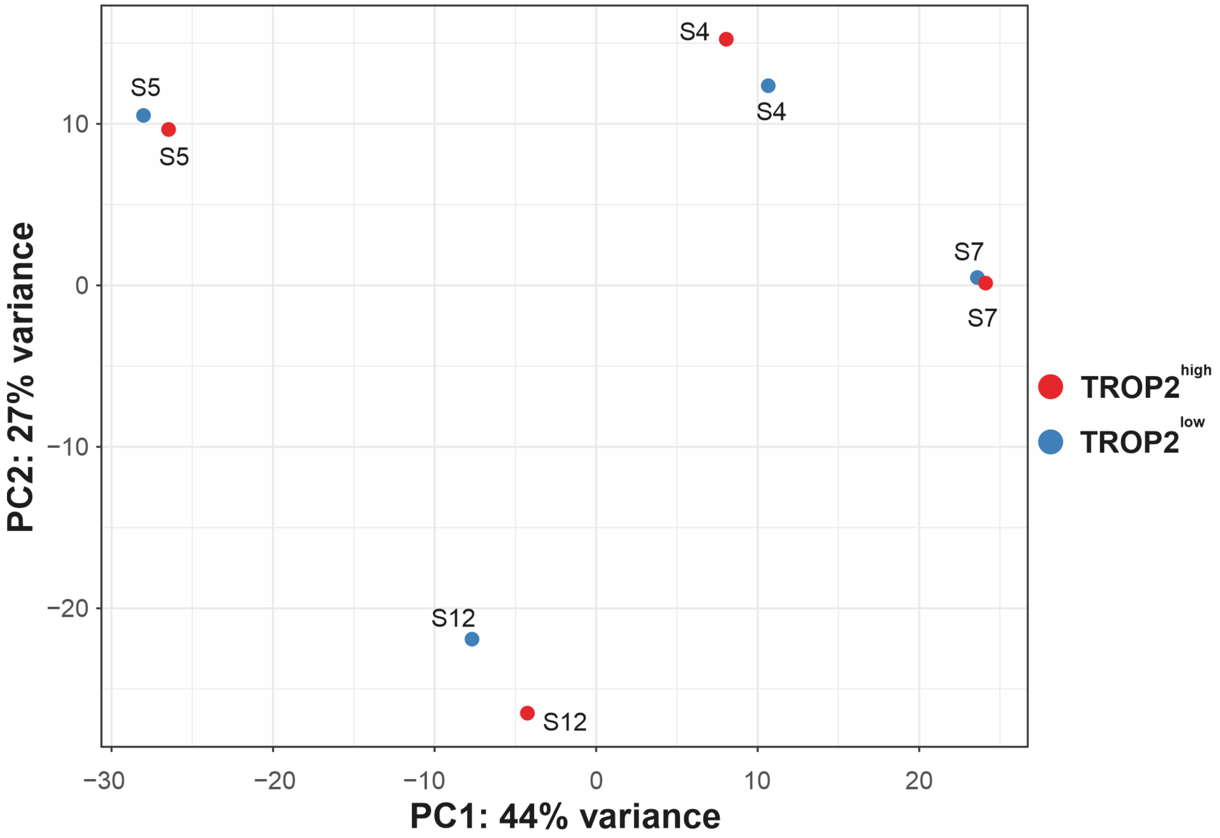


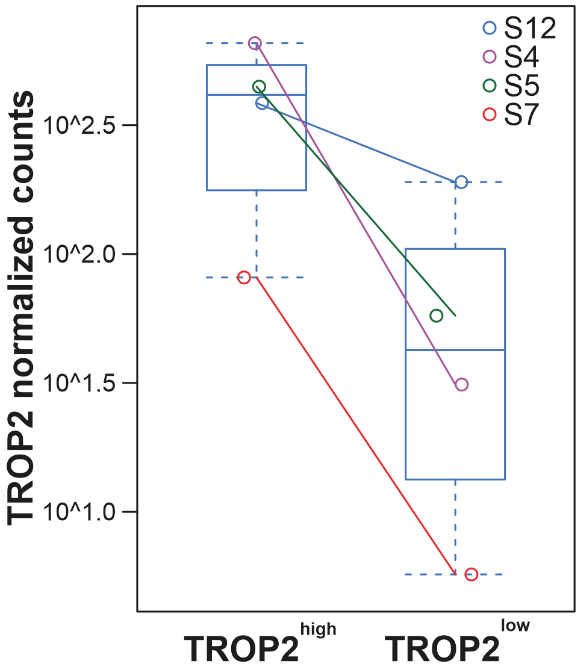

**Supplementary Figure S5**

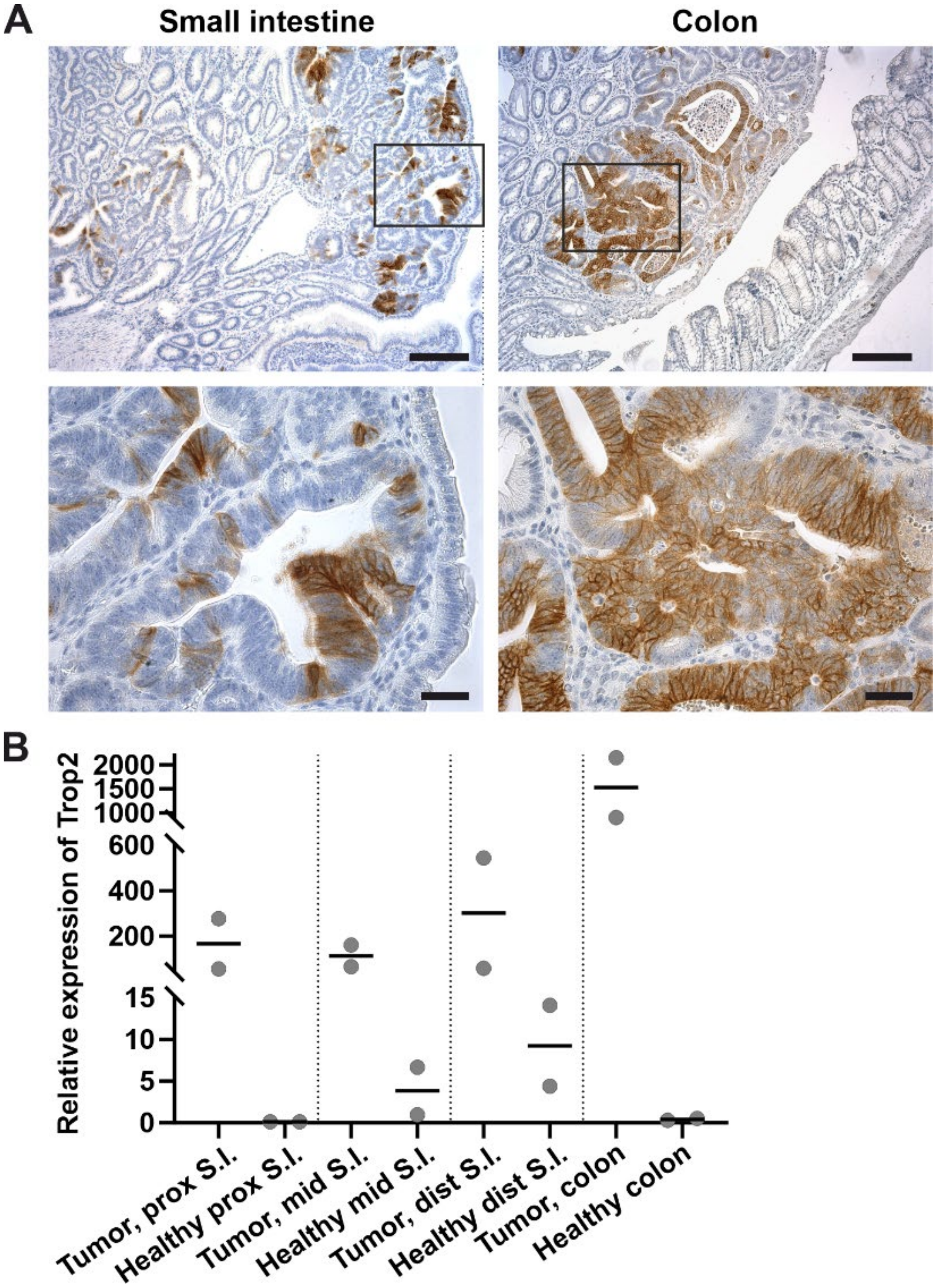

**Supplementary Figure S6**

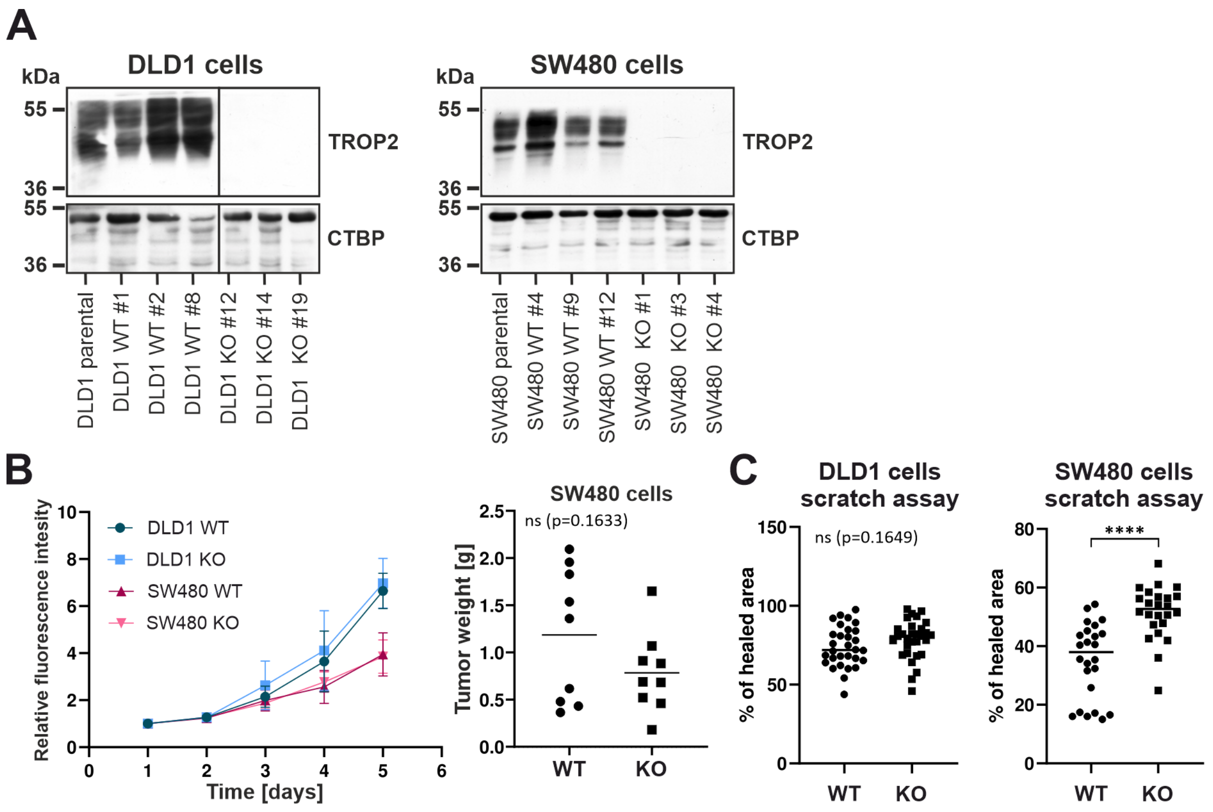

Supplementary Table S1

| **Histology** | **Age (median, min.-max.)** | **Gender (M/F)** |
|---|---|---|
| HYP | 69 (59-77) | 6/3 |
| LGD | 67 (53-89) | 12/15 |
| HGD | 64 (36-85) | 15/9 |
| CRC | 82 (63-90) | 6/6 |

### Supplementary Table S2

**Primers for qRT-PCR on mouse tissue**

| Gene symbol | Organism | Forward primer sequence (5´ to 3´ orientation) | Reverse primer sequence (5´ to 3´ orientation) |
|---|---|---|---|
| β-actin | Mouse | GATCTGGCACCACACCTTCT | GGGGTGTTGAAGGTCTCAAA |
| Cck | Mouse | TACATCCAGCAGGTCCGCAAAG | CGATGGGTATTCGTAGTCCTCGG |
| Cdh1 | Mouse | AGCCATTGCCAAGTACATCC | AAAGACCGGCTGGGTAAACT |
| Clu | Mouse | CACGCCATGAAGATTCTCCT | ATGTGCTTCACTCCCTGGAC |
| Gcg | Mouse | CGCTGATGGCTCCTTCTCTGAC | CAAGTGACTGGCACGAGATGTTG |
| GAPDH | Mouse | AACTTTGGCATTGTGGAAGG | ATCCACAGTCTTCTGGGTGG |
| Hopx | Mouse | TTAGTCAGACGCGCACGGACC | CACTCTGCCAGGCGCTGCTT |
| Ly6a/Sca1 | Mouse | TCTTGTGGCCCTACTGTGTG | GGCAGATGGGTAAGCAAAGA |
| Ngfr | Mouse | CAACCAGACCGTGTGTGAAC | GGAGAACACGAGTCCTGAGC |
| Olfm4 | Mouse | TGGGCAGAAGGTGGGACTGTGT | TGTCAGCGGGAAAGGCGGTA |
| Pgf | Mouse | CTGCTGGGAACAACTCAACA | GCTGTCTTTATCGGCACACA |
| Sox11 | Mouse | CTTCATGGTGTGGTCCAAGA | GTCGGGATAATCAGCCATGT |
| Trop2 | Mouse | GAACCCACCACATCCTCATT | ATGGTGGGCTCCTCATAGTG |
| Vegfr1/Flt1 | Mouse | ACCTCCGTGCATGTGTATGA | TCGTATAGTCCCCTGCATCC |
| Vim | Human | ACTGCTGCCCTGCGTGATGTG | GGTACTCGTTTGACTCCTGCTTGG |
| Ubb | Mouse | ATGTGAAGGCCAAGATCCAG | TAATAGCCACCCCTCAGACG |

**Primers for qRT-PCR on human organoids and cells**

| Gene symbol | Organism | Forward primer sequence (5´ to 3´ orientation) | Reverse primer sequence (5´ to 3´ orientation) |
|---|---|---|---|
| ALPI | Human | GCGGCTTCTACCTCTTTGTG | AAGCCACACCCTCATGATGA |
| ANKRD1 | Human | AGTAGAGGAACTGGTCACTGG | TGGGCTAGAAGTGTCTTCAGAT |
| ANXA1 | Human | GGTGACCGATCTGAGGACTT | ACCTCTGCGAAGTTGTGGAT |
| ASCL2 | Human | GCGAGCTACTCGACTTCTCC | CTCGGCTTCCGGGGCTGAGG |
| AXIN2 | Human | TGAGGTCCACGGAAACTGTTGACAGT | CCCTCCCGCGAATTGAGTGTGA |
| β-ACTIN | Human | GGCATCCTCACCCTGAAGTA | AGGTGTGGTGCCAGATTTTC |
| CTGF | Human | CCTGCAGGCTAGAGAAGCAG | TGGAGATTTTGGGAGTACGG |
| CTNNB | Human | TTCCAGACACGCTATCATGC | AATCCACTGGTGAACCAAGC |
| CTSE | Human | GGAATTCTGGGCCTGGGATA | TCTGACTGCCAAGCCTTGAT |
| CYR61 | Human | CCTTGTGGACAGCCAGTGTA | ACTTGGGCCGGTATTTCTTC |
| E-cadherin | Human | CAGTCAAAAGGCCTCTACGG | GGCGTAGACCAAGAAATGGA |
| E2F1 | Human | TCACCACCACCATCATCTCC | TGGGGAAAGGCTGATGAACT |
| LGR5 | Human | CTCTTCCTCAAACCGTCTGC | GATCGGAGGCTAAGCAACTG |
| SCA1/LY6a | Human | TTTCTTCGCTTGCTCCTCAC | CCACTGTTTCATCCCTTGGC |
| SIM | Human | ATGTGAAGGTTGCCCAAAAC | AAAATTGGCCATGTTTTCCA |
| TAZ | Human | AGCAGTCCTACGACGTGACC | CTAGGGTCTTGCCATGTGGT |
| TBP | Human | AGTTCTGGGATTGTACCGCA | TGTGCACACCATTTTCCCAG |
| TEAD2 | Human | GCCTCCGAGAGCTATATGATCG | TCACTCCGTAGAAGCCACCA |
| TROP2 | Human | ACAACGATGGCCTCTACGAC | TCCAGGTCTGAGTGGTTGAA |
| Vimentin | Human | TGCCCTTAAAGGAACCAATG | TCCAGCAGCTTCCTGTAGGT |
| YAP | Human | CTCGAACCCCAGATGACTTC | CCAGGAATGGCTTCAAGGTA |

**Oligos for single-guide RNA sequences**

| **Oligo name** | **5´- 3´ primer sequence** |
|---|---|
| FhTROP2 lentiCRISPR_a | CACCGTGGGGCGGAGGAACGCGGAC |
| RhTROP2 lentiCRISPR_a | AAACGTCCGCGTTCCTCCGCCCCAC |
| FhTROP2 lentiCRISPR_b | CACCGACACGTGCAGTTGTCCTGCG |
| RhTROP2 lentiCRISPR_b | AAACCGCAGGACAACTGCACGTGTC |
| FhTROP2 pARv-RFP_a | GTGGGGCGGAGGAACGCGGACCGGAT |
| RhTROP2 pARv-RFP_a | CCGGTCCGCGTTCCTCCGCCCCAC |
| FhTROP2 pARv-RFP_b | GACACGTGCAGTTGTCCTGCGCGGAT |
| RhTROP2 pARv-RFP_b | CCGCGCAGGACAACTGCACGTGTC |

**TROP2 genotyping PCR primers**

| **Oligo name** | **5´- 3´ primer sequence** |
|---|---|
| FhTROP2 genotyping | CCTGCAGACCATCCCAGAC |
| RhTROP2 genotyping | TGCACACGGTCATCTTGTTG |

**Sequencing primers**

| **Vector** | **5´- 3´ primer sequence** |
|---|---|
| pARv-RFP | CACCGCTAATTCAAAGCAACCG |
| lentiCRISPRv2 | CTACTATTCTTTCCCCTGCACTGTAC |
| pGEM T-easy | GATTTAGGTGACACTATAG |

**Quantitative RT-PCR primers and UPL probes for analysis of human tissue samples**

| Gene | 5´- 3´ primer sequence | UPL- probe No. |
|---|---|---|
| FhTROP2 | cgcaaaggagacgtttatcc | 8 |
| RhTROP2 | ctggtgtgtgcgcaaaag | |
| FhUBB | aggatcctggtatccgctaac | 39 |
| RhUBB | tcacattttcgatggtgtcact | |
| FhTBP | gaacatcatggatcagaacaaca | 87 |
| RhTBP | atagggattccgggagtcat | |

#### Supplementary Table S3

| | | TROP2 expression (%) | | Univariate analysis by logistic regression | |
|---|---|---|---|---|---|
| **Variable** | **Number (%)** | **Low/Medium score (0 - 8)** | **High score (9 - 12)** | **Odds ratio (95% CI)** | **p-value** |
| **Age at diagnosis (n = 292; 100%)** | | | | | |
| mean ± SD | 68.5 ± 11 | | | | |
| < 75 years | 205 (70.2%) | 156 (76.0%) | 49 (24%) | | |
| ≥ 75 years | 87 (29,8%) | 67 (77%) | 20 (23%) | 0.97 (0.51 - 1.69) | 0.85 |
| **Gender (n = 292; 100%)** | | | | | |
| female | 129 (44.2%) | 93 (72.1%) | 36 (27.9%) | | |
| male | 163 (55.8%) | 130 (79.8%) | 33 (20.2%) | 0.66 (0.38 - 1.13) | 0.127 |
| **Site (n = 291; 99.7%)** | | | | | |
| left colon | 178 (61%) | 143 (80.8%) | 34 (19.2%) | | |
| right colon | 113 (38.7%) | 79 (69.3%) | 35 (30.7%) | 1.85 (1.08 - 3.23) | 0.026 |
| **Morphology (n = 292; 100%)** | | | | | |
| adenocarcinoma, NOS | 275 (94.2%) | 214 (77.8%) | 61 (22.2%) | | |
| mucinous + signet ring cell | 17 (5.8%) | 9 (52.9%) | 8 (47.1%) | 3.12 (1.13 - 8.50) | 0.025 |
| **Grade (n = 288; 98.7%)** | | | | | |
| 1+2 | 211 (72.3%) | 172 (81.5%) | 39 (18.5%) | | |
| 3 | 77 (26.4%) | 48 (62.3%) | 29 (37.7%) | 2.67 (1.50 - 4.75) | <0.001 |
| **Lymph node status (n = 292; 100%)** | | | | | |
| N0 | 153 (52.4%) | 126 (82.4%) | 27 (17.6%) | | |
| N1+N2 | 139 (47.6%) | 97 (69.8%) | 42 (30.2%) | 2.02 (1.17 - 3.54) | 0.012 |
| **UICC stage (n = 292; 100%)** | | | | | |
| I+II | 145 (49.7%) | 118 (81.4%) | 27 (18.6%) | | |
| III+IV | 147 (50.3%) | 105 (71.4%) | 42 (28.6%) | 1.75 (1.01 - 3. 06) | 0.047 |
| **Therapy (n = 291; 99.7%)** | | | | | |
| none | 132 (45.2%) | 104 (78.8%) | 28 (21.2%) | | |
| neoadjuvant | 40 (13,7%) | 33 (82.5%) | 7 (17.5%) | | |
| adjuvant and/or paliative | 119 (40.8%) | 85 (71.4%) | 34 (28.6%) | 1.29 (0.75 - 2.25) | 0.362 |
| **PD-L1 (n = 284; 98%)** | | | | | |
| < 1% | 258 (88.4%) | 201 (77.9%) | 57 (22.1%) | | |
| ≥ 1% | 28 (9.6%) | 16 (57.1%) | 12 (42.9%) | 2.65 (1.16 - 5.89) | 0.018 |
| **CK7 (n = 289; 98.9%)** | | | | | |
| < 10% | 269 (92.1%) | 209 (77.7'%) | 60 (22.3%) | | |
| ≥ 10% | 20 (6.8%) | 11 (55%) | 9 (45%) | 2.85 (1.10 - 7.21) | 0.027 |
| **CK20 (n = 289; 98,9%)** | | | | | |
| < 25% | 67 (22.9%) | 45 (67.2%) | 22 (32.8%) | | |
| ≥ 25% | 222 (76%) | 175 (78.8%) | 47 (21.2%) | 0.55 (0.30 - 1.01) | 0.052 |

**Supplementary Table S4**

Differentially expressed genes (DEGs) in Trop2$^{high}$ compared to Trop2$^{low}$ tumor cells.
Genes with significantly altered expression in hyperplastic epithelium 7 days and in adenomas 6 weeks after Apc inactivation are listed on the appropriate sheet according to whether they were upregulated (72 genes) or downregulated (31 genes). Genes with an adjusted p-value < 0.05 in both experiments were considered significant; pseudogenes were removed from the lists.

**Upregulated genes**

| Ensembl gene ID | Symbol | | log2FC | p-value | FDR | Mean expression |
|---|---|---|---|---|---|---|
| ENSG00000198211 | AC092143.1 | novel protein (MC1R-TUBB3 readthrough) | 3.69E+00 | 7.15E-06 | 2.35E-03 | 1.21 |
| ENSG00000184292 | TACSTD2 | tumor associated calcium signal transducer 2 | 2.89E+00 | 7.75E-09 | 5.68E-06 | 2.36 |
| ENSG00000131016 | Up | A-kinase anchoring protein 12 | 2.59E+00 | 1.94E-09 | 1.65E-06 | 2.31 |
| ENSG00000277991 | FP236241.1 | uncharacterized LOC102723360 | 2.12E+00 | 2.81E-05 | 7.07E-03 | 1.64 |
| ENSG00000284419 | MIR663A | microRNA 663a | 1.55E+00 | 2.28E-18 | 3.68E-14 | 3.58 |
| ENSG00000196188 | CTSE | cathepsin E | 1.52E+00 | 2.30E-07 | 1.16E-04 | 2.52 |
| ENSG00000249624 | AP000295.1 | novel protein | 1.41E+00 | 5.75E-05 | 1.27E-02 | 1.76 |
| ENSG00000212443 | SNORA53 | small nucleolar RNA, H/ACA box 53 | 1.33E+00 | 5.50E-10 | 5.21E-07 | 2.6 |
| ENSG00000199270 | RNA5S12 | RNA, 5S ribosomal 12 | 1.28E+00 | 4.03E-12 | 6.88E-09 | 3.19 |
| ENSG00000199396 | RNA5S5 | RNA, 5S ribosomal 5 | 1.28E+00 | 4.03E-12 | 6.88E-09 | 3.19 |
| ENSG00000201925 | RNA5S15 | RNA, 5S ribosomal 15 | 1.28E+00 | 4.03E-12 | 6.88E-09 | 3.19 |
| ENSG00000137673 | MMP7 | matrix metallopeptidase 7 | 1.27E+00 | 2.70E-06 | 1.06E-03 | 2.72 |
| ENSG00000206941 | SNORD15A | small nucleolar RNA, C/D box 15A | 1.24E+00 | 6.29E-13 | 3.38E-09 | 3.38 |
| ENSG00000264462 | MIR3648-2 | microRNA 3648-2 | 1.22E+00 | 1.58E-12 | 6.03E-09 | 4.36 |
| ENSG00000198695 | MT-ND6 | mitochondrially encoded NADH:ubiquinone oxidoreductase core subunit 6 | 1.07E+00 | 5.01E-07 | 2.35E-04 | 3.32 |
| ENSG00000201998 | SNORA23 | small nucleolar RNA, H/ACA box 23 | 1.06E+00 | 7.82E-08 | 4.50E-05 | 2.69 |
| ENSG00000227195 | MIR663AHG | MIR663A host gene | 1.06E+00 | 1.19E-05 | 3.62E-03 | 4.89 |
| ENSG00000201321 | RNA5S9 | RNA, 5S ribosomal 9 | 1.05E+00 | 5.26E-06 | 1.97E-03 | 2.54 |
| ENSG00000201403 | SNORD14B | small nucleolar RNA, C/D box 14B | 1.00E+00 | 3.13E-05 | 7.63E-03 | 2.45 |
| ENSG00000122861 | PLAU | plasminogen activator, urokinase | 9.71E-01 | 6.09E-05 | 1.31E-02 | 1.95 |
| ENSG00000023445 | BIRC3 | baculoviral IAP repeat containing 3 | 9.56E-01 | 2.22E-06 | 9.16E-04 | 2.71 |
| ENSG00000196924 | FLNA | filamin A | 9.50E-01 | 3.26E-08 | 1.94E-05 | 3.62 |
| ENSG00000202252 | SNORD14C | small nucleolar RNA, C/D box 14C | 9.45E-01 | 2.05E-05 | 5.69E-03 | 2.73 |
| ENSG00000011422 | PLAUR | plasminogen activator, urokinase receptor | 9.41E-01 | 5.53E-06 | 1.98E-03 | 2.78 |
| ENSG00000198888 | MT-ND1 | mitochondrially encoded NADH:ubiquinone oxidoreductase core subunit 1 | 9.33E-01 | 1.87E-12 | 6.03E-09 | 5.21 |
| ENSG00000212907 | MT-ND4L | mitochondrially encoded NADH:ubiquinone oxidoreductase core subunit 4L | 9.23E-01 | 2.36E-11 | 3.16E-08 | 4.3 |
| ENSG00000140279 | DUOX2 | dual oxidase 2 | 8.90E-01 | 1.21E-08 | 7.83E-06 | 3.66 |
| ENSG00000277947 | SNORD3D | small nucleolar RNA, C/D box 3D | 8.76E-01 | 1.38E-07 | 7.43E-05 | 4.39 |
| ENSG00000198727 | MT-CYB | mitochondrially encoded cytochrome b | 8.75E-01 | 4.47E-10 | 4.80E-07 | 5.15 |
| ENSG00000228253 | MT-ATP8 | mitochondrially encoded ATP synthase membrane subunit 8 | 8.67E-01 | 1.56E-09 | 1.39E-06 | 4.54 |
| ENSG00000135046 | ANXA1 | annexin A1 | 8.67E-01 | 5.92E-05 | 1.29E-02 | 2.7 |
| ENSG00000129451 | KLK10 | kallikrein related peptidase 10 | 8.52E-01 | 9.48E-05 | 1.86E-02 | 1.87 |
| ENSG00000053747 | LAMA3 | laminin subunit alpha 3 | 8.15E-01 | 2.84E-06 | 1.09E-03 | 3.32 |
| ENSG00000169908 | TM4SF1 | transmembrane 4 L six family member 1 | 8.03E-01 | 1.32E-06 | 5.77E-04 | 2.92 |
| ENSG00000134531 | EMP1 | epithelial membrane protein 1 | 7.90E-01 | 9.91E-05 | 1.92E-02 | 2.95 |
| ENSG00000135074 | ADAM19 | ADAM metallopeptidase domain 19 | 7.80E-01 | 2.38E-05 | 6.29E-03 | 2.99 |
| ENSG00000222041 | CYTOR | cytoskeleton regulator RNA | 7.75E-01 | 1.67E-04 | 2.89E-02 | 2.15 |
| ENSG00000074276 | CDHR2 | cadherin related family member 2 | 7.36E-01 | 8.66E-06 | 2.73E-03 | 2.92 |
| ENSG00000058085 | LAMC2 | laminin subunit gamma 2 | 7.28E-01 | 5.63E-06 | 1.98E-03 | 3.57 |
| ENSG00000102287 | GABRE | gamma-aminobutyric acid type A receptor subunit epsilon | 7.03E-01 | 2.80E-05 | 7.07E-03 | 3.01 |
| ENSG00000207304 | SNORA8 | small nucleolar RNA, H/ACA box 8 | 7.02E-01 | 8.51E-05 | 1.71E-02 | 2.61 |
| ENSG00000263740 | RN7SL4P | RNA, 7SL, cytoplasmic 4, pseudogene | 6.94E-01 | 3.26E-05 | 7.78E-03 | 3.56 |
| ENSG00000198840 | MT-ND3 | mitochondrially encoded NADH:ubiquinone oxidoreductase core subunit 3 | 6.85E-01 | 5.11E-07 | 2.35E-04 | 4.26 |
| ENSG00000200959 | SNORA74A | small nucleolar RNA, H/ACA box 74A | 6.83E-01 | 2.84E-04 | 4.19E-02 | 2.35 |
| ENSG00000245532 | NEAT1 | nuclear paraspeckle assembly transcript 1 | 6.70E-01 | 8.80E-09 | 6.03E-06 | 4.31 |
| ENSG00000252010 | SCARNA5 | small Cajal body-specific RNA 5 | 6.53E-01 | 1.44E-04 | 2.58E-02 | 2.93 |
| ENSG00000163739 | CXCL1 | C-X-C motif chemokine ligand 1 | 6.52E-01 | 1.27E-05 | 3.78E-03 | 3.01 |
| ENSG00000207241 | SNORD45A | small nucleolar RNA, C/D box 45A | 6.38E-01 | 3.44E-04 | 4.84E-02 | 2.51 |
| ENSG00000172927 | MYEOV | myeloma overexpressed | 6.31E-01 | 2.09E-04 | 3.43E-02 | 2.55 |
| ENSG00000254341 | SNORD87 | small nucleolar RNA, C/D box 87 | 6.26E-01 | 1.99E-04 | 3.31E-02 | 2.66 |
| ENSG00000267322 | SNHG22 | small nucleolar RNA host gene 22 | 6.19E-01 | 1.20E-04 | 2.24E-02 | 2.8 |
| ENSG00000142627 | EPHA2 | EPH receptor A2 | 5.86E-01 | 5.65E-06 | 1.98E-03 | 3 |
| ENSG00000204616 | TRIM31 | tripartite motif containing 31 | 5.68E-01 | 1.68E-05 | 4.83E-03 | 3.15 |
| ENSG00000238917 | SNORD10 | small nucleolar RNA, C/D box 10 | 5.63E-01 | 8.45E-05 | 1.71E-02 | 3.96 |
| ENSG00000280138 | AC027290.2 | novel transcript | 5.40E-01 | 7.89E-05 | 1.63E-02 | 2.96 |
| ENSG00000206588 | RNU1-28P | RNA, U1 small nuclear 28, pseudogene | 5.20E-01 | 3.00E-05 | 7.43E-03 | 3.87 |
| ENSG00000173559 | NABP1 | nucleic acid binding protein 1 | 5.04E-01 | 2.79E-04 | 4.16E-02 | 2.69 |
| ENSG00000130707 | ASS1 | argininosuccinate synthase 1 | 4.73E-01 | 3.61E-04 | 4.93E-02 | 3.38 |
| ENSG00000178209 | PLEC | plectin | 4.53E-01 | 6.96E-07 | 3.11E-04 | 4.3 |
| ENSG00000130164 | LDLR | low density lipoprotein receptor | 4.44E-01 | 2.56E-05 | 6.66E-03 | 3.34 |
| ENSG00000210082 | MT-RNR2 | mitochondrially encoded 16S rRNA | 4.23E-01 | 1.51E-04 | 2.68E-02 | 5.62 |
| ENSG00000090382 | LYZ | lysozyme | 4.19E-01 | 1.21E-04 | 2.24E-02 | 3.55 |
| ENSG00000124942 | AHNAK | AHNAK nucleoprotein | 4.08E-01 | 9.60E-06 | 2.97E-03 | 4.26 |
| ENSG00000123358 | NR4A1 | nuclear receptor subfamily 4 group A member 1 | 4.07E-01 | 2.70E-04 | 4.06E-02 | 3.28 |
| ENSG00000182718 | ANXA2 | annexin A2 | 3.95E-01 | 2.31E-04 | 3.65E-02 | 3.84 |
| ENSG00000267519 | AC020916.1 | novel transcript | 3.85E-01 | 8.89E-05 | 1.77E-02 | 3.34 |
| ENSG00000173898 | SPTBN2 | spectrin beta, non-erythrocytic 2 | 3.80E-01 | 1.60E-04 | 2.80E-02 | 3.2 |
| ENSG00000112715 | VEGFA | vascular endothelial growth factor A | 3.77E-01 | 3.81E-05 | 8.77E-03 | 3.68 |
| ENSG00000092820 | EZR | ezrin | 3.41E-01 | 4.20E-05 | 9.39E-03 | 3.81 |
| ENSG00000062716 | VMP1 | vacuole membrane protein 1 | 3.05E-01 | 3.30E-04 | 4.70E-02 | 3.68 |
| ENSG00000151914 | DST | dystonin | 3.03E-01 | 2.59E-04 | 3.98E-02 | 3.83 |
| ENSG00000197956 | S100A6 | S100 calcium binding protein A6 | 2.95E-01 | 3.53E-04 | 4.85E-02 | 4.07 |

**Downregulated genes**

| Ensembl gene ID | Symbol | | log2FC | p-value | adjusted p-value | Mean expression |
|---|---|---|---|---|---|---|
| ENSG00000276345 | AC004556.3 | 39S ribosomal protein L23, mitochondrial | -2.87E-01 | 3.49E-04 | 4.84E-02 | 2.29 |
| ENSG00000169100 | SLC25A6 | solute carrier family 25 member 6 | -2.88E-01 | 1.02E-04 | 1.95E-02 | 3.71 |
| ENSG00000165238 | WNK2 | WNK lysine deficient protein kinase 2 | -2.97E-01 | 2.56E-04 | 3.98E-02 | 3.64 |
| ENSG00000135916 | ITM2C | integral membrane protein 2C | -3.23E-01 | 1.29E-04 | 2.36E-02 | 3.45 |
| ENSG00000198643 | FAM3D | FAM3 metabolism regulating signaling molecule D | -4.39E-01 | 2.62E-04 | 3.98E-02 | 3.17 |
| ENSG00000267022 | AC067968.1 | novel protein | -4.72E-01 | 3.25E-04 | 4.67E-02 | 1.78 |
| ENSG00000135503 | ACVR1B | activin A receptor type 1B | -4.84E-01 | 1.88E-04 | 3.18E-02 | 2.99 |
| ENSG00000237356 | AL365295.1 | novel transcript | -4.88E-01 | 2.94E-04 | 4.30E-02 | 1.33 |
| ENSG00000204219 | TCEA3 | transcription elongation factor A3 | -5.12E-01 | 2.33E-05 | 6.29E-03 | 3.06 |
| ENSG00000188803 | SHISA6 | shisa family member 6 | -5.27E-01 | 3.05E-04 | 4.43E-02 | 2.11 |
| ENSG00000101333 | PLCB4 | phospholipase C beta 4 | -5.32E-01 | 6.29E-05 | 1.32E-02 | 3.58 |
| ENSG00000174992 | ZG16 | zymogen granule protein 16 | -5.74E-01 | 2.19E-04 | 3.49E-02 | 2.24 |
| ENSG00000256028 | AC026362.1 | novel transcript | -5.89E-01 | 2.14E-04 | 3.46E-02 | 1.47 |
| ENSG00000148057 | IDNK | IDNK gluconokinase | -6.18E-01 | 2.15E-04 | 3.46E-02 | 1.51 |
| ENSG00000116299 | ELAPOR1 | endosome-lysosome associated apoptosis and autophagy regulator 1 | -7.16E-01 | 3.28E-05 | 7.78E-03 | 3.22 |
| ENSG00000162896 | PIGR | polymeric immunoglobulin receptor | -7.17E-01 | 2.21E-11 | 3.16E-08 | 4.85 |
| ENSG00000074370 | ATP2A3 | ATPase sarcoplasmic/endoplasmic reticulum Ca2+ transporting 3 | -8.34E-01 | 1.43E-08 | 8.88E-06 | 3.18 |
| ENSG00000117983 | MUC5B | mucin 5B, oligomeric mucus/gel-forming | -8.54E-01 | 8.28E-08 | 4.60E-05 | 4.47 |
| ENSG00000206127 | GOLGA8O | golgin A8 family member O | -9.63E-01 | 1.11E-04 | 2.10E-02 | 1.41 |
| ENSG00000167080 | B4GALNT2 | beta-1,4-N-acetyl-galactosaminyltransferase 2 | -9.65E-01 | 6.18E-05 | 1.31E-02 | 2.51 |
| ENSG00000198788 | MUC2 | mucin 2, oligomeric mucus/gel-forming | -9.73E-01 | 8.99E-09 | 6.03E-06 | 5.3 |
| ENSG00000128917 | DLL4 | delta like canonical Notch ligand 4 | -9.81E-01 | 7.62E-06 | 2.45E-03 | 2.53 |
| ENSG00000174136 | RGMB | repulsive guidance molecule BMP co-receptor b | -9.86E-01 | 5.94E-06 | 2.04E-03 | 2.9 |
| ENSG00000073849 | ST6GAL1 | ST6 beta-galactoside alpha-2,6-sialyltransferase 1 | -1.21E+00 | 4.09E-10 | 4.70E-07 | 3.03 |
| ENSG00000112562 | SMOC2 | SPARC related modular calcium binding 2 | -1.64E+00 | 5.24E-10 | 5.21E-07 | 2.65 |
| ENSG00000267881 | AC243967.1 | novel protein, readthrough between CEACAM5-CEACAM6 | -1.88E+00 | 3.50E-05 | 8.16E-03 | 1.66 |
| ENSG00000261771 | DNAAF4-CCPG1 | DNAAF4-CCPG1 readthrough (NMD candidate) | -2.26E+00 | 2.37E-05 | 6.29E-03 | 1.27 |
| ENSG00000277150 | F8A3 | coagulation factor VIII associated 3 | -2.46E+00 | 1.77E-05 | 5.00E-03 | 1.57 |
| ENSG00000118785 | SPP1 | secreted phosphoprotein 1 | -5.06E+00 | 4.30E-09 | 3.30E-06 | 1.73 |
| ENSG00000250151 | ARPC4-TTLL3 | ARPC4-TTLL3 readthrough | -5.53E+00 | 1.47E-07 | 7.61E-05 | 1.67 |
| ENSG00000100604 | CHGA | chromogranin A | -6.32E+00 | 3.12E-07 | 1.52E-04 | 2.13 |

### Supplementary Table S5

Differentially expressed genes (DEGs) in two populations of RFP-positive, i.e. proliferating cells sorted for Trop2 expression, and in a population of RFP-negative, i.e. differentiated cells sorted RFP dataset from hyperplastic epithelium 7 days after Apc inactivation. Genes with an adjusted p-value < 0.05 and a log change $|\log_2 FC| \geq 1$ were considered significant. DEGs between $Trop2^{+}RFP^{+}$ and $Trop2^{-}RFP^{+}$ cells and between the two $RFP^{+}$ cell-derived gene expression profiles the $RFP^{-}$ cells dataset and the are shown on the corresponding sheet.

**RFP+Trop2+ vs. RFP+Trop2- cells**

| Ensemble gene code | Symbol | Gene name | logFC |
|---|---|---|---|
| ENSMUSG00000029304 | Spp1 | secreted phosphoprotein 1 | 7.42 |
| ENSMUSG00000027485 | Bpifb1 | BPI fold containing family B, member 1 | 6.53 |
| ENSMUSG00000051397 | Tacstd2 | tumor-associated calcium signal transducer 2 | 6.26 |
| ENSMUSG00000041523 | Upk2 | uroplakin 2 | 5.85 |
| ENSMUSG00000032473 | Cldn18 | claudin 18 | 5.76 |
| ENSMUSG00000074625 | Arhgap40 | Rho GTPase activating protein 40 | 5.61 |
| ENSMUSG00000049001 | Ndnf | neuron-derived neurotrophic factor | 5.46 |
| ENSMUSG00000059900 | Tmem40 | transmembrane protein 40 | 5.40 |
| ENSMUSG00000039005 | Tlr4 | toll-like receptor 4 | 5.32 |
| ENSMUSG00000021950 | Anxa8 | annexin A8 | 4.98 |
| ENSMUSG00000059668 | Krt4 | keratin 4 | 4.76 |
| ENSMUSG00000024659 | Anxa1 | annexin A1 | 4.59 |
| ENSMUSG00000022037 | Clu | clusterin | 4.57 |
| ENSMUSG00000033377 | Palmd | palmdelphin | 4.44 |
| ENSMUSG00000091243 | Vgll3 | vestigial like family member 3 | 4.43 |
| ENSMUSG00000044912 | Syt16 | synaptotagmin XVI | 4.37 |
| ENSMUSG00000031841 | Cdh13 | cadherin 13 | 4.27 |
| ENSMUSG00000092586 | Ly6g6c | lymphocyte antigen 6 complex, locus G6C | 3.91 |
| ENSMUSG00000051076 | Vtcn1 | V-set domain containing T cell activation inhibitor 1 | 3.81 |
| ENSMUSG00000034463 | Scara3 | scavenger receptor class A, member 3 | 3.71 |
| ENSMUSG00000029648 | Flt1 | FMS-like tyrosine kinase 1 | 3.69 |
| ENSMUSG00000021822 | Plau | plasminogen activator, urokinase | 3.66 |
| ENSMUSG00000021765 | Fst | follistatin | 3.64 |
| ENSMUSG00000048376 | F2r | coagulation factor II (thrombin) receptor | 3.59 |
| ENSMUSG00000048915 | Efna5 | ephrin A5 | 3.58 |
| ENSMUSG00000037188 | Grhl3 | grainyhead like transcription factor 3 | 3.54 |
| ENSMUSG00000040998 | Npnt | nephronectin | 3.50 |
| ENSMUSG00000046213 | Cym | chymosin | 3.49 |
| ENSMUSG00000026065 | Slc9a4 | solute carrier family 9 (sodium/hydrogen exchanger), member 4 | 3.37 |
| ENSMUSG00000032327 | Stra6 | stimulated by retinoic acid gene 6 | 3.35 |
| ENSMUSG00000027376 | Prom2 | prominin 2 | 3.33 |
| ENSMUSG00000015134 | Aldh1a3 | aldehyde dehydrogenase family 1, subfamily A3 | 3.30 |
| ENSMUSG00000034573 | Ptpn13 | protein tyrosine phosphatase, non-receptor type 13 | 3.29 |
| ENSMUSG00000037946 | Fgd3 | FYVE, RhoGEF and PH domain containing 3 | 3.28 |
| ENSMUSG00000043088 | Il17re | interleukin 17 receptor E | 3.23 |
| ENSMUSG00000061517 | Sox21 | SRY (sex determining region Y)-box 21 | 3.20 |
| ENSMUSG00000044162 | Tnip3 | TNFAIP3 interacting protein 3 | 3.19 |
| ENSMUSG00000037833 | Sh2d4b | SH2 domain containing 4B | 3.16 |
| ENSMUSG00000056632 | Dsg3 | desmoglein 3 | 3.00 |
| ENSMUSG00000004552 | Ctse | cathepsin E | 2.95 |
| ENSMUSG00000039153 | Runx2 | runt related transcription factor 2 | 2.93 |
| ENSMUSG00000055407 | Map6 | microtubule-associated protein 6 | 2.93 |
| ENSMUSG00000028236 | Sdr16c5 | short chain dehydrogenase/reductase family 16C, member 5 | 2.88 |
| ENSMUSG00000032068 | Plet1 | placenta expressed transcript 1 | 2.88 |
| ENSMUSG00000029154 | Cwh43 | cell wall biogenesis 43 C-terminal homolog | 2.80 |
| ENSMUSG00000056888 | Glipr1 | GLI pathogenesis-related 1 (glioma) | 2.78 |
| ENSMUSG00000013338 | Fer1l4 | fer-1-like 4 (C. elegans) | 2.77 |
| ENSMUSG00000063011 | Msln | mesothelin | 2.77 |
| ENSMUSG00000031430 | Vsig1 | V-set and immunoglobulin domain containing 1 | 2.76 |
| ENSMUSG00000055333 | Fat2 | FAT atypical cadherin 2 | 2.75 |
| ENSMUSG00000045763 | Basp1 | brain abundant, membrane attached signal protein 1 | 2.74 |
| ENSMUSG00000026620 | Mark1 | MAP/microtubule affinity regulating kinase 1 | 2.73 |
| ENSMUSG00000021792 | Fam213a | family with sequence similarity 213, member A | 2.72 |
| ENSMUSG00000001288 | Rarg | retinoic acid receptor, gamma | 2.72 |
| ENSMUSG00000030510 | Cers3 | ceramide synthase 3 | 2.71 |
| ENSMUSG00000031075 | Ano1 | anoctamin 1, calcium activated chloride channel | 2.70 |
| ENSMUSG00000050640 | Tmem150c | transmembrane protein 150C | 2.70 |
| ENSMUSG00000050359 | Sprr1a | small proline-rich protein 1A | 2.70 |
| ENSMUSG00000040663 | Clcf1 | cardiotrophin-like cytokine factor 1 | 2.68 |
| ENSMUSG00000029484 | Anxa3 | annexin A3 | 2.67 |
| ENSMUSG00000040990 | Sh3kbp1 | SH3-domain kinase binding protein 1 | 2.67 |

| | | | |
|---|---|---|---|
| ENSMUSG00000029869 | Ephb6 | Eph receptor B6 | 2.67 |
| ENSMUSG00000048612 | Myof | myoferlin | 2.66 |
| ENSMUSG00000072812 | Ahnak2 | AHNAK nucleoprotein 2 | 2.63 |
| ENSMUSG00000021838 | Samd4 | sterile alpha motif domain containing 4 | 2.60 |
| ENSMUSG00000046402 | Rbp1 | retinol binding protein 1, cellular | 2.60 |
| ENSMUSG00000009687 | Fxyd5 | FXYD domain-containing ion transport regulator 5 | 2.59 |
| ENSMUSG00000033227 | Wnt6 | wingless-type MMTV integration site family, member 6 | 2.59 |
| ENSMUSG00000040488 | Ltbp4 | latent transforming growth factor beta binding protein 4 | 2.58 |
| ENSMUSG00000022754 | Tmem45a | transmembrane protein 45a | 2.56 |
| ENSMUSG00000032348 | Gsta4 | glutathione S-transferase, alpha 4 | 2.55 |
| ENSMUSG00000046449 | Nexmif | neurite extension and migration factor | 2.50 |
| ENSMUSG00000046623 | Gjb4 | gap junction protein, beta 4 | 2.49 |
| ENSMUSG00000063727 | Tnfrsf11b | tumor necrosis factor receptor superfamily, member 11b (osteoprotegerin) | 2.48 |
| ENSMUSG00000025938 | Slco5a1 | solute carrier organic anion transporter family, member 5A1 | 2.46 |
| ENSMUSG00000026193 | Fn1 | fibronectin 1 | 2.46 |
| ENSMUSG00000025473 | Adam8 | a disintegrin and metallopeptidase domain 8 | 2.45 |
| ENSMUSG00000026981 | Il1rn | interleukin 1 receptor antagonist | 2.44 |
| ENSMUSG00000022203 | Efs | embryonal Fyn-associated substrate | 2.44 |
| ENSMUSG00000062591 | Tubb4a | tubulin, beta 4A class IVA | 2.43 |
| ENSMUSG00000028716 | Pdzk1ip1 | PDZK1 interacting protein 1 | 2.43 |
| ENSMUSG00000031098 | Syt8 | synaptotagmin VIII | 2.42 |
| ENSMUSG00000108218 | Olfr1372-ps1 | olfactory receptor 1372, pseudogene 1 | 2.38 |
| ENSMUSG00000026548 | Slamf9 | SLAM family member 9 | 2.36 |
| ENSMUSG00000020651 | Slc26a4 | solute carrier family 26, member 4 | 2.35 |
| ENSMUSG00000001435 | Col18a1 | collagen, type XVIII, alpha 1 | 2.34 |
| ENSMUSG00000031635 | Anxa10 | annexin A10 | 2.32 |
| ENSMUSG00000037060 | Cavin3 | caveolae associated 3 | 2.29 |
| ENSMUSG00000008398 | Elk3 | ELK3, member of ETS oncogene family | 2.27 |
| ENSMUSG00000026405 | C4bp | complement component 4 binding protein | 2.24 |
| ENSMUSG00000024087 | Cyp1b1 | cytochrome P450, family 1, subfamily b, polypeptide 1 | 2.21 |
| ENSMUSG00000038587 | Akap12 | A kinase (PRKA) anchor protein (gravin) 12 | 2.21 |
| ENSMUSG00000051343 | Rab11fip5 | RAB11 family interacting protein 5 (class I) | 2.20 |
| ENSMUSG00000018340 | Anxa6 | annexin A6 | 2.19 |
| ENSMUSG00000034634 | Ly6d | lymphocyte antigen 6 complex, locus D | 2.18 |
| ENSMUSG00000026509 | Capn2 | calpain 2 | 2.17 |
| ENSMUSG00000037716 | Ccdc33 | coiled-coil domain containing 33 | 2.17 |
| ENSMUSG00000029371 | Cxcl5 | chemokine (C-X-C motif) ligand 5 | 2.17 |
| ENSMUSG00000039384 | Dusp10 | dual specificity phosphatase 10 | 2.15 |
| ENSMUSG00000080316 | Spaca6 | sperm acrosome associated 6 | 2.15 |
| ENSMUSG00000021367 | Edn1 | endothelin 1 | 2.13 |
| ENSMUSG00000047501 | Cldn4 | claudin 4 | 2.13 |
| ENSMUSG00000027712 | Anxa5 | annexin A5 | 2.12 |
| ENSMUSG00000026435 | Slc45a3 | solute carrier family 45, member 3 | 2.12 |
| ENSMUSG00000001025 | S100a6 | S100 calcium binding protein A6 (calcyclin) | 2.11 |
| ENSMUSG00000053007 | Creb5 | cAMP responsive element binding protein 5 | 2.11 |
| ENSMUSG00000079330 | Lemd1 | LEM domain containing 1 | 2.11 |
| ENSMUSG00000040152 | Thbs1 | thrombospondin 1 | 2.10 |
| ENSMUSG00000004951 | Hspb1 | heat shock protein 1 | 2.09 |
| ENSMUSG00000027624 | Epb41l1 | erythrocyte membrane protein band 4.1 like 1 | 2.08 |
| ENSMUSG00000019997 | Ctgf | connective tissue growth factor | 2.07 |
| ENSMUSG00000037185 | Krt80 | keratin 80 | 2.07 |
| ENSMUSG00000053137 | Mapk11 | mitogen-activated protein kinase 11 | 2.06 |
| ENSMUSG00000085412 | Halr1 | Hoxa adjacent long noncoding RNA 1 | 2.06 |
| ENSMUSG00000007039 | Ddah2 | dimethylarginine dimethylaminohydrolase 2 | 2.05 |
| ENSMUSG00000024421 | Lama3 | laminin, alpha 3 | 2.05 |
| ENSMUSG00000028031 | Dkk2 | dickkopf WNT signaling pathway inhibitor 2 | 2.02 |
| ENSMUSG00000031586 | Rbpms | RNA binding protein gene with multiple splicing | 2.02 |
| ENSMUSG00000038843 | Gcnt1 | glucosaminyl (N-acetyl) transferase 1, core 2 | 2.02 |
| ENSMUSG00000028128 | F3 | coagulation factor III | 2.01 |
| ENSMUSG00000043430 | Psapl1 | prosaposin-like 1 | 2.01 |
| ENSMUSG00000022055 | Nefl | neurofilament, light polypeptide | 2.01 |
| ENSMUSG00000042734 | Ttc9 | tetratricopeptide repeat domain 9 | 2.00 |
| ENSMUSG00000067889 | Sptbn2 | spectrin beta, non-erythrocytic 2 | 1.98 |
| ENSMUSG00000045027 | Prss22 | protease, serine 22 | 1.98 |

| | | | |
|---|---|---|---|
| ENSMUSG00000038463 | Olfml2b | olfactomedin-like 2B | 1.98 |
| ENSMUSG00000022505 | Emp2 | epithelial membrane protein 2 | 1.97 |
| ENSMUSG00000032013 | Trim29 | tripartite motif-containing 29 | 1.96 |
| ENSMUSG00000000983 | Wfdc18 | WAP four-disulfide core domain 18 | 1.95 |
| ENSMUSG00000034220 | Gpc1 | glypican 1 | 1.95 |
| ENSMUSG00000023073 | Slc10a2 | solute carrier family 10, member 2 | 1.95 |
| ENSMUSG00000031538 | Plat | plasminogen activator, tissue | 1.94 |
| ENSMUSG00000031871 | Cdh5 | cadherin 5 | 1.94 |
| ENSMUSG00000018800 | Abca5 | ATP-binding cassette, sub-family A (ABC1), member 5 | 1.92 |
| ENSMUSG00000024727 | Trpm6 | transient receptor potential cation channel, subfamily M, member 6 | 1.91 |
| ENSMUSG00000040430 | Pitpnc1 | phosphatidylinositol transfer protein, cytoplasmic 1 | 1.91 |
| ENSMUSG00000040836 | Gpr161 | G protein-coupled receptor 161 | 1.91 |
| ENSMUSG00000042306 | S100a14 | S100 calcium binding protein A14 | 1.89 |
| ENSMUSG00000038775 | Vill | villin-like | 1.89 |
| ENSMUSG00000006411 | Nectin4 | nectin cell adhesion molecule 4 | 1.87 |
| ENSMUSG00000028270 | Gbp2 | guanylate binding protein 2 | 1.86 |
| ENSMUSG00000043461 | Sptssb | serine palmitoyltransferase, small subunit B | 1.85 |
| ENSMUSG00000026167 | Wnt10a | wingless-type MMTV integration site family, member 10A | 1.85 |
| ENSMUSG00000024501 | Dpysl3 | dihydropyrimidinase-like 3 | 1.84 |
| ENSMUSG00000025921 | Rdh10 | retinol dehydrogenase 10 (all-trans) | 1.83 |
| ENSMUSG00000041886 | Macc1 | metastasis associated in colon cancer 1 | 1.83 |
| ENSMUSG00000072235 | Tuba1a | tubulin, alpha 1A | 1.83 |
| ENSMUSG00000039232 | Stx11 | syntaxin 11 | 1.83 |
| ENSMUSG00000004044 | Cavin1 | caveolae associated 1 | 1.82 |
| ENSMUSG00000028464 | Tpm2 | tropomyosin 2, beta | 1.82 |
| ENSMUSG00000026678 | Rgs5 | regulator of G-protein signaling 5 | 1.81 |
| ENSMUSG00000047344 | Lancl3 | LanC lantibiotic synthetase component C-like 3 (bacterial) | 1.80 |
| ENSMUSG00000025321 | Itgb8 | integrin beta 8 | 1.80 |
| ENSMUSG00000030259 | Rassf8 | Ras association (RalGDS/AF-6) domain family (N-terminal) member 8 | 1.80 |
| ENSMUSG00000025330 | Padi4 | peptidyl arginine deiminase, type IV | 1.79 |
| ENSMUSG00000072620 | Slfn2 | schlafen 2 | 1.77 |
| ENSMUSG00000020121 | Srgap1 | SLIT-ROBO Rho GTPase activating protein 1 | 1.76 |
| ENSMUSG00000027858 | Tspan2 | tetraspanin 2 | 1.76 |
| ENSMUSG00000044646 | Zbtb7c | zinc finger and BTB domain containing 7C | 1.75 |
| ENSMUSG00000039934 | Gsap | gamma-secretase activating protein | 1.73 |
| ENSMUSG00000009097 | Tbx1 | T-box 1 | 1.73 |
| ENSMUSG00000021493 | Pdlim7 | PDZ and LIM domain 7 | 1.72 |
| ENSMUSG00000024401 | Tnf | tumor necrosis factor | 1.72 |
| ENSMUSG00000029761 | Cald1 | caldesmon 1 | 1.72 |
| ENSMUSG00000027210 | Meis2 | Meis homeobox 2 | 1.71 |
| ENSMUSG00000022178 | Ajuba | ajuba LIM protein | 1.70 |
| ENSMUSG00000039457 | Ppl | periplakin | 1.70 |
| ENSMUSG00000017723 | Wfdc2 | WAP four-disulfide core domain 2 | 1.70 |
| ENSMUSG00000037613 | Tnfrsf23 | tumor necrosis factor receptor superfamily, member 23 | 1.69 |
| ENSMUSG00000032186 | Tmod2 | tropomodulin 2 | 1.68 |
| ENSMUSG00000031372 | Trex2 | three prime repair exonuclease 2 | 1.68 |
| ENSMUSG00000021701 | Plk2 | polo like kinase 2 | 1.67 |
| ENSMUSG00000025432 | Avil | advillin | 1.66 |
| ENSMUSG00000046223 | Plaur | plasminogen activator, urokinase receptor | 1.66 |
| ENSMUSG00000006403 | Adamts4 | a disintegrin-like and metallopeptidase (reprolysin type) with thrombospondin type 1 | 1.66 |
| ENSMUSG00000115009 | G930009F23Rik | RIKEN cDNA G930009F23 gene | 1.66 |
| ENSMUSG00000031328 | Flna | filamin, alpha | 1.65 |
| ENSMUSG00000037995 | Igsf9 | immunoglobulin superfamily, member 9 | 1.65 |
| ENSMUSG00000052105 | Mtcl1 | microtubule crosslinking factor 1 | 1.65 |
| ENSMUSG00000006777 | Krt23 | keratin 23 | 1.63 |
| ENSMUSG00000023008 | Fmnl3 | formin-like 3 | 1.63 |
| ENSMUSG00000029762 | Akr1b8 | aldo-keto reductase family 1, member B8 | 1.63 |
| ENSMUSG00000062661 | Ncs1 | neuronal calcium sensor 1 | 1.63 |
| ENSMUSG00000051043 | Gprc5c | G protein-coupled receptor, family C, group 5, member C | 1.63 |
| ENSMUSG00000090877 | Hspa1b | heat shock protein 1B | 1.63 |
| ENSMUSG00000026604 | Ptpn14 | protein tyrosine phosphatase, non-receptor type 14 | 1.63 |
| ENSMUSG00000018920 | Cxcl16 | chemokine (C-X-C motif) ligand 16 | 1.62 |
| ENSMUSG00000020668 | Kif3c | kinesin family member 3C | 1.62 |
| ENSMUSG00000027737 | Slc7a11 | solute carrier family 7 (cationic amino acid transporter, y+ system), member 11 | 1.61 |

| | | | |
|---|---|---|---|
| ENSMUSG00000116919 | AC165151.1 | conserved helix-loop-helix ubiquitous kinase (Chuk) pseudogene | 1.60 |
| ENSMUSG00000002980 | Bcam | basal cell adhesion molecule | 1.59 |
| ENSMUSG00000023885 | Thbs2 | thrombospondin 2 | 1.59 |
| ENSMUSG00000048960 | Prex2 | phosphatidylinositol-3,4,5-trisphosphate-dependent Rac exchange factor 2 | 1.58 |
| ENSMUSG00000073599 | Ecscr | endothelial cell surface expressed chemotaxis and apoptosis regulator | 1.57 |
| ENSMUSG00000020427 | Igfbp3 | insulin-like growth factor binding protein 3 | 1.57 |
| ENSMUSG00000079018 | Ly6c1 | lymphocyte antigen 6 complex, locus C1 | 1.57 |
| ENSMUSG00000032839 | Trpc1 | transient receptor potential cation channel, subfamily C, member 1 | 1.56 |
| ENSMUSG00000038415 | Foxq1 | forkhead box Q1 | 1.54 |
| ENSMUSG00000026479 | Lamc2 | laminin, gamma 2 | 1.54 |
| ENSMUSG00000046589 | Lrrc8e | leucine rich repeat containing 8 family, member E | 1.54 |
| ENSMUSG00000026639 | Lamb3 | laminin, beta 3 | 1.53 |
| ENSMUSG00000027978 | Prss12 | protease, serine 12 neurotrypsin (motopsin) | 1.53 |
| ENSMUSG00000025931 | Paqr8 | progestin and adipoQ receptor family member VIII | 1.53 |
| ENSMUSG00000103738 | Gm37652 | predicted gene, 37652 | 1.53 |
| ENSMUSG00000017897 | Eya2 | EYA transcriptional coactivator and phosphatase 2 | 1.52 |
| ENSMUSG00000048078 | Tenm4 | teneurin transmembrane protein 4 | 1.51 |
| ENSMUSG00000067006 | Serpinb5 | serine (or cysteine) peptidase inhibitor, clade B, member 5 | 1.50 |
| ENSMUSG00000032496 | Ltf | lactotransferrin | 1.50 |
| ENSMUSG00000074364 | Ehd2 | EH-domain containing 2 | 1.48 |
| ENSMUSG00000075602 | Ly6a | lymphocyte antigen 6 complex, locus A | 1.47 |
| ENSMUSG00000035678 | Tnfsf9 | tumor necrosis factor (ligand) superfamily, member 9 | 1.47 |
| ENSMUSG00000021559 | Dapk1 | death associated protein kinase 1 | 1.46 |
| ENSMUSG00000022371 | Col14a1 | collagen, type XIV, alpha 1 | 1.46 |
| ENSMUSG00000035105 | Egln3 | egl-9 family hypoxia-inducible factor 3 | 1.45 |
| ENSMUSG00000040543 | Pitpnm3 | PITPNM family member 3 | 1.45 |
| ENSMUSG00000027907 | S100a11 | S100 calcium binding protein A11 | 1.44 |
| ENSMUSG00000090394 | 4930523C07Rik | RIKEN cDNA 4930523C07 gene | 1.43 |
| ENSMUSG00000041608 | Entpd3 | ectonucleoside triphosphate diphosphohydrolase 3 | 1.43 |
| ENSMUSG00000004891 | Nes | nestin | 1.43 |
| ENSMUSG00000003541 | Ier3 | immediate early response 3 | 1.41 |
| ENSMUSG00000026655 | Fam107b | family with sequence similarity 107, member B | 1.40 |
| ENSMUSG00000061751 | Kalrn | kalirin, RhoGEF kinase | 1.40 |
| ENSMUSG00000041570 | Camsap2 | calmodulin regulated spectrin-associated protein family, member 2 | 1.40 |
| ENSMUSG00000053617 | Sh3pxd2a | SH3 and PX domains 2A | 1.39 |
| ENSMUSG00000031790 | Mmp15 | matrix metallopeptidase 15 | 1.38 |
| ENSMUSG00000040624 | Plekhg1 | pleckstrin homology domain containing, family G (with RhoGef domain) member 1 | 1.38 |
| ENSMUSG00000030759 | Far1 | fatty acyl CoA reductase 1 | 1.38 |
| ENSMUSG00000020773 | Trim47 | tripartite motif-containing 47 | 1.38 |
| ENSMUSG00000032875 | Arhgef17 | Rho guanine nucleotide exchange factor (GEF) 17 | 1.36 |
| ENSMUSG00000062488 | Ifit3b | interferon-induced protein with tetratricopeptide repeats 3B | 1.36 |
| ENSMUSG00000032717 | Mdfi | MyoD family inhibitor | 1.35 |
| ENSMUSG00000028073 | Pear1 | platelet endothelial aggregation receptor 1 | 1.35 |
| ENSMUSG00000025372 | Baiap2 | brain-specific angiogenesis inhibitor 1-associated protein 2 | 1.34 |
| ENSMUSG00000021477 | Ctsl | cathepsin L | 1.34 |
| ENSMUSG00000034595 | Ppp1r18 | protein phosphatase 1, regulatory subunit 18 | 1.34 |
| ENSMUSG00000023039 | Krt7 | keratin 7 | 1.33 |
| ENSMUSG00000039831 | Arhgap29 | Rho GTPase activating protein 29 | 1.33 |
| ENSMUSG00000034825 | Nrip3 | nuclear receptor interacting protein 3 | 1.33 |
| ENSMUSG00000035697 | Arhgap45 | Rho GTPase activating protein 45 | 1.33 |
| ENSMUSG00000029417 | Cxcl9 | chemokine (C-X-C motif) ligand 9 | 1.33 |
| ENSMUSG00000020230 | Prmt2 | protein arginine N-methyltransferase 2 | 1.32 |
| ENSMUSG00000036832 | Lpar3 | lysophosphatidic acid receptor 3 | 1.32 |
| ENSMUSG00000026335 | Pam | peptidylglycine alpha-amidating monooxygenase | 1.31 |
| ENSMUSG00000089809 | Rasgef1b | RasGEF domain family, member 1B | 1.31 |
| ENSMUSG00000021281 | Tnfaip2 | tumor necrosis factor, alpha-induced protein 2 | 1.31 |
| ENSMUSG00000051951 | Xkr4 | X-linked Kx blood group related 4 | 1.30 |
| ENSMUSG00000017765 | Slc12a4 | solute carrier family 12, member 4 | 1.29 |
| ENSMUSG00000031925 | Maml2 | mastermind like transcriptional coactivator 2 | 1.29 |
| ENSMUSG00000079339 | Ifit1bl1 | interferon induced protein with tetratricpeptide repeats 1B like 1 | 1.29 |
| ENSMUSG00000102460 | Gm38197 | predicted gene, 38197 | 1.29 |
| ENSMUSG00000082361 | Btc | betacellulin, epidermal growth factor family member | 1.28 |
| ENSMUSG00000024810 | Il33 | interleukin 33 | 1.28 |
| ENSMUSG00000039364 | Sectm1b | secreted and transmembrane 1B | 1.27 |

| | | | |
|---|---|---|---|
| ENSMUSG00000048058 | Ldlrad3 | low density lipoprotein receptor class A domain containing 3 | 1.27 |
| ENSMUSG00000035818 | Plekhs1 | pleckstrin homology domain containing, family S member 1 | 1.27 |
| ENSMUSG00000000489 | Pdgfb | platelet derived growth factor, B polypeptide | 1.27 |
| ENSMUSG00000020181 | Nav3 | neuron navigator 3 | 1.27 |
| ENSMUSG00000013076 | Amotl1 | angiomotin-like 1 | 1.26 |
| ENSMUSG00000025287 | Acot9 | acyl-CoA thioesterase 9 | 1.26 |
| ENSMUSG00000040711 | Sh3pxd2b | SH3 and PX domains 2B | 1.26 |
| ENSMUSG00000028378 | Ptgr1 | prostaglandin reductase 1 | 1.25 |
| ENSMUSG00000074796 | Slc4a11 | solute carrier family 4, sodium bicarbonate transporter-like, member 11 | 1.25 |
| ENSMUSG00000041390 | Mdfic | MyoD family inhibitor domain containing | 1.25 |
| ENSMUSG00000032724 | Abtb2 | ankyrin repeat and BTB (POZ) domain containing 2 | 1.24 |
| ENSMUSG00000046694 | Tent5b | terminal nucleotidyltransferase 5B | 1.24 |
| ENSMUSG00000110218 | Gm20219 | predicted gene, 20219 | 1.23 |
| ENSMUSG00000022176 | Rem2 | rad and gem related GTP binding protein 2 | 1.23 |
| ENSMUSG00000030352 | Tspan9 | tetraspanin 9 | 1.23 |
| ENSMUSG00000021108 | Prkch | protein kinase C, eta | 1.23 |
| ENSMUSG00000025278 | Flnb | filamin, beta | 1.23 |
| ENSMUSG00000000706 | Btn1a1 | butyrophilin, subfamily 1, member A1 | 1.23 |
| ENSMUSG00000025161 | Slc16a3 | solute carrier family 16 (monocarboxylic acid transporters), member 3 | 1.22 |
| ENSMUSG00000022636 | Alcam | activated leukocyte cell adhesion molecule | 1.21 |
| ENSMUSG00000024912 | Fosl1 | fos-like antigen 1 | 1.21 |
| ENSMUSG00000009628 | Tex15 | testis expressed gene 15 | 1.21 |
| ENSMUSG00000025383 | Il23a | interleukin 23, alpha subunit p19 | 1.21 |
| ENSMUSG00000024302 | Dtna | dystrobrevin alpha | 1.21 |
| ENSMUSG00000027381 | Bcl2l11 | BCL2-like 11 (apoptosis facilitator) | 1.20 |
| ENSMUSG00000027257 | Pacsin3 | protein kinase C and casein kinase substrate in neurons 3 | 1.20 |
| ENSMUSG00000019647 | Sema6a | sema domain, transmembrane domain (TM), and cytoplasmic domain, (semaphorin) | 1.20 |
| ENSMUSG00000028435 | Aqp3 | aquaporin 3 | 1.20 |
| ENSMUSG00000046731 | Kctd11 | potassium channel tetramerisation domain containing 11 | 1.19 |
| ENSMUSG00000022665 | Ccdc80 | coiled-coil domain containing 80 | 1.19 |
| ENSMUSG00000075033 | Nxpe3 | neurexophilin and PC-esterase domain family, member 3 | 1.18 |
| ENSMUSG00000047735 | Samd9l | sterile alpha motif domain containing 9-like | 1.18 |
| ENSMUSG00000060923 | Acyp2 | acylphosphatase 2, muscle type | 1.18 |
| ENSMUSG00000021360 | Gcnt2 | glucosaminyl (N-acetyl) transferase 2, I-branching enzyme | 1.17 |
| ENSMUSG00000041189 | Chrnb1 | cholinergic receptor, nicotinic, beta polypeptide 1 (muscle) | 1.17 |
| ENSMUSG00000019853 | Hebp2 | heme binding protein 2 | 1.16 |
| ENSMUSG00000022865 | Cxadr | coxsackie virus and adenovirus receptor | 1.16 |
| ENSMUSG00000038387 | Rras | related RAS viral (r-ras) oncogene | 1.16 |
| ENSMUSG00000042510 | AA986860 | expressed sequence AA986860 | 1.16 |
| ENSMUSG00000032643 | Fhl3 | four and a half LIM domains 3 | 1.16 |
| ENSMUSG00000069255 | Dusp22 | dual specificity phosphatase 22 | 1.16 |
| ENSMUSG00000037860 | Aim2 | absent in melanoma 2 | 1.14 |
| ENSMUSG00000029377 | Ereg | epiregulin | 1.14 |
| ENSMUSG00000018907 | Alox12e | arachidonate lipoxygenase, epidermal | 1.14 |
| ENSMUSG00000029822 | Osbpl3 | oxysterol binding protein-like 3 | 1.13 |
| ENSMUSG00000034459 | Ifit1 | interferon-induced protein with tetratricopeptide repeats 1 | 1.13 |
| ENSMUSG00000042834 | Nrep | neuronal regeneration related protein | 1.13 |
| ENSMUSG00000049807 | Arhgap23 | Rho GTPase activating protein 23 | 1.12 |
| ENSMUSG00000037606 | Osbpl5 | oxysterol binding protein-like 5 | 1.12 |
| ENSMUSG00000024940 | Ltbp3 | latent transforming growth factor beta binding protein 3 | 1.12 |
| ENSMUSG00000069601 | Ank3 | ankyrin 3, epithelial | 1.11 |
| ENSMUSG00000031844 | Hsd17b2 | hydroxysteroid (17-beta) dehydrogenase 2 | 1.11 |
| ENSMUSG00000037419 | Endod1 | endonuclease domain containing 1 | 1.10 |
| ENSMUSG00000001473 | Tubb6 | tubulin, beta 6 class V | 1.10 |
| ENSMUSG00000055976 | Cldn23 | claudin 23 | 1.10 |
| ENSMUSG00000034295 | Fhod3 | formin homology 2 domain containing 3 | 1.10 |
| ENSMUSG00000020023 | Tmcc3 | transmembrane and coiled coil domains 3 | 1.09 |
| ENSMUSG00000006435 | Neurl1a | neuralized E3 ubiquitin protein ligase 1A | 1.09 |
| ENSMUSG00000029832 | Nfe2l3 | nuclear factor, erythroid derived 2, like 3 | 1.08 |
| ENSMUSG00000027456 | Sdcbp2 | syndecan binding protein (syntenin) 2 | 1.08 |
| ENSMUSG00000020674 | Pxdn | peroxidasin | 1.08 |
| ENSMUSG00000027860 | Vangl1 | VANGL planar cell polarity 1 | 1.07 |
| ENSMUSG00000035547 | Capn5 | calpain 5 | 1.07 |
| ENSMUSG00000029378 | Areg | amphiregulin | 1.07 |

| | | | |
|---|---|---|---|
| ENSMUSG00000038295 | Atg9b | autophagy related 9B | 1.06 |
| ENSMUSG00000028631 | Kcnq4 | potassium voltage-gated channel, subfamily Q, member 4 | 1.06 |
| ENSMUSG00000114996 | Gm48958 | predicted gene, 48958 | 1.06 |
| ENSMUSG00000026797 | Stxbp1 | syntaxin binding protein 1 | 1.05 |
| ENSMUSG00000031441 | Atp11a | ATPase, class VI, type 11A | 1.05 |
| ENSMUSG00000053702 | Nebl | nebulette | 1.05 |
| ENSMUSG00000068566 | Myadm | myeloid-associated differentiation marker | 1.04 |
| ENSMUSG00000036528 | Ppfibp2 | PTPRF interacting protein, binding protein 2 (liprin beta 2) | 1.04 |
| ENSMUSG00000024777 | Ppp2r5b | protein phosphatase 2, regulatory subunit B', beta | 1.04 |
| ENSMUSG00000038400 | Pmepa1 | prostate transmembrane protein, androgen induced 1 | 1.04 |
| ENSMUSG00000049858 | Suox | sulfite oxidase | 1.04 |
| ENSMUSG00000031367 | Ap1s2 | adaptor-related protein complex 1, sigma 2 subunit | 1.04 |
| ENSMUSG00000104415 | Gm37069 | predicted gene, 37069 | 1.04 |
| ENSMUSG00000030342 | Cd9 | CD9 antigen | 1.03 |
| ENSMUSG00000048911 | Rnf24 | ring finger protein 24 | 1.03 |
| ENSMUSG00000026961 | Lrrc26 | leucine rich repeat containing 26 | 1.03 |
| ENSMUSG00000074006 | Omp | olfactory marker protein | 1.03 |
| ENSMUSG00000027602 | Map1lc3a | microtubule-associated protein 1 light chain 3 alpha | 1.02 |
| ENSMUSG00000040447 | Spns2 | spinster homolog 2 | 1.02 |
| ENSMUSG00000069114 | Zbtb10 | zinc finger and BTB domain containing 10 | 1.02 |
| ENSMUSG00000026478 | Lamc1 | laminin, gamma 1 | 1.01 |
| ENSMUSG00000026796 | Fam129b | family with sequence similarity 129, member B | 1.01 |
| ENSMUSG00000049866 | Arl4c | ADP-ribosylation factor-like 4C | 1.01 |
| ENSMUSG00000057193 | Slc44a2 | solute carrier family 44, member 2 | 1.01 |
| ENSMUSG00000038235 | F11r | F11 receptor | 1.01 |
| ENSMUSG00000112198 | Gm4065 | predicted gene 4065 | 1.01 |
| ENSMUSG00000026971 | Itgb6 | integrin beta 6 | 1.00 |
| ENSMUSG00000079547 | H2-DMb1 | histocompatibility 2, class II, locus Mb1 | -1.00 |
| ENSMUSG00000057604 | Lmcd1 | LIM and cysteine-rich domains 1 | -1.00 |
| ENSMUSG00000102380 | Gm38140 | predicted gene, 38140 | -1.00 |
| ENSMUSG00000027452 | Acss1 | acyl-CoA synthetase short-chain family member 1 | -1.01 |
| ENSMUSG00000028415 | Spink4 | serine peptidase inhibitor, Kazal type 4 | -1.01 |
| ENSMUSG00000038255 | Neurod2 | neurogenic differentiation 2 | -1.01 |
| ENSMUSG00000025738 | Fbxl16 | F-box and leucine-rich repeat protein 16 | -1.02 |
| ENSMUSG00000031453 | Rasa3 | RAS p21 protein activator 3 | -1.03 |
| ENSMUSG00000074653 | Lrrc31 | leucine rich repeat containing 31 | -1.03 |
| ENSMUSG00000095328 | Defa-ps6 | defensin, alpha, pseudogene 6 | -1.03 |
| ENSMUSG00000055197 | Fev | FEV (ETS oncogene family) | -1.03 |
| ENSMUSG00000060586 | H2-Eb1 | histocompatibility 2, class II antigen E beta | -1.04 |
| ENSMUSG00000031891 | Hsd11b2 | hydroxysteroid 11-beta dehydrogenase 2 | -1.04 |
| ENSMUSG00000051079 | Rgs13 | regulator of G-protein signaling 13 | -1.04 |
| ENSMUSG00000036006 | Ripor2 | RHO family interacting cell polarization regulator 2 | -1.05 |
| ENSMUSG00000024975 | Pdcd4 | programmed cell death 4 | -1.06 |
| ENSMUSG00000046613 | Vwa5b2 | von Willebrand factor A domain containing 5B2 | -1.06 |
| ENSMUSG00000003545 | Fosb | FBJ osteosarcoma oncogene B | -1.07 |
| ENSMUSG00000063895 | Nupl1 | nucleoporin like 1 | -1.07 |
| ENSMUSG00000063142 | Kcnma1 | potassium large conductance calcium-activated channel, subfamily M, alpha membe | -1.08 |
| ENSMUSG00000021830 | Txndc16 | thioredoxin domain containing 16 | -1.09 |
| ENSMUSG00000108961 | Gm32540 | predicted gene, 32540 | -1.09 |
| ENSMUSG00000030934 | Oat | ornithine aminotransferase | -1.10 |
| ENSMUSG00000022816 | Fstl1 | follistatin-like 1 | -1.11 |
| ENSMUSG00000097924 | A730020E08Rik | RIKEN cDNA A730020E08 gene | -1.11 |
| ENSMUSG00000043487 | Acot6 | acyl-CoA thioesterase 6 | -1.12 |
| ENSMUSG00000026610 | Esrrg | estrogen-related receptor gamma | -1.13 |
| ENSMUSG00000078586 | Gm11735 | predicted gene 11735 | -1.13 |
| ENSMUSG00000033705 | Stard9 | START domain containing 9 | -1.14 |
| ENSMUSG00000009378 | Slc16a12 | solute carrier family 16 (monocarboxylic acid transporters), member 12 | -1.15 |
| ENSMUSG00000033715 | Akr1c14 | aldo-keto reductase family 1, member C14 | -1.15 |
| ENSMUSG00000038745 | Nlrp6 | NLR family, pyrin domain containing 6 | -1.16 |
| ENSMUSG00000031451 | Gas6 | growth arrest specific 6 | -1.16 |
| ENSMUSG00000052026 | Slc6a7 | solute carrier family 6 (neurotransmitter transporter, L-proline), member 7 | -1.16 |
| ENSMUSG00000058740 | Kcnt1 | potassium channel, subfamily T, member 1 | -1.16 |
| ENSMUSG00000097494 | 4933406C10Rik | RIKEN cDNA 4933406C10 gene | -1.17 |
| ENSMUSG00000028838 | Extl1 | exostoses (multiple)-like 1 | -1.18 |

| | | | |
|---|---|---|---|
| ENSMUSG00000024225 | Clps | colipase, pancreatic | -1.18 |
| ENSMUSG00000074715 | Ccl28 | chemokine (C-C motif) ligand 28 | -1.19 |
| ENSMUSG00000060208 | Defa17 | defensin, alpha, 17 | -1.19 |
| ENSMUSG00000026398 | Nr5a2 | nuclear receptor subfamily 5, group A, member 2 | -1.21 |
| ENSMUSG00000002346 | Slc25a42 | solute carrier family 25, member 42 | -1.21 |
| ENSMUSG00000040412 | 5330417C22Rik | RIKEN cDNA 5330417C22 gene | -1.25 |
| ENSMUSG00000056553 | Ptprn2 | protein tyrosine phosphatase, receptor type, N polypeptide 2 | -1.25 |
| ENSMUSG00000109936 | Gm45889 | predicted gene 45889 | -1.25 |
| ENSMUSG00000033740 | St18 | suppression of tumorigenicity 18 | -1.25 |
| ENSMUSG00000110170 | St6galnac2 | ST6 (alpha-N-acetyl-neuraminyl-2,3-beta-galactosyl-1,3)-N-acetylgalactosaminide alp | -1.25 |
| ENSMUSG00000000416 | Cttnbp2 | cortactin binding protein 2 | -1.26 |
| ENSMUSG00000026442 | Nfasc | neurofascin | -1.26 |
| ENSMUSG00000005089 | Slc1a2 | solute carrier family 1 (glial high affinity glutamate transporter), member 2 | -1.26 |
| ENSMUSG00000110622 | Iqcn | IQ motif containing N | -1.26 |
| ENSMUSG00000079180 | Mptx2 | mucosal pentraxin 2 | -1.27 |
| ENSMUSG00000043648 | Pld6 | phospholipase D family, member 6 | -1.27 |
| ENSMUSG00000045709 | Smkr-ps | smal lysine rich protein 1, pseudogene | -1.28 |
| ENSMUSG00000031173 | Otc | ornithine transcarbamylase | -1.29 |
| ENSMUSG00000106944 | Gm43843 | predicted gene 43843 | -1.29 |
| ENSMUSG00000040706 | Agmat | agmatine ureohydrolase (agmatinase) | -1.30 |
| ENSMUSG00000028327 | Stra6l | STRA6-like | -1.30 |
| ENSMUSG00000048027 | Rgmb | repulsive guidance molecule family member B | -1.31 |
| ENSMUSG00000022018 | Rgcc | regulator of cell cycle | -1.32 |
| ENSMUSG00000025790 | Slco3a1 | solute carrier organic anion transporter family, member 3a1 | -1.32 |
| ENSMUSG00000033066 | Gas7 | growth arrest specific 7 | -1.32 |
| ENSMUSG00000023886 | Smoc2 | SPARC related modular calcium binding 2 | -1.33 |
| ENSMUSG00000025196 | Cpn1 | carboxypeptidase N, polypeptide 1 | -1.33 |
| ENSMUSG00000022650 | Retnlb | resistin like beta | -1.33 |
| ENSMUSG00000058618 | AY761184 | cDNA sequence AY761184 | -1.33 |
| ENSMUSG00000009394 | Syn2 | synapsin II | -1.33 |
| ENSMUSG00000025082 | Vwa2 | von Willebrand factor A domain containing 2 | -1.34 |
| ENSMUSG00000033721 | Vav3 | vav 3 oncogene | -1.34 |
| ENSMUSG00000004655 | Aqp1 | aquaporin 1 | -1.36 |
| ENSMUSG00000025867 | Cplx2 | complexin 2 | -1.36 |
| ENSMUSG00000052271 | Bhlha15 | basic helix-loop-helix family, member a15 | -1.38 |
| ENSMUSG00000079559 | Colca2 | COLCA2 homolog | -1.38 |
| ENSMUSG00000005268 | Prlr | prolactin receptor | -1.39 |
| ENSMUSG00000074437 | Defa29 | defensin, alpha, 29 | -1.39 |
| ENSMUSG00000090105 | Gm15890 | predicted gene 15890 | -1.39 |
| ENSMUSG00000070337 | Gpr179 | G protein-coupled receptor 179 | -1.40 |
| ENSMUSG00000032452 | Clstn2 | calsyntenin 2 | -1.40 |
| ENSMUSG00000038058 | Nod1 | nucleotide-binding oligomerization domain containing 1 | -1.43 |
| ENSMUSG00000022504 | Ciita | class II transactivator | -1.43 |
| ENSMUSG00000032561 | Acpp | acid phosphatase, prostate | -1.44 |
| ENSMUSG00000041193 | Pla2g5 | phospholipase A2, group V | -1.45 |
| ENSMUSG00000010021 | Kif19a | kinesin family member 19A | -1.45 |
| ENSMUSG00000084939 | Gm830 | predicted gene 830 | -1.46 |
| ENSMUSG00000044694 | 2010007H06Rik | RIKEN cDNA 2010007H06 gene | -1.47 |
| ENSMUSG00000005360 | Slc1a3 | solute carrier family 1 (glial high affinity glutamate transporter), member 3 | -1.48 |
| ENSMUSG00000019900 | Rfx6 | regulatory factor X, 6 | -1.51 |
| ENSMUSG00000020140 | Lgr5 | leucine rich repeat containing G protein coupled receptor 5 | -1.52 |
| ENSMUSG00000033717 | Adra2a | adrenergic receptor, alpha 2a | -1.53 |
| ENSMUSG00000026786 | Apbb1ip | amyloid beta (A4) precursor protein-binding, family B, member 1 interacting protein | -1.56 |
| ENSMUSG00000112816 | Gm48798 | predicted gene, 48798 | -1.56 |
| ENSMUSG00000025991 | Cps1 | carbamoyl-phosphate synthetase 1 | -1.60 |
| ENSMUSG00000028602 | Tnfrsf8 | tumor necrosis factor receptor superfamily, member 8 | -1.60 |
| ENSMUSG00000040528 | Milr1 | mast cell immunoglobulin like receptor 1 | -1.60 |
| ENSMUSG00000035561 | Aldh1b1 | aldehyde dehydrogenase 1 family, member B1 | -1.61 |
| ENSMUSG00000026489 | Coq8a | coenzyme Q8A | -1.61 |
| ENSMUSG00000073424 | Cyp4f15 | cytochrome P450, family 4, subfamily f, polypeptide 15 | -1.61 |
| ENSMUSG00000086513 | 9130208D14Rik | RIKEN cDNA 9130208D14 gene | -1.65 |
| ENSMUSG00000027359 | Slc27a2 | solute carrier family 27 (fatty acid transporter), member 2 | -1.66 |
| ENSMUSG00000031428 | Zcchc18 | zinc finger, CCHC domain containing 18 | -1.67 |
| ENSMUSG00000021719 | Rgs7bp | regulator of G-protein signalling 7 binding protein | -1.68 |

| | | | |
|---|---|---|---|
| ENSMUSG00000095649 | Gm8979 | predicted gene 8979 | -1.75 |
| ENSMUSG00000029445 | Hpd | 4-hydroxyphenylpyruvic acid dioxygenase | -1.76 |
| ENSMUSG00000030207 | Fam234b | family with sequence similarity 234, member B | -1.77 |
| ENSMUSG00000018341 | Il12rb2 | interleukin 12 receptor, beta 2 | -1.80 |
| ENSMUSG00000000031 | H19 | H19, imprinted maternally expressed transcript | -1.81 |
| ENSMUSG00000037953 | A4gnt | alpha-1,4-N-acetylglucosaminyltransferase | -1.81 |
| ENSMUSG00000109685 | Gm45912 | predicted gene 45912 | -1.85 |
| ENSMUSG00000023236 | Scg5 | secretogranin V | -1.86 |
| ENSMUSG00000117254 | AC171111.1 | novel transcript | -1.87 |
| ENSMUSG00000066687 | Zbtb16 | zinc finger and BTB domain containing 16 | -1.89 |
| ENSMUSG00000036585 | Fgf1 | fibroblast growth factor 1 | -1.95 |
| ENSMUSG00000036526 | Card11 | caspase recruitment domain family, member 11 | -1.99 |
| ENSMUSG00000024411 | Aqp4 | aquaporin 4 | -2.00 |
| ENSMUSG00000023828 | Slc22a3 | solute carrier family 22 (organic cation transporter), member 3 | -2.08 |
| ENSMUSG00000041577 | Prelp | proline arginine-rich end leucine-rich repeat | -2.09 |
| ENSMUSG00000034472 | Rasd2 | RASD family, member 2 | -2.09 |
| ENSMUSG00000000402 | Egfl6 | EGF-like-domain, multiple 6 | -2.31 |
| ENSMUSG00000110266 | Gm32742 | predicted gene, 32742 | -2.31 |
| ENSMUSG00000036422 | Pcdh8 | protocadherin 8 | -2.60 |
| ENSMUSG00000022026 | Olfm4 | olfactomedin 4 | -2.76 |
| ENSMUSG00000038765 | Lmx1b | LIM homeobox transcription factor 1 beta | -2.92 |

**RFP+Trop2+ vs. differentiated cells**

| Ensemble gene code | Symbol | Gene name | logFC |
|---|---|---|---|
| ENSMUSG00000029304 | Spp1 | secreted phosphoprotein 1 | 5.80 |
| ENSMUSG00000051397 | Tacstd2 | tumor-associated calcium signal transducer 2 | 5.16 |
| ENSMUSG00000041523 | Upk2 | uroplakin 2 | 4.78 |
| ENSMUSG00000022754 | Tmem45a | transmembrane protein 45a | 4.67 |
| ENSMUSG00000059668 | Krt4 | keratin 4 | 4.65 |
| ENSMUSG00000037060 | Cavin3 | caveolae associated 3 | 4.58 |
| ENSMUSG00000024087 | Cyp1b1 | cytochrome P450, family 1, subfamily b, polypeptide 1 | 4.51 |
| ENSMUSG00000021950 | Anxa8 | annexin A8 | 4.23 |
| ENSMUSG00000024659 | Anxa1 | annexin A1 | 4.20 |
| ENSMUSG00000029648 | Flt1 | FMS-like tyrosine kinase 1 | 4.20 |
| ENSMUSG00000031430 | Vsig1 | V-set and immunoglobulin domain containing 1 | 4.17 |
| ENSMUSG00000033377 | Palmd | palmdelphin | 3.89 |
| ENSMUSG00000022037 | Clu | clusterin | 3.82 |
| ENSMUSG00000079330 | Lemd1 | LEM domain containing 1 | 3.76 |
| ENSMUSG00000049128 | Ivl | involucrin | 3.75 |
| ENSMUSG00000091243 | Vgll3 | vestigial like family member 3 | 3.68 |
| ENSMUSG00000037124 | Trim58 | tripartite motif-containing 58 | 3.63 |
| ENSMUSG00000061517 | Sox21 | SRY (sex determining region Y)-box 21 | 3.58 |
| ENSMUSG00000032327 | Stra6 | stimulated by retinoic acid gene 6 | 3.57 |
| ENSMUSG00000058354 | Krt6a | keratin 6A | 3.40 |
| ENSMUSG00000037188 | Grhl3 | grainyhead like transcription factor 3 | 3.39 |
| ENSMUSG00000009097 | Tbx1 | T-box 1 | 3.34 |
| ENSMUSG00000039153 | Runx2 | runt related transcription factor 2 | 3.28 |
| ENSMUSG00000056632 | Dsg3 | desmoglein 3 | 3.27 |
| ENSMUSG00000027485 | Bpifb1 | BPI fold containing family B, member 1 | 3.23 |
| ENSMUSG00000020773 | Trim47 | tripartite motif-containing 47 | 3.21 |
| ENSMUSG00000030510 | Cers3 | ceramide synthase 3 | 3.19 |
| ENSMUSG00000097519 | 4930558J18Rik | RIKEN cDNA 4930558J18 gene | 3.17 |
| ENSMUSG00000022203 | Efs | embryonal Fyn-associated substrate | 3.09 |
| ENSMUSG00000051076 | Vtcn1 | V-set domain containing T cell activation inhibitor 1 | 2.93 |
| ENSMUSG00000034463 | Scara3 | scavenger receptor class A, member 3 | 2.91 |
| ENSMUSG00000063011 | Msln | mesothelin | 2.91 |
| ENSMUSG00000048915 | Efna5 | ephrin A5 | 2.91 |
| ENSMUSG00000041886 | Macc1 | metastasis associated in colon cancer 1 | 2.89 |
| ENSMUSG00000025473 | Adam8 | a disintegrin and metallopeptidase domain 8 | 2.86 |
| ENSMUSG00000022371 | Col14a1 | collagen, type XIV, alpha 1 | 2.85 |
| ENSMUSG00000063060 | Sox7 | SRY (sex determining region Y)-box 7 | 2.85 |
| ENSMUSG00000046623 | Gjb4 | gap junction protein, beta 4 | 2.80 |
| ENSMUSG00000035818 | Plekhs1 | pleckstrin homology domain containing, family S member 1 | 2.79 |
| ENSMUSG00000021838 | Samd4 | sterile alpha motif domain containing 4 | 2.78 |
| ENSMUSG00000082706 | Gm11663 | predicted gene 11663 | 2.77 |
| ENSMUSG00000074625 | Arhgap40 | Rho GTPase activating protein 40 | 2.76 |
| ENSMUSG00000042734 | Ttc9 | tetratricopeptide repeat domain 9 | 2.75 |
| ENSMUSG00000028236 | Sdr16c5 | short chain dehydrogenase/reductase family 16C, member 5 | 2.75 |
| ENSMUSG00000031097 | Tnni2 | troponin I, skeletal, fast 2 | 2.75 |
| ENSMUSG00000092586 | Ly6g6c | lymphocyte antigen 6 complex, locus G6C | 2.73 |
| ENSMUSG00000113168 | Gm7614 | predicted gene 7614 | 2.70 |
| ENSMUSG00000013338 | Fer1l4 | fer-1-like 4 (C. elegans) | 2.67 |
| ENSMUSG00000040488 | Ltbp4 | latent transforming growth factor beta binding protein 4 | 2.66 |
| ENSMUSG00000040543 | Pitpnm3 | PITPNM family member 3 | 2.63 |
| ENSMUSG00000046213 | Cym | chymosin | 2.60 |
| ENSMUSG00000079018 | Ly6c1 | lymphocyte antigen 6 complex, locus C1 | 2.52 |
| ENSMUSG00000085412 | Halr1 | Hoxa adjacent long noncoding RNA 1 | 2.51 |
| ENSMUSG00000031871 | Cdh5 | cadherin 5 | 2.46 |
| ENSMUSG00000031841 | Cdh13 | cadherin 13 | 2.42 |
| ENSMUSG00000028464 | Tpm2 | tropomyosin 2, beta | 2.41 |
| ENSMUSG00000107350 | Gm19610 | predicted gene, 19610 | 2.41 |
| ENSMUSG00000006411 | Nectin4 | nectin cell adhesion molecule 4 | 2.40 |
| ENSMUSG00000037833 | Sh2d4b | SH2 domain containing 4B | 2.40 |
| ENSMUSG00000021367 | Edn1 | endothelin 1 | 2.39 |
| ENSMUSG00000015134 | Aldh1a3 | aldehyde dehydrogenase family 1, subfamily A3 | 2.39 |
| ENSMUSG00000034573 | Ptpn13 | protein tyrosine phosphatase, non-receptor type 13 | 2.38 |
| ENSMUSG00000114996 | Gm48958 | predicted gene, 48958 | 2.38 |
| ENSMUSG00000041482 | Piezo2 | piezo-type mechanosensitive ion channel component 2 | 2.38 |

| | | | |
|---|---|---|---|
| ENSMUSG00000029371 | Cxcl5 | chemokine (C-X-C motif) ligand 5 | 2.35 |
| ENSMUSG00000026981 | Il1rn | interleukin 1 receptor antagonist | 2.33 |
| ENSMUSG00000029484 | Anxa3 | annexin A3 | 2.29 |
| ENSMUSG00000026065 | Slc9a4 | solute carrier family 9 (sodium/hydrogen exchanger), member 4 | 2.29 |
| ENSMUSG00000025902 | Sox17 | SRY (sex determining region Y)-box 17 | 2.27 |
| ENSMUSG00000030259 | Rassf8 | Ras association (RalGDS/AF-6) domain family (N-terminal) member 8 | 2.27 |
| ENSMUSG00000055333 | Fat2 | FAT atypical cadherin 2 | 2.25 |
| ENSMUSG00000046402 | Rbp1 | retinol binding protein 1, cellular | 2.23 |
| ENSMUSG00000026327 | Serpinb11 | serine (or cysteine) peptidase inhibitor, clade B (ovalbumin), member 11 | 2.23 |
| ENSMUSG00000001288 | Rarg | retinoic acid receptor, gamma | 2.22 |
| ENSMUSG00000037628 | Cdkn3 | cyclin-dependent kinase inhibitor 3 | 2.22 |
| ENSMUSG00000000706 | Btn1a1 | butyrophilin, subfamily 1, member A1 | 2.22 |
| ENSMUSG00000022178 | Ajuba | ajuba LIM protein | 2.20 |
| ENSMUSG00000028068 | Iqgap3 | IQ motif containing GTPase activating protein 3 | 2.18 |
| ENSMUSG00000082095 | Gm11991 | predicted gene 11991 | 2.18 |
| ENSMUSG00000031098 | Syt8 | synaptotagmin VIII | 2.17 |
| ENSMUSG00000025321 | Itgb8 | integrin beta 8 | 2.17 |
| ENSMUSG00000032254 | Kif23 | kinesin family member 23 | 2.16 |
| ENSMUSG00000022505 | Emp2 | epithelial membrane protein 2 | 2.16 |
| ENSMUSG00000013766 | Ly6g6e | lymphocyte antigen 6 complex, locus G6E | 2.16 |
| ENSMUSG00000031538 | Plat | plasminogen activator, tissue | 2.15 |
| ENSMUSG00000020427 | Igfbp3 | insulin-like growth factor binding protein 3 | 2.14 |
| ENSMUSG00000094248 | Hist1h2ao | histone cluster 1, H2ao | 2.10 |
| ENSMUSG00000031328 | Flna | filamin, alpha | 2.08 |
| ENSMUSG00000027375 | Mal | myelin and lymphocyte protein, T cell differentiation protein | 2.08 |
| ENSMUSG00000053007 | Creb5 | cAMP responsive element binding protein 5 | 2.08 |
| ENSMUSG00000038415 | Foxq1 | forkhead box Q1 | 2.06 |
| ENSMUSG00000040998 | Npnt | nephronectin | 2.06 |
| ENSMUSG00000021176 | Efcab11 | EF-hand calcium binding domain 11 | 2.05 |
| ENSMUSG00000051378 | Kif18b | kinesin family member 18B | 2.04 |
| ENSMUSG00000075602 | Ly6a | lymphocyte antigen 6 complex, locus A | 2.04 |
| ENSMUSG00000006398 | Cdc20 | cell division cycle 20 | 2.02 |
| ENSMUSG00000056888 | Glipr1 | GLI pathogenesis-related 1 (glioma) | 2.02 |
| ENSMUSG00000021822 | Plau | plasminogen activator, urokinase | 2.02 |
| ENSMUSG00000027861 | Casq2 | calsequestrin 2 | 2.02 |
| ENSMUSG00000027469 | Tpx2 | TPX2, microtubule-associated | 2.01 |
| ENSMUSG00000063727 | Tnfrsf11b | tumor necrosis factor receptor superfamily, member 11b (osteoprotegerin) | 2.01 |
| ENSMUSG00000044201 | Cdc25c | cell division cycle 25C | 2.01 |
| ENSMUSG00000029380 | Cxcl1 | chemokine (C-X-C motif) ligand 1 | 2.01 |
| ENSMUSG00000035365 | Parpbp | PARP1 binding protein | 2.00 |
| ENSMUSG00000037946 | Fgd3 | FYVE, RhoGEF and PH domain containing 3 | 2.00 |
| ENSMUSG00000027699 | Ect2 | ect2 oncogene | 1.99 |
| ENSMUSG00000020330 | Hmmr | hyaluronan mediated motility receptor (RHAMM) | 1.99 |
| ENSMUSG00000079553 | Kifc1 | kinesin family member C1 | 1.99 |
| ENSMUSG00000040711 | Sh3pxd2b | SH3 and PX domains 2B | 1.99 |
| ENSMUSG00000101355 | Hist1h3h | histone cluster 1, H3h | 1.99 |
| ENSMUSG00000032783 | Troap | trophinin associated protein | 1.98 |
| ENSMUSG00000020808 | Pimreg | PICALM interacting mitotic regulator | 1.98 |
| ENSMUSG00000017716 | Birc5 | baculoviral IAP repeat-containing 5 | 1.97 |
| ENSMUSG00000041498 | Kif14 | kinesin family member 14 | 1.96 |
| ENSMUSG00000003541 | Ier3 | immediate early response 3 | 1.95 |
| ENSMUSG00000028678 | Kif2c | kinesin family member 2C | 1.95 |
| ENSMUSG00000106296 | 4632404M16Rik | RIKEN cDNA 4632404M16 gene | 1.95 |
| ENSMUSG00000027715 | Ccna2 | cyclin A2 | 1.94 |
| ENSMUSG00000059900 | Tmem40 | transmembrane protein 40 | 1.94 |
| ENSMUSG00000024989 | Cep55 | centrosomal protein 55 | 1.93 |
| ENSMUSG00000003779 | Kif20a | kinesin family member 20A | 1.93 |
| ENSMUSG00000027379 | Bub1 | BUB1, mitotic checkpoint serine/threonine kinase | 1.93 |
| ENSMUSG00000024056 | Ndc80 | NDC80 kinetochore complex component | 1.92 |
| ENSMUSG00000027860 | Vangl1 | VANGL planar cell polarity 1 | 1.91 |
| ENSMUSG00000032875 | Arhgef17 | Rho guanine nucleotide exchange factor (GEF) 17 | 1.91 |
| ENSMUSG00000028175 | Depdc1a | DEP domain containing 1a | 1.91 |
| ENSMUSG00000041431 | Ccnb1 | cyclin B1 | 1.90 |
| ENSMUSG00000037544 | Dlgap5 | DLG associated protein 5 | 1.89 |
| ENSMUSG00000036777 | Anln | anillin, actin binding protein | 1.89 |
| ENSMUSG00000023348 | Trip6 | thyroid hormone receptor interactor 6 | 1.89 |
| ENSMUSG00000072082 | Ccnf | cyclin F | 1.89 |

| | | | |
|---|---|---|---|
| ENSMUSG00000004044 | Cavin1 | caveolae associated 1 | 1.89 |
| ENSMUSG00000026622 | Nek2 | NIMA (never in mitosis gene a)-related expressed kinase 2 | 1.88 |
| ENSMUSG00000033952 | Aspm | abnormal spindle microtubule assembly | 1.88 |
| ENSMUSG00000023505 | Cdca3 | cell division cycle associated 3 | 1.87 |
| ENSMUSG00000069310 | Hist1h3c | histone cluster 1, H3c | 1.86 |
| ENSMUSG00000090394 | 4930523C07Rik | RIKEN cDNA 4930523C07 gene | 1.86 |
| ENSMUSG00000034311 | Kif4 | kinesin family member 4 | 1.85 |
| ENSMUSG00000035683 | Melk | maternal embryonic leucine zipper kinase | 1.84 |
| ENSMUSG00000041984 | Rptn | repetin | 1.84 |
| ENSMUSG00000114456 | Hist1h2bh | histone cluster 1, H2bh | 1.83 |
| ENSMUSG00000094338 | Hist1h2bl | histone cluster 1, H2bl | 1.83 |
| ENSMUSG00000001435 | Col18a1 | collagen, type XVIII, alpha 1 | 1.82 |
| ENSMUSG00000019832 | Rab32 | RAB32, member RAS oncogene family | 1.82 |
| ENSMUSG00000069267 | Hist1h3b | histone cluster 1, H3b | 1.81 |
| ENSMUSG00000022322 | Shcbp1 | Shc SH2-domain binding protein 1 | 1.81 |
| ENSMUSG00000001403 | Ube2c | ubiquitin-conjugating enzyme E2C | 1.81 |
| ENSMUSG00000032218 | Ccnb2 | cyclin B2 | 1.80 |
| ENSMUSG00000019942 | Cdk1 | cyclin-dependent kinase 1 | 1.80 |
| ENSMUSG00000081740 | Gm14279 | predicted gene 14279 | 1.80 |
| ENSMUSG00000023015 | Racgap1 | Rac GTPase-activating protein 1 | 1.79 |
| ENSMUSG00000099583 | Hist1h3d | histone cluster 1, H3d | 1.79 |
| ENSMUSG00000042306 | S100a14 | S100 calcium binding protein A14 | 1.79 |
| ENSMUSG00000027737 | Slc7a11 | solute carrier family 7 (cationic amino acid transporter, y+ system), member 11 | 1.79 |
| ENSMUSG00000024727 | Trpm6 | transient receptor potential cation channel, subfamily M, member 6 | 1.79 |
| ENSMUSG00000026683 | Nuf2 | NUF2, NDC80 kinetochore complex component | 1.78 |
| ENSMUSG00000022096 | Hr | hairless | 1.77 |
| ENSMUSG00000028873 | Cdca8 | cell division cycle associated 8 | 1.77 |
| ENSMUSG00000021965 | Ska3 | spindle and kinetochore associated complex subunit 3 | 1.77 |
| ENSMUSG00000021493 | Pdlim7 | PDZ and LIM domain 7 | 1.77 |
| ENSMUSG00000020493 | Prr11 | proline rich 11 | 1.77 |
| ENSMUSG00000048376 | F2r | coagulation factor II (thrombin) receptor | 1.77 |
| ENSMUSG00000038943 | Prc1 | protein regulator of cytokinesis 1 | 1.76 |
| ENSMUSG00000029177 | Cenpa | centromere protein A | 1.76 |
| ENSMUSG00000031262 | Cenpi | centromere protein I | 1.76 |
| ENSMUSG00000099517 | Hist1h3g | histone cluster 1, H3g | 1.75 |
| ENSMUSG00000026605 | Cenpf | centromere protein F | 1.74 |
| ENSMUSG00000098985 | Gm27219 | predicted gene 27219 | 1.74 |
| ENSMUSG00000037313 | Tacc3 | transforming, acidic coiled-coil containing protein 3 | 1.73 |
| ENSMUSG00000029910 | Mad2l1 | MAD2 mitotic arrest deficient-like 1 | 1.73 |
| ENSMUSG00000022021 | Diaph3 | diaphanous related formin 3 | 1.73 |
| ENSMUSG00000035783 | Acta2 | actin, alpha 2, smooth muscle, aorta | 1.73 |
| ENSMUSG00000045328 | Cenpe | centromere protein E | 1.72 |
| ENSMUSG00000069910 | Spdl1 | spindle apparatus coiled-coil protein 1 | 1.72 |
| ENSMUSG00000036111 | Lmo1 | LIM domain only 1 | 1.72 |
| ENSMUSG00000048327 | Ckap2l | cytoskeleton associated protein 2-like | 1.71 |
| ENSMUSG00000022034 | Esco2 | establishment of sister chromatid cohesion N-acetyltransferase 2 | 1.71 |
| ENSMUSG00000027115 | Kif18a | kinesin family member 18A | 1.71 |
| ENSMUSG00000041064 | Pif1 | PIF1 5'-to-3' DNA helicase | 1.71 |
| ENSMUSG00000067818 | Myl9 | myosin, light polypeptide 9, regulatory | 1.71 |
| ENSMUSG00000027496 | Aurka | aurora kinase A | 1.70 |
| ENSMUSG00000114279 | Hist1h2bm | histone cluster 1, H2bm | 1.70 |
| ENSMUSG00000114374 | Gm7911 | predicted gene 7911 | 1.70 |
| ENSMUSG00000075031 | Hist1h2bb | histone cluster 1, H2bb | 1.69 |
| ENSMUSG00000031257 | Nox1 | NADPH oxidase 1 | 1.69 |
| ENSMUSG00000074364 | Ehd2 | EH-domain containing 2 | 1.69 |
| ENSMUSG00000027331 | Knstrn | kinetochore-localized astrin/SPAG5 binding | 1.68 |
| ENSMUSG00000023940 | Sgo1 | shugoshin 1 | 1.68 |
| ENSMUSG00000062661 | Ncs1 | neuronal calcium sensor 1 | 1.68 |
| ENSMUSG00000110644 | Gm7390 | predicted gene 7390 | 1.68 |
| ENSMUSG00000017861 | Mybl2 | myeloblastosis oncogene-like 2 | 1.67 |
| ENSMUSG00000024912 | Fosl1 | fos-like antigen 1 | 1.67 |
| ENSMUSG00000004552 | Ctse | cathepsin E | 1.67 |
| ENSMUSG00000097651 | 4930461G14Rik | RIKEN cDNA 4930461G14 gene | 1.67 |
| ENSMUSG00000026435 | Slc45a3 | solute carrier family 45, member 3 | 1.67 |
| ENSMUSG00000028073 | Pear1 | platelet endothelial aggregation receptor 1 | 1.66 |
| ENSMUSG00000033031 | Cip2a | cell proliferation regulating inhibitor of protein phosphatase 2A | 1.65 |
| ENSMUSG00000030677 | Kif22 | kinesin family member 22 | 1.64 |

| | | | |
|---|---|---|---|
| ENSMUSG00000014444 | Piezo1 | piezo-type mechanosensitive ion channel component 1 | 1.64 |
| ENSMUSG00000069184 | Zfp72 | zinc finger protein 72 | 1.64 |
| ENSMUSG00000029377 | Ereg | epiregulin | 1.64 |
| ENSMUSG00000090877 | Hspa1b | heat shock protein 1B | 1.63 |
| ENSMUSG00000027306 | Nusap1 | nucleolar and spindle associated protein 1 | 1.62 |
| ENSMUSG00000037725 | Ckap2 | cytoskeleton associated protein 2 | 1.62 |
| ENSMUSG00000021701 | Plk2 | polo like kinase 2 | 1.62 |
| ENSMUSG00000024795 | Kif20b | kinesin family member 20B | 1.62 |
| ENSMUSG00000021569 | Trip13 | thyroid hormone receptor interactor 13 | 1.62 |
| ENSMUSG00000031075 | Ano1 | anoctamin 1, calcium activated chloride channel | 1.62 |
| ENSMUSG00000103445 | Gm36948 | predicted gene, 36948 | 1.62 |
| ENSMUSG00000026970 | Rbms1 | RNA binding motif, single stranded interacting protein 1 | 1.61 |
| ENSMUSG00000069300 | Hist1h2bj | histone cluster 1, H2bj | 1.61 |
| ENSMUSG00000062727 | Hist1h2bk | histone cluster 1, H2bk | 1.61 |
| ENSMUSG00000051439 | Cd14 | CD14 antigen | 1.61 |
| ENSMUSG00000067006 | Serpinb5 | serine (or cysteine) peptidase inhibitor, clade B, member 5 | 1.61 |
| ENSMUSG00000037716 | Ccdc33 | coiled-coil domain containing 33 | 1.61 |
| ENSMUSG00000032446 | Eomes | eomesodermin | 1.61 |
| ENSMUSG00000020914 | Top2a | topoisomerase (DNA) II alpha | 1.60 |
| ENSMUSG00000015880 | Ncapg | non-SMC condensin I complex, subunit G | 1.60 |
| ENSMUSG00000062937 | Mtap | methylthioadenosine phosphorylase | 1.60 |
| ENSMUSG00000012443 | Kif11 | kinesin family member 11 | 1.60 |
| ENSMUSG00000032717 | Mdfi | MyoD family inhibitor | 1.60 |
| ENSMUSG00000058773 | Hist1h1b | histone cluster 1, H1b | 1.59 |
| ENSMUSG00000052565 | Hist1h1d | histone cluster 1, H1d | 1.59 |
| ENSMUSG00000058290 | Espl1 | extra spindle pole bodies 1, separase | 1.59 |
| ENSMUSG00000020897 | Aurkb | aurora kinase B | 1.59 |
| ENSMUSG00000040084 | Bub1b | BUB1B, mitotic checkpoint serine/threonine kinase | 1.59 |
| ENSMUSG00000038295 | Atg9b | autophagy related 9B | 1.59 |
| ENSMUSG00000040836 | Gpr161 | G protein-coupled receptor 161 | 1.59 |
| ENSMUSG00000049866 | Arl4c | ADP-ribosylation factor-like 4C | 1.58 |
| ENSMUSG00000025372 | Baiap2 | brain-specific angiogenesis inhibitor 1-associated protein 2 | 1.58 |
| ENSMUSG00000022883 | Robo1 | roundabout guidance receptor 1 | 1.58 |
| ENSMUSG00000037071 | Scd1 | stearoyl-Coenzyme A desaturase 1 | 1.58 |
| ENSMUSG00000038252 | Ncapd2 | non-SMC condensin I complex, subunit D2 | 1.57 |
| ENSMUSG00000001517 | Foxm1 | forkhead box M1 | 1.57 |
| ENSMUSG00000059791 | Nrm | nurim (nuclear envelope membrane protein) | 1.57 |
| ENSMUSG00000089762 | Ier5l | immediate early response 5-like | 1.57 |
| ENSMUSG00000073599 | Ecscr | endothelial cell surface expressed chemotaxis and apoptosis regulator | 1.56 |
| ENSMUSG00000078521 | Aunip | aurora kinase A and ninein interacting protein | 1.56 |
| ENSMUSG00000082485 | Gm15693 | predicted gene 15693 | 1.56 |
| ENSMUSG00000021604 | Irx4 | Iroquois homeobox 4 | 1.56 |
| ENSMUSG00000026039 | Sgo2a | shugoshin 2A | 1.55 |
| ENSMUSG00000069265 | Hist1h3a | histone cluster 1, H3a | 1.55 |
| ENSMUSG00000004891 | Nes | nestin | 1.55 |
| ENSMUSG00000023931 | Efhb | EF hand domain family, member B | 1.55 |
| ENSMUSG00000104238 | Gm37587 | predicted gene, 37587 | 1.55 |
| ENSMUSG00000022033 | Pbk | PDZ binding kinase | 1.54 |
| ENSMUSG00000095217 | Hist1h2bn | histone cluster 1, H2bn | 1.54 |
| ENSMUSG00000025921 | Rdh10 | retinol dehydrogenase 10 (all-trans) | 1.54 |
| ENSMUSG00000034773 | BC030867 | cDNA sequence BC030867 | 1.54 |
| ENSMUSG00000032348 | Gsta4 | glutathione S-transferase, alpha 4 | 1.54 |
| ENSMUSG00000043430 | Psapl1 | prosaposin-like 1 | 1.54 |
| ENSMUSG00000027323 | Rad51 | RAD51 recombinase | 1.53 |
| ENSMUSG00000069274 | Hist1h4f | histone cluster 1, H4f | 1.53 |
| ENSMUSG00000032221 | Mns1 | meiosis-specific nuclear structural protein 1 | 1.53 |
| ENSMUSG00000097755 | 2010110K18Rik | RIKEN cDNA 2010110K18 gene | 1.53 |
| ENSMUSG00000025279 | Dnase1l3 | deoxyribonuclease 1-like 3 | 1.53 |
| ENSMUSG00000058385 | Hist1h2bg | histone cluster 1, H2bg | 1.52 |
| ENSMUSG00000075033 | Nxpe3 | neurexophilin and PC-esterase domain family, member 3 | 1.52 |
| ENSMUSG00000022218 | Tgm1 | transglutaminase 1, K polypeptide | 1.52 |
| ENSMUSG00000061991 | Hist1h2af | histone cluster 1, H2af | 1.52 |
| ENSMUSG00000025154 | Arhgap19 | Rho GTPase activating protein 19 | 1.51 |
| ENSMUSG00000028312 | Smc2 | structural maintenance of chromosomes 2 | 1.51 |
| ENSMUSG00000057193 | Slc44a2 | solute carrier family 44, member 2 | 1.51 |
| ENSMUSG00000051220 | Ercc6l | excision repair cross-complementing rodent repair deficiency complementation grc | 1.51 |
| ENSMUSG00000045273 | Cenph | centromere protein H | 1.51 |

| | | | |
|---|---|---|---|
| ENSMUSG00000020492 | Ska2 | spindle and kinetochore associated complex subunit 2 | 1.51 |
| ENSMUSG00000006777 | Krt23 | keratin 23 | 1.50 |
| ENSMUSG00000027326 | Knl1 | kinetochore scaffold 1 | 1.50 |
| ENSMUSG00000068101 | Cenpm | centromere protein M | 1.50 |
| ENSMUSG00000072980 | Oip5 | Opa interacting protein 5 | 1.50 |
| ENSMUSG00000000489 | Pdgfb | platelet derived growth factor, B polypeptide | 1.50 |
| ENSMUSG00000034906 | Ncaph | non-SMC condensin I complex, subunit H | 1.49 |
| ENSMUSG00000082361 | Btc | betacellulin, epidermal growth factor family member | 1.49 |
| ENSMUSG00000022385 | Gtse1 | G two S phase expressed protein 1 | 1.49 |
| ENSMUSG00000026278 | Bok | BCL2-related ovarian killer | 1.49 |
| ENSMUSG00000024590 | Lmnb1 | lamin B1 | 1.48 |
| ENSMUSG00000064288 | Hist1h4k | histone cluster 1, H4k | 1.48 |
| ENSMUSG00000043439 | Epop | elongin BC and polycomb repressive complex 2 associated protein | 1.48 |
| ENSMUSG00000001473 | Tubb6 | tubulin, beta 6 class V | 1.48 |
| ENSMUSG00000030641 | Ddias | DNA damage-induced apoptosis suppressor | 1.48 |
| ENSMUSG00000061615 | Hist1h2ab | histone cluster 1, H2ab | 1.48 |
| ENSMUSG00000040663 | Clcf1 | cardiotrophin-like cytokine factor 1 | 1.48 |
| ENSMUSG00000002297 | Dbf4 | DBF4 zinc finger | 1.47 |
| ENSMUSG00000041859 | Mcm3 | minichromosome maintenance complex component 3 | 1.47 |
| ENSMUSG00000038379 | Ttk | Ttk protein kinase | 1.47 |
| ENSMUSG00000090534 | Gm4675 | predicted gene 4675 | 1.47 |
| ENSMUSG00000025758 | Plk4 | polo like kinase 4 | 1.46 |
| ENSMUSG00000049539 | Hist1h1a | histone cluster 1, H1a | 1.46 |
| ENSMUSG00000016382 | Pls3 | plastin 3 (T-isoform) | 1.46 |
| ENSMUSG00000047534 | Mis18bp1 | MIS18 binding protein 1 | 1.46 |
| ENSMUSG00000055945 | Prr18 | proline rich 18 | 1.46 |
| ENSMUSG00000011267 | Zfp296 | zinc finger protein 296 | 1.46 |
| ENSMUSG00000116919 | AC165151.1 | conserved helix-loop-helix ubiquitous kinase (Chuk) pseudogene | 1.46 |
| ENSMUSG00000005233 | Spc25 | SPC25, NDC80 kinetochore complex component, homolog (S. cerevisiae) | 1.45 |
| ENSMUSG00000074825 | Itpripl1 | inositol 1,4,5-triphosphate receptor interacting protein-like 1 | 1.45 |
| ENSMUSG00000023008 | Fmnl3 | formin-like 3 | 1.45 |
| ENSMUSG00000116102 | Gm20082 | predicted gene, 20082 | 1.45 |
| ENSMUSG00000036768 | Kif15 | kinesin family member 15 | 1.44 |
| ENSMUSG00000023004 | Tuba1b | tubulin, alpha 1B | 1.44 |
| ENSMUSG00000029762 | Akr1b8 | aldo-keto reductase family 1, member B8 | 1.44 |
| ENSMUSG00000042367 | Gjb3 | gap junction protein, beta 3 | 1.44 |
| ENSMUSG00000052852 | Reep1 | receptor accessory protein 1 | 1.44 |
| ENSMUSG00000071516 | Hist1h2ai | histone cluster 1, H2ai | 1.44 |
| ENSMUSG00000062310 | Glrp1 | glutamine repeat protein 1 | 1.44 |
| ENSMUSG00000081943 | Gm9078 | predicted gene 9078 | 1.44 |
| ENSMUSG00000069266 | Hist1h4b | histone cluster 1, H4b | 1.43 |
| ENSMUSG00000034295 | Fhod3 | formin homology 2 domain containing 3 | 1.43 |
| ENSMUSG00000024660 | Incenp | inner centromere protein | 1.42 |
| ENSMUSG00000100210 | Hist1h3f | histone cluster 1, H3f | 1.42 |
| ENSMUSG00000036223 | Ska1 | spindle and kinetochore associated complex subunit 1 | 1.42 |
| ENSMUSG00000019966 | Kitl | kit ligand | 1.41 |
| ENSMUSG00000022881 | Rfc4 | replication factor C (activator 1) 4 | 1.41 |
| ENSMUSG00000048922 | Cdca2 | cell division cycle associated 2 | 1.41 |
| ENSMUSG00000053398 | Phgdh | 3-phosphoglycerate dehydrogenase | 1.41 |
| ENSMUSG00000031756 | Cenpn | centromere protein N | 1.41 |
| ENSMUSG00000056665 | Them6 | thioesterase superfamily member 6 | 1.41 |
| ENSMUSG00000025938 | Slco5a1 | solute carrier organic anion transporter family, member 5A1 | 1.41 |
| ENSMUSG00000001228 | Uhrf1 | ubiquitin-like, containing PHD and RING finger domains, 1 | 1.40 |
| ENSMUSG00000020185 | E2f7 | E2F transcription factor 7 | 1.40 |
| ENSMUSG00000024791 | Cdca5 | cell division cycle associated 5 | 1.40 |
| ENSMUSG00000026955 | Sapcd2 | suppressor APC domain containing 2 | 1.40 |
| ENSMUSG00000101972 | Hist1h3i | histone cluster 1, H3i | 1.40 |
| ENSMUSG00000068855 | Hist2h2ac | histone cluster 2, H2ac | 1.40 |
| ENSMUSG00000004665 | Cnn2 | calponin 2 | 1.40 |
| ENSMUSG00000105987 | AI506816 | expressed sequence AI506816 | 1.39 |
| ENSMUSG00000002055 | Spag5 | sperm associated antigen 5 | 1.39 |
| ENSMUSG00000027803 | Wwtr1 | WW domain containing transcription regulator 1 | 1.39 |
| ENSMUSG00000026193 | Fn1 | fibronectin 1 | 1.39 |
| ENSMUSG00000027811 | 4930579G24Rik | RIKEN cDNA 4930579G24 gene | 1.39 |
| ENSMUSG00000030342 | Cd9 | CD9 antigen | 1.38 |
| ENSMUSG00000026779 | Mastl | microtubule associated serine/threonine kinase-like | 1.38 |
| ENSMUSG00000019773 | Fbxo5 | F-box protein 5 | 1.38 |

| | | | |
|---|---|---|---|
| ENSMUSG00000076437 | Selenoh | selenoprotein H | 1.37 |
| ENSMUSG00000046591 | Ticrr | TOPBP1-interacting checkpoint and replication regulator | 1.37 |
| ENSMUSG00000078952 | Lncenc1 | long non-coding RNA, embryonic stem cells expressed 1 | 1.37 |
| ENSMUSG00000109404 | Gm19963 | predicted gene, 19963 | 1.37 |
| ENSMUSG00000030978 | Rrm1 | ribonucleotide reductase M1 | 1.36 |
| ENSMUSG00000029869 | Ephb6 | Eph receptor B6 | 1.36 |
| ENSMUSG00000022582 | Ly6g | lymphocyte antigen 6 complex, locus G | 1.36 |
| ENSMUSG00000005410 | Mcm5 | minichromosome maintenance complex component 5 | 1.35 |
| ENSMUSG00000053604 | Rpia | ribose 5-phosphate isomerase A | 1.35 |
| ENSMUSG00000042489 | Clspn | claspin | 1.35 |
| ENSMUSG00000029521 | Chek2 | checkpoint kinase 2 | 1.35 |
| ENSMUSG00000030513 | Pcsk6 | proprotein convertase subtilisin/kexin type 6 | 1.35 |
| ENSMUSG00000051343 | Rab11fip5 | RAB11 family interacting protein 5 (class I) | 1.35 |
| ENSMUSG00000063021 | Hist1h2ak | histone cluster 1, H2ak | 1.35 |
| ENSMUSG00000069272 | Hist1h2ae | histone cluster 1, H2ae | 1.35 |
| ENSMUSG00000021765 | Fst | follistatin | 1.35 |
| ENSMUSG00000028718 | Stil | Scl/Tal1 interrupting locus | 1.34 |
| ENSMUSG00000032092 | Mpzl2 | myelin protein zero-like 2 | 1.34 |
| ENSMUSG00000028378 | Ptgr1 | prostaglandin reductase 1 | 1.34 |
| ENSMUSG00000025934 | Gsta3 | glutathione S-transferase, alpha 3 | 1.34 |
| ENSMUSG00000073184 | Gm10479 | predicted gene 10479 | 1.34 |
| ENSMUSG00000001525 | Tubb5 | tubulin, beta 5 class I | 1.33 |
| ENSMUSG00000028614 | Ndc1 | NDC1 transmembrane nucleoporin | 1.33 |
| ENSMUSG00000048779 | P2ry6 | pyrimidinergic receptor P2Y, G-protein coupled, 6 | 1.33 |
| ENSMUSG00000034883 | Lrr1 | leucine rich repeat protein 1 | 1.33 |
| ENSMUSG00000049001 | Ndnf | neuron-derived neurotrophic factor | 1.33 |
| ENSMUSG00000034349 | Smc4 | structural maintenance of chromosomes 4 | 1.32 |
| ENSMUSG00000049932 | H2afx | H2A histone family, member X | 1.32 |
| ENSMUSG00000027381 | Bcl2l11 | BCL2-like 11 (apoptosis facilitator) | 1.32 |
| ENSMUSG00000022325 | Pop1 | processing of precursor 1, ribonuclease P/MRP family, (S. cerevisiae) | 1.32 |
| ENSMUSG00000030254 | Rad18 | RAD18 E3 ubiquitin protein ligase | 1.32 |
| ENSMUSG00000098132 | Rassf10 | Ras association (RalGDS/AF-6) domain family (N-terminal) member 10 | 1.32 |
| ENSMUSG00000091021 | Gm17300 | predicted gene, 17300 | 1.32 |
| ENSMUSG00000037185 | Krt80 | keratin 80 | 1.32 |
| ENSMUSG00000000320 | Alox12 | arachidonate 12-lipoxygenase | 1.32 |
| ENSMUSG00000031004 | Mki67 | antigen identified by monoclonal antibody Ki 67 | 1.31 |
| ENSMUSG00000062248 | Cks2 | CDC28 protein kinase regulatory subunit 2 | 1.31 |
| ENSMUSG00000001056 | Nhp2 | NHP2 ribonucleoprotein | 1.31 |
| ENSMUSG00000033307 | Mif | macrophage migration inhibitory factor (glycosylation-inhibiting factor) | 1.31 |
| ENSMUSG00000035351 | Nup37 | nucleoporin 37 | 1.31 |
| ENSMUSG00000040658 | Dnph1 | 2'-deoxynucleoside 5'-phosphate N-hydrolase 1 | 1.31 |
| ENSMUSG00000031861 | Lpar2 | lysophosphatidic acid receptor 2 | 1.31 |
| ENSMUSG00000098090 | 2700099C18Rik | RIKEN cDNA 2700099C18 gene | 1.31 |
| ENSMUSG00000037148 | Arhgap10 | Rho GTPase activating protein 10 | 1.31 |
| ENSMUSG00000010760 | Phlda2 | pleckstrin homology like domain, family A, member 2 | 1.31 |
| ENSMUSG00000028128 | F3 | coagulation factor III | 1.31 |
| ENSMUSG00000108218 | Olfr1372-ps1 | olfactory receptor 1372, pseudogene 1 | 1.31 |
| ENSMUSG00000022346 | Myc | myelocytomatosis oncogene | 1.30 |
| ENSMUSG00000028599 | Tnfrsf1b | tumor necrosis factor receptor superfamily, member 1b | 1.30 |
| ENSMUSG00000060093 | Hist1h4a | histone cluster 1, H4a | 1.30 |
| ENSMUSG00000062510 | Nsl1 | NSL1, MIS12 kinetochore complex component | 1.30 |
| ENSMUSG00000078773 | Rad54b | RAD54 homolog B (S. cerevisiae) | 1.30 |
| ENSMUSG00000031383 | Dusp9 | dual specificity phosphatase 9 | 1.30 |
| ENSMUSG00000027210 | Meis2 | Meis homeobox 2 | 1.30 |
| ENSMUSG00000083618 | Gm11771 | predicted gene 11771 | 1.30 |
| ENSMUSG00000028780 | Sema3c | sema domain, immunoglobulin domain (Ig), short basic domain, secreted, (semaph | 1.29 |
| ENSMUSG00000005732 | Ranbp1 | RAN binding protein 1 | 1.29 |
| ENSMUSG00000035455 | Fignl1 | fidgetin-like 1 | 1.29 |
| ENSMUSG00000029414 | Kntc1 | kinetochore associated 1 | 1.29 |
| ENSMUSG00000078532 | Nkain1 | Na+/K+ transporting ATPase interacting 1 | 1.29 |
| ENSMUSG00000056904 | Gm5620 | predicted gene 5620 | 1.29 |
| ENSMUSG00000062007 | Hsh2d | hematopoietic SH2 domain containing | 1.29 |
| ENSMUSG00000032108 | D730048I06Rik | RIKEN cDNA D730048I06 gene | 1.29 |
| ENSMUSG00000000028 | Cdc45 | cell division cycle 45 | 1.28 |
| ENSMUSG00000031697 | Orc6 | origin recognition complex, subunit 6 | 1.28 |
| ENSMUSG00000027752 | Exosc8 | exosome component 8 | 1.28 |
| ENSMUSG00000024613 | Tcof1 | treacle ribosome biogenesis factor 1 | 1.28 |

| | | | |
|---|---|---|---|
| ENSMUSG00000051235 | Gen1 | GEN1, Holliday junction 5' flap endonuclease | 1.28 |
| ENSMUSG00000030929 | Eri2 | exoribonuclease 2 | 1.28 |
| ENSMUSG00000032586 | Traip | TRAF-interacting protein | 1.28 |
| ENSMUSG00000019997 | Ctgf | connective tissue growth factor | 1.28 |
| ENSMUSG00000079357 | Gm11100 | predicted gene 11100 | 1.28 |
| ENSMUSG00000022673 | Mcm4 | minichromosome maintenance complex component 4 | 1.27 |
| ENSMUSG00000005470 | Asf1b | anti-silencing function 1B histone chaperone | 1.27 |
| ENSMUSG00000039994 | Timeless | timeless circadian clock 1 | 1.27 |
| ENSMUSG00000028031 | Dkk2 | dickkopf WNT signaling pathway inhibitor 2 | 1.27 |
| ENSMUSG00000035104 | Eva1a | eva-1 homolog A (C. elegans) | 1.27 |
| ENSMUSG00000098113 | Gm2445 | predicted gene 2445 | 1.27 |
| ENSMUSG00000041219 | Arhgap11a | Rho GTPase activating protein 11A | 1.26 |
| ENSMUSG00000028896 | Rcc1 | regulator of chromosome condensation 1 | 1.26 |
| ENSMUSG00000021115 | Vrk1 | vaccinia related kinase 1 | 1.26 |
| ENSMUSG00000040034 | Nup43 | nucleoporin 43 | 1.26 |
| ENSMUSG00000022978 | Mis18a | MIS18 kinetochore protein A | 1.26 |
| ENSMUSG00000027353 | Mcm8 | minichromosome maintenance 8 homologous recombination repair factor | 1.26 |
| ENSMUSG00000027907 | S100a11 | S100 calcium binding protein A11 | 1.26 |
| ENSMUSG00000038508 | Gdf15 | growth differentiation factor 15 | 1.26 |
| ENSMUSG00000040204 | Pclaf | PCNA clamp associated factor | 1.25 |
| ENSMUSG00000028044 | Cks1b | CDC28 protein kinase 1b | 1.25 |
| ENSMUSG00000028587 | Orc1 | origin recognition complex, subunit 1 | 1.25 |
| ENSMUSG00000068270 | Shroom4 | shroom family member 4 | 1.25 |
| ENSMUSG00000013415 | Igf2bp1 | insulin-like growth factor 2 mRNA binding protein 1 | 1.25 |
| ENSMUSG00000032013 | Trim29 | tripartite motif-containing 29 | 1.25 |
| ENSMUSG00000054717 | Hmgb2 | high mobility group box 2 | 1.24 |
| ENSMUSG00000047443 | Erfe | erythroferrone | 1.24 |
| ENSMUSG00000069268 | Hist1h2bf | histone cluster 1, H2bf | 1.24 |
| ENSMUSG00000030352 | Tspan9 | tetraspanin 9 | 1.24 |
| ENSMUSG00000074796 | Slc4a11 | solute carrier family 4, sodium bicarbonate transporter-like, member 11 | 1.24 |
| ENSMUSG00000108528 | Gm8319 | predicted gene 8319 | 1.24 |
| ENSMUSG00000006218 | Fam131c | family with sequence similarity 131, member C | 1.24 |
| ENSMUSG00000085627 | Gm11222 | predicted gene 11222 | 1.24 |
| ENSMUSG00000030162 | Olr1 | oxidized low density lipoprotein (lectin-like) receptor 1 | 1.24 |
| ENSMUSG00000040681 | Hmgn1 | high mobility group nucleosomal binding domain 1 | 1.23 |
| ENSMUSG00000007050 | Lsm2 | LSM2 homolog, U6 small nuclear RNA and mRNA degradation associated | 1.23 |
| ENSMUSG00000032232 | Cgnl1 | cingulin-like 1 | 1.23 |
| ENSMUSG00000054342 | Kcnn4 | potassium intermediate/small conductance calcium-activated channel, subfamily N | 1.23 |
| ENSMUSG00000039509 | Nup133 | nucleoporin 133 | 1.23 |
| ENSMUSG00000020534 | Shmt1 | serine hydroxymethyltransferase 1 (soluble) | 1.23 |
| ENSMUSG00000037206 | Islr | immunoglobulin superfamily containing leucine-rich repeat | 1.23 |
| ENSMUSG00000047832 | Cdca4 | cell division cycle associated 4 | 1.22 |
| ENSMUSG00000025403 | Shmt2 | serine hydroxymethyltransferase 2 (mitochondrial) | 1.22 |
| ENSMUSG00000069273 | Hist1h3e | histone cluster 1, H3e | 1.22 |
| ENSMUSG00000022246 | Rai14 | retinoic acid induced 14 | 1.22 |
| ENSMUSG00000028702 | Rad54l | RAD54 like (S. cerevisiae) | 1.22 |
| ENSMUSG00000037466 | Tedc1 | tubulin epsilon and delta complex 1 | 1.22 |
| ENSMUSG00000032400 | Zwilch | zwilch kinetochore protein | 1.22 |
| ENSMUSG00000026196 | Bard1 | BRCA1 associated RING domain 1 | 1.22 |
| ENSMUSG00000027635 | Dsn1 | DSN1 homolog, MIS12 kinetochore complex component | 1.22 |
| ENSMUSG00000031629 | Cenpu | centromere protein U | 1.22 |
| ENSMUSG00000031351 | Zfp185 | zinc finger protein 185 | 1.22 |
| ENSMUSG00000030935 | Acsm3 | acyl-CoA synthetase medium-chain family member 3 | 1.22 |
| ENSMUSG00000062588 | Gm6104 | predicted gene 6104 | 1.22 |
| ENSMUSG00000021509 | Slc25a48 | solute carrier family 25, member 48 | 1.22 |
| ENSMUSG00000111729 | Gm48038 | predicted gene, 48038 | 1.22 |
| ENSMUSG00000029430 | Ran | RAN, member RAS oncogene family | 1.21 |
| ENSMUSG00000026434 | Nucks1 | nuclear casein kinase and cyclin-dependent kinase substrate 1 | 1.21 |
| ENSMUSG00000005087 | Cd44 | CD44 antigen | 1.21 |
| ENSMUSG00000020592 | Sdc1 | syndecan 1 | 1.21 |
| ENSMUSG00000031146 | Plp2 | proteolipid protein 2 | 1.21 |
| ENSMUSG00000073705 | Cenps | centromere protein S | 1.21 |
| ENSMUSG00000096726 | Gm5558 | predicted gene 5558 | 1.21 |
| ENSMUSG00000024891 | Slc29a2 | solute carrier family 29 (nucleoside transporters), member 2 | 1.21 |
| ENSMUSG00000024401 | Tnf | tumor necrosis factor | 1.21 |
| ENSMUSG00000018566 | Slc2a4 | solute carrier family 2 (facilitated glucose transporter), member 4 | 1.21 |
| ENSMUSG00000037824 | Tspan14 | tetraspanin 14 | 1.20 |

| | | | |
|---|---|---|---|
| ENSMUSG00000030796 | Tead2 | TEA domain family member 2 | 1.20 |
| ENSMUSG00000030528 | Blm | Bloom syndrome, RecQ like helicase | 1.20 |
| ENSMUSG00000037860 | Aim2 | absent in melanoma 2 | 1.20 |
| ENSMUSG00000034205 | Loxl2 | lysyl oxidase-like 2 | 1.20 |
| ENSMUSG00000053297 | AI854703 | expressed sequence AI854703 | 1.20 |
| ENSMUSG00000005481 | Ddx39 | DEAD (Asp-Glu-Ala-Asp) box polypeptide 39 | 1.19 |
| ENSMUSG00000029730 | Mcm7 | minichromosome maintenance complex component 7 | 1.19 |
| ENSMUSG00000043336 | Filip1l | filamin A interacting protein 1-like | 1.19 |
| ENSMUSG00000041506 | Rrp9 | RRP9, small subunit (SSU) processome component, homolog (yeast) | 1.19 |
| ENSMUSG00000025747 | Tyms | thymidylate synthase | 1.19 |
| ENSMUSG00000026999 | Nup35 | nucleoporin 35 | 1.19 |
| ENSMUSG00000016526 | Dyrk3 | dual-specificity tyrosine-(Y)-phosphorylation regulated kinase 3 | 1.19 |
| ENSMUSG00000028693 | Nasp | nuclear autoantigenic sperm protein (histone-binding) | 1.18 |
| ENSMUSG00000032300 | 1700017B05Rik | RIKEN cDNA 1700017B05 gene | 1.18 |
| ENSMUSG00000063018 | 2010204K13Rik | RIKEN cDNA 2010204K13 gene | 1.18 |
| ENSMUSG00000028199 | Cryz | crystallin, zeta | 1.18 |
| ENSMUSG00000069307 | Hist1h2bq | histone cluster 1, H2bq | 1.18 |
| ENSMUSG00000084353 | Gm15452 | predicted gene 15452 | 1.18 |
| ENSMUSG00000032940 | Rbm11 | RNA binding motif protein 11 | 1.18 |
| ENSMUSG00000006585 | Cdt1 | chromatin licensing and DNA replication factor 1 | 1.17 |
| ENSMUSG00000002870 | Mcm2 | minichromosome maintenance complex component 2 | 1.17 |
| ENSMUSG00000091625 | Lsm5 | LSM5 homolog, U6 small nuclear RNA and mRNA degradation associated | 1.17 |
| ENSMUSG00000022945 | Chaf1b | chromatin assembly factor 1, subunit B (p60) | 1.17 |
| ENSMUSG00000026646 | Suv39h2 | suppressor of variegation 3-9 2 | 1.17 |
| ENSMUSG00000022422 | Dscc1 | DNA replication and sister chromatid cohesion 1 | 1.17 |
| ENSMUSG00000080186 | Gm14448 | predicted gene 14448 | 1.17 |
| ENSMUSG00000046589 | Lrrc8e | leucine rich repeat containing 8 family, member E | 1.17 |
| ENSMUSG00000082163 | Gm14276 | predicted gene 14276 | 1.17 |
| ENSMUSG00000087201 | Gm15261 | predicted gene 15261 | 1.17 |
| ENSMUSG00000019961 | Tmpo | thymopoietin | 1.16 |
| ENSMUSG00000027405 | Nop56 | NOP56 ribonucleoprotein | 1.16 |
| ENSMUSG00000056394 | Lig1 | ligase I, DNA, ATP-dependent | 1.16 |
| ENSMUSG00000024732 | Ccdc86 | coiled-coil domain containing 86 | 1.16 |
| ENSMUSG00000026669 | Mcm10 | minichromosome maintenance 10 replication initiation factor | 1.16 |
| ENSMUSG00000024844 | Banf1 | barrier to autointegration factor 1 | 1.16 |
| ENSMUSG00000053801 | Grwd1 | glutamate-rich WD repeat containing 1 | 1.16 |
| ENSMUSG00000055760 | Gemin6 | gem nuclear organelle associated protein 6 | 1.16 |
| ENSMUSG00000020937 | Plcd3 | phospholipase C, delta 3 | 1.16 |
| ENSMUSG00000041801 | Phlda3 | pleckstrin homology like domain, family A, member 3 | 1.16 |
| ENSMUSG00000063605 | Ccdc102a | coiled-coil domain containing 102A | 1.16 |
| ENSMUSG00000102840 | Gm38037 | predicted gene, 38037 | 1.16 |
| ENSMUSG00000047246 | Hist1h2be | histone cluster 1, H2be | 1.15 |
| ENSMUSG00000032640 | Chsy1 | chondroitin sulfate synthase 1 | 1.15 |
| ENSMUSG00000027203 | Dut | deoxyuridine triphosphatase | 1.15 |
| ENSMUSG00000074476 | Spc24 | SPC24, NDC80 kinetochore complex component, homolog (S. cerevisiae) | 1.15 |
| ENSMUSG00000023919 | Cenpq | centromere protein Q | 1.15 |
| ENSMUSG00000027257 | Pacsin3 | protein kinase C and casein kinase substrate in neurons 3 | 1.15 |
| ENSMUSG00000032215 | Rsl24d1 | ribosomal L24 domain containing 1 | 1.15 |
| ENSMUSG00000017491 | Rarb | retinoic acid receptor, beta | 1.15 |
| ENSMUSG00000001025 | S100a6 | S100 calcium binding protein A6 (calcyclin) | 1.15 |
| ENSMUSG00000106358 | Gm7047 | predicted gene 7047 | 1.15 |
| ENSMUSG00000082878 | Gm12583 | predicted gene 12583 | 1.15 |
| ENSMUSG00000017499 | Cdc6 | cell division cycle 6 | 1.14 |
| ENSMUSG00000091405 | Hist2h4 | histone cluster 2, H4 | 1.14 |
| ENSMUSG00000029516 | Cit | citron | 1.14 |
| ENSMUSG00000032815 | Fanca | Fanconi anemia, complementation group A | 1.14 |
| ENSMUSG00000044037 | Als2cl | ALS2 C-terminal like | 1.14 |
| ENSMUSG00000040187 | Arntl2 | aryl hydrocarbon receptor nuclear translocator-like 2 | 1.14 |
| ENSMUSG00000063628 | Gm7665 | predicted pseudogene 7665 | 1.14 |
| ENSMUSG00000042272 | Sestd1 | SEC14 and spectrin domains 1 | 1.13 |
| ENSMUSG00000044254 | Pcsk9 | proprotein convertase subtilisin/kexin type 9 | 1.13 |
| ENSMUSG00000027878 | Notch2 | notch 2 | 1.13 |
| ENSMUSG00000021958 | Pinx1 | PIN2/TERF1 interacting, telomerase inhibitor 1 | 1.13 |
| ENSMUSG00000037419 | Endod1 | endonuclease domain containing 1 | 1.13 |
| ENSMUSG00000069793 | Slfn9 | schlafen 9 | 1.13 |
| ENSMUSG00000039748 | Exo1 | exonuclease 1 | 1.13 |
| ENSMUSG00000000555 | Itga5 | integrin alpha 5 (fibronectin receptor alpha) | 1.13 |

| | | | |
|---|---|---|---|
| ENSMUSG00000030346 | Rad51ap1 | RAD51 associated protein 1 | 1.13 |
| ENSMUSG00000051517 | Arhgef39 | Rho guanine nucleotide exchange factor (GEF) 39 | 1.13 |
| ENSMUSG00000031379 | Pir | pirin | 1.13 |
| ENSMUSG00000075266 | Cenpw | centromere protein W | 1.13 |
| ENSMUSG00000049916 | 2610318N02Rik | RIKEN cDNA 2610318N02 gene | 1.13 |
| ENSMUSG00000006403 | Adamts4 | a disintegrin-like and metallopeptidase (reprolysin type) with thrombospondin type | 1.13 |
| ENSMUSG00000099902 | Gm12115 | predicted gene 12115 | 1.13 |
| ENSMUSG00000109186 | Gm34821 | predicted gene, 34821 | 1.13 |
| ENSMUSG00000034317 | Trim59 | tripartite motif-containing 59 | 1.12 |
| ENSMUSG00000032184 | Lysmd2 | LysM, putative peptidoglycan-binding, domain containing 2 | 1.12 |
| ENSMUSG00000047044 | D030056L22Rik | RIKEN cDNA D030056L22 gene | 1.12 |
| ENSMUSG00000034206 | Polq | polymerase (DNA directed), theta | 1.12 |
| ENSMUSG00000043924 | Ncmap | noncompact myelin associated protein | 1.12 |
| ENSMUSG00000042215 | Bag2 | BCL2-associated athanogene 2 | 1.12 |
| ENSMUSG00000110218 | Gm20219 | predicted gene, 20219 | 1.12 |
| ENSMUSG00000039187 | Fanci | Fanconi anemia, complementation group I | 1.12 |
| ENSMUSG00000040990 | Sh3kbp1 | SH3-domain kinase binding protein 1 | 1.12 |
| ENSMUSG00000033508 | Asprv1 | aspartic peptidase, retroviral-like 1 | 1.12 |
| ENSMUSG00000052833 | Sae1 | SUMO1 activating enzyme subunit 1 | 1.11 |
| ENSMUSG00000022678 | Nde1 | nudE neurodevelopment protein 1 | 1.11 |
| ENSMUSG00000057113 | Npm1 | nucleophosmin 1 | 1.11 |
| ENSMUSG00000015749 | Anp32e | acidic (leucine-rich) nuclear phosphoprotein 32 family, member E | 1.11 |
| ENSMUSG00000002718 | Cse1l | chromosome segregation 1-like (S. cerevisiae) | 1.11 |
| ENSMUSG00000026234 | Ncl | nucleolin | 1.11 |
| ENSMUSG00000026355 | Mcm6 | minichromosome maintenance complex component 6 | 1.11 |
| ENSMUSG00000061458 | Nol10 | nucleolar protein 10 | 1.11 |
| ENSMUSG00000022636 | Alcam | activated leukocyte cell adhesion molecule | 1.11 |
| ENSMUSG00000038047 | Haus6 | HAUS augmin-like complex, subunit 6 | 1.11 |
| ENSMUSG00000064326 | Siva1 | SIVA1, apoptosis-inducing factor | 1.11 |
| ENSMUSG00000027478 | Dnmt3b | DNA methyltransferase 3B | 1.11 |
| ENSMUSG00000036992 | Nxt1 | NTF2-related export protein 1 | 1.11 |
| ENSMUSG00000044948 | Cfap43 | cilia and flagella associated protein 43 | 1.11 |
| ENSMUSG00000060639 | Hist1h4i | histone cluster 1, H4i | 1.11 |
| ENSMUSG00000006356 | Crip2 | cysteine rich protein 2 | 1.11 |
| ENSMUSG00000055653 | Gpc3 | glypican 3 | 1.11 |
| ENSMUSG00000114515 | Aldoa | aldolase A, fructose-bisphosphate | 1.11 |
| ENSMUSG00000066551 | Hmgb1 | high mobility group box 1 | 1.10 |
| ENSMUSG00000042029 | Ncapg2 | non-SMC condensin II complex, subunit G2 | 1.10 |
| ENSMUSG00000021193 | Pitrm1 | pitrilysin metallepetidase 1 | 1.10 |
| ENSMUSG00000109511 | Nup62 | nucleoporin 62 | 1.10 |
| ENSMUSG00000032939 | Nup93 | nucleoporin 93 | 1.10 |
| ENSMUSG00000070348 | Ccnd1 | cyclin D1 | 1.10 |
| ENSMUSG00000026020 | Nop58 | NOP58 ribonucleoprotein | 1.10 |
| ENSMUSG00000024925 | Rnaseh2c | ribonuclease H2, subunit C | 1.10 |
| ENSMUSG00000021131 | Erh | ERH mRNA splicing and mitosis factor | 1.10 |
| ENSMUSG00000028010 | Gar1 | GAR1 ribonucleoprotein | 1.10 |
| ENSMUSG00000032344 | Mb21d1 | Mab-21 domain containing 1 | 1.10 |
| ENSMUSG00000022464 | Slc38a4 | solute carrier family 38, member 4 | 1.10 |
| ENSMUSG00000024479 | Mal2 | mal, T cell differentiation protein 2 | 1.10 |
| ENSMUSG00000014075 | Tctex1d2 | Tctex1 domain containing 2 | 1.10 |
| ENSMUSG00000078249 | Hmga1b | high mobility group AT-hook 1B | 1.10 |
| ENSMUSG00000115009 | G930009F23Rik | RIKEN cDNA G930009F23 gene | 1.10 |
| ENSMUSG00000025134 | Alyref | Aly/REF export factor | 1.09 |
| ENSMUSG00000058799 | Nap1l1 | nucleosome assembly protein 1-like 1 | 1.09 |
| ENSMUSG00000034192 | Lsm3 | LSM3 homolog, U6 small nuclear RNA and mRNA degradation associated | 1.09 |
| ENSMUSG00000096010 | Hist4h4 | histone cluster 4, H4 | 1.09 |
| ENSMUSG00000001707 | Eef1e1 | eukaryotic translation elongation factor 1 epsilon 1 | 1.09 |
| ENSMUSG00000024800 | Rpp30 | ribonuclease P/MRP 30 subunit | 1.09 |
| ENSMUSG00000032643 | Fhl3 | four and a half LIM domains 3 | 1.09 |
| ENSMUSG00000024301 | Kifc5b | kinesin family member C5B | 1.09 |
| ENSMUSG00000021391 | Cenpp | centromere protein P | 1.09 |
| ENSMUSG00000021418 | Rpp40 | ribonuclease P 40 subunit | 1.09 |
| ENSMUSG00000027454 | Gins1 | GINS complex subunit 1 (Psf1 homolog) | 1.09 |
| ENSMUSG00000031112 | Stk26 | serine/threonine kinase 26 | 1.09 |
| ENSMUSG00000022177 | Haus4 | HAUS augmin-like complex, subunit 4 | 1.08 |
| ENSMUSG00000027018 | Hat1 | histone aminotransferase 1 | 1.08 |
| ENSMUSG00000030079 | Ruvbl1 | RuvB-like protein 1 | 1.08 |

| | | | |
|---|---|---|---|
| ENSMUSG00000031403 | Dkc1 | dyskeratosis congenita 1, dyskerin | 1.08 |
| ENSMUSG00000037020 | Wdr62 | WD repeat domain 62 | 1.08 |
| ENSMUSG00000012483 | Rpa3 | replication protein A3 | 1.08 |
| ENSMUSG00000067367 | Lyar | Ly1 antibody reactive clone | 1.08 |
| ENSMUSG00000039231 | Suv39h1 | suppressor of variegation 3-9 1 | 1.08 |
| ENSMUSG00000039457 | Ppl | periplakin | 1.08 |
| ENSMUSG00000110631 | Gm42047 | predicted gene, 42047 | 1.08 |
| ENSMUSG00000048612 | Myof | myoferlin | 1.08 |
| ENSMUSG00000050520 | Cldn8 | claudin 8 | 1.08 |
| ENSMUSG00000031400 | G6pdx | glucose-6-phosphate dehydrogenase X-linked | 1.07 |
| ENSMUSG00000033032 | Afap1l1 | actin filament associated protein 1-like 1 | 1.07 |
| ENSMUSG00000002835 | Chaf1a | chromatin assembly factor 1, subunit A (p150) | 1.07 |
| ENSMUSG00000049521 | Cdc42ep1 | CDC42 effector protein (Rho GTPase binding) 1 | 1.07 |
| ENSMUSG00000028633 | Ctps | cytidine 5'-triphosphate synthase | 1.07 |
| ENSMUSG00000037275 | Gemin5 | gem nuclear organelle associated protein 5 | 1.07 |
| ENSMUSG00000007080 | Pole | polymerase (DNA directed), epsilon | 1.07 |
| ENSMUSG00000025289 | Prdx4 | peroxiredoxin 4 | 1.07 |
| ENSMUSG00000056305 | Usp39 | ubiquitin specific peptidase 39 | 1.07 |
| ENSMUSG00000024691 | Fam111a | family with sequence similarity 111, member A | 1.07 |
| ENSMUSG00000039055 | Eme1 | essential meiotic structure-specific endonuclease 1 | 1.07 |
| ENSMUSG00000039200 | Atf7ip2 | activating transcription factor 7 interacting protein 2 | 1.07 |
| ENSMUSG00000023885 | Thbs2 | thrombospondin 2 | 1.07 |
| ENSMUSG00000027342 | Pcna | proliferating cell nuclear antigen | 1.06 |
| ENSMUSG00000026547 | Tagln2 | transgelin 2 | 1.06 |
| ENSMUSG00000030189 | Ybx3 | Y box protein 3 | 1.06 |
| ENSMUSG00000109324 | Prmt1 | protein arginine N-methyltransferase 1 | 1.06 |
| ENSMUSG00000032834 | Pwp2 | PWP2 periodic tryptophan protein homolog (yeast) | 1.06 |
| ENSMUSG00000095567 | Noc2l | NOC2 like nucleolar associated transcriptional repressor | 1.06 |
| ENSMUSG00000058392 | Rrp1b | ribosomal RNA processing 1 homolog B (S. cerevisiae) | 1.06 |
| ENSMUSG00000046179 | E2f8 | E2F transcription factor 8 | 1.06 |
| ENSMUSG00000028741 | Mrto4 | mRNA turnover 4, ribosome maturation factor | 1.06 |
| ENSMUSG00000025001 | Hells | helicase, lymphoid specific | 1.06 |
| ENSMUSG00000020974 | Pole2 | polymerase (DNA directed), epsilon 2 (p59 subunit) | 1.06 |
| ENSMUSG00000031563 | Wwc2 | WW, C2 and coiled-coil domain containing 2 | 1.06 |
| ENSMUSG00000027624 | Epb41l1 | erythrocyte membrane protein band 4.1 like 1 | 1.06 |
| ENSMUSG00000021697 | Depdc1b | DEP domain containing 1B | 1.06 |
| ENSMUSG00000117256 | AC154779.4 | peptidylprolyl isomerase A (Ppia) pseudogene | 1.06 |
| ENSMUSG00000097797 | Gm26901 | predicted gene, 26901 | 1.06 |
| ENSMUSG00000041390 | Mdfic | MyoD family inhibitor domain containing | 1.06 |
| ENSMUSG00000041688 | Amot | angiomotin | 1.05 |
| ENSMUSG00000026238 | Ptma | prothymosin alpha | 1.05 |
| ENSMUSG00000116564 | Riok2 | RIO kinase 2 | 1.05 |
| ENSMUSG00000026192 | Atic | 5-aminoimidazole-4-carboxamide ribonucleotide formyltransferase/IMP cyclohydrc | 1.05 |
| ENSMUSG00000002477 | Snrpd1 | small nuclear ribonucleoprotein D1 | 1.05 |
| ENSMUSG00000028884 | Rpa2 | replication protein A2 | 1.05 |
| ENSMUSG00000024640 | Psat1 | phosphoserine aminotransferase 1 | 1.05 |
| ENSMUSG00000026796 | Fam129b | family with sequence similarity 129, member B | 1.05 |
| ENSMUSG00000030512 | Snrpa1 | small nuclear ribonucleoprotein polypeptide A' | 1.05 |
| ENSMUSG00000021645 | Smn1 | survival motor neuron 1 | 1.05 |
| ENSMUSG00000040463 | Mybbp1a | MYB binding protein (P160) 1a | 1.05 |
| ENSMUSG00000026235 | Epha4 | Eph receptor A4 | 1.05 |
| ENSMUSG00000034595 | Ppp1r18 | protein phosphatase 1, regulatory subunit 18 | 1.05 |
| ENSMUSG00000010751 | Tnfrsf22 | tumor necrosis factor receptor superfamily, member 22 | 1.05 |
| ENSMUSG00000096403 | Gm9825 | predicted gene 9825 | 1.05 |
| ENSMUSG00000007033 | Hspa1l | heat shock protein 1-like | 1.05 |
| ENSMUSG00000003868 | Ruvbl2 | RuvB-like protein 2 | 1.04 |
| ENSMUSG00000001674 | Ddx18 | DEAD (Asp-Glu-Ala-Asp) box polypeptide 18 | 1.04 |
| ENSMUSG00000060860 | Ube2s | ubiquitin-conjugating enzyme E2S | 1.04 |
| ENSMUSG00000022391 | Rangap1 | RAN GTPase activating protein 1 | 1.04 |
| ENSMUSG00000020929 | Eftud2 | elongation factor Tu GTP binding domain containing 2 | 1.04 |
| ENSMUSG00000015176 | Nolc1 | nucleolar and coiled-body phosphoprotein 1 | 1.04 |
| ENSMUSG00000022360 | Atad2 | ATPase family, AAA domain containing 2 | 1.04 |
| ENSMUSG00000060981 | Hist1h4h | histone cluster 1, H4h | 1.04 |
| ENSMUSG00000014907 | Naf1 | nuclear assembly factor 1 ribonucleoprotein | 1.04 |
| ENSMUSG00000035960 | Apex1 | apurinic/apyrimidinic endonuclease 1 | 1.04 |
| ENSMUSG00000021714 | Cenpk | centromere protein K | 1.04 |
| ENSMUSG00000067455 | Hist1h4j | histone cluster 1, H4j | 1.04 |

| | | | |
|---|---|---|---|
| ENSMUSG00000095677 | Dynlt1f | dynein light chain Tctex-type 1F | 1.04 |
| ENSMUSG00000039232 | Stx11 | syntaxin 11 | 1.04 |
| ENSMUSG00000019992 | Mtfr2 | mitochondrial fission regulator 2 | 1.04 |
| ENSMUSG00000032724 | Abtb2 | ankyrin repeat and BTB (POZ) domain containing 2 | 1.04 |
| ENSMUSG00000001131 | Timp1 | tissue inhibitor of metalloproteinase 1 | 1.04 |
| ENSMUSG00000116640 | AC166114.1 | twist basic helix-loop-helix transcription factor 1 neighbor (Twistnb) pseudogene | 1.04 |
| ENSMUSG00000025364 | Pa2g4 | proliferation-associated 2G4 | 1.03 |
| ENSMUSG00000029014 | Dnajc2 | DnaJ heat shock protein family (Hsp40) member C2 | 1.03 |
| ENSMUSG00000033166 | Dis3 | DIS3 homolog, exosome endoribonuclease and 3'-5' exoribonuclease | 1.03 |
| ENSMUSG00000025980 | Hspd1 | heat shock protein 1 (chaperonin) | 1.03 |
| ENSMUSG00000029247 | Paics | phosphoribosylaminoimidazole carboxylase, phosphoribosylaminoribosylaminoimic | 1.03 |
| ENSMUSG00000020706 | Ftsj3 | FtsJ RNA methyltransferase homolog 3 (E. coli) | 1.03 |
| ENSMUSG00000027330 | Cdc25b | cell division cycle 25B | 1.03 |
| ENSMUSG00000031434 | Morc4 | microrchidia 4 | 1.03 |
| ENSMUSG00000023988 | Bysl | bystin-like | 1.03 |
| ENSMUSG00000035165 | Kcne3 | potassium voltage-gated channel, Isk-related subfamily, gene 3 | 1.03 |
| ENSMUSG00000015217 | Hmgb3 | high mobility group box 3 | 1.03 |
| ENSMUSG00000033294 | Noc4l | NOC4 like | 1.03 |
| ENSMUSG00000025395 | Prim1 | DNA primase, p49 subunit | 1.03 |
| ENSMUSG00000079478 | Sssca1 | Sjogren syndrome/scleroderma autoantigen 1 | 1.03 |
| ENSMUSG00000038323 | 1700066M21Rik | RIKEN cDNA 1700066M21 gene | 1.03 |
| ENSMUSG00000028933 | Xrcc2 | X-ray repair complementing defective repair in Chinese hamster cells 2 | 1.03 |
| ENSMUSG00000033752 | Mnd1 | meiotic nuclear divisions 1 | 1.03 |
| ENSMUSG00000020656 | Grhl1 | grainyhead like transcription factor 1 | 1.03 |
| ENSMUSG00000046295 | Ankle1 | ankyrin repeat and LEM domain containing 1 | 1.03 |
| ENSMUSG00000079685 | Ulbp1 | UL16 binding protein 1 | 1.03 |
| ENSMUSG00000035678 | Tnfsf9 | tumor necrosis factor (ligand) superfamily, member 9 | 1.03 |
| ENSMUSG00000058656 | Samd12 | sterile alpha motif domain containing 12 | 1.03 |
| ENSMUSG00000080928 | Hmgb1-ps6 | high mobility group box 1, pseudogene 6 | 1.03 |
| ENSMUSG00000030098 | Grip2 | glutamate receptor interacting protein 2 | 1.03 |
| ENSMUSG00000071724 | Smpd5 | sphingomyelin phosphodiesterase 5 | 1.03 |
| ENSMUSG00000054766 | Set | SET nuclear oncogene | 1.02 |
| ENSMUSG00000061482 | Hist1h4d | histone cluster 1, H4d | 1.02 |
| ENSMUSG00000054321 | Taf4b | TATA-box binding protein associated factor 4b | 1.02 |
| ENSMUSG00000028333 | Anp32b | acidic (leucine-rich) nuclear phosphoprotein 32 family, member B | 1.02 |
| ENSMUSG00000039298 | Cdk5rap2 | CDK5 regulatory subunit associated protein 2 | 1.02 |
| ENSMUSG00000068856 | Sf3b4 | splicing factor 3b, subunit 4 | 1.02 |
| ENSMUSG00000060288 | Ppih | peptidyl prolyl isomerase H | 1.02 |
| ENSMUSG00000034612 | Chst11 | carbohydrate sulfotransferase 11 | 1.02 |
| ENSMUSG00000050244 | Heatr1 | HEAT repeat containing 1 | 1.02 |
| ENSMUSG00000001761 | Smo | smoothened, frizzled class receptor | 1.02 |
| ENSMUSG00000022792 | Yars2 | tyrosyl-tRNA synthetase 2 (mitochondrial) | 1.02 |
| ENSMUSG00000019853 | Hebp2 | heme binding protein 2 | 1.02 |
| ENSMUSG00000050107 | Haspin | histone H3 associated protein kinase | 1.02 |
| ENSMUSG00000034023 | Fancd2 | Fanconi anemia, complementation group D2 | 1.02 |
| ENSMUSG00000097415 | AU020206 | expressed sequence AU020206 | 1.02 |
| ENSMUSG00000026404 | Ddx59 | DEAD (Asp-Glu-Ala-Asp) box polypeptide 59 | 1.02 |
| ENSMUSG00000039103 | Nexn | nexilin | 1.02 |
| ENSMUSG00000017765 | Slc12a4 | solute carrier family 12, member 4 | 1.02 |
| ENSMUSG00000020834 | Dhrs13 | dehydrogenase/reductase (SDR family) member 13 | 1.02 |
| ENSMUSG00000020358 | Hnrnpab | heterogeneous nuclear ribonucleoprotein A/B | 1.01 |
| ENSMUSG00000022962 | Gart | phosphoribosylglycinamide formyltransferase | 1.01 |
| ENSMUSG00000028184 | Adgrl2 | adhesion G protein-coupled receptor L2 | 1.01 |
| ENSMUSG00000017607 | Tns4 | tensin 4 | 1.01 |
| ENSMUSG00000046711 | Hmga1 | high mobility group AT-hook 1 | 1.01 |
| ENSMUSG00000001053 | N4bp3 | NEDD4 binding protein 3 | 1.01 |
| ENSMUSG00000019214 | Chtf18 | CTF18, chromosome transmission fidelity factor 18 | 1.01 |
| ENSMUSG00000027978 | Prss12 | protease, serine 12 neurotrypsin (motopsin) | 1.01 |
| ENSMUSG00000029761 | Cald1 | caldesmon 1 | 1.01 |
| ENSMUSG00000044966 | Fbxo48 | F-box protein 48 | 1.01 |
| ENSMUSG00000032473 | Cldn18 | claudin 18 | 1.01 |
| ENSMUSG00000078157 | 4931440F15Rik | RIKEN cDNA 4931440F15 gene | 1.01 |
| ENSMUSG00000044912 | Syt16 | synaptotagmin XVI | 1.01 |
| ENSMUSG00000025613 | Cct8 | chaperonin containing Tcp1, subunit 8 (theta) | 1.00 |
| ENSMUSG00000014226 | Cacybp | calcyclin binding protein | 1.00 |
| ENSMUSG00000025534 | Gusb | glucuronidase, beta | 1.00 |
| ENSMUSG00000022247 | Brix1 | BRX1, biogenesis of ribosomes | 1.00 |

| | | | |
|---|---|---|---|
| ENSMUSG00000005846 | Rsl1d1 | ribosomal L1 domain containing 1 | 1.00 |
| ENSMUSG00000006715 | Gmnn | geminin | 1.00 |
| ENSMUSG00000023110 | Prmt5 | protein arginine N-methyltransferase 5 | 1.00 |
| ENSMUSG00000031790 | Mmp15 | matrix metallopeptidase 15 | 1.00 |
| ENSMUSG00000062127 | Cttnbp2nl | CTTNBP2 N-terminal like | 1.00 |
| ENSMUSG00000105140 | Gm43127 | predicted gene 43127 | 1.00 |
| ENSMUSG00000103576 | Gm38188 | predicted gene, 38188 | 1.00 |
| ENSMUSG00000042251 | Pm20d1 | peptidase M20 domain containing 1 | -1.00 |
| ENSMUSG00000073402 | Gm8909 | predicted gene 8909 | -1.00 |
| ENSMUSG00000110773 | AC061963.1 | novel transcript, antisense to Kmt2a | -1.00 |
| ENSMUSG00000104852 | Gm43201 | predicted gene 43201 | -1.00 |
| ENSMUSG00000032417 | Rwdd2a | RWD domain containing 2A | -1.00 |
| ENSMUSG00000004902 | Slc25a18 | solute carrier family 25 (mitochondrial carrier), member 18 | -1.00 |
| ENSMUSG00000107428 | Gm44154 | predicted gene, 44154 | -1.00 |
| ENSMUSG00000032527 | Pccb | propionyl Coenzyme A carboxylase, beta polypeptide | -1.01 |
| ENSMUSG00000068587 | Mgam | maltase-glucoamylase | -1.01 |
| ENSMUSG00000005686 | Ampd3 | adenosine monophosphate deaminase 3 | -1.01 |
| ENSMUSG00000025467 | Prap1 | proline-rich acidic protein 1 | -1.01 |
| ENSMUSG00000024055 | Cyp4f13 | cytochrome P450, family 4, subfamily f, polypeptide 13 | -1.01 |
| ENSMUSG00000019278 | Dpep1 | dipeptidase 1 (renal) | -1.01 |
| ENSMUSG00000023959 | Clic5 | chloride intracellular channel 5 | -1.01 |
| ENSMUSG00000039199 | Zdhhc1 | zinc finger, DHHC domain containing 1 | -1.01 |
| ENSMUSG00000054999 | Naaladl1 | N-acetylated alpha-linked acidic dipeptidase-like 1 | -1.01 |
| ENSMUSG00000021684 | Pde8b | phosphodiesterase 8B | -1.01 |
| ENSMUSG00000002944 | Cd36 | CD36 molecule | -1.01 |
| ENSMUSG00000112481 | Gm29684 | predicted gene, 29684 | -1.01 |
| ENSMUSG00000082148 | Gm12266 | predicted gene 12266 | -1.01 |
| ENSMUSG00000026395 | Ptprc | protein tyrosine phosphatase, receptor type, C | -1.01 |
| ENSMUSG00000095440 | Fignl2 | fidgetin-like 2 | -1.01 |
| ENSMUSG00000060402 | Chst8 | carbohydrate (N-acetylgalactosamine 4-0) sulfotransferase 8 | -1.01 |
| ENSMUSG00000045689 | Pcdhb4 | protocadherin beta 4 | -1.01 |
| ENSMUSG00000114401 | Gm38604 | predicted gene, 38604 | -1.01 |
| ENSMUSG00000040712 | Camta2 | calmodulin binding transcription activator 2 | -1.02 |
| ENSMUSG00000030102 | Itpr1 | inositol 1,4,5-trisphosphate receptor 1 | -1.02 |
| ENSMUSG00000068742 | Cry2 | cryptochrome 2 (photolyase-like) | -1.02 |
| ENSMUSG00000066800 | Rnasel | ribonuclease L (2', 5'-oligoisoadenylate synthetase-dependent) | -1.02 |
| ENSMUSG00000036833 | Pnpla7 | patatin-like phospholipase domain containing 7 | -1.02 |
| ENSMUSG00000015202 | Cnksr3 | Cnksr family member 3 | -1.02 |
| ENSMUSG00000028211 | Trp53inp1 | transformation related protein 53 inducible nuclear protein 1 | -1.02 |
| ENSMUSG00000000823 | Zfp512b | zinc finger protein 512B | -1.02 |
| ENSMUSG00000019989 | Enpp3 | ectonucleotide pyrophosphatase/phosphodiesterase 3 | -1.02 |
| ENSMUSG00000017390 | Aldoc | aldolase C, fructose-bisphosphate | -1.02 |
| ENSMUSG00000017204 | Gsdma | gasdermin A | -1.02 |
| ENSMUSG00000029188 | Slc34a2 | solute carrier family 34 (sodium phosphate), member 2 | -1.02 |
| ENSMUSG00000022408 | Fam83f | family with sequence similarity 83, member F | -1.02 |
| ENSMUSG00000027296 | Itpka | inositol 1,4,5-trisphosphate 3-kinase A | -1.02 |
| ENSMUSG00000025396 | Hsd17b6 | hydroxysteroid (17-beta) dehydrogenase 6 | -1.02 |
| ENSMUSG00000026692 | Fmo4 | flavin containing monooxygenase 4 | -1.02 |
| ENSMUSG00000086946 | Gm15527 | predicted gene 15527 | -1.02 |
| ENSMUSG00000037003 | Tns2 | tensin 2 | -1.02 |
| ENSMUSG00000046794 | Ppp1r3b | protein phosphatase 1, regulatory subunit 3B | -1.02 |
| ENSMUSG00000064455 | n-R5s56 | nuclear encoded rRNA 5S 56 | -1.02 |
| ENSMUSG00000097258 | Gm26767 | predicted gene, 26767 | -1.02 |
| ENSMUSG00000097006 | 9530082P21Rik | RIKEN cDNA 9530082P21 gene | -1.02 |
| ENSMUSG00000115918 | A930001M01Rik | RIKEN cDNA A930001M01 gene | -1.02 |
| ENSMUSG00000015966 | Il17rb | interleukin 17 receptor B | -1.02 |
| ENSMUSG00000086766 | Gm15969 | predicted gene 15969 | -1.02 |
| ENSMUSG00000021566 | Slc6a19os | solute carrier family 6 (neurotransmitter transporter), member 19, opposite strand | -1.02 |
| ENSMUSG00000039963 | Ccdc40 | coiled-coil domain containing 40 | -1.02 |
| ENSMUSG00000038011 | Dnah10 | dynein, axonemal, heavy chain 10 | -1.02 |
| ENSMUSG00000034786 | Gpsm3 | G-protein signalling modulator 3 (AGS3-like, C. elegans) | -1.02 |
| ENSMUSG00000108818 | Gm44869 | predicted gene 44869 | -1.02 |
| ENSMUSG00000030329 | Pianp | PILR alpha associated neural protein | -1.02 |
| ENSMUSG00000079497 | Gm13420 | predicted gene 13420 | -1.02 |
| ENSMUSG00000018796 | Acsl1 | acyl-CoA synthetase long-chain family member 1 | -1.03 |
| ENSMUSG00000030465 | Psd3 | pleckstrin and Sec7 domain containing 3 | -1.03 |
| ENSMUSG00000020894 | Vamp2 | vesicle-associated membrane protein 2 | -1.03 |

| | | | |
|---|---|---|---|
| ENSMUSG00000039686 | Zer1 | zyg-11 related, cell cycle regulator | -1.03 |
| ENSMUSG00000031808 | Slc27a1 | solute carrier family 27 (fatty acid transporter), member 1 | -1.03 |
| ENSMUSG00000029161 | Cgref1 | cell growth regulator with EF hand domain 1 | -1.03 |
| ENSMUSG00000003363 | Pld3 | phospholipase D family, member 3 | -1.03 |
| ENSMUSG00000034947 | Tmem106a | transmembrane protein 106A | -1.03 |
| ENSMUSG00000112944 | Gm48885 | predicted gene, 48885 | -1.03 |
| ENSMUSG00000026826 | Nr4a2 | nuclear receptor subfamily 4, group A, member 2 | -1.03 |
| ENSMUSG00000044080 | S100a1 | S100 calcium binding protein A1 | -1.03 |
| ENSMUSG00000108954 | Gm44901 | predicted gene 44901 | -1.03 |
| ENSMUSG00000054598 | 9130230L23Rik | RIKEN cDNA 9130230L23 gene | -1.03 |
| ENSMUSG00000074195 | Clca4b | chloride channel accessory 4B | -1.03 |
| ENSMUSG00000029082 | Bst1 | bone marrow stromal cell antigen 1 | -1.03 |
| ENSMUSG00000028785 | Hpca | hippocalcin | -1.03 |
| ENSMUSG00000058163 | Gm5431 | predicted gene 5431 | -1.03 |
| ENSMUSG00000097321 | 1700028E10Rik | RIKEN cDNA 1700028E10 gene | -1.03 |
| ENSMUSG00000067049 | Unc93a | unc-93 homolog A | -1.03 |
| ENSMUSG00000103037 | Pcdhgb1 | protocadherin gamma subfamily B, 1 | -1.03 |
| ENSMUSG00000019055 | Plod1 | procollagen-lysine, 2-oxoglutarate 5-dioxygenase 1 | -1.03 |
| ENSMUSG00000104576 | F830115B05Rik | RIKEN cDNA F830115B05 gene | -1.03 |
| ENSMUSG00000107227 | Gm42559 | predicted gene 42559 | -1.03 |
| ENSMUSG00000072974 | Gm4787 | predicted gene 4787 | -1.03 |
| ENSMUSG00000114891 | Gm47272 | predicted gene, 47272 | -1.03 |
| ENSMUSG00000111447 | Gm48249 | predicted gene, 48249 | -1.03 |
| ENSMUSG00000002033 | Cd3g | CD3 antigen, gamma polypeptide | -1.03 |
| ENSMUSG00000087055 | Gm11948 | predicted gene 11948 | -1.03 |
| ENSMUSG00000054277 | Arfgap3 | ADP-ribosylation factor GTPase activating protein 3 | -1.04 |
| ENSMUSG00000036687 | Tmem184a | transmembrane protein 184a | -1.04 |
| ENSMUSG00000061455 | Stx17 | syntaxin 17 | -1.04 |
| ENSMUSG00000029727 | Cyp3a13 | cytochrome P450, family 3, subfamily a, polypeptide 13 | -1.04 |
| ENSMUSG00000032010 | Usp2 | ubiquitin specific peptidase 2 | -1.04 |
| ENSMUSG00000037826 | Ppm1k | protein phosphatase 1K (PP2C domain containing) | -1.04 |
| ENSMUSG00000029335 | Bmp3 | bone morphogenetic protein 3 | -1.04 |
| ENSMUSG00000042041 | 2010003K11Rik | RIKEN cDNA 2010003K11 gene | -1.04 |
| ENSMUSG00000068349 | Gml | glycosylphosphatidylinositol anchored molecule like | -1.04 |
| ENSMUSG00000019762 | Iyd | iodotyrosine deiodinase | -1.04 |
| ENSMUSG00000047394 | Odf3b | outer dense fiber of sperm tails 3B | -1.04 |
| ENSMUSG00000104535 | Gm42686 | predicted gene 42686 | -1.04 |
| ENSMUSG00000113853 | Gm47583 | predicted gene, 47583 | -1.04 |
| ENSMUSG00000057054 | Inca1 | inhibitor of CDK, cyclin A1 interacting protein 1 | -1.04 |
| ENSMUSG00000083149 | Gm11380 | predicted gene 11380 | -1.04 |
| ENSMUSG00000052364 | B630019K06Rik | RIKEN cDNA B630019K06 gene | -1.04 |
| ENSMUSG00000115186 | Gm49417 | predicted gene, 49417 | -1.04 |
| ENSMUSG00000097566 | A930024N18Rik | RIKEN cDNA A930024N18 gene | -1.04 |
| ENSMUSG00000042106 | Inka1 | inka box actin regulator 1 | -1.04 |
| ENSMUSG00000034563 | Ccpg1 | cell cycle progression 1 | -1.04 |
| ENSMUSG00000099310 | Gm19427 | predicted gene, 19427 | -1.04 |
| ENSMUSG00000049538 | Adamts16 | a disintegrin-like and metallopeptidase (reprolysin type) with thrombospondin type | -1.04 |
| ENSMUSG00000038648 | Creb3l2 | cAMP responsive element binding protein 3-like 2 | -1.05 |
| ENSMUSG00000022575 | Gsdmd | gasdermin D | -1.05 |
| ENSMUSG00000044786 | Zfp36 | zinc finger protein 36 | -1.05 |
| ENSMUSG00000002227 | Mov10 | Moloney leukemia virus 10 | -1.05 |
| ENSMUSG00000025557 | Slc15a1 | solute carrier family 15 (oligopeptide transporter), member 1 | -1.05 |
| ENSMUSG00000028517 | Plpp3 | phospholipid phosphatase 3 | -1.05 |
| ENSMUSG00000060487 | Samd5 | sterile alpha motif domain containing 5 | -1.05 |
| ENSMUSG00000042439 | Zfp532 | zinc finger protein 532 | -1.05 |
| ENSMUSG00000055114 | Anxa13 | annexin A13 | -1.05 |
| ENSMUSG00000047617 | Paxx | non-homologous end joining factor | -1.05 |
| ENSMUSG00000078650 | G6pc | glucose-6-phosphatase, catalytic | -1.05 |
| ENSMUSG00000020121 | Srgap1 | SLIT-ROBO Rho GTPase activating protein 1 | -1.05 |
| ENSMUSG00000043432 | Leng9 | leukocyte receptor cluster (LRC) member 9 | -1.05 |
| ENSMUSG00000030930 | Chst15 | carbohydrate (N-acetylgalactosamine 4-sulfate 6-O) sulfotransferase 15 | -1.05 |
| ENSMUSG00000108181 | C030015A19Rik | RIKEN cDNA C030015A19 gene | -1.05 |
| ENSMUSG00000050640 | Tmem150c | transmembrane protein 150C | -1.05 |
| ENSMUSG00000107549 | Gm43961 | predicted gene, 43961 | -1.05 |
| ENSMUSG00000089636 | 1700058P15Rik | RIKEN cDNA 1700058P15 gene | -1.05 |
| ENSMUSG00000040856 | Dlk1 | delta like non-canonical Notch ligand 1 | -1.05 |
| ENSMUSG00000102148 | Gm38059 | predicted gene, 38059 | -1.05 |

| | | | |
|---|---|---|---|
| ENSMUSG00000103701 | Gm10728 | predicted gene 10728 | -1.05 |
| ENSMUSG00000105083 | Gm42699 | predicted gene 42699 | -1.05 |
| ENSMUSG00000032065 | Tex12 | testis expressed 12 | -1.05 |
| ENSMUSG00000095041 | AC149090.1 | NULL | -1.06 |
| ENSMUSG00000060227 | Casc4 | cancer susceptibility candidate 4 | -1.06 |
| ENSMUSG00000043668 | Tox3 | TOX high mobility group box family member 3 | -1.06 |
| ENSMUSG00000079507 | H2-Q1 | histocompatibility 2, Q region locus 1 | -1.06 |
| ENSMUSG00000028307 | Aldob | aldolase B, fructose-bisphosphate | -1.06 |
| ENSMUSG00000047193 | Dync2h1 | dynein cytoplasmic 2 heavy chain 1 | -1.06 |
| ENSMUSG00000030492 | Slc7a9 | solute carrier family 7 (cationic amino acid transporter, y+ system), member 9 | -1.06 |
| ENSMUSG00000096929 | A330023F24Rik | RIKEN cDNA A330023F24 gene | -1.06 |
| ENSMUSG00000091387 | Gcnt4 | glucosaminyl (N-acetyl) transferase 4, core 2 (beta-1,6-N-acetylglucosaminyltransfe | -1.06 |
| ENSMUSG00000013643 | Lypd8 | LY6/PLAUR domain containing 8 | -1.06 |
| ENSMUSG00000054723 | Vmac | vimentin-type intermediate filament associated coiled-coil protein | -1.06 |
| ENSMUSG00000020641 | Rsad2 | radical S-adenosyl methionine domain containing 2 | -1.06 |
| ENSMUSG00000110277 | Gm45871 | predicted gene 45871 | -1.06 |
| ENSMUSG00000023017 | Asic1 | acid-sensing (proton-gated) ion channel 1 | -1.06 |
| ENSMUSG00000039339 | Mfsd4b2 | major facilitator superfamily domain containing 4B2 | -1.06 |
| ENSMUSG00000050944 | Efcab5 | EF-hand calcium binding domain 5 | -1.06 |
| ENSMUSG00000026888 | Grb14 | growth factor receptor bound protein 14 | -1.06 |
| ENSMUSG00000030107 | Usp18 | ubiquitin specific peptidase 18 | -1.06 |
| ENSMUSG00000022840 | Adcy5 | adenylate cyclase 5 | -1.06 |
| ENSMUSG00000075604 | Cyp11b1 | cytochrome P450, family 11, subfamily b, polypeptide 1 | -1.06 |
| ENSMUSG00000043811 | Rtn4r | reticulon 4 receptor | -1.06 |
| ENSMUSG00000093668 | Pou5f2 | POU domain class 5, transcription factor 2 | -1.06 |
| ENSMUSG00000000686 | Abhd15 | abhydrolase domain containing 15 | -1.06 |
| ENSMUSG00000103632 | Gm38329 | predicted gene, 38329 | -1.06 |
| ENSMUSG00000020609 | Apob | apolipoprotein B | -1.07 |
| ENSMUSG00000033618 | Map3k13 | mitogen-activated protein kinase kinase kinase 13 | -1.07 |
| ENSMUSG00000051359 | Ncald | neurocalcin delta | -1.07 |
| ENSMUSG00000014361 | Mertk | c-mer proto-oncogene tyrosine kinase | -1.07 |
| ENSMUSG00000020142 | Slc1a4 | solute carrier family 1 (glutamate/neutral amino acid transporter), member 4 | -1.07 |
| ENSMUSG00000032080 | Apoa4 | apolipoprotein A-IV | -1.07 |
| ENSMUSG00000052595 | A1cf | APOBEC1 complementation factor | -1.07 |
| ENSMUSG00000102752 | Gm7694 | predicted gene 7694 | -1.07 |
| ENSMUSG00000055897 | Ppp4r1l-ps | protein phosphatase 4, regulatory subunit 1-like, pseudogene | -1.07 |
| ENSMUSG00000017697 | Ada | adenosine deaminase | -1.07 |
| ENSMUSG00000032593 | Amigo3 | adhesion molecule with Ig like domain 3 | -1.07 |
| ENSMUSG00000031886 | Ces2e | carboxylesterase 2E | -1.07 |
| ENSMUSG00000096243 | Gm24265 | predicted gene, 24265 | -1.07 |
| ENSMUSG00000029001 | Fbxo44 | F-box protein 44 | -1.07 |
| ENSMUSG00000079038 | D130040H23Rik | RIKEN cDNA D130040H23 gene | -1.07 |
| ENSMUSG00000045775 | Slc16a5 | solute carrier family 16 (monocarboxylic acid transporters), member 5 | -1.07 |
| ENSMUSG00000031173 | Otc | ornithine transcarbamylase | -1.07 |
| ENSMUSG00000036216 | Leap2 | liver-expressed antimicrobial peptide 2 | -1.07 |
| ENSMUSG00000109852 | Gm45360 | predicted gene 45360 | -1.07 |
| ENSMUSG00000027438 | Napb | N-ethylmaleimide sensitive fusion protein attachment protein beta | -1.07 |
| ENSMUSG00000031445 | Proz | protein Z, vitamin K-dependent plasma glycoprotein | -1.07 |
| ENSMUSG00000103183 | Gm37090 | predicted gene, 37090 | -1.07 |
| ENSMUSG00000112990 | Gm47372 | predicted gene, 47372 | -1.07 |
| ENSMUSG00000114443 | Gm19241 | predicted gene, 19241 | -1.07 |
| ENSMUSG00000041323 | Ak7 | adenylate kinase 7 | -1.07 |
| ENSMUSG00000005493 | Msh4 | mutS homolog 4 | -1.07 |
| ENSMUSG00000095280 | Gm21738 | predicted gene, 21738 | -1.07 |
| ENSMUSG00000086133 | Gm16331 | predicted gene 16331 | -1.07 |
| ENSMUSG00000054901 | Arhgef33 | Rho guanine nucleotide exchange factor (GEF) 33 | -1.07 |
| ENSMUSG00000030275 | Etnk1 | ethanolamine kinase 1 | -1.08 |
| ENSMUSG00000067212 | H2-T23 | histocompatibility 2, T region locus 23 | -1.08 |
| ENSMUSG00000028088 | Fmo5 | flavin containing monooxygenase 5 | -1.08 |
| ENSMUSG00000030155 | Clec2e | C-type lectin domain family 2, member e | -1.08 |
| ENSMUSG00000054422 | Fabp1 | fatty acid binding protein 1, liver | -1.08 |
| ENSMUSG00000039853 | Trim14 | tripartite motif-containing 14 | -1.08 |
| ENSMUSG00000028024 | Enpep | glutamyl aminopeptidase | -1.08 |
| ENSMUSG00000029269 | Sult1b1 | sulfotransferase family 1B, member 1 | -1.08 |
| ENSMUSG00000022938 | Fam3b | family with sequence similarity 3, member B | -1.08 |
| ENSMUSG00000035429 | Ptprh | protein tyrosine phosphatase, receptor type, H | -1.08 |
| ENSMUSG00000039911 | Spsb1 | splA/ryanodine receptor domain and SOCS box containing 1 | -1.08 |

| | | | |
|---|---|---|---|
| ENSMUSG00000005237 | Dnah2 | dynein, axonemal, heavy chain 2 | -1.08 |
| ENSMUSG00000075010 | AW112010 | expressed sequence AW112010 | -1.08 |
| ENSMUSG00000105358 | Gm43099 | predicted gene 43099 | -1.08 |
| ENSMUSG00000050357 | Carmil2 | capping protein regulator and myosin 1 linker 2 | -1.08 |
| ENSMUSG00000023132 | Gzma | granzyme A | -1.08 |
| ENSMUSG00000110928 | Gm48114 | predicted gene, 48114 | -1.08 |
| ENSMUSG00000027489 | Necab3 | N-terminal EF-hand calcium binding protein 3 | -1.08 |
| ENSMUSG00000029054 | Gabrd | gamma-aminobutyric acid (GABA) A receptor, subunit delta | -1.08 |
| ENSMUSG00000103222 | Gm37729 | predicted gene, 37729 | -1.08 |
| ENSMUSG00000022629 | Kif21a | kinesin family member 21A | -1.09 |
| ENSMUSG00000018126 | Baiap2l2 | BAI1-associated protein 2-like 2 | -1.09 |
| ENSMUSG00000035112 | Wnk4 | WNK lysine deficient protein kinase 4 | -1.09 |
| ENSMUSG00000005958 | Ephb3 | Eph receptor B3 | -1.09 |
| ENSMUSG00000006395 | Hyi | hydroxypyruvate isomerase (putative) | -1.09 |
| ENSMUSG00000021676 | Iqgap2 | IQ motif containing GTPase activating protein 2 | -1.09 |
| ENSMUSG00000001225 | Slc26a3 | solute carrier family 26, member 3 | -1.09 |
| ENSMUSG00000043831 | Lysmd4 | LysM, putative peptidoglycan-binding, domain containing 4 | -1.09 |
| ENSMUSG00000038178 | Slc43a2 | solute carrier family 43, member 2 | -1.09 |
| ENSMUSG00000037736 | Limch1 | LIM and calponin homology domains 1 | -1.09 |
| ENSMUSG00000026170 | Cyp27a1 | cytochrome P450, family 27, subfamily a, polypeptide 1 | -1.09 |
| ENSMUSG00000042286 | Stab1 | stabilin 1 | -1.09 |
| ENSMUSG00000025743 | Sdc3 | syndecan 3 | -1.09 |
| ENSMUSG00000040740 | Slc25a34 | solute carrier family 25, member 34 | -1.09 |
| ENSMUSG00000049536 | Tceal1 | transcription elongation factor A (SII)-like 1 | -1.09 |
| ENSMUSG00000026586 | Prrx1 | paired related homeobox 1 | -1.09 |
| ENSMUSG00000038527 | C1rl | complement component 1, r subcomponent-like | -1.09 |
| ENSMUSG00000042417 | Ccno | cyclin O | -1.09 |
| ENSMUSG00000059852 | Kcng2 | potassium voltage-gated channel, subfamily G, member 2 | -1.09 |
| ENSMUSG00000001542 | Ell2 | elongation factor RNA polymerase II 2 | -1.10 |
| ENSMUSG00000027799 | Nbea | neurobeachin | -1.10 |
| ENSMUSG00000029470 | P2rx4 | purinergic receptor P2X, ligand-gated ion channel 4 | -1.10 |
| ENSMUSG00000071356 | Reg3b | regenerating islet-derived 3 beta | -1.10 |
| ENSMUSG00000030650 | Tmc5 | transmembrane channel-like gene family 5 | -1.10 |
| ENSMUSG00000074715 | Ccl28 | chemokine (C-C motif) ligand 28 | -1.10 |
| ENSMUSG00000049858 | Suox | sulfite oxidase | -1.10 |
| ENSMUSG00000024140 | Epas1 | endothelial PAS domain protein 1 | -1.10 |
| ENSMUSG00000022383 | Ppara | peroxisome proliferator activated receptor alpha | -1.10 |
| ENSMUSG00000058063 | Trim31 | tripartite motif-containing 31 | -1.10 |
| ENSMUSG00000068341 | Reg3d | regenerating islet-derived 3 delta | -1.10 |
| ENSMUSG00000032489 | Kif9 | kinesin family member 9 | -1.10 |
| ENSMUSG00000024253 | Dync2li1 | dynein cytoplasmic 2 light intermediate chain 1 | -1.10 |
| ENSMUSG00000100094 | 1810008I18Rik | RIKEN cDNA 1810008I18 gene | -1.10 |
| ENSMUSG00000059013 | Sh2d3c | SH2 domain containing 3C | -1.10 |
| ENSMUSG00000047242 | Taf9b | TATA-box binding protein associated factor 9B | -1.10 |
| ENSMUSG00000087047 | 1700110K17Rik | RIKEN cDNA 1700110K17 gene | -1.10 |
| ENSMUSG00000087691 | Gm15674 | predicted gene 15674 | -1.10 |
| ENSMUSG00000029352 | Crybb3 | crystallin, beta B3 | -1.10 |
| ENSMUSG00000020836 | Coro6 | coronin 6 | -1.10 |
| ENSMUSG00000033487 | Fndc3a | fibronectin type III domain containing 3A | -1.11 |
| ENSMUSG00000028555 | Ttc39a | tetratricopeptide repeat domain 39A | -1.11 |
| ENSMUSG00000020246 | Hcfc2 | host cell factor C2 | -1.11 |
| ENSMUSG00000022040 | Ephx2 | epoxide hydrolase 2, cytoplasmic | -1.11 |
| ENSMUSG00000039899 | Fgl2 | fibrinogen-like protein 2 | -1.11 |
| ENSMUSG00000024190 | Dusp1 | dual specificity phosphatase 1 | -1.11 |
| ENSMUSG00000062638 | Btnl1 | butyrophilin-like 1 | -1.11 |
| ENSMUSG00000010601 | Apol7a | apolipoprotein L 7a | -1.11 |
| ENSMUSG00000036989 | Trim3 | tripartite motif-containing 3 | -1.11 |
| ENSMUSG00000022180 | Slc7a8 | solute carrier family 7 (cationic amino acid transporter, y+ system), member 8 | -1.11 |
| ENSMUSG00000021751 | Acox2 | acyl-Coenzyme A oxidase 2, branched chain | -1.11 |
| ENSMUSG00000099746 | Ppnr | per-pentamer repeat gene | -1.11 |
| ENSMUSG00000037049 | Smpd1 | sphingomyelin phosphodiesterase 1, acid lysosomal | -1.11 |
| ENSMUSG00000024503 | Spink1 | serine peptidase inhibitor, Kazal type 1 | -1.11 |
| ENSMUSG00000038496 | Slc19a3 | solute carrier family 19, member 3 | -1.11 |
| ENSMUSG00000023987 | Pgc | progastricsin (pepsinogen C) | -1.11 |
| ENSMUSG00000029156 | Sgcb | sarcoglycan, beta (dystrophin-associated glycoprotein) | -1.11 |
| ENSMUSG00000039239 | Tgfb2 | transforming growth factor, beta 2 | -1.11 |
| ENSMUSG00000110276 | Gm45330 | predicted gene 45330 | -1.11 |

| | | | |
|---|---|---|---|
| ENSMUSG00000107456 | Gm10400 | predicted gene 10400 | -1.11 |
| ENSMUSG00000090062 | Galnt6os | polypeptide N-acetylgalactosaminyltransferase 6, opposite strand | -1.11 |
| ENSMUSG00000117182 | CT025731.2 | ribosomal protein S6 (Rps6) pseudogene | -1.11 |
| ENSMUSG00000035969 | Rusc2 | RUN and SH3 domain containing 2 | -1.11 |
| ENSMUSG00000023036 | Pcdhgc4 | protocadherin gamma subfamily C, 4 | -1.11 |
| ENSMUSG00000093916 | Gm379 | predicted gene 379 | -1.11 |
| ENSMUSG00000097578 | Gm26798 | predicted gene, 26798 | -1.11 |
| ENSMUSG00000046056 | Sbsn | suprabasin | -1.11 |
| ENSMUSG00000033191 | Tie1 | tyrosine kinase with immunoglobulin-like and EGF-like domains 1 | -1.11 |
| ENSMUSG00000100572 | 2410021H03Rik | RIKEN cDNA 2410021H03 gene | -1.11 |
| ENSMUSG00000086991 | Gm15334 | predicted gene 15334 | -1.11 |
| ENSMUSG00000113250 | Gm48585 | predicted gene, 48585 | -1.11 |
| ENSMUSG00000113864 | Gm48381 | predicted gene, 48381 | -1.11 |
| ENSMUSG00000036151 | Tm6sf2 | transmembrane 6 superfamily member 2 | -1.12 |
| ENSMUSG00000038375 | Trp53inp2 | transformation related protein 53 inducible nuclear protein 2 | -1.12 |
| ENSMUSG00000045038 | Prkce | protein kinase C, epsilon | -1.12 |
| ENSMUSG00000021565 | Slc6a19 | solute carrier family 6 (neurotransmitter transporter), member 19 | -1.12 |
| ENSMUSG00000027219 | Slc28a2 | solute carrier family 28 (sodium-coupled nucleoside transporter), member 2 | -1.12 |
| ENSMUSG00000074604 | Mgst2 | microsomal glutathione S-transferase 2 | -1.12 |
| ENSMUSG00000021062 | Rab15 | RAB15, member RAS oncogene family | -1.12 |
| ENSMUSG00000005107 | Slc2a9 | solute carrier family 2 (facilitated glucose transporter), member 9 | -1.12 |
| ENSMUSG00000097222 | 1010001N08Rik | RIKEN cDNA 1010001N08 gene | -1.12 |
| ENSMUSG00000111375 | Btbd8 | BTB (POZ) domain containing 8 | -1.12 |
| ENSMUSG00000028519 | Dab1 | disabled 1 | -1.12 |
| ENSMUSG00000069805 | Fbp1 | fructose bisphosphatase 1 | -1.12 |
| ENSMUSG00000017453 | Pipox | pipecolic acid oxidase | -1.12 |
| ENSMUSG00000049134 | Nrap | nebulin-related anchoring protein | -1.12 |
| ENSMUSG00000103159 | F830112A20Rik | RIKEN cDNA F830112A20 gene | -1.12 |
| ENSMUSG00000059540 | Tcea2 | transcription elongation factor A (SII), 2 | -1.12 |
| ENSMUSG00000027843 | Ptpn22 | protein tyrosine phosphatase, non-receptor type 22 (lymphoid) | -1.12 |
| ENSMUSG00000040147 | Maob | monoamine oxidase B | -1.12 |
| ENSMUSG00000020573 | Pik3cg | phosphatidylinositol-4,5-bisphosphate 3-kinase catalytic subunit gamma | -1.12 |
| ENSMUSG00000041117 | Ccdc8 | coiled-coil domain containing 8 | -1.12 |
| ENSMUSG00000024901 | Peli3 | pellino 3 | -1.12 |
| ENSMUSG00000104943 | Gm42868 | predicted gene 42868 | -1.12 |
| ENSMUSG00000107878 | Gm44106 | predicted gene, 44106 | -1.12 |
| ENSMUSG00000038418 | Egr1 | early growth response 1 | -1.13 |
| ENSMUSG00000035506 | Slc12a8 | solute carrier family 12 (potassium/chloride transporters), member 8 | -1.13 |
| ENSMUSG00000065954 | Tacc1 | transforming, acidic coiled-coil containing protein 1 | -1.13 |
| ENSMUSG00000025408 | Ddit3 | DNA-damage inducible transcript 3 | -1.13 |
| ENSMUSG00000006342 | Susd2 | sushi domain containing 2 | -1.13 |
| ENSMUSG00000035919 | Bbs9 | Bardet-Biedl syndrome 9 (human) | -1.13 |
| ENSMUSG00000074622 | Mafb | v-maf musculoaponeurotic fibrosarcoma oncogene family, protein B (avian) | -1.13 |
| ENSMUSG00000077167 | Gm24119 | predicted gene, 24119 | -1.13 |
| ENSMUSG00000110803 | Gm20275 | predicted gene, 20275 | -1.13 |
| ENSMUSG00000069713 | 4933406P04Rik | RIKEN cDNA 4933406P04 gene | -1.13 |
| ENSMUSG00000030306 | Tmtc1 | transmembrane and tetratricopeptide repeat containing 1 | -1.13 |
| ENSMUSG00000052125 | F730043M19Rik | RIKEN cDNA F730043M19 gene | -1.13 |
| ENSMUSG00000104211 | Gm37985 | predicted gene, 37985 | -1.13 |
| ENSMUSG00000022564 | Grina | glutamate receptor, ionotropic, N-methyl D-aspartate-associated protein 1 (glutam | -1.14 |
| ENSMUSG00000028207 | Asph | aspartate-beta-hydroxylase | -1.14 |
| ENSMUSG00000037280 | Galnt6 | polypeptide N-acetylgalactosaminyltransferase 6 | -1.14 |
| ENSMUSG00000022750 | Klhl22 | kelch-like 22 | -1.14 |
| ENSMUSG00000036745 | Ttll7 | tubulin tyrosine ligase-like family, member 7 | -1.14 |
| ENSMUSG00000029811 | Aoc1 | amine oxidase, copper-containing 1 | -1.14 |
| ENSMUSG00000022574 | Naprt | nicotinate phosphoribosyltransferase | -1.14 |
| ENSMUSG00000067813 | Xkr9 | X-linked Kx blood group related 9 | -1.14 |
| ENSMUSG00000040389 | Wdr47 | WD repeat domain 47 | -1.14 |
| ENSMUSG00000001156 | Mxd1 | MAX dimerization protein 1 | -1.14 |
| ENSMUSG00000035121 | Neil2 | nei like 2 (E. coli) | -1.14 |
| ENSMUSG00000110170 | St6galnac2 | ST6 (alpha-N-acetyl-neuraminyl-2,3-beta-galactosyl-1,3)-N-acetylgalactosaminide a | -1.14 |
| ENSMUSG00000059824 | Dbp | D site albumin promoter binding protein | -1.14 |
| ENSMUSG00000112276 | 5033421B08Rik | RIKEN cDNA 5033421B08 gene | -1.14 |
| ENSMUSG00000098439 | Hm629797 | cDNA sequence HM629797 | -1.14 |
| ENSMUSG00000081245 | Gm6587 | predicted gene 6587 | -1.14 |
| ENSMUSG00000041180 | Hectd2 | HECT domain E3 ubiquitin protein ligase 2 | -1.14 |
| ENSMUSG00000042066 | Tmcc2 | transmembrane and coiled-coil domains 2 | -1.14 |

| | | | |
|---|---|---|---|
| ENSMUSG00000034377 | Tulp4 | tubby like protein 4 | -1.15 |
| ENSMUSG00000026589 | Sec16b | SEC16 homolog B (S. cerevisiae) | -1.15 |
| ENSMUSG00000019970 | Sgk1 | serum/glucocorticoid regulated kinase 1 | -1.15 |
| ENSMUSG00000031378 | Abcd1 | ATP-binding cassette, sub-family D (ALD), member 1 | -1.15 |
| ENSMUSG00000041695 | Kcnj2 | potassium inwardly-rectifying channel, subfamily J, member 2 | -1.15 |
| ENSMUSG00000021620 | Acot12 | acyl-CoA thioesterase 12 | -1.15 |
| ENSMUSG00000108460 | Gm39041 | predicted gene, 39041 | -1.15 |
| ENSMUSG00000074196 | Clca4c-ps | chloride channel accessory 4C, pseudogene | -1.15 |
| ENSMUSG00000024885 | Aldh3b1 | aldehyde dehydrogenase 3 family, member B1 | -1.15 |
| ENSMUSG00000025064 | Col17a1 | collagen, type XVII, alpha 1 | -1.15 |
| ENSMUSG00000040441 | Slc26a10 | solute carrier family 26, member 10 | -1.15 |
| ENSMUSG00000062064 | Slc2a7 | solute carrier family 2 (facilitated glucose transporter), member 7 | -1.15 |
| ENSMUSG00000033847 | Pla2g4c | phospholipase A2, group IVC (cytosolic, calcium-independent) | -1.15 |
| ENSMUSG00000112592 | Gm19972 | predicted gene, 19972 | -1.15 |
| ENSMUSG00000045062 | Pcdhb7 | protocadherin beta 7 | -1.15 |
| ENSMUSG00000038860 | Garnl3 | GTPase activating RANGAP domain-like 3 | -1.15 |
| ENSMUSG00000024451 | Arap3 | ArfGAP with RhoGAP domain, ankyrin repeat and PH domain 3 | -1.15 |
| ENSMUSG00000102967 | Gm37946 | predicted gene, 37946 | -1.15 |
| ENSMUSG00000022783 | Spag6l | sperm associated antigen 6-like | -1.15 |
| ENSMUSG00000108693 | Gm45153 | predicted gene 45153 | -1.15 |
| ENSMUSG00000070436 | Serpinh1 | serine (or cysteine) peptidase inhibitor, clade H, member 1 | -1.15 |
| ENSMUSG00000030838 | Ush1c | USH1 protein network component harmonin | -1.16 |
| ENSMUSG00000020600 | Slc7a15 | solute carrier family 7 (cationic amino acid transporter, y+ system), member 15 | -1.16 |
| ENSMUSG00000020681 | Ace | angiotensin I converting enzyme (peptidyl-dipeptidase A) 1 | -1.16 |
| ENSMUSG00000020261 | Slc36a1 | solute carrier family 36 (proton/amino acid symporter), member 1 | -1.16 |
| ENSMUSG00000023960 | Enpp5 | ectonucleotide pyrophosphatase/phosphodiesterase 5 | -1.16 |
| ENSMUSG00000032454 | Rbp2 | retinol binding protein 2, cellular | -1.16 |
| ENSMUSG00000033917 | Gde1 | glycerophosphodiester phosphodiesterase 1 | -1.16 |
| ENSMUSG00000032105 | Pdzd3 | PDZ domain containing 3 | -1.16 |
| ENSMUSG00000039878 | Slc39a5 | solute carrier family 39 (metal ion transporter), member 5 | -1.16 |
| ENSMUSG00000107109 | Gm42571 | predicted gene 42571 | -1.16 |
| ENSMUSG00000109585 | Gm45358 | predicted gene 45358 | -1.16 |
| ENSMUSG00000034450 | Gulo | gulonolactone (L-) oxidase | -1.16 |
| ENSMUSG00000029168 | Dpysl5 | dihydropyrimidinase-like 5 | -1.16 |
| ENSMUSG00000000253 | Gmpr | guanosine monophosphate reductase | -1.16 |
| ENSMUSG00000112955 | Gm48889 | predicted gene, 48889 | -1.16 |
| ENSMUSG00000085774 | Gm13055 | predicted gene 13055 | -1.16 |
| ENSMUSG00000113102 | A930040O22Rik | RIKEN cDNA A930040O22 gene | -1.16 |
| ENSMUSG00000029162 | Khk | ketohexokinase | -1.17 |
| ENSMUSG00000035847 | Ids | iduronate 2-sulfatase | -1.17 |
| ENSMUSG00000024131 | Slc3a1 | solute carrier family 3, member 1 | -1.17 |
| ENSMUSG00000043822 | Adamtsl5 | ADAMTS-like 5 | -1.17 |
| ENSMUSG00000079445 | B3gnt7 | UDP-GlcNAc:betaGal beta-1,3-N-acetylglucosaminyltransferase 7 | -1.17 |
| ENSMUSG00000055148 | Klf2 | Kruppel-like factor 2 (lung) | -1.17 |
| ENSMUSG00000097277 | 2900076A07Rik | RIKEN cDNA 2900076A07 gene | -1.17 |
| ENSMUSG00000102423 | Gm37465 | predicted gene, 37465 | -1.17 |
| ENSMUSG00000104693 | Gm42941 | predicted gene 42941 | -1.17 |
| ENSMUSG00000042770 | Hebp1 | heme binding protein 1 | -1.17 |
| ENSMUSG00000043760 | Pkhd1 | polycystic kidney and hepatic disease 1 | -1.17 |
| ENSMUSG00000051503 | Gm6583 | predicted gene 6583 | -1.17 |
| ENSMUSG00000084798 | 4930533B01Rik | RIKEN cDNA 4930533B01 gene | -1.17 |
| ENSMUSG00000054822 | 1700041G16Rik | RIKEN cDNA 1700041G16 gene | -1.17 |
| ENSMUSG00000113679 | Gm10432 | predicted gene 10432 | -1.17 |
| ENSMUSG00000079993 | Gm11799 | predicted gene 11799 | -1.17 |
| ENSMUSG00000039545 | Flicr | Foxp3 regulating long intergenic noncoding RNA | -1.17 |
| ENSMUSG00000100009 | Gm7967 | predicted gene 7967 | -1.17 |
| ENSMUSG00000048970 | C1galt1c1 | C1GALT1-specific chaperone 1 | -1.18 |
| ENSMUSG00000032120 | C2cd2l | C2 calcium-dependent domain containing 2-like | -1.18 |
| ENSMUSG00000025497 | Cdhr5 | cadherin-related family member 5 | -1.18 |
| ENSMUSG00000023886 | Smoc2 | SPARC related modular calcium binding 2 | -1.18 |
| ENSMUSG00000002228 | Ppm1j | protein phosphatase 1J | -1.18 |
| ENSMUSG00000037139 | Myom3 | myomesin family, member 3 | -1.18 |
| ENSMUSG00000042684 | Npl | N-acetylneuraminate pyruvate lyase | -1.18 |
| ENSMUSG00000022861 | Dgkg | diacylglycerol kinase, gamma | -1.18 |
| ENSMUSG00000106596 | Gm42822 | predicted gene 42822 | -1.18 |
| ENSMUSG00000044906 | 4930503L19Rik | RIKEN cDNA 4930503L19 gene | -1.18 |
| ENSMUSG00000026969 | Fam166a | family with sequence similarity 166, member A | -1.18 |

| | | | |
|---|---|---|---|
| ENSMUSG00000032482 | Cspg5 | chondroitin sulfate proteoglycan 5 | -1.18 |
| ENSMUSG00000029830 | Svopl | SV2 related protein homolog (rat)-like | -1.18 |
| ENSMUSG00000036960 | Clca2 | chloride channel accessory 2 | -1.18 |
| ENSMUSG00000092268 | Gm20540 | predicted gene 20540 | -1.18 |
| ENSMUSG00000018672 | Copz2 | coatomer protein complex, subunit zeta 2 | -1.18 |
| ENSMUSG00000062542 | Syt9 | synaptotagmin IX | -1.18 |
| ENSMUSG00000021070 | Bdkrb2 | bradykinin receptor, beta 2 | -1.18 |
| ENSMUSG00000090061 | Nwd2 | NACHT and WD repeat domain containing 2 | -1.18 |
| ENSMUSG00000020484 | Xbp1 | X-box binding protein 1 | -1.19 |
| ENSMUSG00000015094 | Npdc1 | neural proliferation, differentiation and control 1 | -1.19 |
| ENSMUSG00000019960 | Dusp6 | dual specificity phosphatase 6 | -1.19 |
| ENSMUSG00000091705 | H2-Q2 | histocompatibility 2, Q region locus 2 | -1.19 |
| ENSMUSG00000024313 | Mep1b | meprin 1 beta | -1.19 |
| ENSMUSG00000029802 | Abcg2 | ATP binding cassette subfamily G member 2 (Junior blood group) | -1.19 |
| ENSMUSG00000032098 | Treh | trehalase (brush-border membrane glycoprotein) | -1.19 |
| ENSMUSG00000038751 | Ptk6 | PTK6 protein tyrosine kinase 6 | -1.19 |
| ENSMUSG00000035713 | Usp35 | ubiquitin specific peptidase 35 | -1.19 |
| ENSMUSG00000029630 | Cyp3a25 | cytochrome P450, family 3, subfamily a, polypeptide 25 | -1.19 |
| ENSMUSG00000027983 | Cyp2u1 | cytochrome P450, family 2, subfamily u, polypeptide 1 | -1.19 |
| ENSMUSG00000039084 | Chad | chondroadherin | -1.19 |
| ENSMUSG00000038074 | Fkbp14 | FK506 binding protein 14 | -1.19 |
| ENSMUSG00000025092 | Hspa12a | heat shock protein 12A | -1.19 |
| ENSMUSG00000105701 | Gm42587 | predicted gene 42587 | -1.19 |
| ENSMUSG00000029369 | Afm | afamin | -1.19 |
| ENSMUSG00000026211 | Obsl1 | obscurin-like 1 | -1.19 |
| ENSMUSG00000103785 | Gm35025 | predicted gene, 35025 | -1.19 |
| ENSMUSG00000085900 | A930041C12Rik | RIKEN cDNA A930041C12 gene | -1.19 |
| ENSMUSG00000003469 | Phyhip | phytanoyl-CoA hydroxylase interacting protein | -1.19 |
| ENSMUSG00000022708 | Zbtb20 | zinc finger and BTB domain containing 20 | -1.20 |
| ENSMUSG00000022844 | Pdia5 | protein disulfide isomerase associated 5 | -1.20 |
| ENSMUSG00000027792 | Bche | butyrylcholinesterase | -1.20 |
| ENSMUSG00000021238 | Aldh6a1 | aldehyde dehydrogenase family 6, subfamily A1 | -1.20 |
| ENSMUSG00000032842 | Abcc10 | ATP-binding cassette, sub-family C (CFTR/MRP), member 10 | -1.20 |
| ENSMUSG00000039620 | Trmt9b | tRNA methyltransferase 9B | -1.20 |
| ENSMUSG00000107171 | Gm42572 | predicted gene 42572 | -1.20 |
| ENSMUSG00000056133 | Unc93a2 | unc-93 homolog A2 | -1.20 |
| ENSMUSG00000057265 | Bbof1 | basal body orientation factor 1 | -1.20 |
| ENSMUSG00000102440 | Pcdhga9 | protocadherin gamma subfamily A, 9 | -1.20 |
| ENSMUSG00000085615 | A330035P11Rik | RIKEN cDNA A330035P11 gene | -1.20 |
| ENSMUSG00000040473 | Cfap69 | cilia and flagella associated protein 69 | -1.20 |
| ENSMUSG00000028681 | Ptch2 | patched 2 | -1.20 |
| ENSMUSG00000038700 | Hoxb5 | homeobox B5 | -1.20 |
| ENSMUSG00000014529 | Tmbim7 | transmembrane BAX inhibitor motif containing 7 | -1.20 |
| ENSMUSG00000060988 | Galnt13 | polypeptide N-acetylgalactosaminyltransferase 13 | -1.20 |
| ENSMUSG00000108489 | Gm45025 | predicted gene 45025 | -1.20 |
| ENSMUSG00000040963 | Asgr2 | asialoglycoprotein receptor 2 | -1.20 |
| ENSMUSG00000019564 | Arid3a | AT rich interactive domain 3A (BRIGHT-like) | -1.21 |
| ENSMUSG00000032741 | Tpcn1 | two pore channel 1 | -1.21 |
| ENSMUSG00000034918 | Cdhr2 | cadherin-related family member 2 | -1.21 |
| ENSMUSG00000052302 | Tbc1d30 | TBC1 domain family, member 30 | -1.21 |
| ENSMUSG00000070461 | 9230112E08Rik | RIKEN cDNA 9230112E08 gene | -1.21 |
| ENSMUSG00000084911 | Gm16185 | predicted gene 16185 | -1.21 |
| ENSMUSG00000086844 | B230206H07Rik | RIKEN cDNA B230206H07 gene | -1.21 |
| ENSMUSG00000032081 | Apoc3 | apolipoprotein C-III | -1.21 |
| ENSMUSG00000031972 | Acta1 | actin, alpha 1, skeletal muscle | -1.21 |
| ENSMUSG00000104156 | Gm38102 | predicted gene, 38102 | -1.21 |
| ENSMUSG00000086199 | Bcas3os1 | breast carcinoma amplified sequence 3, opposite strand 1 | -1.21 |
| ENSMUSG00000089028 | Gm25010 | predicted gene, 25010 | -1.21 |
| ENSMUSG00000087143 | A830082K12Rik | RIKEN cDNA A830082K12 gene | -1.21 |
| ENSMUSG00000047495 | Dlgap2 | DLG associated protein 2 | -1.21 |
| ENSMUSG00000046447 | Camk2n1 | calcium/calmodulin-dependent protein kinase II inhibitor 1 | -1.22 |
| ENSMUSG00000035722 | Abca7 | ATP-binding cassette, sub-family A (ABC1), member 7 | -1.22 |
| ENSMUSG00000037685 | Atp8a1 | ATPase, aminophospholipid transporter (APLT), class I, type 8A, member 1 | -1.22 |
| ENSMUSG00000052512 | Nav2 | neuron navigator 2 | -1.22 |
| ENSMUSG00000073399 | Trim40 | tripartite motif-containing 40 | -1.22 |
| ENSMUSG00000026222 | Sp100 | nuclear antigen Sp100 | -1.22 |
| ENSMUSG00000029607 | Ankrd61 | ankyrin repeat domain 61 | -1.22 |

| | | | |
|---|---|---|---|
| ENSMUSG00000034687 | Fras1 | Fraser extracellular matrix complex subunit 1 | -1.22 |
| ENSMUSG00000019359 | Gdpd2 | glycerophosphodiester phosphodiesterase domain containing 2 | -1.22 |
| ENSMUSG00000030600 | Lrfn1 | leucine rich repeat and fibronectin type III domain containing 1 | -1.22 |
| ENSMUSG00000031766 | Slc12a3 | solute carrier family 12, member 3 | -1.22 |
| ENSMUSG00000007783 | Cpt1c | carnitine palmitoyltransferase 1c | -1.22 |
| ENSMUSG00000101462 | Gm3052 | predicted gene 3052 | -1.22 |
| ENSMUSG00000064413 | Gm22245 | predicted gene, 22245 | -1.22 |
| ENSMUSG00000110767 | Gm31432 | predicted gene, 31432 | -1.22 |
| ENSMUSG00000084052 | Gm11745 | predicted gene 11745 | -1.22 |
| ENSMUSG00000105344 | Gm43556 | predicted gene 43556 | -1.22 |
| ENSMUSG00000025469 | Msx3 | msh homeobox 3 | -1.22 |
| ENSMUSG00000025498 | Irf7 | interferon regulatory factor 7 | -1.23 |
| ENSMUSG00000026113 | Inpp4a | inositol polyphosphate-4-phosphatase, type I | -1.23 |
| ENSMUSG00000048249 | Crebrf | CREB3 regulatory factor | -1.23 |
| ENSMUSG00000044860 | Gm1123 | predicted gene 1123 | -1.23 |
| ENSMUSG00000024254 | Abcg8 | ATP binding cassette subfamily G member 8 | -1.23 |
| ENSMUSG00000029370 | Rassf6 | Ras association (RalGDS/AF-6) domain family member 6 | -1.23 |
| ENSMUSG00000026614 | Slc30a10 | solute carrier family 30, member 10 | -1.23 |
| ENSMUSG00000047641 | Krt87 | keratin 87 | -1.23 |
| ENSMUSG00000085747 | Slc13a2os | solute carrier family 13 (sodium-dependent dicarboxylate transporter), member 2, | -1.23 |
| ENSMUSG00000042010 | Acacb | acetyl-Coenzyme A carboxylase beta | -1.23 |
| ENSMUSG00000103693 | Gm37529 | predicted gene, 37529 | -1.23 |
| ENSMUSG00000048347 | Pcdhb18 | protocadherin beta 18 | -1.23 |
| ENSMUSG00000021268 | Meg3 | maternally expressed 3 | -1.23 |
| ENSMUSG00000022766 | Serpind1 | serine (or cysteine) peptidase inhibitor, clade D, member 1 | -1.23 |
| ENSMUSG00000111063 | Zkscan7 | zinc finger with KRAB and SCAN domains 7 | -1.23 |
| ENSMUSG00000113780 | Gm33195 | predicted gene, 33195 | -1.23 |
| ENSMUSG00000047033 | Pcdhb15 | protocadherin beta 15 | -1.23 |
| ENSMUSG00000030865 | Chp2 | calcineurin-like EF hand protein 2 | -1.24 |
| ENSMUSG00000050747 | Trim15 | tripartite motif-containing 15 | -1.24 |
| ENSMUSG00000025140 | Pycr1 | pyrroline-5-carboxylate reductase 1 | -1.24 |
| ENSMUSG00000073889 | Il11ra1 | interleukin 11 receptor, alpha chain 1 | -1.24 |
| ENSMUSG00000032289 | Thsd4 | thrombospondin, type I, domain containing 4 | -1.24 |
| ENSMUSG00000032036 | Kirrel3 | kirre like nephrin family adhesion molecule 3 | -1.24 |
| ENSMUSG00000103472 | Pcdhga7 | protocadherin gamma subfamily A, 7 | -1.24 |
| ENSMUSG00000103567 | Pcdhga5 | protocadherin gamma subfamily A, 5 | -1.24 |
| ENSMUSG00000113918 | Gm6566 | predicted gene 6566 | -1.24 |
| ENSMUSG00000086507 | Adap2os | ArfGAP with dual PH domains 2, opposite strand | -1.24 |
| ENSMUSG00000084890 | A830036E02Rik | RIKEN cDNA A830036E02 gene | -1.24 |
| ENSMUSG00000000392 | Fap | fibroblast activation protein | -1.24 |
| ENSMUSG00000086533 | Mypopos | Myb-related transcription factor, partner of profilin, opposite strand | -1.24 |
| ENSMUSG00000109780 | Gm45447 | predicted gene 45447 | -1.24 |
| ENSMUSG00000109869 | Gm39158 | predicted gene, 39158 | -1.24 |
| ENSMUSG00000021919 | Chat | choline acetyltransferase | -1.24 |
| ENSMUSG00000048562 | Sp8 | trans-acting transcription factor 8 | -1.24 |
| ENSMUSG00000106450 | Gm43018 | predicted gene 43018 | -1.24 |
| ENSMUSG00000108076 | 9530086O07Rik | RIKEN cDNA 9530086O07 gene | -1.24 |
| ENSMUSG00000038894 | Irs2 | insulin receptor substrate 2 | -1.25 |
| ENSMUSG00000005580 | Adcy9 | adenylate cyclase 9 | -1.25 |
| ENSMUSG00000037822 | Smim14 | small integral membrane protein 14 | -1.25 |
| ENSMUSG00000078439 | Smim24 | small integral membrane protein 24 | -1.25 |
| ENSMUSG00000048473 | Sult6b2 | sulfotransferase family 6B, member 2 | -1.25 |
| ENSMUSG00000022540 | Rogdi | rogdi homolog | -1.25 |
| ENSMUSG00000043621 | Ubxn10 | UBX domain protein 10 | -1.25 |
| ENSMUSG00000094830 | n-R5s194 | nuclear encoded rRNA 5S 194 | -1.25 |
| ENSMUSG00000040432 | Ltb4r2 | leukotriene B4 receptor 2 | -1.25 |
| ENSMUSG00000107813 | Gm44434 | predicted gene, 44434 | -1.25 |
| ENSMUSG00000103432 | 6720464F23Rik | RIKEN cDNA 6720464F23 gene | -1.25 |
| ENSMUSG00000055493 | Epm2a | epilepsy, progressive myoclonic epilepsy, type 2 gene alpha | -1.25 |
| ENSMUSG00000073680 | Tmem88b | transmembrane protein 88B | -1.25 |
| ENSMUSG00000038143 | Stox2 | storkhead box 2 | -1.25 |
| ENSMUSG00000113041 | Gm48302 | predicted gene, 48302 | -1.25 |
| ENSMUSG00000113852 | Gm48062 | predicted gene, 48062 | -1.25 |
| ENSMUSG00000033768 | Nrxn2 | neurexin II | -1.25 |
| ENSMUSG00000038576 | Susd4 | sushi domain containing 4 | -1.25 |
| ENSMUSG00000032083 | Apoa1 | apolipoprotein A-I | -1.26 |
| ENSMUSG00000028015 | Ctso | cathepsin O | -1.26 |

| | | | |
|---|---|---|---|
| ENSMUSG00000032596 | Uba7 | ubiquitin-like modifier activating enzyme 7 | -1.26 |
| ENSMUSG00000033147 | Slc22a15 | solute carrier family 22 (organic anion/cation transporter), member 15 | -1.26 |
| ENSMUSG00000036473 | Tbc1d24 | TBC1 domain family, member 24 | -1.26 |
| ENSMUSG00000033987 | Dnah17 | dynein, axonemal, heavy chain 17 | -1.26 |
| ENSMUSG00000074653 | Lrrc31 | leucine rich repeat containing 31 | -1.26 |
| ENSMUSG00000051043 | Gprc5c | G protein-coupled receptor, family C, group 5, member C | -1.26 |
| ENSMUSG00000033721 | Vav3 | vav 3 oncogene | -1.26 |
| ENSMUSG00000097443 | Gm17529 | predicted gene, 17529 | -1.26 |
| ENSMUSG00000034579 | Pla2g3 | phospholipase A2, group III | -1.26 |
| ENSMUSG00000073403 | Gm10499 | predicted gene 10499 | -1.26 |
| ENSMUSG00000054477 | Kcnn2 | potassium intermediate/small conductance calcium-activated channel, subfamily N | -1.26 |
| ENSMUSG00000052767 | Gm12703 | predicted gene 12703 | -1.26 |
| ENSMUSG00000113370 | Gm46348 | predicted gene, 46348 | -1.26 |
| ENSMUSG00000110366 | Gm45885 | predicted gene 45885 | -1.26 |
| ENSMUSG00000025207 | Sema4g | sema domain, immunoglobulin domain (Ig), transmembrane domain (TM) and shor | -1.27 |
| ENSMUSG00000022546 | Gpt | glutamic pyruvic transaminase, soluble | -1.27 |
| ENSMUSG00000032038 | St3gal4 | ST3 beta-galactoside alpha-2,3-sialyltransferase 4 | -1.27 |
| ENSMUSG00000032652 | Crebl2 | cAMP responsive element binding protein-like 2 | -1.27 |
| ENSMUSG00000056602 | Fry | FRY microtubule binding protein | -1.27 |
| ENSMUSG00000027513 | Pck1 | phosphoenolpyruvate carboxykinase 1, cytosolic | -1.27 |
| ENSMUSG00000108291 | Gm44292 | predicted gene, 44292 | -1.27 |
| ENSMUSG00000094377 | Gm24407 | predicted gene, 24407 | -1.27 |
| ENSMUSG00000074261 | Erich4 | glutamate rich 4 | -1.27 |
| ENSMUSG00000028427 | Aqp7 | aquaporin 7 | -1.27 |
| ENSMUSG00000048895 | Cdk5r1 | cyclin-dependent kinase 5, regulatory subunit 1 (p35) | -1.27 |
| ENSMUSG00000032420 | Nt5e | 5' nucleotidase, ecto | -1.27 |
| ENSMUSG00000098269 | Mir8094 | microRNA 8094 | -1.27 |
| ENSMUSG00000085903 | Gm15340 | predicted gene 15340 | -1.27 |
| ENSMUSG00000042532 | Golga7b | golgi autoantigen, golgin subfamily a, 7B | -1.27 |
| ENSMUSG00000102466 | Gm38200 | predicted gene, 38200 | -1.27 |
| ENSMUSG00000103897 | Pcdhga8 | protocadherin gamma subfamily A, 8 | -1.27 |
| ENSMUSG00000108075 | Gm45061 | predicted gene 45061 | -1.27 |
| ENSMUSG00000041481 | Serpina3g | serine (or cysteine) peptidase inhibitor, clade A, member 3G | -1.27 |
| ENSMUSG00000086708 | Gm15577 | predicted gene 15577 | -1.27 |
| ENSMUSG00000106516 | Gm42441 | predicted gene 42441 | -1.27 |
| ENSMUSG00000113535 | AU015791 | expressed sequence AU015791 | -1.27 |
| ENSMUSG00000029992 | Gfpt1 | glutamine fructose-6-phosphate transaminase 1 | -1.28 |
| ENSMUSG00000092341 | Malat1 | metastasis associated lung adenocarcinoma transcript 1 (non-coding RNA) | -1.28 |
| ENSMUSG00000060002 | Chpt1 | choline phosphotransferase 1 | -1.28 |
| ENSMUSG00000004655 | Aqp1 | aquaporin 1 | -1.28 |
| ENSMUSG00000006435 | Neurl1a | neuralized E3 ubiquitin protein ligase 1A | -1.28 |
| ENSMUSG00000028017 | Egf | epidermal growth factor | -1.28 |
| ENSMUSG00000029163 | Emilin1 | elastin microfibril interfacer 1 | -1.28 |
| ENSMUSG00000079355 | Ackr4 | atypical chemokine receptor 4 | -1.28 |
| ENSMUSG00000085925 | Rtl1 | retrotransposon Gaglike 1 | -1.28 |
| ENSMUSG00000009654 | Oit3 | oncoprotein induced transcript 3 | -1.28 |
| ENSMUSG00000038522 | Mfsd4b1 | major facilitator superfamily domain containing 4B1 | -1.28 |
| ENSMUSG00000020169 | Best3 | bestrophin 3 | -1.28 |
| ENSMUSG00000052469 | Tcp10c | t-complex protein 10c | -1.28 |
| ENSMUSG00000038305 | Spats2l | spermatogenesis associated, serine-rich 2-like | -1.29 |
| ENSMUSG00000005373 | Mlxipl | MLX interacting protein-like | -1.29 |
| ENSMUSG00000013275 | Slc41a1 | solute carrier family 41, member 1 | -1.29 |
| ENSMUSG00000102964 | 9430034N14Rik | RIKEN cDNA 9430034N14 gene | -1.29 |
| ENSMUSG00000051606 | 2010001K21Rik | RIKEN cDNA 2010001K21 gene | -1.29 |
| ENSMUSG00000110068 | Gm45294 | predicted gene 45294 | -1.29 |
| ENSMUSG00000040794 | C1qtnf4 | C1q and tumor necrosis factor related protein 4 | -1.29 |
| ENSMUSG00000021725 | Parp8 | poly (ADP-ribose) polymerase family, member 8 | -1.29 |
| ENSMUSG00000087530 | Gm15533 | predicted gene 15533 | -1.29 |
| ENSMUSG00000112234 | Gm48427 | predicted gene, 48427 | -1.29 |
| ENSMUSG00000040265 | Dnm3 | dynamin 3 | -1.29 |
| ENSMUSG00000105470 | Gm42589 | predicted gene 42589 | -1.29 |
| ENSMUSG00000112462 | Gm48772 | predicted gene, 48772 | -1.29 |
| ENSMUSG00000028777 | Gnat3 | guanine nucleotide binding protein, alpha transducing 3 | -1.29 |
| ENSMUSG00000026807 | Ak8 | adenylate kinase 8 | -1.29 |
| ENSMUSG00000021190 | Lgmn | legumain | -1.30 |
| ENSMUSG00000023045 | Soat2 | sterol O-acyltransferase 2 | -1.30 |
| ENSMUSG00000040270 | Bach2 | BTB and CNC homology, basic leucine zipper transcription factor 2 | -1.30 |

| | | | |
|---|---|---|---|
| ENSMUSG00000066258 | Trim12a | tripartite motif-containing 12A | -1.30 |
| ENSMUSG00000040721 | Zfhx2 | zinc finger homeobox 2 | -1.30 |
| ENSMUSG00000059708 | Akap17b | A kinase (PRKA) anchor protein 17B | -1.30 |
| ENSMUSG00000038058 | Nod1 | nucleotide-binding oligomerization domain containing 1 | -1.30 |
| ENSMUSG00000037940 | Inpp4b | inositol polyphosphate-4-phosphatase, type II | -1.30 |
| ENSMUSG00000110187 | Gm45496 | predicted gene 45496 | -1.30 |
| ENSMUSG00000068600 | Gml2 | glycosylphosphatidylinositol anchored molecule like 2 | -1.30 |
| ENSMUSG00000085884 | Gm15342 | predicted gene 15342 | -1.30 |
| ENSMUSG00000084329 | Gm6733 | predicted gene 6733 | -1.30 |
| ENSMUSG00000116180 | Gm49492 | predicted gene, 49492 | -1.30 |
| ENSMUSG00000036437 | Npy1r | neuropeptide Y receptor Y1 | -1.30 |
| ENSMUSG00000083603 | Gm14810 | predicted gene 14810 | -1.30 |
| ENSMUSG00000014905 | Dnajb9 | DnaJ heat shock protein family (Hsp40) member B9 | -1.31 |
| ENSMUSG00000030017 | Reg3g | regenerating islet-derived 3 gamma | -1.31 |
| ENSMUSG00000031570 | Plpp5 | phospholipid phosphatase 5 | -1.31 |
| ENSMUSG00000034449 | Dhrs11 | dehydrogenase/reductase (SDR family) member 11 | -1.31 |
| ENSMUSG00000004952 | Rasa4 | RAS p21 protein activator 4 | -1.31 |
| ENSMUSG00000053334 | Ficd | FIC domain containing | -1.31 |
| ENSMUSG00000095915 | n-R5s115 | nuclear encoded rRNA 5S 115 | -1.31 |
| ENSMUSG00000023092 | Fhl1 | four and a half LIM domains 1 | -1.31 |
| ENSMUSG00000097147 | Gm26780 | predicted gene, 26780 | -1.31 |
| ENSMUSG00000019737 | Syne4 | spectrin repeat containing, nuclear envelope family member 4 | -1.31 |
| ENSMUSG00000047115 | Fam221a | family with sequence similarity 221, member A | -1.31 |
| ENSMUSG00000052783 | Grk4 | G protein-coupled receptor kinase 4 | -1.31 |
| ENSMUSG00000029195 | Klb | klotho beta | -1.31 |
| ENSMUSG00000062017 | Abca14 | ATP-binding cassette, sub-family A (ABC1), member 14 | -1.31 |
| ENSMUSG00000113811 | Gm47882 | predicted gene, 47882 | -1.31 |
| ENSMUSG00000036995 | Asap3 | ArfGAP with SH3 domain, ankyrin repeat and PH domain 3 | -1.31 |
| ENSMUSG00000108246 | Gm43896 | predicted gene, 43896 | -1.31 |
| ENSMUSG00000096549 | Prickle4 | prickle planar cell polarity protein 4 | -1.31 |
| ENSMUSG00000026070 | Il18r1 | interleukin 18 receptor 1 | -1.31 |
| ENSMUSG00000028185 | Dnase2b | deoxyribonuclease II beta | -1.31 |
| ENSMUSG00000036587 | Fut7 | fucosyltransferase 7 | -1.31 |
| ENSMUSG00000102404 | 5530400K19Rik | RIKEN cDNA 5530400K19 gene | -1.31 |
| ENSMUSG00000026825 | Dnm1 | dynamin 1 | -1.32 |
| ENSMUSG00000019066 | Rab3d | RAB3D, member RAS oncogene family | -1.32 |
| ENSMUSG00000027790 | Sis | sucrase isomaltase (alpha-glucosidase) | -1.32 |
| ENSMUSG00000015944 | Castor2 | cytosolic arginine sensor for mTORC1 subunit 2 | -1.32 |
| ENSMUSG00000002504 | Slc9a3r2 | solute carrier family 9 (sodium/hydrogen exchanger), member 3 regulator 2 | -1.32 |
| ENSMUSG00000079516 | Reg3a | regenerating islet-derived 3 alpha | -1.32 |
| ENSMUSG00000031155 | Pim2 | proviral integration site 2 | -1.32 |
| ENSMUSG00000046387 | Pcdhb17 | protocadherin beta 17 | -1.32 |
| ENSMUSG00000036298 | Slc2a13 | solute carrier family 2 (facilitated glucose transporter), member 13 | -1.32 |
| ENSMUSG00000046191 | Pcdhb20 | protocadherin beta 20 | -1.32 |
| ENSMUSG00000032656 | 37681 | membrane-associated ring finger (C3HC4) 3 | -1.32 |
| ENSMUSG00000049246 | Hspe1-ps2 | heat shock protein 1 (chaperonin 10), pseudogene 2 | -1.32 |
| ENSMUSG00000096039 | D830030K20Rik | RIKEN cDNA D830030K20 gene | -1.32 |
| ENSMUSG00000031230 | Fgf16 | fibroblast growth factor 16 | -1.32 |
| ENSMUSG00000035561 | Aldh1b1 | aldehyde dehydrogenase 1 family, member B1 | -1.33 |
| ENSMUSG00000023951 | Vegfa | vascular endothelial growth factor A | -1.33 |
| ENSMUSG00000053054 | Adh6a | alcohol dehydrogenase 6A (class V) | -1.33 |
| ENSMUSG00000042675 | Ypel3 | yippee like 3 | -1.33 |
| ENSMUSG00000046546 | Fam43a | family with sequence similarity 43, member A | -1.33 |
| ENSMUSG00000020709 | Adap2 | ArfGAP with dual PH domains 2 | -1.33 |
| ENSMUSG00000022197 | Pdzd2 | PDZ domain containing 2 | -1.33 |
| ENSMUSG00000052135 | Foxo6 | forkhead box O6 | -1.33 |
| ENSMUSG00000000402 | Egfl6 | EGF-like-domain, multiple 6 | -1.33 |
| ENSMUSG00000079499 | 6530402F18Rik | RIKEN cDNA 6530402F18 gene | -1.33 |
| ENSMUSG00000033029 | 1700088E04Rik | RIKEN cDNA 1700088E04 gene | -1.33 |
| ENSMUSG00000112338 | Gm47693 | predicted gene, 47693 | -1.33 |
| ENSMUSG00000024965 | Fermt3 | fermitin family member 3 | -1.33 |
| ENSMUSG00000025083 | Afap1l2 | actin filament associated protein 1-like 2 | -1.33 |
| ENSMUSG00000112596 | Gm48804 | predicted gene, 48804 | -1.33 |
| ENSMUSG00000084137 | Gm9085 | predicted gene 9085 | -1.33 |
| ENSMUSG00000035279 | Ssc5d | scavenger receptor cysteine rich family, 5 domains | -1.33 |
| ENSMUSG00000078451 | Ppil6 | peptidylprolyl isomerase (cyclophilin)-like 6 | -1.33 |
| ENSMUSG00000073242 | Dnmt3aos | DNA methyltransferase 3A, opposite strand | -1.33 |

| | | | |
|---|---|---|---|
| ENSMUSG00000052188 | Gm14964 | predicted gene 14964 | -1.33 |
| ENSMUSG00000001334 | Fndc5 | fibronectin type III domain containing 5 | -1.33 |
| ENSMUSG00000041642 | Kif21b | kinesin family member 21B | -1.34 |
| ENSMUSG00000062995 | Ica1 | islet cell autoantigen 1 | -1.34 |
| ENSMUSG00000032776 | Mctp2 | multiple C2 domains, transmembrane 2 | -1.34 |
| ENSMUSG00000042428 | Mgat3 | mannoside acetylglucosaminyltransferase 3 | -1.34 |
| ENSMUSG00000020447 | Npc1l1 | NPC1 like intracellular cholesterol transporter 1 | -1.34 |
| ENSMUSG00000026610 | Esrrg | estrogen-related receptor gamma | -1.34 |
| ENSMUSG00000035493 | Tgfbi | transforming growth factor, beta induced | -1.34 |
| ENSMUSG00000020614 | Fam20a | family with sequence similarity 20, member A | -1.34 |
| ENSMUSG00000045094 | Arhgef37 | Rho guanine nucleotide exchange factor (GEF) 37 | -1.34 |
| ENSMUSG00000042312 | S100a13 | S100 calcium binding protein A13 | -1.34 |
| ENSMUSG00000101365 | Gm19325 | predicted gene, 19325 | -1.34 |
| ENSMUSG00000086189 | Gm15462 | predicted gene 15462 | -1.34 |
| ENSMUSG00000047227 | Gm527 | predicted gene 527 | -1.34 |
| ENSMUSG00000107094 | Gm43294 | predicted gene 43294 | -1.34 |
| ENSMUSG00000104020 | Gm37215 | predicted gene, 37215 | -1.34 |
| ENSMUSG00000043719 | Col6a6 | collagen, type VI, alpha 6 | -1.34 |
| ENSMUSG00000113203 | Gm47587 | predicted gene, 47587 | -1.34 |
| ENSMUSG00000032899 | Styk1 | serine/threonine/tyrosine kinase 1 | -1.35 |
| ENSMUSG00000038312 | Edem2 | ER degradation enhancer, mannosidase alpha-like 2 | -1.35 |
| ENSMUSG00000042632 | Pla2g6 | phospholipase A2, group VI | -1.35 |
| ENSMUSG00000028943 | Espn | espin | -1.35 |
| ENSMUSG00000034312 | Iqsec1 | IQ motif and Sec7 domain 1 | -1.35 |
| ENSMUSG00000058921 | Slc10a5 | solute carrier family 10 (sodium/bile acid cotransporter family), member 5 | -1.35 |
| ENSMUSG00000030657 | Xylt1 | xylosyltransferase 1 | -1.35 |
| ENSMUSG00000051427 | Ccdc157 | coiled-coil domain containing 157 | -1.35 |
| ENSMUSG00000056025 | Clca3a1 | chloride channel accessory 3A1 | -1.35 |
| ENSMUSG00000033066 | Gas7 | growth arrest specific 7 | -1.35 |
| ENSMUSG00000113119 | Gm48883 | predicted gene, 48883 | -1.35 |
| ENSMUSG00000049556 | Lingo1 | leucine rich repeat and Ig domain containing 1 | -1.35 |
| ENSMUSG00000063564 | Col23a1 | collagen, type XXIII, alpha 1 | -1.35 |
| ENSMUSG00000020793 | Galr2 | galanin receptor 2 | -1.35 |
| ENSMUSG00000110357 | A030001D20Rik | RIKEN cDNA A030001D20 gene | -1.35 |
| ENSMUSG00000038311 | 2410017I17Rik | RIKEN cDNA 2410017I17 gene | -1.35 |
| ENSMUSG00000028782 | Adgrb2 | adhesion G protein-coupled receptor B2 | -1.35 |
| ENSMUSG00000041559 | Fmod | fibromodulin | -1.35 |
| ENSMUSG00000082207 | Gm12998 | predicted gene 12998 | -1.35 |
| ENSMUSG00000040929 | Rfx3 | regulatory factor X, 3 (influences HLA class II expression) | -1.36 |
| ENSMUSG00000024713 | Pcsk5 | proprotein convertase subtilisin/kexin type 5 | -1.36 |
| ENSMUSG00000053862 | Slc51b | solute carrier family 51, beta subunit | -1.36 |
| ENSMUSG00000055730 | Ces2a | carboxylesterase 2A | -1.36 |
| ENSMUSG00000044626 | Liph | lipase, member H | -1.36 |
| ENSMUSG00000008999 | Bmp7 | bone morphogenetic protein 7 | -1.36 |
| ENSMUSG00000068735 | Trp53i11 | transformation related protein 53 inducible protein 11 | -1.36 |
| ENSMUSG00000102732 | Gm37342 | predicted gene, 37342 | -1.36 |
| ENSMUSG00000023467 | Tulp2 | tubby-like protein 2 | -1.36 |
| ENSMUSG00000116506 | 5730414N17Rik | RIKEN cDNA 5730414N17 gene | -1.36 |
| ENSMUSG00000093580 | Gm20706 | predicted gene 20706 | -1.36 |
| ENSMUSG00000114302 | Gm48683 | predicted gene, 48683 | -1.36 |
| ENSMUSG00000021250 | Fos | FBJ osteosarcoma oncogene | -1.37 |
| ENSMUSG00000052085 | Dock8 | dedicator of cytokinesis 8 | -1.37 |
| ENSMUSG00000037005 | Xpnpep2 | X-prolyl aminopeptidase (aminopeptidase P) 2, membrane-bound | -1.37 |
| ENSMUSG00000028051 | Hcn3 | hyperpolarization-activated, cyclic nucleotide-gated K+ 3 | -1.37 |
| ENSMUSG00000032839 | Trpc1 | transient receptor potential cation channel, subfamily C, member 1 | -1.37 |
| ENSMUSG00000075270 | Pde11a | phosphodiesterase 11A | -1.37 |
| ENSMUSG00000022686 | B3gnt5 | UDP-GlcNAc:betaGal beta-1,3-N-acetylglucosaminyltransferase 5 | -1.37 |
| ENSMUSG00000003282 | Plag1 | pleiomorphic adenoma gene 1 | -1.37 |
| ENSMUSG00000107197 | Gm43312 | predicted gene 43312 | -1.37 |
| ENSMUSG00000075014 | Gm10800 | predicted gene 10800 | -1.37 |
| ENSMUSG00000041729 | Coro2b | coronin, actin binding protein, 2B | -1.37 |
| ENSMUSG00000084744 | Gm25291 | predicted gene, 25291 | -1.37 |
| ENSMUSG00000063929 | Cyp4a32 | cytochrome P450, family 4, subfamily a, polypeptide 32 | -1.37 |
| ENSMUSG00000087341 | 0610040F04Rik | RIKEN cDNA 0610040F04 gene | -1.37 |
| ENSMUSG00000042800 | Spata46 | spermatogenesis associated 46 | -1.37 |
| ENSMUSG00000052889 | Prkcb | protein kinase C, beta | -1.37 |
| ENSMUSG00000038540 | Tmc3 | transmembrane channel-like gene family 3 | -1.37 |

| | | | |
|---|---|---|---|
| ENSMUSG00000104343 | 5730408A14Rik | RIKEN cDNA 5730408A14 gene | -1.37 |
| ENSMUSG00000105975 | Gm42609 | predicted gene 42609 | -1.37 |
| ENSMUSG00000032564 | Cpne4 | copine IV | -1.37 |
| ENSMUSG00000019789 | Hey2 | hairy/enhancer-of-split related with YRPW motif 2 | -1.37 |
| ENSMUSG00000062991 | Nrg1 | neuregulin 1 | -1.37 |
| ENSMUSG00000023259 | Slc26a6 | solute carrier family 26, member 6 | -1.38 |
| ENSMUSG00000020080 | Hkdc1 | hexokinase domain containing 1 | -1.38 |
| ENSMUSG00000024340 | Btnl2 | butyrophilin-like 2 | -1.38 |
| ENSMUSG00000073420 | Btnl5-ps | butyrophilin-like 5, pseudogene | -1.38 |
| ENSMUSG00000003032 | Klf4 | Kruppel-like factor 4 (gut) | -1.38 |
| ENSMUSG00000052353 | Cemip | cell migration inducing protein, hyaluronan binding | -1.38 |
| ENSMUSG00000037921 | Ddx60 | DEAD (Asp-Glu-Ala-Asp) box polypeptide 60 | -1.38 |
| ENSMUSG00000096349 | Gm22513 | predicted gene, 22513 | -1.38 |
| ENSMUSG00000000296 | Tpd52l1 | tumor protein D52-like 1 | -1.38 |
| ENSMUSG00000025268 | Maged2 | melanoma antigen, family D, 2 | -1.38 |
| ENSMUSG00000045312 | Lhfpl2 | lipoma HMGIC fusion partner-like 2 | -1.38 |
| ENSMUSG00000068323 | Slc4a5 | solute carrier family 4, sodium bicarbonate cotransporter, member 5 | -1.38 |
| ENSMUSG00000020604 | Arsg | arylsulfatase G | -1.38 |
| ENSMUSG00000038541 | Srd5a2 | steroid 5 alpha-reductase 2 | -1.38 |
| ENSMUSG00000018620 | Mmp20 | matrix metallopeptidase 20 (enamelysin) | -1.38 |
| ENSMUSG00000103793 | Pcdhga6 | protocadherin gamma subfamily A, 6 | -1.38 |
| ENSMUSG00000009092 | Derl3 | Der1-like domain family, member 3 | -1.38 |
| ENSMUSG00000006784 | Ttc25 | tetratricopeptide repeat domain 25 | -1.38 |
| ENSMUSG00000102224 | 4930447F24Rik | RIKEN cDNA 4930447F24 gene | -1.38 |
| ENSMUSG00000021101 | 4930408O17Rik | RIKEN cDNA 4930408O17 gene | -1.38 |
| ENSMUSG00000045518 | Onecut3 | one cut domain, family member 3 | -1.38 |
| ENSMUSG00000027634 | Ndrg3 | N-myc downstream regulated gene 3 | -1.39 |
| ENSMUSG00000028836 | Slc30a2 | solute carrier family 30 (zinc transporter), member 2 | -1.39 |
| ENSMUSG00000034981 | Parm1 | prostate androgen-regulated mucin-like protein 1 | -1.39 |
| ENSMUSG00000078612 | Fyb2 | FYN binding protein 2 | -1.39 |
| ENSMUSG00000034614 | Pik3ip1 | phosphoinositide-3-kinase interacting protein 1 | -1.39 |
| ENSMUSG00000061462 | Obscn | obscurin, cytoskeletal calmodulin and titin-interacting RhoGEF | -1.39 |
| ENSMUSG00000033389 | Arhgap44 | Rho GTPase activating protein 44 | -1.39 |
| ENSMUSG00000114576 | Naip3-ps1 | NLR family, apoptosis inhibitory protein 3, pseudogene 1 | -1.39 |
| ENSMUSG00000032679 | Cd59a | CD59a antigen | -1.39 |
| ENSMUSG00000116262 | Gm49544 | predicted gene, 49544 | -1.39 |
| ENSMUSG00000039543 | Cfap70 | cilia and flagella associated protein 70 | -1.39 |
| ENSMUSG00000086258 | Gm15239 | predicted gene 15239 | -1.39 |
| ENSMUSG00000097911 | Gm26691 | predicted gene, 26691 | -1.39 |
| ENSMUSG00000048540 | Nhlh2 | nescient helix loop helix 2 | -1.39 |
| ENSMUSG00000026815 | Gfi1b | growth factor independent 1B | -1.39 |
| ENSMUSG00000025006 | Sorbs1 | sorbin and SH3 domain containing 1 | -1.40 |
| ENSMUSG00000031618 | Nr3c2 | nuclear receptor subfamily 3, group C, member 2 | -1.40 |
| ENSMUSG00000014773 | Dll1 | delta like canonical Notch ligand 1 | -1.40 |
| ENSMUSG00000035390 | Brsk1 | BR serine/threonine kinase 1 | -1.40 |
| ENSMUSG00000042589 | Cux2 | cut-like homeobox 2 | -1.40 |
| ENSMUSG00000060224 | Pyroxd2 | pyridine nucleotide-disulphide oxidoreductase domain 2 | -1.40 |
| ENSMUSG00000063239 | Grm4 | glutamate receptor, metabotropic 4 | -1.40 |
| ENSMUSG00000022340 | Sybu | syntabulin (syntaxin-interacting) | -1.40 |
| ENSMUSG00000002633 | Shh | sonic hedgehog | -1.40 |
| ENSMUSG00000036611 | Eepd1 | endonuclease/exonuclease/phosphatase family domain containing 1 | -1.41 |
| ENSMUSG00000022048 | Dpysl2 | dihydropyrimidinase-like 2 | -1.41 |
| ENSMUSG00000038473 | Nos1ap | nitric oxide synthase 1 (neuronal) adaptor protein | -1.41 |
| ENSMUSG00000030659 | Nucb2 | nucleobindin 2 | -1.41 |
| ENSMUSG00000057880 | Abat | 4-aminobutyrate aminotransferase | -1.41 |
| ENSMUSG00000041420 | Meis3 | Meis homeobox 3 | -1.41 |
| ENSMUSG00000020882 | Cacnb1 | calcium channel, voltage-dependent, beta 1 subunit | -1.41 |
| ENSMUSG00000073733 | Rsg1 | REM2 and RAB-like small GTPase 1 | -1.41 |
| ENSMUSG00000005986 | Ankrd13d | ankyrin repeat domain 13 family, member D | -1.41 |
| ENSMUSG00000105873 | Gm43708 | predicted gene 43708 | -1.41 |
| ENSMUSG00000020715 | Ern1 | endoplasmic reticulum (ER) to nucleus signalling 1 | -1.42 |
| ENSMUSG00000038967 | Pdk2 | pyruvate dehydrogenase kinase, isoenzyme 2 | -1.42 |
| ENSMUSG00000054708 | Ankrd24 | ankyrin repeat domain 24 | -1.42 |
| ENSMUSG00000047496 | Rnf152 | ring finger protein 152 | -1.42 |
| ENSMUSG00000040181 | Fmo1 | flavin containing monooxygenase 1 | -1.42 |
| ENSMUSG00000026452 | Syt2 | synaptotagmin II | -1.42 |
| ENSMUSG00000028750 | Pla2g2c | phospholipase A2, group IIC | -1.42 |

| | | | |
|---|---|---|---|
| ENSMUSG00000086391 | 1700042O10Rik | RIKEN cDNA 1700042O10 gene | -1.42 |
| ENSMUSG00000106384 | Gm43188 | predicted gene 43188 | -1.42 |
| ENSMUSG00000074439 | Defa5 | defensin, alpha, 5 | -1.42 |
| ENSMUSG00000028167 | Bdh2 | 3-hydroxybutyrate dehydrogenase, type 2 | -1.42 |
| ENSMUSG00000036452 | Arhgap26 | Rho GTPase activating protein 26 | -1.43 |
| ENSMUSG00000028150 | Rorc | RAR-related orphan receptor gamma | -1.43 |
| ENSMUSG00000110060 | Gm9860 | predicted gene 9860 | -1.43 |
| ENSMUSG00000061728 | Btnl7-ps | butyrophilin-like 7, pseudogene | -1.43 |
| ENSMUSG00000073406 | H2-Bl | histocompatibility 2, blastocyst | -1.43 |
| ENSMUSG00000103194 | Gm37643 | predicted gene, 37643 | -1.43 |
| ENSMUSG00000104351 | Gm37125 | predicted gene, 37125 | -1.43 |
| ENSMUSG00000113966 | Gm47980 | predicted gene, 47980 | -1.43 |
| ENSMUSG00000090214 | Gm15657 | predicted gene 15657 | -1.44 |
| ENSMUSG00000092618 | Btnl6 | butyrophilin-like 6 | -1.44 |
| ENSMUSG00000021208 | Ifi27l2b | interferon, alpha-inducible protein 27 like 2B | -1.44 |
| ENSMUSG00000048027 | Rgmb | repulsive guidance molecule family member B | -1.44 |
| ENSMUSG00000026347 | Tmem163 | transmembrane protein 163 | -1.44 |
| ENSMUSG00000038042 | Ptpdc1 | protein tyrosine phosphatase domain containing 1 | -1.44 |
| ENSMUSG00000111271 | Gm48127 | predicted gene, 48127 | -1.44 |
| ENSMUSG00000079157 | Fam155a | family with sequence similarity 155, member A | -1.44 |
| ENSMUSG00000022946 | Dopey2 | dopey family member 2 | -1.45 |
| ENSMUSG00000022026 | Olfm4 | olfactomedin 4 | -1.45 |
| ENSMUSG00000026131 | Dst | dystonin | -1.45 |
| ENSMUSG00000035041 | Creb3l3 | cAMP responsive element binding protein 3-like 3 | -1.45 |
| ENSMUSG00000040505 | Abcg5 | ATP binding cassette subfamily G member 5 | -1.45 |
| ENSMUSG00000029273 | Sult1d1 | sulfotransferase family 1D, member 1 | -1.45 |
| ENSMUSG00000031613 | Hpgd | hydroxyprostaglandin dehydrogenase 15 (NAD) | -1.45 |
| ENSMUSG00000034570 | Inpp5j | inositol polyphosphate 5-phosphatase J | -1.45 |
| ENSMUSG00000094651 | Gal3st2 | galactose-3-O-sulfotransferase 2 | -1.45 |
| ENSMUSG00000029650 | Slc46a3 | solute carrier family 46, member 3 | -1.45 |
| ENSMUSG00000051146 | Camk2n2 | calcium/calmodulin-dependent protein kinase II inhibitor 2 | -1.45 |
| ENSMUSG00000045348 | Nyap1 | neuronal tyrosine-phosphorylated phosphoinositide 3-kinase adaptor 1 | -1.45 |
| ENSMUSG00000011256 | Adam19 | a disintegrin and metallopeptidase domain 19 (meltrin beta) | -1.45 |
| ENSMUSG00000089874 | 9230117E06Rik | RIKEN cDNA 9230117E06 gene | -1.45 |
| ENSMUSG00000015243 | Abca1 | ATP-binding cassette, sub-family A (ABC1), member 1 | -1.46 |
| ENSMUSG00000066621 | Tecpr1 | tectonin beta-propeller repeat containing 1 | -1.46 |
| ENSMUSG00000037390 | Muc3 | mucin 3, intestinal | -1.46 |
| ENSMUSG00000043079 | Synpo | synaptopodin | -1.46 |
| ENSMUSG00000048572 | Tmem252 | transmembrane protein 252 | -1.46 |
| ENSMUSG00000025082 | Vwa2 | von Willebrand factor A domain containing 2 | -1.46 |
| ENSMUSG00000027805 | Pfn2 | profilin 2 | -1.46 |
| ENSMUSG00000037493 | Cib2 | calcium and integrin binding family member 2 | -1.46 |
| ENSMUSG00000108476 | Gm44974 | predicted gene 44974 | -1.46 |
| ENSMUSG00000074115 | Saa1 | serum amyloid A 1 | -1.46 |
| ENSMUSG00000072915 | Gm12258 | predicted gene 12258 | -1.46 |
| ENSMUSG00000024986 | Hhex | hematopoietically expressed homeobox | -1.46 |
| ENSMUSG00000078949 | R3hdml | R3H domain containing-like | -1.46 |
| ENSMUSG00000042564 | Fam227a | family with sequence similarity 227, member A | -1.46 |
| ENSMUSG00000084950 | Gm5577 | predicted gene 5577 | -1.46 |
| ENSMUSG00000072683 | Gm7457 | predicted gene 7457 | -1.46 |
| ENSMUSG00000092482 | Gm20531 | predicted gene 20531 | -1.46 |
| ENSMUSG00000046242 | Nme9 | NME/NM23 family member 9 | -1.46 |
| ENSMUSG00000028139 | Riiad1 | regulatory subunit of type II PKA R-subunit (RIIa) domain containing 1 | -1.46 |
| ENSMUSG00000063873 | Slc24a3 | solute carrier family 24 (sodium/potassium/calcium exchanger), member 3 | -1.46 |
| ENSMUSG00000022416 | Cacna1i | calcium channel, voltage-dependent, alpha 1I subunit | -1.46 |
| ENSMUSG00000032925 | Itgbl1 | integrin, beta-like 1 | -1.46 |
| ENSMUSG00000034731 | Dgkh | diacylglycerol kinase, eta | -1.47 |
| ENSMUSG00000010307 | Tmem86a | transmembrane protein 86A | -1.47 |
| ENSMUSG00000084803 | 5830444B04Rik | RIKEN cDNA 5830444B04 gene | -1.47 |
| ENSMUSG00000100183 | Gm28512 | predicted gene 28512 | -1.47 |
| ENSMUSG00000112501 | Gm33091 | predicted gene, 33091 | -1.47 |
| ENSMUSG00000032690 | Oas2 | 2'-5' oligoadenylate synthetase 2 | -1.47 |
| ENSMUSG00000073400 | Trim10 | tripartite motif-containing 10 | -1.47 |
| ENSMUSG00000037235 | Mxd4 | Max dimerization protein 4 | -1.48 |
| ENSMUSG00000031770 | Herpud1 | homocysteine-inducible, endoplasmic reticulum stress-inducible, ubiquitin-like dom | -1.48 |
| ENSMUSG00000036813 | Entpd8 | ectonucleoside triphosphate diphosphohydrolase 8 | -1.48 |
| ENSMUSG00000040327 | Cul9 | cullin 9 | -1.48 |

| | | | |
|---|---|---|---|
| ENSMUSG00000075590 | Nrbp2 | nuclear receptor binding protein 2 | -1.48 |
| ENSMUSG00000030717 | Nupr1 | nuclear protein transcription regulator 1 | -1.48 |
| ENSMUSG00000025196 | Cpn1 | carboxypeptidase N, polypeptide 1 | -1.48 |
| ENSMUSG00000037664 | Cdkn1c | cyclin-dependent kinase inhibitor 1C (P57) | -1.48 |
| ENSMUSG00000090293 | Gm17034 | predicted gene 17034 | -1.48 |
| ENSMUSG00000015879 | Fam184b | family with sequence similarity 184, member B | -1.48 |
| ENSMUSG00000018341 | Il12rb2 | interleukin 12 receptor, beta 2 | -1.48 |
| ENSMUSG00000112887 | Gm47961 | predicted gene, 47961 | -1.48 |
| ENSMUSG00000112556 | Gm47339 | predicted gene, 47339 | -1.48 |
| ENSMUSG00000007021 | Syngr3 | synaptogyrin 3 | -1.48 |
| ENSMUSG00000052520 | Cyp2j5 | cytochrome P450, family 2, subfamily j, polypeptide 5 | -1.48 |
| ENSMUSG00000036782 | Klhl13 | kelch-like 13 | -1.48 |
| ENSMUSG00000071019 | Sdr16c6 | short chain dehydrogenase/reductase family 16C, member 6 | -1.48 |
| ENSMUSG00000037977 | 6430571L13Rik | RIKEN cDNA 6430571L13 gene | -1.48 |
| ENSMUSG00000039234 | Sec24d | Sec24 related gene family, member D (S. cerevisiae) | -1.49 |
| ENSMUSG00000034427 | Myo15b | myosin XVB | -1.49 |
| ENSMUSG00000052117 | D630039A03Rik | RIKEN cDNA D630039A03 gene | -1.49 |
| ENSMUSG00000036814 | Slc6a20a | solute carrier family 6 (neurotransmitter transporter), member 20A | -1.49 |
| ENSMUSG00000005268 | Prlr | prolactin receptor | -1.49 |
| ENSMUSG00000085333 | 1700030A11Rik | RIKEN cDNA 1700030A11 gene | -1.49 |
| ENSMUSG00000036882 | Arhgap33 | Rho GTPase activating protein 33 | -1.49 |
| ENSMUSG00000005267 | Zfp287 | zinc finger protein 287 | -1.49 |
| ENSMUSG00000048481 | Mypop | Myb-related transcription factor, partner of profilin | -1.49 |
| ENSMUSG00000074981 | Dcdc5 | doublecortin domain containing 5 | -1.49 |
| ENSMUSG00000029185 | Fam114a1 | family with sequence similarity 114, member A1 | -1.50 |
| ENSMUSG00000009739 | Pou6f1 | POU domain, class 6, transcription factor 1 | -1.50 |
| ENSMUSG00000070315 | 4930581F22Rik | RIKEN cDNA 4930581F22 gene | -1.50 |
| ENSMUSG00000015468 | Notch4 | notch 4 | -1.50 |
| ENSMUSG00000047656 | Trpt1 | tRNA phosphotransferase 1 | -1.50 |
| ENSMUSG00000055805 | Fmnl1 | formin-like 1 | -1.50 |
| ENSMUSG00000078161 | Erich3 | glutamate rich 3 | -1.50 |
| ENSMUSG00000040016 | Ptger3 | prostaglandin E receptor 3 (subtype EP3) | -1.50 |
| ENSMUSG00000026940 | Ccdc183 | coiled-coil domain containing 183 | -1.50 |
| ENSMUSG00000106542 | Gm43410 | predicted gene 43410 | -1.50 |
| ENSMUSG00000011751 | Sptbn4 | spectrin beta, non-erythrocytic 4 | -1.51 |
| ENSMUSG00000047910 | Pcdhb16 | protocadherin beta 16 | -1.51 |
| ENSMUSG00000033276 | Stk36 | serine/threonine kinase 36 | -1.51 |
| ENSMUSG00000024065 | Ehd3 | EH-domain containing 3 | -1.51 |
| ENSMUSG00000103144 | Pcdhga1 | protocadherin gamma subfamily A, 1 | -1.51 |
| ENSMUSG00000100071 | Gm28707 | predicted gene 28707 | -1.51 |
| ENSMUSG00000059991 | Nptx2 | neuronal pentraxin 2 | -1.52 |
| ENSMUSG00000027950 | Chrnb2 | cholinergic receptor, nicotinic, beta polypeptide 2 (neuronal) | -1.52 |
| ENSMUSG00000044681 | Cnpy1 | canopy FGF signaling regulator 1 | -1.52 |
| ENSMUSG00000103332 | Pcdhga2 | protocadherin gamma subfamily A, 2 | -1.52 |
| ENSMUSG00000052584 | Serp2 | stress-associated endoplasmic reticulum protein family member 2 | -1.52 |
| ENSMUSG00000106045 | Gm42996 | predicted gene 42996 | -1.52 |
| ENSMUSG00000041119 | Pde9a | phosphodiesterase 9A | -1.53 |
| ENSMUSG00000041351 | Rap1gap | Rap1 GTPase-activating protein | -1.53 |
| ENSMUSG00000022537 | Tmem44 | transmembrane protein 44 | -1.53 |
| ENSMUSG00000044067 | Gpr22 | G protein-coupled receptor 22 | -1.53 |
| ENSMUSG00000087651 | 1500009L16Rik | RIKEN cDNA 1500009L16 gene | -1.53 |
| ENSMUSG00000055862 | Izumo4 | IZUMO family member 4 | -1.53 |
| ENSMUSG00000027801 | Tm4sf4 | transmembrane 4 superfamily member 4 | -1.53 |
| ENSMUSG00000111296 | 2010001M07Rik | RIKEN cDNA 2010001M07 gene | -1.53 |
| ENSMUSG00000102533 | Gm37226 | predicted gene, 37226 | -1.53 |
| ENSMUSG00000021219 | Rgs6 | regulator of G-protein signaling 6 | -1.53 |
| ENSMUSG00000114626 | Gm48266 | predicted gene, 48266 | -1.53 |
| ENSMUSG00000027227 | Sord | sorbitol dehydrogenase | -1.54 |
| ENSMUSG00000026380 | Tfcp2l1 | transcription factor CP2-like 1 | -1.54 |
| ENSMUSG00000106847 | Peg13 | paternally expressed 13 | -1.54 |
| ENSMUSG00000006345 | Ggt1 | gamma-glutamyltransferase 1 | -1.54 |
| ENSMUSG00000049690 | Nckap5 | NCK-associated protein 5 | -1.54 |
| ENSMUSG00000037818 | Abhd18 | abhydrolase domain containing 18 | -1.54 |
| ENSMUSG00000020072 | Pbld2 | phenazine biosynthesis-like protein domain containing 2 | -1.54 |
| ENSMUSG00000031840 | Rab3a | RAB3A, member RAS oncogene family | -1.54 |
| ENSMUSG00000033898 | Cfhr2 | complement factor H-related 2 | -1.54 |
| ENSMUSG00000094605 | Gm25873 | predicted gene, 25873 | -1.54 |

| | | | |
|---|---|---|---|
| ENSMUSG00000028713 | Cyp4b1 | cytochrome P450, family 4, subfamily b, polypeptide 1 | -1.55 |
| ENSMUSG00000021336 | Slc17a4 | solute carrier family 17 (sodium phosphate), member 4 | -1.55 |
| ENSMUSG00000030364 | Clec2h | C-type lectin domain family 2, member h | -1.55 |
| ENSMUSG00000033308 | Dpyd | dihydropyrimidine dehydrogenase | -1.55 |
| ENSMUSG00000035274 | Tpbg | trophoblast glycoprotein | -1.55 |
| ENSMUSG00000046314 | Stxbp6 | syntaxin binding protein 6 (amisyn) | -1.55 |
| ENSMUSG00000005873 | Reep5 | receptor accessory protein 5 | -1.55 |
| ENSMUSG00000103585 | Pcdhgb4 | protocadherin gamma subfamily B, 4 | -1.55 |
| ENSMUSG00000015053 | Gata2 | GATA binding protein 2 | -1.55 |
| ENSMUSG00000092035 | Peg10 | paternally expressed 10 | -1.55 |
| ENSMUSG00000020732 | Rab37 | RAB37, member RAS oncogene family | -1.55 |
| ENSMUSG00000070777 | Ceacam20 | carcinoembryonic antigen-related cell adhesion molecule 20 | -1.56 |
| ENSMUSG00000030223 | Ptpro | protein tyrosine phosphatase, receptor type, O | -1.56 |
| ENSMUSG00000024924 | Vldlr | very low density lipoprotein receptor | -1.56 |
| ENSMUSG00000078490 | Cfap74 | cilia and flagella associated protein 74 | -1.56 |
| ENSMUSG00000072572 | Slc39a2 | solute carrier family 39 (zinc transporter), member 2 | -1.56 |
| ENSMUSG00000018378 | Cuedc1 | CUE domain containing 1 | -1.57 |
| ENSMUSG00000002565 | Scin | scinderin | -1.57 |
| ENSMUSG00000074207 | Adh1 | alcohol dehydrogenase 1 (class I) | -1.57 |
| ENSMUSG00000102758 | Naaladl2 | N-acetylated alpha-linked acidic dipeptidase-like 2 | -1.57 |
| ENSMUSG00000097494 | 4933406C10Rik | RIKEN cDNA 4933406C10 gene | -1.57 |
| ENSMUSG00000022683 | Pla2g10 | phospholipase A2, group X | -1.57 |
| ENSMUSG00000074892 | B3galt5 | UDP-Gal:betaGlcNAc beta 1,3-galactosyltransferase, polypeptide 5 | -1.58 |
| ENSMUSG00000023809 | Rps6ka2 | ribosomal protein S6 kinase, polypeptide 2 | -1.58 |
| ENSMUSG00000020284 | 1810043G02Rik | RIKEN cDNA 1810043G02 gene | -1.58 |
| ENSMUSG00000052131 | Akr1b7 | aldo-keto reductase family 1, member B7 | -1.58 |
| ENSMUSG00000030834 | Abcc6 | ATP-binding cassette, sub-family C (CFTR/MRP), member 6 | -1.58 |
| ENSMUSG00000105762 | Gm43605 | predicted gene 43605 | -1.58 |
| ENSMUSG00000029428 | Stx2 | syntaxin 2 | -1.58 |
| ENSMUSG00000074461 | Gm10699 | predicted gene 10699 | -1.58 |
| ENSMUSG00000075520 | Malrd1 | MAM and LDL receptor class A domain containing 1 | -1.59 |
| ENSMUSG00000003134 | Tbc1d8 | TBC1 domain family, member 8 | -1.59 |
| ENSMUSG00000028860 | Sytl1 | synaptotagmin-like 1 | -1.59 |
| ENSMUSG00000042523 | Dnal1 | dynein, axonemal, light chain 1 | -1.59 |
| ENSMUSG00000030621 | Me3 | malic enzyme 3, NADP(+)-dependent, mitochondrial | -1.59 |
| ENSMUSG00000052605 | Isoc2b | isochorismatase domain containing 2b | -1.59 |
| ENSMUSG00000021364 | Elovl2 | elongation of very long chain fatty acids (FEN1/Elo2, SUR4/Elo3, yeast)-like 2 | -1.59 |
| ENSMUSG00000115098 | Gm49134 | predicted gene, 49134 | -1.59 |
| ENSMUSG00000000120 | Ngfr | nerve growth factor receptor (TNFR superfamily, member 16) | -1.59 |
| ENSMUSG00000105302 | Gm19817 | predicted gene, 19817 | -1.59 |
| ENSMUSG00000099893 | Gm29245 | predicted gene 29245 | -1.59 |
| ENSMUSG00000005800 | Mmp8 | matrix metallopeptidase 8 | -1.59 |
| ENSMUSG00000000202 | Btbd17 | BTB (POZ) domain containing 17 | -1.60 |
| ENSMUSG00000067203 | H2-K2 | histocompatibility 2, K region locus 2 | -1.60 |
| ENSMUSG00000102723 | Gm37936 | predicted gene, 37936 | -1.60 |
| ENSMUSG00000050600 | Zfp831 | zinc finger protein 831 | -1.60 |
| ENSMUSG00000030731 | Syt3 | synaptotagmin III | -1.60 |
| ENSMUSG00000074912 | Gm14207 | predicted gene 14207 | -1.61 |
| ENSMUSG00000103284 | 3110080O07Rik | RIKEN cDNA 3110080O07 gene | -1.61 |
| ENSMUSG00000017740 | Slc12a5 | solute carrier family 12, member 5 | -1.61 |
| ENSMUSG00000097768 | 2310043M15Rik | RIKEN cDNA 2310043M15 gene | -1.61 |
| ENSMUSG00000107182 | Gm43268 | predicted gene 43268 | -1.61 |
| ENSMUSG00000019429 | Ffar3 | free fatty acid receptor 3 | -1.61 |
| ENSMUSG00000110301 | Gm35363 | predicted gene, 35363 | -1.61 |
| ENSMUSG00000001604 | Tcea3 | transcription elongation factor A (SII), 3 | -1.62 |
| ENSMUSG00000018740 | Slc25a35 | solute carrier family 25, member 35 | -1.62 |
| ENSMUSG00000035930 | Chst4 | carbohydrate (chondroitin 6/keratan) sulfotransferase 4 | -1.62 |
| ENSMUSG00000036492 | Rnf39 | ring finger protein 39 | -1.62 |
| ENSMUSG00000048960 | Prex2 | phosphatidylinositol-3,4,5-trisphosphate-dependent Rac exchange factor 2 | -1.62 |
| ENSMUSG00000043313 | Pcdhb19 | protocadherin beta 19 | -1.62 |
| ENSMUSG00000018581 | Dnah11 | dynein, axonemal, heavy chain 11 | -1.62 |
| ENSMUSG00000103577 | 9330162B11Rik | RIKEN cDNA 9330162B11 gene | -1.62 |
| ENSMUSG00000027514 | Zbp1 | Z-DNA binding protein 1 | -1.63 |
| ENSMUSG00000022219 | Cideb | cell death-inducing DNA fragmentation factor, alpha subunit-like effector B | -1.63 |
| ENSMUSG00000095105 | Edaradd | EDAR (ectodysplasin-A receptor)-associated death domain | -1.63 |
| ENSMUSG00000025422 | Agap2 | ArfGAP with GTPase domain, ankyrin repeat and PH domain 2 | -1.63 |
| ENSMUSG00000030882 | Dnhd1 | dynein heavy chain domain 1 | -1.63 |

| | | | |
|---|---|---|---|
| ENSMUSG00000035849 | Krt222 | keratin 222 | -1.63 |
| ENSMUSG00000041696 | Rasl12 | RAS-like, family 12 | -1.63 |
| ENSMUSG00000068859 | Sp9 | trans-acting transcription factor 9 | -1.63 |
| ENSMUSG00000028845 | Tekt2 | tektin 2 | -1.63 |
| ENSMUSG00000064373 | Selenop | selenoprotein P | -1.64 |
| ENSMUSG00000022332 | Khdrbs3 | KH domain containing, RNA binding, signal transduction associated 3 | -1.64 |
| ENSMUSG00000036585 | Fgf1 | fibroblast growth factor 1 | -1.64 |
| ENSMUSG00000029700 | Slc13a1 | solute carrier family 13 (sodium/sulfate symporters), member 1 | -1.64 |
| ENSMUSG00000049971 | Glt1d1 | glycosyltransferase 1 domain containing 1 | -1.64 |
| ENSMUSG00000050295 | Foxc1 | forkhead box C1 | -1.64 |
| ENSMUSG00000001249 | Hpn | hepsin | -1.64 |
| ENSMUSG00000061578 | Ksr2 | kinase suppressor of ras 2 | -1.64 |
| ENSMUSG00000068196 | Col8a1 | collagen, type VIII, alpha 1 | -1.64 |
| ENSMUSG00000036040 | Adamtsl2 | ADAMTS-like 2 | -1.64 |
| ENSMUSG00000078942 | Naip6 | NLR family, apoptosis inhibitory protein 6 | -1.65 |
| ENSMUSG00000108427 | Gm36696 | predicted gene, 36696 | -1.65 |
| ENSMUSG00000037942 | Crp | C-reactive protein, pentraxin-related | -1.65 |
| ENSMUSG00000024193 | Phf1 | PHD finger protein 1 | -1.66 |
| ENSMUSG00000033705 | Stard9 | START domain containing 9 | -1.66 |
| ENSMUSG00000028795 | Ccdc28b | coiled coil domain containing 28B | -1.66 |
| ENSMUSG00000026728 | Vim | vimentin | -1.66 |
| ENSMUSG00000006241 | Ccdc159 | coiled-coil domain containing 159 | -1.66 |
| ENSMUSG00000092210 | A930009A15Rik | RIKEN cDNA A930009A15 gene | -1.66 |
| ENSMUSG00000037418 | Best1 | bestrophin 1 | -1.66 |
| ENSMUSG00000064140 | Trim38 | tripartite motif-containing 38 | -1.66 |
| ENSMUSG00000033684 | Qsox1 | quiescin Q6 sulfhydryl oxidase 1 | -1.67 |
| ENSMUSG00000039578 | Ccser1 | coiled-coil serine rich 1 | -1.67 |
| ENSMUSG00000048232 | Fbxo10 | F-box protein 10 | -1.67 |
| ENSMUSG00000035357 | Pdzrn3 | PDZ domain containing RING finger 3 | -1.67 |
| ENSMUSG00000007950 | Abhd8 | abhydrolase domain containing 8 | -1.67 |
| ENSMUSG00000018339 | Gpx3 | glutathione peroxidase 3 | -1.67 |
| ENSMUSG00000045912 | C2cd4c | C2 calcium-dependent domain containing 4C | -1.67 |
| ENSMUSG00000079114 | Gm7849 | predicted gene 7849 | -1.67 |
| ENSMUSG00000025479 | Cyp2e1 | cytochrome P450, family 2, subfamily e, polypeptide 1 | -1.68 |
| ENSMUSG00000085873 | Ttc39aos1 | Ttc39a opposite strand RNA 1 | -1.68 |
| ENSMUSG00000104966 | Gm43273 | predicted gene 43273 | -1.68 |
| ENSMUSG00000043614 | Vps37d | vacuolar protein sorting 37D | -1.68 |
| ENSMUSG00000032238 | Rora | RAR-related orphan receptor alpha | -1.68 |
| ENSMUSG00000039236 | Isg20 | interferon-stimulated protein | -1.68 |
| ENSMUSG00000085003 | Pip5k1bos | phosphatidylinositol-4-phosphate 5-kinase, type 1 beta, opposite strand | -1.68 |
| ENSMUSG00000024391 | Apom | apolipoprotein M | -1.68 |
| ENSMUSG00000032251 | Irak1bp1 | interleukin-1 receptor-associated kinase 1 binding protein 1 | -1.68 |
| ENSMUSG00000039168 | Dap | death-associated protein | -1.69 |
| ENSMUSG00000038167 | Plekhg6 | pleckstrin homology domain containing, family G (with RhoGef domain) member 6 | -1.69 |
| ENSMUSG00000068551 | Zfp467 | zinc finger protein 467 | -1.69 |
| ENSMUSG00000051435 | Fhad1 | forkhead-associated (FHA) phosphopeptide binding domain 1 | -1.69 |
| ENSMUSG00000052921 | Arhgef15 | Rho guanine nucleotide exchange factor (GEF) 15 | -1.69 |
| ENSMUSG00000040061 | Plcb2 | phospholipase C, beta 2 | -1.69 |
| ENSMUSG00000086970 | Bcas1os1 | breast carcinoma amplified sequence 1, opposite strand 1 | -1.69 |
| ENSMUSG00000075015 | Gm10801 | predicted gene 10801 | -1.69 |
| ENSMUSG00000023781 | Hes7 | hes family bHLH transcription factor 7 | -1.69 |
| ENSMUSG00000081059 | Gm11945 | predicted gene 11945 | -1.69 |
| ENSMUSG00000016942 | Tmprss6 | transmembrane serine protease 6 | -1.69 |
| ENSMUSG00000091043 | Glyatl3 | glycine-N-acyltransferase-like 3 | -1.69 |
| ENSMUSG00000036377 | C530008M17Rik | RIKEN cDNA C530008M17 gene | -1.70 |
| ENSMUSG00000078486 | Perm1 | PPARGC1 and ESRR induced regulator, muscle 1 | -1.70 |
| ENSMUSG00000039976 | Tbc1d16 | TBC1 domain family, member 16 | -1.70 |
| ENSMUSG00000097924 | A730020E08Rik | RIKEN cDNA A730020E08 gene | -1.70 |
| ENSMUSG00000022199 | Slc22a17 | solute carrier family 22 (organic cation transporter), member 17 | -1.70 |
| ENSMUSG00000092323 | BB365896 | expressed sequence BB365896 | -1.70 |
| ENSMUSG00000084433 | Gm25945 | predicted gene, 25945 | -1.70 |
| ENSMUSG00000055003 | Lrtm2 | leucine-rich repeats and transmembrane domains 2 | -1.70 |
| ENSMUSG00000041261 | Car8 | carbonic anhydrase 8 | -1.71 |
| ENSMUSG00000010830 | Kdelr3 | KDEL (Lys-Asp-Glu-Leu) endoplasmic reticulum protein retention receptor 3 | -1.71 |
| ENSMUSG00000027612 | Mmp24 | matrix metallopeptidase 24 | -1.71 |
| ENSMUSG00000022440 | C1qtnf6 | C1q and tumor necrosis factor related protein 6 | -1.71 |
| ENSMUSG00000047473 | Zfp30 | zinc finger protein 30 | -1.71 |

| | | | |
|---|---|---|---|
| ENSMUSG00000073418 | C4b | complement component 4B (Chido blood group) | -1.71 |
| ENSMUSG00000079471 | Mymx | myomixer, myoblast fusion factor | -1.71 |
| ENSMUSG00000032226 | Gcnt3 | glucosaminyl (N-acetyl) transferase 3, mucin type | -1.72 |
| ENSMUSG00000027457 | Snph | syntaphilin | -1.72 |
| ENSMUSG00000038497 | Tmco3 | transmembrane and coiled-coil domains 3 | -1.73 |
| ENSMUSG00000037025 | Foxa2 | forkhead box A2 | -1.73 |
| ENSMUSG00000030861 | Acadsb | acyl-Coenzyme A dehydrogenase, short/branched chain | -1.73 |
| ENSMUSG00000054252 | Fgfr3 | fibroblast growth factor receptor 3 | -1.73 |
| ENSMUSG00000003355 | Fkbp11 | FK506 binding protein 11 | -1.73 |
| ENSMUSG00000032661 | Oas3 | 2'-5' oligoadenylate synthetase 3 | -1.73 |
| ENSMUSG00000033460 | Armcx1 | armadillo repeat containing, X-linked 1 | -1.73 |
| ENSMUSG00000039114 | Nrn1 | neuritin 1 | -1.73 |
| ENSMUSG00000030630 | Fah | fumarylacetoacetate hydrolase | -1.73 |
| ENSMUSG00000055497 | Gm9974 | predicted gene 9974 | -1.73 |
| ENSMUSG00000085589 | A430078I02Rik | RIKEN cDNA A430078I02 gene | -1.73 |
| ENSMUSG00000032353 | Tmed3 | transmembrane p24 trafficking protein 3 | -1.74 |
| ENSMUSG00000037709 | Fam13a | family with sequence similarity 13, member A | -1.74 |
| ENSMUSG00000033717 | Adra2a | adrenergic receptor, alpha 2a | -1.74 |
| ENSMUSG00000030483 | Cyp2b10 | cytochrome P450, family 2, subfamily b, polypeptide 10 | -1.74 |
| ENSMUSG00000117286 | Gm1043 | predicted 1043 | -1.74 |
| ENSMUSG00000047878 | A4galt | alpha 1,4-galactosyltransferase | -1.74 |
| ENSMUSG00000079174 | Gm3054 | predicted gene 3054 | -1.75 |
| ENSMUSG00000028838 | Extl1 | exostoses (multiple)-like 1 | -1.75 |
| ENSMUSG00000085852 | Gm13807 | predicted gene 13807 | -1.75 |
| ENSMUSG00000107076 | Gm43480 | predicted gene 43480 | -1.75 |
| ENSMUSG00000003410 | Elavl3 | ELAV like RNA binding protein 3 | -1.75 |
| ENSMUSG00000085941 | Gm11201 | predicted gene 11201 | -1.75 |
| ENSMUSG00000109093 | Gm19950 | predicted gene, 19950 | -1.75 |
| ENSMUSG00000104362 | Gm37928 | predicted gene, 37928 | -1.75 |
| ENSMUSG00000039943 | Plcb4 | phospholipase C, beta 4 | -1.76 |
| ENSMUSG00000003378 | Grik5 | glutamate receptor, ionotropic, kainate 5 (gamma 2) | -1.76 |
| ENSMUSG00000110397 | Gm45540 | predicted gene 45540 | -1.76 |
| ENSMUSG00000004933 | Matk | megakaryocyte-associated tyrosine kinase | -1.76 |
| ENSMUSG00000081270 | Gm11653 | predicted gene 11653 | -1.76 |
| ENSMUSG00000024530 | Prelid3a | PRELI domain containing 3A | -1.76 |
| ENSMUSG00000045875 | Adra1a | adrenergic receptor, alpha 1a | -1.76 |
| ENSMUSG00000031451 | Gas6 | growth arrest specific 6 | -1.77 |
| ENSMUSG00000049307 | Fut4 | fucosyltransferase 4 | -1.77 |
| ENSMUSG00000024063 | Lbh | limb-bud and heart | -1.77 |
| ENSMUSG00000035246 | Pcyt1b | phosphate cytidylyltransferase 1, choline, beta isoform | -1.77 |
| ENSMUSG00000024778 | Fas | Fas (TNF receptor superfamily member 6) | -1.77 |
| ENSMUSG00000033287 | Kctd17 | potassium channel tetramerisation domain containing 17 | -1.77 |
| ENSMUSG00000094286 | Gm12424 | predicted gene 12424 | -1.77 |
| ENSMUSG00000110666 | Gm9172 | predicted gene 9172 | -1.77 |
| ENSMUSG00000023328 | Ache | acetylcholinesterase | -1.78 |
| ENSMUSG00000073067 | 9130019P16Rik | RIKEN cDNA 9130019P16 gene | -1.78 |
| ENSMUSG00000032740 | Ccdc88a | coiled coil domain containing 88A | -1.78 |
| ENSMUSG00000087670 | 9530036M11Rik | RIKEN cDNA 9530036M11 gene | -1.78 |
| ENSMUSG00000038777 | Sema6c | sema domain, transmembrane domain (TM), and cytoplasmic domain, (semaphorin | -1.78 |
| ENSMUSG00000033209 | Ttc28 | tetratricopeptide repeat domain 28 | -1.78 |
| ENSMUSG00000012017 | Scarf2 | scavenger receptor class F, member 2 | -1.78 |
| ENSMUSG00000074052 | BC048644 | cDNA sequence BC048644 | -1.78 |
| ENSMUSG00000028758 | Kif17 | kinesin family member 17 | -1.78 |
| ENSMUSG00000069456 | Rdh16 | retinol dehydrogenase 16 | -1.79 |
| ENSMUSG00000032053 | Pou2af1 | POU domain, class 2, associating factor 1 | -1.79 |
| ENSMUSG00000028186 | Uox | urate oxidase | -1.79 |
| ENSMUSG00000021414 | Fam217a | family with sequence similarity 217, member A | -1.79 |
| ENSMUSG00000021057 | Akap5 | A kinase (PRKA) anchor protein 5 | -1.80 |
| ENSMUSG00000027075 | Slc43a1 | solute carrier family 43, member 1 | -1.80 |
| ENSMUSG00000112307 | Gm48751 | predicted gene, 48751 | -1.80 |
| ENSMUSG00000056856 | Jakmip3 | janus kinase and microtubule interacting protein 3 | -1.80 |
| ENSMUSG00000056648 | Hoxb8 | homeobox B8 | -1.80 |
| ENSMUSG00000092384 | Gm4189 | predicted gene 4189 | -1.80 |
| ENSMUSG00000028524 | Sgip1 | SH3-domain GRB2-like (endophilin) interacting protein 1 | -1.80 |
| ENSMUSG00000027435 | Cd93 | CD93 antigen | -1.80 |
| ENSMUSG00000036390 | Gadd45a | growth arrest and DNA-damage-inducible 45 alpha | -1.81 |
| ENSMUSG00000011832 | Evi5l | ecotropic viral integration site 5 like | -1.81 |

| | | | |
|---|---|---|---|
| ENSMUSG00000015981 | Stk32c | serine/threonine kinase 32C | -1.81 |
| ENSMUSG00000039485 | Tspyl4 | TSPY-like 4 | -1.81 |
| ENSMUSG00000055370 | Gm9968 | predicted gene 9968 | -1.81 |
| ENSMUSG00000040387 | Klhl32 | kelch-like 32 | -1.81 |
| ENSMUSG00000031133 | Arhgef6 | Rac/Cdc42 guanine nucleotide exchange factor (GEF) 6 | -1.81 |
| ENSMUSG00000111323 | Gm48129 | predicted gene, 48129 | -1.81 |
| ENSMUSG00000099826 | Scgb2b10 | secretoglobin, family 2B, member 10 | -1.81 |
| ENSMUSG00000040987 | Mill2 | MHC I like leukocyte 2 | -1.82 |
| ENSMUSG00000110540 | Gm45743 | predicted gene 45743 | -1.82 |
| ENSMUSG00000004630 | Pcp2 | Purkinje cell protein 2 (L7) | -1.82 |
| ENSMUSG00000027314 | Dll4 | delta like canonical Notch ligand 4 | -1.83 |
| ENSMUSG00000051335 | Gfod1 | glucose-fructose oxidoreductase domain containing 1 | -1.83 |
| ENSMUSG00000006711 | D130043K22Rik | RIKEN cDNA D130043K22 gene | -1.83 |
| ENSMUSG00000020034 | Tcp11l2 | t-complex 11 (mouse) like 2 | -1.83 |
| ENSMUSG00000025221 | Kcnip2 | Kv channel-interacting protein 2 | -1.83 |
| ENSMUSG00000078091 | Gm10912 | predicted gene 10912 | -1.83 |
| ENSMUSG00000091189 | Ear-ps3 | eosinophil-associated, ribonuclease A family, pseudogene 3 | -1.83 |
| ENSMUSG00000024109 | Nrxn1 | neurexin I | -1.83 |
| ENSMUSG00000071203 | Naip5 | NLR family, apoptosis inhibitory protein 5 | -1.84 |
| ENSMUSG00000028634 | Hivep3 | human immunodeficiency virus type I enhancer binding protein 3 | -1.84 |
| ENSMUSG00000040350 | Trim7 | tripartite motif-containing 7 | -1.84 |
| ENSMUSG00000049804 | Armcx4 | armadillo repeat containing, X-linked 4 | -1.84 |
| ENSMUSG00000025104 | Hdgfl3 | HDGF like 3 | -1.84 |
| ENSMUSG00000087049 | Bach2it1 | BTB and CNC homology 2, intronic transcript 1 | -1.84 |
| ENSMUSG00000063522 | 2010109I03Rik | RIKEN cDNA 2010109I03 gene | -1.85 |
| ENSMUSG00000025352 | Gdf11 | growth differentiation factor 11 | -1.85 |
| ENSMUSG00000056185 | Snx32 | sorting nexin 32 | -1.85 |
| ENSMUSG00000037902 | Sirpa | signal-regulatory protein alpha | -1.85 |
| ENSMUSG00000053675 | Tgm5 | transglutaminase 5 | -1.85 |
| ENSMUSG00000032101 | Ddx25 | DEAD (Asp-Glu-Ala-Asp) box polypeptide 25 | -1.85 |
| ENSMUSG00000029563 | Foxp2 | forkhead box P2 | -1.85 |
| ENSMUSG00000034112 | Atp2c2 | ATPase, Ca++ transporting, type 2C, member 2 | -1.86 |
| ENSMUSG00000036422 | Pcdh8 | protocadherin 8 | -1.86 |
| ENSMUSG00000040896 | Kcnd3 | potassium voltage-gated channel, Shal-related family, member 3 | -1.86 |
| ENSMUSG00000042388 | Dlgap3 | DLG associated protein 3 | -1.86 |
| ENSMUSG00000047904 | Sstr2 | somatostatin receptor 2 | -1.86 |
| ENSMUSG00000044250 | Pced1b | PC-esterase domain containing 1B | -1.86 |
| ENSMUSG00000038608 | Dock10 | dedicator of cytokinesis 10 | -1.86 |
| ENSMUSG00000110131 | Gm18066 | predicted gene, 18066 | -1.86 |
| ENSMUSG00000022012 | Enox1 | ecto-NOX disulfide-thiol exchanger 1 | -1.86 |
| ENSMUSG00000082443 | Gm12604 | predicted gene 12604 | -1.86 |
| ENSMUSG00000033715 | Akr1c14 | aldo-keto reductase family 1, member C14 | -1.87 |
| ENSMUSG00000039115 | Itga9 | integrin alpha 9 | -1.87 |
| ENSMUSG00000029769 | Ccdc136 | coiled-coil domain containing 136 | -1.87 |
| ENSMUSG00000041710 | Trpc5 | transient receptor potential cation channel, subfamily C, member 5 | -1.87 |
| ENSMUSG00000087623 | Gm12404 | predicted gene 12404 | -1.87 |
| ENSMUSG00000030207 | Fam234b | family with sequence similarity 234, member B | -1.88 |
| ENSMUSG00000084939 | Gm830 | predicted gene 830 | -1.88 |
| ENSMUSG00000029919 | Hpgds | hematopoietic prostaglandin D synthase | -1.88 |
| ENSMUSG00000054434 | Tmem120b | transmembrane protein 120B | -1.88 |
| ENSMUSG00000072720 | Myo18b | myosin XVIIIb | -1.88 |
| ENSMUSG00000014158 | Trpv4 | transient receptor potential cation channel, subfamily V, member 4 | -1.88 |
| ENSMUSG00000031561 | Tenm3 | teneurin transmembrane protein 3 | -1.88 |
| ENSMUSG00000117328 | AC169675.1 | novel transcript, antisense to Smoc2 | -1.88 |
| ENSMUSG00000027188 | Pamr1 | peptidase domain containing associated with muscle regeneration 1 | -1.88 |
| ENSMUSG00000065145 | Vaultrc5 | vault RNA component 5 | -1.89 |
| ENSMUSG00000038893 | Fam117a | family with sequence similarity 117, member A | -1.89 |
| ENSMUSG00000023927 | Satb1 | special AT-rich sequence binding protein 1 | -1.89 |
| ENSMUSG00000032978 | Guca2b | guanylate cyclase activator 2b (retina) | -1.90 |
| ENSMUSG00000039652 | Cpeb3 | cytoplasmic polyadenylation element binding protein 3 | -1.90 |
| ENSMUSG00000025656 | Arhgef9 | CDC42 guanine nucleotide exchange factor (GEF) 9 | -1.90 |
| ENSMUSG00000026959 | Grin1 | glutamate receptor, ionotropic, NMDA1 (zeta 1) | -1.90 |
| ENSMUSG00000027217 | Tspan18 | tetraspanin 18 | -1.90 |
| ENSMUSG00000050132 | Sarm1 | sterile alpha and HEAT/Armadillo motif containing 1 | -1.90 |
| ENSMUSG00000089712 | Gm15889 | predicted gene 15889 | -1.91 |
| ENSMUSG00000035067 | Xkr6 | X-linked Kx blood group related 6 | -1.92 |
| ENSMUSG00000073424 | Cyp4f15 | cytochrome P450, family 4, subfamily f, polypeptide 15 | -1.92 |

| | | | |
|---|---|---|---|
| ENSMUSG00000041789 | 2700046A07Rik | RIKEN cDNA 2700046A07 gene | -1.92 |
| ENSMUSG00000048078 | Tenm4 | teneurin transmembrane protein 4 | -1.93 |
| ENSMUSG00000057069 | Ero1lb | ERO1-like beta (S. cerevisiae) | -1.93 |
| ENSMUSG00000026360 | Rgs2 | regulator of G-protein signaling 2 | -1.93 |
| ENSMUSG00000103579 | Gm37113 | predicted gene, 37113 | -1.93 |
| ENSMUSG00000027339 | Rassf2 | Ras association (RalGDS/AF-6) domain family member 2 | -1.93 |
| ENSMUSG00000029797 | Sspo | SCO-spondin | -1.93 |
| ENSMUSG00000047161 | Chst9 | carbohydrate (N-acetylgalactosamine 4-0) sulfotransferase 9 | -1.93 |
| ENSMUSG00000024039 | Cbs | cystathionine beta-synthase | -1.94 |
| ENSMUSG00000062859 | Tcp11 | t-complex protein 11 | -1.94 |
| ENSMUSG00000030228 | Pik3c2g | phosphatidylinositol-4-phosphate 3-kinase catalytic subunit type 2 gamma | -1.94 |
| ENSMUSG00000031202 | Rab39b | RAB39B, member RAS oncogene family | -1.94 |
| ENSMUSG00000020423 | Btg2 | B cell translocation gene 2, anti-proliferative | -1.95 |
| ENSMUSG00000041132 | N4bp2l1 | NEDD4 binding protein 2-like 1 | -1.95 |
| ENSMUSG00000018623 | Mmp7 | matrix metallopeptidase 7 | -1.95 |
| ENSMUSG00000054150 | Syne3 | spectrin repeat containing, nuclear envelope family member 3 | -1.95 |
| ENSMUSG00000009628 | Tex15 | testis expressed gene 15 | -1.95 |
| ENSMUSG00000016356 | Col20a1 | collagen, type XX, alpha 1 | -1.95 |
| ENSMUSG00000103220 | Gm37728 | predicted gene, 37728 | -1.95 |
| ENSMUSG00000039419 | Cntnap2 | contactin associated protein-like 2 | -1.95 |
| ENSMUSG00000067642 | Adgrf3 | adhesion G protein-coupled receptor F3 | -1.95 |
| ENSMUSG00000055737 | Ghr | growth hormone receptor | -1.96 |
| ENSMUSG00000018427 | Ypel2 | yippee like 2 | -1.96 |
| ENSMUSG00000034591 | Slc41a2 | solute carrier family 41, member 2 | -1.96 |
| ENSMUSG00000024124 | Prss30 | protease, serine 30 | -1.96 |
| ENSMUSG00000006464 | Bbs1 | Bardet-Biedl syndrome 1 (human) | -1.96 |
| ENSMUSG00000041912 | Tdrkh | tudor and KH domain containing protein | -1.96 |
| ENSMUSG00000043020 | Wdr63 | WD repeat domain 63 | -1.96 |
| ENSMUSG00000115360 | Gm48932 | predicted gene, 48932 | -1.96 |
| ENSMUSG00000032609 | Klhdc8b | kelch domain containing 8B | -1.97 |
| ENSMUSG00000021675 | F2rl2 | coagulation factor II (thrombin) receptor-like 2 | -1.97 |
| ENSMUSG00000074782 | 4833422C13Rik | RIKEN cDNA 4833422C13 gene | -1.97 |
| ENSMUSG00000025978 | Rftn2 | raftlin family member 2 | -1.97 |
| ENSMUSG00000036887 | C1qa | complement component 1, q subcomponent, alpha polypeptide | -1.97 |
| ENSMUSG00000040867 | Begain | brain-enriched guanylate kinase-associated | -1.97 |
| ENSMUSG00000042269 | Fam92b | family with sequence similarity 92, member B | -1.97 |
| ENSMUSG00000072115 | Ang | angiogenin, ribonuclease, RNase A family, 5 | -1.98 |
| ENSMUSG00000060935 | Tmem263 | transmembrane protein 263 | -1.98 |
| ENSMUSG00000006930 | Hap1 | huntingtin-associated protein 1 | -1.98 |
| ENSMUSG00000061808 | Ttr | transthyretin | -1.98 |
| ENSMUSG00000111282 | Gm47528 | predicted gene, 47528 | -1.98 |
| ENSMUSG00000117050 | AC167363.2 | TEC | -1.98 |
| ENSMUSG00000020486 | 38231 | septin 4 | -1.98 |
| ENSMUSG00000057400 | Ces1c | carboxylesterase 1C | -1.98 |
| ENSMUSG00000066516 | Klk1b21 | kallikrein 1-related peptidase b21 | -1.98 |
| ENSMUSG00000025427 | Rnf165 | ring finger protein 165 | -1.98 |
| ENSMUSG00000052387 | Trpm3 | transient receptor potential cation channel, subfamily M, member 3 | -1.98 |
| ENSMUSG00000051910 | Sox6 | SRY (sex determining region Y)-box 6 | -2.00 |
| ENSMUSG00000049191 | Rtl5 | retrotransposon Gag like 5 | -2.00 |
| ENSMUSG00000031791 | Tmem38a | transmembrane protein 38A | -2.00 |
| ENSMUSG00000034949 | Zfr2 | zinc finger RNA binding protein 2 | -2.00 |
| ENSMUSG00000039629 | Strip2 | striatin interacting protein 2 | -2.00 |
| ENSMUSG00000023966 | Rsph9 | radial spoke head 9 homolog (Chlamydomonas) | -2.00 |
| ENSMUSG00000085664 | Atxn7l1os2 | ataxin 7-like 1, opposite strand 2 | -2.00 |
| ENSMUSG00000036196 | Slc26a8 | solute carrier family 26, member 8 | -2.00 |
| ENSMUSG00000036892 | Prodh2 | proline dehydrogenase (oxidase) 2 | -2.00 |
| ENSMUSG00000061576 | Dpp6 | dipeptidylpeptidase 6 | -2.01 |
| ENSMUSG00000090812 | Samd15 | sterile alpha motif domain containing 15 | -2.01 |
| ENSMUSG00000106749 | Gm43281 | predicted gene 43281 | -2.01 |
| ENSMUSG00000088891 | Gm22516 | predicted gene, 22516 | -2.01 |
| ENSMUSG00000058897 | Col25a1 | collagen, type XXV, alpha 1 | -2.01 |
| ENSMUSG00000105495 | Gm42995 | predicted gene 42995 | -2.01 |
| ENSMUSG00000029166 | Mapre3 | microtubule-associated protein, RP/EB family, member 3 | -2.02 |
| ENSMUSG00000032548 | Slco2a1 | solute carrier organic anion transporter family, member 2a1 | -2.02 |
| ENSMUSG00000040415 | Dtx3 | deltex 3, E3 ubiquitin ligase | -2.02 |
| ENSMUSG00000035211 | Xrra1 | X-ray radiation resistance associated 1 | -2.02 |
| ENSMUSG00000031303 | Map3k15 | mitogen-activated protein kinase kinase kinase 15 | -2.03 |

| | | | |
|---|---|---|---|
| ENSMUSG00000047146 | Tet1 | tet methylcytosine dioxygenase 1 | -2.03 |
| ENSMUSG00000028076 | Cd1d1 | CD1d1 antigen | -2.03 |
| ENSMUSG00000112601 | Gm47590 | predicted gene, 47590 | -2.03 |
| ENSMUSG00000023906 | Cldn6 | claudin 6 | -2.04 |
| ENSMUSG00000046378 | Asphd1 | aspartate beta-hydroxylase domain containing 1 | -2.04 |
| ENSMUSG00000032356 | Rasgrf1 | RAS protein-specific guanine nucleotide-releasing factor 1 | -2.04 |
| ENSMUSG00000106350 | Gm4869 | predicted gene 4869 | -2.04 |
| ENSMUSG00000045658 | Pid1 | phosphotyrosine interaction domain containing 1 | -2.04 |
| ENSMUSG00000056679 | Gpr173 | G-protein coupled receptor 173 | -2.04 |
| ENSMUSG00000038271 | Iffo1 | intermediate filament family orphan 1 | -2.05 |
| ENSMUSG00000094840 | Muc3a | mucin 3A, cell surface associated | -2.06 |
| ENSMUSG00000040852 | Plekhh2 | pleckstrin homology domain containing, family H (with MyTH4 domain) member 2 | -2.06 |
| ENSMUSG00000059898 | Dsc3 | desmocollin 3 | -2.06 |
| ENSMUSG00000104776 | Gm43691 | predicted gene 43691 | -2.06 |
| ENSMUSG00000073102 | Drc1 | dynein regulatory complex subunit 1 | -2.06 |
| ENSMUSG00000027297 | Ltk | leukocyte tyrosine kinase | -2.07 |
| ENSMUSG00000053141 | Ptprt | protein tyrosine phosphatase, receptor type, T | -2.07 |
| ENSMUSG00000040495 | Chrm4 | cholinergic receptor, muscarinic 4 | -2.07 |
| ENSMUSG00000042109 | Csdc2 | cold shock domain containing C2, RNA binding | -2.07 |
| ENSMUSG00000020140 | Lgr5 | leucine rich repeat containing G protein coupled receptor 5 | -2.08 |
| ENSMUSG00000027223 | Mapk8ip1 | mitogen-activated protein kinase 8 interacting protein 1 | -2.08 |
| ENSMUSG00000051166 | Eml5 | echinoderm microtubule associated protein like 5 | -2.08 |
| ENSMUSG00000054893 | Zfp667 | zinc finger protein 667 | -2.08 |
| ENSMUSG00000044726 | Erich5 | glutamate rich 5 | -2.08 |
| ENSMUSG00000097328 | Tnfsf12 | tumor necrosis factor (ligand) superfamily, member 12 | -2.08 |
| ENSMUSG00000107451 | Gm44421 | predicted gene, 44421 | -2.08 |
| ENSMUSG00000045498 | Pcdhb3 | protocadherin beta 3 | -2.08 |
| ENSMUSG00000095066 | Defa20 | defensin, alpha, 20 | -2.08 |
| ENSMUSG00000031824 | 6430548M08Rik | RIKEN cDNA 6430548M08 gene | -2.09 |
| ENSMUSG00000085363 | Gm15478 | predicted gene 15478 | -2.09 |
| ENSMUSG00000026841 | Fibcd1 | fibrinogen C domain containing 1 | -2.09 |
| ENSMUSG00000029359 | Tesc | tescalcin | -2.10 |
| ENSMUSG00000038793 | Lefty1 | left right determination factor 1 | -2.10 |
| ENSMUSG00000030406 | Gipr | gastric inhibitory polypeptide receptor | -2.10 |
| ENSMUSG00000020953 | Coch | cochlin | -2.10 |
| ENSMUSG00000105720 | Gm42440 | predicted gene 42440 | -2.10 |
| ENSMUSG00000050963 | Kcns2 | K+ voltage-gated channel, subfamily S, 2 | -2.10 |
| ENSMUSG00000102112 | 1810041H14Rik | RIKEN cDNA 1810041H14 gene | -2.11 |
| ENSMUSG00000003411 | Rab3b | RAB3B, member RAS oncogene family | -2.11 |
| ENSMUSG00000020930 | Ccdc103 | coiled-coil domain containing 103 | -2.11 |
| ENSMUSG00000038175 | Mylip | myosin regulatory light chain interacting protein | -2.12 |
| ENSMUSG00000112035 | Gm49335 | predicted gene, 49335 | -2.12 |
| ENSMUSG00000027931 | Npr1 | natriuretic peptide receptor 1 | -2.13 |
| ENSMUSG00000040367 | Lrrd1 | leucine rich repeats and death domain containing 1 | -2.13 |
| ENSMUSG00000104936 | 2610011E03Rik | RIKEN cDNA 2610011E03 gene | -2.13 |
| ENSMUSG00000103331 | Gm37995 | predicted gene, 37995 | -2.14 |
| ENSMUSG00000017639 | Rab11fip4 | RAB11 family interacting protein 4 (class II) | -2.15 |
| ENSMUSG00000034116 | Vav1 | vav 1 oncogene | -2.15 |
| ENSMUSG00000110887 | Cbx3-ps8 | chromobox 3, pseudogene 8 | -2.15 |
| ENSMUSG00000063160 | Numbl | numb-like | -2.15 |
| ENSMUSG00000031919 | Tmed6 | transmembrane p24 trafficking protein 6 | -2.16 |
| ENSMUSG00000049892 | Rasd1 | RAS, dexamethasone-induced 1 | -2.16 |
| ENSMUSG00000111086 | Gm48671 | predicted gene, 48671 | -2.16 |
| ENSMUSG00000031523 | Dlc1 | deleted in liver cancer 1 | -2.16 |
| ENSMUSG00000034115 | Scn11a | sodium channel, voltage-gated, type XI, alpha | -2.16 |
| ENSMUSG00000027359 | Slc27a2 | solute carrier family 27 (fatty acid transporter), member 2 | -2.17 |
| ENSMUSG00000110622 | Iqcn | IQ motif containing N | -2.17 |
| ENSMUSG00000006958 | Chrd | chordin | -2.17 |
| ENSMUSG00000108961 | Gm32540 | predicted gene, 32540 | -2.17 |
| ENSMUSG00000038807 | Rap1gap2 | RAP1 GTPase activating protein 2 | -2.18 |
| ENSMUSG00000062785 | Kcnc3 | potassium voltage gated channel, Shaw-related subfamily, member 3 | -2.18 |
| ENSMUSG00000004187 | Kifc2 | kinesin family member C2 | -2.18 |
| ENSMUSG00000021379 | Id4 | inhibitor of DNA binding 4 | -2.18 |
| ENSMUSG00000105289 | Gm43483 | predicted gene 43483 | -2.18 |
| ENSMUSG00000020182 | Ddc | dopa decarboxylase | -2.19 |
| ENSMUSG00000072294 | Klf12 | Kruppel-like factor 12 | -2.19 |
| ENSMUSG00000055214 | Pld5 | phospholipase D family, member 5 | -2.20 |

| | | | |
|---|---|---|---|
| ENSMUSG00000029544 | Cabp1 | calcium binding protein 1 | -2.20 |
| ENSMUSG00000036027 | 1810046K07Rik | RIKEN cDNA 1810046K07 gene | -2.20 |
| ENSMUSG00000078964 | Ces1b | carboxylesterase 1B | -2.20 |
| ENSMUSG00000105485 | Gm42447 | predicted gene 42447 | -2.20 |
| ENSMUSG00000047787 | Flrt1 | fibronectin leucine rich transmembrane protein 1 | -2.22 |
| ENSMUSG00000017652 | Cd40 | CD40 antigen | -2.22 |
| ENSMUSG00000023828 | Slc22a3 | solute carrier family 22 (organic cation transporter), member 3 | -2.22 |
| ENSMUSG00000073293 | Nudt10 | nudix (nucleoside diphosphate linked moiety X)-type motif 10 | -2.22 |
| ENSMUSG00000040310 | Alx4 | aristaless-like homeobox 4 | -2.23 |
| ENSMUSG00000089371 | Mir1938 | microRNA 1938 | -2.24 |
| ENSMUSG00000021453 | Gadd45g | growth arrest and DNA-damage-inducible 45 gamma | -2.25 |
| ENSMUSG00000071489 | Ptgdr | prostaglandin D receptor | -2.25 |
| ENSMUSG00000094305 | Scgb2b20 | secretoglobin, family 2B, member 20 | -2.25 |
| ENSMUSG00000027938 | Creb3l4 | cAMP responsive element binding protein 3-like 4 | -2.26 |
| ENSMUSG00000034675 | Dbn1 | drebrin 1 | -2.26 |
| ENSMUSG00000020227 | Irak3 | interleukin-1 receptor-associated kinase 3 | -2.26 |
| ENSMUSG00000037738 | Nek5 | NIMA (never in mitosis gene a)-related expressed kinase 5 | -2.26 |
| ENSMUSG00000037217 | Syn1 | synapsin I | -2.26 |
| ENSMUSG00000097294 | Gm26888 | predicted gene, 26888 | -2.26 |
| ENSMUSG00000046668 | Cxxc5 | CXXC finger 5 | -2.27 |
| ENSMUSG00000050100 | Hmx2 | H6 homeobox 2 | -2.27 |
| ENSMUSG00000032549 | Rab6b | RAB6B, member RAS oncogene family | -2.28 |
| ENSMUSG00000052726 | Kcnt2 | potassium channel, subfamily T, member 2 | -2.28 |
| ENSMUSG00000049811 | Fam161a | family with sequence similarity 161, member A | -2.28 |
| ENSMUSG00000108348 | Gm42372 | predicted gene, 42372 | -2.28 |
| ENSMUSG00000070385 | Ampd1 | adenosine monophosphate deaminase 1 | -2.28 |
| ENSMUSG00000112805 | Gm34574 | predicted gene, 34574 | -2.28 |
| ENSMUSG00000024232 | Bambi | BMP and activin membrane-bound inhibitor | -2.29 |
| ENSMUSG00000041000 | Trim62 | tripartite motif-containing 62 | -2.29 |
| ENSMUSG00000048330 | Ric3 | RIC3 acetylcholine receptor chaperone | -2.29 |
| ENSMUSG00000047181 | Samd14 | sterile alpha motif domain containing 14 | -2.29 |
| ENSMUSG00000035863 | Palm | paralemmin | -2.29 |
| ENSMUSG00000028327 | Stra6l | STRA6-like | -2.30 |
| ENSMUSG00000057335 | Cep170 | centrosomal protein 170 | -2.30 |
| ENSMUSG00000000530 | Acvrl1 | activin A receptor, type II-like 1 | -2.30 |
| ENSMUSG00000097194 | 9330175E14Rik | RIKEN cDNA 9330175E14 gene | -2.30 |
| ENSMUSG00000052861 | Dnah6 | dynein, axonemal, heavy chain 6 | -2.30 |
| ENSMUSG00000025576 | Rbfox3 | RNA binding protein, fox-1 homolog (C. elegans) 3 | -2.31 |
| ENSMUSG00000037492 | Zmat4 | zinc finger, matrin type 4 | -2.31 |
| ENSMUSG00000029490 | Mfsd7a | major facilitator superfamily domain containing 7A | -2.32 |
| ENSMUSG00000000305 | Cdh4 | cadherin 4 | -2.32 |
| ENSMUSG00000089678 | Agxt2 | alanine-glyoxylate aminotransferase 2 | -2.32 |
| ENSMUSG00000066687 | Zbtb16 | zinc finger and BTB domain containing 16 | -2.33 |
| ENSMUSG00000052942 | Glis3 | GLIS family zinc finger 3 | -2.33 |
| ENSMUSG00000025787 | Tgm4 | transglutaminase 4 (prostate) | -2.33 |
| ENSMUSG00000079559 | Colca2 | COLCA2 homolog | -2.33 |
| ENSMUSG00000034758 | Tle6 | transducin-like enhancer of split 6 | -2.33 |
| ENSMUSG00000020651 | Slc26a4 | solute carrier family 26, member 4 | -2.33 |
| ENSMUSG00000046719 | Nxph3 | neurexophilin 3 | -2.33 |
| ENSMUSG00000060508 | Nlrp9b | NLR family, pyrin domain containing 9B | -2.34 |
| ENSMUSG00000025969 | Nrp2 | neuropilin 2 | -2.34 |
| ENSMUSG00000024500 | Ppp2r2b | protein phosphatase 2, regulatory subunit B, beta | -2.34 |
| ENSMUSG00000008193 | Spib | Spi-B transcription factor (Spi-1/PU.1 related) | -2.34 |
| ENSMUSG00000102620 | Gm37675 | predicted gene, 37675 | -2.34 |
| ENSMUSG00000022438 | Parvb | parvin, beta | -2.34 |
| ENSMUSG00000106825 | 2510016D11Rik | RIKEN cDNA 2510016D11 gene | -2.35 |
| ENSMUSG00000028909 | Ptpru | protein tyrosine phosphatase, receptor type, U | -2.35 |
| ENSMUSG00000083718 | Ccnb2-ps | cyclin B2, pseudogene | -2.35 |
| ENSMUSG00000008461 | Fut1 | fucosyltransferase 1 | -2.35 |
| ENSMUSG00000044022 | Pcdhb21 | protocadherin beta 21 | -2.35 |
| ENSMUSG00000042249 | Grk3 | G protein-coupled receptor kinase 3 | -2.36 |
| ENSMUSG00000052026 | Slc6a7 | solute carrier family 6 (neurotransmitter transporter, L-proline), member 7 | -2.36 |
| ENSMUSG00000037754 | Ppp1r16b | protein phosphatase 1, regulatory subunit 16B | -2.36 |
| ENSMUSG00000002578 | Ikzf4 | IKAROS family zinc finger 4 | -2.36 |
| ENSMUSG00000041577 | Prelp | proline arginine-rich end leucine-rich repeat | -2.37 |
| ENSMUSG00000006651 | Aplp1 | amyloid beta (A4) precursor-like protein 1 | -2.38 |
| ENSMUSG00000039126 | Prune2 | prune homolog 2 | -2.38 |

| | | | |
|---|---|---|---|
| ENSMUSG00000029408 | Abcb9 | ATP-binding cassette, sub-family B (MDR/TAP), member 9 | -2.38 |
| ENSMUSG00000033427 | Upb1 | ureidopropionase, beta | -2.38 |
| ENSMUSG00000029275 | Gfi1 | growth factor independent 1 | -2.38 |
| ENSMUSG00000019986 | Ahi1 | Abelson helper integration site 1 | -2.38 |
| ENSMUSG00000022661 | Cd200 | CD200 antigen | -2.38 |
| ENSMUSG00000037341 | Slc9a7 | solute carrier family 9 (sodium/hydrogen exchanger), member 7 | -2.38 |
| ENSMUSG00000021876 | Rnase4 | ribonuclease, RNase A family 4 | -2.39 |
| ENSMUSG00000021986 | Amer2 | APC membrane recruitment 2 | -2.39 |
| ENSMUSG00000035486 | Plk5 | polo like kinase 5 | -2.39 |
| ENSMUSG00000043789 | Vwce | von Willebrand factor C and EGF domains | -2.40 |
| ENSMUSG00000001663 | Gstt1 | glutathione S-transferase, theta 1 | -2.40 |
| ENSMUSG00000043458 | Pcdhb12 | protocadherin beta 12 | -2.40 |
| ENSMUSG00000103466 | Gm37247 | predicted gene, 37247 | -2.40 |
| ENSMUSG00000036295 | Lrrn3 | leucine rich repeat protein 3, neuronal | -2.41 |
| ENSMUSG00000022208 | Jph4 | junctophilin 4 | -2.41 |
| ENSMUSG00000042096 | Dao | D-amino acid oxidase | -2.41 |
| ENSMUSG00000056031 | 9330154J02Rik | RIKEN cDNA 9330154J02 gene | -2.41 |
| ENSMUSG00000055430 | Nap1l5 | nucleosome assembly protein 1-like 5 | -2.41 |
| ENSMUSG00000069227 | Gprin1 | G protein-regulated inducer of neurite outgrowth 1 | -2.42 |
| ENSMUSG00000020599 | Rgs9 | regulator of G-protein signaling 9 | -2.42 |
| ENSMUSG00000112124 | Gm19056 | predicted gene, 19056 | -2.42 |
| ENSMUSG00000116287 | Gm3924 | predicted gene 3924 | -2.42 |
| ENSMUSG00000003153 | Slc2a3 | solute carrier family 2 (facilitated glucose transporter), member 3 | -2.42 |
| ENSMUSG00000020990 | Cdkl1 | cyclin-dependent kinase-like 1 (CDC2-related kinase) | -2.42 |
| ENSMUSG00000028972 | Car6 | carbonic anhydrase 6 | -2.42 |
| ENSMUSG00000021913 | Ogdhl | oxoglutarate dehydrogenase-like | -2.43 |
| ENSMUSG00000039579 | Grin3a | glutamate receptor ionotropic, NMDA3A | -2.43 |
| ENSMUSG00000038555 | Reep2 | receptor accessory protein 2 | -2.43 |
| ENSMUSG00000062296 | Trank1 | tetratricopeptide repeat and ankyrin repeat containing 1 | -2.43 |
| ENSMUSG00000019467 | Arhgef25 | Rho guanine nucleotide exchange factor (GEF) 25 | -2.43 |
| ENSMUSG00000005672 | Kit | KIT proto-oncogene receptor tyrosine kinase | -2.44 |
| ENSMUSG00000027230 | Creb3l1 | cAMP responsive element binding protein 3-like 1 | -2.44 |
| ENSMUSG00000041268 | Dmxl2 | Dmx-like 2 | -2.44 |
| ENSMUSG00000044694 | 2010007H06Rik | RIKEN cDNA 2010007H06 gene | -2.44 |
| ENSMUSG00000028439 | Fam219a | family with sequence similarity 219, member A | -2.44 |
| ENSMUSG00000091735 | Gpr62 | G protein-coupled receptor 62 | -2.44 |
| ENSMUSG00000087675 | Gm11762 | predicted gene 11762 | -2.44 |
| ENSMUSG00000102481 | Gm37925 | predicted gene, 37925 | -2.44 |
| ENSMUSG00000033730 | Egr3 | early growth response 3 | -2.45 |
| ENSMUSG00000062044 | Lmtk3 | lemur tyrosine kinase 3 | -2.45 |
| ENSMUSG00000030911 | Zp2 | zona pellucida glycoprotein 2 | -2.45 |
| ENSMUSG00000006575 | Rundc3a | RUN domain containing 3A | -2.46 |
| ENSMUSG00000035580 | Kcnh8 | potassium voltage-gated channel, subfamily H (eag-related), member 8 | -2.46 |
| ENSMUSG00000074657 | Kif5a | kinesin family member 5A | -2.46 |
| ENSMUSG00000045657 | Pcdhb10 | protocadherin beta 10 | -2.46 |
| ENSMUSG00000117254 | AC171111.1 | novel transcript | -2.47 |
| ENSMUSG00000034438 | Gbp8 | guanylate-binding protein 8 | -2.47 |
| ENSMUSG00000097587 | 4930578M01Rik | RIKEN cDNA 4930578M01 gene | -2.47 |
| ENSMUSG00000022580 | Rhpn1 | rhophilin, Rho GTPase binding protein 1 | -2.47 |
| ENSMUSG00000102424 | Paupar | Pax6 upstream antisense RNA | -2.47 |
| ENSMUSG00000069378 | Prdm6 | PR domain containing 6 | -2.47 |
| ENSMUSG00000038692 | Hoxb4 | homeobox B4 | -2.48 |
| ENSMUSG00000112433 | Gm30122 | predicted gene, 30122 | -2.48 |
| ENSMUSG00000045467 | Ttll13 | tubulin tyrosine ligase-like family, member 13 | -2.48 |
| ENSMUSG00000042351 | Grap2 | GRB2-related adaptor protein 2 | -2.48 |
| ENSMUSG00000034111 | Tmed8 | transmembrane p24 trafficking protein 8 | -2.49 |
| ENSMUSG00000029102 | Hgfac | hepatocyte growth factor activator | -2.49 |
| ENSMUSG00000074882 | Cyp2c68 | cytochrome P450, family 2, subfamily c, polypeptide 68 | -2.49 |
| ENSMUSG00000001248 | Gramd1a | GRAM domain containing 1A | -2.49 |
| ENSMUSG00000029298 | Gbp9 | guanylate-binding protein 9 | -2.49 |
| ENSMUSG00000025348 | Itga7 | integrin alpha 7 | -2.49 |
| ENSMUSG00000060843 | Ctnna3 | catenin (cadherin associated protein), alpha 3 | -2.49 |
| ENSMUSG00000050822 | Slc29a4 | solute carrier family 29 (nucleoside transporters), member 4 | -2.50 |
| ENSMUSG00000079442 | St6galnac4 | ST6 (alpha-N-acetyl-neuraminyl-2,3-beta-galactosyl-1,3)-N-acetylgalactosaminide a | -2.50 |
| ENSMUSG00000070498 | Tmem132b | transmembrane protein 132B | -2.50 |
| ENSMUSG00000049551 | Fzd9 | frizzled class receptor 9 | -2.51 |
| ENSMUSG00000038370 | Pcp4l1 | Purkinje cell protein 4-like 1 | -2.51 |

| | | | |
|---|---|---|---|
| ENSMUSG00000040183 | Ankrd6 | ankyrin repeat domain 6 | -2.51 |
| ENSMUSG00000063428 | Ddo | D-aspartate oxidase | -2.51 |
| ENSMUSG00000023826 | Prkn | parkin RBR E3 ubiquitin protein ligase | -2.51 |
| ENSMUSG00000051486 | Pcdhb11 | protocadherin beta 11 | -2.51 |
| ENSMUSG00000032500 | Dclk3 | doublecortin-like kinase 3 | -2.51 |
| ENSMUSG00000020847 | Rph3al | rabphilin 3A-like (without C2 domains) | -2.52 |
| ENSMUSG00000053046 | Brsk2 | BR serine/threonine kinase 2 | -2.52 |
| ENSMUSG00000048782 | Insc | INSC spindle orientation adaptor protein | -2.52 |
| ENSMUSG00000040428 | Plekha4 | pleckstrin homology domain containing, family A (phosphoinositide binding specific | -2.52 |
| ENSMUSG00000030317 | Timp4 | tissue inhibitor of metalloproteinase 4 | -2.52 |
| ENSMUSG00000111792 | Gm33858 | predicted gene, 33858 | -2.52 |
| ENSMUSG00000018830 | Myh11 | myosin, heavy polypeptide 11, smooth muscle | -2.53 |
| ENSMUSG00000103529 | A730089K16Rik | RIKEN cDNA A730089K16 gene | -2.53 |
| ENSMUSG00000089829 | Gm16565 | predicted gene 16565 | -2.53 |
| ENSMUSG00000003418 | St8sia6 | ST8 alpha-N-acetyl-neuraminide alpha-2,8-sialyltransferase 6 | -2.53 |
| ENSMUSG00000039330 | Tsga10ip | testis specific 10 interacting protein | -2.54 |
| ENSMUSG00000020589 | Fam49a | family with sequence similarity 49, member A | -2.55 |
| ENSMUSG00000074899 | Sptbn5 | spectrin beta, non-erythrocytic 5 | -2.55 |
| ENSMUSG00000097391 | Mirg | miRNA containing gene | -2.55 |
| ENSMUSG00000047250 | Ptgs1 | prostaglandin-endoperoxide synthase 1 | -2.55 |
| ENSMUSG00000030110 | Ret | ret proto-oncogene | -2.56 |
| ENSMUSG00000000632 | Sez6 | seizure related gene 6 | -2.56 |
| ENSMUSG00000048965 | Mrgpre | MAS-related GPR, member E | -2.57 |
| ENSMUSG00000050321 | Neto1 | neuropilin (NRP) and tolloid (TLL)-like 1 | -2.57 |
| ENSMUSG00000006457 | Actn3 | actinin alpha 3 | -2.57 |
| ENSMUSG00000074274 | D930028M14Rik | RIKEN cDNA D930028M14 gene | -2.57 |
| ENSMUSG00000030866 | Ern2 | endoplasmic reticulum (ER) to nucleus signalling 2 | -2.58 |
| ENSMUSG00000038065 | Mturn | maturin, neural progenitor differentiation regulator homolog (Xenopus) | -2.58 |
| ENSMUSG00000057614 | Gnai1 | guanine nucleotide binding protein (G protein), alpha inhibiting 1 | -2.58 |
| ENSMUSG00000073608 | Gal3st2c | galactose-3-O-sulfotransferase 2C | -2.58 |
| ENSMUSG00000079502 | Cfap77 | cilia and flagella associated protein 77 | -2.58 |
| ENSMUSG00000053580 | Tanc2 | tetratricopeptide repeat, ankyrin repeat and coiled-coil containing 2 | -2.59 |
| ENSMUSG00000057897 | Camk2b | calcium/calmodulin-dependent protein kinase II, beta | -2.59 |
| ENSMUSG00000030772 | Dkk3 | dickkopf WNT signaling pathway inhibitor 3 | -2.59 |
| ENSMUSG00000111293 | Gm34006 | predicted gene, 34006 | -2.59 |
| ENSMUSG00000039954 | Stk32a | serine/threonine kinase 32A | -2.59 |
| ENSMUSG00000024215 | Spdef | SAM pointed domain containing ets transcription factor | -2.60 |
| ENSMUSG00000004709 | Cd244a | CD244 molecule A | -2.60 |
| ENSMUSG00000108688 | Gm44985 | predicted gene 44985 | -2.61 |
| ENSMUSG00000029279 | Brdt | bromodomain, testis-specific | -2.61 |
| ENSMUSG00000073295 | Nudt11 | nudix (nucleoside diphosphate linked moiety X)-type motif 11 | -2.61 |
| ENSMUSG00000044288 | Cnr1 | cannabinoid receptor 1 (brain) | -2.62 |
| ENSMUSG00000006362 | Cbfa2t3 | core-binding factor, runt domain, alpha subunit 2, translocated to, 3 (human) | -2.63 |
| ENSMUSG00000020581 | Agr2 | anterior gradient 2 | -2.63 |
| ENSMUSG00000073043 | Atoh1 | atonal bHLH transcription factor 1 | -2.63 |
| ENSMUSG00000001281 | Itgb7 | integrin beta 7 | -2.63 |
| ENSMUSG00000042763 | Maneal | mannosidase, endo-alpha-like | -2.63 |
| ENSMUSG00000112197 | Gm47337 | predicted gene, 47337 | -2.63 |
| ENSMUSG00000110394 | Gm18991 | predicted gene, 18991 | -2.63 |
| ENSMUSG00000025701 | Alox5 | arachidonate 5-lipoxygenase | -2.64 |
| ENSMUSG00000092517 | Art2a-ps | ADP-ribosyltransferase 2a, pseudogene | -2.64 |
| ENSMUSG00000051652 | Lrrc3 | leucine rich repeat containing 3 | -2.64 |
| ENSMUSG00000005045 | Chd5 | chromodomain helicase DNA binding protein 5 | -2.65 |
| ENSMUSG00000032085 | Tagln | transgelin | -2.65 |
| ENSMUSG00000086219 | Srrm4os | serine/arginine repetitive matrix 4, opposite strand | -2.65 |
| ENSMUSG00000032816 | Igdcc4 | immunoglobulin superfamily, DCC subclass, member 4 | -2.66 |
| ENSMUSG00000058488 | Kl | klotho | -2.66 |
| ENSMUSG00000050395 | Tnfsf15 | tumor necrosis factor (ligand) superfamily, member 15 | -2.67 |
| ENSMUSG00000036862 | Dchs1 | dachsous cadherin related 1 | -2.67 |
| ENSMUSG00000020788 | Atp2a3 | ATPase, Ca++ transporting, ubiquitous | -2.68 |
| ENSMUSG00000021730 | Hcn1 | hyperpolarization-activated, cyclic nucleotide-gated K+ 1 | -2.68 |
| ENSMUSG00000008153 | Clstn3 | calsyntenin 3 | -2.68 |
| ENSMUSG00000019890 | Nts | neurotensin | -2.68 |
| ENSMUSG00000031442 | Mcf2l | mcf.2 transforming sequence-like | -2.70 |
| ENSMUSG00000022469 | Rapgef3 | Rap guanine nucleotide exchange factor (GEF) 3 | -2.70 |
| ENSMUSG00000038517 | Tbkbp1 | TBK1 binding protein 1 | -2.70 |
| ENSMUSG00000015222 | Map2 | microtubule-associated protein 2 | -2.71 |

| | | | |
|---|---|---|---|
| ENSMUSG00000046999 | 1110032F04Rik | RIKEN cDNA 1110032F04 gene | -2.71 |
| ENSMUSG00000045215 | Asxl3 | additional sex combs like 3, transcriptional regulator | -2.71 |
| ENSMUSG00000110266 | Gm32742 | predicted gene, 32742 | -2.72 |
| ENSMUSG00000021013 | Ttc8 | tetratricopeptide repeat domain 8 | -2.72 |
| ENSMUSG00000085189 | Gm11963 | predicted gene 11963 | -2.72 |
| ENSMUSG00000047298 | Kcnv2 | potassium channel, subfamily V, member 2 | -2.72 |
| ENSMUSG00000001670 | Tat | tyrosine aminotransferase | -2.73 |
| ENSMUSG00000038349 | Plcl1 | phospholipase C-like 1 | -2.73 |
| ENSMUSG00000051177 | Plcb1 | phospholipase C, beta 1 | -2.74 |
| ENSMUSG00000062861 | Zfp28 | zinc finger protein 28 | -2.74 |
| ENSMUSG00000035551 | Igfbpl1 | insulin-like growth factor binding protein-like 1 | -2.74 |
| ENSMUSG00000053049 | Gm15413 | predicted gene 15413 | -2.74 |
| ENSMUSG00000032087 | Dscaml1 | DS cell adhesion molecule like 1 | -2.74 |
| ENSMUSG00000036533 | Cdc42ep3 | CDC42 effector protein (Rho GTPase binding) 3 | -2.75 |
| ENSMUSG00000019906 | Lin7a | lin-7 homolog A (C. elegans) | -2.75 |
| ENSMUSG00000037386 | Rims2 | regulating synaptic membrane exocytosis 2 | -2.76 |
| ENSMUSG00000102380 | Gm38140 | predicted gene, 38140 | -2.77 |
| ENSMUSG00000037159 | Wee2 | WEE1 homolog 2 (S. pombe) | -2.78 |
| ENSMUSG00000023009 | Nckap5l | NCK-associated protein 5-like | -2.79 |
| ENSMUSG00000048070 | Pirt | phosphoinositide-interacting regulator of transient receptor potential channels | -2.79 |
| ENSMUSG00000035509 | Fbxl21 | F-box and leucine-rich repeat protein 21 | -2.79 |
| ENSMUSG00000031374 | Zfp92 | zinc finger protein 92 | -2.80 |
| ENSMUSG00000027199 | Gatm | glycine amidinotransferase (L-arginine:glycine amidinotransferase) | -2.80 |
| ENSMUSG00000034336 | Ina | internexin neuronal intermediate filament protein, alpha | -2.81 |
| ENSMUSG00000003992 | Ssbp2 | single-stranded DNA binding protein 2 | -2.81 |
| ENSMUSG00000061947 | Serpina10 | serine (or cysteine) peptidase inhibitor, clade A (alpha-1 antiproteinase, antitrypsin | -2.81 |
| ENSMUSG00000037259 | Dzank1 | double zinc ribbon and ankyrin repeat domains 1 | -2.82 |
| ENSMUSG00000049281 | Scn3b | sodium channel, voltage-gated, type III, beta | -2.82 |
| ENSMUSG00000026828 | Galnt5 | polypeptide N-acetylgalactosaminyltransferase 5 | -2.83 |
| ENSMUSG00000034958 | Atcay | ataxia, cerebellar, Cayman type | -2.83 |
| ENSMUSG00000049044 | Rapgef4 | Rap guanine nucleotide exchange factor (GEF) 4 | -2.84 |
| ENSMUSG00000035200 | Chrnb4 | cholinergic receptor, nicotinic, beta polypeptide 4 | -2.85 |
| ENSMUSG00000037973 | Ccdc129 | coiled-coil domain containing 129 | -2.85 |
| ENSMUSG00000028626 | Col9a2 | collagen, type IX, alpha 2 | -2.87 |
| ENSMUSG00000047238 | Mageh1 | melanoma antigen, family H, 1 | -2.88 |
| ENSMUSG00000035095 | Fam167a | family with sequence similarity 167, member A | -2.88 |
| ENSMUSG00000025584 | Pde8a | phosphodiesterase 8A | -2.89 |
| ENSMUSG00000034648 | Lrrn1 | leucine rich repeat protein 1, neuronal | -2.89 |
| ENSMUSG00000059136 | Olfr539 | olfactory receptor 539 | -2.89 |
| ENSMUSG00000005503 | Evx1 | even-skipped homeobox 1 | -2.90 |
| ENSMUSG00000063919 | Srrm4 | serine/arginine repetitive matrix 4 | -2.90 |
| ENSMUSG00000026482 | Rgl1 | ral guanine nucleotide dissociation stimulator,-like 1 | -2.90 |
| ENSMUSG00000112816 | Gm48798 | predicted gene, 48798 | -2.90 |
| ENSMUSG00000111014 | Gm47795 | predicted gene, 47795 | -2.90 |
| ENSMUSG00000045382 | Cxcr4 | chemokine (C-X-C motif) receptor 4 | -2.91 |
| ENSMUSG00000017446 | C1qtnf1 | C1q and tumor necrosis factor related protein 1 | -2.91 |
| ENSMUSG00000031845 | Bco1 | beta-carotene oxygenase 1 | -2.92 |
| ENSMUSG00000037868 | Egr2 | early growth response 2 | -2.92 |
| ENSMUSG00000014602 | Kif1a | kinesin family member 1A | -2.92 |
| ENSMUSG00000036915 | Kirrel2 | kirre like nephrin family adhesion molecule 2 | -2.92 |
| ENSMUSG00000044468 | Tent5c | terminal nucleotidyltransferase 5C | -2.93 |
| ENSMUSG00000058420 | Syt17 | synaptotagmin XVII | -2.93 |
| ENSMUSG00000003545 | Fosb | FBJ osteosarcoma oncogene B | -2.94 |
| ENSMUSG00000103732 | Gm38315 | predicted gene, 38315 | -2.94 |
| ENSMUSG00000030664 | Sox6os | SRY (sex determining region Y)-box 6, opposite strand | -2.94 |
| ENSMUSG00000020019 | Ntn4 | netrin 4 | -2.95 |
| ENSMUSG00000029343 | Crybb1 | crystallin, beta B1 | -2.95 |
| ENSMUSG00000005973 | Rcn1 | reticulocalbin 1 | -2.96 |
| ENSMUSG00000025091 | Pnliprp2 | pancreatic lipase-related protein 2 | -2.96 |
| ENSMUSG00000094818 | Defa32 | defensin, alpha, 32 | -2.96 |
| ENSMUSG00000028600 | Podn | podocan | -2.97 |
| ENSMUSG00000086126 | Evx1os | even skipped homeotic gene 1, opposite strand | -2.97 |
| ENSMUSG00000032773 | Chrm1 | cholinergic receptor, muscarinic 1, CNS | -2.98 |
| ENSMUSG00000102581 | Gm37443 | predicted gene, 37443 | -2.99 |
| ENSMUSG00000030401 | Rtn2 | reticulon 2 (Z-band associated protein) | -3.00 |
| ENSMUSG00000032015 | Pou2f3 | POU domain, class 2, transcription factor 3 | -3.00 |
| ENSMUSG00000051251 | Nhlh1 | nescient helix loop helix 1 | -3.00 |

| | | | |
|---|---|---|---|
| ENSMUSG00000064043 | Trerf1 | transcriptional regulating factor 1 | -3.01 |
| ENSMUSG00000046159 | Chrm3 | cholinergic receptor, muscarinic 3, cardiac | -3.03 |
| ENSMUSG00000075702 | Selenom | selenoprotein M | -3.04 |
| ENSMUSG00000003436 | Dll3 | delta like canonical Notch ligand 3 | -3.04 |
| ENSMUSG00000087326 | Gm12503 | predicted gene 12503 | -3.04 |
| ENSMUSG00000049515 | Espnl | espin-like | -3.04 |
| ENSMUSG00000019775 | Rgs17 | regulator of G-protein signaling 17 | -3.04 |
| ENSMUSG00000021719 | Rgs7bp | regulator of G-protein signalling 7 binding protein | -3.05 |
| ENSMUSG00000064262 | Gimap8 | GTPase, IMAP family member 8 | -3.05 |
| ENSMUSG00000044043 | Pcdhb14 | protocadherin beta 14 | -3.05 |
| ENSMUSG00000034771 | Tle2 | transducin-like enhancer of split 2 | -3.05 |
| ENSMUSG00000035158 | Mitf | melanogenesis associated transcription factor | -3.05 |
| ENSMUSG00000052087 | Rgs14 | regulator of G-protein signaling 14 | -3.06 |
| ENSMUSG00000024238 | Zeb1 | zinc finger E-box binding homeobox 1 | -3.06 |
| ENSMUSG00000085329 | 2810404F17Rik | RIKEN cDNA 2810404F17 gene | -3.07 |
| ENSMUSG00000078670 | Fam174b | family with sequence similarity 174, member B | -3.08 |
| ENSMUSG00000038594 | Cep85l | centrosomal protein 85-like | -3.08 |
| ENSMUSG00000042846 | Lrrtm3 | leucine rich repeat transmembrane neuronal 3 | -3.08 |
| ENSMUSG00000017211 | Gsdma2 | gasdermin A2 | -3.09 |
| ENSMUSG00000032452 | Clstn2 | calsyntenin 2 | -3.09 |
| ENSMUSG00000028364 | Tnc | tenascin C | -3.09 |
| ENSMUSG00000036764 | Dnajc12 | DnaJ heat shock protein family (Hsp40) member C12 | -3.10 |
| ENSMUSG00000036872 | Abcc12 | ATP-binding cassette, sub-family C (CFTR/MRP), member 12 | -3.10 |
| ENSMUSG00000071424 | Grid2 | glutamate receptor, ionotropic, delta 2 | -3.10 |
| ENSMUSG00000035864 | Syt1 | synaptotagmin I | -3.10 |
| ENSMUSG00000039601 | Rcan2 | regulator of calcineurin 2 | -3.11 |
| ENSMUSG00000038319 | Kcnh2 | potassium voltage-gated channel, subfamily H (eag-related), member 2 | -3.12 |
| ENSMUSG00000096146 | Kcnj11 | potassium inwardly rectifying channel, subfamily J, member 11 | -3.12 |
| ENSMUSG00000033510 | Otud7a | OTU domain containing 7A | -3.13 |
| ENSMUSG00000039013 | Siglecf | sialic acid binding Ig-like lectin F | -3.14 |
| ENSMUSG00000010064 | Slc38a3 | solute carrier family 38, member 3 | -3.14 |
| ENSMUSG00000032936 | Camkv | CaM kinase-like vesicle-associated | -3.15 |
| ENSMUSG00000099148 | Gm3331 | predicted gene 3331 | -3.15 |
| ENSMUSG00000001333 | Sync | syncoilin | -3.16 |
| ENSMUSG00000026764 | Kif5c | kinesin family member 5C | -3.16 |
| ENSMUSG00000042757 | Tmem108 | transmembrane protein 108 | -3.16 |
| ENSMUSG00000095079 | Igha | immunoglobulin heavy constant alpha | -3.16 |
| ENSMUSG00000043391 | 2510009E07Rik | RIKEN cDNA 2510009E07 gene | -3.17 |
| ENSMUSG00000056812 | St8sia3 | ST8 alpha-N-acetyl-neuraminide alpha-2,8-sialyltransferase 3 | -3.17 |
| ENSMUSG00000106547 | B230303O12Rik | RIKEN cDNA B230303O12 gene | -3.17 |
| ENSMUSG00000113502 | Gm47411 | predicted gene, 47411 | -3.17 |
| ENSMUSG00000022441 | Efcab6 | EF-hand calcium binding domain 6 | -3.17 |
| ENSMUSG00000066191 | Anks6 | ankyrin repeat and sterile alpha motif domain containing 6 | -3.18 |
| ENSMUSG00000072623 | Zfp9 | zinc finger protein 9 | -3.18 |
| ENSMUSG00000025876 | Unc5a | unc-5 netrin receptor A | -3.19 |
| ENSMUSG00000090105 | Gm15890 | predicted gene 15890 | -3.19 |
| ENSMUSG00000102498 | Gm19445 | predicted gene, 19445 | -3.19 |
| ENSMUSG00000000214 | Th | tyrosine hydroxylase | -3.20 |
| ENSMUSG00000023247 | Guca2a | guanylate cyclase activator 2a (guanylin) | -3.21 |
| ENSMUSG00000038486 | Sv2a | synaptic vesicle glycoprotein 2 a | -3.21 |
| ENSMUSG00000063704 | Mapk15 | mitogen-activated protein kinase 15 | -3.22 |
| ENSMUSG00000035459 | Stab2 | stabilin 2 | -3.22 |
| ENSMUSG00000038599 | Capn8 | calpain 8 | -3.23 |
| ENSMUSG00000060671 | Atp8b2 | ATPase, class I, type 8B, member 2 | -3.25 |
| ENSMUSG00000033220 | Rac2 | Rac family small GTPase 2 | -3.25 |
| ENSMUSG00000018470 | Kcnab3 | potassium voltage-gated channel, shaker-related subfamily, beta member 3 | -3.26 |
| ENSMUSG00000030616 | Sytl2 | synaptotagmin-like 2 | -3.27 |
| ENSMUSG00000079037 | Prnp | prion protein | -3.27 |
| ENSMUSG00000044548 | Dact1 | dishevelled-binding antagonist of beta-catenin 1 | -3.27 |
| ENSMUSG00000086141 | 9030622O22Rik | RIKEN cDNA 9030622O22 gene | -3.27 |
| ENSMUSG00000042515 | Mum1l1 | melanoma associated antigen (mutated) 1-like 1 | -3.27 |
| ENSMUSG00000114985 | 1810028F09Rik | RIKEN cDNA 1810028F09 gene | -3.28 |
| ENSMUSG00000040146 | Rgl3 | ral guanine nucleotide dissociation stimulator-like 3 | -3.29 |
| ENSMUSG00000086231 | Rapgef4os3 | Rap guanine nucleotide exchange factor (GEF) 4, opposite strand 3 | -3.29 |
| ENSMUSG00000074436 | Defa-ps18 | defensin, alpha, pseudogene 18 | -3.29 |
| ENSMUSG00000029134 | Plb1 | phospholipase B1 | -3.30 |
| ENSMUSG00000041673 | Lrrc18 | leucine rich repeat containing 18 | -3.30 |

| | | | |
|---|---|---|---|
| ENSMUSG00000026950 | Neb | nebulin | -3.31 |
| ENSMUSG00000013846 | St3gal1 | ST3 beta-galactoside alpha-2,3-sialyltransferase 1 | -3.31 |
| ENSMUSG00000013523 | Bcas1 | breast carcinoma amplified sequence 1 | -3.31 |
| ENSMUSG00000038132 | Rbm24 | RNA binding motif protein 24 | -3.31 |
| ENSMUSG00000048763 | Hoxb3 | homeobox B3 | -3.32 |
| ENSMUSG00000107061 | Gm19590 | predicted gene, 19590 | -3.32 |
| ENSMUSG00000024743 | Syt7 | synaptotagmin VII | -3.33 |
| ENSMUSG00000015968 | Cacna1d | calcium channel, voltage-dependent, L type, alpha 1D subunit | -3.33 |
| ENSMUSG00000113680 | Gm47405 | predicted gene, 47405 | -3.34 |
| ENSMUSG00000038738 | Shank1 | SH3 and multiple ankyrin repeat domains 1 | -3.34 |
| ENSMUSG00000060735 | Rxfp3 | relaxin family peptide receptor 3 | -3.34 |
| ENSMUSG00000061414 | Cracr2a | calcium release activated channel regulator 2A | -3.35 |
| ENSMUSG00000024548 | Setbp1 | SET binding protein 1 | -3.35 |
| ENSMUSG00000040528 | Milr1 | mast cell immunoglobulin like receptor 1 | -3.35 |
| ENSMUSG00000037106 | Fer1l6 | fer-1-like 6 (C. elegans) | -3.36 |
| ENSMUSG00000034818 | Celf5 | CUGBP, Elav-like family member 5 | -3.36 |
| ENSMUSG00000020340 | Cyfip2 | cytoplasmic FMR1 interacting protein 2 | -3.37 |
| ENSMUSG00000086245 | Gm16170 | predicted gene 16170 | -3.37 |
| ENSMUSG00000031510 | 1700128E19Rik | RIKEN cDNA 1700128E19 gene | -3.37 |
| ENSMUSG00000032502 | Stac | src homology three (SH3) and cysteine rich domain | -3.38 |
| ENSMUSG00000018486 | Wnt9b | wingless-type MMTV integration site family, member 9B | -3.38 |
| ENSMUSG00000026475 | Rgs16 | regulator of G-protein signaling 16 | -3.38 |
| ENSMUSG00000074344 | Tmigd3 | transmembrane and immunoglobulin domain containing 3 | -3.38 |
| ENSMUSG00000028528 | Dnajc6 | DnaJ heat shock protein family (Hsp40) member C6 | -3.38 |
| ENSMUSG00000032202 | Rab27a | RAB27A, member RAS oncogene family | -3.39 |
| ENSMUSG00000050821 | Fam131a | family with sequence similarity 131, member A | -3.39 |
| ENSMUSG00000026285 | Pdcd1 | programmed cell death 1 | -3.39 |
| ENSMUSG00000016763 | Scube1 | signal peptide, CUB domain, EGF-like 1 | -3.40 |
| ENSMUSG00000036306 | Lzts1 | leucine zipper, putative tumor suppressor 1 | -3.40 |
| ENSMUSG00000023868 | Pde10a | phosphodiesterase 10A | -3.40 |
| ENSMUSG00000057716 | Tmem178b | transmembrane protein 178B | -3.40 |
| ENSMUSG00000020218 | Wif1 | Wnt inhibitory factor 1 | -3.40 |
| ENSMUSG00000042078 | Svop | SV2 related protein | -3.40 |
| ENSMUSG00000021609 | Slc6a3 | solute carrier family 6 (neurotransmitter transporter, dopamine), member 3 | -3.41 |
| ENSMUSG00000059588 | Calcrl | calcitonin receptor-like | -3.41 |
| ENSMUSG00000033900 | Map9 | microtubule-associated protein 9 | -3.43 |
| ENSMUSG00000093805 | Gal3st2b | galactose-3-O-sulfotransferase 2B | -3.43 |
| ENSMUSG00000091017 | Fam71a | family with sequence similarity 71, member A | -3.43 |
| ENSMUSG00000072966 | Gprasp2 | G protein-coupled receptor associated sorting protein 2 | -3.43 |
| ENSMUSG00000040121 | Rep15 | RAB15 effector protein | -3.44 |
| ENSMUSG00000037239 | Spred3 | sprouty-related, EVH1 domain containing 3 | -3.44 |
| ENSMUSG00000026288 | Inpp5d | inositol polyphosphate-5-phosphatase D | -3.44 |
| ENSMUSG00000007594 | Hapln4 | hyaluronan and proteoglycan link protein 4 | -3.44 |
| ENSMUSG00000084048 | Gm12042 | predicted gene 12042 | -3.44 |
| ENSMUSG00000044748 | Defb1 | defensin beta 1 | -3.45 |
| ENSMUSG00000000197 | Nalcn | sodium leak channel, non-selective | -3.45 |
| ENSMUSG00000027661 | Slc2a10 | solute carrier family 2 (facilitated glucose transporter), member 10 | -3.46 |
| ENSMUSG00000032826 | Ank2 | ankyrin 2, brain | -3.46 |
| ENSMUSG00000000125 | Wnt3 | wingless-type MMTV integration site family, member 3 | -3.47 |
| ENSMUSG00000026463 | Atp2b4 | ATPase, Ca++ transporting, plasma membrane 4 | -3.47 |
| ENSMUSG00000047730 | Fcgbp | Fc fragment of IgG binding protein | -3.47 |
| ENSMUSG00000030087 | Klf15 | Kruppel-like factor 15 | -3.47 |
| ENSMUSG00000034460 | Six4 | sine oculis-related homeobox 4 | -3.47 |
| ENSMUSG00000018593 | Sparc | secreted acidic cysteine rich glycoprotein | -3.48 |
| ENSMUSG00000085247 | 4930545L23Rik | RIKEN cDNA 4930545L23 gene | -3.48 |
| ENSMUSG00000021559 | Dapk1 | death associated protein kinase 1 | -3.49 |
| ENSMUSG00000023032 | Slc4a8 | solute carrier family 4 (anion exchanger), member 8 | -3.49 |
| ENSMUSG00000021614 | Vcan | versican | -3.50 |
| ENSMUSG00000098557 | Kctd12 | potassium channel tetramerisation domain containing 12 | -3.51 |
| ENSMUSG00000020102 | Slc16a7 | solute carrier family 16 (monocarboxylic acid transporters), member 7 | -3.51 |
| ENSMUSG00000042804 | Gpr153 | G protein-coupled receptor 153 | -3.51 |
| ENSMUSG00000022456 | 37865 | septin 3 | -3.52 |
| ENSMUSG00000041681 | Iapp | islet amyloid polypeptide | -3.52 |
| ENSMUSG00000010086 | Rnf112 | ring finger protein 112 | -3.52 |
| ENSMUSG00000035896 | Rnase1 | ribonuclease, RNase A family, 1 (pancreatic) | -3.53 |
| ENSMUSG00000029875 | Ccdc184 | coiled-coil domain containing 184 | -3.53 |
| ENSMUSG00000070576 | Mn1 | meningioma 1 | -3.53 |

| | | | |
|---|---|---|---|
| ENSMUSG00000025272 | Tro | trophinin | -3.54 |
| ENSMUSG00000030428 | Ttyh1 | tweety family member 1 | -3.55 |
| ENSMUSG00000015484 | Fam163a | family with sequence similarity 163, member A | -3.55 |
| ENSMUSG00000013921 | Clip3 | CAP-GLY domain containing linker protein 3 | -3.55 |
| ENSMUSG00000078776 | 9530053A07Rik | RIKEN cDNA 9530053A07 gene | -3.56 |
| ENSMUSG00000089995 | Gm15716 | predicted gene 15716 | -3.56 |
| ENSMUSG00000032268 | Tmprss5 | transmembrane protease, serine 5 (spinesin) | -3.56 |
| ENSMUSG00000024330 | Col11a2 | collagen, type XI, alpha 2 | -3.57 |
| ENSMUSG00000026834 | Acvr1c | activin A receptor, type IC | -3.57 |
| ENSMUSG00000049047 | Armcx3 | armadillo repeat containing, X-linked 3 | -3.57 |
| ENSMUSG00000030411 | Nova2 | neuro-oncological ventral antigen 2 | -3.57 |
| ENSMUSG00000032035 | Ets1 | E26 avian leukemia oncogene 1, 5' domain | -3.58 |
| ENSMUSG00000024029 | Tff3 | trefoil factor 3, intestinal | -3.59 |
| ENSMUSG00000080715 | Gm5406 | predicted gene 5406 | -3.59 |
| ENSMUSG00000036912 | Piwil4 | piwi-like RNA-mediated gene silencing 4 | -3.59 |
| ENSMUSG00000059921 | Unc5c | unc-5 netrin receptor C | -3.59 |
| ENSMUSG00000034107 | Ano7 | anoctamin 7 | -3.60 |
| ENSMUSG00000018927 | Ccl6 | chemokine (C-C motif) ligand 6 | -3.60 |
| ENSMUSG00000043518 | Rai2 | retinoic acid induced 2 | -3.60 |
| ENSMUSG00000047642 | D930020B18Rik | RIKEN cDNA D930020B18 gene | -3.60 |
| ENSMUSG00000055963 | Triqk | triple QxxK/R motif containing | -3.61 |
| ENSMUSG00000001120 | Pcbp3 | poly(rC) binding protein 3 | -3.62 |
| ENSMUSG00000024897 | Apba1 | amyloid beta (A4) precursor protein binding, family A, member 1 | -3.62 |
| ENSMUSG00000031028 | Tub | tubby bipartite transcription factor | -3.62 |
| ENSMUSG00000042179 | Pnliprp1 | pancreatic lipase related protein 1 | -3.63 |
| ENSMUSG00000022425 | Enpp2 | ectonucleotide pyrophosphatase/phosphodiesterase 2 | -3.63 |
| ENSMUSG00000020704 | Asic2 | acid-sensing (proton-gated) ion channel 2 | -3.63 |
| ENSMUSG00000001827 | Folr1 | folate receptor 1 (adult) | -3.63 |
| ENSMUSG00000032744 | Heyl | hairy/enhancer-of-split related with YRPW motif-like | -3.63 |
| ENSMUSG00000104283 | Gm37459 | predicted gene, 37459 | -3.63 |
| ENSMUSG00000035964 | Tmem59l | transmembrane protein 59-like | -3.64 |
| ENSMUSG00000028445 | Enho | energy homeostasis associated | -3.64 |
| ENSMUSG00000053552 | Ebf4 | early B cell factor 4 | -3.64 |
| ENSMUSG00000036330 | Slc18a1 | solute carrier family 18 (vesicular monoamine), member 1 | -3.65 |
| ENSMUSG00000086999 | Bcas1os2 | breast carcinoma amplified sequence 1, opposite strand 2 | -3.65 |
| ENSMUSG00000040136 | Abcc8 | ATP-binding cassette, sub-family C (CFTR/MRP), member 8 | -3.66 |
| ENSMUSG00000032064 | Dixdc1 | DIX domain containing 1 | -3.67 |
| ENSMUSG00000001802 | Lrp3 | low density lipoprotein receptor-related protein 3 | -3.67 |
| ENSMUSG00000031428 | Zcchc18 | zinc finger, CCHC domain containing 18 | -3.67 |
| ENSMUSG00000020120 | Plek | pleckstrin | -3.67 |
| ENSMUSG00000024598 | Fbn2 | fibrillin 2 | -3.68 |
| ENSMUSG00000054169 | Ceacam10 | carcinoembryonic antigen-related cell adhesion molecule 10 | -3.69 |
| ENSMUSG00000015396 | Cd83 | CD83 antigen | -3.69 |
| ENSMUSG00000040563 | Plppr2 | phospholipid phosphatase related 2 | -3.69 |
| ENSMUSG00000003283 | Hck | hemopoietic cell kinase | -3.69 |
| ENSMUSG00000097311 | Gm26871 | predicted gene, 26871 | -3.69 |
| ENSMUSG00000020181 | Nav3 | neuron navigator 3 | -3.70 |
| ENSMUSG00000031760 | Mt3 | metallothionein 3 | -3.70 |
| ENSMUSG00000020601 | Trib2 | tribbles pseudokinase 2 | -3.70 |
| ENSMUSG00000034472 | Rasd2 | RASD family, member 2 | -3.71 |
| ENSMUSG00000028749 | Pla2g2f | phospholipase A2, group IIF | -3.72 |
| ENSMUSG00000018862 | Otop3 | otopetrin 3 | -3.72 |
| ENSMUSG00000027520 | Zdbf2 | zinc finger, DBF-type containing 2 | -3.72 |
| ENSMUSG00000004961 | Syt5 | synaptotagmin V | -3.73 |
| ENSMUSG00000056004 | 9330182L06Rik | RIKEN cDNA 9330182L06 gene | -3.73 |
| ENSMUSG00000044017 | Adgrd1 | adhesion G protein-coupled receptor D1 | -3.73 |
| ENSMUSG00000044312 | Neurog3 | neurogenin 3 | -3.73 |
| ENSMUSG00000035513 | Ntng2 | netrin G2 | -3.73 |
| ENSMUSG00000032377 | Plscr4 | phospholipid scramblase 4 | -3.73 |
| ENSMUSG00000028354 | Fmn2 | formin 2 | -3.73 |
| ENSMUSG00000053182 | Gm609 | predicted gene 609 | -3.74 |
| ENSMUSG00000025075 | Habp2 | hyaluronic acid binding protein 2 | -3.74 |
| ENSMUSG00000027674 | Pex5l | peroxisomal biogenesis factor 5-like | -3.74 |
| ENSMUSG00000021587 | Pcsk1 | proprotein convertase subtilisin/kexin type 1 | -3.75 |
| ENSMUSG00000064125 | Prr36 | proline rich 36 | -3.75 |
| ENSMUSG00000026870 | Cutal | cutA divalent cation tolerance homolog-like | -3.75 |
| ENSMUSG00000021539 | Lect2 | leukocyte cell-derived chemotaxin 2 | -3.76 |

| | | | |
|---|---|---|---|
| ENSMUSG00000054453 | Sytl5 | synaptotagmin-like 5 | -3.76 |
| ENSMUSG00000018169 | Mfng | MFNG O-fucosylpeptide 3-beta-N-acetylglucosaminyltransferase | -3.76 |
| ENSMUSG00000047867 | Gimap6 | GTPase, IMAP family member 6 | -3.76 |
| ENSMUSG00000031377 | Bmx | BMX non-receptor tyrosine kinase | -3.77 |
| ENSMUSG00000022829 | Stxbp5l | syntaxin binding protein 5-like | -3.77 |
| ENSMUSG00000009246 | Trpm5 | transient receptor potential cation channel, subfamily M, member 5 | -3.78 |
| ENSMUSG00000062444 | Ap3b2 | adaptor-related protein complex 3, beta 2 subunit | -3.78 |
| ENSMUSG00000037035 | Inhbb | inhibin beta-B | -3.78 |
| ENSMUSG00000072875 | Gpr27 | G protein-coupled receptor 27 | -3.78 |
| ENSMUSG00000025551 | Fgf14 | fibroblast growth factor 14 | -3.78 |
| ENSMUSG00000020435 | Osbp2 | oxysterol binding protein 2 | -3.79 |
| ENSMUSG00000029223 | Uchl1 | ubiquitin carboxy-terminal hydrolase L1 | -3.79 |
| ENSMUSG00000101126 | Gm10538 | predicted gene 10538 | -3.79 |
| ENSMUSG00000040170 | Fmo2 | flavin containing monooxygenase 2 | -3.80 |
| ENSMUSG00000037579 | Kcnh3 | potassium voltage-gated channel, subfamily H (eag-related), member 3 | -3.81 |
| ENSMUSG00000069171 | Nr2f1 | nuclear receptor subfamily 2, group F, member 1 | -3.81 |
| ENSMUSG00000022995 | Enah | ENAH actin regulator | -3.82 |
| ENSMUSG00000037306 | Man1c1 | mannosidase, alpha, class 1C, member 1 | -3.82 |
| ENSMUSG00000019122 | Ccl9 | chemokine (C-C motif) ligand 9 | -3.83 |
| ENSMUSG00000020101 | Vsir | V-set immunoregulatory receptor | -3.83 |
| ENSMUSG00000020312 | Shc2 | SHC (Src homology 2 domain containing) transforming protein 2 | -3.84 |
| ENSMUSG00000027947 | Il6ra | interleukin 6 receptor, alpha | -3.84 |
| ENSMUSG00000080115 | Eef1akmt3 | EEF1A lysine methyltransferase 3 | -3.84 |
| ENSMUSG00000017692 | Rhbdl3 | rhomboid like 3 | -3.84 |
| ENSMUSG00000048216 | Gpr85 | G protein-coupled receptor 85 | -3.84 |
| ENSMUSG00000102123 | Gm4319 | predicted gene 4319 | -3.84 |
| ENSMUSG00000038916 | Soga3 | SOGA family member 3 | -3.85 |
| ENSMUSG00000034799 | Unc13a | unc-13 homolog A | -3.86 |
| ENSMUSG00000046922 | Gpr6 | G protein-coupled receptor 6 | -3.86 |
| ENSMUSG00000081303 | Gm16011 | predicted gene 16011 | -3.87 |
| ENSMUSG00000051832 | E230016K23Rik | RIKEN cDNA E230016K23 gene | -3.87 |
| ENSMUSG00000027249 | F2 | coagulation factor II | -3.87 |
| ENSMUSG00000038665 | Dgki | diacylglycerol kinase, iota | -3.87 |
| ENSMUSG00000060063 | Alox5ap | arachidonate 5-lipoxygenase activating protein | -3.87 |
| ENSMUSG00000005883 | Spo11 | SPO11 meiotic protein covalently bound to DSB | -3.87 |
| ENSMUSG00000074442 | Defa31 | defensin, alpha, 31 | -3.87 |
| ENSMUSG00000055733 | Nap1l3 | nucleosome assembly protein 1-like 3 | -3.87 |
| ENSMUSG00000022148 | Fyb | FYN binding protein | -3.88 |
| ENSMUSG00000031551 | Ido1 | indoleamine 2,3-dioxygenase 1 | -3.88 |
| ENSMUSG00000037610 | Kcnmb2 | potassium large conductance calcium-activated channel, subfamily M, beta membe | -3.88 |
| ENSMUSG00000040148 | Hmx3 | H6 homeobox 3 | -3.89 |
| ENSMUSG00000052271 | Bhlha15 | basic helix-loop-helix family, member a15 | -3.90 |
| ENSMUSG00000033615 | Cplx1 | complexin 1 | -3.91 |
| ENSMUSG00000114751 | 5430425E15Rik | RIKEN cDNA 5430425E15 gene | -3.91 |
| ENSMUSG00000063903 | Klk1 | kallikrein 1 | -3.92 |
| ENSMUSG00000024030 | Abcg1 | ATP binding cassette subfamily G member 1 | -3.92 |
| ENSMUSG00000043388 | Tmem130 | transmembrane protein 130 | -3.92 |
| ENSMUSG00000001985 | Grik3 | glutamate receptor, ionotropic, kainate 3 | -3.92 |
| ENSMUSG00000031981 | Capn9 | calpain 9 | -3.95 |
| ENSMUSG00000042115 | Klhdc8a | kelch domain containing 8A | -3.95 |
| ENSMUSG00000044933 | Sstr3 | somatostatin receptor 3 | -3.96 |
| ENSMUSG00000057914 | Cacnb2 | calcium channel, voltage-dependent, beta 2 subunit | -3.96 |
| ENSMUSG00000074434 | Defa28 | defensin, alpha, 28 | -3.96 |
| ENSMUSG00000027570 | Col9a3 | collagen, type IX, alpha 3 | -3.97 |
| ENSMUSG00000024013 | Fgd2 | FYVE, RhoGEF and PH domain containing 2 | -3.97 |
| ENSMUSG00000028255 | Clca1 | chloride channel accessory 1 | -3.98 |
| ENSMUSG00000052212 | Cd177 | CD177 antigen | -3.99 |
| ENSMUSG00000057315 | Arhgap24 | Rho GTPase activating protein 24 | -3.99 |
| ENSMUSG00000060615 | Ang4 | angiogenin, ribonuclease A family, member 4 | -3.99 |
| ENSMUSG00000058589 | Anks1b | ankyrin repeat and sterile alpha motif domain containing 1B | -3.99 |
| ENSMUSG00000028865 | Cd164l2 | CD164 sialomucin-like 2 | -3.99 |
| ENSMUSG00000022211 | Carmil3 | capping protein regulator and myosin 1 linker 3 | -4.00 |
| ENSMUSG00000097266 | Gm26802 | predicted gene, 26802 | -4.00 |
| ENSMUSG00000028078 | Dclk2 | doublecortin-like kinase 2 | -4.01 |
| ENSMUSG00000029245 | Epha5 | Eph receptor A5 | -4.01 |
| ENSMUSG00000038112 | AW551984 | expressed sequence AW551984 | -4.01 |
| ENSMUSG00000070687 | Htr1d | 5-hydroxytryptamine (serotonin) receptor 1D | -4.01 |

| | | | |
|---|---|---|---|
| ENSMUSG00000041794 | Myrip | myosin VIIA and Rab interacting protein | -4.02 |
| ENSMUSG00000024011 | Pi16 | peptidase inhibitor 16 | -4.02 |
| ENSMUSG00000071550 | Cfap44 | cilia and flagella associated protein 44 | -4.03 |
| ENSMUSG00000029368 | Alb | albumin | -4.03 |
| ENSMUSG00000019232 | Etnppl | ethanolamine phosphate phospholyase | -4.04 |
| ENSMUSG00000042766 | Trim46 | tripartite motif-containing 46 | -4.04 |
| ENSMUSG00000021087 | Rtn1 | reticulon 1 | -4.04 |
| ENSMUSG00000041544 | Disp3 | dispatched RND transporter family member 3 | -4.04 |
| ENSMUSG00000079481 | Nhsl2 | NHS-like 2 | -4.05 |
| ENSMUSG00000069515 | Lyz1 | lysozyme 1 | -4.06 |
| ENSMUSG00000042873 | Lhfpl4 | lipoma HMGIC fusion partner-like protein 4 | -4.08 |
| ENSMUSG00000030074 | Gxylt2 | glucoside xylosyltransferase 2 | -4.08 |
| ENSMUSG00000030302 | Atp2b2 | ATPase, Ca++ transporting, plasma membrane 2 | -4.09 |
| ENSMUSG00000084934 | Gm16035 | predicted gene 16035 | -4.09 |
| ENSMUSG00000040723 | Rcsd1 | RCSD domain containing 1 | -4.10 |
| ENSMUSG00000038580 | Sct | secretin | -4.11 |
| ENSMUSG00000031980 | Agt | angiotensinogen (serpin peptidase inhibitor, clade A, member 8) | -4.11 |
| ENSMUSG00000044177 | Wfikkn2 | WAP, follistatin/kazal, immunoglobulin, kunitz and netrin domain containing 2 | -4.11 |
| ENSMUSG00000021069 | Pygl | liver glycogen phosphorylase | -4.11 |
| ENSMUSG00000040412 | 5330417C22Rik | RIKEN cDNA 5330417C22 gene | -4.12 |
| ENSMUSG00000019846 | Lama4 | laminin, alpha 4 | -4.12 |
| ENSMUSG00000024084 | Qpct | glutaminyl-peptide cyclotransferase (glutaminyl cyclase) | -4.12 |
| ENSMUSG00000097520 | 4930488L21Rik | RIKEN cDNA 4930488L21 gene | -4.12 |
| ENSMUSG00000016200 | Syt14 | synaptotagmin XIV | -4.13 |
| ENSMUSG00000034353 | Ramp1 | receptor (calcitonin) activity modifying protein 1 | -4.13 |
| ENSMUSG00000025582 | Nptx1 | neuronal pentraxin 1 | -4.13 |
| ENSMUSG00000022650 | Retnlb | resistin like beta | -4.14 |
| ENSMUSG00000033200 | Tpsg1 | tryptase gamma 1 | -4.14 |
| ENSMUSG00000114501 | Gm48582 | predicted gene, 48582 | -4.14 |
| ENSMUSG00000025867 | Cplx2 | complexin 2 | -4.16 |
| ENSMUSG00000031748 | Gnao1 | guanine nucleotide binding protein, alpha O | -4.16 |
| ENSMUSG00000070337 | Gpr179 | G protein-coupled receptor 179 | -4.16 |
| ENSMUSG00000056296 | Synpr | synaptoporin | -4.16 |
| ENSMUSG00000026527 | Rgs7 | regulator of G protein signaling 7 | -4.17 |
| ENSMUSG00000022054 | Nefm | neurofilament, medium polypeptide | -4.18 |
| ENSMUSG00000099025 | Gm27162 | predicted gene 27162 | -4.18 |
| ENSMUSG00000020175 | Rab36 | RAB36, member RAS oncogene family | -4.18 |
| ENSMUSG00000057060 | Slc35f3 | solute carrier family 35, member F3 | -4.18 |
| ENSMUSG00000071113 | Mboat4 | membrane bound O-acyltransferase domain containing 4 | -4.19 |
| ENSMUSG00000049148 | Plcxd3 | phosphatidylinositol-specific phospholipase C, X domain containing 3 | -4.19 |
| ENSMUSG00000091890 | A830073O21Rik | RIKEN cDNA A830073O21 gene | -4.19 |
| ENSMUSG00000024299 | Adamts10 | a disintegrin-like and metallopeptidase (reprolysin type) with thrombospondin type | -4.20 |
| ENSMUSG00000054932 | Afp | alpha fetoprotein | -4.20 |
| ENSMUSG00000030518 | Fam189a1 | family with sequence similarity 189, member A1 | -4.20 |
| ENSMUSG00000023216 | Epb42 | erythrocyte membrane protein band 4.2 | -4.21 |
| ENSMUSG00000028415 | Spink4 | serine peptidase inhibitor, Kazal type 4 | -4.21 |
| ENSMUSG00000049122 | Frmd3 | FERM domain containing 3 | -4.21 |
| ENSMUSG00000044156 | Hepacam2 | HEPACAM family member 2 | -4.22 |
| ENSMUSG00000074446 | Defa23 | defensin, alpha, 23 | -4.23 |
| ENSMUSG00000032625 | Thsd7a | thrombospondin, type I, domain containing 7A | -4.23 |
| ENSMUSG00000033174 | Mgll | monoglyceride lipase | -4.23 |
| ENSMUSG00000029778 | Adcyap1r1 | adenylate cyclase activating polypeptide 1 receptor 1 | -4.23 |
| ENSMUSG00000036699 | Zcchc12 | zinc finger, CCHC domain containing 12 | -4.23 |
| ENSMUSG00000109561 | Ankrd31 | ankyrin repeat domain 31 | -4.24 |
| ENSMUSG00000027273 | Snap25 | synaptosomal-associated protein 25 | -4.26 |
| ENSMUSG00000004151 | Etv1 | ets variant 1 | -4.27 |
| ENSMUSG00000025515 | Muc2 | mucin 2 | -4.28 |
| ENSMUSG00000023473 | Celsr3 | cadherin, EGF LAG seven-pass G-type receptor 3 | -4.29 |
| ENSMUSG00000049350 | Zg16 | zymogen granule protein 16 | -4.29 |
| ENSMUSG00000106386 | Gm42991 | predicted gene 42991 | -4.29 |
| ENSMUSG00000044988 | Ucn3 | urocortin 3 | -4.29 |
| ENSMUSG00000025855 | Prkar1b | protein kinase, cAMP dependent regulatory, type I beta | -4.30 |
| ENSMUSG00000068617 | Efcab1 | EF-hand calcium binding domain 1 | -4.30 |
| ENSMUSG00000082211 | Defa27 | defensin, alpha, 27 | -4.30 |
| ENSMUSG00000051111 | Sv2c | synaptic vesicle glycoprotein 2c | -4.30 |
| ENSMUSG00000026817 | Ak1 | adenylate kinase 1 | -4.30 |
| ENSMUSG00000069516 | Lyz2 | lysozyme 2 | -4.31 |

| | | | |
|---|---|---|---|
| ENSMUSG00000034656 | Cacna1a | calcium channel, voltage-dependent, P/Q type, alpha 1A subunit | -4.31 |
| ENSMUSG00000062760 | Shisal1 | shisa like 1 | -4.31 |
| ENSMUSG00000038248 | Sobp | sine oculis binding protein | -4.32 |
| ENSMUSG00000038209 | Itln1 | intelectin 1 (galactofuranose binding) | -4.33 |
| ENSMUSG00000015829 | Tnr | tenascin R | -4.33 |
| ENSMUSG00000048022 | Tmem229a | transmembrane protein 229A | -4.34 |
| ENSMUSG00000056973 | Ces1d | carboxylesterase 1D | -4.34 |
| ENSMUSG00000071226 | Cecr2 | CECR2, histone acetyl-lysine reader | -4.35 |
| ENSMUSG00000103761 | Gm37859 | predicted gene, 37859 | -4.35 |
| ENSMUSG00000020717 | Pecam1 | platelet/endothelial cell adhesion molecule 1 | -4.36 |
| ENSMUSG00000046204 | Pnma2 | paraneoplastic antigen MA2 | -4.36 |
| ENSMUSG00000066176 | Gm12511 | predicted gene 12511 | -4.36 |
| ENSMUSG00000109305 | 1810010D01Rik | RIKEN cDNA 1810010D01 gene | -4.38 |
| ENSMUSG00000069814 | Ccdc92b | coiled-coil domain containing 92B | -4.39 |
| ENSMUSG00000063142 | Kcnma1 | potassium large conductance calcium-activated channel, subfamily M, alpha memb | -4.40 |
| ENSMUSG00000027797 | Dclk1 | doublecortin-like kinase 1 | -4.40 |
| ENSMUSG00000058297 | Spock2 | sparc/osteonectin, cwcv and kazal-like domains proteoglycan 2 | -4.41 |
| ENSMUSG00000029193 | Cckar | cholecystokinin A receptor | -4.41 |
| ENSMUSG00000020901 | Pik3r5 | phosphoinositide-3-kinase regulatory subunit 5 | -4.42 |
| ENSMUSG00000030263 | Lrmp | lymphoid-restricted membrane protein | -4.42 |
| ENSMUSG00000056752 | Dnah9 | dynein, axonemal, heavy chain 9 | -4.42 |
| ENSMUSG00000004707 | Ly9 | lymphocyte antigen 9 | -4.42 |
| ENSMUSG00000111443 | Gm46123 | predicted gene, 46123 | -4.42 |
| ENSMUSG00000086029 | Pax6os1 | paired box 6 opposite strand 1 | -4.42 |
| ENSMUSG00000087211 | Lhx1os | LIM homeobox 1, opposite strand | -4.42 |
| ENSMUSG00000051314 | Ffar2 | free fatty acid receptor 2 | -4.43 |
| ENSMUSG00000049154 | Fam183b | family with sequence similarity 183, member B | -4.44 |
| ENSMUSG00000015599 | Ttbk1 | tau tubulin kinase 1 | -4.44 |
| ENSMUSG00000033152 | Podxl2 | podocalyxin-like 2 | -4.44 |
| ENSMUSG00000002900 | Lamb1 | laminin B1 | -4.45 |
| ENSMUSG00000046460 | Sh2d7 | SH2 domain containing 7 | -4.45 |
| ENSMUSG00000018822 | Sfrp5 | secreted frizzled-related sequence protein 5 | -4.46 |
| ENSMUSG00000028926 | Cdk14 | cyclin-dependent kinase 14 | -4.47 |
| ENSMUSG00000040978 | Gm11992 | predicted gene 11992 | -4.47 |
| ENSMUSG00000061958 | Gm14851 | predicted gene 14851 | -4.49 |
| ENSMUSG00000025175 | Fn3k | fructosamine 3 kinase | -4.49 |
| ENSMUSG00000030020 | Prickle2 | prickle planar cell polarity protein 2 | -4.49 |
| ENSMUSG00000066705 | Fxyd6 | FXYD domain-containing ion transport regulator 6 | -4.50 |
| ENSMUSG00000004113 | Cacna1b | calcium channel, voltage-dependent, N type, alpha 1B subunit | -4.51 |
| ENSMUSG00000032297 | Celf6 | CUGBP, Elav-like family member 6 | -4.51 |
| ENSMUSG00000056492 | Adgrf5 | adhesion G protein-coupled receptor F5 | -4.52 |
| ENSMUSG00000024304 | Cdh2 | cadherin 2 | -4.52 |
| ENSMUSG00000106491 | Gm42446 | predicted gene 42446 | -4.53 |
| ENSMUSG00000050994 | Adgb | androglobin | -4.53 |
| ENSMUSG00000071178 | Serpina1b | serine (or cysteine) preptidase inhibitor, clade A, member 1B | -4.55 |
| ENSMUSG00000052572 | Dlg2 | discs large MAGUK scaffold protein 2 | -4.55 |
| ENSMUSG00000064213 | Defa24 | defensin, alpha, 24 | -4.57 |
| ENSMUSG00000052631 | Sh2d6 | SH2 domain containing 6 | -4.57 |
| ENSMUSG00000037627 | Rgs22 | regulator of G-protein signalling 22 | -4.57 |
| ENSMUSG00000029445 | Hpd | 4-hydroxyphenylpyruvic acid dioxygenase | -4.58 |
| ENSMUSG00000002265 | Peg3 | paternally expressed 3 | -4.58 |
| ENSMUSG00000009394 | Syn2 | synapsin II | -4.58 |
| ENSMUSG00000022449 | Adamts20 | a disintegrin-like and metallopeptidase (reprolysin type) with thrombospondin type | -4.58 |
| ENSMUSG00000040653 | Ppp1r14c | protein phosphatase 1, regulatory inhibitor subunit 14C | -4.58 |
| ENSMUSG00000037953 | A4gnt | alpha-1,4-N-acetylglucosaminyltransferase | -4.59 |
| ENSMUSG00000074441 | Gm15292 | predicted gene 15292 | -4.59 |
| ENSMUSG00000010066 | Cacna2d2 | calcium channel, voltage-dependent, alpha 2/delta subunit 2 | -4.59 |
| ENSMUSG00000000031 | H19 | H19, imprinted maternally expressed transcript | -4.60 |
| ENSMUSG00000023033 | Scn8a | sodium channel, voltage-gated, type VIII, alpha | -4.60 |
| ENSMUSG00000044807 | Zfp354c | zinc finger protein 354C | -4.61 |
| ENSMUSG00000032852 | Rspo4 | R-spondin 4 | -4.62 |
| ENSMUSG00000075517 | Cyp2d37-ps | cytochrome P450, family 2, subfamily d, polypeptide 37, pseudogene | -4.62 |
| ENSMUSG00000021071 | Trim9 | tripartite motif-containing 9 | -4.63 |
| ENSMUSG00000074440 | Defa3 | defensin, alpha, 3 | -4.64 |
| ENSMUSG00000000394 | Gcg | glucagon | -4.65 |
| ENSMUSG00000060070 | Defa26 | defensin, alpha, 26 | -4.66 |
| ENSMUSG00000074444 | Defa30 | defensin, alpha, 30 | -4.66 |

| | | | |
|---|---|---|---|
| ENSMUSG00000026247 | Ecel1 | endothelin converting enzyme-like 1 | -4.66 |
| ENSMUSG00000041078 | Grid1 | glutamate receptor, ionotropic, delta 1 | -4.68 |
| ENSMUSG00000074443 | Defa22 | defensin, alpha, 22 | -4.69 |
| ENSMUSG00000074437 | Defa29 | defensin, alpha, 29 | -4.70 |
| ENSMUSG00000019124 | Scrn1 | secernin 1 | -4.71 |
| ENSMUSG00000079015 | Serpina1c | serine (or cysteine) peptidase inhibitor, clade A, member 1C | -4.71 |
| ENSMUSG00000021647 | Cartpt | CART prepropeptide | -4.73 |
| ENSMUSG00000046215 | Rprml | reprimo-like | -4.73 |
| ENSMUSG00000060208 | Defa17 | defensin, alpha, 17 | -4.74 |
| ENSMUSG00000111977 | Gm47163 | predicted gene, 47163 | -4.74 |
| ENSMUSG00000094662 | Defa36 | defensin, alpha, 36 | -4.74 |
| ENSMUSG00000030683 | Sez6l2 | seizure related 6 homolog like 2 | -4.75 |
| ENSMUSG00000039809 | Gabbr2 | gamma-aminobutyric acid (GABA) B receptor, 2 | -4.75 |
| ENSMUSG00000074447 | Defa21 | defensin, alpha, 21 | -4.75 |
| ENSMUSG00000065956 | Defa37 | defensin, alpha, 37 | -4.75 |
| ENSMUSG00000097927 | Gm6999 | predicted gene 6999 | -4.76 |
| ENSMUSG00000027434 | Nkx2-2 | NK2 homeobox 2 | -4.77 |
| ENSMUSG00000028137 | Celf3 | CUGBP, Elav-like family member 3 | -4.77 |
| ENSMUSG00000040035 | Disp2 | dispatched RND tramsporter family member 2 | -4.78 |
| ENSMUSG00000024225 | Clps | colipase, pancreatic | -4.78 |
| ENSMUSG00000061911 | Myt1l | myelin transcription factor 1-like | -4.79 |
| ENSMUSG00000061845 | Defa35 | defensin, alpha, 35 | -4.80 |
| ENSMUSG00000064177 | Ghrl | ghrelin | -4.82 |
| ENSMUSG00000027254 | Map1a | microtubule-associated protein 1 A | -4.82 |
| ENSMUSG00000036913 | Trim67 | tripartite motif-containing 67 | -4.82 |
| ENSMUSG00000056069 | Otulinl | OTU deubiquitinase with linear linkage specificity like | -4.83 |
| ENSMUSG00000020787 | P2rx1 | purinergic receptor P2X, ligand-gated ion channel, 1 | -4.83 |
| ENSMUSG00000004347 | Pde1c | phosphodiesterase 1C | -4.84 |
| ENSMUSG00000063206 | Defa34 | defensin, alpha, 34 | -4.84 |
| ENSMUSG00000029005 | Draxin | dorsal inhibitory axon guidance protein | -4.84 |
| ENSMUSG00000079056 | Kcnip3 | Kv channel interacting protein 3, calsenilin | -4.85 |
| ENSMUSG00000050824 | Sstr5 | somatostatin receptor 5 | -4.85 |
| ENSMUSG00000031144 | Syp | synaptophysin | -4.86 |
| ENSMUSG00000058618 | AY761184 | cDNA sequence AY761184 | -4.86 |
| ENSMUSG00000020701 | Tmem132e | transmembrane protein 132E | -4.88 |
| ENSMUSG00000025964 | Adam23 | a disintegrin and metallopeptidase domain 23 | -4.88 |
| ENSMUSG00000025020 | Slit1 | slit guidance ligand 1 | -4.89 |
| ENSMUSG00000025089 | Gfra1 | glial cell line derived neurotrophic factor family receptor alpha 1 | -4.89 |
| ENSMUSG00000037428 | Vgf | VGF nerve growth factor inducible | -4.90 |
| ENSMUSG00000021763 | BC067074 | cDNA sequence BC067074 | -4.90 |
| ENSMUSG00000096295 | Defa2 | defensin, alpha, 2 | -4.91 |
| ENSMUSG00000064080 | Fbln2 | fibulin 2 | -4.91 |
| ENSMUSG00000031284 | Pak3 | p21 (RAC1) activated kinase 3 | -4.91 |
| ENSMUSG00000004110 | Cacna1e | calcium channel, voltage-dependent, R type, alpha 1E subunit | -4.91 |
| ENSMUSG00000051331 | Cacna1c | calcium channel, voltage-dependent, L type, alpha 1C subunit | -4.92 |
| ENSMUSG00000020848 | Doc2b | double C2, beta | -4.92 |
| ENSMUSG00000032769 | Trpa1 | transient receptor potential cation channel, subfamily A, member 1 | -4.92 |
| ENSMUSG00000026833 | Olfm1 | olfactomedin 1 | -4.93 |
| ENSMUSG00000043301 | Kcnj6 | potassium inwardly-rectifying channel, subfamily J, member 6 | -4.93 |
| ENSMUSG00000019796 | Lrp11 | low density lipoprotein receptor-related protein 11 | -4.95 |
| ENSMUSG00000038224 | Serpinf2 | serine (or cysteine) peptidase inhibitor, clade F, member 2 | -4.95 |
| ENSMUSG00000051079 | Rgs13 | regulator of G-protein signaling 13 | -4.97 |
| ENSMUSG00000044365 | Cxxc4 | CXXC finger 4 | -4.98 |
| ENSMUSG00000041633 | Kctd12b | potassium channel tetramerisation domain containing 12b | -4.99 |
| ENSMUSG00000099907 | Gm10421 | predicted gene 10421 | -5.00 |
| ENSMUSG00000037852 | Cpe | carboxypeptidase E | -5.02 |
| ENSMUSG00000083282 | Ctsf | cathepsin F | -5.02 |
| ENSMUSG00000033061 | Resp18 | regulated endocrine-specific protein 18 | -5.02 |
| ENSMUSG00000035967 | Ints6l | integrator complex subunit 6 like | -5.02 |
| ENSMUSG00000058248 | Kcnh1 | potassium voltage-gated channel, subfamily H (eag-related), member 1 | -5.02 |
| ENSMUSG00000020734 | Grin2c | glutamate receptor, ionotropic, NMDA2C (epsilon 3) | -5.02 |
| ENSMUSG00000075012 | Fjx1 | four jointed box 1 | -5.03 |
| ENSMUSG00000020090 | Npffr1 | neuropeptide FF receptor 1 | -5.04 |
| ENSMUSG00000068154 | Insm1 | insulinoma-associated 1 | -5.05 |
| ENSMUSG00000020723 | Cacng4 | calcium channel, voltage-dependent, gamma subunit 4 | -5.06 |
| ENSMUSG00000047507 | Baiap3 | BAI1-associated protein 3 | -5.07 |
| ENSMUSG00000028008 | Asic5 | acid-sensing (proton-gated) ion channel family member 5 | -5.09 |

| | | | |
|---|---|---|---|
| ENSMUSG00000020333 | Acsl6 | acyl-CoA synthetase long-chain family member 6 | -5.10 |
| ENSMUSG00000091345 | Col6a5 | collagen, type VI, alpha 5 | -5.10 |
| ENSMUSG00000105271 | Gm42875 | predicted gene 42875 | -5.10 |
| ENSMUSG00000049605 | Olfr418 | olfactory receptor 418 | -5.11 |
| ENSMUSG00000028339 | Col15a1 | collagen, type XV, alpha 1 | -5.13 |
| ENSMUSG00000106320 | Gm42992 | predicted gene 42992 | -5.13 |
| ENSMUSG00000008932 | Slc1a7 | solute carrier family 1 (glutamate transporter), member 7 | -5.13 |
| ENSMUSG00000016346 | Kcnq2 | potassium voltage-gated channel, subfamily Q, member 2 | -5.13 |
| ENSMUSG00000006546 | Cryba2 | crystallin, beta A2 | -5.14 |
| ENSMUSG00000062380 | Tubb3 | tubulin, beta 3 class III | -5.15 |
| ENSMUSG00000020811 | Wscd1 | WSC domain containing 1 | -5.15 |
| ENSMUSG00000051980 | Casr | calcium-sensing receptor | -5.15 |
| ENSMUSG00000020886 | Dlg4 | discs large MAGUK scaffold protein 4 | -5.16 |
| ENSMUSG00000057182 | Scn3a | sodium channel, voltage-gated, type III, alpha | -5.17 |
| ENSMUSG00000090291 | Lrrc10b | leucine rich repeat containing 10B | -5.17 |
| ENSMUSG00000017311 | Pyy | peptide YY | -5.17 |
| ENSMUSG00000015401 | Cltrn | collectrin, amino acid transport regulator | -5.18 |
| ENSMUSG00000039037 | St6galnac5 | ST6 (alpha-N-acetyl-neuraminyl-2,3-beta-galactosyl-1,3)-N-acetylgalactosaminide a | -5.19 |
| ENSMUSG00000036766 | Dner | delta/notch-like EGF repeat containing | -5.19 |
| ENSMUSG00000017943 | Gdap1l1 | ganglioside-induced differentiation-associated protein 1-like 1 | -5.19 |
| ENSMUSG00000047798 | Cd300lf | CD300 molecule like family member F | -5.19 |
| ENSMUSG00000005640 | Insrr | insulin receptor-related receptor | -5.19 |
| ENSMUSG00000105985 | Gm42993 | predicted gene 42993 | -5.20 |
| ENSMUSG00000104886 | Gm43000 | predicted gene 43000 | -5.20 |
| ENSMUSG00000055567 | Unc80 | unc-80, NALCN activator | -5.21 |
| ENSMUSG00000070577 | Gm572 | predicted gene 572 | -5.21 |
| ENSMUSG00000040276 | Pacsin1 | protein kinase C and casein kinase substrate in neurons 1 | -5.22 |
| ENSMUSG00000020703 | 5530401A14Rik | RIKEN cDNA 5530401A14 gene | -5.22 |
| ENSMUSG00000027834 | Serpini1 | serine (or cysteine) peptidase inhibitor, clade I, member 1 | -5.23 |
| ENSMUSG00000027876 | Reg4 | regenerating islet-derived family, member 4 | -5.23 |
| ENSMUSG00000095328 | Defa-ps6 | defensin, alpha, pseudogene 6 | -5.23 |
| ENSMUSG00000114624 | Gm48267 | predicted gene, 48267 | -5.23 |
| ENSMUSG00000037624 | Kcnk2 | potassium channel, subfamily K, member 2 | -5.23 |
| ENSMUSG00000079180 | Mptx2 | mucosal pentraxin 2 | -5.25 |
| ENSMUSG00000044359 | P2ry4 | pyrimidinergic receptor P2Y, G-protein coupled, 4 | -5.25 |
| ENSMUSG00000050751 | Pgbd5 | piggyBac transposable element derived 5 | -5.25 |
| ENSMUSG00000048148 | Nwd1 | NACHT and WD repeat domain containing 1 | -5.26 |
| ENSMUSG00000058975 | Kcnc1 | potassium voltage gated channel, Shaw-related subfamily, member 1 | -5.27 |
| ENSMUSG00000040118 | Cacna2d1 | calcium channel, voltage-dependent, alpha2/delta subunit 1 | -5.28 |
| ENSMUSG00000032338 | Hcn4 | hyperpolarization-activated, cyclic nucleotide-gated K+ 4 | -5.28 |
| ENSMUSG00000106048 | Gm42444 | predicted gene 42444 | -5.28 |
| ENSMUSG00000010505 | Myt1 | myelin transcription factor 1 | -5.29 |
| ENSMUSG00000028469 | Npr2 | natriuretic peptide receptor 2 | -5.29 |
| ENSMUSG00000025810 | Nrp1 | neuropilin 1 | -5.30 |
| ENSMUSG00000115190 | Gm49542 | predicted gene, 49542 | -5.30 |
| ENSMUSG00000023224 | Serping1 | serine (or cysteine) peptidase inhibitor, clade G, member 1 | -5.32 |
| ENSMUSG00000052727 | Map1b | microtubule-associated protein 1B | -5.32 |
| ENSMUSG00000031740 | Mmp2 | matrix metallopeptidase 2 | -5.32 |
| ENSMUSG00000035431 | Sstr1 | somatostatin receptor 1 | -5.35 |
| ENSMUSG00000114245 | Gm38655 | predicted gene, 38655 | -5.35 |
| ENSMUSG00000043004 | Gng2 | guanine nucleotide binding protein (G protein), gamma 2 | -5.36 |
| ENSMUSG00000040009 | Gnaz | guanine nucleotide binding protein, alpha z subunit | -5.36 |
| ENSMUSG00000006586 | Runx1t1 | runt-related transcription factor 1; translocated to, 1 (cyclin D-related) | -5.36 |
| ENSMUSG00000031837 | Necab2 | N-terminal EF-hand calcium binding protein 2 | -5.36 |
| ENSMUSG00000028360 | Slc44a5 | solute carrier family 44, member 5 | -5.37 |
| ENSMUSG00000038456 | Dennd2a | DENN/MADD domain containing 2A | -5.38 |
| ENSMUSG00000024682 | Gif | gastric intrinsic factor | -5.39 |
| ENSMUSG00000075256 | Cerkl | ceramide kinase-like | -5.40 |
| ENSMUSG00000032394 | Igdcc3 | immunoglobulin superfamily, DCC subclass, member 3 | -5.40 |
| ENSMUSG00000046613 | Vwa5b2 | von Willebrand factor A domain containing 5B2 | -5.41 |
| ENSMUSG00000026872 | Zeb2 | zinc finger E-box binding homeobox 2 | -5.41 |
| ENSMUSG00000021303 | Gng4 | guanine nucleotide binding protein (G protein), gamma 4 | -5.43 |
| ENSMUSG00000031099 | Smarca1 | SWI/SNF related, matrix associated, actin dependent regulator of chromatin, subfa | -5.43 |
| ENSMUSG00000096847 | Tmem151b | transmembrane protein 151B | -5.44 |
| ENSMUSG00000046743 | Fat4 | FAT atypical cadherin 4 | -5.47 |
| ENSMUSG00000048038 | Ccdc187 | coiled-coil domain containing 187 | -5.47 |
| ENSMUSG00000097767 | Miat | myocardial infarction associated transcript (non-protein coding) | -5.48 |

| | | | |
|---|---|---|---|
| ENSMUSG00000105526 | Gm43490 | predicted gene 43490 | -5.48 |
| ENSMUSG00000054667 | Irs4 | insulin receptor substrate 4 | -5.48 |
| ENSMUSG00000028356 | Ambp | alpha 1 microglobulin/bikunin | -5.49 |
| ENSMUSG00000050473 | Slc35d3 | solute carrier family 35, member D3 | -5.49 |
| ENSMUSG00000058740 | Kcnt1 | potassium channel, subfamily T, member 1 | -5.50 |
| ENSMUSG00000051985 | Igfn1 | immunoglobulin-like and fibronectin type III domain containing 1 | -5.50 |
| ENSMUSG00000104301 | Wdr49 | WD repeat domain 49 | -5.51 |
| ENSMUSG00000092083 | Kcnb2 | potassium voltage gated channel, Shab-related subfamily, member 2 | -5.52 |
| ENSMUSG00000029706 | Pax4 | paired box 4 | -5.53 |
| ENSMUSG00000026443 | Lrrn2 | leucine rich repeat protein 2, neuronal | -5.53 |
| ENSMUSG00000027009 | Itga4 | integrin alpha 4 | -5.54 |
| ENSMUSG00000042750 | Bex2 | brain expressed X-linked 2 | -5.55 |
| ENSMUSG00000100182 | 1810006J02Rik | RIKEN cDNA 1810006J02 gene | -5.58 |
| ENSMUSG00000026442 | Nfasc | neurofascin | -5.59 |
| ENSMUSG00000038530 | Rgs4 | regulator of G-protein signaling 4 | -5.59 |
| ENSMUSG00000040046 | Tph1 | tryptophan hydroxylase 1 | -5.59 |
| ENSMUSG00000034145 | Tmem63c | transmembrane protein 63c | -5.61 |
| ENSMUSG00000001901 | Kcnh6 | potassium voltage-gated channel, subfamily H (eag-related), member 6 | -5.62 |
| ENSMUSG00000039323 | Igfbp2 | insulin-like growth factor binding protein 2 | -5.62 |
| ENSMUSG00000043857 | Mgat5b | mannoside acetylglucosaminyltransferase 5, isoenzyme B | -5.62 |
| ENSMUSG00000104149 | Gm37138 | predicted gene, 37138 | -5.63 |
| ENSMUSG00000027360 | Hdc | histidine decarboxylase | -5.67 |
| ENSMUSG00000026163 | Sphkap | SPHK1 interactor, AKAP domain containing | -5.69 |
| ENSMUSG00000090071 | Cdk5r2 | cyclin-dependent kinase 5, regulatory subunit 2 (p39) | -5.70 |
| ENSMUSG00000035277 | Arx | aristaless related homeobox | -5.73 |
| ENSMUSG00000039059 | Hrh3 | histamine receptor H3 | -5.73 |
| ENSMUSG00000041798 | Gck | glucokinase | -5.74 |
| ENSMUSG00000026432 | Avpr1b | arginine vasopressin receptor 1B | -5.74 |
| ENSMUSG00000024366 | Gfra3 | glial cell line derived neurotrophic factor family receptor alpha 3 | -5.75 |
| ENSMUSG00000033676 | Gabrb3 | gamma-aminobutyric acid (GABA) A receptor, subunit beta 3 | -5.75 |
| ENSMUSG00000086067 | Gm16183 | predicted gene 16183 | -5.76 |
| ENSMUSG00000034701 | Neurod1 | neurogenic differentiation 1 | -5.77 |
| ENSMUSG00000070802 | Pnmal2 | PNMA-like 2 | -5.77 |
| ENSMUSG00000095813 | Defa-ps1 | defensin, alpha, pseudogene 1 | -5.78 |
| ENSMUSG00000021536 | Adcy2 | adenylate cyclase 2 | -5.78 |
| ENSMUSG00000061171 | Slc38a11 | solute carrier family 38, member 11 | -5.79 |
| ENSMUSG00000094626 | Tmem121b | transmembrane protein 121B | -5.79 |
| ENSMUSG00000061762 | Tac1 | tachykinin 1 | -5.80 |
| ENSMUSG00000024990 | Rbp4 | retinol binding protein 4, plasma | -5.81 |
| ENSMUSG00000021194 | Chga | chromogranin A | -5.82 |
| ENSMUSG00000032181 | Scg3 | secretogranin III | -5.82 |
| ENSMUSG00000053166 | Cdh22 | cadherin 22 | -5.82 |
| ENSMUSG00000028546 | Elavl4 | ELAV like RNA binding protein 4 | -5.83 |
| ENSMUSG00000047842 | Diras2 | DIRAS family, GTP-binding RAS-like 2 | -5.83 |
| ENSMUSG00000101814 | Gm17807 | predicted gene, 17807 | -5.83 |
| ENSMUSG00000024112 | Cacna1h | calcium channel, voltage-dependent, T type, alpha 1H subunit | -5.87 |
| ENSMUSG00000056553 | Ptprn2 | protein tyrosine phosphatase, receptor type, N polypeptide 2 | -5.88 |
| ENSMUSG00000054640 | Slc8a1 | solute carrier family 8 (sodium/calcium exchanger), member 1 | -5.88 |
| ENSMUSG00000020177 | 9530003J23Rik | RIKEN cDNA 9530003J23 gene | -5.88 |
| ENSMUSG00000027350 | Chgb | chromogranin B | -5.92 |
| ENSMUSG00000010021 | Kif19a | kinesin family member 19A | -5.93 |
| ENSMUSG00000033740 | St18 | suppression of tumorigenicity 18 | -5.94 |
| ENSMUSG00000026204 | Ptprn | protein tyrosine phosphatase, receptor type, N | -5.96 |
| ENSMUSG00000066456 | Hmgn3 | high mobility group nucleosomal binding domain 3 | -5.96 |
| ENSMUSG00000042631 | Xkr7 | X-linked Kx blood group related 7 | -5.96 |
| ENSMUSG00000039860 | Srrm3 | serine/arginine repetitive matrix 3 | -5.97 |
| ENSMUSG00000043496 | Tril | TLR4 interactor with leucine-rich repeats | -5.97 |
| ENSMUSG00000110557 | Gm5159 | predicted gene 5159 | -5.97 |
| ENSMUSG00000025375 | Aatk | apoptosis-associated tyrosine kinase | -5.99 |
| ENSMUSG00000021700 | Rab3c | RAB3C, member RAS oncogene family | -6.02 |
| ENSMUSG00000029420 | Rimbp2 | RIMS binding protein 2 | -6.03 |
| ENSMUSG00000072849 | Serpina1e | serine (or cysteine) peptidase inhibitor, clade A, member 1E | -6.03 |
| ENSMUSG00000033981 | Gria2 | glutamate receptor, ionotropic, AMPA2 (alpha 2) | -6.04 |
| ENSMUSG00000021904 | Sema3g | sema domain, immunoglobulin domain (Ig), short basic domain, secreted, (semaph | -6.07 |
| ENSMUSG00000004366 | Sst | somatostatin | -6.08 |
| ENSMUSG00000071265 | 1700086L19Rik | RIKEN cDNA 1700086L19 gene | -6.10 |
| ENSMUSG00000019900 | Rfx6 | regulatory factor X, 6 | -6.12 |

| | | | |
|---|---|---|---|
| ENSMUSG00000027168 | Pax6 | paired box 6 | -6.12 |
| ENSMUSG00000039683 | Sdk1 | sidekick cell adhesion molecule 1 | -6.13 |
| ENSMUSG00000101930 | Gm5441 | predicted gene 5441 | -6.13 |
| ENSMUSG00000053545 | 6430503K07Rik | RIKEN cDNA 6430503K07 gene | -6.16 |
| ENSMUSG00000079116 | Gm15293 | predicted gene 15293 | -6.18 |
| ENSMUSG00000019876 | Pkib | protein kinase inhibitor beta, cAMP dependent, testis specific | -6.18 |
| ENSMUSG00000044453 | Ffar1 | free fatty acid receptor 1 | -6.21 |
| ENSMUSG00000039278 | Pcsk1n | proprotein convertase subtilisin/kexin type 1 inhibitor | -6.23 |
| ENSMUSG00000040972 | Igsf21 | immunoglobulin superfamily, member 21 | -6.23 |
| ENSMUSG00000026824 | Kcnj3 | potassium inwardly-rectifying channel, subfamily J, member 3 | -6.24 |
| ENSMUSG00000027220 | Syt13 | synaptotagmin XIII | -6.25 |
| ENSMUSG00000023236 | Scg5 | secretogranin V | -6.28 |
| ENSMUSG00000061601 | Pclo | piccolo (presynaptic cytomatrix protein) | -6.30 |
| ENSMUSG00000050556 | Kcnb1 | potassium voltage gated channel, Shab-related subfamily, member 1 | -6.30 |
| ENSMUSG00000021198 | Unc79 | unc-79 homolog | -6.30 |
| ENSMUSG00000048483 | Zdhhc22 | zinc finger, DHHC-type containing 22 | -6.30 |
| ENSMUSG00000038255 | Neurod2 | neurogenic differentiation 2 | -6.37 |
| ENSMUSG00000068263 | Efcc1 | EF hand and coiled-coil domain containing 1 | -6.38 |
| ENSMUSG00000048215 | A630023P12Rik | RIKEN cDNA A630023P12 gene | -6.38 |
| ENSMUSG00000026686 | Lmx1a | LIM homeobox transcription factor 1 alpha | -6.40 |
| ENSMUSG00000023387 | Kcnk16 | potassium channel, subfamily K, member 16 | -6.43 |
| ENSMUSG00000048218 | Amigo2 | adhesion molecule with Ig like domain 2 | -6.43 |
| ENSMUSG00000054423 | Cadps | Ca2+-dependent secretion activator | -6.45 |
| ENSMUSG00000050711 | Scg2 | secretogranin II | -6.48 |
| ENSMUSG00000037032 | Apbb1 | amyloid beta (A4) precursor protein-binding, family B, member 1 | -6.48 |
| ENSMUSG00000114230 | Gm48239 | predicted gene, 48239 | -6.48 |
| ENSMUSG00000052951 | C130021I20Rik | Riken cDNA C130021I20 gene | -6.49 |
| ENSMUSG00000106139 | Gm30648 | predicted gene, 30648 | -6.51 |
| ENSMUSG00000042258 | Isl1 | ISL1 transcription factor, LIM/homeodomain | -6.57 |
| ENSMUSG00000014351 | Gip | gastric inhibitory polypeptide | -6.58 |
| ENSMUSG00000032532 | Cck | cholecystokinin | -6.58 |
| ENSMUSG00000029878 | Dbpht2 | DNA binding protein with his-thr domain | -6.59 |
| ENSMUSG00000071719 | Tmem28 | transmembrane protein 28 | -6.59 |
| ENSMUSG00000025738 | Fbxl16 | F-box and leucine-rich repeat protein 16 | -6.70 |
| ENSMUSG00000102937 | Gm38116 | predicted gene, 38116 | -6.72 |
| ENSMUSG00000021337 | Scgn | secretagogin, EF-hand calcium binding protein | -6.74 |
| ENSMUSG00000069072 | Slc7a14 | solute carrier family 7 (cationic amino acid transporter, y+ system), member 14 | -6.80 |
| ENSMUSG00000073530 | Pappa2 | pappalysin 2 | -6.88 |
| ENSMUSG00000029121 | Crmp1 | collapsin response mediator protein 1 | -6.94 |
| ENSMUSG00000024553 | Galr1 | galanin receptor 1 | -6.94 |
| ENSMUSG00000055409 | Nell1 | NEL-like 1 | -7.00 |
| ENSMUSG00000028226 | Mmp16 | matrix metallopeptidase 16 | -7.10 |
| ENSMUSG00000055197 | Fev | FEV (ETS oncogene family) | -7.25 |
| ENSMUSG00000009292 | Trpm2 | transient receptor potential cation channel, subfamily M, member 2 | -7.40 |
| ENSMUSG00000022762 | Ncam2 | neural cell adhesion molecule 2 | -7.43 |
| ENSMUSG00000039372 | 38047 | membrane-associated ring finger (C3HC4) 4 | -7.51 |
| ENSMUSG00000041141 | Pnmal1 | PNMA-like 1 | -7.61 |
| ENSMUSG00000018698 | Lhx1 | LIM homeobox protein 1 | -7.68 |
| ENSMUSG00000030359 | Pzp | PZP, alpha-2-macroglobulin like | -7.76 |
| ENSMUSG00000053852 | Adgrg4 | adhesion G protein-coupled receptor G4 | -7.97 |
| ENSMUSG00000038765 | Lmx1b | LIM homeobox transcription factor 1 beta | -8.47 |
| ENSMUSG00000022619 | Mapk8ip2 | mitogen-activated protein kinase 8 interacting protein 2 | -9.32 |

**RFP+Trop2- vs. differentiated cells**

| Ensemble gene code | Symbol | Gene name | logFC |
|---|---|---|---|
| ENSMUSG00000097519 | 4930558J18Rik | RIKEN cDNA 4930558J18 gene | 2.88 |
| ENSMUSG00000082095 | Gm11991 | predicted gene 11991 | 2.55 |
| ENSMUSG00000082706 | Gm11663 | predicted gene 11663 | 2.35 |
| ENSMUSG00000037124 | Trim58 | tripartite motif-containing 58 | 2.33 |
| ENSMUSG00000033952 | Aspm | abnormal spindle microtubule assembly | 2.26 |
| ENSMUSG00000032783 | Troap | trophinin associated protein | 2.26 |
| ENSMUSG00000041498 | Kif14 | kinesin family member 14 | 2.25 |
| ENSMUSG00000044201 | Cdc25c | cell division cycle 25C | 2.21 |
| ENSMUSG00000061991 | Hist1h2af | histone cluster 1, H2af | 2.20 |
| ENSMUSG00000037628 | Cdkn3 | cyclin-dependent kinase inhibitor 3 | 2.19 |
| ENSMUSG00000108528 | Gm8319 | predicted gene 8319 | 2.19 |
| ENSMUSG00000023505 | Cdca3 | cell division cycle associated 3 | 2.17 |
| ENSMUSG00000028678 | Kif2c | kinesin family member 2C | 2.17 |
| ENSMUSG00000027715 | Ccna2 | cyclin A2 | 2.16 |
| ENSMUSG00000102359 | Gm38351 | predicted gene, 38351 | 2.16 |
| ENSMUSG00000028068 | Iqgap3 | IQ motif containing GTPase activating protein 3 | 2.15 |
| ENSMUSG00000035365 | Parpbp | PARP1 binding protein | 2.13 |
| ENSMUSG00000017716 | Birc5 | baculoviral IAP repeat-containing 5 | 2.11 |
| ENSMUSG00000026605 | Cenpf | centromere protein F | 2.11 |
| ENSMUSG00000026622 | Nek2 | NIMA (never in mitosis gene a)-related expressed kinase 2 | 2.11 |
| ENSMUSG00000026683 | Nuf2 | NUF2, NDC80 kinetochore complex component | 2.11 |
| ENSMUSG00000041431 | Ccnb1 | cyclin B1 | 2.11 |
| ENSMUSG00000006398 | Cdc20 | cell division cycle 20 | 2.10 |
| ENSMUSG00000039814 | Xkr5 | X-linked Kx blood group related 5 | 2.10 |
| ENSMUSG00000003779 | Kif20a | kinesin family member 20A | 2.08 |
| ENSMUSG00000051378 | Kif18b | kinesin family member 18B | 2.08 |
| ENSMUSG00000020330 | Hmmr | hyaluronan mediated motility receptor (RHAMM) | 2.07 |
| ENSMUSG00000045328 | Cenpe | centromere protein E | 2.07 |
| ENSMUSG00000041064 | Pif1 | PIF1 5'-to-3' DNA helicase | 2.07 |
| ENSMUSG00000032218 | Ccnb2 | cyclin B2 | 2.06 |
| ENSMUSG00000058290 | Espl1 | extra spindle pole bodies 1, separase | 2.06 |
| ENSMUSG00000079553 | Kifc1 | kinesin family member C1 | 2.06 |
| ENSMUSG00000027469 | Tpx2 | TPX2, microtubule-associated | 2.05 |
| ENSMUSG00000027379 | Bub1 | BUB1, mitotic checkpoint serine/threonine kinase | 2.04 |
| ENSMUSG00000114456 | Hist1h2bh | histone cluster 1, H2bh | 2.04 |
| ENSMUSG00000029177 | Cenpa | centromere protein A | 2.02 |
| ENSMUSG00000001403 | Ube2c | ubiquitin-conjugating enzyme E2C | 2.01 |
| ENSMUSG00000031262 | Cenpi | centromere protein I | 2.01 |
| ENSMUSG00000094248 | Hist1h2ao | histone cluster 1, H2ao | 2.01 |
| ENSMUSG00000113168 | Gm7614 | predicted gene 7614 | 2.00 |
| ENSMUSG00000027115 | Kif18a | kinesin family member 18A | 1.99 |
| ENSMUSG00000098985 | Gm27219 | predicted gene 27219 | 1.99 |
| ENSMUSG00000028175 | Depdc1a | DEP domain containing 1a | 1.98 |
| ENSMUSG00000027699 | Ect2 | ect2 oncogene | 1.97 |
| ENSMUSG00000027331 | Knstrn | kinetochore-localized astrin/SPAG5 binding | 1.97 |
| ENSMUSG00000037725 | Ckap2 | cytoskeleton associated protein 2 | 1.97 |
| ENSMUSG00000024795 | Kif20b | kinesin family member 20B | 1.97 |
| ENSMUSG00000012443 | Kif11 | kinesin family member 11 | 1.95 |
| ENSMUSG00000022322 | Shcbp1 | Shc SH2-domain binding protein 1 | 1.94 |
| ENSMUSG00000072082 | Ccnf | cyclin F | 1.94 |
| ENSMUSG00000027326 | Knl1 | kinetochore scaffold 1 | 1.92 |
| ENSMUSG00000019942 | Cdk1 | cyclin-dependent kinase 1 | 1.92 |
| ENSMUSG00000023015 | Racgap1 | Rac GTPase-activating protein 1 | 1.92 |
| ENSMUSG00000088956 | Gm24945 | predicted gene, 24945 | 1.92 |
| ENSMUSG00000027496 | Aurka | aurora kinase A | 1.91 |
| ENSMUSG00000021176 | Efcab11 | EF-hand calcium binding domain 11 | 1.91 |
| ENSMUSG00000034311 | Kif4 | kinesin family member 4 | 1.90 |
| ENSMUSG00000027811 | 4930579G24Rik | RIKEN cDNA 4930579G24 gene | 1.90 |
| ENSMUSG00000020914 | Top2a | topoisomerase (DNA) II alpha | 1.89 |
| ENSMUSG00000069910 | Spdl1 | spindle apparatus coiled-coil protein 1 | 1.89 |
| ENSMUSG00000099583 | Hist1h3d | histone cluster 1, H3d | 1.89 |
| ENSMUSG00000021965 | Ska3 | spindle and kinetochore associated complex subunit 3 | 1.88 |
| ENSMUSG00000023940 | Sgo1 | shugoshin 1 | 1.88 |
| ENSMUSG00000030677 | Kif22 | kinesin family member 22 | 1.87 |
| ENSMUSG00000048327 | Ckap2l | cytoskeleton associated protein 2-like | 1.87 |
| ENSMUSG00000069300 | Hist1h2bj | histone cluster 1, H2bj | 1.87 |
| ENSMUSG00000024056 | Ndc80 | NDC80 kinetochore complex component | 1.87 |
| ENSMUSG00000032254 | Kif23 | kinesin family member 23 | 1.86 |
| ENSMUSG00000037544 | Dlgap5 | DLG associated protein 5 | 1.86 |
| ENSMUSG00000035683 | Melk | maternal embryonic leucine zipper kinase | 1.86 |

| | | | |
|---|---|---|---|
| ENSMUSG00000099517 | Hist1h3g | histone cluster 1, H3g | 1.86 |
| ENSMUSG00000033031 | Cip2a | cell proliferation regulating inhibitor of protein phosphatase 2A | 1.85 |
| ENSMUSG00000114279 | Hist1h2bm | histone cluster 1, H2bm | 1.85 |
| ENSMUSG00000032221 | Mns1 | meiosis-specific nuclear structural protein 1 | 1.85 |
| ENSMUSG00000055653 | Gpc3 | glypican 3 | 1.85 |
| ENSMUSG00000028873 | Cdca8 | cell division cycle associated 8 | 1.84 |
| ENSMUSG00000069267 | Hist1h3b | histone cluster 1, H3b | 1.84 |
| ENSMUSG00000040084 | Bub1b | BUB1B, mitotic checkpoint serine/threonine kinase | 1.84 |
| ENSMUSG00000026955 | Sapcd2 | suppressor APC domain containing 2 | 1.83 |
| ENSMUSG00000024989 | Cep55 | centrosomal protein 55 | 1.82 |
| ENSMUSG00000037313 | Tacc3 | transforming, acidic coiled-coil containing protein 3 | 1.82 |
| ENSMUSG00000038252 | Ncapd2 | non-SMC condensin I complex, subunit D2 | 1.82 |
| ENSMUSG00000101355 | Hist1h3h | histone cluster 1, H3h | 1.82 |
| ENSMUSG00000031004 | Mki67 | antigen identified by monoclonal antibody Ki 67 | 1.81 |
| ENSMUSG00000022033 | Pbk | PDZ binding kinase | 1.81 |
| ENSMUSG00000036768 | Kif15 | kinesin family member 15 | 1.81 |
| ENSMUSG00000020897 | Aurkb | aurora kinase B | 1.80 |
| ENSMUSG00000020493 | Prr11 | proline rich 11 | 1.79 |
| ENSMUSG00000024087 | Cyp1b1 | cytochrome P450, family 1, subfamily b, polypeptide 1 | 1.79 |
| ENSMUSG00000058773 | Hist1h1b | histone cluster 1, H1b | 1.78 |
| ENSMUSG00000015880 | Ncapg | non-SMC condensin I complex, subunit G | 1.78 |
| ENSMUSG00000075031 | Hist1h2bb | histone cluster 1, H2bb | 1.78 |
| ENSMUSG00000069274 | Hist1h4f | histone cluster 1, H4f | 1.77 |
| ENSMUSG00000069310 | Hist1h3c | histone cluster 1, H3c | 1.77 |
| ENSMUSG00000062727 | Hist1h2bk | histone cluster 1, H2bk | 1.76 |
| ENSMUSG00000038379 | Ttk | Ttk protein kinase | 1.76 |
| ENSMUSG00000046295 | Ankle1 | ankyrin repeat and LEM domain containing 1 | 1.76 |
| ENSMUSG00000044966 | Fbxo48 | F-box protein 48 | 1.76 |
| ENSMUSG00000082605 | Gm16148 | predicted gene 16148 | 1.76 |
| ENSMUSG00000048922 | Cdca2 | cell division cycle associated 2 | 1.75 |
| ENSMUSG00000027306 | Nusap1 | nucleolar and spindle associated protein 1 | 1.74 |
| ENSMUSG00000034906 | Ncaph | non-SMC condensin I complex, subunit H | 1.74 |
| ENSMUSG00000095217 | Hist1h2bn | histone cluster 1, H2bn | 1.74 |
| ENSMUSG00000028312 | Smc2 | structural maintenance of chromosomes 2 | 1.74 |
| ENSMUSG00000031756 | Cenpn | centromere protein N | 1.74 |
| ENSMUSG00000038943 | Prc1 | protein regulator of cytokinesis 1 | 1.73 |
| ENSMUSG00000024301 | Kifc5b | kinesin family member C5B | 1.73 |
| ENSMUSG00000085627 | Gm11222 | predicted gene 11222 | 1.73 |
| ENSMUSG00000051517 | Arhgef39 | Rho guanine nucleotide exchange factor (GEF) 39 | 1.72 |
| ENSMUSG00000017969 | Ptgis | prostaglandin I2 (prostacyclin) synthase | 1.71 |
| ENSMUSG00000001517 | Foxm1 | forkhead box M1 | 1.70 |
| ENSMUSG00000022385 | Gtse1 | G two S phase expressed protein 1 | 1.70 |
| ENSMUSG00000064288 | Hist1h4k | histone cluster 1, H4k | 1.70 |
| ENSMUSG00000021569 | Trip13 | thyroid hormone receptor interactor 13 | 1.70 |
| ENSMUSG00000027323 | Rad51 | RAD51 recombinase | 1.70 |
| ENSMUSG00000051220 | Ercc6l | excision repair cross-complementing rodent repair deficiency complementation group 6 like | 1.70 |
| ENSMUSG00000034883 | Lrr1 | leucine rich repeat protein 1 | 1.70 |
| ENSMUSG00000094338 | Hist1h2bl | histone cluster 1, H2bl | 1.70 |
| ENSMUSG00000047534 | Mis18bp1 | MIS18 binding protein 1 | 1.68 |
| ENSMUSG00000049916 | 2610318N02Rik | RIKEN cDNA 2610318N02 gene | 1.68 |
| ENSMUSG00000022021 | Diaph3 | diaphanous related formin 3 | 1.67 |
| ENSMUSG00000036223 | Ska1 | spindle and kinetochore associated complex subunit 1 | 1.67 |
| ENSMUSG00000067818 | Myl9 | myosin, light polypeptide 9, regulatory | 1.67 |
| ENSMUSG00000022034 | Esco2 | establishment of sister chromatid cohesion N-acetyltransferase 2 | 1.66 |
| ENSMUSG00000002055 | Spag5 | sperm associated antigen 5 | 1.66 |
| ENSMUSG00000078773 | Rad54b | RAD54 homolog B (S. cerevisiae) | 1.66 |
| ENSMUSG00000024660 | Incenp | inner centromere protein | 1.65 |
| ENSMUSG00000040204 | Pclaf | PCNA clamp associated factor | 1.65 |
| ENSMUSG00000036777 | Anln | anillin, actin binding protein | 1.65 |
| ENSMUSG00000058385 | Hist1h2bg | histone cluster 1, H2bg | 1.64 |
| ENSMUSG00000029910 | Mad2l1 | MAD2 mitotic arrest deficient-like 1 | 1.64 |
| ENSMUSG00000005233 | Spc25 | SPC25, NDC80 kinetochore complex component, homolog (S. cerevisiae) | 1.63 |
| ENSMUSG00000026039 | Sgo2a | shugoshin 2A | 1.63 |
| ENSMUSG00000051235 | Gen1 | GEN1, Holliday junction 5' flap endonuclease | 1.63 |
| ENSMUSG00000025154 | Arhgap19 | Rho GTPase activating protein 19 | 1.62 |
| ENSMUSG00000034773 | BC030867 | cDNA sequence BC030867 | 1.62 |
| ENSMUSG00000074476 | Spc24 | SPC24, NDC80 kinetochore complex component, homolog (S. cerevisiae) | 1.61 |
| ENSMUSG00000021697 | Depdc1b | DEP domain containing 1B | 1.61 |
| ENSMUSG00000017861 | Mybl2 | myeloblastosis oncogene-like 2 | 1.60 |
| ENSMUSG00000029516 | Cit | citron | 1.60 |
| ENSMUSG00000023004 | Tuba1b | tubulin, alpha 1B | 1.60 |
| ENSMUSG00000081670 | Gm15697 | predicted gene 15697 | 1.60 |
| ENSMUSG00000046591 | Ticrr | TOPBP1-interacting checkpoint and replication regulator | 1.59 |

| | | | |
|---|---|---|---|
| ENSMUSG00000062248 | Cks2 | CDC28 protein kinase regulatory subunit 2 | 1.58 |
| ENSMUSG00000029521 | Chek2 | checkpoint kinase 2 | 1.58 |
| ENSMUSG00000031257 | Nox1 | NADPH oxidase 1 | 1.58 |
| ENSMUSG00000034349 | Smc4 | structural maintenance of chromosomes 4 | 1.57 |
| ENSMUSG00000025902 | Sox17 | SRY (sex determining region Y)-box 17 | 1.57 |
| ENSMUSG00000011267 | Zfp296 | zinc finger protein 296 | 1.57 |
| ENSMUSG00000020808 | Pimreg | PICALM interacting mitotic regulator | 1.57 |
| ENSMUSG00000000028 | Cdc45 | cell division cycle 45 | 1.56 |
| ENSMUSG00000005470 | Asf1b | anti-silencing function 1B histone chaperone | 1.56 |
| ENSMUSG00000035455 | Fignl1 | fidgetin-like 1 | 1.56 |
| ENSMUSG00000026669 | Mcm10 | minichromosome maintenance 10 replication initiation factor | 1.54 |
| ENSMUSG00000063021 | Hist1h2ak | histone cluster 1, H2ak | 1.54 |
| ENSMUSG00000078521 | Aunip | aurora kinase A and ninein interacting protein | 1.54 |
| ENSMUSG00000079357 | Gm11100 | predicted gene 11100 | 1.54 |
| ENSMUSG00000054717 | Hmgb2 | high mobility group box 2 | 1.53 |
| ENSMUSG00000025758 | Plk4 | polo like kinase 4 | 1.53 |
| ENSMUSG00000042489 | Clspn | claspin | 1.53 |
| ENSMUSG00000060093 | Hist1h4a | histone cluster 1, H4a | 1.53 |
| ENSMUSG00000068744 | Psrc1 | proline/serine-rich coiled-coil 1 | 1.53 |
| ENSMUSG00000052565 | Hist1h1d | histone cluster 1, H1d | 1.52 |
| ENSMUSG00000002297 | Dbf4 | DBF4 zinc finger | 1.52 |
| ENSMUSG00000024791 | Cdca5 | cell division cycle associated 5 | 1.52 |
| ENSMUSG00000045273 | Cenph | centromere protein H | 1.52 |
| ENSMUSG00000102840 | Gm38037 | predicted gene, 38037 | 1.52 |
| ENSMUSG00000024590 | Lmnb1 | lamin B1 | 1.51 |
| ENSMUSG00000005410 | Mcm5 | minichromosome maintenance complex component 5 | 1.51 |
| ENSMUSG00000020185 | E2f7 | E2F transcription factor 7 | 1.51 |
| ENSMUSG00000039994 | Timeless | timeless circadian clock 1 | 1.51 |
| ENSMUSG00000049539 | Hist1h1a | histone cluster 1, H1a | 1.50 |
| ENSMUSG00000027203 | Dut | deoxyuridine triphosphatase | 1.50 |
| ENSMUSG00000034023 | Fancd2 | Fanconi anemia, complementation group D2 | 1.50 |
| ENSMUSG00000072980 | Oip5 | Opa interacting protein 5 | 1.50 |
| ENSMUSG00000028933 | Xrcc2 | X-ray repair complementing defective repair in Chinese hamster cells 2 | 1.50 |
| ENSMUSG00000069307 | Hist1h2bq | histone cluster 1, H2bq | 1.50 |
| ENSMUSG00000049932 | H2afx | H2A histone family, member X | 1.49 |
| ENSMUSG00000030978 | Rrm1 | ribonucleotide reductase M1 | 1.49 |
| ENSMUSG00000046179 | E2f8 | E2F transcription factor 8 | 1.49 |
| ENSMUSG00000062937 | Mtap | methylthioadenosine phosphorylase | 1.49 |
| ENSMUSG00000039509 | Nup133 | nucleoporin 133 | 1.49 |
| ENSMUSG00000032400 | Zwilch | zwilch kinetochore protein | 1.49 |
| ENSMUSG00000061615 | Hist1h2ab | histone cluster 1, H2ab | 1.49 |
| ENSMUSG00000039187 | Fanci | Fanconi anemia, complementation group I | 1.49 |
| ENSMUSG00000069266 | Hist1h4b | histone cluster 1, H4b | 1.48 |
| ENSMUSG00000040658 | Dnph1 | 2'-deoxynucleoside 5'-phosphate N-hydrolase 1 | 1.48 |
| ENSMUSG00000110644 | Gm7390 | predicted gene 7390 | 1.48 |
| ENSMUSG00000062510 | Nsl1 | NSL1, MIS12 kinetochore complex component | 1.47 |
| ENSMUSG00000027635 | Dsn1 | DSN1 homolog, MIS12 kinetochore complex component | 1.47 |
| ENSMUSG00000069268 | Hist1h2bf | histone cluster 1, H2bf | 1.47 |
| ENSMUSG00000059791 | Nrm | nurim (nuclear envelope membrane protein) | 1.47 |
| ENSMUSG00000054342 | Kcnn4 | potassium intermediate/small conductance calcium-activated channel, subfamily N, member 4 | 1.46 |
| ENSMUSG00000028702 | Rad54l | RAD54 like (S. cerevisiae) | 1.46 |
| ENSMUSG00000035351 | Nup37 | nucleoporin 37 | 1.46 |
| ENSMUSG00000073705 | Cenps | centromere protein S | 1.46 |
| ENSMUSG00000032586 | Traip | TRAF-interacting protein | 1.46 |
| ENSMUSG00000110631 | Gm42047 | predicted gene, 42047 | 1.46 |
| ENSMUSG00000080186 | Gm14448 | predicted gene 14448 | 1.46 |
| ENSMUSG00000001228 | Uhrf1 | ubiquitin-like, containing PHD and RING finger domains, 1 | 1.45 |
| ENSMUSG00000025747 | Tyms | thymidylate synthase | 1.45 |
| ENSMUSG00000039396 | Neil3 | nei like 3 (E. coli) | 1.45 |
| ENSMUSG00000069184 | Zfp72 | zinc finger protein 72 | 1.45 |
| ENSMUSG00000028614 | Ndc1 | NDC1 transmembrane nucleoporin | 1.44 |
| ENSMUSG00000030346 | Rad51ap1 | RAD51 associated protein 1 | 1.44 |
| ENSMUSG00000069272 | Hist1h2ae | histone cluster 1, H2ae | 1.44 |
| ENSMUSG00000031112 | Stk26 | serine/threonine kinase 26 | 1.44 |
| ENSMUSG00000007050 | Lsm2 | LSM2 homolog, U6 small nuclear RNA and mRNA degradation associated | 1.43 |
| ENSMUSG00000019773 | Fbxo5 | F-box protein 5 | 1.43 |
| ENSMUSG00000041859 | Mcm3 | minichromosome maintenance complex component 3 | 1.43 |
| ENSMUSG00000024613 | Tcof1 | treacle ribosome biogenesis factor 1 | 1.43 |
| ENSMUSG00000069265 | Hist1h3a | histone cluster 1, H3a | 1.43 |
| ENSMUSG00000024891 | Slc29a2 | solute carrier family 29 (nucleoside transporters), member 2 | 1.43 |
| ENSMUSG00000026779 | Mastl | microtubule associated serine/threonine kinase-like | 1.42 |
| ENSMUSG00000031629 | Cenpu | centromere protein U | 1.42 |
| ENSMUSG00000028718 | Stil | Scl/Tal1 interrupting locus | 1.41 |

| | | | |
|---|---|---|---|
| ENSMUSG00000021175 | Cdca7l | cell division cycle associated 7 like | 1.41 |
| ENSMUSG00000032344 | Mb21d1 | Mab-21 domain containing 1 | 1.41 |
| ENSMUSG00000037466 | Tedc1 | tubulin epsilon and delta complex 1 | 1.41 |
| ENSMUSG00000039055 | Eme1 | essential meiotic structure-specific endonuclease 1 | 1.41 |
| ENSMUSG00000075266 | Cenpw | centromere protein W | 1.41 |
| ENSMUSG00000056394 | Lig1 | ligase I, DNA, ATP-dependent | 1.40 |
| ENSMUSG00000020534 | Shmt1 | serine hydroxymethyltransferase 1 (soluble) | 1.40 |
| ENSMUSG00000068101 | Cenpm | centromere protein M | 1.40 |
| ENSMUSG00000027353 | Mcm8 | minichromosome maintenance 8 homologous recombination repair factor | 1.40 |
| ENSMUSG00000063018 | 2010204K13Rik | RIKEN cDNA 2010204K13 gene | 1.39 |
| ENSMUSG00000091021 | Gm17300 | predicted gene, 17300 | 1.39 |
| ENSMUSG00000039200 | Atf7ip2 | activating transcription factor 7 interacting protein 2 | 1.39 |
| ENSMUSG00000106883 | Gm34091 | predicted gene, 34091 | 1.39 |
| ENSMUSG00000022177 | Haus4 | HAUS augmin-like complex, subunit 4 | 1.38 |
| ENSMUSG00000038644 | Pold1 | polymerase (DNA directed), delta 1, catalytic subunit | 1.38 |
| ENSMUSG00000034206 | Polq | polymerase (DNA directed), theta | 1.38 |
| ENSMUSG00000082163 | Gm14276 | predicted gene 14276 | 1.38 |
| ENSMUSG00000006585 | Cdt1 | chromatin licensing and DNA replication factor 1 | 1.37 |
| ENSMUSG00000027018 | Hat1 | histone aminotransferase 1 | 1.37 |
| ENSMUSG00000007080 | Pole | polymerase (DNA directed), epsilon | 1.37 |
| ENSMUSG00000022881 | Rfc4 | replication factor C (activator 1) 4 | 1.37 |
| ENSMUSG00000074825 | Itpripl1 | inositol 1,4,5-triphosphate receptor interacting protein-like 1 | 1.37 |
| ENSMUSG00000116542 | Gm17783 | predicted, 17783 | 1.37 |
| ENSMUSG00000030528 | Blm | Bloom syndrome, RecQ like helicase | 1.37 |
| ENSMUSG00000098090 | 2700099C18Rik | RIKEN cDNA 2700099C18 gene | 1.37 |
| ENSMUSG00000041219 | Arhgap11a | Rho GTPase activating protein 11A | 1.36 |
| ENSMUSG00000025403 | Shmt2 | serine hydroxymethyltransferase 2 (mitochondrial) | 1.36 |
| ENSMUSG00000091405 | Hist2h4 | histone cluster 2, H4 | 1.36 |
| ENSMUSG00000030641 | Ddias | DNA damage-induced apoptosis suppressor | 1.36 |
| ENSMUSG00000050107 | Haspin | histone H3 associated protein kinase | 1.35 |
| ENSMUSG00000026196 | Bard1 | BRCA1 associated RING domain 1 | 1.35 |
| ENSMUSG00000029414 | Kntc1 | kinetochore associated 1 | 1.35 |
| ENSMUSG00000044005 | Gls2 | glutaminase 2 (liver, mitochondrial) | 1.35 |
| ENSMUSG00000098318 | Lockd | lncRNA downstream of Cdkn1b | 1.35 |
| ENSMUSG00000022673 | Mcm4 | minichromosome maintenance complex component 4 | 1.34 |
| ENSMUSG00000032815 | Fanca | Fanconi anemia, complementation group A | 1.34 |
| ENSMUSG00000022364 | Tbc1d31 | TBC1 domain family, member 31 | 1.34 |
| ENSMUSG00000036875 | Dna2 | DNA replication helicase/nuclease 2 | 1.34 |
| ENSMUSG00000087060 | Eldr | Egfr long non-coding downstream RNA | 1.34 |
| ENSMUSG00000028602 | Tnfrsf8 | tumor necrosis factor receptor superfamily, member 8 | 1.34 |
| ENSMUSG00000019961 | Tmpo | thymopoietin | 1.33 |
| ENSMUSG00000028896 | Rcc1 | regulator of chromosome condensation 1 | 1.33 |
| ENSMUSG00000029730 | Mcm7 | minichromosome maintenance complex component 7 | 1.33 |
| ENSMUSG00000001056 | Nhp2 | NHP2 ribonucleoprotein | 1.33 |
| ENSMUSG00000040034 | Nup43 | nucleoporin 43 | 1.33 |
| ENSMUSG00000022945 | Chaf1b | chromatin assembly factor 1, subunit B (p60) | 1.33 |
| ENSMUSG00000028066 | Pmf1 | polyamine-modulated factor 1 | 1.32 |
| ENSMUSG00000026434 | Nucks1 | nuclear casein kinase and cyclin-dependent kinase substrate 1 | 1.32 |
| ENSMUSG00000005732 | Ranbp1 | RAN binding protein 1 | 1.32 |
| ENSMUSG00000004880 | Lbr | lamin B receptor | 1.32 |
| ENSMUSG00000100210 | Hist1h3f | histone cluster 1, H3f | 1.32 |
| ENSMUSG00000044702 | Palb2 | partner and localizer of BRCA2 | 1.32 |
| ENSMUSG00000033752 | Mnd1 | meiotic nuclear divisions 1 | 1.32 |
| ENSMUSG00000083921 | Gm15750 | predicted gene 15750 | 1.32 |
| ENSMUSG00000097180 | 2700038G22Rik | RIKEN cDNA 2700038G22 gene | 1.32 |
| ENSMUSG00000099474 | 1700097N02Rik | RIKEN cDNA 1700097N02 gene | 1.32 |
| ENSMUSG00000078586 | Gm11735 | predicted gene 11735 | 1.32 |
| ENSMUSG00000086721 | Gm13750 | predicted gene 13750 | 1.32 |
| ENSMUSG00000028693 | Nasp | nuclear autoantigenic sperm protein (histone-binding) | 1.31 |
| ENSMUSG00000025980 | Hspd1 | heat shock protein 1 (chaperonin) | 1.31 |
| ENSMUSG00000002870 | Mcm2 | minichromosome maintenance complex component 2 | 1.31 |
| ENSMUSG00000067455 | Hist1h4j | histone cluster 1, H4j | 1.31 |
| ENSMUSG00000021391 | Cenpp | centromere protein P | 1.31 |
| ENSMUSG00000073184 | Gm10479 | predicted gene 10479 | 1.31 |
| ENSMUSG00000028884 | Rpa2 | replication protein A2 | 1.30 |
| ENSMUSG00000002835 | Chaf1a | chromatin assembly factor 1, subunit A (p150) | 1.30 |
| ENSMUSG00000025001 | Hells | helicase, lymphoid specific | 1.30 |
| ENSMUSG00000037020 | Wdr62 | WD repeat domain 62 | 1.30 |
| ENSMUSG00000022325 | Pop1 | processing of precursor 1, ribonuclease P/MRP family, (S. cerevisiae) | 1.30 |
| ENSMUSG00000029283 | Cdc7 | cell division cycle 7 (S. cerevisiae) | 1.30 |
| ENSMUSG00000053398 | Phgdh | 3-phosphoglycerate dehydrogenase | 1.30 |
| ENSMUSG00000030929 | Eri2 | exoribonuclease 2 | 1.30 |
| ENSMUSG00000027654 | Fam83d | family with sequence similarity 83, member D | 1.30 |

| | | | |
|---|---|---|---|
| ENSMUSG00000097651 | 4930461G14Rik | RIKEN cDNA 4930461G14 gene | 1.30 |
| ENSMUSG00000031762 | Mt2 | metallothionein 2 | 1.30 |
| ENSMUSG00000109245 | Gm44860 | predicted gene 44860 | 1.30 |
| ENSMUSG00000112406 | 4921515L22Rik | RIKEN cDNA 4921515L22 gene | 1.30 |
| ENSMUSG00000021115 | Vrk1 | vaccinia related kinase 1 | 1.29 |
| ENSMUSG00000026355 | Mcm6 | minichromosome maintenance complex component 6 | 1.29 |
| ENSMUSG00000060981 | Hist1h4h | histone cluster 1, H4h | 1.29 |
| ENSMUSG00000027752 | Exosc8 | exosome component 8 | 1.29 |
| ENSMUSG00000025395 | Prim1 | DNA primase, p49 subunit | 1.29 |
| ENSMUSG00000019992 | Mtfr2 | mitochondrial fission regulator 2 | 1.29 |
| ENSMUSG00000081740 | Gm14279 | predicted gene 14279 | 1.29 |
| ENSMUSG00000047246 | Hist1h2be | histone cluster 1, H2be | 1.28 |
| ENSMUSG00000101972 | Hist1h3i | histone cluster 1, H3i | 1.28 |
| ENSMUSG00000001525 | Tubb5 | tubulin, beta 5 class I | 1.27 |
| ENSMUSG00000034317 | Trim59 | tripartite motif-containing 59 | 1.27 |
| ENSMUSG00000076437 | Selenoh | selenoprotein H | 1.27 |
| ENSMUSG00000031403 | Dkc1 | dyskeratosis congenita 1, dyskerin | 1.27 |
| ENSMUSG00000105987 | AI506816 | expressed sequence AI506816 | 1.27 |
| ENSMUSG00000022360 | Atad2 | ATPase family, AAA domain containing 2 | 1.27 |
| ENSMUSG00000047443 | Erfe | erythroferrone | 1.27 |
| ENSMUSG00000028044 | Cks1b | CDC28 protein kinase 1b | 1.27 |
| ENSMUSG00000030254 | Rad18 | RAD18 E3 ubiquitin protein ligase | 1.27 |
| ENSMUSG00000020649 | Rrm2 | ribonucleotide reductase M2 | 1.27 |
| ENSMUSG00000009248 | Ascl2 | achaete-scute family bHLH transcription factor 2 | 1.27 |
| ENSMUSG00000048779 | P2ry6 | pyrimidinergic receptor P2Y, G-protein coupled, 6 | 1.27 |
| ENSMUSG00000012483 | Rpa3 | replication protein A3 | 1.26 |
| ENSMUSG00000029366 | Dck | deoxycytidine kinase | 1.26 |
| ENSMUSG00000030079 | Ruvbl1 | RuvB-like protein 1 | 1.25 |
| ENSMUSG00000035165 | Kcne3 | potassium voltage-gated channel, Isk-related subfamily, gene 3 | 1.25 |
| ENSMUSG00000024691 | Fam111a | family with sequence similarity 111, member A | 1.25 |
| ENSMUSG00000022978 | Mis18a | MIS18 kinetochore protein A | 1.25 |
| ENSMUSG00000038046 | Mrm3 | mitochondrial rRNA methyltransferase 3 | 1.25 |
| ENSMUSG00000036526 | Card11 | caspase recruitment domain family, member 11 | 1.25 |
| ENSMUSG00000097412 | 1810014B01Rik | RIKEN cDNA 1810014B01 gene | 1.25 |
| ENSMUSG00000078952 | Lncenc1 | long non-coding RNA, embryonic stem cells expressed 1 | 1.25 |
| ENSMUSG00000066551 | Hmgb1 | high mobility group box 1 | 1.24 |
| ENSMUSG00000042029 | Ncapg2 | non-SMC condensin II complex, subunit G2 | 1.24 |
| ENSMUSG00000057113 | Npm1 | nucleophosmin 1 | 1.24 |
| ENSMUSG00000021048 | Mthfd1 | methylenetetrahydrofolate dehydrogenase (NADP+ dependent), methenyltetrahydrofolate cyclohyc | 1.24 |
| ENSMUSG00000022792 | Yars2 | tyrosyl-tRNA synthetase 2 (mitochondrial) | 1.24 |
| ENSMUSG00000067367 | Lyar | Ly1 antibody reactive clone | 1.24 |
| ENSMUSG00000028587 | Orc1 | origin recognition complex, subunit 1 | 1.24 |
| ENSMUSG00000109936 | Gm45889 | predicted gene 45889 | 1.24 |
| ENSMUSG00000044320 | 1700001O22Rik | RIKEN cDNA 1700001O22 gene | 1.24 |
| ENSMUSG00000060794 | Tssk5 | testis-specific serine kinase 5 | 1.24 |
| ENSMUSG00000022962 | Gart | phosphoribosylglycinamide formyltransferase | 1.23 |
| ENSMUSG00000027405 | Nop56 | NOP56 ribonucleoprotein | 1.23 |
| ENSMUSG00000026234 | Ncl | nucleolin | 1.23 |
| ENSMUSG00000031697 | Orc6 | origin recognition complex, subunit 6 | 1.23 |
| ENSMUSG00000022026 | Olfm4 | olfactomedin 4 | 1.23 |
| ENSMUSG00000039748 | Exo1 | exonuclease 1 | 1.23 |
| ENSMUSG00000026785 | Pkn3 | protein kinase N3 | 1.23 |
| ENSMUSG00000045709 | Smkr-ps | smal lysine rich protein 1, pseudogene | 1.23 |
| ENSMUSG00000116009 | Gm49503 | predicted gene, 49503 | 1.23 |
| ENSMUSG00000025991 | Cps1 | carbamoyl-phosphate synthetase 1 | 1.22 |
| ENSMUSG00000096010 | Hist4h4 | histone cluster 4, H4 | 1.22 |
| ENSMUSG00000039298 | Cdk5rap2 | CDK5 regulatory subunit associated protein 2 | 1.22 |
| ENSMUSG00000032555 | Topbp1 | topoisomerase (DNA) II binding protein 1 | 1.22 |
| ENSMUSG00000027424 | Mgme1 | mitochondrial genome maintenance exonuclease 1 | 1.22 |
| ENSMUSG00000086513 | 9130208D14Rik | RIKEN cDNA 9130208D14 gene | 1.22 |
| ENSMUSG00000025912 | Mybl1 | myeloblastosis oncogene-like 1 | 1.22 |
| ENSMUSG00000022754 | Tmem45a | transmembrane protein 45a | 1.22 |
| ENSMUSG00000081151 | Gm11448 | predicted gene 11448 | 1.22 |
| ENSMUSG00000052833 | Sae1 | SUMO1 activating enzyme subunit 1 | 1.21 |
| ENSMUSG00000004099 | Dnmt1 | DNA methyltransferase (cytosine-5) 1 | 1.21 |
| ENSMUSG00000050410 | Tcf19 | transcription factor 19 | 1.21 |
| ENSMUSG00000020974 | Pole2 | polymerase (DNA directed), epsilon 2 (p59 subunit) | 1.21 |
| ENSMUSG00000014633 | Cmc2 | COX assembly mitochondrial protein 2 | 1.21 |
| ENSMUSG00000047757 | Fancb | Fanconi anemia, complementation group B | 1.21 |
| ENSMUSG00000114515 | Aldoa | aldolase A, fructose-bisphosphate | 1.21 |
| ENSMUSG00000060777 | Gm10087 | predicted gene 10087 | 1.21 |
| ENSMUSG00000029430 | Ran | RAN, member RAS oncogene family | 1.20 |
| ENSMUSG00000044254 | Pcsk9 | proprotein convertase subtilisin/kexin type 9 | 1.20 |

| | | | |
|---|---|---|---|
| ENSMUSG00000015749 | Anp32e | acidic (leucine-rich) nuclear phosphoprotein 32 family, member E | 1.20 |
| ENSMUSG00000003868 | Ruvbl2 | RuvB-like protein 2 | 1.20 |
| ENSMUSG00000017493 | Igfbp4 | insulin-like growth factor binding protein 4 | 1.20 |
| ENSMUSG00000039356 | Exosc2 | exosome component 2 | 1.20 |
| ENSMUSG00000022797 | Tfrc | transferrin receptor | 1.20 |
| ENSMUSG00000069793 | Slfn9 | schlafen 9 | 1.20 |
| ENSMUSG00000041193 | Pla2g5 | phospholipase A2, group V | 1.20 |
| ENSMUSG00000019214 | Chtf18 | CTF18, chromosome transmission fidelity factor 18 | 1.20 |
| ENSMUSG00000027454 | Gins1 | GINS complex subunit 1 (Psf1 homolog) | 1.20 |
| ENSMUSG00000073415 | Gm10501 | predicted gene 10501 | 1.20 |
| ENSMUSG00000018446 | C1qbp | complement component 1, q subcomponent binding protein | 1.19 |
| ENSMUSG00000034192 | Lsm3 | LSM3 homolog, U6 small nuclear RNA and mRNA degradation associated | 1.19 |
| ENSMUSG00000006715 | Gmnn | geminin | 1.19 |
| ENSMUSG00000053801 | Grwd1 | glutamate-rich WD repeat containing 1 | 1.19 |
| ENSMUSG00000097415 | AU020206 | expressed sequence AU020206 | 1.19 |
| ENSMUSG00000056904 | Gm5620 | predicted gene 5620 | 1.19 |
| ENSMUSG00000037148 | Arhgap10 | Rho GTPase activating protein 10 | 1.19 |
| ENSMUSG00000035818 | Plekhs1 | pleckstrin homology domain containing, family S member 1 | 1.19 |
| ENSMUSG00000068220 | Lgals1 | lectin, galactose binding, soluble 1 | 1.19 |
| ENSMUSG00000071516 | Hist1h2ai | histone cluster 1, H2ai | 1.19 |
| ENSMUSG00000097755 | 2010110K18Rik | RIKEN cDNA 2010110K18 gene | 1.19 |
| ENSMUSG00000042606 | Hirip3 | HIRA interacting protein 3 | 1.18 |
| ENSMUSG00000039231 | Suv39h1 | suppressor of variegation 3-9 1 | 1.18 |
| ENSMUSG00000038323 | 1700066M21Rik | RIKEN cDNA 1700066M21 gene | 1.18 |
| ENSMUSG00000021418 | Rpp40 | ribonuclease P 40 subunit | 1.18 |
| ENSMUSG00000079139 | Gm4204 | predicted gene 4204 | 1.18 |
| ENSMUSG00000022096 | Hr | hairless | 1.17 |
| ENSMUSG00000002477 | Snrpd1 | small nuclear ribonucleoprotein D1 | 1.17 |
| ENSMUSG00000002718 | Cse1l | chromosome segregation 1-like (S. cerevisiae) | 1.17 |
| ENSMUSG00000032939 | Nup93 | nucleoporin 93 | 1.17 |
| ENSMUSG00000068394 | Cep152 | centrosomal protein 152 | 1.17 |
| ENSMUSG00000025507 | Pidd1 | p53 induced death domain protein 1 | 1.17 |
| ENSMUSG00000027395 | Polr1b | polymerase (RNA) I polypeptide B | 1.17 |
| ENSMUSG00000014907 | Naf1 | nuclear assembly factor 1 ribonucleoprotein | 1.17 |
| ENSMUSG00000025742 | Prps2 | phosphoribosyl pyrophosphate synthetase 2 | 1.17 |
| ENSMUSG00000005360 | Slc1a3 | solute carrier family 1 (glial high affinity glutamate transporter), member 3 | 1.17 |
| ENSMUSG00000041840 | Haus1 | HAUS augmin-like complex, subunit 1 | 1.17 |
| ENSMUSG00000063445 | Nmral1 | NmrA-like family domain containing 1 | 1.17 |
| ENSMUSG00000060950 | Trmt61a | tRNA methyltransferase 61A | 1.17 |
| ENSMUSG00000003824 | Syce2 | synaptonemal complex central element protein 2 | 1.17 |
| ENSMUSG00000078502 | Gm13212 | predicted gene 13212 | 1.17 |
| ENSMUSG00000042854 | Trp53rkb | transformation related protein 53 regulating kinase B | 1.17 |
| ENSMUSG00000062588 | Gm6104 | predicted gene 6104 | 1.17 |
| ENSMUSG00000064672 | Gm22806 | predicted gene, 22806 | 1.17 |
| ENSMUSG00000069273 | Hist1h3e | histone cluster 1, H3e | 1.16 |
| ENSMUSG00000041506 | Rrp9 | RRP9, small subunit (SSU) processome component, homolog (yeast) | 1.16 |
| ENSMUSG00000017146 | Brca1 | breast cancer 1, early onset | 1.16 |
| ENSMUSG00000061533 | Cep128 | centrosomal protein 128 | 1.16 |
| ENSMUSG00000086421 | Gm14091 | predicted gene 14091 | 1.16 |
| ENSMUSG00000057244 | Gm6139 | predicted gene 6139 | 1.16 |
| ENSMUSG00000022346 | Myc | myelocytomatosis oncogene | 1.15 |
| ENSMUSG00000005481 | Ddx39 | DEAD (Asp-Glu-Ala-Asp) box polypeptide 39 | 1.15 |
| ENSMUSG00000028333 | Anp32b | acidic (leucine-rich) nuclear phosphoprotein 32 family, member B | 1.15 |
| ENSMUSG00000024925 | Rnaseh2c | ribonuclease H2, subunit C | 1.15 |
| ENSMUSG00000026646 | Suv39h2 | suppressor of variegation 3-9 2 | 1.15 |
| ENSMUSG00000059939 | 9430015G10Rik | RIKEN cDNA 9430015G10 gene | 1.15 |
| ENSMUSG00000060639 | Hist1h4i | histone cluster 1, H4i | 1.15 |
| ENSMUSG00000079685 | Ulbp1 | UL16 binding protein 1 | 1.15 |
| ENSMUSG00000020773 | Trim47 | tripartite motif-containing 47 | 1.15 |
| ENSMUSG00000017607 | Tns4 | tensin 4 | 1.14 |
| ENSMUSG00000024844 | Banf1 | barrier to autointegration factor 1 | 1.14 |
| ENSMUSG00000032373 | Car12 | carbonic anhydrase 12 | 1.14 |
| ENSMUSG00000095567 | Noc2l | NOC2 like nucleolar associated transcriptional repressor | 1.14 |
| ENSMUSG00000033307 | Mif | macrophage migration inhibitory factor (glycosylation-inhibiting factor) | 1.14 |
| ENSMUSG00000022913 | Psmg1 | proteasome (prosome, macropain) assembly chaperone 1 | 1.14 |
| ENSMUSG00000025732 | Mcrip2 | MAPK regulated corepressor interacting protein 2 | 1.14 |
| ENSMUSG00000021714 | Cenpk | centromere protein K | 1.14 |
| ENSMUSG00000109685 | Gm45912 | predicted gene 45912 | 1.14 |
| ENSMUSG00000044827 | Tlr1 | toll-like receptor 1 | 1.14 |
| ENSMUSG00000069308 | Hist1h2bp | histone cluster 1, H2bp | 1.14 |
| ENSMUSG00000028560 | Usp1 | ubiquitin specific peptidase 1 | 1.13 |
| ENSMUSG00000060860 | Ube2s | ubiquitin-conjugating enzyme E2S | 1.13 |
| ENSMUSG00000070348 | Ccnd1 | cyclin D1 | 1.13 |

| | | | |
|---|---|---|---|
| ENSMUSG00000015176 | Nolc1 | nucleolar and coiled-body phosphoprotein 1 | 1.13 |
| ENSMUSG00000030662 | Ipo5 | importin 5 | 1.13 |
| ENSMUSG00000037275 | Gemin5 | gem nuclear organelle associated protein 5 | 1.13 |
| ENSMUSG00000022070 | Bora | bora, aurora kinase A activator | 1.13 |
| ENSMUSG00000015217 | Hmgb3 | high mobility group box 3 | 1.13 |
| ENSMUSG00000057541 | Pus7 | pseudouridylate synthase 7 | 1.13 |
| ENSMUSG00000013629 | Cad | carbamoyl-phosphate synthetase 2, aspartate transcarbamylase, and dihydroorotase | 1.13 |
| ENSMUSG00000097769 | Snhg4 | small nucleolar RNA host gene 4 | 1.13 |
| ENSMUSG00000035960 | Apex1 | apurinic/apyrimidinic endonuclease 1 | 1.13 |
| ENSMUSG00000025665 | Rps6ka6 | ribosomal protein S6 kinase polypeptide 6 | 1.13 |
| ENSMUSG00000026238 | Ptma | prothymosin alpha | 1.12 |
| ENSMUSG00000027342 | Pcna | proliferating cell nuclear antigen | 1.12 |
| ENSMUSG00000054766 | Set | SET nuclear oncogene | 1.12 |
| ENSMUSG00000061482 | Hist1h4d | histone cluster 1, H4d | 1.12 |
| ENSMUSG00000036109 | Mbnl3 | muscleblind like splicing factor 3 | 1.12 |
| ENSMUSG00000028633 | Ctps | cytidine 5'-triphosphate synthase | 1.12 |
| ENSMUSG00000023908 | Pkmyt1 | protein kinase, membrane associated tyrosine/threonine 1 | 1.12 |
| ENSMUSG00000055760 | Gemin6 | gem nuclear organelle associated protein 6 | 1.12 |
| ENSMUSG00000059493 | Nhs | NHS actin remodeling regulator | 1.12 |
| ENSMUSG00000035439 | Haus8 | 4HAUS augmin-like complex, subunit 8 | 1.12 |
| ENSMUSG00000052852 | Reep1 | receptor accessory protein 1 | 1.12 |
| ENSMUSG00000026192 | Atic | 5-aminoimidazole-4-carboxamide ribonucleotide formyltransferase/IMP cyclohydrolase | 1.11 |
| ENSMUSG00000054321 | Taf4b | TATA-box binding protein associated factor 4b | 1.11 |
| ENSMUSG00000020706 | Ftsj3 | FtsJ RNA methyltransferase homolog 3 (E. coli) | 1.11 |
| ENSMUSG00000046711 | Hmga1 | high mobility group AT-hook 1 | 1.11 |
| ENSMUSG00000026281 | Dtymk | deoxythymidylate kinase | 1.11 |
| ENSMUSG00000111877 | Gm6477 | predicted gene 6477 | 1.11 |
| ENSMUSG00000040463 | Mybbp1a | MYB binding protein (P160) 1a | 1.11 |
| ENSMUSG00000033706 | Smyd5 | SET and MYND domain containing 5 | 1.11 |
| ENSMUSG00000036678 | Aaas | achalasia, adrenocortical insufficiency, alacrimia | 1.11 |
| ENSMUSG00000023348 | Trip6 | thyroid hormone receptor interactor 6 | 1.11 |
| ENSMUSG00000080928 | Hmgb1-ps6 | high mobility group box 1, pseudogene 6 | 1.11 |
| ENSMUSG00000103576 | Gm38188 | predicted gene, 38188 | 1.11 |
| ENSMUSG00000024640 | Psat1 | phosphoserine aminotransferase 1 | 1.10 |
| ENSMUSG00000020739 | Nup85 | nucleoporin 85 | 1.10 |
| ENSMUSG00000003438 | Timm50 | translocase of inner mitochondrial membrane 50 | 1.10 |
| ENSMUSG00000021377 | Dek | DEK oncogene (DNA binding) | 1.10 |
| ENSMUSG00000023110 | Prmt5 | protein arginine N-methyltransferase 5 | 1.10 |
| ENSMUSG00000050244 | Heatr1 | HEAT repeat containing 1 | 1.10 |
| ENSMUSG00000091625 | Lsm5 | LSM5 homolog, U6 small nuclear RNA and mRNA degradation associated | 1.10 |
| ENSMUSG00000037474 | Dtl | denticleless E3 ubiquitin protein ligase | 1.10 |
| ENSMUSG00000054115 | Skp2 | S-phase kinase-associated protein 2 (p45) | 1.10 |
| ENSMUSG00000026404 | Ddx59 | DEAD (Asp-Glu-Ala-Asp) box polypeptide 59 | 1.10 |
| ENSMUSG00000073627 | C130036L24Rik | RIKEN cDNA C130036L24 gene | 1.10 |
| ENSMUSG00000106358 | Gm7047 | predicted gene 7047 | 1.10 |
| ENSMUSG00000097177 | 9330159M07Rik | RIKEN cDNA 9330159M07 gene | 1.10 |
| ENSMUSG00000038685 | Rtel1 | regulator of telomere elongation helicase 1 | 1.09 |
| ENSMUSG00000030512 | Snrpa1 | small nuclear ribonucleoprotein polypeptide A' | 1.09 |
| ENSMUSG00000109511 | Nup62 | nucleoporin 62 | 1.09 |
| ENSMUSG00000027067 | Ssrp1 | structure specific recognition protein 1 | 1.09 |
| ENSMUSG00000004356 | Utp20 | UTP20 small subunit processome component | 1.09 |
| ENSMUSG00000027160 | Ccdc34 | coiled-coil domain containing 34 | 1.09 |
| ENSMUSG00000026020 | Nop58 | NOP58 ribonucleoprotein | 1.09 |
| ENSMUSG00000070304 | Scn2b | sodium channel, voltage-gated, type II, beta | 1.09 |
| ENSMUSG00000019982 | Myb | myeloblastosis oncogene | 1.09 |
| ENSMUSG00000053553 | 3110082I17Rik | RIKEN cDNA 3110082I17 gene | 1.09 |
| ENSMUSG00000024800 | Rpp30 | ribonuclease P/MRP 30 subunit | 1.09 |
| ENSMUSG00000097493 | 9930014A18Rik | RIKEN cDNA 9930014A18 gene | 1.09 |
| ENSMUSG00000039254 | Pomt1 | protein-O-mannosyltransferase 1 | 1.09 |
| ENSMUSG00000022748 | Cmss1 | cms small ribosomal subunit 1 | 1.09 |
| ENSMUSG00000078941 | Ak6 | adenylate kinase 6 | 1.09 |
| ENSMUSG00000068855 | Hist2h2ac | histone cluster 2, H2ac | 1.09 |
| ENSMUSG00000115306 | Gm49159 | predicted gene, 49159 | 1.09 |
| ENSMUSG00000025364 | Pa2g4 | proliferation-associated 2G4 | 1.08 |
| ENSMUSG00000033166 | Dis3 | DIS3 homolog, exosome endoribonuclease and 3'-5' exoribonuclease | 1.08 |
| ENSMUSG00000025134 | Alyref | Aly/REF export factor | 1.08 |
| ENSMUSG00000014226 | Cacybp | calcyclin binding protein | 1.08 |
| ENSMUSG00000058799 | Nap1l1 | nucleosome assembly protein 1-like 1 | 1.08 |
| ENSMUSG00000040681 | Hmgn1 | high mobility group nucleosomal binding domain 1 | 1.08 |
| ENSMUSG00000032459 | Mrps22 | mitochondrial ribosomal protein S22 | 1.08 |
| ENSMUSG00000097195 | Snhg5 | small nucleolar RNA host gene 5 | 1.08 |
| ENSMUSG00000036285 | Noa1 | nitric oxide associated 1 | 1.08 |
| ENSMUSG00000026999 | Nup35 | nucleoporin 35 | 1.08 |

| | | | |
|---|---|---|---|
| ENSMUSG00000021177 | Tdp1 | tyrosyl-DNA phosphodiesterase 1 | 1.08 |
| ENSMUSG00000036639 | Nudt1 | nudix (nucleoside diphosphate linked moiety X)-type motif 1 | 1.08 |
| ENSMUSG00000021572 | Cep72 | centrosomal protein 72 | 1.08 |
| ENSMUSG00000017832 | Hspb9 | heat shock protein, alpha-crystallin-related, B9 | 1.08 |
| ENSMUSG00000034424 | Gcsh | glycine cleavage system protein H (aminomethyl carrier) | 1.07 |
| ENSMUSG00000021193 | Pitrm1 | pitrilysin metallepetidase 1 | 1.07 |
| ENSMUSG00000018921 | Pelp1 | proline, glutamic acid and leucine rich protein 1 | 1.07 |
| ENSMUSG00000024833 | Pola2 | polymerase (DNA directed), alpha 2 | 1.07 |
| ENSMUSG00000020899 | Pfas | phosphoribosylformylglycinamidine synthase (FGAR amidotransferase) | 1.07 |
| ENSMUSG00000031453 | Rasa3 | RAS p21 protein activator 3 | 1.07 |
| ENSMUSG00000109324 | Prmt1 | protein arginine N-methyltransferase 1 | 1.07 |
| ENSMUSG00000023066 | Rttn | rotatin | 1.07 |
| ENSMUSG00000042215 | Bag2 | BCL2-associated athanogene 2 | 1.07 |
| ENSMUSG00000028391 | Wdr31 | WD repeat domain 31 | 1.07 |
| ENSMUSG00000062480 | Acat3 | acetyl-Coenzyme A acetyltransferase 3 | 1.07 |
| ENSMUSG00000057469 | E2f6 | E2F transcription factor 6 | 1.06 |
| ENSMUSG00000025869 | Nop16 | NOP16 nucleolar protein | 1.06 |
| ENSMUSG00000026709 | Dars2 | aspartyl-tRNA synthetase 2 (mitochondrial) | 1.06 |
| ENSMUSG00000025962 | Fastkd2 | FAST kinase domains 2 | 1.06 |
| ENSMUSG00000060288 | Ppih | peptidyl prolyl isomerase H | 1.06 |
| ENSMUSG00000017499 | Cdc6 | cell division cycle 6 | 1.06 |
| ENSMUSG00000027076 | Timm10 | translocase of inner mitochondrial membrane 10 | 1.06 |
| ENSMUSG00000056209 | Npm3 | nucleoplasmin 3 | 1.06 |
| ENSMUSG00000031821 | Gins2 | GINS complex subunit 2 (Psf2 homolog) | 1.06 |
| ENSMUSG00000024525 | Impa2 | inositol (myo)-1(or 4)-monophosphatase 2 | 1.06 |
| ENSMUSG00000079478 | Sssca1 | Sjogren syndrome/scleroderma autoantigen 1 | 1.06 |
| ENSMUSG00000014075 | Tctex1d2 | Tctex1 domain containing 2 | 1.06 |
| ENSMUSG00000026322 | Htr4 | 5 hydroxytryptamine (serotonin) receptor 4 | 1.06 |
| ENSMUSG00000103445 | Gm36948 | predicted gene, 36948 | 1.06 |
| ENSMUSG00000022698 | Naa50 | N(alpha)-acetyltransferase 50, NatE catalytic subunit | 1.05 |
| ENSMUSG00000005899 | Smpd4 | sphingomyelin phosphodiesterase 4 | 1.05 |
| ENSMUSG00000047832 | Cdca4 | cell division cycle associated 4 | 1.05 |
| ENSMUSG00000044934 | Zfp367 | zinc finger protein 367 | 1.05 |
| ENSMUSG00000026019 | Wdr12 | WD repeat domain 12 | 1.05 |
| ENSMUSG00000024411 | Aqp4 | aquaporin 4 | 1.05 |
| ENSMUSG00000031669 | Gins3 | GINS complex subunit 3 (Psf3 homolog) | 1.05 |
| ENSMUSG00000025086 | Trub1 | TruB pseudouridine (psi) synthase family member 1 | 1.05 |
| ENSMUSG00000085156 | Snhg15 | small nucleolar RNA host gene 15 | 1.05 |
| ENSMUSG00000084129 | Hmgb1-ps1 | high mobility group box 1, pseudogene 1 | 1.05 |
| ENSMUSG00000032446 | Eomes | eomesodermin | 1.05 |
| ENSMUSG00000025026 | Add3 | adducin 3 (gamma) | 1.04 |
| ENSMUSG00000029036 | Atad3a | ATPase family, AAA domain containing 3A | 1.04 |
| ENSMUSG00000062867 | Impdh2 | inosine monophosphate dehydrogenase 2 | 1.04 |
| ENSMUSG00000024732 | Ccdc86 | coiled-coil domain containing 86 | 1.04 |
| ENSMUSG00000031672 | Got2 | glutamatic-oxaloacetic transaminase 2, mitochondrial | 1.04 |
| ENSMUSG00000029246 | Ppat | phosphoribosyl pyrophosphate amidotransferase | 1.04 |
| ENSMUSG00000024785 | Rcl1 | RNA terminal phosphate cyclase-like 1 | 1.04 |
| ENSMUSG00000020471 | Pold2 | polymerase (DNA directed), delta 2, regulatory subunit | 1.04 |
| ENSMUSG00000058392 | Rrp1b | ribosomal RNA processing 1 homolog B (S. cerevisiae) | 1.04 |
| ENSMUSG00000002068 | Ccne1 | cyclin E1 | 1.04 |
| ENSMUSG00000026676 | Ccdc3 | coiled-coil domain containing 3 | 1.04 |
| ENSMUSG00000034329 | Brip1 | BRCA1 interacting protein C-terminal helicase 1 | 1.04 |
| ENSMUSG00000024078 | Ttc27 | tetratricopeptide repeat domain 27 | 1.04 |
| ENSMUSG00000032113 | Chek1 | checkpoint kinase 1 | 1.04 |
| ENSMUSG00000056305 | Usp39 | ubiquitin specific peptidase 39 | 1.04 |
| ENSMUSG00000028212 | Ccne2 | cyclin E2 | 1.04 |
| ENSMUSG00000023919 | Cenpq | centromere protein Q | 1.04 |
| ENSMUSG00000030513 | Pcsk6 | proprotein convertase subtilisin/kexin type 6 | 1.04 |
| ENSMUSG00000056531 | Ccdc18 | coiled-coil domain containing 18 | 1.04 |
| ENSMUSG00000095677 | Dynlt1f | dynein light chain Tctex-type 1F | 1.04 |
| ENSMUSG00000040321 | Zfp770 | zinc finger protein 770 | 1.04 |
| ENSMUSG00000090602 | Gm5611 | predicted gene 5611 | 1.04 |
| ENSMUSG00000106926 | Rpl7-ps7 | ribosomal protein L7, pseudogene 7 | 1.04 |
| ENSMUSG00000043424 | Eif3j2 | eukaryotic translation initiation factor 3, subunit J2 | 1.04 |
| ENSMUSG00000020290 | Xpo1 | exportin 1 | 1.03 |
| ENSMUSG00000061479 | Snrpa | small nuclear ribonucleoprotein polypeptide A | 1.03 |
| ENSMUSG00000038759 | Nup205 | nucleoporin 205 | 1.03 |
| ENSMUSG00000061718 | Ppp1r1b | protein phosphatase 1, regulatory inhibitor subunit 1B | 1.03 |
| ENSMUSG00000034203 | Chchd4 | coiled-coil-helix-coiled-coil-helix domain containing 4 | 1.03 |
| ENSMUSG00000024542 | Cep192 | centrosomal protein 192 | 1.03 |
| ENSMUSG00000038510 | Rpf2 | ribosome production factor 2 homolog | 1.03 |
| ENSMUSG00000020918 | Kat2a | K(lysine) acetyltransferase 2A | 1.03 |
| ENSMUSG00000038845 | Phb | prohibitin | 1.03 |

| | | | |
|---|---|---|---|
| ENSMUSG00000055612 | Cdca7 | cell division cycle associated 7 | 1.03 |
| ENSMUSG00000033294 | Noc4l | NOC4 like | 1.03 |
| ENSMUSG00000063480 | Snu13 | SNU13 homolog, small nuclear ribonucleoprotein (U4/U6.U5) | 1.03 |
| ENSMUSG00000105703 | Gm43305 | predicted gene 43305 | 1.03 |
| ENSMUSG00000056032 | BC018473 | cDNA sequence BC018473 | 1.03 |
| ENSMUSG00000105077 | Gm4859 | predicted gene 4859 | 1.03 |
| ENSMUSG00000068631 | Gm7676 | predicted gene 7676 | 1.03 |
| ENSMUSG00000114374 | Gm7911 | predicted gene 7911 | 1.03 |
| ENSMUSG00000042354 | Gnl3 | guanine nucleotide binding protein-like 3 (nucleolar) | 1.02 |
| ENSMUSG00000022678 | Nde1 | nudE neurodevelopment protein 1 | 1.02 |
| ENSMUSG00000020741 | Cluh | clustered mitochondria (cluA/CLU1) homolog | 1.02 |
| ENSMUSG00000063052 | Lrrc40 | leucine rich repeat containing 40 | 1.02 |
| ENSMUSG00000018583 | G3bp1 | GTPase activating protein (SH3 domain) binding protein 1 | 1.02 |
| ENSMUSG00000020929 | Eftud2 | elongation factor Tu GTP binding domain containing 2 | 1.02 |
| ENSMUSG00000014444 | Piezo1 | piezo-type mechanosensitive ion channel component 1 | 1.02 |
| ENSMUSG00000032834 | Pwp2 | PWP2 periodic tryptophan protein homolog (yeast) | 1.02 |
| ENSMUSG00000046791 | Riox1 | ribosomal oxygenase 1 | 1.02 |
| ENSMUSG00000028741 | Mrto4 | mRNA turnover 4, ribosome maturation factor | 1.02 |
| ENSMUSG00000040749 | Siah1b | siah E3 ubiquitin protein ligase 1B | 1.02 |
| ENSMUSG00000035606 | Ky | kyphoscoliosis peptidase | 1.02 |
| ENSMUSG00000081205 | Gm5940 | predicted gene 5940 | 1.02 |
| ENSMUSG00000041057 | Wdr43 | WD repeat domain 43 | 1.01 |
| ENSMUSG00000073676 | Hspe1 | heat shock protein 1 (chaperonin 10) | 1.01 |
| ENSMUSG00000019297 | Nop9 | NOP9 nucleolar protein | 1.01 |
| ENSMUSG00000022391 | Rangap1 | RAN GTPase activating protein 1 | 1.01 |
| ENSMUSG00000001785 | Pwp1 | PWP1 homolog, endonuclein | 1.01 |
| ENSMUSG00000038047 | Haus6 | HAUS augmin-like complex, subunit 6 | 1.01 |
| ENSMUSG00000078713 | Tomm5 | translocase of outer mitochondrial membrane 5 | 1.01 |
| ENSMUSG00000032092 | Mpzl2 | myelin protein zero-like 2 | 1.01 |
| ENSMUSG00000029703 | Lrwd1 | leucine-rich repeats and WD repeat domain containing 1 | 1.01 |
| ENSMUSG00000031388 | Naa10 | N(alpha)-acetyltransferase 10, NatA catalytic subunit | 1.01 |
| ENSMUSG00000032397 | Tipin | timeless interacting protein | 1.01 |
| ENSMUSG00000086290 | Snhg12 | small nucleolar RNA host gene 12 | 1.01 |
| ENSMUSG00000073771 | Btbd19 | BTB (POZ) domain containing 19 | 1.01 |
| ENSMUSG00000035215 | Lsm7 | LSM7 homolog, U6 small nuclear RNA and mRNA degradation associated | 1.01 |
| ENSMUSG00000062981 | Mrpl42 | mitochondrial ribosomal protein L42 | 1.00 |
| ENSMUSG00000001674 | Ddx18 | DEAD (Asp-Glu-Ala-Asp) box polypeptide 18 | 1.00 |
| ENSMUSG00000031146 | Plp2 | proteolipid protein 2 | 1.00 |
| ENSMUSG00000033222 | Ttf2 | transcription termination factor, RNA polymerase II | 1.00 |
| ENSMUSG00000016984 | Etaa1 | Ewing tumor-associated antigen 1 | 1.00 |
| ENSMUSG00000028551 | Cdkn2c | cyclin dependent kinase inhibitor 2C | 1.00 |
| ENSMUSG00000104350 | Gm38244 | predicted gene, 38244 | -1.00 |
| ENSMUSG00000027313 | Chac1 | ChaC, cation transport regulator 1 | -1.00 |
| ENSMUSG00000092185 | Gm20416 | predicted gene 20416 | -1.00 |
| ENSMUSG00000028059 | Arhgef2 | rho/rac guanine nucleotide exchange factor (GEF) 2 | -1.01 |
| ENSMUSG00000005580 | Adcy9 | adenylate cyclase 9 | -1.01 |
| ENSMUSG00000021097 | Clmn | calmin | -1.01 |
| ENSMUSG00000028699 | Tspan1 | tetraspanin 1 | -1.01 |
| ENSMUSG00000002227 | Mov10 | Moloney leukemia virus 10 | -1.01 |
| ENSMUSG00000028755 | Cda | cytidine deaminase | -1.01 |
| ENSMUSG00000057092 | Fxyd3 | FXYD domain-containing ion transport regulator 3 | -1.01 |
| ENSMUSG00000096243 | Gm24265 | predicted gene, 24265 | -1.01 |
| ENSMUSG00000025059 | Gk | glycerol kinase | -1.01 |
| ENSMUSG00000002083 | Bbc3 | BCL2 binding component 3 | -1.01 |
| ENSMUSG00000050592 | Fam78a | family with sequence similarity 78, member A | -1.01 |
| ENSMUSG00000093815 | Gm26444 | predicted gene, 26444 | -1.01 |
| ENSMUSG00000029163 | Emilin1 | elastin microfibril interfacer 1 | -1.01 |
| ENSMUSG00000045382 | Cxcr4 | chemokine (C-X-C motif) receptor 4 | -1.01 |
| ENSMUSG00000083240 | Gm13453 | predicted gene 13453 | -1.01 |
| ENSMUSG00000075420 | Smim6 | small integral membrane protein 6 | -1.02 |
| ENSMUSG00000026628 | Atf3 | activating transcription factor 3 | -1.02 |
| ENSMUSG00000037580 | Gch1 | GTP cyclohydrolase 1 | -1.02 |
| ENSMUSG00000024777 | Ppp2r5b | protein phosphatase 2, regulatory subunit B', beta | -1.02 |
| ENSMUSG00000029162 | Khk | ketohexokinase | -1.02 |
| ENSMUSG00000024937 | Ehbp1l1 | EH domain binding protein 1-like 1 | -1.02 |
| ENSMUSG00000023045 | Soat2 | sterol O-acyltransferase 2 | -1.02 |
| ENSMUSG00000112944 | Gm48885 | predicted gene, 48885 | -1.02 |
| ENSMUSG00000027456 | Sdcbp2 | syndecan binding protein (syntenin) 2 | -1.02 |
| ENSMUSG00000043621 | Ubxn10 | UBX domain protein 10 | -1.02 |
| ENSMUSG00000024462 | Gabbr1 | gamma-aminobutyric acid (GABA) B receptor, 1 | -1.02 |
| ENSMUSG00000089712 | Gm15889 | predicted gene 15889 | -1.02 |
| ENSMUSG00000035969 | Rusc2 | RUN and SH3 domain containing 2 | -1.02 |
| ENSMUSG00000098224 | Gm7985 | predicted gene 7985 | -1.02 |

| | | | |
|---|---|---|---|
| ENSMUSG00000083813 | Gm15502 | predicted gene 15502 | -1.02 |
| ENSMUSG00000052783 | Grk4 | G protein-coupled receptor kinase 4 | -1.02 |
| ENSMUSG00000036036 | Zfp57 | zinc finger protein 57 | -1.02 |
| ENSMUSG00000042417 | Ccno | cyclin O | -1.02 |
| ENSMUSG00000044034 | Npb | neuropeptide B | -1.02 |
| ENSMUSG00000042632 | Pla2g6 | phospholipase A2, group VI | -1.03 |
| ENSMUSG00000013160 | Atp6v0d1 | ATPase, H+ transporting, lysosomal V0 subunit D1 | -1.03 |
| ENSMUSG00000037025 | Foxa2 | forkhead box A2 | -1.03 |
| ENSMUSG00000040111 | Gramd1b | GRAM domain containing 1B | -1.03 |
| ENSMUSG00000036813 | Entpd8 | ectonucleoside triphosphate diphosphohydrolase 8 | -1.03 |
| ENSMUSG00000022445 | Cyp2d26 | cytochrome P450, family 2, subfamily d, polypeptide 26 | -1.03 |
| ENSMUSG00000053113 | Socs3 | suppressor of cytokine signaling 3 | -1.03 |
| ENSMUSG00000033147 | Slc22a15 | solute carrier family 22 (organic anion/cation transporter), member 15 | -1.03 |
| ENSMUSG00000029674 | Limk1 | LIM-domain containing, protein kinase | -1.03 |
| ENSMUSG00000031378 | Abcd1 | ATP-binding cassette, sub-family D (ALD), member 1 | -1.03 |
| ENSMUSG00000055675 | Kbtbd11 | kelch repeat and BTB (POZ) domain containing 11 | -1.03 |
| ENSMUSG00000020774 | Aspa | aspartoacylase | -1.03 |
| ENSMUSG00000022176 | Rem2 | rad and gem related GTP binding protein 2 | -1.03 |
| ENSMUSG00000096349 | Gm22513 | predicted gene, 22513 | -1.03 |
| ENSMUSG00000093880 | Tmem181c-ps | transmembrane protein 181C, pseudogene | -1.03 |
| ENSMUSG00000064215 | Ifi27 | interferon, alpha-inducible protein 27 | -1.03 |
| ENSMUSG00000034687 | Fras1 | Fraser extracellular matrix complex subunit 1 | -1.03 |
| ENSMUSG00000018620 | Mmp20 | matrix metallopeptidase 20 (enamelysin) | -1.03 |
| ENSMUSG00000034156 | Tspoap1 | TSPO associated protein 1 | -1.03 |
| ENSMUSG00000109679 | Gm45342 | predicted gene 45342 | -1.03 |
| ENSMUSG00000040134 | Rdh7 | retinol dehydrogenase 7 | -1.03 |
| ENSMUSG00000112955 | Gm48889 | predicted gene, 48889 | -1.03 |
| ENSMUSG00000060441 | Trim5 | tripartite motif-containing 5 | -1.03 |
| ENSMUSG00000021363 | Mak | male germ cell-associated kinase | -1.03 |
| ENSMUSG00000107182 | Gm43268 | predicted gene 43268 | -1.03 |
| ENSMUSG00000086544 | Chn1os3 | chimerin 1, opposite strand 3 | -1.03 |
| ENSMUSG00000084154 | Gm15644 | predicted gene 15644 | -1.03 |
| ENSMUSG00000030275 | Etnk1 | ethanolamine kinase 1 | -1.04 |
| ENSMUSG00000032899 | Styk1 | serine/threonine/tyrosine kinase 1 | -1.04 |
| ENSMUSG00000038418 | Egr1 | early growth response 1 | -1.04 |
| ENSMUSG00000020777 | Acox1 | acyl-Coenzyme A oxidase 1, palmitoyl | -1.04 |
| ENSMUSG00000023055 | Calcoco1 | calcium binding and coiled coil domain 1 | -1.04 |
| ENSMUSG00000032098 | Treh | trehalase (brush-border membrane glycoprotein) | -1.04 |
| ENSMUSG00000070287 | Slc35g2 | solute carrier family 35, member G2 | -1.04 |
| ENSMUSG00000051427 | Ccdc157 | coiled-coil domain containing 157 | -1.04 |
| ENSMUSG00000032105 | Pdzd3 | PDZ domain containing 3 | -1.04 |
| ENSMUSG00000043789 | Vwce | von Willebrand factor C and EGF domains | -1.04 |
| ENSMUSG00000034220 | Gpc1 | glypican 1 | -1.04 |
| ENSMUSG00000031586 | Rbpms | RNA binding protein gene with multiple splicing | -1.04 |
| ENSMUSG00000041298 | Katnal1 | katanin p60 subunit A-like 1 | -1.04 |
| ENSMUSG00000108181 | C030015A19Rik | RIKEN cDNA C030015A19 gene | -1.04 |
| ENSMUSG00000072915 | Gm12258 | predicted gene 12258 | -1.04 |
| ENSMUSG00000037348 | Paqr7 | progestin and adipoQ receptor family member VII | -1.04 |
| ENSMUSG00000021057 | Akap5 | A kinase (PRKA) anchor protein 5 | -1.04 |
| ENSMUSG00000055862 | Izumo4 | IZUMO family member 4 | -1.04 |
| ENSMUSG00000024778 | Fas | Fas (TNF receptor superfamily member 6) | -1.04 |
| ENSMUSG00000021675 | F2rl2 | coagulation factor II (thrombin) receptor-like 2 | -1.04 |
| ENSMUSG00000000901 | Mmp11 | matrix metallopeptidase 11 | -1.04 |
| ENSMUSG00000029195 | Klb | klotho beta | -1.04 |
| ENSMUSG00000096992 | Gm26788 | predicted gene, 26788 | -1.04 |
| ENSMUSG00000114150 | Gm46367 | predicted gene, 46367 | -1.04 |
| ENSMUSG00000116180 | Gm49492 | predicted gene, 49492 | -1.04 |
| ENSMUSG00000040462 | Os9 | amplified in osteosarcoma | -1.05 |
| ENSMUSG00000031532 | Saraf | store-operated calcium entry-associated regulatory factor | -1.05 |
| ENSMUSG00000036246 | Gmip | Gem-interacting protein | -1.05 |
| ENSMUSG00000020284 | 1810043G02Rik | RIKEN cDNA 1810043G02 gene | -1.05 |
| ENSMUSG00000107225 | Gm43637 | predicted gene 43637 | -1.05 |
| ENSMUSG00000037139 | Myom3 | myomesin family, member 3 | -1.05 |
| ENSMUSG00000018931 | Natd1 | N-acetyltransferase domain containing 1 | -1.05 |
| ENSMUSG00000050957 | Insl6 | insulin-like 6 | -1.05 |
| ENSMUSG00000017314 | Mpp2 | membrane protein, palmitoylated 2 (MAGUK p55 subfamily member 2) | -1.05 |
| ENSMUSG00000097006 | 9530082P21Rik | RIKEN cDNA 9530082P21 gene | -1.05 |
| ENSMUSG00000107741 | Gm2011 | predicted gene 2011 | -1.05 |
| ENSMUSG00000096924 | Gm26824 | predicted gene, 26824 | -1.05 |
| ENSMUSG00000035279 | Ssc5d | scavenger receptor cysteine rich family, 5 domains | -1.05 |
| ENSMUSG00000020120 | Plek | pleckstrin | -1.05 |
| ENSMUSG00000044505 | Lingo4 | leucine rich repeat and Ig domain containing 4 | -1.05 |
| ENSMUSG00000054901 | Arhgef33 | Rho guanine nucleotide exchange factor (GEF) 33 | -1.05 |

| | | | |
|---|---|---|---|
| ENSMUSG00000038507 | Parp12 | poly (ADP-ribose) polymerase family, member 12 | -1.06 |
| ENSMUSG00000073420 | Btnl5-ps | butyrophilin-like 5, pseudogene | -1.06 |
| ENSMUSG00000022565 | Plec | plectin | -1.06 |
| ENSMUSG00000034312 | Iqsec1 | IQ motif and Sec7 domain 1 | -1.06 |
| ENSMUSG00000044080 | S100a1 | S100 calcium binding protein A1 | -1.06 |
| ENSMUSG00000041548 | Hspb8 | heat shock protein 8 | -1.06 |
| ENSMUSG00000019577 | Pdk4 | pyruvate dehydrogenase kinase, isoenzyme 4 | -1.06 |
| ENSMUSG00000027524 | Edn3 | endothelin 3 | -1.06 |
| ENSMUSG00000099021 | Rn7s1 | 7S RNA 1 | -1.06 |
| ENSMUSG00000113119 | Gm48883 | predicted gene, 48883 | -1.06 |
| ENSMUSG00000035692 | Isg15 | ISG15 ubiquitin-like modifier | -1.06 |
| ENSMUSG00000115918 | A930001M01Rik | RIKEN cDNA A930001M01 gene | -1.06 |
| ENSMUSG00000103037 | Pcdhgb1 | protocadherin gamma subfamily B, 1 | -1.06 |
| ENSMUSG00000053985 | Zfp14 | zinc finger protein 14 | -1.06 |
| ENSMUSG00000026347 | Tmem163 | transmembrane protein 163 | -1.06 |
| ENSMUSG00000104830 | 5830487J09Rik | RIKEN cDNA 5830487J09 gene | -1.06 |
| ENSMUSG00000026475 | Rgs16 | regulator of G-protein signaling 16 | -1.06 |
| ENSMUSG00000097855 | A930007I19Rik | RIKEN cDNA A930007I19 gene | -1.06 |
| ENSMUSG00000113076 | Gm47088 | predicted gene, 47088 | -1.06 |
| ENSMUSG00000096883 | Shisa8 | shisa family member 8 | -1.06 |
| ENSMUSG00000038528 | Mfsd4b5 | major facilitator superfamily domain containing 4B5 | -1.06 |
| ENSMUSG00000022783 | Spag6l | sperm associated antigen 6-like | -1.06 |
| ENSMUSG00000033898 | Cfhr2 | complement factor H-related 2 | -1.06 |
| ENSMUSG00000032198 | Dock6 | dedicator of cytokinesis 6 | -1.07 |
| ENSMUSG00000018750 | Zbtb4 | zinc finger and BTB domain containing 4 | -1.07 |
| ENSMUSG00000025192 | Entpd7 | ectonucleoside triphosphate diphosphohydrolase 7 | -1.07 |
| ENSMUSG00000033538 | Casp4 | caspase 4, apoptosis-related cysteine peptidase | -1.07 |
| ENSMUSG00000033308 | Dpyd | dihydropyrimidine dehydrogenase | -1.07 |
| ENSMUSG00000022098 | Bmp1 | bone morphogenetic protein 1 | -1.07 |
| ENSMUSG00000001604 | Tcea3 | transcription elongation factor A (SII), 3 | -1.07 |
| ENSMUSG00000025429 | Pstpip2 | proline-serine-threonine phosphatase-interacting protein 2 | -1.07 |
| ENSMUSG00000053702 | Nebl | nebulette | -1.07 |
| ENSMUSG00000020407 | Upp1 | uridine phosphorylase 1 | -1.07 |
| ENSMUSG00000054708 | Ankrd24 | ankyrin repeat domain 24 | -1.07 |
| ENSMUSG00000038298 | Pdzk1 | PDZ domain containing 1 | -1.07 |
| ENSMUSG00000109585 | Gm45358 | predicted gene 45358 | -1.07 |
| ENSMUSG00000085873 | Ttc39aos1 | Ttc39a opposite strand RNA 1 | -1.07 |
| ENSMUSG00000020032 | Nuak1 | NUAK family, SNF1-like kinase, 1 | -1.07 |
| ENSMUSG00000110187 | Gm45496 | predicted gene 45496 | -1.07 |
| ENSMUSG00000002107 | Celf2 | CUGBP, Elav-like family member 2 | -1.07 |
| ENSMUSG00000104576 | F830115B05Rik | RIKEN cDNA F830115B05 gene | -1.07 |
| ENSMUSG00000049536 | Tceal1 | transcription elongation factor A (SII)-like 1 | -1.07 |
| ENSMUSG00000038860 | Garnl3 | GTPase activating RANGAP domain-like 3 | -1.07 |
| ENSMUSG00000073608 | Gal3st2c | galactose-3-O-sulfotransferase 2C | -1.07 |
| ENSMUSG00000059852 | Kcng2 | potassium voltage-gated channel, subfamily G, member 2 | -1.07 |
| ENSMUSG00000105212 | Gm43557 | predicted gene 43557 | -1.07 |
| ENSMUSG00000085709 | Gm14211 | predicted gene 14211 | -1.07 |
| ENSMUSG00000015291 | Gdi1 | guanosine diphosphate (GDP) dissociation inhibitor 1 | -1.08 |
| ENSMUSG00000023951 | Vegfa | vascular endothelial growth factor A | -1.08 |
| ENSMUSG00000020577 | Tspan13 | tetraspanin 13 | -1.08 |
| ENSMUSG00000028555 | Ttc39a | tetratricopeptide repeat domain 39A | -1.08 |
| ENSMUSG00000002885 | Adgre5 | adhesion G protein-coupled receptor E5 | -1.08 |
| ENSMUSG00000014361 | Mertk | c-mer proto-oncogene tyrosine kinase | -1.08 |
| ENSMUSG00000045038 | Prkce | protein kinase C, epsilon | -1.08 |
| ENSMUSG00000025648 | Pfkfb4 | 6-phosphofructo-2-kinase/fructose-2,6-biphosphatase 4 | -1.08 |
| ENSMUSG00000030747 | Dgat2 | diacylglycerol O-acyltransferase 2 | -1.08 |
| ENSMUSG00000026773 | Pfkfb3 | 6-phosphofructo-2-kinase/fructose-2,6-biphosphatase 3 | -1.08 |
| ENSMUSG00000019558 | Slc6a8 | solute carrier family 6 (neurotransmitter transporter, creatine), member 8 | -1.08 |
| ENSMUSG00000102752 | Gm7694 | predicted gene 7694 | -1.08 |
| ENSMUSG00000028542 | Slc6a9 | solute carrier family 6 (neurotransmitter transporter, glycine), member 9 | -1.08 |
| ENSMUSG00000031919 | Tmed6 | transmembrane p24 trafficking protein 6 | -1.08 |
| ENSMUSG00000073889 | Il11ra1 | interleukin 11 receptor, alpha chain 1 | -1.08 |
| ENSMUSG00000027078 | Ube2l6 | ubiquitin-conjugating enzyme E2L 6 | -1.08 |
| ENSMUSG00000071551 | Akr1c19 | aldo-keto reductase family 1, member C19 | -1.08 |
| ENSMUSG00000086844 | B230206H07Rik | RIKEN cDNA B230206H07 gene | -1.08 |
| ENSMUSG00000031844 | Hsd17b2 | hydroxysteroid (17-beta) dehydrogenase 2 | -1.08 |
| ENSMUSG00000032726 | Bmp8a | bone morphogenetic protein 8a | -1.08 |
| ENSMUSG00000049612 | Omg | oligodendrocyte myelin glycoprotein | -1.08 |
| ENSMUSG00000057137 | Tmem140 | transmembrane protein 140 | -1.08 |
| ENSMUSG00000055214 | Pld5 | phospholipase D family, member 5 | -1.08 |
| ENSMUSG00000086825 | Gm15675 | predicted gene 15675 | -1.08 |
| ENSMUSG00000074628 | Tldc2 | TBC/LysM associated domain containing 2 | -1.08 |
| ENSMUSG00000106547 | B230303O12Rik | RIKEN cDNA B230303O12 gene | -1.08 |

| | | | |
|---|---|---|---|
| ENSMUSG00000109899 | Gm6012 | predicted gene 6012 | -1.08 |
| ENSMUSG00000081516 | Gm12470 | predicted gene 12470 | -1.08 |
| ENSMUSG00000049515 | Espnl | espin-like | -1.08 |
| ENSMUSG00000018830 | Myh11 | myosin, heavy polypeptide 11, smooth muscle | -1.08 |
| ENSMUSG00000097294 | Gm26888 | predicted gene, 26888 | -1.08 |
| ENSMUSG00000036862 | Dchs1 | dachsous cadherin related 1 | -1.08 |
| ENSMUSG00000001542 | Ell2 | elongation factor RNA polymerase II 2 | -1.09 |
| ENSMUSG00000048970 | C1galt1c1 | C1GALT1-specific chaperone 1 | -1.09 |
| ENSMUSG00000056116 | H2-T22 | histocompatibility 2, T region locus 22 | -1.09 |
| ENSMUSG00000028413 | B4galt1 | UDP-Gal:betaGlcNAc beta 1,4- galactosyltransferase, polypeptide 1 | -1.09 |
| ENSMUSG00000015202 | Cnksr3 | Cnksr family member 3 | -1.09 |
| ENSMUSG00000012123 | Crybg2 | crystallin beta-gamma domain containing 2 | -1.09 |
| ENSMUSG00000041608 | Entpd3 | ectonucleoside triphosphate diphosphohydrolase 3 | -1.09 |
| ENSMUSG00000102440 | Pcdhga9 | protocadherin gamma subfamily A, 9 | -1.09 |
| ENSMUSG00000044681 | Cnpy1 | canopy FGF signaling regulator 1 | -1.09 |
| ENSMUSG00000089960 | Ugt1a1 | UDP glucuronosyltransferase 1 family, polypeptide A1 | -1.09 |
| ENSMUSG00000054404 | Slfn5 | schlafen 5 | -1.09 |
| ENSMUSG00000112459 | Gm47689 | predicted gene, 47689 | -1.09 |
| ENSMUSG00000112303 | Gm48759 | predicted gene, 48759 | -1.09 |
| ENSMUSG00000046242 | Nme9 | NME/NM23 family member 9 | -1.09 |
| ENSMUSG00000051736 | Fam229b | family with sequence similarity 229, member B | -1.09 |
| ENSMUSG00000096107 | Gm16505 | predicted gene 16505 | -1.09 |
| ENSMUSG00000022844 | Pdia5 | protein disulfide isomerase associated 5 | -1.10 |
| ENSMUSG00000021360 | Gcnt2 | glucosaminyl (N-acetyl) transferase 2, I-branching enzyme | -1.10 |
| ENSMUSG00000038781 | Stap2 | signal transducing adaptor family member 2 | -1.10 |
| ENSMUSG00000099746 | Ppnr | per-pentamer repeat gene | -1.10 |
| ENSMUSG00000022540 | Rogdi | rogdi homolog | -1.10 |
| ENSMUSG00000094830 | n-R5s194 | nuclear encoded rRNA 5S 194 | -1.10 |
| ENSMUSG00000024774 | Ankrd22 | ankyrin repeat domain 22 | -1.10 |
| ENSMUSG00000028631 | Kcnq4 | potassium voltage-gated channel, subfamily Q, member 4 | -1.10 |
| ENSMUSG00000061451 | Tmem151a | transmembrane protein 151A | -1.10 |
| ENSMUSG00000030882 | Dnhd1 | dynein heavy chain domain 1 | -1.10 |
| ENSMUSG00000026678 | Rgs5 | regulator of G-protein signaling 5 | -1.10 |
| ENSMUSG00000030589 | Rasgrp4 | RAS guanyl releasing protein 4 | -1.10 |
| ENSMUSG00000060988 | Galnt13 | polypeptide N-acetylgalactosaminyltransferase 13 | -1.10 |
| ENSMUSG00000036760 | Kcnk9 | potassium channel, subfamily K, member 9 | -1.10 |
| ENSMUSG00000104429 | 2610034O05Rik | RIKEN cDNA 2610034O05 gene | -1.10 |
| ENSMUSG00000043556 | Fbxl7 | F-box and leucine-rich repeat protein 7 | -1.10 |
| ENSMUSG00000078627 | 40238 | membrane-associated ring finger (C3HC4) 10 | -1.10 |
| ENSMUSG00000039050 | Osbpl2 | oxysterol binding protein-like 2 | -1.11 |
| ENSMUSG00000037012 | Hk1 | hexokinase 1 | -1.11 |
| ENSMUSG00000024066 | Xdh | xanthine dehydrogenase | -1.11 |
| ENSMUSG00000035847 | Ids | iduronate 2-sulfatase | -1.11 |
| ENSMUSG00000035699 | Slc51a | solute carrier family 51, alpha subunit | -1.11 |
| ENSMUSG00000041845 | Rhod | ras homolog family member D | -1.11 |
| ENSMUSG00000095915 | n-R5s115 | nuclear encoded rRNA 5S 115 | -1.11 |
| ENSMUSG00000097615 | Gm2061 | predicted gene 2061 | -1.11 |
| ENSMUSG00000111539 | Gm48372 | predicted gene, 48372 | -1.11 |
| ENSMUSG00000027075 | Slc43a1 | solute carrier family 43, member 1 | -1.11 |
| ENSMUSG00000052605 | Isoc2b | isochorismatase domain containing 2b | -1.11 |
| ENSMUSG00000046733 | Gprc5a | G protein-coupled receptor, family C, group 5, member A | -1.11 |
| ENSMUSG00000087691 | Gm15674 | predicted gene 15674 | -1.11 |
| ENSMUSG00000116862 | AC114990.2 | novel transcript | -1.11 |
| ENSMUSG00000049670 | Morn4 | MORN repeat containing 4 | -1.11 |
| ENSMUSG00000020486 | 38231 | septin 4 | -1.11 |
| ENSMUSG00000081683 | Fzd10 | frizzled class receptor 10 | -1.11 |
| ENSMUSG00000104346 | Pcdhga3 | protocadherin gamma subfamily A, 3 | -1.11 |
| ENSMUSG00000037784 | Dzip1l | DAZ interacting protein 1-like | -1.11 |
| ENSMUSG00000087143 | A830082K12Rik | RIKEN cDNA A830082K12 gene | -1.11 |
| ENSMUSG00000047495 | Dlgap2 | DLG associated protein 2 | -1.11 |
| ENSMUSG00000081854 | Gm11447 | predicted gene 11447 | -1.11 |
| ENSMUSG00000106478 | Gm36551 | predicted gene, 36551 | -1.11 |
| ENSMUSG00000025469 | Msx3 | msh homeobox 3 | -1.11 |
| ENSMUSG00000112532 | Gm36283 | predicted gene, 36283 | -1.11 |
| ENSMUSG00000024892 | Pcx | pyruvate carboxylase | -1.12 |
| ENSMUSG00000028307 | Aldob | aldolase B, fructose-bisphosphate | -1.12 |
| ENSMUSG00000032596 | Uba7 | ubiquitin-like modifier activating enzyme 7 | -1.12 |
| ENSMUSG00000038963 | Slco4a1 | solute carrier organic anion transporter family, member 4a1 | -1.12 |
| ENSMUSG00000022090 | Pdlim2 | PDZ and LIM domain 2 | -1.12 |
| ENSMUSG00000024885 | Aldh3b1 | aldehyde dehydrogenase 3 family, member B1 | -1.12 |
| ENSMUSG00000042684 | Npl | N-acetylneuraminate pyruvate lyase | -1.12 |
| ENSMUSG00000051146 | Camk2n2 | calcium/calmodulin-dependent protein kinase II inhibitor 2 | -1.12 |
| ENSMUSG00000042942 | Greb1l | growth regulation by estrogen in breast cancer-like | -1.12 |

| | | | |
|---|---|---|---|
| ENSMUSG00000028076 | Cd1d1 | CD1d1 antigen | -1.12 |
| ENSMUSG00000037613 | Tnfrsf23 | tumor necrosis factor receptor superfamily, member 23 | -1.12 |
| ENSMUSG00000034648 | Lrrn1 | leucine rich repeat protein 1, neuronal | -1.12 |
| ENSMUSG00000043811 | Rtn4r | reticulon 4 receptor | -1.12 |
| ENSMUSG00000019699 | Akt3 | thymoma viral proto-oncogene 3 | -1.12 |
| ENSMUSG00000006542 | Prkag3 | protein kinase, AMP-activated, gamma 3 non-catalytic subunit | -1.12 |
| ENSMUSG00000086507 | Adap2os | ArfGAP with dual PH domains 2, opposite strand | -1.12 |
| ENSMUSG00000111951 | Gm48755 | predicted gene, 48755 | -1.12 |
| ENSMUSG00000030402 | Ppm1n | protein phosphatase, Mg2+/Mn2+ dependent, 1N (putative) | -1.12 |
| ENSMUSG00000103194 | Gm37643 | predicted gene, 37643 | -1.12 |
| ENSMUSG00000051652 | Lrrc3 | leucine rich repeat containing 3 | -1.12 |
| ENSMUSG00000030651 | Art2b | ADP-ribosyltransferase 2b | -1.12 |
| ENSMUSG00000033510 | Otud7a | OTU domain containing 7A | -1.12 |
| ENSMUSG00000034377 | Tulp4 | tubby like protein 4 | -1.13 |
| ENSMUSG00000042207 | Kdm5b | lysine (K)-specific demethylase 5B | -1.13 |
| ENSMUSG00000027215 | Cd82 | CD82 antigen | -1.13 |
| ENSMUSG00000028207 | Asph | aspartate-beta-hydroxylase | -1.13 |
| ENSMUSG00000025357 | Dgka | diacylglycerol kinase, alpha | -1.13 |
| ENSMUSG00000037280 | Galnt6 | polypeptide N-acetylgalactosaminyltransferase 6 | -1.13 |
| ENSMUSG00000023952 | Gtpbp2 | GTP binding protein 2 | -1.13 |
| ENSMUSG00000029470 | P2rx4 | purinergic receptor P2X, ligand-gated ion channel 4 | -1.13 |
| ENSMUSG00000021749 | Oit1 | oncoprotein induced transcript 1 | -1.13 |
| ENSMUSG00000028158 | Mttp | microsomal triglyceride transfer protein | -1.13 |
| ENSMUSG00000024292 | Cyp4f14 | cytochrome P450, family 4, subfamily f, polypeptide 14 | -1.13 |
| ENSMUSG00000048911 | Rnf24 | ring finger protein 24 | -1.13 |
| ENSMUSG00000031886 | Ces2e | carboxylesterase 2E | -1.13 |
| ENSMUSG00000026354 | Lct | lactase | -1.13 |
| ENSMUSG00000004552 | Ctse | cathepsin E | -1.13 |
| ENSMUSG00000038751 | Ptk6 | PTK6 protein tyrosine kinase 6 | -1.13 |
| ENSMUSG00000042523 | Dnal1 | dynein, axonemal, light chain 1 | -1.13 |
| ENSMUSG00000038843 | Gcnt1 | glucosaminyl (N-acetyl) transferase 1, core 2 | -1.13 |
| ENSMUSG00000033029 | 1700088E04Rik | RIKEN cDNA 1700088E04 gene | -1.13 |
| ENSMUSG00000006958 | Chrd | chordin | -1.13 |
| ENSMUSG00000103897 | Pcdhga8 | protocadherin gamma subfamily A, 8 | -1.13 |
| ENSMUSG00000086847 | Tbx3os2 | T-box 3, opposite strand 2 | -1.13 |
| ENSMUSG00000102742 | Pcdhga11 | protocadherin gamma subfamily A, 11 | -1.13 |
| ENSMUSG00000020701 | Tmem132e | transmembrane protein 132E | -1.13 |
| ENSMUSG00000010342 | Tex14 | testis expressed gene 14 | -1.13 |
| ENSMUSG00000047298 | Kcnv2 | potassium channel, subfamily V, member 2 | -1.13 |
| ENSMUSG00000020432 | Tcn2 | transcobalamin 2 | -1.14 |
| ENSMUSG00000040929 | Rfx3 | regulatory factor X, 3 (influences HLA class II expression) | -1.14 |
| ENSMUSG00000037995 | Igsf9 | immunoglobulin superfamily, member 9 | -1.14 |
| ENSMUSG00000024854 | Pold4 | polymerase (DNA-directed), delta 4 | -1.14 |
| ENSMUSG00000034449 | Dhrs11 | dehydrogenase/reductase (SDR family) member 11 | -1.14 |
| ENSMUSG00000008398 | Elk3 | ELK3, member of ETS oncogene family | -1.14 |
| ENSMUSG00000056602 | Fry | FRY microtubule binding protein | -1.14 |
| ENSMUSG00000017390 | Aldoc | aldolase C, fructose-bisphosphate | -1.14 |
| ENSMUSG00000029919 | Hpgds | hematopoietic prostaglandin D synthase | -1.14 |
| ENSMUSG00000071847 | Apcdd1 | adenomatosis polyposis coli down-regulated 1 | -1.14 |
| ENSMUSG00000037868 | Egr2 | early growth response 2 | -1.14 |
| ENSMUSG00000062064 | Slc2a7 | solute carrier family 2 (facilitated glucose transporter), member 7 | -1.14 |
| ENSMUSG00000079339 | Ifit1bl1 | interferon induced protein with tetratricpeptide repeats 1B like 1 | -1.14 |
| ENSMUSG00000047473 | Zfp30 | zinc finger protein 30 | -1.14 |
| ENSMUSG00000104612 | Gm42449 | predicted gene 42449 | -1.14 |
| ENSMUSG00000003992 | Ssbp2 | single-stranded DNA binding protein 2 | -1.14 |
| ENSMUSG00000091898 | Tnnc1 | troponin C, cardiac/slow skeletal | -1.14 |
| ENSMUSG00000033498 | Strc | stereocilin | -1.14 |
| ENSMUSG00000009687 | Fxyd5 | FXYD domain-containing ion transport regulator 5 | -1.14 |
| ENSMUSG00000087938 | Gm26049 | predicted gene, 26049 | -1.14 |
| ENSMUSG00000079497 | Gm13420 | predicted gene 13420 | -1.14 |
| ENSMUSG00000025221 | Kcnip2 | Kv channel-interacting protein 2 | -1.14 |
| ENSMUSG00000021091 | Serpina3n | serine (or cysteine) peptidase inhibitor, clade A, member 3N | -1.14 |
| ENSMUSG00000113370 | Gm46348 | predicted gene, 46348 | -1.14 |
| ENSMUSG00000033487 | Fndc3a | fibronectin type III domain containing 3A | -1.15 |
| ENSMUSG00000039286 | Fndc3b | fibronectin type III domain containing 3B | -1.15 |
| ENSMUSG00000020376 | Rnf130 | ring finger protein 130 | -1.15 |
| ENSMUSG00000001225 | Slc26a3 | solute carrier family 26, member 3 | -1.15 |
| ENSMUSG00000058624 | Gda | guanine deaminase | -1.15 |
| ENSMUSG00000031840 | Rab3a | RAB3A, member RAS oncogene family | -1.15 |
| ENSMUSG00000048191 | Muc6 | mucin 6, gastric | -1.15 |
| ENSMUSG00000074115 | Saa1 | serum amyloid A 1 | -1.15 |
| ENSMUSG00000103472 | Pcdhga7 | protocadherin gamma subfamily A, 7 | -1.15 |
| ENSMUSG00000097990 | Gm19557 | predicted gene, 19557 | -1.15 |

| | | | |
|---|---|---|---|
| ENSMUSG00000113918 | Gm6566 | predicted gene 6566 | -1.15 |
| ENSMUSG00000002769 | Gnmt | glycine N-methyltransferase | -1.15 |
| ENSMUSG00000072572 | Slc39a2 | solute carrier family 39 (zinc transporter), member 2 | -1.15 |
| ENSMUSG00000114302 | Gm48683 | predicted gene, 48683 | -1.15 |
| ENSMUSG00000026247 | Ecel1 | endothelin converting enzyme-like 1 | -1.15 |
| ENSMUSG00000092116 | Gm10320 | predicted pseudogene 10320 | -1.15 |
| ENSMUSG00000016942 | Tmprss6 | transmembrane serine protease 6 | -1.15 |
| ENSMUSG00000073295 | Nudt11 | nudix (nucleoside diphosphate linked moiety X)-type motif 11 | -1.15 |
| ENSMUSG00000022270 | Retreg1 | reticulophagy regulator 1 | -1.16 |
| ENSMUSG00000020246 | Hcfc2 | host cell factor C2 | -1.16 |
| ENSMUSG00000039782 | Cpeb2 | cytoplasmic polyadenylation element binding protein 2 | -1.16 |
| ENSMUSG00000024887 | Asah2 | N-acylsphingosine amidohydrolase 2 | -1.16 |
| ENSMUSG00000039270 | Megf9 | multiple EGF-like-domains 9 | -1.16 |
| ENSMUSG00000025467 | Prap1 | proline-rich acidic protein 1 | -1.16 |
| ENSMUSG00000039062 | Anpep | alanyl (membrane) aminopeptidase | -1.16 |
| ENSMUSG00000024731 | Ms4a10 | membrane-spanning 4-domains, subfamily A, member 10 | -1.16 |
| ENSMUSG00000046410 | Kcnk6 | potassium inwardly-rectifying channel, subfamily K, member 6 | -1.16 |
| ENSMUSG00000024140 | Epas1 | endothelial PAS domain protein 1 | -1.16 |
| ENSMUSG00000032515 | Csrnp1 | cysteine-serine-rich nuclear protein 1 | -1.16 |
| ENSMUSG00000016496 | Cd274 | CD274 antigen | -1.16 |
| ENSMUSG00000027297 | Ltk | leukocyte tyrosine kinase | -1.16 |
| ENSMUSG00000028909 | Ptpru | protein tyrosine phosphatase, receptor type, U | -1.16 |
| ENSMUSG00000040856 | Dlk1 | delta like non-canonical Notch ligand 1 | -1.16 |
| ENSMUSG00000044726 | Erich5 | glutamate rich 5 | -1.16 |
| ENSMUSG00000068859 | Sp9 | trans-acting transcription factor 9 | -1.16 |
| ENSMUSG00000105986 | Gm43065 | predicted gene 43065 | -1.16 |
| ENSMUSG00000015568 | Lpl | lipoprotein lipase | -1.16 |
| ENSMUSG00000038070 | Cntln | centlein, centrosomal protein | -1.16 |
| ENSMUSG00000087675 | Gm11762 | predicted gene 11762 | -1.16 |
| ENSMUSG00000078439 | Smim24 | small integral membrane protein 24 | -1.17 |
| ENSMUSG00000043831 | Lysmd4 | LysM, putative peptidoglycan-binding, domain containing 4 | -1.17 |
| ENSMUSG00000015647 | Lama5 | laminin, alpha 5 | -1.17 |
| ENSMUSG00000025140 | Pycr1 | pyrroline-5-carboxylate reductase 1 | -1.17 |
| ENSMUSG00000049971 | Glt1d1 | glycosyltransferase 1 domain containing 1 | -1.17 |
| ENSMUSG00000068349 | Gml | glycosylphosphatidylinositol anchored molecule like | -1.17 |
| ENSMUSG00000022683 | Pla2g10 | phospholipase A2, group X | -1.17 |
| ENSMUSG00000062743 | Zfp677 | zinc finger protein 677 | -1.17 |
| ENSMUSG00000027377 | Mall | mal, T cell differentiation protein-like | -1.17 |
| ENSMUSG00000001248 | Gramd1a | GRAM domain containing 1A | -1.17 |
| ENSMUSG00000067049 | Unc93a | unc-93 homolog A | -1.17 |
| ENSMUSG00000087242 | C78197 | expressed sequence C78197 | -1.17 |
| ENSMUSG00000095440 | Fignl2 | fidgetin-like 2 | -1.17 |
| ENSMUSG00000106417 | Gm42628 | predicted gene 42628 | -1.17 |
| ENSMUSG00000052469 | Tcp10c | t-complex protein 10c | -1.17 |
| ENSMUSG00000086708 | Gm15577 | predicted gene 15577 | -1.17 |
| ENSMUSG00000026223 | Itm2c | integral membrane protein 2C | -1.18 |
| ENSMUSG00000036151 | Tm6sf2 | transmembrane 6 superfamily member 2 | -1.18 |
| ENSMUSG00000030650 | Tmc5 | transmembrane channel-like gene family 5 | -1.18 |
| ENSMUSG00000062638 | Btnl1 | butyrophilin-like 1 | -1.18 |
| ENSMUSG00000026170 | Cyp27a1 | cytochrome P450, family 27, subfamily a, polypeptide 1 | -1.18 |
| ENSMUSG00000079445 | B3gnt7 | UDP-GlcNAc:betaGal beta-1,3-N-acetylglucosaminyltransferase 7 | -1.18 |
| ENSMUSG00000042743 | Sgtb | small glutamine-rich tetratricopeptide repeat (TPR)-containing, beta | -1.18 |
| ENSMUSG00000061878 | Sphk1 | sphingosine kinase 1 | -1.18 |
| ENSMUSG00000039339 | Mfsd4b2 | major facilitator superfamily domain containing 4B2 | -1.18 |
| ENSMUSG00000022857 | Tmprss15 | transmembrane protease, serine 15 | -1.18 |
| ENSMUSG00000056486 | Chn1 | chimerin 1 | -1.18 |
| ENSMUSG00000108991 | Gm45807 | predicted gene 45807 | -1.18 |
| ENSMUSG00000046449 | Nexmif | neurite extension and migration factor | -1.18 |
| ENSMUSG00000031133 | Arhgef6 | Rac/Cdc42 guanine nucleotide exchange factor (GEF) 6 | -1.18 |
| ENSMUSG00000085500 | Gm16976 | predicted gene, 16976 | -1.18 |
| ENSMUSG00000112548 | Gm47338 | predicted gene, 47338 | -1.18 |
| ENSMUSG00000097834 | Gm26911 | predicted gene, 26911 | -1.18 |
| ENSMUSG00000038894 | Irs2 | insulin receptor substrate 2 | -1.19 |
| ENSMUSG00000029119 | Man2b2 | mannosidase 2, alpha B2 | -1.19 |
| ENSMUSG00000042428 | Mgat3 | mannoside acetylglucosaminyltransferase 3 | -1.19 |
| ENSMUSG00000024340 | Btnl2 | butyrophilin-like 2 | -1.19 |
| ENSMUSG00000004661 | Arid3b | AT rich interactive domain 3B (BRIGHT-like) | -1.19 |
| ENSMUSG00000022043 | Trim35 | tripartite motif-containing 35 | -1.19 |
| ENSMUSG00000042249 | Grk3 | G protein-coupled receptor kinase 3 | -1.19 |
| ENSMUSG00000022900 | Ildr1 | immunoglobulin-like domain containing receptor 1 | -1.19 |
| ENSMUSG00000074063 | Osgin1 | oxidative stress induced growth inhibitor 1 | -1.19 |
| ENSMUSG00000083138 | Cyp4a29 | cytochrome P450, family 4, subfamily a, polypeptide 29 | -1.19 |
| ENSMUSG00000025017 | Pik3ap1 | phosphoinositide-3-kinase adaptor protein 1 | -1.19 |

| | | | |
|---|---|---|---|
| ENSMUSG00000026452 | Syt2 | synaptotagmin II | -1.19 |
| ENSMUSG00000054417 | Cyp3a44 | cytochrome P450, family 3, subfamily a, polypeptide 44 | -1.19 |
| ENSMUSG00000038540 | Tmc3 | transmembrane channel-like gene family 3 | -1.19 |
| ENSMUSG00000036587 | Fut7 | fucosyltransferase 7 | -1.19 |
| ENSMUSG00000024907 | Gal | galanin | -1.19 |
| ENSMUSG00000038312 | Edem2 | ER degradation enhancer, mannosidase alpha-like 2 | -1.20 |
| ENSMUSG00000041132 | N4bp2l1 | NEDD4 binding protein 2-like 1 | -1.20 |
| ENSMUSG00000031570 | Plpp5 | phospholipid phosphatase 5 | -1.20 |
| ENSMUSG00000005107 | Slc2a9 | solute carrier family 2 (facilitated glucose transporter), member 9 | -1.20 |
| ENSMUSG00000030223 | Ptpro | protein tyrosine phosphatase, receptor type, O | -1.20 |
| ENSMUSG00000017697 | Ada | adenosine deaminase | -1.20 |
| ENSMUSG00000028214 | Gem | GTP binding protein (gene overexpressed in skeletal muscle) | -1.20 |
| ENSMUSG00000027339 | Rassf2 | Ras association (RalGDS/AF-6) domain family member 2 | -1.20 |
| ENSMUSG00000112700 | Gm48757 | predicted gene, 48757 | -1.20 |
| ENSMUSG00000111528 | Gm39460 | predicted gene, 39460 | -1.20 |
| ENSMUSG00000049001 | Ndnf | neuron-derived neurotrophic factor | -1.20 |
| ENSMUSG00000027559 | Car3 | carbonic anhydrase 3 | -1.20 |
| ENSMUSG00000111731 | Gm48727 | predicted gene, 48727 | -1.20 |
| ENSMUSG00000086689 | Gm16876 | predicted gene, 16876 | -1.20 |
| ENSMUSG00000066621 | Tecpr1 | tectonin beta-propeller repeat containing 1 | -1.21 |
| ENSMUSG00000019302 | Atp6v0a1 | ATPase, H+ transporting, lysosomal V0 subunit A1 | -1.21 |
| ENSMUSG00000039007 | Cpq | carboxypeptidase Q | -1.21 |
| ENSMUSG00000054422 | Fabp1 | fatty acid binding protein 1, liver | -1.21 |
| ENSMUSG00000030865 | Chp2 | calcineurin-like EF hand protein 2 | -1.21 |
| ENSMUSG00000047501 | Cldn4 | claudin 4 | -1.21 |
| ENSMUSG00000029269 | Sult1b1 | sulfotransferase family 1B, member 1 | -1.21 |
| ENSMUSG00000040721 | Zfhx2 | zinc finger homeobox 2 | -1.21 |
| ENSMUSG00000050777 | Tmem37 | transmembrane protein 37 | -1.21 |
| ENSMUSG00000051606 | 2010001K21Rik | RIKEN cDNA 2010001K21 gene | -1.21 |
| ENSMUSG00000074912 | Gm14207 | predicted gene 14207 | -1.21 |
| ENSMUSG00000026879 | Gsn | gelsolin | -1.21 |
| ENSMUSG00000046694 | Tent5b | terminal nucleotidyltransferase 5B | -1.21 |
| ENSMUSG00000024299 | Adamts10 | a disintegrin-like and metallopeptidase (reprolysin type) with thrombospondin type 1 motif, 10 | -1.21 |
| ENSMUSG00000021306 | Gpr137b | G protein-coupled receptor 137B | -1.21 |
| ENSMUSG00000048562 | Sp8 | trans-acting transcription factor 8 | -1.21 |
| ENSMUSG00000037747 | Phyhipl | phytanoyl-CoA hydroxylase interacting protein-like | -1.21 |
| ENSMUSG00000006395 | Hyi | hydroxypyruvate isomerase (putative) | -1.22 |
| ENSMUSG00000051910 | Sox6 | SRY (sex determining region Y)-box 6 | -1.22 |
| ENSMUSG00000059316 | Slc27a4 | solute carrier family 27 (fatty acid transporter), member 4 | -1.22 |
| ENSMUSG00000023959 | Clic5 | chloride intracellular channel 5 | -1.22 |
| ENSMUSG00000018919 | Tm4sf5 | transmembrane 4 superfamily member 5 | -1.22 |
| ENSMUSG00000096215 | Smim22 | small integral membrane protein 22 | -1.22 |
| ENSMUSG00000047641 | Krt87 | keratin 87 | -1.22 |
| ENSMUSG00000059540 | Tcea2 | transcription elongation factor A (SII), 2 | -1.22 |
| ENSMUSG00000049556 | Lingo1 | leucine rich repeat and Ig domain containing 1 | -1.22 |
| ENSMUSG00000112276 | 5033421B08Rik | RIKEN cDNA 5033421B08 gene | -1.22 |
| ENSMUSG00000045027 | Prss22 | protease, serine 22 | -1.22 |
| ENSMUSG00000052125 | F730043M19Rik | RIKEN cDNA F730043M19 gene | -1.22 |
| ENSMUSG00000106871 | Gm3289 | predicted gene 3289 | -1.22 |
| ENSMUSG00000068699 | Flnc | filamin C, gamma | -1.22 |
| ENSMUSG00000052188 | Gm14964 | predicted gene 14964 | -1.22 |
| ENSMUSG00000048249 | Crebrf | CREB3 regulatory factor | -1.23 |
| ENSMUSG00000031618 | Nr3c2 | nuclear receptor subfamily 3, group C, member 2 | -1.23 |
| ENSMUSG00000024131 | Slc3a1 | solute carrier family 3, member 1 | -1.23 |
| ENSMUSG00000034858 | Fam214a | family with sequence similarity 214, member A | -1.23 |
| ENSMUSG00000023495 | Pcbp4 | poly(rC) binding protein 4 | -1.23 |
| ENSMUSG00000060429 | Sntb1 | syntrophin, basic 1 | -1.23 |
| ENSMUSG00000078612 | Fyb2 | FYN binding protein 2 | -1.23 |
| ENSMUSG00000022219 | Cideb | cell death-inducing DNA fragmentation factor, alpha subunit-like effector B | -1.23 |
| ENSMUSG00000041540 | Sox5 | SRY (sex determining region Y)-box 5 | -1.23 |
| ENSMUSG00000001249 | Hpn | hepsin | -1.23 |
| ENSMUSG00000027438 | Napb | N-ethylmaleimide sensitive fusion protein attachment protein beta | -1.23 |
| ENSMUSG00000115062 | 4930544F09Rik | RIKEN cDNA 4930544F09 gene | -1.23 |
| ENSMUSG00000074896 | Ifit3 | interferon-induced protein with tetratricopeptide repeats 3 | -1.23 |
| ENSMUSG00000091017 | Fam71a | family with sequence similarity 71, member A | -1.23 |
| ENSMUSG00000105985 | Gm42993 | predicted gene 42993 | -1.23 |
| ENSMUSG00000102620 | Gm37675 | predicted gene, 37675 | -1.23 |
| ENSMUSG00000038608 | Dock10 | dedicator of cytokinesis 10 | -1.23 |
| ENSMUSG00000089861 | Gm16139 | predicted gene 16139 | -1.23 |
| ENSMUSG00000071356 | Reg3b | regenerating islet-derived 3 beta | -1.24 |
| ENSMUSG00000020829 | Slc46a1 | solute carrier family 46, member 1 | -1.24 |
| ENSMUSG00000034947 | Tmem106a | transmembrane protein 106A | -1.24 |
| ENSMUSG00000001095 | Slc13a2 | solute carrier family 13 (sodium-dependent dicarboxylate transporter), member 2 | -1.24 |

| | | | |
|---|---|---|---|
| ENSMUSG00000097277 | 2900076A07Rik | RIKEN cDNA 2900076A07 gene | -1.24 |
| ENSMUSG00000033460 | Armcx1 | armadillo repeat containing, X-linked 1 | -1.24 |
| ENSMUSG00000034450 | Gulo | gulonolactone (L-) oxidase | -1.24 |
| ENSMUSG00000103693 | Gm37529 | predicted gene, 37529 | -1.24 |
| ENSMUSG00000097601 | Gm26660 | predicted gene, 26660 | -1.24 |
| ENSMUSG00000038777 | Sema6c | sema domain, transmembrane domain (TM), and cytoplasmic domain, (semaphorin) 6C | -1.24 |
| ENSMUSG00000033276 | Stk36 | serine/threonine kinase 36 | -1.24 |
| ENSMUSG00000035551 | Igfbpl1 | insulin-like growth factor binding protein-like 1 | -1.24 |
| ENSMUSG00000033227 | Wnt6 | wingless-type MMTV integration site family, member 6 | -1.24 |
| ENSMUSG00000084137 | Gm9085 | predicted gene 9085 | -1.24 |
| ENSMUSG00000041696 | Rasl12 | RAS-like, family 12 | -1.24 |
| ENSMUSG00000057060 | Slc35f3 | solute carrier family 35, member F3 | -1.24 |
| ENSMUSG00000085085 | 1700086P04Rik | RIKEN cDNA 1700086P04 gene | -1.24 |
| ENSMUSG00000105302 | Gm19817 | predicted gene, 19817 | -1.24 |
| ENSMUSG00000040296 | Ddx58 | DEAD (Asp-Glu-Ala-Asp) box polypeptide 58 | -1.25 |
| ENSMUSG00000038244 | Mical2 | microtubule associated monooxygenase, calponin and LIM domain containing 2 | -1.25 |
| ENSMUSG00000091705 | H2-Q2 | histocompatibility 2, Q region locus 2 | -1.25 |
| ENSMUSG00000032120 | C2cd2l | C2 calcium-dependent domain containing 2-like | -1.25 |
| ENSMUSG00000079507 | H2-Q1 | histocompatibility 2, Q region locus 1 | -1.25 |
| ENSMUSG00000040327 | Cul9 | cullin 9 | -1.25 |
| ENSMUSG00000039943 | Plcb4 | phospholipase C, beta 4 | -1.25 |
| ENSMUSG00000042675 | Ypel3 | yippee like 3 | -1.25 |
| ENSMUSG00000032593 | Amigo3 | adhesion molecule with Ig like domain 3 | -1.25 |
| ENSMUSG00000068923 | Syt11 | synaptotagmin XI | -1.25 |
| ENSMUSG00000058063 | Trim31 | tripartite motif-containing 31 | -1.25 |
| ENSMUSG00000047617 | Paxx | non-homologous end joining factor | -1.25 |
| ENSMUSG00000057706 | Mex3b | mex3 RNA binding family member B | -1.25 |
| ENSMUSG00000021061 | Sptb | spectrin beta, erythrocytic | -1.25 |
| ENSMUSG00000024899 | Papss2 | 3'-phosphoadenosine 5'-phosphosulfate synthase 2 | -1.25 |
| ENSMUSG00000040483 | Xaf1 | XIAP associated factor 1 | -1.25 |
| ENSMUSG00000110276 | Gm45330 | predicted gene 45330 | -1.25 |
| ENSMUSG00000031841 | Cdh13 | cadherin 13 | -1.25 |
| ENSMUSG00000038354 | Ankrd35 | ankyrin repeat domain 35 | -1.25 |
| ENSMUSG00000104020 | Gm37215 | predicted gene, 37215 | -1.25 |
| ENSMUSG00000021414 | Fam217a | family with sequence similarity 217, member A | -1.25 |
| ENSMUSG00000031647 | Mfap3l | microfibrillar-associated protein 3-like | -1.25 |
| ENSMUSG00000115360 | Gm48932 | predicted gene, 48932 | -1.25 |
| ENSMUSG00000054277 | Arfgap3 | ADP-ribosylation factor GTPase activating protein 3 | -1.26 |
| ENSMUSG00000075520 | Malrd1 | MAM and LDL receptor class A domain containing 1 | -1.26 |
| ENSMUSG00000035506 | Slc12a8 | solute carrier family 12 (potassium/chloride transporters), member 8 | -1.26 |
| ENSMUSG00000062995 | Ica1 | islet cell autoantigen 1 | -1.26 |
| ENSMUSG00000031613 | Hpgd | hydroxyprostaglandin dehydrogenase 15 (NAD) | -1.26 |
| ENSMUSG00000042041 | 2010003K11Rik | RIKEN cDNA 2010003K11 gene | -1.26 |
| ENSMUSG00000085333 | 1700030A11Rik | RIKEN cDNA 1700030A11 gene | -1.26 |
| ENSMUSG00000040990 | Sh3kbp1 | SH3-domain kinase binding protein 1 | -1.26 |
| ENSMUSG00000031155 | Pim2 | proviral integration site 2 | -1.26 |
| ENSMUSG00000089817 | Gm7162 | predicted gene 7162 | -1.26 |
| ENSMUSG00000103749 | Pcdhgb5 | protocadherin gamma subfamily B, 5 | -1.26 |
| ENSMUSG00000096995 | 2810029C07Rik | RIKEN cDNA 2810029C07 gene | -1.26 |
| ENSMUSG00000000126 | Wnt9a | wingless-type MMTV integration site family, member 9A | -1.26 |
| ENSMUSG00000038623 | Tm6sf1 | transmembrane 6 superfamily member 1 | -1.26 |
| ENSMUSG00000025207 | Sema4g | sema domain, immunoglobulin domain (Ig), transmembrane domain (TM) and short cytoplasmic dor | -1.27 |
| ENSMUSG00000032776 | Mctp2 | multiple C2 domains, transmembrane 2 | -1.27 |
| ENSMUSG00000015405 | Ace2 | angiotensin I converting enzyme (peptidyl-dipeptidase A) 2 | -1.27 |
| ENSMUSG00000024254 | Abcg8 | ATP binding cassette subfamily G member 8 | -1.27 |
| ENSMUSG00000039899 | Fgl2 | fibrinogen-like protein 2 | -1.27 |
| ENSMUSG00000071862 | Lrrtm2 | leucine rich repeat transmembrane neuronal 2 | -1.27 |
| ENSMUSG00000037818 | Abhd18 | abhydrolase domain containing 18 | -1.27 |
| ENSMUSG00000054999 | Naaladl1 | N-acetylated alpha-linked acidic dipeptidase-like 1 | -1.27 |
| ENSMUSG00000068341 | Reg3d | regenerating islet-derived 3 delta | -1.27 |
| ENSMUSG00000102428 | Pcdhga12 | protocadherin gamma subfamily A, 12 | -1.27 |
| ENSMUSG00000040350 | Trim7 | tripartite motif-containing 7 | -1.27 |
| ENSMUSG00000074782 | 4833422C13Rik | RIKEN cDNA 4833422C13 gene | -1.27 |
| ENSMUSG00000033177 | Tmprss7 | transmembrane serine protease 7 | -1.27 |
| ENSMUSG00000049811 | Fam161a | family with sequence similarity 161, member A | -1.27 |
| ENSMUSG00000024682 | Gif | gastric intrinsic factor | -1.27 |
| ENSMUSG00000107619 | Gm32479 | predicted gene, 32479 | -1.27 |
| ENSMUSG00000069378 | Prdm6 | PR domain containing 6 | -1.27 |
| ENSMUSG00000025129 | Ppp1r27 | protein phosphatase 1, regulatory subunit 27 | -1.27 |
| ENSMUSG00000047307 | Pcdhb13 | protocadherin beta 13 | -1.27 |
| ENSMUSG00000014905 | Dnajb9 | DnaJ heat shock protein family (Hsp40) member B9 | -1.28 |
| ENSMUSG00000023067 | Cdkn1a | cyclin-dependent kinase inhibitor 1A (P21) | -1.28 |
| ENSMUSG00000059991 | Nptx2 | neuronal pentraxin 2 | -1.28 |

| | | | |
|---|---|---|---|
| ENSMUSG00000028015 | Ctso | cathepsin O | -1.28 |
| ENSMUSG00000024713 | Pcsk5 | proprotein convertase subtilisin/kexin type 5 | -1.28 |
| ENSMUSG00000020261 | Slc36a1 | solute carrier family 36 (proton/amino acid symporter), member 1 | -1.28 |
| ENSMUSG00000029188 | Slc34a2 | solute carrier family 34 (sodium phosphate), member 2 | -1.28 |
| ENSMUSG00000021125 | Arg2 | arginase type II | -1.28 |
| ENSMUSG00000018740 | Slc25a35 | solute carrier family 25, member 35 | -1.28 |
| ENSMUSG00000002603 | Tgfb1 | transforming growth factor, beta 1 | -1.28 |
| ENSMUSG00000073403 | Gm10499 | predicted gene 10499 | -1.28 |
| ENSMUSG00000031637 | Lrp2bp | Lrp2 binding protein | -1.28 |
| ENSMUSG00000031298 | Adgrg2 | adhesion G protein-coupled receptor G2 | -1.28 |
| ENSMUSG00000035964 | Tmem59l | transmembrane protein 59-like | -1.28 |
| ENSMUSG00000028435 | Aqp3 | aquaporin 3 | -1.28 |
| ENSMUSG00000070576 | Mn1 | meningioma 1 | -1.28 |
| ENSMUSG00000086701 | Gm13595 | predicted gene 13595 | -1.28 |
| ENSMUSG00000020609 | Apob | apolipoprotein B | -1.29 |
| ENSMUSG00000068742 | Cry2 | cryptochrome 2 (photolyase-like) | -1.29 |
| ENSMUSG00000049307 | Fut4 | fucosyltransferase 4 | -1.29 |
| ENSMUSG00000021062 | Rab15 | RAB15, member RAS oncogene family | -1.29 |
| ENSMUSG00000068735 | Trp53i11 | transformation related protein 53 inducible protein 11 | -1.29 |
| ENSMUSG00000039579 | Grin3a | glutamate receptor ionotropic, NMDA3A | -1.29 |
| ENSMUSG00000037493 | Cib2 | calcium and integrin binding family member 2 | -1.29 |
| ENSMUSG00000042564 | Fam227a | family with sequence similarity 227, member A | -1.29 |
| ENSMUSG00000002997 | Prkar2b | protein kinase, cAMP dependent regulatory, type II beta | -1.29 |
| ENSMUSG00000041912 | Tdrkh | tudor and KH domain containing protein | -1.29 |
| ENSMUSG00000047669 | Msl3l2 | MSL3 like 2 | -1.29 |
| ENSMUSG00000022041 | Chrna2 | cholinergic receptor, nicotinic, alpha polypeptide 2 (neuronal) | -1.29 |
| ENSMUSG00000074121 | Ntf5 | neurotrophin 5 | -1.29 |
| ENSMUSG00000020169 | Best3 | bestrophin 3 | -1.29 |
| ENSMUSG00000070436 | Serpinh1 | serine (or cysteine) peptidase inhibitor, clade H, member 1 | -1.29 |
| ENSMUSG00000112805 | Gm34574 | predicted gene, 34574 | -1.29 |
| ENSMUSG00000005125 | Ndrg1 | N-myc downstream regulated gene 1 | -1.30 |
| ENSMUSG00000029370 | Rassf6 | Ras association (RalGDS/AF-6) domain family member 6 | -1.30 |
| ENSMUSG00000047728 | BC025446 | cDNA sequence BC025446 | -1.30 |
| ENSMUSG00000032192 | Gnb5 | guanine nucleotide binding protein (G protein), beta 5 | -1.30 |
| ENSMUSG00000067297 | Ifit1bl2 | interferon induced protein with tetratricopeptide repeats 1B like 2 | -1.30 |
| ENSMUSG00000021411 | Pxdc1 | PX domain containing 1 | -1.30 |
| ENSMUSG00000052364 | B630019K06Rik | RIKEN cDNA B630019K06 gene | -1.30 |
| ENSMUSG00000032053 | Pou2af1 | POU domain, class 2, associating factor 1 | -1.30 |
| ENSMUSG00000087623 | Gm12404 | predicted gene 12404 | -1.30 |
| ENSMUSG00000109136 | Gm45114 | predicted gene 45114 | -1.30 |
| ENSMUSG00000085723 | Gm15915 | predicted gene 15915 | -1.30 |
| ENSMUSG00000022040 | Ephx2 | epoxide hydrolase 2, cytoplasmic | -1.31 |
| ENSMUSG00000052302 | Tbc1d30 | TBC1 domain family, member 30 | -1.31 |
| ENSMUSG00000029561 | Oasl2 | 2'-5' oligoadenylate synthetase-like 2 | -1.31 |
| ENSMUSG00000022377 | Asap1 | ArfGAP with SH3 domain, ankyrin repeat and PH domain1 | -1.31 |
| ENSMUSG00000102758 | Naaladl2 | N-acetylated alpha-linked acidic dipeptidase-like 2 | -1.31 |
| ENSMUSG00000070315 | 4930581F22Rik | RIKEN cDNA 4930581F22 gene | -1.31 |
| ENSMUSG00000074261 | Erich4 | glutamate rich 4 | -1.31 |
| ENSMUSG00000075270 | Pde11a | phosphodiesterase 11A | -1.31 |
| ENSMUSG00000021759 | Plpp1 | phospholipid phosphatase 1 | -1.31 |
| ENSMUSG00000027805 | Pfn2 | profilin 2 | -1.31 |
| ENSMUSG00000084803 | 5830444B04Rik | RIKEN cDNA 5830444B04 gene | -1.31 |
| ENSMUSG00000027225 | Duoxa2 | dual oxidase maturation factor 2 | -1.31 |
| ENSMUSG00000029925 | Tbxas1 | thromboxane A synthase 1, platelet | -1.31 |
| ENSMUSG00000021268 | Meg3 | maternally expressed 3 | -1.31 |
| ENSMUSG00000054477 | Kcnn2 | potassium intermediate/small conductance calcium-activated channel, subfamily N, member 2 | -1.31 |
| ENSMUSG00000062859 | Tcp11 | t-complex protein 11 | -1.31 |
| ENSMUSG00000087341 | 0610040F04Rik | RIKEN cDNA 0610040F04 gene | -1.31 |
| ENSMUSG00000106709 | Gm30270 | predicted gene, 30270 | -1.31 |
| ENSMUSG00000019906 | Lin7a | lin-7 homolog A (C. elegans) | -1.31 |
| ENSMUSG00000110802 | Gm47141 | predicted gene, 47141 | -1.31 |
| ENSMUSG00000062991 | Nrg1 | neuregulin 1 | -1.31 |
| ENSMUSG00000027820 | Mme | membrane metallo endopeptidase | -1.32 |
| ENSMUSG00000038473 | Nos1ap | nitric oxide synthase 1 (neuronal) adaptor protein | -1.32 |
| ENSMUSG00000053334 | Ficd | FIC domain containing | -1.32 |
| ENSMUSG00000039911 | Spsb1 | splA/ryanodine receptor domain and SOCS box containing 1 | -1.32 |
| ENSMUSG00000028389 | Zfp37 | zinc finger protein 37 | -1.32 |
| ENSMUSG00000030340 | Scnn1a | sodium channel, nonvoltage-gated 1 alpha | -1.32 |
| ENSMUSG00000020034 | Tcp11l2 | t-complex 11 (mouse) like 2 | -1.32 |
| ENSMUSG00000029279 | Brdt | bromodomain, testis-specific | -1.32 |
| ENSMUSG00000000805 | Car4 | carbonic anhydrase 4 | -1.32 |
| ENSMUSG00000033191 | Tie1 | tyrosine kinase with immunoglobulin-like and EGF-like domains 1 | -1.32 |
| ENSMUSG00000044250 | Pced1b | PC-esterase domain containing 1B | -1.32 |

| | | | |
|---|---|---|---|
| ENSMUSG00000079157 | Fam155a | family with sequence similarity 155, member A | -1.32 |
| ENSMUSG00000046447 | Camk2n1 | calcium/calmodulin-dependent protein kinase II inhibitor 1 | -1.33 |
| ENSMUSG00000041609 | Bicdl1 | BICD family like cargo adaptor 1 | -1.33 |
| ENSMUSG00000090214 | Gm15657 | predicted gene 15657 | -1.33 |
| ENSMUSG00000024730 | Ms4a8a | membrane-spanning 4-domains, subfamily A, member 8A | -1.33 |
| ENSMUSG00000028150 | Rorc | RAR-related orphan receptor gamma | -1.33 |
| ENSMUSG00000026614 | Slc30a10 | solute carrier family 30, member 10 | -1.33 |
| ENSMUSG00000028860 | Sytl1 | synaptotagmin-like 1 | -1.33 |
| ENSMUSG00000074207 | Adh1 | alcohol dehydrogenase 1 (class I) | -1.33 |
| ENSMUSG00000085091 | Egfros | epidermal growth factor receptor, opposite strand | -1.33 |
| ENSMUSG00000024503 | Spink1 | serine peptidase inhibitor, Kazal type 1 | -1.33 |
| ENSMUSG00000032238 | Rora | RAR-related orphan receptor alpha | -1.33 |
| ENSMUSG00000110206 | Flt3l | FMS-like tyrosine kinase 3 ligand | -1.33 |
| ENSMUSG00000089028 | Gm25010 | predicted gene, 25010 | -1.33 |
| ENSMUSG00000081968 | Rpl23a-ps2 | ribosomal protein L23A, pseudogene 2 | -1.33 |
| ENSMUSG00000042066 | Tmcc2 | transmembrane and coiled-coil domains 2 | -1.33 |
| ENSMUSG00000098843 | Gm27240 | predicted gene 27240 | -1.33 |
| ENSMUSG00000022512 | Cldn1 | claudin 1 | -1.33 |
| ENSMUSG00000038150 | Ormdl3 | ORM1-like 3 (S. cerevisiae) | -1.34 |
| ENSMUSG00000036452 | Arhgap26 | Rho GTPase activating protein 26 | -1.34 |
| ENSMUSG00000061825 | Ces2c | carboxylesterase 2C | -1.34 |
| ENSMUSG00000032038 | St3gal4 | ST3 beta-galactoside alpha-2,3-sialyltransferase 4 | -1.34 |
| ENSMUSG00000040998 | Npnt | nephronectin | -1.34 |
| ENSMUSG00000034919 | Ttc22 | tetratricopeptide repeat domain 22 | -1.34 |
| ENSMUSG00000037126 | Psd | pleckstrin and Sec7 domain containing | -1.34 |
| ENSMUSG00000032081 | Apoc3 | apolipoprotein C-III | -1.34 |
| ENSMUSG00000072620 | Slfn2 | schlafen 2 | -1.34 |
| ENSMUSG00000064262 | Gimap8 | GTPase, IMAP family member 8 | -1.34 |
| ENSMUSG00000092323 | BB365896 | expressed sequence BB365896 | -1.34 |
| ENSMUSG00000067261 | Foxd3 | forkhead box D3 | -1.34 |
| ENSMUSG00000097983 | Gm26971 | predicted gene, 26971 | -1.34 |
| ENSMUSG00000056296 | Synpr | synaptoporin | -1.34 |
| ENSMUSG00000022416 | Cacna1i | calcium channel, voltage-dependent, alpha 1I subunit | -1.34 |
| ENSMUSG00000019823 | Mical1 | microtubule associated monooxygenase, calponin and LIM domain containing 1 | -1.35 |
| ENSMUSG00000059713 | Rcan3 | regulator of calcineurin 3 | -1.35 |
| ENSMUSG00000020600 | Slc7a15 | solute carrier family 7 (cationic amino acid transporter, y+ system), member 15 | -1.35 |
| ENSMUSG00000024313 | Mep1b | meprin 1 beta | -1.35 |
| ENSMUSG00000043029 | Trpv3 | transient receptor potential cation channel, subfamily V, member 3 | -1.35 |
| ENSMUSG00000040147 | Maob | monoamine oxidase B | -1.35 |
| ENSMUSG00000024386 | Proc | protein C | -1.35 |
| ENSMUSG00000113864 | Gm48381 | predicted gene, 48381 | -1.35 |
| ENSMUSG00000112198 | Gm4065 | predicted gene 4065 | -1.35 |
| ENSMUSG00000072966 | Gprasp2 | G protein-coupled receptor associated sorting protein 2 | -1.35 |
| ENSMUSG00000026938 | Fcna | ficolin A | -1.35 |
| ENSMUSG00000035722 | Abca7 | ATP-binding cassette, sub-family A (ABC1), member 7 | -1.36 |
| ENSMUSG00000056035 | Cyp3a11 | cytochrome P450, family 3, subfamily a, polypeptide 11 | -1.36 |
| ENSMUSG00000039770 | Ypel5 | yippee like 5 | -1.36 |
| ENSMUSG00000052105 | Mtcl1 | microtubule crosslinking factor 1 | -1.36 |
| ENSMUSG00000041361 | Myzap | myocardial zonula adherens protein | -1.36 |
| ENSMUSG00000019989 | Enpp3 | ectonucleotide pyrophosphatase/phosphodiesterase 3 | -1.36 |
| ENSMUSG00000079440 | Alpi | alkaline phosphatase, intestinal | -1.36 |
| ENSMUSG00000028051 | Hcn3 | hyperpolarization-activated, cyclic nucleotide-gated K+ 3 | -1.36 |
| ENSMUSG00000039384 | Dusp10 | dual specificity phosphatase 10 | -1.36 |
| ENSMUSG00000022661 | Cd200 | CD200 antigen | -1.36 |
| ENSMUSG00000054150 | Syne3 | spectrin repeat containing, nuclear envelope family member 3 | -1.36 |
| ENSMUSG00000089857 | Zfp882 | zinc finger protein 882 | -1.36 |
| ENSMUSG00000019831 | Wasf1 | WAS protein family, member 1 | -1.36 |
| ENSMUSG00000111293 | Gm34006 | predicted gene, 34006 | -1.36 |
| ENSMUSG00000034774 | Dsg1c | desmoglein 1 gamma | -1.36 |
| ENSMUSG00000056091 | St3gal5 | ST3 beta-galactoside alpha-2,3-sialyltransferase 5 | -1.36 |
| ENSMUSG00000005640 | Insrr | insulin receptor-related receptor | -1.36 |
| ENSMUSG00000037235 | Mxd4 | Max dimerization protein 4 | -1.37 |
| ENSMUSG00000020300 | Cpeb4 | cytoplasmic polyadenylation element binding protein 4 | -1.37 |
| ENSMUSG00000030017 | Reg3g | regenerating islet-derived 3 gamma | -1.37 |
| ENSMUSG00000024193 | Phf1 | PHD finger protein 1 | -1.37 |
| ENSMUSG00000028976 | Slc2a5 | solute carrier family 2 (facilitated glucose transporter), member 5 | -1.37 |
| ENSMUSG00000019278 | Dpep1 | dipeptidase 1 (renal) | -1.37 |
| ENSMUSG00000029082 | Bst1 | bone marrow stromal cell antigen 1 | -1.37 |
| ENSMUSG00000103579 | Gm37113 | predicted gene, 37113 | -1.37 |
| ENSMUSG00000110754 | 4930405J17Rik | RIKEN cDNA 4930405J17 gene | -1.37 |
| ENSMUSG00000019762 | Iyd | iodotyrosine deiodinase | -1.37 |
| ENSMUSG00000059013 | Sh2d3c | SH2 domain containing 3C | -1.37 |
| ENSMUSG00000026586 | Prrx1 | paired related homeobox 1 | -1.37 |

| | | | |
|---|---|---|---|
| ENSMUSG00000035158 | Mitf | melanogenesis associated transcription factor | -1.37 |
| ENSMUSG00000044912 | Syt16 | synaptotagmin XVI | -1.37 |
| ENSMUSG00000109093 | Gm19950 | predicted gene, 19950 | -1.37 |
| ENSMUSG00000022946 | Dopey2 | dopey family member 2 | -1.38 |
| ENSMUSG00000092618 | Btnl6 | butyrophilin-like 6 | -1.38 |
| ENSMUSG00000040505 | Abcg5 | ATP binding cassette subfamily G member 5 | -1.38 |
| ENSMUSG00000021751 | Acox2 | acyl-Coenzyme A oxidase 2, branched chain | -1.38 |
| ENSMUSG00000007989 | Fzd3 | frizzled class receptor 3 | -1.38 |
| ENSMUSG00000057596 | Trim30d | tripartite motif-containing 30D | -1.38 |
| ENSMUSG00000019647 | Sema6a | sema domain, transmembrane domain (TM), and cytoplasmic domain, (semaphorin) 6A | -1.38 |
| ENSMUSG00000027358 | Bmp2 | bone morphogenetic protein 2 | -1.38 |
| ENSMUSG00000037738 | Nek5 | NIMA (never in mitosis gene a)-related expressed kinase 5 | -1.38 |
| ENSMUSG00000020604 | Arsg | arylsulfatase G | -1.38 |
| ENSMUSG00000097443 | Gm17529 | predicted gene, 17529 | -1.38 |
| ENSMUSG00000060244 | Alyref2 | Aly/REF export factor 2 | -1.38 |
| ENSMUSG00000000386 | Mx1 | MX dynamin-like GTPase 1 | -1.38 |
| ENSMUSG00000111425 | Gm47324 | predicted gene, 47324 | -1.38 |
| ENSMUSG00000043313 | Pcdhb19 | protocadherin beta 19 | -1.38 |
| ENSMUSG00000095066 | Defa20 | defensin, alpha, 20 | -1.38 |
| ENSMUSG00000026303 | Mlph | melanophilin | -1.39 |
| ENSMUSG00000028088 | Fmo5 | flavin containing monooxygenase 5 | -1.39 |
| ENSMUSG00000020447 | Npc1l1 | NPC1 like intracellular cholesterol transporter 1 | -1.39 |
| ENSMUSG00000018427 | Ypel2 | yippee like 2 | -1.39 |
| ENSMUSG00000048612 | Myof | myoferlin | -1.39 |
| ENSMUSG00000048232 | Fbxo10 | F-box protein 10 | -1.39 |
| ENSMUSG00000047910 | Pcdhb16 | protocadherin beta 16 | -1.39 |
| ENSMUSG00000025348 | Itga7 | integrin alpha 7 | -1.39 |
| ENSMUSG00000062017 | Abca14 | ATP-binding cassette, sub-family A (ABC1), member 14 | -1.39 |
| ENSMUSG00000086231 | Rapgef4os3 | Rap guanine nucleotide exchange factor (GEF) 4, opposite strand 3 | -1.39 |
| ENSMUSG00000042567 | Nek10 | NIMA (never in mitosis gene a)- related kinase 10 | -1.39 |
| ENSMUSG00000038700 | Hoxb5 | homeobox B5 | -1.39 |
| ENSMUSG00000026509 | Capn2 | calpain 2 | -1.40 |
| ENSMUSG00000024190 | Dusp1 | dual specificity phosphatase 1 | -1.40 |
| ENSMUSG00000021453 | Gadd45g | growth arrest and DNA-damage-inducible 45 gamma | -1.40 |
| ENSMUSG00000031303 | Map3k15 | mitogen-activated protein kinase kinase kinase 15 | -1.40 |
| ENSMUSG00000002944 | Cd36 | CD36 molecule | -1.40 |
| ENSMUSG00000037049 | Smpd1 | sphingomyelin phosphodiesterase 1, acid lysosomal | -1.40 |
| ENSMUSG00000045838 | Ccdc9b | coiled-coil domain containing 9B | -1.40 |
| ENSMUSG00000068452 | Duox2 | dual oxidase 2 | -1.40 |
| ENSMUSG00000034685 | Fam171a2 | family with sequence similarity 171, member A2 | -1.40 |
| ENSMUSG00000000253 | Gmpr | guanosine monophosphate reductase | -1.40 |
| ENSMUSG00000034634 | Ly6d | lymphocyte antigen 6 complex, locus D | -1.40 |
| ENSMUSG00000015745 | Plekho1 | pleckstrin homology domain containing, family O member 1 | -1.40 |
| ENSMUSG00000024421 | Lama3 | laminin, alpha 3 | -1.40 |
| ENSMUSG00000033847 | Pla2g4c | phospholipase A2, group IVC (cytosolic, calcium-independent) | -1.40 |
| ENSMUSG00000074217 | 2210011C24Rik | RIKEN cDNA 2210011C24 gene | -1.40 |
| ENSMUSG00000010066 | Cacna2d2 | calcium channel, voltage-dependent, alpha 2/delta subunit 2 | -1.40 |
| ENSMUSG00000056162 | Cndp1 | carnosine dipeptidase 1 (metallopeptidase M20 family) | -1.40 |
| ENSMUSG00000027314 | Dll4 | delta like canonical Notch ligand 4 | -1.41 |
| ENSMUSG00000019970 | Sgk1 | serum/glucocorticoid regulated kinase 1 | -1.41 |
| ENSMUSG00000066258 | Trim12a | tripartite motif-containing 12A | -1.41 |
| ENSMUSG00000032289 | Thsd4 | thrombospondin, type I, domain containing 4 | -1.41 |
| ENSMUSG00000022885 | St6gal1 | beta galactoside alpha 2,6 sialyltransferase 1 | -1.41 |
| ENSMUSG00000087331 | 1810021B22Rik | RIKEN cDNA 1810021B22 gene | -1.41 |
| ENSMUSG00000005986 | Ankrd13d | ankyrin repeat domain 13 family, member D | -1.41 |
| ENSMUSG00000074657 | Kif5a | kinesin family member 5A | -1.41 |
| ENSMUSG00000030897 | Cnga4 | cyclic nucleotide gated channel alpha 4 | -1.41 |
| ENSMUSG00000020423 | Btg2 | B cell translocation gene 2, anti-proliferative | -1.42 |
| ENSMUSG00000025498 | Irf7 | interferon regulatory factor 7 | -1.42 |
| ENSMUSG00000020715 | Ern1 | endoplasmic reticulum (ER) to nucleus signalling 1 | -1.42 |
| ENSMUSG00000034918 | Cdhr2 | cadherin-related family member 2 | -1.42 |
| ENSMUSG00000030155 | Clec2e | C-type lectin domain family 2, member e | -1.42 |
| ENSMUSG00000022048 | Dpysl2 | dihydropyrimidinase-like 2 | -1.42 |
| ENSMUSG00000036989 | Trim3 | tripartite motif-containing 3 | -1.42 |
| ENSMUSG00000065145 | Vaultrc5 | vault RNA component 5 | -1.42 |
| ENSMUSG00000025035 | Arl3 | ADP-ribosylation factor-like 3 | -1.42 |
| ENSMUSG00000051452 | Gm11437 | predicted gene 11437 | -1.42 |
| ENSMUSG00000048895 | Cdk5r1 | cyclin-dependent kinase 5, regulatory subunit 1 (p35) | -1.42 |
| ENSMUSG00000027931 | Npr1 | natriuretic peptide receptor 1 | -1.42 |
| ENSMUSG00000100593 | 1700119H24Rik | RIKEN cDNA 1700119H24 gene | -1.42 |
| ENSMUSG00000096054 | Syne1 | spectrin repeat containing, nuclear envelope 1 | -1.42 |
| ENSMUSG00000097572 | Gm26797 | predicted gene, 26797 | -1.42 |
| ENSMUSG00000103738 | Gm37652 | predicted gene, 37652 | -1.42 |

| | | | |
|---|---|---|---|
| ENSMUSG00000100747 | 1700084E18Rik | RIKEN cDNA 1700084E18 gene | -1.42 |
| ENSMUSG00000081245 | Gm6587 | predicted gene 6587 | -1.42 |
| ENSMUSG00000021850 | ccdc198 | coiled-coil domain containing 198 | -1.42 |
| ENSMUSG00000020836 | Coro6 | coronin 6 | -1.42 |
| ENSMUSG00000032892 | Rangrf | RAN guanine nucleotide release factor | -1.42 |
| ENSMUSG00000029992 | Gfpt1 | glutamine fructose-6-phosphate transaminase 1 | -1.43 |
| ENSMUSG00000052085 | Dock8 | dedicator of cytokinesis 8 | -1.43 |
| ENSMUSG00000034981 | Parm1 | prostate androgen-regulated mucin-like protein 1 | -1.43 |
| ENSMUSG00000020709 | Adap2 | ArfGAP with dual PH domains 2 | -1.43 |
| ENSMUSG00000061808 | Ttr | transthyretin | -1.43 |
| ENSMUSG00000022686 | B3gnt5 | UDP-GlcNAc:betaGal beta-1,3-N-acetylglucosaminyltransferase 5 | -1.43 |
| ENSMUSG00000079499 | 6530402F18Rik | RIKEN cDNA 6530402F18 gene | -1.43 |
| ENSMUSG00000042109 | Csdc2 | cold shock domain containing C2, RNA binding | -1.43 |
| ENSMUSG00000105720 | Gm42440 | predicted gene 42440 | -1.43 |
| ENSMUSG00000078942 | Naip6 | NLR family, apoptosis inhibitory protein 6 | -1.44 |
| ENSMUSG00000020021 | Fgd6 | FYVE, RhoGEF and PH domain containing 6 | -1.44 |
| ENSMUSG00000028024 | Enpep | glutamyl aminopeptidase | -1.44 |
| ENSMUSG00000041827 | Oasl1 | 2'-5' oligoadenylate synthetase-like 1 | -1.44 |
| ENSMUSG00000085148 | Mir22hg | Mir22 host gene (non-protein coding) | -1.44 |
| ENSMUSG00000040389 | Wdr47 | WD repeat domain 47 | -1.44 |
| ENSMUSG00000054793 | Cadm4 | cell adhesion molecule 4 | -1.44 |
| ENSMUSG00000116940 | CT033750.3 | antizyme inhibitor 2 (Azin2) pseudogene | -1.44 |
| ENSMUSG00000097081 | Gm10425 | predicted gene 10425 | -1.44 |
| ENSMUSG00000011256 | Adam19 | a disintegrin and metallopeptidase domain 19 (meltrin beta) | -1.44 |
| ENSMUSG00000103726 | Gm30074 | predicted gene, 30074 | -1.44 |
| ENSMUSG00000037822 | Smim14 | small integral membrane protein 14 | -1.45 |
| ENSMUSG00000041119 | Pde9a | phosphodiesterase 9A | -1.45 |
| ENSMUSG00000053054 | Adh6a | alcohol dehydrogenase 6A (class V) | -1.45 |
| ENSMUSG00000027219 | Slc28a2 | solute carrier family 28 (sodium-coupled nucleoside transporter), member 2 | -1.45 |
| ENSMUSG00000019066 | Rab3d | RAB3D, member RAS oncogene family | -1.45 |
| ENSMUSG00000044734 | Serpinb1a | serine (or cysteine) peptidase inhibitor, clade B, member 1a | -1.45 |
| ENSMUSG00000055730 | Ces2a | carboxylesterase 2A | -1.45 |
| ENSMUSG00000041695 | Kcnj2 | potassium inwardly-rectifying channel, subfamily J, member 2 | -1.45 |
| ENSMUSG00000030643 | Rab30 | RAB30, member RAS oncogene family | -1.45 |
| ENSMUSG00000027950 | Chrnb2 | cholinergic receptor, nicotinic, beta polypeptide 2 (neuronal) | -1.45 |
| ENSMUSG00000024846 | Cst6 | cystatin E/M | -1.45 |
| ENSMUSG00000020900 | Myh10 | myosin, heavy polypeptide 10, non-muscle | -1.45 |
| ENSMUSG00000024039 | Cbs | cystathionine beta-synthase | -1.45 |
| ENSMUSG00000097147 | Gm26780 | predicted gene, 26780 | -1.45 |
| ENSMUSG00000022091 | Sorbs3 | sorbin and SH3 domain containing 3 | -1.45 |
| ENSMUSG00000097391 | Mirg | miRNA containing gene | -1.45 |
| ENSMUSG00000014158 | Trpv4 | transient receptor potential cation channel, subfamily V, member 4 | -1.45 |
| ENSMUSG00000040428 | Plekha4 | pleckstrin homology domain containing, family A (phosphoinositide binding specific) member 4 | -1.45 |
| ENSMUSG00000107583 | Gm44104 | predicted gene, 44104 | -1.45 |
| ENSMUSG00000056418 | BC043934 | cDNA sequence BC043934 | -1.45 |
| ENSMUSG00000102882 | Gm2065 | predicted gene 2065 | -1.45 |
| ENSMUSG00000038178 | Slc43a2 | solute carrier family 43, member 2 | -1.46 |
| ENSMUSG00000048905 | 4930539E08Rik | RIKEN cDNA 4930539E08 gene | -1.46 |
| ENSMUSG00000022197 | Pdzd2 | PDZ domain containing 2 | -1.46 |
| ENSMUSG00000110397 | Gm45540 | predicted gene 45540 | -1.46 |
| ENSMUSG00000061740 | Cyp2d22 | cytochrome P450, family 2, subfamily d, polypeptide 22 | -1.46 |
| ENSMUSG00000105374 | Gm42551 | predicted gene 42551 | -1.46 |
| ENSMUSG00000029838 | Ptn | pleiotrophin | -1.46 |
| ENSMUSG00000094840 | Muc3a | mucin 3A, cell surface associated | -1.47 |
| ENSMUSG00000036390 | Gadd45a | growth arrest and DNA-damage-inducible 45 alpha | -1.47 |
| ENSMUSG00000039217 | Il18 | interleukin 18 | -1.47 |
| ENSMUSG00000032652 | Crebl2 | cAMP responsive element binding protein-like 2 | -1.47 |
| ENSMUSG00000058921 | Slc10a5 | solute carrier family 10 (sodium/bile acid cotransporter family), member 5 | -1.47 |
| ENSMUSG00000048473 | Sult6b2 | sulfotransferase family 6B, member 2 | -1.47 |
| ENSMUSG00000055148 | Klf2 | Kruppel-like factor 2 (lung) | -1.47 |
| ENSMUSG00000074622 | Mafb | v-maf musculoaponeurotic fibrosarcoma oncogene family, protein B (avian) | -1.47 |
| ENSMUSG00000110887 | Cbx3-ps8 | chromobox 3, pseudogene 8 | -1.47 |
| ENSMUSG00000057933 | Gsta2 | glutathione S-transferase, alpha 2 (Yc2) | -1.47 |
| ENSMUSG00000023393 | Slc17a9 | solute carrier family 17, member 9 | -1.47 |
| ENSMUSG00000032965 | Ift57 | intraflagellar transport 57 | -1.47 |
| ENSMUSG00000051579 | Tceal8 | transcription elongation factor A (SII)-like 8 | -1.47 |
| ENSMUSG00000050132 | Sarm1 | sterile alpha and HEAT/Armadillo motif containing 1 | -1.47 |
| ENSMUSG00000029707 | Fscn3 | fascin actin-bundling protein 3 | -1.47 |
| ENSMUSG00000038648 | Creb3l2 | cAMP responsive element binding protein 3-like 2 | -1.48 |
| ENSMUSG00000041644 | Slc5a12 | solute carrier family 5 (sodium/glucose cotransporter), member 12 | -1.48 |
| ENSMUSG00000062785 | Kcnc3 | potassium voltage gated channel, Shaw-related subfamily, member 3 | -1.48 |
| ENSMUSG00000038893 | Fam117a | family with sequence similarity 117, member A | -1.48 |
| ENSMUSG00000103432 | 6720464F23Rik | RIKEN cDNA 6720464F23 gene | -1.48 |

| | | | |
|---|---|---|---|
| ENSMUSG00000022440 | C1qtnf6 | C1q and tumor necrosis factor related protein 6 | -1.48 |
| ENSMUSG00000006675 | P4htm | prolyl 4-hydroxylase, transmembrane (endoplasmic reticulum) | -1.48 |
| ENSMUSG00000035513 | Ntng2 | netrin G2 | -1.48 |
| ENSMUSG00000026380 | Tfcp2l1 | transcription factor CP2-like 1 | -1.49 |
| ENSMUSG00000036611 | Eepd1 | endonuclease/exonuclease/phosphatase family domain containing 1 | -1.49 |
| ENSMUSG00000000204 | Slfn4 | schlafen 4 | -1.49 |
| ENSMUSG00000050295 | Foxc1 | forkhead box C1 | -1.49 |
| ENSMUSG00000073406 | H2-Bl | histocompatibility 2, blastocyst | -1.49 |
| ENSMUSG00000020178 | Adora2a | adenosine A2a receptor | -1.49 |
| ENSMUSG00000101587 | Gm29036 | predicted gene 29036 | -1.49 |
| ENSMUSG00000085003 | Pip5k1bos | phosphatidylinositol-4-phosphate 5-kinase, type 1 beta, opposite strand | -1.49 |
| ENSMUSG00000036872 | Abcc12 | ATP-binding cassette, sub-family C (CFTR/MRP), member 12 | -1.49 |
| ENSMUSG00000110497 | 9230110F11Rik | RIKEN cDNA 9230110F11 gene | -1.49 |
| ENSMUSG00000097194 | 9330175E14Rik | RIKEN cDNA 9330175E14 gene | -1.49 |
| ENSMUSG00000087625 | 4930419G24Rik | RIKEN cDNA 4930419G24 gene | -1.49 |
| ENSMUSG00000058488 | Kl | klotho | -1.49 |
| ENSMUSG00000021565 | Slc6a19 | solute carrier family 6 (neurotransmitter transporter), member 19 | -1.50 |
| ENSMUSG00000053862 | Slc51b | solute carrier family 51, beta subunit | -1.50 |
| ENSMUSG00000024232 | Bambi | BMP and activin membrane-bound inhibitor | -1.50 |
| ENSMUSG00000004187 | Kifc2 | kinesin family member C2 | -1.50 |
| ENSMUSG00000058163 | Gm5431 | predicted gene 5431 | -1.50 |
| ENSMUSG00000024940 | Ltbp3 | latent transforming growth factor beta binding protein 3 | -1.50 |
| ENSMUSG00000113680 | Gm47405 | predicted gene, 47405 | -1.50 |
| ENSMUSG00000085852 | Gm13807 | predicted gene 13807 | -1.50 |
| ENSMUSG00000055805 | Fmnl1 | formin-like 1 | -1.50 |
| ENSMUSG00000022340 | Sybu | syntabulin (syntaxin-interacting) | -1.50 |
| ENSMUSG00000032085 | Tagln | transgelin | -1.50 |
| ENSMUSG00000102147 | Gm38055 | predicted gene, 38055 | -1.50 |
| ENSMUSG00000036040 | Adamtsl2 | ADAMTS-like 2 | -1.50 |
| ENSMUSG00000039168 | Dap | death-associated protein | -1.51 |
| ENSMUSG00000003363 | Pld3 | phospholipase D family, member 3 | -1.51 |
| ENSMUSG00000027602 | Map1lc3a | microtubule-associated protein 1 light chain 3 alpha | -1.51 |
| ENSMUSG00000032080 | Apoa4 | apolipoprotein A-IV | -1.51 |
| ENSMUSG00000003032 | Klf4 | Kruppel-like factor 4 (gut) | -1.51 |
| ENSMUSG00000032454 | Rbp2 | retinol binding protein 2, cellular | -1.51 |
| ENSMUSG00000028519 | Dab1 | disabled 1 | -1.51 |
| ENSMUSG00000067813 | Xkr9 | X-linked Kx blood group related 9 | -1.51 |
| ENSMUSG00000036002 | Fam214b | family with sequence similarity 214, member B | -1.51 |
| ENSMUSG00000094584 | Ms4a18 | membrane-spanning 4-domains, subfamily A, member 18 | -1.51 |
| ENSMUSG00000039934 | Gsap | gamma-secretase activating protein | -1.51 |
| ENSMUSG00000028427 | Aqp7 | aquaporin 7 | -1.51 |
| ENSMUSG00000039963 | Ccdc40 | coiled-coil domain containing 40 | -1.51 |
| ENSMUSG00000037605 | Adgrl3 | adhesion G protein-coupled receptor L3 | -1.51 |
| ENSMUSG00000115555 | Gm49307 | predicted gene, 49307 | -1.51 |
| ENSMUSG00000003418 | St8sia6 | ST8 alpha-N-acetyl-neuraminide alpha-2,8-sialyltransferase 6 | -1.51 |
| ENSMUSG00000037762 | Slc16a9 | solute carrier family 16 (monocarboxylic acid transporters), member 9 | -1.51 |
| ENSMUSG00000046876 | Atxn1 | ataxin 1 | -1.52 |
| ENSMUSG00000013643 | Lypd8 | LY6/PLAUR domain containing 8 | -1.52 |
| ENSMUSG00000032226 | Gcnt3 | glucosaminyl (N-acetyl) transferase 3, mucin type | -1.52 |
| ENSMUSG00000036377 | C530008M17Rik | RIKEN cDNA C530008M17 gene | -1.52 |
| ENSMUSG00000022938 | Fam3b | family with sequence similarity 3, member B | -1.52 |
| ENSMUSG00000108291 | Gm44292 | predicted gene, 44292 | -1.52 |
| ENSMUSG00000026222 | Sp100 | nuclear antigen Sp100 | -1.52 |
| ENSMUSG00000021281 | Tnfaip2 | tumor necrosis factor, alpha-induced protein 2 | -1.52 |
| ENSMUSG00000019359 | Gdpd2 | glycerophosphodiester phosphodiesterase domain containing 2 | -1.52 |
| ENSMUSG00000039004 | Bmp6 | bone morphogenetic protein 6 | -1.52 |
| ENSMUSG00000074183 | Gsta1 | glutathione S-transferase, alpha 1 (Ya) | -1.52 |
| ENSMUSG00000045664 | Cdc42ep2 | CDC42 effector protein (Rho GTPase binding) 2 | -1.52 |
| ENSMUSG00000104776 | Gm43691 | predicted gene 43691 | -1.52 |
| ENSMUSG00000113404 | Gm48622 | predicted gene, 48622 | -1.52 |
| ENSMUSG00000102481 | Gm37925 | predicted gene, 37925 | -1.52 |
| ENSMUSG00000040093 | Bmf | BCL2 modifying factor | -1.53 |
| ENSMUSG00000020847 | Rph3al | rabphilin 3A-like (without C2 domains) | -1.53 |
| ENSMUSG00000047496 | Rnf152 | ring finger protein 152 | -1.53 |
| ENSMUSG00000074195 | Clca4b | chloride channel accessory 4B | -1.53 |
| ENSMUSG00000085747 | Slc13a2os | solute carrier family 13 (sodium-dependent dicarboxylate transporter), member 2, opposite strand | -1.53 |
| ENSMUSG00000045349 | Sh2d5 | SH2 domain containing 5 | -1.53 |
| ENSMUSG00000106229 | Gm19409 | predicted gene, 19409 | -1.53 |
| ENSMUSG00000070687 | Htr1d | 5-hydroxytryptamine (serotonin) receptor 1D | -1.53 |
| ENSMUSG00000056596 | Trnp1 | TMF1-regulated nuclear protein 1 | -1.53 |
| ENSMUSG00000029343 | Crybb1 | crystallin, beta B1 | -1.53 |
| ENSMUSG00000043822 | Adamtsl5 | ADAMTS-like 5 | -1.54 |
| ENSMUSG00000068547 | Clca4a | chloride channel accessory 4A | -1.54 |

| | | | |
|---|---|---|---|
| ENSMUSG00000025528 | 2010106E10Rik | RIKEN cDNA 2010106E10 gene | -1.54 |
| ENSMUSG00000053158 | Fes | feline sarcoma oncogene | -1.54 |
| ENSMUSG00000079038 | D130040H23Rik | RIKEN cDNA D130040H23 gene | -1.54 |
| ENSMUSG00000035390 | Brsk1 | BR serine/threonine kinase 1 | -1.54 |
| ENSMUSG00000025854 | Fam20c | family with sequence similarity 20, member C | -1.54 |
| ENSMUSG00000042096 | Dao | D-amino acid oxidase | -1.54 |
| ENSMUSG00000085830 | Grin1os | glutamate receptor, ionotropic, NMDA1 (zeta 1), opposite strand | -1.54 |
| ENSMUSG00000041710 | Trpc5 | transient receptor potential cation channel, subfamily C, member 5 | -1.54 |
| ENSMUSG00000051678 | Pcdhb6 | protocadherin beta 6 | -1.54 |
| ENSMUSG00000022564 | Grina | glutamate receptor, ionotropic, N-methyl D-aspartate-associated protein 1 (glutamate binding) | -1.55 |
| ENSMUSG00000092341 | Malat1 | metastasis associated lung adenocarcinoma transcript 1 (non-coding RNA) | -1.55 |
| ENSMUSG00000037390 | Muc3 | mucin 3, intestinal | -1.55 |
| ENSMUSG00000018378 | Cuedc1 | CUE domain containing 1 | -1.55 |
| ENSMUSG00000040447 | Spns2 | spinster homolog 2 | -1.55 |
| ENSMUSG00000006464 | Bbs1 | Bardet-Biedl syndrome 1 (human) | -1.55 |
| ENSMUSG00000040432 | Ltb4r2 | leukotriene B4 receptor 2 | -1.55 |
| ENSMUSG00000056973 | Ces1d | carboxylesterase 1D | -1.55 |
| ENSMUSG00000028758 | Kif17 | kinesin family member 17 | -1.55 |
| ENSMUSG00000108603 | 5330411O13Rik | RIKEN cDNA 5330411O13 gene | -1.55 |
| ENSMUSG00000020080 | Hkdc1 | hexokinase domain containing 1 | -1.56 |
| ENSMUSG00000006930 | Hap1 | huntingtin-associated protein 1 | -1.56 |
| ENSMUSG00000039652 | Cpeb3 | cytoplasmic polyadenylation element binding protein 3 | -1.56 |
| ENSMUSG00000049551 | Fzd9 | frizzled class receptor 9 | -1.56 |
| ENSMUSG00000074196 | Clca4c-ps | chloride channel accessory 4C, pseudogene | -1.56 |
| ENSMUSG00000019916 | P4ha1 | procollagen-proline, 2-oxoglutarate 4-dioxygenase (proline 4-hydroxylase), alpha 1 polypeptide | -1.56 |
| ENSMUSG00000026620 | Mark1 | MAP/microtubule affinity regulating kinase 1 | -1.56 |
| ENSMUSG00000022146 | Osmr | oncostatin M receptor | -1.56 |
| ENSMUSG00000030317 | Timp4 | tissue inhibitor of metalloproteinase 4 | -1.56 |
| ENSMUSG00000022575 | Gsdmd | gasdermin D | -1.57 |
| ENSMUSG00000039621 | Prex1 | phosphatidylinositol-3,4,5-trisphosphate-dependent Rac exchange factor 1 | -1.57 |
| ENSMUSG00000035493 | Tgfbi | transforming growth factor, beta induced | -1.57 |
| ENSMUSG00000073399 | Trim40 | tripartite motif-containing 40 | -1.57 |
| ENSMUSG00000014303 | Glis2 | GLIS family zinc finger 2 | -1.57 |
| ENSMUSG00000040714 | Klc3 | kinesin light chain 3 | -1.57 |
| ENSMUSG00000085363 | Gm15478 | predicted gene 15478 | -1.57 |
| ENSMUSG00000036912 | Piwil4 | piwi-like RNA-mediated gene silencing 4 | -1.57 |
| ENSMUSG00000117182 | CT025731.2 | ribosomal protein S6 (Rps6) pseudogene | -1.57 |
| ENSMUSG00000105326 | 6030400A10Rik | RIKEN cDNA 6030400A10 gene | -1.57 |
| ENSMUSG00000116506 | 5730414N17Rik | RIKEN cDNA 5730414N17 gene | -1.57 |
| ENSMUSG00000086039 | Gm12227 | predicted gene 12227 | -1.57 |
| ENSMUSG00000031523 | Dlc1 | deleted in liver cancer 1 | -1.57 |
| ENSMUSG00000087082 | Gm15423 | predicted gene 15423 | -1.57 |
| ENSMUSG00000023259 | Slc26a6 | solute carrier family 26, member 6 | -1.58 |
| ENSMUSG00000025497 | Cdhr5 | cadherin-related family member 5 | -1.58 |
| ENSMUSG00000037005 | Xpnpep2 | X-prolyl aminopeptidase (aminopeptidase P) 2, membrane-bound | -1.58 |
| ENSMUSG00000061728 | Btnl7-ps | butyrophilin-like 7, pseudogene | -1.58 |
| ENSMUSG00000046794 | Ppp1r3b | protein phosphatase 1, regulatory subunit 3B | -1.58 |
| ENSMUSG00000081219 | Bambi-ps1 | BMP and activin membrane-bound inhibitor, pseudogene (Xenopus laevis) | -1.58 |
| ENSMUSG00000051951 | Xkr4 | X-linked Kx blood group related 4 | -1.58 |
| ENSMUSG00000061576 | Dpp6 | dipeptidylpeptidase 6 | -1.58 |
| ENSMUSG00000027199 | Gatm | glycine amidinotransferase (L-arginine:glycine amidinotransferase) | -1.58 |
| ENSMUSG00000104519 | Gm37161 | predicted gene, 37161 | -1.58 |
| ENSMUSG00000003153 | Slc2a3 | solute carrier family 2 (facilitated glucose transporter), member 3 | -1.58 |
| ENSMUSG00000073293 | Nudt10 | nudix (nucleoside diphosphate linked moiety X)-type motif 10 | -1.58 |
| ENSMUSG00000041261 | Car8 | carbonic anhydrase 8 | -1.59 |
| ENSMUSG00000034586 | Hid1 | HID1 domain containing | -1.59 |
| ENSMUSG00000029727 | Cyp3a13 | cytochrome P450, family 3, subfamily a, polypeptide 13 | -1.59 |
| ENSMUSG00000006345 | Ggt1 | gamma-glutamyltransferase 1 | -1.59 |
| ENSMUSG00000041351 | Rap1gap | Rap1 GTPase-activating protein | -1.59 |
| ENSMUSG00000024511 | Rab27b | RAB27B, member RAS oncogene family | -1.59 |
| ENSMUSG00000025787 | Tgm4 | transglutaminase 4 (prostate) | -1.59 |
| ENSMUSG00000016356 | Col20a1 | collagen, type XX, alpha 1 | -1.59 |
| ENSMUSG00000074067 | Gm10619 | predicted gene 10619 | -1.59 |
| ENSMUSG00000024011 | Pi16 | peptidase inhibitor 16 | -1.59 |
| ENSMUSG00000117246 | CT025671.2 | periphilin 1 (Pphln1) pseudogene | -1.59 |
| ENSMUSG00000006342 | Susd2 | sushi domain containing 2 | -1.60 |
| ENSMUSG00000003378 | Grik5 | glutamate receptor, ionotropic, kainate 5 (gamma 2) | -1.60 |
| ENSMUSG00000036295 | Lrrn3 | leucine rich repeat protein 3, neuronal | -1.60 |
| ENSMUSG00000035775 | Krt20 | keratin 20 | -1.60 |
| ENSMUSG00000028737 | Aldh4a1 | aldehyde dehydrogenase 4 family, member A1 | -1.60 |
| ENSMUSG00000017667 | Zfp334 | zinc finger protein 334 | -1.60 |
| ENSMUSG00000061947 | Serpina10 | serine (or cysteine) peptidase inhibitor, clade A (alpha-1 antiproteinase, antitrypsin), member 10 | -1.60 |
| ENSMUSG00000117286 | Gm1043 | predicted 1043 | -1.60 |

| | | | |
|---|---|---|---|
| ENSMUSG00000055357 | 4933400A11Rik | RIKEN cDNA 4933400A11 gene | -1.60 |
| ENSMUSG00000024027 | Glp1r | glucagon-like peptide 1 receptor | -1.60 |
| ENSMUSG00000066392 | Nrxn3 | neurexin III | -1.60 |
| ENSMUSG00000107427 | Gm43948 | predicted gene, 43948 | -1.60 |
| ENSMUSG00000042846 | Lrrtm3 | leucine rich repeat transmembrane neuronal 3 | -1.60 |
| ENSMUSG00000019960 | Dusp6 | dual specificity phosphatase 6 | -1.61 |
| ENSMUSG00000055978 | Fut2 | fucosyltransferase 2 | -1.61 |
| ENSMUSG00000034427 | Myo15b | myosin XVB | -1.61 |
| ENSMUSG00000022546 | Gpt | glutamic pyruvic transaminase, soluble | -1.61 |
| ENSMUSG00000055114 | Anxa13 | annexin A13 | -1.61 |
| ENSMUSG00000047787 | Flrt1 | fibronectin leucine rich transmembrane protein 1 | -1.61 |
| ENSMUSG00000100094 | 1810008I18Rik | RIKEN cDNA 1810008I18 gene | -1.61 |
| ENSMUSG00000040473 | Cfap69 | cilia and flagella associated protein 69 | -1.61 |
| ENSMUSG00000062488 | Ifit3b | interferon-induced protein with tetratricopeptide repeats 3B | -1.61 |
| ENSMUSG00000024526 | Cidea | cell death-inducing DNA fragmentation factor, alpha subunit-like effector A | -1.61 |
| ENSMUSG00000031074 | Fgf3 | fibroblast growth factor 3 | -1.61 |
| ENSMUSG00000045875 | Adra1a | adrenergic receptor, alpha 1a | -1.61 |
| ENSMUSG00000020182 | Ddc | dopa decarboxylase | -1.62 |
| ENSMUSG00000038903 | Ccdc68 | coiled-coil domain containing 68 | -1.62 |
| ENSMUSG00000114601 | Gm48188 | predicted gene, 48188 | -1.62 |
| ENSMUSG00000052921 | Arhgef15 | Rho guanine nucleotide exchange factor (GEF) 15 | -1.62 |
| ENSMUSG00000025104 | Hdgfl3 | HDGF like 3 | -1.62 |
| ENSMUSG00000026765 | Lypd6b | LY6/PLAUR domain containing 6B | -1.62 |
| ENSMUSG00000027107 | Chrna1 | cholinergic receptor, nicotinic, alpha polypeptide 1 (muscle) | -1.62 |
| ENSMUSG00000104362 | Gm37928 | predicted gene, 37928 | -1.62 |
| ENSMUSG00000022358 | Fbxo32 | F-box protein 32 | -1.63 |
| ENSMUSG00000010307 | Tmem86a | transmembrane protein 86A | -1.63 |
| ENSMUSG00000102224 | 4930447F24Rik | RIKEN cDNA 4930447F24 gene | -1.63 |
| ENSMUSG00000025175 | Fn3k | fructosamine 3 kinase | -1.63 |
| ENSMUSG00000090743 | Gm17213 | predicted gene 17213 | -1.63 |
| ENSMUSG00000023328 | Ache | acetylcholinesterase | -1.64 |
| ENSMUSG00000061751 | Kalrn | kalirin, RhoGEF kinase | -1.64 |
| ENSMUSG00000040624 | Plekhg1 | pleckstrin homology domain containing, family G (with RhoGef domain) member 1 | -1.64 |
| ENSMUSG00000035429 | Ptprh | protein tyrosine phosphatase, receptor type, H | -1.64 |
| ENSMUSG00000040613 | Apobec1 | apolipoprotein B mRNA editing enzyme, catalytic polypeptide 1 | -1.64 |
| ENSMUSG00000031635 | Anxa10 | annexin A10 | -1.64 |
| ENSMUSG00000020573 | Pik3cg | phosphatidylinositol-4,5-bisphosphate 3-kinase catalytic subunit gamma | -1.64 |
| ENSMUSG00000074461 | Gm10699 | predicted gene 10699 | -1.64 |
| ENSMUSG00000030911 | Zp2 | zona pellucida glycoprotein 2 | -1.64 |
| ENSMUSG00000039234 | Sec24d | Sec24 related gene family, member D (S. cerevisiae) | -1.65 |
| ENSMUSG00000002233 | Rhoc | ras homolog family member C | -1.65 |
| ENSMUSG00000036553 | Sh3tc1 | SH3 domain and tetratricopeptide repeats 1 | -1.65 |
| ENSMUSG00000102423 | Gm37465 | predicted gene, 37465 | -1.65 |
| ENSMUSG00000055745 | Rtl6 | retrotransposon Gag like 6 | -1.65 |
| ENSMUSG00000111282 | Gm47528 | predicted gene, 47528 | -1.65 |
| ENSMUSG00000040310 | Alx4 | aristaless-like homeobox 4 | -1.65 |
| ENSMUSG00000099826 | Scgb2b10 | secretoglobin, family 2B, member 10 | -1.65 |
| ENSMUSG00000028713 | Cyp4b1 | cytochrome P450, family 4, subfamily b, polypeptide 1 | -1.66 |
| ENSMUSG00000021336 | Slc17a4 | solute carrier family 17 (sodium phosphate), member 4 | -1.66 |
| ENSMUSG00000044626 | Liph | lipase, member H | -1.66 |
| ENSMUSG00000003545 | Fosb | FBJ osteosarcoma oncogene B | -1.66 |
| ENSMUSG00000027230 | Creb3l1 | cAMP responsive element binding protein 3-like 1 | -1.66 |
| ENSMUSG00000021620 | Acot12 | acyl-CoA thioesterase 12 | -1.66 |
| ENSMUSG00000026405 | C4bp | complement component 4 binding protein | -1.66 |
| ENSMUSG00000010830 | Kdelr3 | KDEL (Lys-Asp-Glu-Leu) endoplasmic reticulum protein retention receptor 3 | -1.66 |
| ENSMUSG00000051166 | Eml5 | echinoderm microtubule associated protein like 5 | -1.66 |
| ENSMUSG00000037902 | Sirpa | signal-regulatory protein alpha | -1.66 |
| ENSMUSG00000097566 | A930024N18Rik | RIKEN cDNA A930024N18 gene | -1.66 |
| ENSMUSG00000035041 | Creb3l3 | cAMP responsive element binding protein 3-like 3 | -1.67 |
| ENSMUSG00000028943 | Espn | espin | -1.67 |
| ENSMUSG00000015944 | Castor2 | cytosolic arginine sensor for mTORC1 subunit 2 | -1.67 |
| ENSMUSG00000025268 | Maged2 | melanoma antigen, family D, 2 | -1.67 |
| ENSMUSG00000030930 | Chst15 | carbohydrate (N-acetylgalactosamine 4-sulfate 6-O) sulfotransferase 15 | -1.67 |
| ENSMUSG00000023966 | Rsph9 | radial spoke head 9 homolog (Chlamydomonas) | -1.67 |
| ENSMUSG00000005045 | Chd5 | chromodomain helicase DNA binding protein 5 | -1.67 |
| ENSMUSG00000085925 | Rtl1 | retrotransposon Gaglike 1 | -1.67 |
| ENSMUSG00000036923 | Stox1 | storkhead box 1 | -1.67 |
| ENSMUSG00000095280 | Gm21738 | predicted gene, 21738 | -1.67 |
| ENSMUSG00000100522 | Gm19280 | predicted gene, 19280 | -1.67 |
| ENSMUSG00000108693 | Gm45153 | predicted gene 45153 | -1.67 |
| ENSMUSG00000010064 | Slc38a3 | solute carrier family 38, member 3 | -1.67 |
| ENSMUSG00000024109 | Nrxn1 | neurexin I | -1.67 |
| ENSMUSG00000043458 | Pcdhb12 | protocadherin beta 12 | -1.67 |

| | | | |
|---|---|---|---|
| ENSMUSG00000009216 | Fam163b | family with sequence similarity 163, member B | -1.67 |
| ENSMUSG00000067212 | H2-T23 | histocompatibility 2, T region locus 23 | -1.68 |
| ENSMUSG00000043885 | Slc36a4 | solute carrier family 36 (proton/amino acid symporter), member 4 | -1.68 |
| ENSMUSG00000003863 | Ppfia3 | protein tyrosine phosphatase, receptor type, f polypeptide (PTPRF), interacting protein (liprin), alpha | -1.68 |
| ENSMUSG00000039021 | Ttc16 | tetratricopeptide repeat domain 16 | -1.68 |
| ENSMUSG00000039485 | Tspyl4 | TSPY-like 4 | -1.68 |
| ENSMUSG00000104140 | Gm37140 | predicted gene, 37140 | -1.68 |
| ENSMUSG00000031202 | Rab39b | RAB39B, member RAS oncogene family | -1.68 |
| ENSMUSG00000072115 | Ang | angiogenin, ribonuclease, RNase A family, 5 | -1.69 |
| ENSMUSG00000020681 | Ace | angiotensin I converting enzyme (peptidyl-dipeptidase A) 1 | -1.69 |
| ENSMUSG00000021208 | Ifi27l2b | interferon, alpha-inducible protein 27 like 2B | -1.69 |
| ENSMUSG00000025557 | Slc15a1 | solute carrier family 15 (oligopeptide transporter), member 1 | -1.69 |
| ENSMUSG00000042439 | Zfp532 | zinc finger protein 532 | -1.69 |
| ENSMUSG00000031824 | 6430548M08Rik | RIKEN cDNA 6430548M08 gene | -1.69 |
| ENSMUSG00000000627 | Sema4f | sema domain, immunoglobulin domain (Ig), TM domain, and short cytoplasmic domain | -1.69 |
| ENSMUSG00000027188 | Pamr1 | peptidase domain containing associated with muscle regeneration 1 | -1.69 |
| ENSMUSG00000020908 | Myh3 | myosin, heavy polypeptide 3, skeletal muscle, embryonic | -1.69 |
| ENSMUSG00000021876 | Rnase4 | ribonuclease, RNase A family 4 | -1.70 |
| ENSMUSG00000021792 | Fam213a | family with sequence similarity 213, member A | -1.70 |
| ENSMUSG00000036814 | Slc6a20a | solute carrier family 6 (neurotransmitter transporter), member 20A | -1.70 |
| ENSMUSG00000048070 | Pirt | phosphoinositide-interacting regulator of transient receptor potential channels | -1.70 |
| ENSMUSG00000068600 | Gml2 | glycosylphosphatidylinositol anchored molecule like 2 | -1.70 |
| ENSMUSG00000043614 | Vps37d | vacuolar protein sorting 37D | -1.70 |
| ENSMUSG00000034758 | Tle6 | transducin-like enhancer of split 6 | -1.70 |
| ENSMUSG00000020151 | Ptprr | protein tyrosine phosphatase, receptor type, R | -1.70 |
| ENSMUSG00000022594 | Lynx1 | Ly6/neurotoxin 1 | -1.70 |
| ENSMUSG00000009092 | Derl3 | Der1-like domain family, member 3 | -1.70 |
| ENSMUSG00000006457 | Actn3 | actinin alpha 3 | -1.70 |
| ENSMUSG00000055733 | Nap1l3 | nucleosome assembly protein 1-like 3 | -1.70 |
| ENSMUSG00000027068 | Dhrs9 | dehydrogenase/reductase (SDR family) member 9 | -1.71 |
| ENSMUSG00000019055 | Plod1 | procollagen-lysine, 2-oxoglutarate 5-dioxygenase 1 | -1.71 |
| ENSMUSG00000038042 | Ptpdc1 | protein tyrosine phosphatase domain containing 1 | -1.71 |
| ENSMUSG00000017446 | C1qtnf1 | C1q and tumor necrosis factor related protein 1 | -1.71 |
| ENSMUSG00000115834 | Gm10385 | predicted gene 10385 | -1.71 |
| ENSMUSG00000110767 | Gm31432 | predicted gene, 31432 | -1.71 |
| ENSMUSG00000034677 | Gpr142 | G protein-coupled receptor 142 | -1.71 |
| ENSMUSG00000038497 | Tmco3 | transmembrane and coiled-coil domains 3 | -1.72 |
| ENSMUSG00000033213 | AA467197 | expressed sequence AA467197 | -1.72 |
| ENSMUSG00000100183 | Gm28512 | predicted gene 28512 | -1.72 |
| ENSMUSG00000018486 | Wnt9b | wingless-type MMTV integration site family, member 9B | -1.72 |
| ENSMUSG00000005267 | Zfp287 | zinc finger protein 287 | -1.72 |
| ENSMUSG00000000214 | Th | tyrosine hydroxylase | -1.72 |
| ENSMUSG00000002228 | Ppm1j | protein phosphatase 1J | -1.73 |
| ENSMUSG00000068551 | Zfp467 | zinc finger protein 467 | -1.73 |
| ENSMUSG00000110405 | Gm45534 | predicted gene 45534 | -1.73 |
| ENSMUSG00000069456 | Rdh16 | retinol dehydrogenase 16 | -1.73 |
| ENSMUSG00000107497 | Gm44957 | predicted gene 44957 | -1.73 |
| ENSMUSG00000106825 | 2510016D11Rik | RIKEN cDNA 2510016D11 gene | -1.74 |
| ENSMUSG00000029154 | Cwh43 | cell wall biogenesis 43 C-terminal homolog | -1.74 |
| ENSMUSG00000075014 | Gm10800 | predicted gene 10800 | -1.74 |
| ENSMUSG00000040016 | Ptger3 | prostaglandin E receptor 3 (subtype EP3) | -1.74 |
| ENSMUSG00000111077 | Gm48743 | predicted gene, 48743 | -1.74 |
| ENSMUSG00000074344 | Tmigd3 | transmembrane and immunoglobulin domain containing 3 | -1.74 |
| ENSMUSG00000090706 | Gm17233 | predicted gene 17233 | -1.74 |
| ENSMUSG00000079174 | Gm3054 | predicted gene 3054 | -1.75 |
| ENSMUSG00000015094 | Npdc1 | neural proliferation, differentiation and control 1 | -1.75 |
| ENSMUSG00000032083 | Apoa1 | apolipoprotein A-I | -1.75 |
| ENSMUSG00000090891 | D6Ertd527e | DNA segment, Chr 6, ERATO Doi 527, expressed | -1.75 |
| ENSMUSG00000050447 | Lypd6 | LY6/PLAUR domain containing 6 | -1.75 |
| ENSMUSG00000032353 | Tmed3 | transmembrane p24 trafficking protein 3 | -1.76 |
| ENSMUSG00000019564 | Arid3a | AT rich interactive domain 3A (BRIGHT-like) | -1.76 |
| ENSMUSG00000060487 | Samd5 | sterile alpha motif domain containing 5 | -1.76 |
| ENSMUSG00000009739 | Pou6f1 | POU domain, class 6, transcription factor 1 | -1.76 |
| ENSMUSG00000034949 | Zfr2 | zinc finger RNA binding protein 2 | -1.76 |
| ENSMUSG00000000392 | Fap | fibroblast activation protein | -1.76 |
| ENSMUSG00000060063 | Alox5ap | arachidonate 5-lipoxygenase activating protein | -1.76 |
| ENSMUSG00000001827 | Folr1 | folate receptor 1 (adult) | -1.76 |
| ENSMUSG00000110301 | Gm35363 | predicted gene, 35363 | -1.76 |
| ENSMUSG00000047161 | Chst9 | carbohydrate (N-acetylgalactosamine 4-0) sulfotransferase 9 | -1.76 |
| ENSMUSG00000038807 | Rap1gap2 | RAP1 GTPase activating protein 2 | -1.77 |
| ENSMUSG00000043079 | Synpo | synaptopodin | -1.77 |
| ENSMUSG00000040270 | Bach2 | BTB and CNC homology, basic leucine zipper transcription factor 2 | -1.77 |
| ENSMUSG00000040415 | Dtx3 | deltex 3, E3 ubiquitin ligase | -1.77 |

| | | | |
|---|---|---|---|
| ENSMUSG00000029630 | Cyp3a25 | cytochrome P450, family 3, subfamily a, polypeptide 25 | -1.77 |
| ENSMUSG00000038305 | Spats2l | spermatogenesis associated, serine-rich 2-like | -1.78 |
| ENSMUSG00000026639 | Lamb3 | laminin, beta 3 | -1.78 |
| ENSMUSG00000032420 | Nt5e | 5' nucleotidase, ecto | -1.78 |
| ENSMUSG00000032549 | Rab6b | RAB6B, member RAS oncogene family | -1.78 |
| ENSMUSG00000055737 | Ghr | growth hormone receptor | -1.79 |
| ENSMUSG00000030364 | Clec2h | C-type lectin domain family 2, member h | -1.79 |
| ENSMUSG00000074892 | B3galt5 | UDP-Gal:betaGlcNAc beta 1,3-galactosyltransferase, polypeptide 5 | -1.79 |
| ENSMUSG00000071203 | Naip5 | NLR family, apoptosis inhibitory protein 5 | -1.79 |
| ENSMUSG00000027790 | Sis | sucrase isomaltase (alpha-glucosidase) | -1.79 |
| ENSMUSG00000087651 | 1500009L16Rik | RIKEN cDNA 1500009L16 gene | -1.79 |
| ENSMUSG00000034438 | Gbp8 | guanylate-binding protein 8 | -1.79 |
| ENSMUSG00000030228 | Pik3c2g | phosphatidylinositol-4-phosphate 3-kinase catalytic subunit type 2 gamma | -1.79 |
| ENSMUSG00000081270 | Gm11653 | predicted gene 11653 | -1.79 |
| ENSMUSG00000024530 | Prelid3a | PRELI domain containing 3A | -1.79 |
| ENSMUSG00000114113 | Gm47639 | predicted gene, 47639 | -1.79 |
| ENSMUSG00000038375 | Trp53inp2 | transformation related protein 53 inducible nuclear protein 2 | -1.80 |
| ENSMUSG00000020646 | Mboat2 | membrane bound O-acyltransferase domain containing 2 | -1.80 |
| ENSMUSG00000033427 | Upb1 | ureidopropionase, beta | -1.80 |
| ENSMUSG00000020062 | Slc5a8 | solute carrier family 5 (iodide transporter), member 8 | -1.80 |
| ENSMUSG00000017868 | Sgk2 | serum/glucocorticoid regulated kinase 2 | -1.80 |
| ENSMUSG00000060735 | Rxfp3 | relaxin family peptide receptor 3 | -1.80 |
| ENSMUSG00000006576 | Slc4a3 | solute carrier family 4 (anion exchanger), member 3 | -1.80 |
| ENSMUSG00000023826 | Prkn | parkin RBR E3 ubiquitin protein ligase | -1.80 |
| ENSMUSG00000104211 | Gm37985 | predicted gene, 37985 | -1.80 |
| ENSMUSG00000102424 | Paupar | Pax6 upstream antisense RNA | -1.80 |
| ENSMUSG00000042269 | Fam92b | family with sequence similarity 92, member B | -1.80 |
| ENSMUSG00000029185 | Fam114a1 | family with sequence similarity 114, member A1 | -1.81 |
| ENSMUSG00000049690 | Nckap5 | NCK-associated protein 5 | -1.81 |
| ENSMUSG00000018623 | Mmp7 | matrix metallopeptidase 7 | -1.81 |
| ENSMUSG00000029273 | Sult1d1 | sulfotransferase family 1D, member 1 | -1.81 |
| ENSMUSG00000030659 | Nucb2 | nucleobindin 2 | -1.81 |
| ENSMUSG00000029156 | Sgcb | sarcoglycan, beta (dystrophin-associated glycoprotein) | -1.81 |
| ENSMUSG00000044072 | Eml6 | echinoderm microtubule associated protein like 6 | -1.81 |
| ENSMUSG00000044748 | Defb1 | defensin beta 1 | -1.81 |
| ENSMUSG00000052387 | Trpm3 | transient receptor potential cation channel, subfamily M, member 3 | -1.81 |
| ENSMUSG00000025006 | Sorbs1 | sorbin and SH3 domain containing 1 | -1.82 |
| ENSMUSG00000017652 | Cd40 | CD40 antigen | -1.82 |
| ENSMUSG00000047143 | Dmrta2 | doublesex and mab-3 related transcription factor like family A2 | -1.82 |
| ENSMUSG00000087326 | Gm12503 | predicted gene 12503 | -1.82 |
| ENSMUSG00000017692 | Rhbdl3 | rhomboid like 3 | -1.82 |
| ENSMUSG00000020363 | Gfpt2 | glutamine fructose-6-phosphate transaminase 2 | -1.82 |
| ENSMUSG00000046169 | Adamts6 | a disintegrin-like and metallopeptidase (reprolysin type) with thrombospondin type 1 motif, 6 | -1.82 |
| ENSMUSG00000050600 | Zfp831 | zinc finger protein 831 | -1.82 |
| ENSMUSG00000060935 | Tmem263 | transmembrane protein 263 | -1.83 |
| ENSMUSG00000037606 | Osbpl5 | oxysterol binding protein-like 5 | -1.83 |
| ENSMUSG00000050747 | Trim15 | tripartite motif-containing 15 | -1.83 |
| ENSMUSG00000032186 | Tmod2 | tropomodulin 2 | -1.83 |
| ENSMUSG00000017211 | Gsdma2 | gasdermin A2 | -1.83 |
| ENSMUSG00000028716 | Pdzk1ip1 | PDZK1 interacting protein 1 | -1.83 |
| ENSMUSG00000032773 | Chrm1 | cholinergic receptor, muscarinic 1, CNS | -1.83 |
| ENSMUSG00000046908 | Ltb4r1 | leukotriene B4 receptor 1 | -1.83 |
| ENSMUSG00000009394 | Syn2 | synapsin II | -1.83 |
| ENSMUSG00000061958 | Gm14851 | predicted gene 14851 | -1.83 |
| ENSMUSG00000102184 | Gm37192 | predicted gene, 37192 | -1.83 |
| ENSMUSG00000034731 | Dgkh | diacylglycerol kinase, eta | -1.84 |
| ENSMUSG00000022602 | Arc | activity regulated cytoskeletal-associated protein | -1.84 |
| ENSMUSG00000001663 | Gstt1 | glutathione S-transferase, theta 1 | -1.84 |
| ENSMUSG00000028246 | Faxc | failed axon connections homolog | -1.84 |
| ENSMUSG00000084934 | Gm16035 | predicted gene 16035 | -1.84 |
| ENSMUSG00000028845 | Tekt2 | tektin 2 | -1.84 |
| ENSMUSG00000063873 | Slc24a3 | solute carrier family 24 (sodium/potassium/calcium exchanger), member 3 | -1.84 |
| ENSMUSG00000103284 | 3110080O07Rik | RIKEN cDNA 3110080O07 gene | -1.85 |
| ENSMUSG00000108427 | Gm36696 | predicted gene, 36696 | -1.85 |
| ENSMUSG00000045518 | Onecut3 | one cut domain, family member 3 | -1.85 |
| ENSMUSG00000054934 | Kcnmb4 | potassium large conductance calcium-activated channel, subfamily M, beta member 4 | -1.85 |
| ENSMUSG00000060962 | Dmkn | dermokine | -1.85 |
| ENSMUSG00000112601 | Gm47590 | predicted gene, 47590 | -1.85 |
| ENSMUSG00000020990 | Cdkl1 | cyclin-dependent kinase-like 1 (CDC2-related kinase) | -1.85 |
| ENSMUSG00000029275 | Gfi1 | growth factor independent 1 | -1.86 |
| ENSMUSG00000032816 | Igdcc4 | immunoglobulin superfamily, DCC subclass, member 4 | -1.86 |
| ENSMUSG00000015981 | Stk32c | serine/threonine kinase 32C | -1.86 |
| ENSMUSG00000050821 | Fam131a | family with sequence similarity 131, member A | -1.86 |

| | | | |
|---|---|---|---|
| ENSMUSG00000015243 | Abca1 | ATP-binding cassette, sub-family A (ABC1), member 1 | -1.87 |
| ENSMUSG00000033917 | Gde1 | glycerophosphodiester phosphodiesterase 1 | -1.87 |
| ENSMUSG00000049191 | Rtl5 | retrotransposon Gag like 5 | -1.87 |
| ENSMUSG00000021765 | Fst | follistatin | -1.87 |
| ENSMUSG00000033209 | Ttc28 | tetratricopeptide repeat domain 28 | -1.87 |
| ENSMUSG00000103701 | Gm10728 | predicted gene 10728 | -1.87 |
| ENSMUSG00000006711 | D130043K22Rik | RIKEN cDNA D130043K22 gene | -1.88 |
| ENSMUSG00000029798 | Herc6 | hect domain and RLD 6 | -1.88 |
| ENSMUSG00000008153 | Clstn3 | calsyntenin 3 | -1.88 |
| ENSMUSG00000040387 | Klhl32 | kelch-like 32 | -1.88 |
| ENSMUSG00000020953 | Coch | cochlin | -1.88 |
| ENSMUSG00000046159 | Chrm3 | cholinergic receptor, muscarinic 3, cardiac | -1.89 |
| ENSMUSG00000103585 | Pcdhgb4 | protocadherin gamma subfamily B, 4 | -1.89 |
| ENSMUSG00000030087 | Klf15 | Kruppel-like factor 15 | -1.89 |
| ENSMUSG00000024065 | Ehd3 | EH-domain containing 3 | -1.89 |
| ENSMUSG00000047904 | Sstr2 | somatostatin receptor 2 | -1.89 |
| ENSMUSG00000108348 | Gm42372 | predicted gene, 42372 | -1.89 |
| ENSMUSG00000026070 | Il18r1 | interleukin 18 receptor 1 | -1.89 |
| ENSMUSG00000002633 | Shh | sonic hedgehog | -1.89 |
| ENSMUSG00000026335 | Pam | peptidylglycine alpha-amidating monooxygenase | -1.90 |
| ENSMUSG00000033684 | Qsox1 | quiescin Q6 sulfhydryl oxidase 1 | -1.90 |
| ENSMUSG00000034459 | Ifit1 | interferon-induced protein with tetratricopeptide repeats 1 | -1.90 |
| ENSMUSG00000031791 | Tmem38a | transmembrane protein 38A | -1.90 |
| ENSMUSG00000025701 | Alox5 | arachidonate 5-lipoxygenase | -1.90 |
| ENSMUSG00000104136 | Gm36955 | predicted gene, 36955 | -1.90 |
| ENSMUSG00000029428 | Stx2 | syntaxin 2 | -1.90 |
| ENSMUSG00000040148 | Hmx3 | H6 homeobox 3 | -1.90 |
| ENSMUSG00000114282 | 5330431K02Rik | RIKEN cDNA 5330431K02 gene | -1.90 |
| ENSMUSG00000085329 | 2810404F17Rik | RIKEN cDNA 2810404F17 gene | -1.91 |
| ENSMUSG00000038541 | Srd5a2 | steroid 5 alpha-reductase 2 | -1.91 |
| ENSMUSG00000072623 | Zfp9 | zinc finger protein 9 | -1.91 |
| ENSMUSG00000038271 | Iffo1 | intermediate filament family orphan 1 | -1.91 |
| ENSMUSG00000106320 | Gm42992 | predicted gene 42992 | -1.91 |
| ENSMUSG00000039419 | Cntnap2 | contactin associated protein-like 2 | -1.91 |
| ENSMUSG00000010601 | Apol7a | apolipoprotein L 7a | -1.92 |
| ENSMUSG00000005672 | Kit | KIT proto-oncogene receptor tyrosine kinase | -1.92 |
| ENSMUSG00000029408 | Abcb9 | ATP-binding cassette, sub-family B (MDR/TAP), member 9 | -1.92 |
| ENSMUSG00000073802 | Cdkn2b | cyclin dependent kinase inhibitor 2B | -1.92 |
| ENSMUSG00000004933 | Matk | megakaryocyte-associated tyrosine kinase | -1.92 |
| ENSMUSG00000023868 | Pde10a | phosphodiesterase 10A | -1.92 |
| ENSMUSG00000020882 | Cacnb1 | calcium channel, voltage-dependent, beta 1 subunit | -1.92 |
| ENSMUSG00000111014 | Gm47795 | predicted gene, 47795 | -1.92 |
| ENSMUSG00000022580 | Rhpn1 | rhophilin, Rho GTPase binding protein 1 | -1.92 |
| ENSMUSG00000021379 | Id4 | inhibitor of DNA binding 4 | -1.93 |
| ENSMUSG00000051832 | E230016K23Rik | RIKEN cDNA E230016K23 gene | -1.93 |
| ENSMUSG00000020435 | Osbp2 | oxysterol binding protein 2 | -1.93 |
| ENSMUSG00000068196 | Col8a1 | collagen, type VIII, alpha 1 | -1.93 |
| ENSMUSG00000114626 | Gm48266 | predicted gene, 48266 | -1.93 |
| ENSMUSG00000067889 | Sptbn2 | spectrin beta, non-erythrocytic 2 | -1.94 |
| ENSMUSG00000035930 | Chst4 | carbohydrate (chondroitin 6/keratan) sulfotransferase 4 | -1.94 |
| ENSMUSG00000034116 | Vav1 | vav 1 oncogene | -1.94 |
| ENSMUSG00000042312 | S100a13 | S100 calcium binding protein A13 | -1.94 |
| ENSMUSG00000050212 | Eva1b | eva-1 homolog B (C. elegans) | -1.94 |
| ENSMUSG00000084950 | Gm5577 | predicted gene 5577 | -1.94 |
| ENSMUSG00000027249 | F2 | coagulation factor II | -1.94 |
| ENSMUSG00000053168 | 9030619P08Rik | RIKEN cDNA 9030619P08 gene | -1.94 |
| ENSMUSG00000032101 | Ddx25 | DEAD (Asp-Glu-Ala-Asp) box polypeptide 25 | -1.94 |
| ENSMUSG00000018217 | Pmp22 | peripheral myelin protein 22 | -1.95 |
| ENSMUSG00000041120 | Nbl1 | neuroblastoma, suppression of tumorigenicity 1 | -1.95 |
| ENSMUSG00000027612 | Mmp24 | matrix metallopeptidase 24 | -1.95 |
| ENSMUSG00000080115 | Eef1akmt3 | EEF1A lysine methyltransferase 3 | -1.95 |
| ENSMUSG00000018822 | Sfrp5 | secreted frizzled-related sequence protein 5 | -1.95 |
| ENSMUSG00000059900 | Tmem40 | transmembrane protein 40 | -1.95 |
| ENSMUSG00000029359 | Tesc | tescalcin | -1.96 |
| ENSMUSG00000074442 | Defa31 | defensin, alpha, 31 | -1.96 |
| ENSMUSG00000110131 | Gm18066 | predicted gene, 18066 | -1.96 |
| ENSMUSG00000103331 | Gm37995 | predicted gene, 37995 | -1.96 |
| ENSMUSG00000002980 | Bcam | basal cell adhesion molecule | -1.97 |
| ENSMUSG00000026961 | Lrrc26 | leucine rich repeat containing 26 | -1.97 |
| ENSMUSG00000078670 | Fam174b | family with sequence similarity 174, member B | -1.97 |
| ENSMUSG00000027801 | Tm4sf4 | transmembrane 4 superfamily member 4 | -1.97 |
| ENSMUSG00000073043 | Atoh1 | atonal bHLH transcription factor 1 | -1.98 |
| ENSMUSG00000000202 | Btbd17 | BTB (POZ) domain containing 17 | -1.98 |

| | | | |
|---|---|---|---|
| ENSMUSG00000066516 | Klk1b21 | kallikrein 1-related peptidase b21 | -1.98 |
| ENSMUSG00000069114 | Zbtb10 | zinc finger and BTB domain containing 10 | -1.99 |
| ENSMUSG00000031845 | Bco1 | beta-carotene oxygenase 1 | -1.99 |
| ENSMUSG00000035509 | Fbxl21 | F-box and leucine-rich repeat protein 21 | -1.99 |
| ENSMUSG00000078161 | Erich3 | glutamate rich 3 | -1.99 |
| ENSMUSG00000034460 | Six4 | sine oculis-related homeobox 4 | -1.99 |
| ENSMUSG00000007783 | Cpt1c | carnitine palmitoyltransferase 1c | -1.99 |
| ENSMUSG00000027175 | Tcp11l1 | t-complex 11 like 1 | -2.00 |
| ENSMUSG00000035067 | Xkr6 | X-linked Kx blood group related 6 | -2.00 |
| ENSMUSG00000038793 | Lefty1 | left right determination factor 1 | -2.00 |
| ENSMUSG00000040181 | Fmo1 | flavin containing monooxygenase 1 | -2.00 |
| ENSMUSG00000094818 | Defa32 | defensin, alpha, 32 | -2.00 |
| ENSMUSG00000017943 | Gdap1l1 | ganglioside-induced differentiation-associated protein 1-like 1 | -2.00 |
| ENSMUSG00000078650 | G6pc | glucose-6-phosphatase, catalytic | -2.01 |
| ENSMUSG00000057069 | Ero1lb | ERO1-like beta (S. cerevisiae) | -2.01 |
| ENSMUSG00000025743 | Sdc3 | syndecan 3 | -2.01 |
| ENSMUSG00000049804 | Armcx4 | armadillo repeat containing, X-linked 4 | -2.01 |
| ENSMUSG00000026797 | Stxbp1 | syntaxin binding protein 1 | -2.02 |
| ENSMUSG00000053617 | Sh3pxd2a | SH3 and PX domains 2A | -2.02 |
| ENSMUSG00000048330 | Ric3 | RIC3 acetylcholine receptor chaperone | -2.02 |
| ENSMUSG00000038587 | Akap12 | A kinase (PRKA) anchor protein (gravin) 12 | -2.02 |
| ENSMUSG00000063687 | Pcdhb5 | protocadherin beta 5 | -2.02 |
| ENSMUSG00000016179 | Camk1g | calcium/calmodulin-dependent protein kinase I gamma | -2.02 |
| ENSMUSG00000027938 | Creb3l4 | cAMP responsive element binding protein 3-like 4 | -2.03 |
| ENSMUSG00000094651 | Gal3st2 | galactose-3-O-sulfotransferase 2 | -2.03 |
| ENSMUSG00000053046 | Brsk2 | BR serine/threonine kinase 2 | -2.03 |
| ENSMUSG00000032661 | Oas3 | 2'-5' oligoadenylate synthetase 3 | -2.03 |
| ENSMUSG00000019944 | Rhobtb1 | Rho-related BTB domain containing 1 | -2.03 |
| ENSMUSG00000060508 | Nlrp9b | NLR family, pyrin domain containing 9B | -2.03 |
| ENSMUSG00000046191 | Pcdhb20 | protocadherin beta 20 | -2.03 |
| ENSMUSG00000048481 | Mypop | Myb-related transcription factor, partner of profilin | -2.03 |
| ENSMUSG00000028865 | Cd164l2 | CD164 sialomucin-like 2 | -2.03 |
| ENSMUSG00000117220 | CT025671.1 | novel transcript | -2.03 |
| ENSMUSG00000064373 | Selenop | selenoprotein P | -2.04 |
| ENSMUSG00000020230 | Prmt2 | protein arginine N-methyltransferase 2 | -2.04 |
| ENSMUSG00000045312 | Lhfpl2 | lipoma HMGIC fusion partner-like 2 | -2.04 |
| ENSMUSG00000042286 | Stab1 | stabilin 1 | -2.04 |
| ENSMUSG00000026817 | Ak1 | adenylate kinase 1 | -2.04 |
| ENSMUSG00000093580 | Gm20706 | predicted gene 20706 | -2.04 |
| ENSMUSG00000037973 | Ccdc129 | coiled-coil domain containing 129 | -2.04 |
| ENSMUSG00000031028 | Tub | tubby bipartite transcription factor | -2.04 |
| ENSMUSG00000052584 | Serp2 | stress-associated endoplasmic reticulum protein family member 2 | -2.04 |
| ENSMUSG00000019933 | Mrln | myoregulin | -2.04 |
| ENSMUSG00000026834 | Acvr1c | activin A receptor, type IC | -2.05 |
| ENSMUSG00000049281 | Scn3b | sodium channel, voltage-gated, type III, beta | -2.05 |
| ENSMUSG00000032268 | Tmprss5 | transmembrane protease, serine 5 (spinesin) | -2.05 |
| ENSMUSG00000021219 | Rgs6 | regulator of G-protein signaling 6 | -2.05 |
| ENSMUSG00000045094 | Arhgef37 | Rho guanine nucleotide exchange factor (GEF) 37 | -2.06 |
| ENSMUSG00000032609 | Klhdc8b | kelch domain containing 8B | -2.06 |
| ENSMUSG00000032740 | Ccdc88a | coiled coil domain containing 88A | -2.06 |
| ENSMUSG00000073680 | Tmem88b | transmembrane protein 88B | -2.06 |
| ENSMUSG00000038738 | Shank1 | SH3 and multiple ankyrin repeat domains 1 | -2.06 |
| ENSMUSG00000033805 | Ephx4 | epoxide hydrolase 4 | -2.06 |
| ENSMUSG00000053675 | Tgm5 | transglutaminase 5 | -2.06 |
| ENSMUSG00000029650 | Slc46a3 | solute carrier family 46, member 3 | -2.07 |
| ENSMUSG00000052131 | Akr1b7 | aldo-keto reductase family 1, member B7 | -2.07 |
| ENSMUSG00000001156 | Mxd1 | MAX dimerization protein 1 | -2.07 |
| ENSMUSG00000038175 | Mylip | myosin regulatory light chain interacting protein | -2.07 |
| ENSMUSG00000018927 | Ccl6 | chemokine (C-C motif) ligand 6 | -2.07 |
| ENSMUSG00000037341 | Slc9a7 | solute carrier family 9 (sodium/hydrogen exchanger), member 7 | -2.07 |
| ENSMUSG00000006784 | Ttc25 | tetratricopeptide repeat domain 25 | -2.07 |
| ENSMUSG00000038011 | Dnah10 | dynein, axonemal, heavy chain 10 | -2.08 |
| ENSMUSG00000029797 | Sspo | SCO-spondin | -2.08 |
| ENSMUSG00000030866 | Ern2 | endoplasmic reticulum (ER) to nucleus signalling 2 | -2.09 |
| ENSMUSG00000020581 | Agr2 | anterior gradient 2 | -2.09 |
| ENSMUSG00000078486 | Perm1 | PPARGC1 and ESRR induced regulator, muscle 1 | -2.09 |
| ENSMUSG00000063704 | Mapk15 | mitogen-activated protein kinase 15 | -2.09 |
| ENSMUSG00000093916 | Gm379 | predicted gene 379 | -2.09 |
| ENSMUSG00000048148 | Nwd1 | NACHT and WD repeat domain containing 1 | -2.09 |
| ENSMUSG00000004951 | Hspb1 | heat shock protein 1 | -2.09 |
| ENSMUSG00000070577 | Gm572 | predicted gene 572 | -2.09 |
| ENSMUSG00000069171 | Nr2f1 | nuclear receptor subfamily 2, group F, member 1 | -2.09 |
| ENSMUSG00000035580 | Kcnh8 | potassium voltage-gated channel, subfamily H (eag-related), member 8 | -2.10 |

| | | | |
|---|---|---|---|
| ENSMUSG00000022650 | Retnlb | resistin like beta | -2.10 |
| ENSMUSG00000035208 | Slfn8 | schlafen 8 | -2.10 |
| ENSMUSG00000097520 | 4930488L21Rik | RIKEN cDNA 4930488L21 gene | -2.10 |
| ENSMUSG00000114751 | 5430425E15Rik | RIKEN cDNA 5430425E15 gene | -2.10 |
| ENSMUSG00000032978 | Guca2b | guanylate cyclase activator 2b (retina) | -2.11 |
| ENSMUSG00000017204 | Gsdma | gasdermin A | -2.11 |
| ENSMUSG00000027513 | Pck1 | phosphoenolpyruvate carboxykinase 1, cytosolic | -2.11 |
| ENSMUSG00000024124 | Prss30 | protease, serine 30 | -2.11 |
| ENSMUSG00000054893 | Zfp667 | zinc finger protein 667 | -2.11 |
| ENSMUSG00000115186 | Gm49417 | predicted gene, 49417 | -2.11 |
| ENSMUSG00000031442 | Mcf2l | mcf.2 transforming sequence-like | -2.12 |
| ENSMUSG00000024063 | Lbh | limb-bud and heart | -2.12 |
| ENSMUSG00000049892 | Rasd1 | RAS, dexamethasone-induced 1 | -2.12 |
| ENSMUSG00000056031 | 9330154J02Rik | RIKEN cDNA 9330154J02 gene | -2.12 |
| ENSMUSG00000023927 | Satb1 | special AT-rich sequence binding protein 1 | -2.12 |
| ENSMUSG00000074439 | Defa5 | defensin, alpha, 5 | -2.12 |
| ENSMUSG00000074882 | Cyp2c68 | cytochrome P450, family 2, subfamily c, polypeptide 68 | -2.13 |
| ENSMUSG00000022469 | Rapgef3 | Rap guanine nucleotide exchange factor (GEF) 3 | -2.13 |
| ENSMUSG00000011832 | Evi5l | ecotropic viral integration site 5 like | -2.13 |
| ENSMUSG00000028634 | Hivep3 | human immunodeficiency virus type I enhancer binding protein 3 | -2.13 |
| ENSMUSG00000045348 | Nyap1 | neuronal tyrosine-phosphorylated phosphoinositide 3-kinase adaptor 1 | -2.13 |
| ENSMUSG00000019817 | Plagl1 | pleiomorphic adenoma gene-like 1 | -2.13 |
| ENSMUSG00000044162 | Tnip3 | TNFAIP3 interacting protein 3 | -2.13 |
| ENSMUSG00000021087 | Rtn1 | reticulon 1 | -2.13 |
| ENSMUSG00000025324 | Atp10a | ATPase, class V, type 10A | -2.13 |
| ENSMUSG00000104886 | Gm43000 | predicted gene 43000 | -2.13 |
| ENSMUSG00000070777 | Ceacam20 | carcinoembryonic antigen-related cell adhesion molecule 20 | -2.14 |
| ENSMUSG00000005973 | Rcn1 | reticulocalbin 1 | -2.14 |
| ENSMUSG00000026888 | Grb14 | growth factor receptor bound protein 14 | -2.14 |
| ENSMUSG00000035200 | Chrnb4 | cholinergic receptor, nicotinic, beta polypeptide 4 | -2.14 |
| ENSMUSG00000020429 | Igfbp1 | insulin-like growth factor binding protein 1 | -2.14 |
| ENSMUSG00000006241 | Ccdc159 | coiled-coil domain containing 159 | -2.14 |
| ENSMUSG00000006362 | Cbfa2t3 | core-binding factor, runt domain, alpha subunit 2, translocated to, 3 (human) | -2.15 |
| ENSMUSG00000020641 | Rsad2 | radical S-adenosyl methionine domain containing 2 | -2.15 |
| ENSMUSG00000030483 | Cyp2b10 | cytochrome P450, family 2, subfamily b, polypeptide 10 | -2.15 |
| ENSMUSG00000074899 | Sptbn5 | spectrin beta, non-erythrocytic 5 | -2.15 |
| ENSMUSG00000092517 | Art2a-ps | ADP-ribosyltransferase 2a, pseudogene | -2.15 |
| ENSMUSG00000042804 | Gpr153 | G protein-coupled receptor 153 | -2.15 |
| ENSMUSG00000021609 | Slc6a3 | solute carrier family 6 (neurotransmitter transporter, dopamine), member 3 | -2.15 |
| ENSMUSG00000024924 | Vldlr | very low density lipoprotein receptor | -2.16 |
| ENSMUSG00000080316 | Spaca6 | sperm acrosome associated 6 | -2.16 |
| ENSMUSG00000022548 | Apod | apolipoprotein D | -2.16 |
| ENSMUSG00000036437 | Npy1r | neuropeptide Y receptor Y1 | -2.16 |
| ENSMUSG00000061414 | Cracr2a | calcium release activated channel regulator 2A | -2.17 |
| ENSMUSG00000025656 | Arhgef9 | CDC42 guanine nucleotide exchange factor (GEF) 9 | -2.17 |
| ENSMUSG00000022012 | Enox1 | ecto-NOX disulfide-thiol exchanger 1 | -2.17 |
| ENSMUSG00000021913 | Ogdhl | oxoglutarate dehydrogenase-like | -2.18 |
| ENSMUSG00000038167 | Plekhg6 | pleckstrin homology domain containing, family G (with RhoGef domain) member 6 | -2.19 |
| ENSMUSG00000075702 | Selenom | selenoprotein M | -2.19 |
| ENSMUSG00000033730 | Egr3 | early growth response 3 | -2.19 |
| ENSMUSG00000000530 | Acvrl1 | activin A receptor, type II-like 1 | -2.19 |
| ENSMUSG00000046387 | Pcdhb17 | protocadherin beta 17 | -2.19 |
| ENSMUSG00000108041 | Gm5154 | predicted gene 5154 | -2.19 |
| ENSMUSG00000103466 | Gm37247 | predicted gene, 37247 | -2.19 |
| ENSMUSG00000112307 | Gm48751 | predicted gene, 48751 | -2.20 |
| ENSMUSG00000051435 | Fhad1 | forkhead-associated (FHA) phosphopeptide binding domain 1 | -2.20 |
| ENSMUSG00000042515 | Mum1l1 | melanoma associated antigen (mutated) 1-like 1 | -2.20 |
| ENSMUSG00000026131 | Dst | dystonin | -2.21 |
| ENSMUSG00000024215 | Spdef | SAM pointed domain containing ets transcription factor | -2.21 |
| ENSMUSG00000037953 | A4gnt | alpha-1,4-N-acetylglucosaminyltransferase | -2.21 |
| ENSMUSG00000097266 | Gm26802 | predicted gene, 26802 | -2.21 |
| ENSMUSG00000037921 | Ddx60 | DEAD (Asp-Glu-Ala-Asp) box polypeptide 60 | -2.22 |
| ENSMUSG00000017639 | Rab11fip4 | RAB11 family interacting protein 4 (class II) | -2.22 |
| ENSMUSG00000025352 | Gdf11 | growth differentiation factor 11 | -2.22 |
| ENSMUSG00000019986 | Ahi1 | Abelson helper integration site 1 | -2.22 |
| ENSMUSG00000035863 | Palm | paralemmin | -2.22 |
| ENSMUSG00000011751 | Sptbn4 | spectrin beta, non-erythrocytic 4 | -2.22 |
| ENSMUSG00000037259 | Dzank1 | double zinc ribbon and ankyrin repeat domains 1 | -2.22 |
| ENSMUSG00000020788 | Atp2a3 | ATPase, Ca++ transporting, ubiquitous | -2.23 |
| ENSMUSG00000026828 | Galnt5 | polypeptide N-acetylgalactosaminyltransferase 5 | -2.23 |
| ENSMUSG00000050395 | Tnfsf15 | tumor necrosis factor (ligand) superfamily, member 15 | -2.23 |
| ENSMUSG00000039236 | Isg20 | interferon-stimulated protein | -2.23 |
| ENSMUSG00000000197 | Nalcn | sodium leak channel, non-selective | -2.23 |

| | | | |
|---|---|---|---|
| ENSMUSG00000036745 | Ttll7 | tubulin tyrosine ligase-like family, member 7 | -2.24 |
| ENSMUSG00000025584 | Pde8a | phosphodiesterase 8A | -2.24 |
| ENSMUSG00000053980 | Gm9930 | predicted gene 9930 | -2.24 |
| ENSMUSG00000089995 | Gm15716 | predicted gene 15716 | -2.24 |
| ENSMUSG00000028637 | Ccdc30 | coiled-coil domain containing 30 | -2.24 |
| ENSMUSG00000097927 | Gm6999 | predicted gene 6999 | -2.24 |
| ENSMUSG00000029700 | Slc13a1 | solute carrier family 13 (sodium/sulfate symporters), member 1 | -2.25 |
| ENSMUSG00000049044 | Rapgef4 | Rap guanine nucleotide exchange factor (GEF) 4 | -2.25 |
| ENSMUSG00000045763 | Basp1 | brain abundant, membrane attached signal protein 1 | -2.25 |
| ENSMUSG00000051335 | Gfod1 | glucose-fructose oxidoreductase domain containing 1 | -2.26 |
| ENSMUSG00000032548 | Slco2a1 | solute carrier organic anion transporter family, member 2a1 | -2.26 |
| ENSMUSG00000027514 | Zbp1 | Z-DNA binding protein 1 | -2.26 |
| ENSMUSG00000021986 | Amer2 | APC membrane recruitment 2 | -2.26 |
| ENSMUSG00000048965 | Mrgpre | MAS-related GPR, member E | -2.26 |
| ENSMUSG00000040061 | Plcb2 | phospholipase C, beta 2 | -2.26 |
| ENSMUSG00000044468 | Tent5c | terminal nucleotidyltransferase 5C | -2.27 |
| ENSMUSG00000075010 | AW112010 | expressed sequence AW112010 | -2.27 |
| ENSMUSG00000050321 | Neto1 | neuropilin (NRP) and tolloid (TLL)-like 1 | -2.27 |
| ENSMUSG00000022208 | Jph4 | junctophilin 4 | -2.27 |
| ENSMUSG00000084890 | A830036E02Rik | RIKEN cDNA A830036E02 gene | -2.27 |
| ENSMUSG00000109561 | Ankrd31 | ankyrin repeat domain 31 | -2.27 |
| ENSMUSG00000029102 | Hgfac | hepatocyte growth factor activator | -2.28 |
| ENSMUSG00000028270 | Gbp2 | guanylate binding protein 2 | -2.28 |
| ENSMUSG00000018169 | Mfng | MFNG O-fucosylpeptide 3-beta-N-acetylglucosaminyltransferase | -2.28 |
| ENSMUSG00000037217 | Syn1 | synapsin I | -2.28 |
| ENSMUSG00000045062 | Pcdhb7 | protocadherin beta 7 | -2.28 |
| ENSMUSG00000041729 | Coro2b | coronin, actin binding protein, 2B | -2.28 |
| ENSMUSG00000111323 | Gm48129 | predicted gene, 48129 | -2.28 |
| ENSMUSG00000060843 | Ctnna3 | catenin (cadherin associated protein), alpha 3 | -2.28 |
| ENSMUSG00000002565 | Scin | scinderin | -2.29 |
| ENSMUSG00000054434 | Tmem120b | transmembrane protein 120B | -2.29 |
| ENSMUSG00000103567 | Pcdhga5 | protocadherin gamma subfamily A, 5 | -2.29 |
| ENSMUSG00000034591 | Slc41a2 | solute carrier family 41, member 2 | -2.30 |
| ENSMUSG00000035274 | Tpbg | trophoblast glycoprotein | -2.30 |
| ENSMUSG00000039126 | Prune2 | prune homolog 2 | -2.31 |
| ENSMUSG00000063522 | 2010109I03Rik | RIKEN cDNA 2010109I03 gene | -2.31 |
| ENSMUSG00000041268 | Dmxl2 | Dmx-like 2 | -2.31 |
| ENSMUSG00000023809 | Rps6ka2 | ribosomal protein S6 kinase, polypeptide 2 | -2.31 |
| ENSMUSG00000038143 | Stox2 | storkhead box 2 | -2.31 |
| ENSMUSG00000074437 | Defa29 | defensin, alpha, 29 | -2.31 |
| ENSMUSG00000030329 | Pianp | PILR alpha associated neural protein | -2.31 |
| ENSMUSG00000029348 | Asphd2 | aspartate beta-hydroxylase domain containing 2 | -2.31 |
| ENSMUSG00000040152 | Thbs1 | thrombospondin 1 | -2.32 |
| ENSMUSG00000051177 | Plcb1 | phospholipase C, beta 1 | -2.32 |
| ENSMUSG00000108688 | Gm44985 | predicted gene 44985 | -2.32 |
| ENSMUSG00000003436 | Dll3 | delta like canonical Notch ligand 3 | -2.32 |
| ENSMUSG00000042388 | Dlgap3 | DLG associated protein 3 | -2.32 |
| ENSMUSG00000036915 | Kirrel2 | kirre like nephrin family adhesion molecule 2 | -2.32 |
| ENSMUSG00000025551 | Fgf14 | fibroblast growth factor 14 | -2.32 |
| ENSMUSG00000099148 | Gm3331 | predicted gene 3331 | -2.32 |
| ENSMUSG00000020599 | Rgs9 | regulator of G-protein signaling 9 | -2.33 |
| ENSMUSG00000047867 | Gimap6 | GTPase, IMAP family member 6 | -2.33 |
| ENSMUSG00000078235 | Fam43b | family with sequence similarity 43, member B | -2.33 |
| ENSMUSG00000036492 | Rnf39 | ring finger protein 39 | -2.34 |
| ENSMUSG00000058618 | AY761184 | cDNA sequence AY761184 | -2.34 |
| ENSMUSG00000044646 | Zbtb7c | zinc finger and BTB domain containing 7C | -2.34 |
| ENSMUSG00000020940 | 1700023F06Rik | RIKEN cDNA 1700023F06 gene | -2.34 |
| ENSMUSG00000033768 | Nrxn2 | neurexin II | -2.34 |
| ENSMUSG00000020019 | Ntn4 | netrin 4 | -2.35 |
| ENSMUSG00000033287 | Kctd17 | potassium channel tetramerisation domain containing 17 | -2.35 |
| ENSMUSG00000045912 | C2cd4c | C2 calcium-dependent domain containing 4C | -2.35 |
| ENSMUSG00000032017 | Grik4 | glutamate receptor, ionotropic, kainate 4 | -2.35 |
| ENSMUSG00000034111 | Tmed8 | transmembrane p24 trafficking protein 8 | -2.36 |
| ENSMUSG00000013846 | St3gal1 | ST3 beta-galactoside alpha-2,3-sialyltransferase 1 | -2.36 |
| ENSMUSG00000042834 | Nrep | neuronal regeneration related protein | -2.36 |
| ENSMUSG00000102732 | Gm37342 | predicted gene, 37342 | -2.36 |
| ENSMUSG00000074625 | Arhgap40 | Rho GTPase activating protein 40 | -2.36 |
| ENSMUSG00000037016 | Frem2 | Fras1 related extracellular matrix protein 2 | -2.36 |
| ENSMUSG00000063239 | Grm4 | glutamate receptor, metabotropic 4 | -2.36 |
| ENSMUSG00000113811 | Gm47882 | predicted gene, 47882 | -2.36 |
| ENSMUSG00000046215 | Rprml | reprimo-like | -2.36 |
| ENSMUSG00000049858 | Suox | sulfite oxidase | -2.37 |
| ENSMUSG00000030406 | Gipr | gastric inhibitory polypeptide receptor | -2.37 |

| | | | |
|---|---|---|---|
| ENSMUSG00000102723 | Gm37936 | predicted gene, 37936 | -2.37 |
| ENSMUSG00000046668 | Cxxc5 | CXXC finger 5 | -2.38 |
| ENSMUSG00000052271 | Bhlha15 | basic helix-loop-helix family, member a15 | -2.38 |
| ENSMUSG00000063160 | Numbl | numb-like | -2.38 |
| ENSMUSG00000007950 | Abhd8 | abhydrolase domain containing 8 | -2.38 |
| ENSMUSG00000041794 | Myrip | myosin VIIA and Rab interacting protein | -2.39 |
| ENSMUSG00000073102 | Drc1 | dynein regulatory complex subunit 1 | -2.39 |
| ENSMUSG00000043020 | Wdr63 | WD repeat domain 63 | -2.39 |
| ENSMUSG00000060402 | Chst8 | carbohydrate (N-acetylgalactosamine 4-0) sulfotransferase 8 | -2.40 |
| ENSMUSG00000053580 | Tanc2 | tetratricopeptide repeat, ankyrin repeat and coiled-coil containing 2 | -2.41 |
| ENSMUSG00000057897 | Camk2b | calcium/calmodulin-dependent protein kinase II, beta | -2.41 |
| ENSMUSG00000036306 | Lzts1 | leucine zipper, putative tumor suppressor 1 | -2.41 |
| ENSMUSG00000112197 | Gm47337 | predicted gene, 47337 | -2.41 |
| ENSMUSG00000091890 | A830073O21Rik | RIKEN cDNA A830073O21 gene | -2.41 |
| ENSMUSG00000043461 | Sptssb | serine palmitoyltransferase, small subunit B | -2.42 |
| ENSMUSG00000030107 | Usp18 | ubiquitin specific peptidase 18 | -2.42 |
| ENSMUSG00000033200 | Tpsg1 | tryptase gamma 1 | -2.42 |
| ENSMUSG00000043760 | Pkhd1 | polycystic kidney and hepatic disease 1 | -2.43 |
| ENSMUSG00000037627 | Rgs22 | regulator of G-protein signalling 22 | -2.43 |
| ENSMUSG00000101126 | Gm10538 | predicted gene 10538 | -2.43 |
| ENSMUSG00000037664 | Cdkn1c | cyclin-dependent kinase inhibitor 1C (P57) | -2.44 |
| ENSMUSG00000017897 | Eya2 | EYA transcriptional coactivator and phosphatase 2 | -2.44 |
| ENSMUSG00000114985 | 1810028F09Rik | RIKEN cDNA 1810028F09 gene | -2.44 |
| ENSMUSG00000024500 | Ppp2r2b | protein phosphatase 2, regulatory subunit B, beta | -2.44 |
| ENSMUSG00000042351 | Grap2 | GRB2-related adaptor protein 2 | -2.44 |
| ENSMUSG00000016763 | Scube1 | signal peptide, CUB domain, EGF-like 1 | -2.45 |
| ENSMUSG00000038594 | Cep85l | centrosomal protein 85-like | -2.45 |
| ENSMUSG00000070056 | Mfhas1 | malignant fibrous histiocytoma amplified sequence 1 | -2.46 |
| ENSMUSG00000086141 | 9030622O22Rik | RIKEN cDNA 9030622O22 gene | -2.46 |
| ENSMUSG00000024238 | Zeb1 | zinc finger E-box binding homeobox 1 | -2.46 |
| ENSMUSG00000035864 | Syt1 | synaptotagmin I | -2.46 |
| ENSMUSG00000039059 | Hrh3 | histamine receptor H3 | -2.47 |
| ENSMUSG00000039976 | Tbc1d16 | TBC1 domain family, member 16 | -2.48 |
| ENSMUSG00000020227 | Irak3 | interleukin-1 receptor-associated kinase 3 | -2.48 |
| ENSMUSG00000023009 | Nckap5l | NCK-associated protein 5-like | -2.48 |
| ENSMUSG00000028439 | Fam219a | family with sequence similarity 219, member A | -2.48 |
| ENSMUSG00000058740 | Kcnt1 | potassium channel, subfamily T, member 1 | -2.48 |
| ENSMUSG00000103001 | A930005N03Rik | RIKEN cDNA A930005N03 gene | -2.48 |
| ENSMUSG00000029166 | Mapre3 | microtubule-associated protein, RP/EB family, member 3 | -2.50 |
| ENSMUSG00000030717 | Nupr1 | nuclear protein transcription regulator 1 | -2.50 |
| ENSMUSG00000070337 | Gpr179 | G protein-coupled receptor 179 | -2.50 |
| ENSMUSG00000038319 | Kcnh2 | potassium voltage-gated channel, subfamily H (eag-related), member 2 | -2.50 |
| ENSMUSG00000004709 | Cd244a | CD244 molecule A | -2.50 |
| ENSMUSG00000000125 | Wnt3 | wingless-type MMTV integration site family, member 3 | -2.51 |
| ENSMUSG00000069227 | Gprin1 | G protein-regulated inducer of neurite outgrowth 1 | -2.51 |
| ENSMUSG00000048763 | Hoxb3 | homeobox B3 | -2.51 |
| ENSMUSG00000041889 | Shisa4 | shisa family member 4 | -2.51 |
| ENSMUSG00000026950 | Neb | nebulin | -2.52 |
| ENSMUSG00000020102 | Slc16a7 | solute carrier family 16 (monocarboxylic acid transporters), member 7 | -2.52 |
| ENSMUSG00000112433 | Gm30122 | predicted gene, 30122 | -2.52 |
| ENSMUSG00000019122 | Ccl9 | chemokine (C-C motif) ligand 9 | -2.53 |
| ENSMUSG00000080715 | Gm5406 | predicted gene 5406 | -2.53 |
| ENSMUSG00000103761 | Gm37859 | predicted gene, 37859 | -2.53 |
| ENSMUSG00000050824 | Sstr5 | somatostatin receptor 5 | -2.53 |
| ENSMUSG00000057182 | Scn3a | sodium channel, voltage-gated, type III, alpha | -2.53 |
| ENSMUSG00000024598 | Fbn2 | fibrillin 2 | -2.53 |
| ENSMUSG00000064413 | Gm22245 | predicted gene, 22245 | -2.53 |
| ENSMUSG00000039115 | Itga9 | integrin alpha 9 | -2.54 |
| ENSMUSG00000021013 | Ttc8 | tetratricopeptide repeat domain 8 | -2.54 |
| ENSMUSG00000038065 | Mturn | maturin, neural progenitor differentiation regulator homolog (Xenopus) | -2.55 |
| ENSMUSG00000057614 | Gnai1 | guanine nucleotide binding protein (G protein), alpha inhibiting 1 | -2.55 |
| ENSMUSG00000040183 | Ankrd6 | ankyrin repeat domain 6 | -2.55 |
| ENSMUSG00000029134 | Plb1 | phospholipase B1 | -2.55 |
| ENSMUSG00000106048 | Gm42444 | predicted gene 42444 | -2.56 |
| ENSMUSG00000036699 | Zcchc12 | zinc finger, CCHC domain containing 12 | -2.56 |
| ENSMUSG00000000031 | H19 | H19, imprinted maternally expressed transcript | -2.57 |
| ENSMUSG00000074444 | Defa30 | defensin, alpha, 30 | -2.57 |
| ENSMUSG00000006575 | Rundc3a | RUN domain containing 3A | -2.58 |
| ENSMUSG00000021725 | Parp8 | poly (ADP-ribose) polymerase family, member 8 | -2.58 |
| ENSMUSG00000062296 | Trank1 | tetratricopeptide repeat and ankyrin repeat containing 1 | -2.58 |
| ENSMUSG00000039543 | Cfap70 | cilia and flagella associated protein 70 | -2.58 |
| ENSMUSG00000005873 | Reep5 | receptor accessory protein 5 | -2.59 |
| ENSMUSG00000050822 | Slc29a4 | solute carrier family 29 (nucleoside transporters), member 4 | -2.60 |

| | | | |
|---|---|---|---|
| ENSMUSG00000024451 | Arap3 | ArfGAP with RhoGAP domain, ankyrin repeat and PH domain 3 | -2.60 |
| ENSMUSG00000049420 | Tmem200a | transmembrane protein 200A | -2.60 |
| ENSMUSG00000039005 | Tlr4 | toll-like receptor 4 | -2.60 |
| ENSMUSG00000029445 | Hpd | 4-hydroxyphenylpyruvic acid dioxygenase | -2.61 |
| ENSMUSG00000001333 | Sync | syncoilin | -2.61 |
| ENSMUSG00000078949 | R3hdml | R3H domain containing-like | -2.61 |
| ENSMUSG00000001281 | Itgb7 | integrin beta 7 | -2.61 |
| ENSMUSG00000026167 | Wnt10a | wingless-type MMTV integration site family, member 10A | -2.61 |
| ENSMUSG00000036960 | Clca2 | chloride channel accessory 2 | -2.61 |
| ENSMUSG00000015396 | Cd83 | CD83 antigen | -2.62 |
| ENSMUSG00000032068 | Plet1 | placenta expressed transcript 1 | -2.62 |
| ENSMUSG00000040367 | Lrrd1 | leucine rich repeats and death domain containing 1 | -2.62 |
| ENSMUSG00000037610 | Kcnmb2 | potassium large conductance calcium-activated channel, subfamily M, beta member 2 | -2.62 |
| ENSMUSG00000001670 | Tat | tyrosine aminotransferase | -2.63 |
| ENSMUSG00000038370 | Pcp4l1 | Purkinje cell protein 4-like 1 | -2.63 |
| ENSMUSG00000069515 | Lyz1 | lysozyme 1 | -2.63 |
| ENSMUSG00000037754 | Ppp1r16b | protein phosphatase 1, regulatory subunit 16B | -2.64 |
| ENSMUSG00000037579 | Kcnh3 | potassium voltage-gated channel, subfamily H (eag-related), member 3 | -2.64 |
| ENSMUSG00000060070 | Defa26 | defensin, alpha, 26 | -2.64 |
| ENSMUSG00000020175 | Rab36 | RAB36, member RAS oncogene family | -2.64 |
| ENSMUSG00000049047 | Armcx3 | armadillo repeat containing, X-linked 3 | -2.65 |
| ENSMUSG00000083718 | Ccnb2-ps | cyclin B2, pseudogene | -2.65 |
| ENSMUSG00000044312 | Neurog3 | neurogenin 3 | -2.65 |
| ENSMUSG00000100396 | Gm29367 | predicted gene 29367 | -2.65 |
| ENSMUSG00000031980 | Agt | angiotensinogen (serpin peptidase inhibitor, clade A, member 8) | -2.66 |
| ENSMUSG00000027223 | Mapk8ip1 | mitogen-activated protein kinase 8 interacting protein 1 | -2.67 |
| ENSMUSG00000029490 | Mfsd7a | major facilitator superfamily domain containing 7A | -2.67 |
| ENSMUSG00000034059 | Ypel4 | yippee like 4 | -2.67 |
| ENSMUSG00000006651 | Aplp1 | amyloid beta (A4) precursor-like protein 1 | -2.68 |
| ENSMUSG00000072235 | Tuba1a | tubulin, alpha 1A | -2.69 |
| ENSMUSG00000040495 | Chrm4 | cholinergic receptor, muscarinic 4 | -2.69 |
| ENSMUSG00000114245 | Gm38655 | predicted gene, 38655 | -2.69 |
| ENSMUSG00000046056 | Sbsn | suprabasin | -2.70 |
| ENSMUSG00000064043 | Trerf1 | transcriptional regulating factor 1 | -2.70 |
| ENSMUSG00000005883 | Spo11 | SPO11 meiotic protein covalently bound to DSB | -2.70 |
| ENSMUSG00000053182 | Gm609 | predicted gene 609 | -2.71 |
| ENSMUSG00000074441 | Gm15292 | predicted gene 15292 | -2.71 |
| ENSMUSG00000027376 | Prom2 | prominin 2 | -2.71 |
| ENSMUSG00000028078 | Dclk2 | doublecortin-like kinase 2 | -2.72 |
| ENSMUSG00000074981 | Dcdc5 | doublecortin domain containing 5 | -2.72 |
| ENSMUSG00000027520 | Zdbf2 | zinc finger, DBF-type containing 2 | -2.73 |
| ENSMUSG00000018340 | Anxa6 | annexin A6 | -2.74 |
| ENSMUSG00000036764 | Dnajc12 | DnaJ heat shock protein family (Hsp40) member C12 | -2.74 |
| ENSMUSG00000065956 | Defa37 | defensin, alpha, 37 | -2.74 |
| ENSMUSG00000025867 | Cplx2 | complexin 2 | -2.75 |
| ENSMUSG00000021730 | Hcn1 | hyperpolarization-activated, cyclic nucleotide-gated K+ 1 | -2.75 |
| ENSMUSG00000051079 | Rgs13 | regulator of G-protein signaling 13 | -2.75 |
| ENSMUSG00000028749 | Pla2g2f | phospholipase A2, group IIF | -2.76 |
| ENSMUSG00000025876 | Unc5a | unc-5 netrin receptor A | -2.76 |
| ENSMUSG00000066191 | Anks6 | ankyrin repeat and sterile alpha motif domain containing 6 | -2.76 |
| ENSMUSG00000021364 | Elovl2 | elongation of very long chain fatty acids (FEN1/Elo2, SUR4/Elo3, yeast)-like 2 | -2.76 |
| ENSMUSG00000015484 | Fam163a | family with sequence similarity 163, member A | -2.76 |
| ENSMUSG00000063903 | Klk1 | kallikrein 1 | -2.77 |
| ENSMUSG00000020668 | Kif3c | kinesin family member 3C | -2.77 |
| ENSMUSG00000057315 | Arhgap24 | Rho GTPase activating protein 24 | -2.77 |
| ENSMUSG00000062591 | Tubb4a | tubulin, beta 4A class IVA | -2.78 |
| ENSMUSG00000074447 | Defa21 | defensin, alpha, 21 | -2.78 |
| ENSMUSG00000109305 | 1810010D01Rik | RIKEN cDNA 1810010D01 gene | -2.78 |
| ENSMUSG00000021069 | Pygl | liver glycogen phosphorylase | -2.78 |
| ENSMUSG00000025091 | Pnliprp2 | pancreatic lipase-related protein 2 | -2.79 |
| ENSMUSG00000040412 | 5330417C22Rik | RIKEN cDNA 5330417C22 gene | -2.80 |
| ENSMUSG00000031981 | Capn9 | calpain 9 | -2.81 |
| ENSMUSG00000109780 | Gm45447 | predicted gene 45447 | -2.81 |
| ENSMUSG00000024743 | Syt7 | synaptotagmin VII | -2.82 |
| ENSMUSG00000069516 | Lyz2 | lysozyme 2 | -2.82 |
| ENSMUSG00000035896 | Rnase1 | ribonuclease, RNase A family, 1 (pancreatic) | -2.83 |
| ENSMUSG00000038209 | Itln1 | intelectin 1 (galactofuranose binding) | -2.83 |
| ENSMUSG00000074446 | Defa23 | defensin, alpha, 23 | -2.83 |
| ENSMUSG00000029778 | Adcyap1r1 | adenylate cyclase activating polypeptide 1 receptor 1 | -2.83 |
| ENSMUSG00000038599 | Capn8 | calpain 8 | -2.84 |
| ENSMUSG00000025969 | Nrp2 | neuropilin 2 | -2.84 |
| ENSMUSG00000037386 | Rims2 | regulating synaptic membrane exocytosis 2 | -2.84 |
| ENSMUSG00000028415 | Spink4 | serine peptidase inhibitor, Kazal type 4 | -2.85 |

| | | | |
|---|---|---|---|
| ENSMUSG00000040121 | Rep15 | RAB15 effector protein | -2.86 |
| ENSMUSG00000048022 | Tmem229a | transmembrane protein 229A | -2.86 |
| ENSMUSG00000071178 | Serpina1b | serine (or cysteine) preptidase inhibitor, clade A, member 1B | -2.86 |
| ENSMUSG00000013921 | Clip3 | CAP-GLY domain containing linker protein 3 | -2.86 |
| ENSMUSG00000075517 | Cyp2d37-ps | cytochrome P450, family 2, subfamily d, polypeptide 37, pseudogene | -2.86 |
| ENSMUSG00000028626 | Col9a2 | collagen, type IX, alpha 2 | -2.87 |
| ENSMUSG00000099025 | Gm27162 | predicted gene 27162 | -2.87 |
| ENSMUSG00000036832 | Lpar3 | lysophosphatidic acid receptor 3 | -2.87 |
| ENSMUSG00000026546 | Cfap45 | cilia and flagella associated protein 45 | -2.87 |
| ENSMUSG00000031377 | Bmx | BMX non-receptor tyrosine kinase | -2.88 |
| ENSMUSG00000026728 | Vim | vimentin | -2.88 |
| ENSMUSG00000072294 | Klf12 | Kruppel-like factor 12 | -2.88 |
| ENSMUSG00000064213 | Defa24 | defensin, alpha, 24 | -2.88 |
| ENSMUSG00000042873 | Lhfpl4 | lipoma HMGIC fusion partner-like protein 4 | -2.88 |
| ENSMUSG00000051242 | Pcdhb9 | protocadherin beta 9 | -2.88 |
| ENSMUSG00000060208 | Defa17 | defensin, alpha, 17 | -2.89 |
| ENSMUSG00000082443 | Gm12604 | predicted gene 12604 | -2.89 |
| ENSMUSG00000044933 | Sstr3 | somatostatin receptor 3 | -2.90 |
| ENSMUSG00000039013 | Siglecf | sialic acid binding Ig-like lectin F | -2.90 |
| ENSMUSG00000026959 | Grin1 | glutamate receptor, ionotropic, NMDA1 (zeta 1) | -2.91 |
| ENSMUSG00000041000 | Trim62 | tripartite motif-containing 62 | -2.92 |
| ENSMUSG00000030110 | Ret | ret proto-oncogene | -2.92 |
| ENSMUSG00000034107 | Ano7 | anoctamin 7 | -2.92 |
| ENSMUSG00000015968 | Cacna1d | calcium channel, voltage-dependent, L type, alpha 1D subunit | -2.93 |
| ENSMUSG00000018593 | Sparc | secreted acidic cysteine rich glycoprotein | -2.94 |
| ENSMUSG00000111086 | Gm48671 | predicted gene, 48671 | -2.94 |
| ENSMUSG00000068617 | Efcab1 | EF-hand calcium binding domain 1 | -2.95 |
| ENSMUSG00000032502 | Stac | src homology three (SH3) and cysteine rich domain | -2.96 |
| ENSMUSG00000032015 | Pou2f3 | POU domain, class 2, transcription factor 3 | -2.96 |
| ENSMUSG00000082211 | Defa27 | defensin, alpha, 27 | -2.96 |
| ENSMUSG00000044548 | Dact1 | dishevelled-binding antagonist of beta-catenin 1 | -2.97 |
| ENSMUSG00000033220 | Rac2 | Rac family small GTPase 2 | -2.97 |
| ENSMUSG00000024330 | Col11a2 | collagen, type XI, alpha 2 | -2.98 |
| ENSMUSG00000030772 | Dkk3 | dickkopf WNT signaling pathway inhibitor 3 | -2.98 |
| ENSMUSG00000002900 | Lamb1 | laminin B1 | -2.98 |
| ENSMUSG00000008932 | Slc1a7 | solute carrier family 1 (glutamate transporter), member 7 | -2.98 |
| ENSMUSG00000026870 | Cutal | cutA divalent cation tolerance homolog-like | -2.99 |
| ENSMUSG00000026482 | Rgl1 | ral guanine nucleotide dissociation stimulator,-like 1 | -2.99 |
| ENSMUSG00000019775 | Rgs17 | regulator of G-protein signaling 17 | -2.99 |
| ENSMUSG00000098557 | Kctd12 | potassium channel tetramerisation domain containing 12 | -3.00 |
| ENSMUSG00000055963 | Triqk | triple QxxK/R motif containing | -3.00 |
| ENSMUSG00000051486 | Pcdhb11 | protocadherin beta 11 | -3.00 |
| ENSMUSG00000051527 | Usp29 | ubiquitin specific peptidase 29 | -3.00 |
| ENSMUSG00000020901 | Pik3r5 | phosphoinositide-3-kinase regulatory subunit 5 | -3.01 |
| ENSMUSG00000093805 | Gal3st2b | galactose-3-O-sulfotransferase 2B | -3.01 |
| ENSMUSG00000000394 | Gcg | glucagon | -3.01 |
| ENSMUSG00000032202 | Rab27a | RAB27A, member RAS oncogene family | -3.02 |
| ENSMUSG00000040723 | Rcsd1 | RCSD domain containing 1 | -3.02 |
| ENSMUSG00000073400 | Trim10 | tripartite motif-containing 10 | -3.02 |
| ENSMUSG00000033174 | Mgll | monoglyceride lipase | -3.02 |
| ENSMUSG00000036412 | Arsi | arylsulfatase i | -3.02 |
| ENSMUSG00000023032 | Slc4a8 | solute carrier family 4 (anion exchanger), member 8 | -3.03 |
| ENSMUSG00000060615 | Ang4 | angiogenin, ribonuclease A family, member 4 | -3.03 |
| ENSMUSG00000032690 | Oas2 | 2'-5' oligoadenylate synthetase 2 | -3.03 |
| ENSMUSG00000086067 | Gm16183 | predicted gene 16183 | -3.03 |
| ENSMUSG00000023247 | Guca2a | guanylate cyclase activator 2a (guanylin) | -3.04 |
| ENSMUSG00000022211 | Carmil3 | capping protein regulator and myosin 1 linker 3 | -3.04 |
| ENSMUSG00000036533 | Cdc42ep3 | CDC42 effector protein (Rho GTPase binding) 3 | -3.04 |
| ENSMUSG00000045672 | Col27a1 | collagen, type XXVII, alpha 1 | -3.04 |
| ENSMUSG00000020121 | Srgap1 | SLIT-ROBO Rho GTPase activating protein 1 | -3.05 |
| ENSMUSG00000047181 | Samd14 | sterile alpha motif domain containing 14 | -3.05 |
| ENSMUSG00000072812 | Ahnak2 | AHNAK nucleoprotein 2 | -3.05 |
| ENSMUSG00000074443 | Defa22 | defensin, alpha, 22 | -3.05 |
| ENSMUSG00000020101 | Vsir | V-set immunoregulatory receptor | -3.05 |
| ENSMUSG00000040978 | Gm11992 | predicted gene 11992 | -3.06 |
| ENSMUSG00000020717 | Pecam1 | platelet/endothelial cell adhesion molecule 1 | -3.06 |
| ENSMUSG00000025964 | Adam23 | a disintegrin and metallopeptidase domain 23 | -3.06 |
| ENSMUSG00000058589 | Anks1b | ankyrin repeat and sterile alpha motif domain containing 1B | -3.06 |
| ENSMUSG00000001120 | Pcbp3 | poly(rC) binding protein 3 | -3.07 |
| ENSMUSG00000090812 | Samd15 | sterile alpha motif domain containing 15 | -3.07 |
| ENSMUSG00000079442 | St6galnac4 | ST6 (alpha-N-acetyl-neuraminyl-2,3-beta-galactosyl-1,3)-N-acetylgalactosaminide alpha-2,6-sialyltra | -3.08 |
| ENSMUSG00000027661 | Slc2a10 | solute carrier family 2 (facilitated glucose transporter), member 10 | -3.08 |
| ENSMUSG00000047642 | D930020B18Rik | RIKEN cDNA D930020B18 gene | -3.08 |

| | | | |
|---|---|---|---|
| ENSMUSG00000025075 | Habp2 | hyaluronic acid binding protein 2 | -3.09 |
| ENSMUSG00000006435 | Neurl1a | neuralized E3 ubiquitin protein ligase 1A | -3.09 |
| ENSMUSG00000026288 | Inpp5d | inositol polyphosphate-5-phosphatase D | -3.09 |
| ENSMUSG00000034818 | Celf5 | CUGBP, Elav-like family member 5 | -3.09 |
| ENSMUSG00000033152 | Podxl2 | podocalyxin-like 2 | -3.09 |
| ENSMUSG00000050359 | Sprr1a | small proline-rich protein 1A | -3.09 |
| ENSMUSG00000079015 | Serpina1c | serine (or cysteine) peptidase inhibitor, clade A, member 1C | -3.09 |
| ENSMUSG00000029563 | Foxp2 | forkhead box P2 | -3.09 |
| ENSMUSG00000024225 | Clps | colipase, pancreatic | -3.11 |
| ENSMUSG00000044022 | Pcdhb21 | protocadherin beta 21 | -3.11 |
| ENSMUSG00000051251 | Nhlh1 | nescient helix loop helix 1 | -3.11 |
| ENSMUSG00000001802 | Lrp3 | low density lipoprotein receptor-related protein 3 | -3.12 |
| ENSMUSG00000018800 | Abca5 | ATP-binding cassette, sub-family A (ABC1), member 5 | -3.12 |
| ENSMUSG00000021071 | Trim9 | tripartite motif-containing 9 | -3.12 |
| ENSMUSG00000063142 | Kcnma1 | potassium large conductance calcium-activated channel, subfamily M, alpha member 1 | -3.13 |
| ENSMUSG00000043391 | 2510009E07Rik | RIKEN cDNA 2510009E07 gene | -3.13 |
| ENSMUSG00000034353 | Ramp1 | receptor (calcitonin) activity modifying protein 1 | -3.13 |
| ENSMUSG00000018470 | Kcnab3 | potassium voltage-gated channel, shaker-related subfamily, beta member 3 | -3.13 |
| ENSMUSG00000103793 | Pcdhga6 | protocadherin gamma subfamily A, 6 | -3.13 |
| ENSMUSG00000059898 | Dsc3 | desmocollin 3 | -3.14 |
| ENSMUSG00000071550 | Cfap44 | cilia and flagella associated protein 44 | -3.14 |
| ENSMUSG00000046922 | Gpr6 | G protein-coupled receptor 6 | -3.14 |
| ENSMUSG00000022148 | Fyb | FYN binding protein | -3.16 |
| ENSMUSG00000052889 | Prkcb | protein kinase C, beta | -3.16 |
| ENSMUSG00000051043 | Gprc5c | G protein-coupled receptor, family C, group 5, member C | -3.17 |
| ENSMUSG00000061845 | Defa35 | defensin, alpha, 35 | -3.17 |
| ENSMUSG00000036062 | Phf24 | PHD finger protein 24 | -3.17 |
| ENSMUSG00000035357 | Pdzrn3 | PDZ domain containing RING finger 3 | -3.18 |
| ENSMUSG00000028600 | Podn | podocan | -3.18 |
| ENSMUSG00000063206 | Defa34 | defensin, alpha, 34 | -3.18 |
| ENSMUSG00000009246 | Trpm5 | transient receptor potential cation channel, subfamily M, member 5 | -3.19 |
| ENSMUSG00000021614 | Vcan | versican | -3.19 |
| ENSMUSG00000095328 | Defa-ps6 | defensin, alpha, pseudogene 6 | -3.19 |
| ENSMUSG00000008193 | Spib | Spi-B transcription factor (Spi-1/PU.1 related) | -3.19 |
| ENSMUSG00000072875 | Gpr27 | G protein-coupled receptor 27 | -3.19 |
| ENSMUSG00000094305 | Scgb2b20 | secretoglobin, family 2B, member 20 | -3.19 |
| ENSMUSG00000087211 | Lhx1os | LIM homeobox 1, opposite strand | -3.19 |
| ENSMUSG00000013523 | Bcas1 | breast carcinoma amplified sequence 1 | -3.20 |
| ENSMUSG00000042115 | Klhdc8a | kelch domain containing 8A | -3.20 |
| ENSMUSG00000032625 | Thsd7a | thrombospondin, type I, domain containing 7A | -3.20 |
| ENSMUSG00000019232 | Etnppl | ethanolamine phosphate phospholyase | -3.21 |
| ENSMUSG00000023216 | Epb42 | erythrocyte membrane protein band 4.2 | -3.22 |
| ENSMUSG00000025855 | Prkar1b | protein kinase, cAMP dependent regulatory, type I beta | -3.22 |
| ENSMUSG00000041420 | Meis3 | Meis homeobox 3 | -3.22 |
| ENSMUSG00000069814 | Ccdc92b | coiled-coil domain containing 92B | -3.22 |
| ENSMUSG00000079180 | Mptx2 | mucosal pentraxin 2 | -3.22 |
| ENSMUSG00000006546 | Cryba2 | crystallin, beta A2 | -3.22 |
| ENSMUSG00000037712 | Fermt2 | fermitin family member 2 | -3.23 |
| ENSMUSG00000032839 | Trpc1 | transient receptor potential cation channel, subfamily C, member 1 | -3.24 |
| ENSMUSG00000081303 | Gm16011 | predicted gene 16011 | -3.25 |
| ENSMUSG00000036913 | Trim67 | tripartite motif-containing 67 | -3.25 |
| ENSMUSG00000056069 | Otulinl | OTU deubiquitinase with linear linkage specificity like | -3.25 |
| ENSMUSG00000074274 | D930028M14Rik | RIKEN cDNA D930028M14 gene | -3.25 |
| ENSMUSG00000022441 | Efcab6 | EF-hand calcium binding domain 6 | -3.26 |
| ENSMUSG00000031760 | Mt3 | metallothionein 3 | -3.27 |
| ENSMUSG00000043088 | Il17re | interleukin 17 receptor E | -3.27 |
| ENSMUSG00000018339 | Gpx3 | glutathione peroxidase 3 | -3.27 |
| ENSMUSG00000033389 | Arhgap44 | Rho GTPase activating protein 44 | -3.28 |
| ENSMUSG00000045689 | Pcdhb4 | protocadherin beta 4 | -3.28 |
| ENSMUSG00000074440 | Defa3 | defensin, alpha, 3 | -3.29 |
| ENSMUSG00000029769 | Ccdc136 | coiled-coil domain containing 136 | -3.29 |
| ENSMUSG00000030411 | Nova2 | neuro-oncological ventral antigen 2 | -3.29 |
| ENSMUSG00000036480 | Prss56 | protease, serine 56 | -3.29 |
| ENSMUSG00000102625 | Gm37672 | predicted gene, 37672 | -3.29 |
| ENSMUSG00000036330 | Slc18a1 | solute carrier family 18 (vesicular monoamine), member 1 | -3.30 |
| ENSMUSG00000034675 | Dbn1 | drebrin 1 | -3.30 |
| ENSMUSG00000047730 | Fcgbp | Fc fragment of IgG binding protein | -3.30 |
| ENSMUSG00000004707 | Ly9 | lymphocyte antigen 9 | -3.30 |
| ENSMUSG00000038555 | Reep2 | receptor accessory protein 2 | -3.30 |
| ENSMUSG00000030074 | Gxylt2 | glucoside xylosyltransferase 2 | -3.30 |
| ENSMUSG00000032473 | Cldn18 | claudin 18 | -3.30 |
| ENSMUSG00000026360 | Rgs2 | regulator of G-protein signaling 2 | -3.31 |
| ENSMUSG00000069255 | Dusp22 | dual specificity phosphatase 22 | -3.31 |

| | | | |
|---|---|---|---|
| ENSMUSG00000023033 | Scn8a | sodium channel, voltage-gated, type VIII, alpha | -3.32 |
| ENSMUSG00000032936 | Camkv | CaM kinase-like vesicle-associated | -3.33 |
| ENSMUSG00000096295 | Defa2 | defensin, alpha, 2 | -3.33 |
| ENSMUSG00000044043 | Pcdhb14 | protocadherin beta 14 | -3.33 |
| ENSMUSG00000054169 | Ceacam10 | carcinoembryonic antigen-related cell adhesion molecule 10 | -3.34 |
| ENSMUSG00000043857 | Mgat5b | mannoside acetylglucosaminyltransferase 5, isoenzyme B | -3.34 |
| ENSMUSG00000086970 | Bcas1os1 | breast carcinoma amplified sequence 1, opposite strand 1 | -3.34 |
| ENSMUSG00000021539 | Lect2 | leukocyte cell-derived chemotaxin 2 | -3.35 |
| ENSMUSG00000027570 | Col9a3 | collagen, type IX, alpha 3 | -3.35 |
| ENSMUSG00000018862 | Otop3 | otopetrin 3 | -3.35 |
| ENSMUSG00000110394 | Gm18991 | predicted gene, 18991 | -3.35 |
| ENSMUSG00000024030 | Abcg1 | ATP binding cassette subfamily G member 1 | -3.37 |
| ENSMUSG00000038255 | Neurod2 | neurogenic differentiation 2 | -3.37 |
| ENSMUSG00000101930 | Gm5441 | predicted gene 5441 | -3.37 |
| ENSMUSG00000053141 | Ptprt | protein tyrosine phosphatase, receptor type, T | -3.38 |
| ENSMUSG00000025515 | Muc2 | mucin 2 | -3.39 |
| ENSMUSG00000030616 | Sytl2 | synaptotagmin-like 2 | -3.39 |
| ENSMUSG00000052942 | Glis3 | GLIS family zinc finger 3 | -3.39 |
| ENSMUSG00000030401 | Rtn2 | reticulon 2 (Z-band associated protein) | -3.39 |
| ENSMUSG00000035246 | Pcyt1b | phosphate cytidylyltransferase 1, choline, beta isoform | -3.40 |
| ENSMUSG00000005503 | Evx1 | even-skipped homeobox 1 | -3.40 |
| ENSMUSG00000020848 | Doc2b | double C2, beta | -3.40 |
| ENSMUSG00000047238 | Mageh1 | melanoma antigen, family H, 1 | -3.40 |
| ENSMUSG00000028356 | Ambp | alpha 1 microglobulin/bikunin | -3.40 |
| ENSMUSG00000015053 | Gata2 | GATA binding protein 2 | -3.40 |
| ENSMUSG00000030428 | Ttyh1 | tweety family member 1 | -3.41 |
| ENSMUSG00000052212 | Cd177 | CD177 antigen | -3.42 |
| ENSMUSG00000054453 | Sytl5 | synaptotagmin-like 5 | -3.42 |
| ENSMUSG00000029298 | Gbp9 | guanylate-binding protein 9 | -3.42 |
| ENSMUSG00000034336 | Ina | internexin neuronal intermediate filament protein, alpha | -3.42 |
| ENSMUSG00000024029 | Tff3 | trefoil factor 3, intestinal | -3.43 |
| ENSMUSG00000044017 | Adgrd1 | adhesion G protein-coupled receptor D1 | -3.43 |
| ENSMUSG00000100182 | 1810006J02Rik | RIKEN cDNA 1810006J02 gene | -3.43 |
| ENSMUSG00000091735 | Gpr62 | G protein-coupled receptor 62 | -3.44 |
| ENSMUSG00000019846 | Lama4 | laminin, alpha 4 | -3.45 |
| ENSMUSG00000031374 | Zfp92 | zinc finger protein 92 | -3.46 |
| ENSMUSG00000026443 | Lrrn2 | leucine rich repeat protein 2, neuronal | -3.46 |
| ENSMUSG00000029193 | Cckar | cholecystokinin A receptor | -3.46 |
| ENSMUSG00000051314 | Ffar2 | free fatty acid receptor 2 | -3.47 |
| ENSMUSG00000051111 | Sv2c | synaptic vesicle glycoprotein 2c | -3.47 |
| ENSMUSG00000104149 | Gm37138 | predicted gene, 37138 | -3.49 |
| ENSMUSG00000035967 | Ints6l | integrator complex subunit 6 like | -3.50 |
| ENSMUSG00000040170 | Fmo2 | flavin containing monooxygenase 2 | -3.50 |
| ENSMUSG00000050473 | Slc35d3 | solute carrier family 35, member D3 | -3.50 |
| ENSMUSG00000055407 | Map6 | microtubule-associated protein 6 | -3.51 |
| ENSMUSG00000057335 | Cep170 | centrosomal protein 170 | -3.52 |
| ENSMUSG00000048347 | Pcdhb18 | protocadherin beta 18 | -3.52 |
| ENSMUSG00000041078 | Grid1 | glutamate receptor, ionotropic, delta 1 | -3.52 |
| ENSMUSG00000034656 | Cacna1a | calcium channel, voltage-dependent, P/Q type, alpha 1A subunit | -3.53 |
| ENSMUSG00000020601 | Trib2 | tribbles pseudokinase 2 | -3.53 |
| ENSMUSG00000042766 | Trim46 | tripartite motif-containing 46 | -3.53 |
| ENSMUSG00000040276 | Pacsin1 | protein kinase C and casein kinase substrate in neurons 1 | -3.54 |
| ENSMUSG00000044988 | Ucn3 | urocortin 3 | -3.54 |
| ENSMUSG00000097587 | 4930578M01Rik | RIKEN cDNA 4930578M01 gene | -3.55 |
| ENSMUSG00000042179 | Pnliprp1 | pancreatic lipase related protein 1 | -3.56 |
| ENSMUSG00000048078 | Tenm4 | teneurin transmembrane protein 4 | -3.58 |
| ENSMUSG00000044156 | Hepacam2 | HEPACAM family member 2 | -3.58 |
| ENSMUSG00000053545 | 6430503K07Rik | RIKEN cDNA 6430503K07 gene | -3.58 |
| ENSMUSG00000038349 | Plcl1 | phospholipase C-like 1 | -3.58 |
| ENSMUSG00000036395 | Glb1l2 | galactosidase, beta 1-like 2 | -3.58 |
| ENSMUSG00000041180 | Hectd2 | HECT domain E3 ubiquitin protein ligase 2 | -3.58 |
| ENSMUSG00000024548 | Setbp1 | SET binding protein 1 | -3.59 |
| ENSMUSG00000046204 | Pnma2 | paraneoplastic antigen MA2 | -3.59 |
| ENSMUSG00000041633 | Kctd12b | potassium channel tetramerisation domain containing 12b | -3.59 |
| ENSMUSG00000047250 | Ptgs1 | prostaglandin-endoperoxide synthase 1 | -3.60 |
| ENSMUSG00000052631 | Sh2d6 | SH2 domain containing 6 | -3.61 |
| ENSMUSG00000040009 | Gnaz | guanine nucleotide binding protein, alpha z subunit | -3.61 |
| ENSMUSG00000021904 | Sema3g | sema domain, immunoglobulin domain (Ig), short basic domain, secreted, (semaphorin) 3G | -3.61 |
| ENSMUSG00000035459 | Stab2 | stabilin 2 | -3.61 |
| ENSMUSG00000020703 | 5530401A14Rik | RIKEN cDNA 5530401A14 gene | -3.62 |
| ENSMUSG00000085247 | 4930545L23Rik | RIKEN cDNA 4930545L23 gene | -3.63 |
| ENSMUSG00000052087 | Rgs14 | regulator of G-protein signaling 14 | -3.64 |
| ENSMUSG00000025810 | Nrp1 | neuropilin 1 | -3.64 |

| | | | |
|---|---|---|---|
| ENSMUSG00000039037 | St6galnac5 | ST6 (alpha-N-acetyl-neuraminyl-2,3-beta-galactosyl-1,3)-N-acetylgalactosaminide alpha-2,6-sialyltra | -3.64 |
| ENSMUSG00000003283 | Hck | hemopoietic cell kinase | -3.64 |
| ENSMUSG00000066705 | Fxyd6 | FXYD domain-containing ion transport regulator 6 | -3.65 |
| ENSMUSG00000056812 | St8sia3 | ST8 alpha-N-acetyl-neuraminide alpha-2,8-sialyltransferase 3 | -3.66 |
| ENSMUSG00000064125 | Prr36 | proline rich 36 | -3.67 |
| ENSMUSG00000086126 | Evx1os | even skipped homeotic gene 1, opposite strand | -3.67 |
| ENSMUSG00000105271 | Gm42875 | predicted gene 42875 | -3.67 |
| ENSMUSG00000021256 | Vash1 | vasohibin 1 | -3.68 |
| ENSMUSG00000021763 | BC067074 | cDNA sequence BC067074 | -3.68 |
| ENSMUSG00000086029 | Pax6os1 | paired box 6 opposite strand 1 | -3.68 |
| ENSMUSG00000032826 | Ank2 | ankyrin 2, brain | -3.69 |
| ENSMUSG00000037239 | Spred3 | sprouty-related, EVH1 domain containing 3 | -3.70 |
| ENSMUSG00000062861 | Zfp28 | zinc finger protein 28 | -3.70 |
| ENSMUSG00000055430 | Nap1l5 | nucleosome assembly protein 1-like 5 | -3.71 |
| ENSMUSG00000029005 | Draxin | dorsal inhibitory axon guidance protein | -3.71 |
| ENSMUSG00000072849 | Serpina1e | serine (or cysteine) peptidase inhibitor, clade A, member 1E | -3.71 |
| ENSMUSG00000032852 | Rspo4 | R-spondin 4 | -3.72 |
| ENSMUSG00000030518 | Fam189a1 | family with sequence similarity 189, member A1 | -3.72 |
| ENSMUSG00000020340 | Cyfip2 | cytoplasmic FMR1 interacting protein 2 | -3.73 |
| ENSMUSG00000024013 | Fgd2 | FYVE, RhoGEF and PH domain containing 2 | -3.74 |
| ENSMUSG00000038224 | Serpinf2 | serine (or cysteine) peptidase inhibitor, clade F, member 2 | -3.74 |
| ENSMUSG00000040852 | Plekhh2 | pleckstrin homology domain containing, family H (with MyTH4 domain) member 2 | -3.74 |
| ENSMUSG00000023473 | Celsr3 | cadherin, EGF LAG seven-pass G-type receptor 3 | -3.75 |
| ENSMUSG00000032064 | Dixdc1 | DIX domain containing 1 | -3.75 |
| ENSMUSG00000002578 | Ikzf4 | IKAROS family zinc finger 4 | -3.75 |
| ENSMUSG00000040563 | Plppr2 | phospholipid phosphatase related 2 | -3.75 |
| ENSMUSG00000034799 | Unc13a | unc-13 homolog A | -3.76 |
| ENSMUSG00000095813 | Defa-ps1 | defensin, alpha, pseudogene 1 | -3.76 |
| ENSMUSG00000022054 | Nefm | neurofilament, medium polypeptide | -3.77 |
| ENSMUSG00000057914 | Cacnb2 | calcium channel, voltage-dependent, beta 2 subunit | -3.77 |
| ENSMUSG00000038486 | Sv2a | synaptic vesicle glycoprotein 2 a | -3.78 |
| ENSMUSG00000078776 | 9530053A07Rik | RIKEN cDNA 9530053A07 gene | -3.79 |
| ENSMUSG00000004347 | Pde1c | phosphodiesterase 1C | -3.79 |
| ENSMUSG00000079481 | Nhsl2 | NHS-like 2 | -3.79 |
| ENSMUSG00000055567 | Unc80 | unc-80, NALCN activator | -3.79 |
| ENSMUSG00000025738 | Fbxl16 | F-box and leucine-rich repeat protein 16 | -3.79 |
| ENSMUSG00000028360 | Slc44a5 | solute carrier family 44, member 5 | -3.79 |
| ENSMUSG00000015222 | Map2 | microtubule-associated protein 2 | -3.80 |
| ENSMUSG00000030263 | Lrmp | lymphoid-restricted membrane protein | -3.80 |
| ENSMUSG00000097767 | Miat | myocardial infarction associated transcript (non-protein coding) | -3.80 |
| ENSMUSG00000026072 | Il1r1 | interleukin 1 receptor, type I | -3.80 |
| ENSMUSG00000027254 | Map1a | microtubule-associated protein 1 A | -3.81 |
| ENSMUSG00000049605 | Olfr418 | olfactory receptor 418 | -3.81 |
| ENSMUSG00000066176 | Gm12511 | predicted gene 12511 | -3.81 |
| ENSMUSG00000023913 | Pla2g7 | phospholipase A2, group VII (platelet-activating factor acetylhydrolase, plasma) | -3.81 |
| ENSMUSG00000102581 | Gm37443 | predicted gene, 37443 | -3.82 |
| ENSMUSG00000034771 | Tle2 | transducin-like enhancer of split 2 | -3.83 |
| ENSMUSG00000015599 | Ttbk1 | tau tubulin kinase 1 | -3.83 |
| ENSMUSG00000030302 | Atp2b2 | ATPase, Ca++ transporting, plasma membrane 2 | -3.83 |
| ENSMUSG00000068154 | Insm1 | insulinoma-associated 1 | -3.85 |
| ENSMUSG00000094662 | Defa36 | defensin, alpha, 36 | -3.85 |
| ENSMUSG00000021587 | Pcsk1 | proprotein convertase subtilisin/kexin type 1 | -3.86 |
| ENSMUSG00000063919 | Srrm4 | serine/arginine repetitive matrix 4 | -3.86 |
| ENSMUSG00000112234 | Gm48427 | predicted gene, 48427 | -3.86 |
| ENSMUSG00000052951 | C130021I20Rik | Riken cDNA C130021I20 gene | -3.86 |
| ENSMUSG00000047507 | Baiap3 | BAI1-associated protein 3 | -3.87 |
| ENSMUSG00000027674 | Pex5l | peroxisomal biogenesis factor 5-like | -3.87 |
| ENSMUSG00000040146 | Rgl3 | ral guanine nucleotide dissociation stimulator-like 3 | -3.88 |
| ENSMUSG00000020090 | Npffr1 | neuropeptide FF receptor 1 | -3.88 |
| ENSMUSG00000022438 | Parvb | parvin, beta | -3.89 |
| ENSMUSG00000111443 | Gm46123 | predicted gene, 46123 | -3.89 |
| ENSMUSG00000034958 | Atcay | ataxia, cerebellar, Cayman type | -3.89 |
| ENSMUSG00000032035 | Ets1 | E26 avian leukemia oncogene 1, 5' domain | -3.90 |
| ENSMUSG00000075256 | Cerkl | ceramide kinase-like | -3.90 |
| ENSMUSG00000037106 | Fer1l6 | fer-1-like 6 (C. elegans) | -3.91 |
| ENSMUSG00000056004 | 9330182L06Rik | RIKEN cDNA 9330182L06 gene | -3.91 |
| ENSMUSG00000009628 | Tex15 | testis expressed gene 15 | -3.92 |
| ENSMUSG00000021536 | Adcy2 | adenylate cyclase 2 | -3.92 |
| ENSMUSG00000058420 | Syt17 | synaptotagmin XVII | -3.92 |
| ENSMUSG00000040035 | Disp2 | dispatched RND tramsporter family member 2 | -3.94 |
| ENSMUSG00000038665 | Dgki | diacylglycerol kinase, iota | -3.94 |
| ENSMUSG00000050994 | Adgb | androglobin | -3.94 |
| ENSMUSG00000027048 | Abcb11 | ATP-binding cassette, sub-family B (MDR/TAP), member 11 | -3.94 |

| | | | |
|---|---|---|---|
| ENSMUSG00000112596 | Gm48804 | predicted gene, 48804 | -3.95 |
| ENSMUSG00000028469 | Npr2 | natriuretic peptide receptor 2 | -3.95 |
| ENSMUSG00000070498 | Tmem132b | transmembrane protein 132B | -3.95 |
| ENSMUSG00000049122 | Frmd3 | FERM domain containing 3 | -3.96 |
| ENSMUSG00000102937 | Gm38116 | predicted gene, 38116 | -3.96 |
| ENSMUSG00000046460 | Sh2d7 | SH2 domain containing 7 | -3.96 |
| ENSMUSG00000000632 | Sez6 | seizure related gene 6 | -3.97 |
| ENSMUSG00000029706 | Pax4 | paired box 4 | -3.98 |
| ENSMUSG00000040136 | Abcc8 | ATP-binding cassette, sub-family C (CFTR/MRP), member 8 | -3.99 |
| ENSMUSG00000020811 | Wscd1 | WSC domain containing 1 | -4.00 |
| ENSMUSG00000096146 | Kcnj11 | potassium inwardly rectifying channel, subfamily J, member 11 | -4.00 |
| ENSMUSG00000027876 | Reg4 | regenerating islet-derived family, member 4 | -4.01 |
| ENSMUSG00000039809 | Gabbr2 | gamma-aminobutyric acid (GABA) B receptor, 2 | -4.02 |
| ENSMUSG00000028255 | Clca1 | chloride channel accessory 1 | -4.02 |
| ENSMUSG00000048960 | Prex2 | phosphatidylinositol-3,4,5-trisphosphate-dependent Rac exchange factor 2 | -4.02 |
| ENSMUSG00000023906 | Cldn6 | claudin 6 | -4.03 |
| ENSMUSG00000028339 | Col15a1 | collagen, type XV, alpha 1 | -4.03 |
| ENSMUSG00000020333 | Acsl6 | acyl-CoA synthetase long-chain family member 6 | -4.04 |
| ENSMUSG00000019124 | Scrn1 | secernin 1 | -4.04 |
| ENSMUSG00000071226 | Cecr2 | CECR2, histone acetyl-lysine reader | -4.05 |
| ENSMUSG00000010505 | Myt1 | myelin transcription factor 1 | -4.05 |
| ENSMUSG00000037035 | Inhbb | inhibin beta-B | -4.06 |
| ENSMUSG00000038248 | Sobp | sine oculis binding protein | -4.06 |
| ENSMUSG00000036766 | Dner | delta/notch-like EGF repeat containing | -4.06 |
| ENSMUSG00000043004 | Gng2 | guanine nucleotide binding protein (G protein), gamma 2 | -4.07 |
| ENSMUSG00000071719 | Tmem28 | transmembrane protein 28 | -4.07 |
| ENSMUSG00000074434 | Defa28 | defensin, alpha, 28 | -4.07 |
| ENSMUSG00000038916 | Soga3 | SOGA family member 3 | -4.07 |
| ENSMUSG00000044177 | Wfikkn2 | WAP, follistatin/kazal, immunoglobulin, kunitz and netrin domain containing 2 | -4.08 |
| ENSMUSG00000023236 | Scg5 | secretogranin V | -4.09 |
| ENSMUSG00000038580 | Sct | secretin | -4.09 |
| ENSMUSG00000062760 | Shisal1 | shisa like 1 | -4.09 |
| ENSMUSG00000044288 | Cnr1 | cannabinoid receptor 1 (brain) | -4.09 |
| ENSMUSG00000003411 | Rab3b | RAB3B, member RAS oncogene family | -4.09 |
| ENSMUSG00000039954 | Stk32a | serine/threonine kinase 32A | -4.09 |
| ENSMUSG00000047798 | Cd300lf | CD300 molecule like family member F | -4.09 |
| ENSMUSG00000096847 | Tmem151b | transmembrane protein 151B | -4.10 |
| ENSMUSG00000025582 | Nptx1 | neuronal pentraxin 1 | -4.10 |
| ENSMUSG00000022055 | Nefl | neurofilament, light polypeptide | -4.10 |
| ENSMUSG00000031748 | Gnao1 | guanine nucleotide binding protein, alpha O | -4.11 |
| ENSMUSG00000049350 | Zg16 | zymogen granule protein 16 | -4.11 |
| ENSMUSG00000079116 | Gm15293 | predicted gene 15293 | -4.11 |
| ENSMUSG00000099907 | Gm10421 | predicted gene 10421 | -4.11 |
| ENSMUSG00000024897 | Apba1 | amyloid beta (A4) precursor protein binding, family A, member 1 | -4.12 |
| ENSMUSG00000027834 | Serpini1 | serine (or cysteine) peptidase inhibitor, clade I, member 1 | -4.13 |
| ENSMUSG00000040653 | Ppp1r14c | protein phosphatase 1, regulatory inhibitor subunit 14C | -4.13 |
| ENSMUSG00000020732 | Rab37 | RAB37, member RAS oncogene family | -4.13 |
| ENSMUSG00000025020 | Slit1 | slit guidance ligand 1 | -4.14 |
| ENSMUSG00000034701 | Neurod1 | neurogenic differentiation 1 | -4.14 |
| ENSMUSG00000038692 | Hoxb4 | homeobox B4 | -4.14 |
| ENSMUSG00000027797 | Dclk1 | doublecortin-like kinase 1 | -4.14 |
| ENSMUSG00000106491 | Gm42446 | predicted gene 42446 | -4.14 |
| ENSMUSG00000029368 | Alb | albumin | -4.14 |
| ENSMUSG00000026442 | Nfasc | neurofascin | -4.15 |
| ENSMUSG00000114624 | Gm48267 | predicted gene, 48267 | -4.15 |
| ENSMUSG00000014602 | Kif1a | kinesin family member 1A | -4.16 |
| ENSMUSG00000105526 | Gm43490 | predicted gene 43490 | -4.16 |
| ENSMUSG00000086245 | Gm16170 | predicted gene 16170 | -4.16 |
| ENSMUSG00000036782 | Klhl13 | kelch-like 13 | -4.16 |
| ENSMUSG00000027434 | Nkx2-2 | NK2 homeobox 2 | -4.18 |
| ENSMUSG00000038765 | Lmx1b | LIM homeobox transcription factor 1 beta | -4.18 |
| ENSMUSG00000022456 | 37865 | septin 3 | -4.18 |
| ENSMUSG00000029245 | Epha5 | Eph receptor A5 | -4.18 |
| ENSMUSG00000032377 | Plscr4 | phospholipid scramblase 4 | -4.18 |
| ENSMUSG00000032394 | Igdcc3 | immunoglobulin superfamily, DCC subclass, member 3 | -4.18 |
| ENSMUSG00000028528 | Dnajc6 | DnaJ heat shock protein family (Hsp40) member C6 | -4.18 |
| ENSMUSG00000010021 | Kif19a | kinesin family member 19A | -4.20 |
| ENSMUSG00000058975 | Kcnc1 | potassium voltage gated channel, Shaw-related subfamily, member 1 | -4.20 |
| ENSMUSG00000070802 | Pnmal2 | PNMA-like 2 | -4.21 |
| ENSMUSG00000022425 | Enpp2 | ectonucleotide pyrophosphatase/phosphodiesterase 2 | -4.21 |
| ENSMUSG00000033615 | Cplx1 | complexin 1 | -4.21 |
| ENSMUSG00000086999 | Bcas1os2 | breast carcinoma amplified sequence 1, opposite strand 2 | -4.21 |
| ENSMUSG00000046613 | Vwa5b2 | von Willebrand factor A domain containing 5B2 | -4.22 |

| | | | |
|---|---|---|---|
| ENSMUSG00000004961 | Syt5 | synaptotagmin V | -4.22 |
| ENSMUSG00000040046 | Tph1 | tryptophan hydroxylase 1 | -4.24 |
| ENSMUSG00000026463 | Atp2b4 | ATPase, Ca++ transporting, plasma membrane 4 | -4.25 |
| ENSMUSG00000106139 | Gm30648 | predicted gene, 30648 | -4.25 |
| ENSMUSG00000019796 | Lrp11 | low density lipoprotein receptor-related protein 11 | -4.26 |
| ENSMUSG00000050640 | Tmem150c | transmembrane protein 150C | -4.26 |
| ENSMUSG00000062044 | Lmtk3 | lemur tyrosine kinase 3 | -4.26 |
| ENSMUSG00000000120 | Ngfr | nerve growth factor receptor (TNFR superfamily, member 16) | -4.26 |
| ENSMUSG00000064177 | Ghrl | ghrelin | -4.27 |
| ENSMUSG00000031551 | Ido1 | indoleamine 2,3-dioxygenase 1 | -4.27 |
| ENSMUSG00000071113 | Mboat4 | membrane bound O-acyltransferase domain containing 4 | -4.27 |
| ENSMUSG00000104301 | Wdr49 | WD repeat domain 49 | -4.27 |
| ENSMUSG00000015401 | Cltrn | collectrin, amino acid transport regulator | -4.28 |
| ENSMUSG00000043388 | Tmem130 | transmembrane protein 130 | -4.28 |
| ENSMUSG00000097311 | Gm26871 | predicted gene, 26871 | -4.28 |
| ENSMUSG00000026285 | Pdcd1 | programmed cell death 1 | -4.29 |
| ENSMUSG00000030020 | Prickle2 | prickle planar cell polarity protein 2 | -4.29 |
| ENSMUSG00000058297 | Spock2 | sparc/osteonectin, cwcv and kazal-like domains proteoglycan 2 | -4.30 |
| ENSMUSG00000061603 | Akap6 | A kinase (PRKA) anchor protein 6 | -4.30 |
| ENSMUSG00000026872 | Zeb2 | zinc finger E-box binding homeobox 2 | -4.32 |
| ENSMUSG00000079037 | Prnp | prion protein | -4.33 |
| ENSMUSG00000030683 | Sez6l2 | seizure related 6 homolog like 2 | -4.33 |
| ENSMUSG00000051331 | Cacna1c | calcium channel, voltage-dependent, L type, alpha 1C subunit | -4.33 |
| ENSMUSG00000028926 | Cdk14 | cyclin-dependent kinase 14 | -4.34 |
| ENSMUSG00000043496 | Tril | TLR4 interactor with leucine-rich repeats | -4.34 |
| ENSMUSG00000112556 | Gm47339 | predicted gene, 47339 | -4.34 |
| ENSMUSG00000032087 | Dscaml1 | DS cell adhesion molecule like 1 | -4.34 |
| ENSMUSG00000001901 | Kcnh6 | potassium voltage-gated channel, subfamily H (eag-related), member 6 | -4.35 |
| ENSMUSG00000020177 | 9530003J23Rik | RIKEN cDNA 9530003J23 gene | -4.35 |
| ENSMUSG00000110557 | Gm5159 | predicted gene 5159 | -4.35 |
| ENSMUSG00000024990 | Rbp4 | retinol binding protein 4, plasma | -4.36 |
| ENSMUSG00000024366 | Gfra3 | glial cell line derived neurotrophic factor family receptor alpha 3 | -4.37 |
| ENSMUSG00000028137 | Celf3 | CUGBP, Elav-like family member 3 | -4.37 |
| ENSMUSG00000021647 | Cartpt | CART prepropeptide | -4.37 |
| ENSMUSG00000038463 | Olfml2b | olfactomedin-like 2B | -4.38 |
| ENSMUSG00000060671 | Atp8b2 | ATPase, class I, type 8B, member 2 | -4.39 |
| ENSMUSG00000056752 | Dnah9 | dynein, axonemal, heavy chain 9 | -4.39 |
| ENSMUSG00000032356 | Rasgrf1 | RAS protein-specific guanine nucleotide-releasing factor 1 | -4.39 |
| ENSMUSG00000071489 | Ptgdr | prostaglandin D receptor | -4.40 |
| ENSMUSG00000092083 | Kcnb2 | potassium voltage gated channel, Shab-related subfamily, member 2 | -4.41 |
| ENSMUSG00000030359 | Pzp | PZP, alpha-2-macroglobulin like | -4.41 |
| ENSMUSG00000037032 | Apbb1 | amyloid beta (A4) precursor protein-binding, family B, member 1 | -4.42 |
| ENSMUSG00000073530 | Pappa2 | pappalysin 2 | -4.42 |
| ENSMUSG00000104283 | Gm37459 | predicted gene, 37459 | -4.42 |
| ENSMUSG00000018698 | Lhx1 | LIM homeobox protein 1 | -4.43 |
| ENSMUSG00000019900 | Rfx6 | regulatory factor X, 6 | -4.44 |
| ENSMUSG00000033740 | St18 | suppression of tumorigenicity 18 | -4.44 |
| ENSMUSG00000044365 | Cxxc4 | CXXC finger 4 | -4.44 |
| ENSMUSG00000016200 | Syt14 | synaptotagmin XIV | -4.44 |
| ENSMUSG00000034145 | Tmem63c | transmembrane protein 63c | -4.44 |
| ENSMUSG00000033900 | Map9 | microtubule-associated protein 9 | -4.45 |
| ENSMUSG00000066456 | Hmgn3 | high mobility group nucleosomal binding domain 3 | -4.46 |
| ENSMUSG00000026833 | Olfm1 | olfactomedin 1 | -4.46 |
| ENSMUSG00000032769 | Trpa1 | transient receptor potential cation channel, subfamily A, member 1 | -4.47 |
| ENSMUSG00000103529 | A730089K16Rik | RIKEN cDNA A730089K16 gene | -4.48 |
| ENSMUSG00000045215 | Asxl3 | additional sex combs like 3, transcriptional regulator | -4.50 |
| ENSMUSG00000027009 | Itga4 | integrin alpha 4 | -4.51 |
| ENSMUSG00000090291 | Lrrc10b | leucine rich repeat containing 10B | -4.51 |
| ENSMUSG00000010086 | Rnf112 | ring finger protein 112 | -4.51 |
| ENSMUSG00000022995 | Enah | ENAH actin regulator | -4.52 |
| ENSMUSG00000031561 | Tenm3 | teneurin transmembrane protein 3 | -4.52 |
| ENSMUSG00000046743 | Fat4 | FAT atypical cadherin 4 | -4.53 |
| ENSMUSG00000047878 | A4galt | alpha 1,4-galactosyltransferase | -4.53 |
| ENSMUSG00000042750 | Bex2 | brain expressed X-linked 2 | -4.54 |
| ENSMUSG00000062444 | Ap3b2 | adaptor-related protein complex 3, beta 2 subunit | -4.54 |
| ENSMUSG00000027360 | Hdc | histidine decarboxylase | -4.54 |
| ENSMUSG00000079056 | Kcnip3 | Kv channel interacting protein 3, calsenilin | -4.54 |
| ENSMUSG00000114501 | Gm48582 | predicted gene, 48582 | -4.54 |
| ENSMUSG00000056553 | Ptprn2 | protein tyrosine phosphatase, receptor type, N polypeptide 2 | -4.55 |
| ENSMUSG00000037306 | Man1c1 | mannosidase, alpha, class 1C, member 1 | -4.55 |
| ENSMUSG00000006586 | Runx1t1 | runt-related transcription factor 1; translocated to, 1 (cyclin D-related) | -4.56 |
| ENSMUSG00000033676 | Gabrb3 | gamma-aminobutyric acid (GABA) A receptor, subunit beta 3 | -4.56 |
| ENSMUSG00000054667 | Irs4 | insulin receptor substrate 4 | -4.56 |

| | | | |
|---|---|---|---|
| ENSMUSG00000042078 | Svop | SV2 related protein | -4.57 |
| ENSMUSG00000101814 | Gm17807 | predicted gene, 17807 | -4.59 |
| ENSMUSG00000039683 | Sdk1 | sidekick cell adhesion molecule 1 | -4.60 |
| ENSMUSG00000051980 | Casr | calcium-sensing receptor | -4.60 |
| ENSMUSG00000028546 | Elavl4 | ELAV like RNA binding protein 4 | -4.61 |
| ENSMUSG00000055197 | Fev | FEV (ETS oncogene family) | -4.61 |
| ENSMUSG00000009292 | Trpm2 | transient receptor potential cation channel, subfamily M, member 2 | -4.61 |
| ENSMUSG00000037428 | Vgf | VGF nerve growth factor inducible | -4.62 |
| ENSMUSG00000031144 | Syp | synaptophysin | -4.63 |
| ENSMUSG00000029420 | Rimbp2 | RIMS binding protein 2 | -4.64 |
| ENSMUSG00000020218 | Wif1 | Wnt inhibitory factor 1 | -4.66 |
| ENSMUSG00000032338 | Hcn4 | hyperpolarization-activated, cyclic nucleotide-gated K+ 4 | -4.67 |
| ENSMUSG00000022762 | Ncam2 | neural cell adhesion molecule 2 | -4.67 |
| ENSMUSG00000032297 | Celf6 | CUGBP, Elav-like family member 6 | -4.67 |
| ENSMUSG00000027858 | Tspan2 | tetraspanin 2 | -4.67 |
| ENSMUSG00000114230 | Gm48239 | predicted gene, 48239 | -4.68 |
| ENSMUSG00000048218 | Amigo2 | adhesion molecule with Ig like domain 2 | -4.69 |
| ENSMUSG00000095079 | Igha | immunoglobulin heavy constant alpha | -4.69 |
| ENSMUSG00000020312 | Shc2 | SHC (Src homology 2 domain containing) transforming protein 2 | -4.71 |
| ENSMUSG00000001985 | Grik3 | glutamate receptor, ionotropic, kainate 3 | -4.71 |
| ENSMUSG00000044359 | P2ry4 | pyrimidinergic receptor P2Y, G-protein coupled, 4 | -4.72 |
| ENSMUSG00000038112 | AW551984 | expressed sequence AW551984 | -4.72 |
| ENSMUSG00000029765 | Plxna4 | plexin A4 | -4.72 |
| ENSMUSG00000035095 | Fam167a | family with sequence similarity 167, member A | -4.74 |
| ENSMUSG00000031837 | Necab2 | N-terminal EF-hand calcium binding protein 2 | -4.76 |
| ENSMUSG00000090071 | Cdk5r2 | cyclin-dependent kinase 5, regulatory subunit 2 (p39) | -4.77 |
| ENSMUSG00000026527 | Rgs7 | regulator of G protein signaling 7 | -4.77 |
| ENSMUSG00000046999 | 1110032F04Rik | RIKEN cDNA 1110032F04 gene | -4.77 |
| ENSMUSG00000026204 | Ptprn | protein tyrosine phosphatase, receptor type, N | -4.78 |
| ENSMUSG00000053049 | Gm15413 | predicted gene 15413 | -4.78 |
| ENSMUSG00000038530 | Rgs4 | regulator of G-protein signaling 4 | -4.79 |
| ENSMUSG00000061762 | Tac1 | tachykinin 1 | -4.79 |
| ENSMUSG00000058248 | Kcnh1 | potassium voltage-gated channel, subfamily H (eag-related), member 1 | -4.79 |
| ENSMUSG00000029875 | Ccdc184 | coiled-coil domain containing 184 | -4.79 |
| ENSMUSG00000023224 | Serping1 | serine (or cysteine) peptidase inhibitor, clade G, member 1 | -4.80 |
| ENSMUSG00000020886 | Dlg4 | discs large MAGUK scaffold protein 4 | -4.81 |
| ENSMUSG00000024084 | Qpct | glutaminyl-peptide cyclotransferase (glutaminyl cyclase) | -4.81 |
| ENSMUSG00000040972 | Igsf21 | immunoglobulin superfamily, member 21 | -4.81 |
| ENSMUSG00000062380 | Tubb3 | tubulin, beta 3 class III | -4.82 |
| ENSMUSG00000083282 | Ctsf | cathepsin F | -4.82 |
| ENSMUSG00000049154 | Fam183b | family with sequence similarity 183, member B | -4.83 |
| ENSMUSG00000021700 | Rab3c | RAB3C, member RAS oncogene family | -4.84 |
| ENSMUSG00000022829 | Stxbp5l | syntaxin binding protein 5-like | -4.84 |
| ENSMUSG00000069072 | Slc7a14 | solute carrier family 7 (cationic amino acid transporter, y+ system), member 14 | -4.86 |
| ENSMUSG00000004113 | Cacna1b | calcium channel, voltage-dependent, N type, alpha 1B subunit | -4.87 |
| ENSMUSG00000031284 | Pak3 | p21 (RAC1) activated kinase 3 | -4.87 |
| ENSMUSG00000094626 | Tmem121b | transmembrane protein 121B | -4.88 |
| ENSMUSG00000040118 | Cacna2d1 | calcium channel, voltage-dependent, alpha2/delta subunit 1 | -4.89 |
| ENSMUSG00000029121 | Crmp1 | collapsin response mediator protein 1 | -4.89 |
| ENSMUSG00000029223 | Uchl1 | ubiquitin carboxy-terminal hydrolase L1 | -4.90 |
| ENSMUSG00000027220 | Syt13 | synaptotagmin XIII | -4.91 |
| ENSMUSG00000028364 | Tnc | tenascin C | -4.91 |
| ENSMUSG00000041681 | Iapp | islet amyloid polypeptide | -4.91 |
| ENSMUSG00000064080 | Fbln2 | fibulin 2 | -4.93 |
| ENSMUSG00000045657 | Pcdhb10 | protocadherin beta 10 | -4.93 |
| ENSMUSG00000021303 | Gng4 | guanine nucleotide binding protein (G protein), gamma 4 | -4.94 |
| ENSMUSG00000050556 | Kcnb1 | potassium voltage gated channel, Shab-related subfamily, member 1 | -4.95 |
| ENSMUSG00000021337 | Scgn | secretagogin, EF-hand calcium binding protein | -4.97 |
| ENSMUSG00000026764 | Kif5c | kinesin family member 5C | -4.97 |
| ENSMUSG00000037852 | Cpe | carboxypeptidase E | -4.98 |
| ENSMUSG00000043518 | Rai2 | retinoic acid induced 2 | -4.98 |
| ENSMUSG00000016346 | Kcnq2 | potassium voltage-gated channel, subfamily Q, member 2 | -4.98 |
| ENSMUSG00000027350 | Chgb | chromogranin B | -4.99 |
| ENSMUSG00000091345 | Col6a5 | collagen, type VI, alpha 5 | -5.00 |
| ENSMUSG00000026686 | Lmx1a | LIM homeobox transcription factor 1 alpha | -5.01 |
| ENSMUSG00000026432 | Avpr1b | arginine vasopressin receptor 1B | -5.01 |
| ENSMUSG00000028226 | Mmp16 | matrix metallopeptidase 16 | -5.02 |
| ENSMUSG00000021194 | Chga | chromogranin A | -5.03 |
| ENSMUSG00000061601 | Pclo | piccolo (presynaptic cytomatrix protein) | -5.04 |
| ENSMUSG00000035431 | Sstr1 | somatostatin receptor 1 | -5.04 |
| ENSMUSG00000017311 | Pyy | peptide YY | -5.05 |
| ENSMUSG00000050751 | Pgbd5 | piggyBac transposable element derived 5 | -5.06 |
| ENSMUSG00000041141 | Pnmal1 | PNMA-like 1 | -5.07 |

| | | | |
|---|---|---|---|
| ENSMUSG00000039323 | Igfbp2 | insulin-like growth factor binding protein 2 | -5.09 |
| ENSMUSG00000024553 | Galr1 | galanin receptor 1 | -5.11 |
| ENSMUSG00000038456 | Dennd2a | DENN/MADD domain containing 2A | -5.12 |
| ENSMUSG00000115190 | Gm49542 | predicted gene, 49542 | -5.12 |
| ENSMUSG00000057716 | Tmem178b | transmembrane protein 178B | -5.13 |
| ENSMUSG00000024112 | Cacna1h | calcium channel, voltage-dependent, T type, alpha 1H subunit | -5.14 |
| ENSMUSG00000037624 | Kcnk2 | potassium channel, subfamily K, member 2 | -5.14 |
| ENSMUSG00000048216 | Gpr85 | G protein-coupled receptor 85 | -5.14 |
| ENSMUSG00000048215 | A630023P12Rik | RIKEN cDNA A630023P12 gene | -5.15 |
| ENSMUSG00000032181 | Scg3 | secretogranin III | -5.16 |
| ENSMUSG00000041798 | Gck | glucokinase | -5.20 |
| ENSMUSG00000102498 | Gm19445 | predicted gene, 19445 | -5.20 |
| ENSMUSG00000020734 | Grin2c | glutamate receptor, ionotropic, NMDA2C (epsilon 3) | -5.21 |
| ENSMUSG00000019876 | Pkib | protein kinase inhibitor beta, cAMP dependent, testis specific | -5.22 |
| ENSMUSG00000023387 | Kcnk16 | potassium channel, subfamily K, member 16 | -5.25 |
| ENSMUSG00000022449 | Adamts20 | a disintegrin-like and metallopeptidase (reprolysin type) with thrombospondin type 1 motif, 20 | -5.26 |
| ENSMUSG00000020704 | Asic2 | acid-sensing (proton-gated) ion channel 2 | -5.26 |
| ENSMUSG00000039601 | Rcan2 | regulator of calcineurin 2 | -5.27 |
| ENSMUSG00000027273 | Snap25 | synaptosomal-associated protein 25 | -5.28 |
| ENSMUSG00000004366 | Sst | somatostatin | -5.28 |
| ENSMUSG00000052572 | Dlg2 | discs large MAGUK scaffold protein 2 | -5.29 |
| ENSMUSG00000021559 | Dapk1 | death associated protein kinase 1 | -5.31 |
| ENSMUSG00000056492 | Adgrf5 | adhesion G protein-coupled receptor F5 | -5.31 |
| ENSMUSG00000039372 | 38047 | membrane-associated ring finger (C3HC4) 4 | -5.34 |
| ENSMUSG00000054932 | Afp | alpha fetoprotein | -5.37 |
| ENSMUSG00000048483 | Zdhhc22 | zinc finger, DHHC-type containing 22 | -5.37 |
| ENSMUSG00000044453 | Ffar1 | free fatty acid receptor 1 | -5.38 |
| ENSMUSG00000054640 | Slc8a1 | solute carrier family 8 (sodium/calcium exchanger), member 1 | -5.39 |
| ENSMUSG00000002265 | Peg3 | paternally expressed 3 | -5.39 |
| ENSMUSG00000075012 | Fjx1 | four jointed box 1 | -5.41 |
| ENSMUSG00000061911 | Myt1l | myelin transcription factor 1-like | -5.41 |
| ENSMUSG00000028354 | Fmn2 | formin 2 | -5.42 |
| ENSMUSG00000020181 | Nav3 | neuron navigator 3 | -5.43 |
| ENSMUSG00000051985 | Igfn1 | immunoglobulin-like and fibronectin type III domain containing 1 | -5.47 |
| ENSMUSG00000061171 | Slc38a11 | solute carrier family 38, member 11 | -5.48 |
| ENSMUSG00000111977 | Gm47163 | predicted gene, 47163 | -5.48 |
| ENSMUSG00000033981 | Gria2 | glutamate receptor, ionotropic, AMPA2 (alpha 2) | -5.49 |
| ENSMUSG00000052727 | Map1b | microtubule-associated protein 1B | -5.50 |
| ENSMUSG00000042631 | Xkr7 | X-linked Kx blood group related 7 | -5.53 |
| ENSMUSG00000035277 | Arx | aristaless related homeobox | -5.55 |
| ENSMUSG00000027947 | Il6ra | interleukin 6 receptor, alpha | -5.55 |
| ENSMUSG00000026163 | Sphkap | SPHK1 interactor, AKAP domain containing | -5.55 |
| ENSMUSG00000049148 | Plcxd3 | phosphatidylinositol-specific phospholipase C, X domain containing 3 | -5.56 |
| ENSMUSG00000020723 | Cacng4 | calcium channel, voltage-dependent, gamma subunit 4 | -5.58 |
| ENSMUSG00000027168 | Pax6 | paired box 6 | -5.59 |
| ENSMUSG00000028445 | Enho | energy homeostasis associated | -5.62 |
| ENSMUSG00000022619 | Mapk8ip2 | mitogen-activated protein kinase 8 interacting protein 2 | -5.66 |
| ENSMUSG00000071265 | 1700086L19Rik | RIKEN cDNA 1700086L19 gene | -5.66 |
| ENSMUSG00000029878 | Dbpht2 | DNA binding protein with his-thr domain | -5.66 |
| ENSMUSG00000028008 | Asic5 | acid-sensing (proton-gated) ion channel family member 5 | -5.67 |
| ENSMUSG00000025089 | Gfra1 | glial cell line derived neurotrophic factor family receptor alpha 1 | -5.68 |
| ENSMUSG00000047842 | Diras2 | DIRAS family, GTP-binding RAS-like 2 | -5.70 |
| ENSMUSG00000053166 | Cdh22 | cadherin 22 | -5.74 |
| ENSMUSG00000059921 | Unc5c | unc-5 netrin receptor C | -5.74 |
| ENSMUSG00000020651 | Slc26a4 | solute carrier family 26, member 4 | -5.76 |
| ENSMUSG00000021198 | Unc79 | unc-79 homolog | -5.79 |
| ENSMUSG00000048038 | Ccdc187 | coiled-coil domain containing 187 | -5.87 |
| ENSMUSG00000019890 | Nts | neurotensin | -5.87 |
| ENSMUSG00000039278 | Pcsk1n | proprotein convertase subtilisin/kexin type 1 inhibitor | -5.89 |
| ENSMUSG00000039860 | Srrm3 | serine/arginine repetitive matrix 3 | -5.90 |
| ENSMUSG00000052726 | Kcnt2 | potassium channel, subfamily T, member 2 | -5.92 |
| ENSMUSG00000054423 | Cadps | Ca2+-dependent secretion activator | -5.94 |
| ENSMUSG00000042258 | Isl1 | ISL1 transcription factor, LIM/homeodomain | -5.97 |
| ENSMUSG00000068263 | Efcc1 | EF hand and coiled-coil domain containing 1 | -5.98 |
| ENSMUSG00000031740 | Mmp2 | matrix metallopeptidase 2 | -5.99 |
| ENSMUSG00000004151 | Etv1 | ets variant 1 | -6.01 |
| ENSMUSG00000053852 | Adgrg4 | adhesion G protein-coupled receptor G4 | -6.14 |
| ENSMUSG00000032532 | Cck | cholecystokinin | -6.27 |
| ENSMUSG00000033061 | Resp18 | regulated endocrine-specific protein 18 | -6.30 |
| ENSMUSG00000026824 | Kcnj3 | potassium inwardly-rectifying channel, subfamily J, member 3 | -6.45 |
| ENSMUSG00000050711 | Scg2 | secretogranin II | -6.48 |
| ENSMUSG00000025375 | Aatk | apoptosis-associated tyrosine kinase | -6.51 |
| ENSMUSG00000024304 | Cdh2 | cadherin 2 | -6.52 |

| | | | |
|---|---|---|---|
| ENSMUSG00000015829 | Tnr | tenascin R | -6.56 |
| ENSMUSG00000004110 | Cacna1e | calcium channel, voltage-dependent, R type, alpha 1E subunit | -6.62 |
| ENSMUSG00000043301 | Kcnj6 | potassium inwardly-rectifying channel, subfamily J, member 6 | -6.73 |
| ENSMUSG00000031099 | Smarca1 | SWI/SNF related, matrix associated, actin dependent regulator of chromatin, subfamily a, member 1 | -7.19 |
| ENSMUSG00000014351 | Gip | gastric inhibitory polypeptide | -7.88 |
| ENSMUSG00000055409 | Nell1 | NEL-like 1 | -8.89 |
| ENSMUSG00000103321 | 4933403L11Rik | RIKEN cDNA 4933403L11 gene | -22.00 |

### Supplementary Table S6

Differentially expressed genes (DEGs) in two populations of tdTomato-positive tumor cells sorted for Trop2 expression and in a population of tdTomato-negative (differentiated) cells obtained from microroadenomas 6 weeks after Apc inactivation. DEGs between $Trop2^{+}$ $tdTomato^{+}$ and $Trop2^{-}tdTomato^{+}$ cells and between the two tdTomato-positive cell groups and tdTomato-negative cells are shown on the corresponding sheets. Genes with an adjusted p-value $< 0.05$ and $|\log_2 FC| \geq 1$ were considered significant. Wnt-responsive genes are highlighted in red.

**Tom+Trop2+ vs. Tom+Trop2- cells**

| Ensemble gene code | Symbol | Gene name | logFC |
|---|---|---|---|
| ENSMUSG00000063632 | Sox11 | SRY (sex determining region Y)-box 11 | 6.73 |
| ENSMUSG00000051397 | Tacstd2 | tumor-associated calcium signal transducer 2 | 6.34 |
| ENSMUSG00000004951 | Hspb1 | heat shock protein 1 | 5.49 |
| ENSMUSG00000000120 | Ngfr | nerve growth factor receptor (TNFR superfamily, member 16) | 5.47 |
| ENSMUSG00000030873 | Scnn1b | sodium channel, nonvoltage-gated 1 beta | 5.45 |
| ENSMUSG00000042179 | Pnliprp1 | pancreatic lipase related protein 1 | 5.39 |
| ENSMUSG00000108218 | Olfr1372-ps1 | olfactory receptor 1372, pseudogene 1 | 5.08 |
| ENSMUSG00000007097 | Atp1a2 | ATPase, Na+/K+ transporting, alpha 2 polypeptide | 5.04 |
| ENSMUSG00000029838 | Ptn | pleiotrophin | 4.85 |
| ENSMUSG00000019577 | Pdk4 | pyruvate dehydrogenase kinase, isoenzyme 4 | 4.77 |
| ENSMUSG00000031284 | Pak3 | p21 (RAC1) activated kinase 3 | 4.68 |
| ENSMUSG00000036480 | Prss56 | protease, serine 56 | 4.61 |
| ENSMUSG00000053007 | Creb5 | cAMP responsive element binding protein 5 | 4.60 |
| ENSMUSG00000081392 | Gm11668 | predicted gene 11668 | 4.58 |
| ENSMUSG00000063919 | Srrm4 | serine/arginine repetitive matrix 4 | 4.58 |
| ENSMUSG00000069892 | 9930111J21Rik2 | RIKEN cDNA 9930111J21 gene 2 | 4.58 |
| ENSMUSG00000028031 | Dkk2 | dickkopf WNT signaling pathway inhibitor 2 | 4.56 |
| ENSMUSG00000000901 | Mmp11 | matrix metallopeptidase 11 | 4.56 |
| ENSMUSG00000029648 | Flt1 | FMS-like tyrosine kinase 1 | 4.52 |
| ENSMUSG00000014599 | Csf1 | colony stimulating factor 1 (macrophage) | 4.47 |
| ENSMUSG00000020363 | Gfpt2 | glutamine fructose-6-phosphate transaminase 2 | 4.46 |
| ENSMUSG00000026822 | Lcn2 | lipocalin 2 | 4.45 |
| ENSMUSG00000062591 | Tubb4a | tubulin, beta 4A class IVA | 4.43 |
| ENSMUSG00000034220 | Gpc1 | glypican 1 | 4.43 |
| ENSMUSG00000020218 | Wif1 | Wnt inhibitory factor 1 | 4.40 |
| ENSMUSG00000027674 | Pex5l | peroxisomal biogenesis factor 5-like | 4.40 |
| ENSMUSG00000025473 | Adam8 | a disintegrin and metallopeptidase domain 8 | 4.39 |
| ENSMUSG00000074766 | Ism1 | isthmin 1, angiogenesis inhibitor | 4.37 |
| ENSMUSG00000063531 | Sema3e | sema domain, immunoglobulin domain (Ig), short basic domain, secreted, (semaphorin) 3E | 4.35 |
| ENSMUSG00000044313 | Mab21l3 | mab-21-like 3 (C. elegans) | 4.35 |
| ENSMUSG00000026167 | Wnt10a | wingless-type MMTV integration site family, member 10A | 4.34 |
| ENSMUSG00000032936 | Camkv | CaM kinase-like vesicle-associated | 4.34 |
| ENSMUSG00000022037 | Clu | clusterin | 4.32 |
| ENSMUSG00000046623 | Gjb4 | gap junction protein, beta 4 | 4.31 |
| ENSMUSG00000028328 | Tmod1 | tropomodulin 1 | 4.31 |
| ENSMUSG00000021986 | Amer2 | APC membrane recruitment 2 | 4.29 |
| ENSMUSG00000034450 | Gulo | gulonolactone (L-) oxidase | 4.25 |
| ENSMUSG00000027803 | Wwtr1 | WW domain containing transcription regulator 1 | 4.24 |
| ENSMUSG00000042804 | Gpr153 | G protein-coupled receptor 153 | 4.23 |
| ENSMUSG00000029380 | Cxcl1 | chemokine (C-X-C motif) ligand 1 | 4.21 |
| ENSMUSG00000072591 | 5930412G12Rik | RIKEN cDNA 5930412G12 gene | 4.21 |
| ENSMUSG00000034463 | Scara3 | scavenger receptor class A, member 3 | 4.20 |
| ENSMUSG00000071847 | Apcdd1 | adenomatosis polyposis coli down-regulated 1 | 4.16 |
| ENSMUSG00000068327 | Tlx2 | T cell leukemia, homeobox 2 | 4.15 |
| ENSMUSG00000037188 | Grhl3 | grainyhead like transcription factor 3 | 4.12 |
| ENSMUSG00000018340 | Anxa6 | annexin A6 | 4.11 |
| ENSMUSG00000025324 | Atp10a | ATPase, class V, type 10A | 4.11 |
| ENSMUSG00000085412 | Halr1 | Hoxa adjacent long noncoding RNA 1 | 4.10 |
| ENSMUSG00000009097 | Tbx1 | T-box 1 | 4.09 |
| ENSMUSG00000053613 | Notumos | notum palmitoleoyl-protein carboxylesterase, opposite strand | 4.06 |
| ENSMUSG00000027210 | Meis2 | Meis homeobox 2 | 4.05 |
| ENSMUSG00000006576 | Slc4a3 | solute carrier family 4 (anion exchanger), member 3 | 4.05 |
| ENSMUSG00000030246 | Ldhb | lactate dehydrogenase B | 4.00 |
| ENSMUSG00000024164 | C3 | complement component 3 | 3.98 |
| ENSMUSG00000031538 | Plat | plasminogen activator, tissue | 3.97 |
| ENSMUSG00000025140 | Pycr1 | pyrroline-5-carboxylate reductase 1 | 3.97 |
| ENSMUSG00000043903 | Zfp469 | zinc finger protein 469 | 3.95 |
| ENSMUSG00000034612 | Chst11 | carbohydrate sulfotransferase 11 | 3.88 |
| ENSMUSG00000039232 | Stx11 | syntaxin 11 | 3.88 |
| ENSMUSG00000087201 | Gm15261 | predicted gene 15261 | 3.88 |
| ENSMUSG00000016763 | Scube1 | signal peptide, CUB domain, EGF-like 1 | 3.86 |
| ENSMUSG00000029108 | Pcdh7 | protocadherin 7 | 3.85 |
| ENSMUSG00000049807 | Arhgap23 | Rho GTPase activating protein 23 | 3.84 |
| ENSMUSG00000054555 | Adam12 | a disintegrin and metallopeptidase domain 12 (meltrin alpha) | 3.81 |
| ENSMUSG00000039959 | Hip1 | huntingtin interacting protein 1 | 3.80 |
| ENSMUSG00000026764 | Kif5c | kinesin family member 5C | 3.80 |
| ENSMUSG00000043430 | Psapl1 | prosaposin-like 1 | 3.79 |
| ENSMUSG00000039153 | Runx2 | runt related transcription factor 2 | 3.79 |
| ENSMUSG00000004791 | Pgf | placental growth factor | 3.79 |

| | | | |
|---|---|---|---|
| ENSMUSG00000014602 | Kif1a | kinesin family member 1A | 3.77 |
| ENSMUSG00000017466 | Timp2 | tissue inhibitor of metalloproteinase 2 | 3.77 |
| ENSMUSG00000032496 | Ltf | lactotransferrin | 3.77 |
| ENSMUSG00000025330 | Padi4 | peptidyl arginine deiminase, type IV | 3.75 |
| ENSMUSG00000078161 | Erich3 | glutamate rich 3 | 3.74 |
| ENSMUSG00000020092 | Pald1 | phosphatase domain containing, paladin 1 | 3.74 |
| ENSMUSG00000104918 | Gm42944 | predicted gene 42944 | 3.74 |
| ENSMUSG00000022665 | Ccdc80 | coiled-coil domain containing 80 | 3.73 |
| ENSMUSG00000048960 | Prex2 | phosphatidylinositol-3,4,5-trisphosphate-dependent Rac exchange factor 2 | 3.73 |
| ENSMUSG00000021765 | Fst | follistatin | 3.70 |
| ENSMUSG00000038572 | Bpifb5 | BPI fold containing family B, member 5 | 3.69 |
| ENSMUSG00000020900 | Myh10 | myosin, heavy polypeptide 10, non-muscle | 3.64 |
| ENSMUSG00000063727 | Tnfrsf11b | tumor necrosis factor receptor superfamily, member 11b (osteoprotegerin) | 3.61 |
| ENSMUSG00000026728 | Vim | vimentin | 3.61 |
| ENSMUSG00000058297 | Spock2 | sparc/osteonectin, cwcv and kazal-like domains proteoglycan 2 | 3.61 |
| ENSMUSG00000011305 | Plin5 | perilipin 5 | 3.61 |
| ENSMUSG00000040093 | Bmf | BCL2 modifying factor | 3.60 |
| ENSMUSG00000098132 | Rassf10 | Ras association (RalGDS/AF-6) domain family (N-terminal) member 10 | 3.58 |
| ENSMUSG00000029368 | Alb | albumin | 3.56 |
| ENSMUSG00000041608 | Entpd3 | ectonucleoside triphosphate diphosphohydrolase 3 | 3.54 |
| ENSMUSG00000033508 | Asprv1 | aspartic peptidase, retroviral-like 1 | 3.54 |
| ENSMUSG00000009628 | Tex15 | testis expressed gene 15 | 3.53 |
| ENSMUSG00000013419 | Zfp651 | zinc finger protein 651 | 3.52 |
| ENSMUSG00000028360 | Slc44a5 | solute carrier family 44, member 5 | 3.51 |
| ENSMUSG00000054932 | Afp | alpha fetoprotein | 3.51 |
| ENSMUSG00000032965 | Ift57 | intraflagellar transport 57 | 3.50 |
| ENSMUSG00000013584 | Aldh1a2 | aldehyde dehydrogenase family 1, subfamily A2 | 3.50 |
| ENSMUSG00000084822 | Myadml2os | myeloid-associated differentiation marker-like 2, opposite strand | 3.50 |
| ENSMUSG00000034764 | 1700006J14Rik | RIKEN cDNA 1700006J14 gene | 3.49 |
| ENSMUSG00000048191 | Muc6 | mucin 6, gastric | 3.48 |
| ENSMUSG00000041293 | Adgrf1 | adhesion G protein-coupled receptor F1 | 3.48 |
| ENSMUSG00000020651 | Slc26a4 | solute carrier family 26, member 4 | 3.47 |
| ENSMUSG00000006205 | Htra1 | HtrA serine peptidase 1 | 3.46 |
| ENSMUSG00000029761 | Cald1 | caldesmon 1 | 3.46 |
| ENSMUSG00000041189 | Chrnb1 | cholinergic receptor, nicotinic, beta polypeptide 1 (muscle) | 3.46 |
| ENSMUSG00000030772 | Dkk3 | dickkopf WNT signaling pathway inhibitor 3 | 3.45 |
| ENSMUSG00000022790 | Igsf11 | immunoglobulin superfamily, member 11 | 3.45 |
| ENSMUSG00000023828 | Slc22a3 | solute carrier family 22 (organic cation transporter), member 3 | 3.45 |
| ENSMUSG00000001864 | Aif1l | allograft inflammatory factor 1-like | 3.44 |
| ENSMUSG00000041046 | Ramp3 | receptor (calcitonin) activity modifying protein 3 | 3.43 |
| ENSMUSG00000085811 | Cep112it | centrosomal protein 112, intronic transcript | 3.43 |
| ENSMUSG00000034634 | Ly6d | lymphocyte antigen 6 complex, locus D | 3.43 |
| ENSMUSG00000025666 | Tmem47 | transmembrane protein 47 | 3.43 |
| ENSMUSG00000005503 | Evx1 | even-skipped homeobox 1 | 3.42 |
| ENSMUSG00000025352 | Gdf11 | growth differentiation factor 11 | 3.41 |
| ENSMUSG00000031595 | Pdgfrl | platelet-derived growth factor receptor-like | 3.41 |
| ENSMUSG00000020086 | H2afy2 | H2A histone family, member Y2 | 3.41 |
| ENSMUSG00000031925 | Maml2 | mastermind like transcriptional coactivator 2 | 3.41 |
| ENSMUSG00000035783 | Acta2 | actin, alpha 2, smooth muscle, aorta | 3.41 |
| ENSMUSG00000017204 | Gsdma | gasdermin A | 3.40 |
| ENSMUSG00000040998 | Npnt | nephronectin | 3.40 |
| ENSMUSG00000049556 | Lingo1 | leucine rich repeat and Ig domain containing 1 | 3.40 |
| ENSMUSG00000025938 | Slco5a1 | solute carrier organic anion transporter family, member 5A1 | 3.39 |
| ENSMUSG00000033763 | Mtss1l | metastasis suppressor 1-like | 3.39 |
| ENSMUSG00000037035 | Inhbb | inhibin beta-B | 3.39 |
| ENSMUSG00000017491 | Rarb | retinoic acid receptor, beta | 3.39 |
| ENSMUSG00000032186 | Tmod2 | tropomodulin 2 | 3.39 |
| ENSMUSG00000054967 | Zfp647 | zinc finger protein 647 | 3.39 |
| ENSMUSG00000070436 | Serpinh1 | serine (or cysteine) peptidase inhibitor, clade H, member 1 | 3.39 |
| ENSMUSG00000022594 | Lynx1 | Ly6/neurotoxin 1 | 3.36 |
| ENSMUSG00000078922 | Tgtp1 | T cell specific GTPase 1 | 3.34 |
| ENSMUSG00000110187 | Gm45496 | predicted gene 45496 | 3.33 |
| ENSMUSG00000053137 | Mapk11 | mitogen-activated protein kinase 11 | 3.32 |
| ENSMUSG00000033209 | Ttc28 | tetratricopeptide repeat domain 28 | 3.31 |
| ENSMUSG00000055980 | Irs1 | insulin receptor substrate 1 | 3.31 |
| ENSMUSG00000066438 | Plekhd1 | pleckstrin homology domain containing, family D (with coiled-coil domains) member 1 | 3.30 |
| ENSMUSG00000024912 | Fosl1 | fos-like antigen 1 | 3.27 |
| ENSMUSG00000028214 | Gem | GTP binding protein (gene overexpressed in skeletal muscle) | 3.27 |
| ENSMUSG00000059854 | Hydin | HYDIN, axonemal central pair apparatus protein | 3.27 |
| ENSMUSG00000046191 | Pcdhb20 | protocadherin beta 20 | 3.27 |
| ENSMUSG00000050640 | Tmem150c | transmembrane protein 150C | 3.26 |
| ENSMUSG00000004552 | Ctse | cathepsin E | 3.26 |

| | | | |
|---|---|---|---|
| ENSMUSG00000041324 | Inhba | inhibin beta-A | 3.26 |
| ENSMUSG00000029769 | Ccdc136 | coiled-coil domain containing 136 | 3.25 |
| ENSMUSG00000110290 | Gm45336 | predicted gene 45336 | 3.25 |
| ENSMUSG00000055333 | Fat2 | FAT atypical cadherin 2 | 3.23 |
| ENSMUSG00000013338 | Fer1l4 | fer-1-like 4 (C. elegans) | 3.23 |
| ENSMUSG00000018211 | Wfdc15b | WAP four-disulfide core domain 15B | 3.23 |
| ENSMUSG00000037129 | Tmprss13 | transmembrane protease, serine 13 | 3.22 |
| ENSMUSG00000051486 | Pcdhb11 | protocadherin beta 11 | 3.21 |
| ENSMUSG00000026620 | Mark1 | MAP/microtubule affinity regulating kinase 1 | 3.20 |
| ENSMUSG00000029757 | Dync1i1 | dynein cytoplasmic 1 intermediate chain 1 | 3.19 |
| ENSMUSG00000025854 | Fam20c | family with sequence similarity 20, member C | 3.18 |
| ENSMUSG00000080316 | Spaca6 | sperm acrosome associated 6 | 3.18 |
| ENSMUSG00000023009 | Nckap5l | NCK-associated protein 5-like | 3.17 |
| ENSMUSG00000086395 | A630014C17Rik | RIKEN cDNA A630014C17 gene | 3.17 |
| ENSMUSG00000031841 | Cdh13 | cadherin 13 | 3.16 |
| ENSMUSG00000038587 | Akap12 | A kinase (PRKA) anchor protein (gravin) 12 | 3.14 |
| ENSMUSG00000004044 | Cavin1 | caveolae associated 1 | 3.13 |
| ENSMUSG00000040152 | Thbs1 | thrombospondin 1 | 3.12 |
| ENSMUSG00000000983 | Wfdc18 | WAP four-disulfide core domain 18 | 3.12 |
| ENSMUSG00000020674 | Pxdn | peroxidasin | 3.11 |
| ENSMUSG00000079363 | Gbp4 | guanylate binding protein 4 | 3.11 |
| ENSMUSG00000027375 | Mal | myelin and lymphocyte protein, T cell differentiation protein | 3.11 |
| ENSMUSG00000032363 | Adamts7 | a disintegrin-like and metallopeptidase (reprolysin type) with thrombospondin type 1 motif, 7 | 3.10 |
| ENSMUSG00000019990 | Pde7b | phosphodiesterase 7B | 3.09 |
| ENSMUSG00000027376 | Prom2 | prominin 2 | 3.09 |
| ENSMUSG00000042988 | Notum | notum palmitoleoyl-protein carboxylesterase | 3.08 |
| ENSMUSG00000068923 | Syt11 | synaptotagmin XI | 3.08 |
| ENSMUSG00000046618 | Olfml2a | olfactomedin-like 2A | 3.08 |
| ENSMUSG00000029287 | Tgfbr3 | transforming growth factor, beta receptor III | 3.08 |
| ENSMUSG00000018012 | Rac3 | Rac family small GTPase 3 | 3.07 |
| ENSMUSG00000022883 | Robo1 | roundabout guidance receptor 1 | 3.07 |
| ENSMUSG00000093908 | Gm5784 | predicted gene 5784 | 3.07 |
| ENSMUSG00000044033 | Ccdc141 | coiled-coil domain containing 141 | 3.06 |
| ENSMUSG00000034295 | Fhod3 | formin homology 2 domain containing 3 | 3.06 |
| ENSMUSG00000058656 | Samd12 | sterile alpha motif domain containing 12 | 3.05 |
| ENSMUSG00000038456 | Dennd2a | DENN/MADD domain containing 2A | 3.05 |
| ENSMUSG00000055799 | Tcf7l1 | transcription factor 7 like 1 (T cell specific, HMG box) | 3.05 |
| ENSMUSG00000033389 | Arhgap44 | Rho GTPase activating protein 44 | 3.05 |
| ENSMUSG00000026494 | Kif26b | kinesin family member 26B | 3.04 |
| ENSMUSG00000048078 | Tenm4 | teneurin transmembrane protein 4 | 3.04 |
| ENSMUSG00000046449 | Nexmif | neurite extension and migration factor | 3.04 |
| ENSMUSG00000049866 | Arl4c | ADP-ribosylation factor-like 4C | 3.03 |
| ENSMUSG00000037016 | Frem2 | Fras1 related extracellular matrix protein 2 | 3.03 |
| ENSMUSG00000022456 | Sept3 | septin 3 | 3.03 |
| ENSMUSG00000031328 | Flna | filamin, alpha | 3.02 |
| ENSMUSG00000040714 | Klc3 | kinesin light chain 3 | 3.02 |
| ENSMUSG00000110666 | Gm9172 | predicted gene 9172 | 3.02 |
| ENSMUSG00000038576 | Susd4 | sushi domain containing 4 | 3.02 |
| ENSMUSG00000068699 | Flnc | filamin C, gamma | 3.01 |
| ENSMUSG00000029298 | Gbp9 | guanylate-binding protein 9 | 2.99 |
| ENSMUSG00000013076 | Amotl1 | angiomotin-like 1 | 2.98 |
| ENSMUSG00000078921 | Tgtp2 | T cell specific GTPase 2 | 2.98 |
| ENSMUSG00000050212 | Eva1b | eva-1 homolog B (C. elegans) | 2.98 |
| ENSMUSG00000044071 | Fam19a2 | family with sequence similarity 19, member A2 | 2.98 |
| ENSMUSG00000039693 | Msantd3 | Myb/SANT-like DNA-binding domain containing 3 | 2.97 |
| ENSMUSG00000042357 | Gjb5 | gap junction protein, beta 5 | 2.96 |
| ENSMUSG00000026193 | Fn1 | fibronectin 1 | 2.95 |
| ENSMUSG00000046731 | Kctd11 | potassium channel tetramerisation domain containing 11 | 2.95 |
| ENSMUSG00000034675 | Dbn1 | drebrin 1 | 2.95 |
| ENSMUSG00000075304 | Sp5 | trans-acting transcription factor 5 | 2.94 |
| ENSMUSG00000037712 | Fermt2 | fermitin family member 2 | 2.94 |
| ENSMUSG00000054263 | Lifr | leukemia inhibitory factor receptor | 2.94 |
| ENSMUSG00000026117 | Zap70 | zeta-chain (TCR) associated protein kinase | 2.94 |
| ENSMUSG00000085111 | Ascl4 | achaete-scute family bHLH transcription factor 4 | 2.94 |
| ENSMUSG00000041889 | Shisa4 | shisa family member 4 | 2.94 |
| ENSMUSG00000024659 | Anxa1 | annexin A1 | 2.93 |
| ENSMUSG00000071531 | Gprin2 | G protein regulated inducer of neurite outgrowth 2 | 2.93 |
| ENSMUSG00000024940 | Ltbp3 | latent transforming growth factor beta binding protein 3 | 2.91 |
| ENSMUSG00000034460 | Six4 | sine oculis-related homeobox 4 | 2.91 |
| ENSMUSG00000037833 | Sh2d4b | SH2 domain containing 4B | 2.91 |
| ENSMUSG00000036599 | Chst12 | carbohydrate sulfotransferase 12 | 2.91 |

| | | | |
|---|---|---|---|
| ENSMUSG00000037813 | D630003M21Rik | RIKEN cDNA D630003M21 gene | 2.90 |
| ENSMUSG00000033227 | Wnt6 | wingless-type MMTV integration site family, member 6 | 2.90 |
| ENSMUSG00000023008 | Fmnl3 | formin-like 3 | 2.89 |
| ENSMUSG00000021835 | Bmp4 | bone morphogenetic protein 4 | 2.89 |
| ENSMUSG00000104612 | Gm42449 | predicted gene 42449 | 2.89 |
| ENSMUSG00000020181 | Nav3 | neuron navigator 3 | 2.89 |
| ENSMUSG00000072437 | Nanos1 | nanos C2HC-type zinc finger 1 | 2.88 |
| ENSMUSG00000029869 | Ephb6 | Eph receptor B6 | 2.88 |
| ENSMUSG00000032717 | Mdfi | MyoD family inhibitor | 2.88 |
| ENSMUSG00000037681 | Esyt3 | extended synaptotagmin-like protein 3 | 2.87 |
| ENSMUSG00000024206 | Rfx2 | regulatory factor X, 2 (influences HLA class II expression) | 2.87 |
| ENSMUSG00000010175 | Prox1 | prospero homeobox 1 | 2.86 |
| ENSMUSG00000046169 | Adamts6 | a disintegrin-like and metallopeptidase (reprolysin type) with thrombospondin type 1 motif, 6 | 2.86 |
| ENSMUSG00000026765 | Lypd6b | LY6/PLAUR domain containing 6B | 2.86 |
| ENSMUSG00000046971 | Pla2g4f | phospholipase A2, group IVF | 2.86 |
| ENSMUSG00000034573 | Ptpn13 | protein tyrosine phosphatase, non-receptor type 13 | 2.86 |
| ENSMUSG00000067889 | Sptbn2 | spectrin beta, non-erythrocytic 2 | 2.85 |
| ENSMUSG00000031661 | Nkd1 | naked cuticle 1 | 2.84 |
| ENSMUSG00000014813 | Stc1 | stanniocalcin 1 | 2.84 |
| ENSMUSG00000039676 | Capsl | calcyphosine-like | 2.84 |
| ENSMUSG00000103189 | Gm37092 | predicted gene, 37092 | 2.84 |
| ENSMUSG00000035357 | Pdzrn3 | PDZ domain containing RING finger 3 | 2.83 |
| ENSMUSG00000029359 | Tesc | tescalcin | 2.83 |
| ENSMUSG00000031871 | Cdh5 | cadherin 5 | 2.83 |
| ENSMUSG00000035246 | Pcyt1b | phosphate cytidylyltransferase 1, choline, beta isoform | 2.83 |
| ENSMUSG00000002980 | Bcam | basal cell adhesion molecule | 2.83 |
| ENSMUSG00000001288 | Rarg | retinoic acid receptor, gamma | 2.82 |
| ENSMUSG00000022176 | Rem2 | rad and gem related GTP binding protein 2 | 2.82 |
| ENSMUSG00000105096 | Gbp10 | guanylate-binding protein 10 | 2.82 |
| ENSMUSG00000020723 | Cacng4 | calcium channel, voltage-dependent, gamma subunit 4 | 2.81 |
| ENSMUSG00000057193 | Slc44a2 | solute carrier family 44, member 2 | 2.80 |
| ENSMUSG00000041390 | Mdfic | MyoD family inhibitor domain containing | 2.80 |
| ENSMUSG00000051243 | Islr2 | immunoglobulin superfamily containing leucine-rich repeat 2 | 2.80 |
| ENSMUSG00000035678 | Tnfsf9 | tumor necrosis factor (ligand) superfamily, member 9 | 2.78 |
| ENSMUSG00000040430 | Pitpnc1 | phosphatidylinositol transfer protein, cytoplasmic 1 | 2.77 |
| ENSMUSG00000070337 | Gpr179 | G protein-coupled receptor 179 | 2.77 |
| ENSMUSG00000031355 | Arhgap6 | Rho GTPase activating protein 6 | 2.76 |
| ENSMUSG00000050052 | Tdrp | testis development related protein | 2.75 |
| ENSMUSG00000027985 | Lef1 | lymphoid enhancer binding factor 1 | 2.74 |
| ENSMUSG00000024909 | Efemp2 | epidermal growth factor-containing fibulin-like extracellular matrix protein 2 | 2.74 |
| ENSMUSG00000044948 | Cfap43 | cilia and flagella associated protein 43 | 2.74 |
| ENSMUSG00000071984 | Fndc1 | fibronectin type III domain containing 1 | 2.74 |
| ENSMUSG00000069255 | Dusp22 | dual specificity phosphatase 22 | 2.73 |
| ENSMUSG00000028883 | Sema3a | sema domain, immunoglobulin domain (Ig), short basic domain, secreted, (semaphorin) 3A | 2.73 |
| ENSMUSG00000068748 | Ptprz1 | protein tyrosine phosphatase, receptor type Z, polypeptide 1 | 2.72 |
| ENSMUSG00000039607 | Rbms3 | RNA binding motif, single stranded interacting protein | 2.71 |
| ENSMUSG00000070304 | Scn2b | sodium channel, voltage-gated, type II, beta | 2.71 |
| ENSMUSG00000035954 | Dock4 | dedicator of cytokinesis 4 | 2.71 |
| ENSMUSG00000028497 | Hacd4 | 3-hydroxyacyl-CoA dehydratase 4 | 2.71 |
| ENSMUSG00000045667 | Smtnl2 | smoothelin-like 2 | 2.71 |
| ENSMUSG00000022435 | Upk3a | uroplakin 3A | 2.71 |
| ENSMUSG00000063851 | Rnf183 | ring finger protein 183 | 2.71 |
| ENSMUSG00000038415 | Foxq1 | forkhead box Q1 | 2.70 |
| ENSMUSG00000070867 | Trabd2b | TraB domain containing 2B | 2.70 |
| ENSMUSG00000035274 | Tpbg | trophoblast glycoprotein | 2.69 |
| ENSMUSG00000015647 | Lama5 | laminin, alpha 5 | 2.68 |
| ENSMUSG00000041134 | Cyyr1 | cysteine and tyrosine-rich protein 1 | 2.68 |
| ENSMUSG00000020427 | Igfbp3 | insulin-like growth factor binding protein 3 | 2.68 |
| ENSMUSG00000032327 | Stra6 | stimulated by retinoic acid gene 6 | 2.68 |
| ENSMUSG00000032875 | Arhgef17 | Rho guanine nucleotide exchange factor (GEF) 17 | 2.67 |
| ENSMUSG00000060429 | Sntb1 | syntrophin, basic 1 | 2.67 |
| ENSMUSG00000020814 | Mxra7 | matrix-remodelling associated 7 | 2.67 |
| ENSMUSG00000040852 | Plekhh2 | pleckstrin homology domain containing, family H (with MyTH4 domain) member 2 | 2.66 |
| ENSMUSG00000111212 | Gm47087 | predicted gene, 47087 | 2.66 |
| ENSMUSG00000026556 | Vangl2 | VANGL planar cell polarity 2 | 2.65 |
| ENSMUSG00000040488 | Ltbp4 | latent transforming growth factor beta binding protein 4 | 2.65 |
| ENSMUSG00000052911 | Lamb2 | laminin, beta 2 | 2.65 |
| ENSMUSG00000032076 | Cadm1 | cell adhesion molecule 1 | 2.65 |
| ENSMUSG00000058317 | Ube2e2 | ubiquitin-conjugating enzyme E2E 2 | 2.65 |
| ENSMUSG00000033377 | Palmd | palmdelphin | 2.65 |
| ENSMUSG00000034684 | Sema3f | sema domain, immunoglobulin domain (Ig), short basic domain, secreted, (semaphorin) 3F | 2.63 |

| | | | |
|---|---|---|---|
| ENSMUSG00000020044 | Timp3 | tissue inhibitor of metalloproteinase 3 | 2.63 |
| ENSMUSG00000036502 | Tmem255a | transmembrane protein 255A | 2.63 |
| ENSMUSG00000015134 | Aldh1a3 | aldehyde dehydrogenase family 1, subfamily A3 | 2.62 |
| ENSMUSG00000022587 | Ly6e | lymphocyte antigen 6 complex, locus E | 2.61 |
| ENSMUSG00000040836 | Gpr161 | G protein-coupled receptor 161 | 2.61 |
| ENSMUSG00000026475 | Rgs16 | regulator of G-protein signaling 16 | 2.61 |
| ENSMUSG00000097344 | Gm26732 | predicted gene, 26732 | 2.61 |
| ENSMUSG00000021493 | Pdlim7 | PDZ and LIM domain 7 | 2.60 |
| ENSMUSG00000031849 | Comp | cartilage oligomeric matrix protein | 2.60 |
| ENSMUSG00000058145 | Adamts17 | a disintegrin-like and metallopeptidase (reprolysin type) with thrombospondin type 1 motif, 17 | 2.59 |
| ENSMUSG00000048644 | Ctxn1 | cortexin 1 | 2.59 |
| ENSMUSG00000000552 | Zfp385a | zinc finger protein 385A | 2.59 |
| ENSMUSG00000020599 | Rgs9 | regulator of G-protein signaling 9 | 2.58 |
| ENSMUSG00000029771 | Irf5 | interferon regulatory factor 5 | 2.58 |
| ENSMUSG00000061517 | Sox21 | SRY (sex determining region Y)-box 21 | 2.57 |
| ENSMUSG00000036053 | Fmnl2 | formin-like 2 | 2.56 |
| ENSMUSG00000047747 | Rnf150 | ring finger protein 150 | 2.56 |
| ENSMUSG00000090394 | 4930523C07Rik | RIKEN cDNA 4930523C07 gene | 2.56 |
| ENSMUSG00000038496 | Slc19a3 | solute carrier family 19, member 3 | 2.56 |
| ENSMUSG00000108833 | Gm17909 | predicted gene, 17909 | 2.56 |
| ENSMUSG00000038552 | Fndc4 | fibronectin type III domain containing 4 | 2.56 |
| ENSMUSG00000022871 | Fetub | fetuin beta | 2.56 |
| ENSMUSG00000024235 | Map3k8 | mitogen-activated protein kinase kinase kinase 8 | 2.55 |
| ENSMUSG00000032719 | Sbspon | somatomedin B and thrombospondin, type 1 domain containing | 2.55 |
| ENSMUSG00000025492 | Ifitm3 | interferon induced transmembrane protein 3 | 2.54 |
| ENSMUSG00000006411 | Nectin4 | nectin cell adhesion molecule 4 | 2.54 |
| ENSMUSG00000022146 | Osmr | oncostatin M receptor | 2.54 |
| ENSMUSG00000032135 | Mcam | melanoma cell adhesion molecule | 2.54 |
| ENSMUSG00000050989 | Selenon | selenoprotein N | 2.53 |
| ENSMUSG00000025855 | Prkar1b | protein kinase, cAMP dependent regulatory, type I beta | 2.53 |
| ENSMUSG00000039601 | Rcan2 | regulator of calcineurin 2 | 2.52 |
| ENSMUSG00000030600 | Lrfn1 | leucine rich repeat and fibronectin type III domain containing 1 | 2.52 |
| ENSMUSG00000013089 | Etv5 | ets variant 5 | 2.51 |
| ENSMUSG00000002633 | Shh | sonic hedgehog | 2.51 |
| ENSMUSG00000030074 | Gxylt2 | glucoside xylosyltransferase 2 | 2.51 |
| ENSMUSG00000030605 | Mfge8 | milk fat globule-EGF factor 8 protein | 2.50 |
| ENSMUSG00000041801 | Phlda3 | pleckstrin homology like domain, family A, member 3 | 2.50 |
| ENSMUSG00000036158 | Prickle1 | prickle planar cell polarity protein 1 | 2.50 |
| ENSMUSG00000026473 | Glul | glutamate-ammonia ligase (glutamine synthetase) | 2.49 |
| ENSMUSG00000049281 | Scn3b | sodium channel, voltage-gated, type III, beta | 2.49 |
| ENSMUSG00000051236 | Msrb3 | methionine sulfoxide reductase B3 | 2.48 |
| ENSMUSG00000013236 | Ptprs | protein tyrosine phosphatase, receptor type, S | 2.47 |
| ENSMUSG00000058952 | Cfi | complement component factor i | 2.47 |
| ENSMUSG00000089715 | Cbx6 | chromobox 6 | 2.45 |
| ENSMUSG00000029101 | Rgs12 | regulator of G-protein signaling 12 | 2.45 |
| ENSMUSG00000037902 | Sirpa | signal-regulatory protein alpha | 2.45 |
| ENSMUSG00000027257 | Pacsin3 | protein kinase C and casein kinase substrate in neurons 3 | 2.44 |
| ENSMUSG00000022464 | Slc38a4 | solute carrier family 38, member 4 | 2.43 |
| ENSMUSG00000034593 | Myo5a | myosin VA | 2.43 |
| ENSMUSG00000027223 | Mapk8ip1 | mitogen-activated protein kinase 8 interacting protein 1 | 2.43 |
| ENSMUSG00000028246 | Faxc | failed axon connections homolog | 2.43 |
| ENSMUSG00000053477 | Tcf4 | transcription factor 4 | 2.42 |
| ENSMUSG00000024349 | Tmem173 | transmembrane protein 173 | 2.42 |
| ENSMUSG00000035376 | Hacd2 | 3-hydroxyacyl-CoA dehydratase 2 | 2.42 |
| ENSMUSG00000031398 | Plxna3 | plexin A3 | 2.42 |
| ENSMUSG00000059668 | Krt4 | keratin 4 | 2.42 |
| ENSMUSG00000009687 | Fxyd5 | FXYD domain-containing ion transport regulator 5 | 2.42 |
| ENSMUSG00000112249 | Gm30262 | predicted gene, 30262 | 2.42 |
| ENSMUSG00000030796 | Tead2 | TEA domain family member 2 | 2.41 |
| ENSMUSG00000009739 | Pou6f1 | POU domain, class 6, transcription factor 1 | 2.41 |
| ENSMUSG00000023972 | Ptk7 | PTK7 protein tyrosine kinase 7 | 2.40 |
| ENSMUSG00000008398 | Elk3 | ELK3, member of ETS oncogene family | 2.40 |
| ENSMUSG00000035640 | Cbarp | calcium channel, voltage-dependent, beta subunit associated regulatory protein | 2.40 |
| ENSMUSG00000024401 | Tnf | tumor necrosis factor | 2.40 |
| ENSMUSG00000053646 | Plxnb1 | plexin B1 | 2.38 |
| ENSMUSG00000002603 | Tgfb1 | transforming growth factor, beta 1 | 2.38 |
| ENSMUSG00000001131 | Timp1 | tissue inhibitor of metalloproteinase 1 | 2.38 |
| ENSMUSG00000020262 | Adarb1 | adenosine deaminase, RNA-specific, B1 | 2.38 |
| ENSMUSG00000110802 | Gm47141 | predicted gene, 47141 | 2.38 |
| ENSMUSG00000026579 | F5 | coagulation factor V | 2.37 |
| ENSMUSG00000045545 | Krt14 | keratin 14 | 2.37 |

| | | | |
|---|---|---|---|
| ENSMUSG00000025202 | Scd3 | stearoyl-coenzyme A desaturase 3 | 2.37 |
| ENSMUSG00000043391 | 2510009E07Rik | RIKEN cDNA 2510009E07 gene | 2.36 |
| ENSMUSG00000019853 | Hebp2 | heme binding protein 2 | 2.36 |
| ENSMUSG00000053049 | Gm15413 | predicted gene 15413 | 2.36 |
| ENSMUSG00000020728 | Cep112 | centrosomal protein 112 | 2.35 |
| ENSMUSG00000014329 | Bicc1 | BicC family RNA binding protein 1 | 2.35 |
| ENSMUSG00000085851 | 4921518K17Rik | RIKEN cDNA 4921518K17 gene | 2.35 |
| ENSMUSG00000074480 | Mex3a | mex3 RNA binding family member A | 2.34 |
| ENSMUSG00000040415 | Dtx3 | deltex 3, E3 ubiquitin ligase | 2.34 |
| ENSMUSG00000026676 | Ccdc3 | coiled-coil domain containing 3 | 2.33 |
| ENSMUSG00000022371 | Col14a1 | collagen, type XIV, alpha 1 | 2.33 |
| ENSMUSG00000028100 | Nudt17 | nudix (nucleoside diphosphate linked moiety X)-type motif 17 | 2.33 |
| ENSMUSG00000034685 | Fam171a2 | family with sequence similarity 171, member A2 | 2.33 |
| ENSMUSG00000000782 | Tcf7 | transcription factor 7, T cell specific | 2.32 |
| ENSMUSG00000037095 | Lrg1 | leucine-rich alpha-2-glycoprotein 1 | 2.32 |
| ENSMUSG00000024810 | Il33 | interleukin 33 | 2.32 |
| ENSMUSG00000052105 | Mtcl1 | microtubule crosslinking factor 1 | 2.31 |
| ENSMUSG00000022421 | Nptxr | neuronal pentraxin receptor | 2.31 |
| ENSMUSG00000049848 | Ceacam19 | carcinoembryonic antigen-related cell adhesion molecule 19 | 2.30 |
| ENSMUSG00000031760 | Mt3 | metallothionein 3 | 2.30 |
| ENSMUSG00000060166 | Zdhhc8 | zinc finger, DHHC domain containing 8 | 2.29 |
| ENSMUSG00000113211 | 4921525O09Rik | RIKEN cDNA 4921525O09 gene | 2.29 |
| ENSMUSG00000016526 | Dyrk3 | dual-specificity tyrosine-(Y)-phosphorylation regulated kinase 3 | 2.27 |
| ENSMUSG00000041961 | Znrf3 | zinc and ring finger 3 | 2.26 |
| ENSMUSG00000041570 | Camsap2 | calmodulin regulated spectrin-associated protein family, member 2 | 2.26 |
| ENSMUSG00000029832 | Nfe2l3 | nuclear factor, erythroid derived 2, like 3 | 2.26 |
| ENSMUSG00000021747 | 4930452B06Rik | RIKEN cDNA 4930452B06 gene | 2.26 |
| ENSMUSG00000037053 | Azgp1 | alpha-2-glycoprotein 1, zinc | 2.25 |
| ENSMUSG00000050071 | Bex1 | brain expressed X-linked 1 | 2.24 |
| ENSMUSG00000028199 | Cryz | crystallin, zeta | 2.24 |
| ENSMUSG00000021256 | Vash1 | vasohibin 1 | 2.24 |
| ENSMUSG00000040690 | Col16a1 | collagen, type XVI, alpha 1 | 2.23 |
| ENSMUSG00000020937 | Plcd3 | phospholipase C, delta 3 | 2.23 |
| ENSMUSG00000039004 | Bmp6 | bone morphogenetic protein 6 | 2.22 |
| ENSMUSG00000032625 | Thsd7a | thrombospondin, type I, domain containing 7A | 2.21 |
| ENSMUSG00000033287 | Kctd17 | potassium channel tetramerisation domain containing 17 | 2.21 |
| ENSMUSG00000082361 | Btc | betacellulin, epidermal growth factor family member | 2.20 |
| ENSMUSG00000063455 | D630045J12Rik | RIKEN cDNA D630045J12 gene | 2.20 |
| ENSMUSG00000025592 | Dach2 | dachshund family transcription factor 2 | 2.20 |
| ENSMUSG00000045136 | Tubb2b | tubulin, beta 2B class IIB | 2.18 |
| ENSMUSG00000020844 | Nxn | nucleoredoxin | 2.18 |
| ENSMUSG00000024399 | Ltb | lymphotoxin B | 2.18 |
| ENSMUSG00000001435 | Col18a1 | collagen, type XVIII, alpha 1 | 2.17 |
| ENSMUSG00000043639 | Rbm20 | RNA binding motif protein 20 | 2.17 |
| ENSMUSG00000038156 | Spon1 | spondin 1, (f-spondin) extracellular matrix protein | 2.17 |
| ENSMUSG00000042306 | S100a14 | S100 calcium binding protein A14 | 2.15 |
| ENSMUSG00000056856 | Jakmip3 | janus kinase and microtubule interacting protein 3 | 2.15 |
| ENSMUSG00000025038 | Efhc2 | EF-hand domain (C-terminal) containing 2 | 2.15 |
| ENSMUSG00000031441 | Atp11a | ATPase, class VI, type 11A | 2.14 |
| ENSMUSG00000022696 | Sidt1 | SID1 transmembrane family, member 1 | 2.14 |
| ENSMUSG00000028698 | Pik3r3 | phosphoinositide-3-kinase regulatory subunit 3 | 2.14 |
| ENSMUSG00000054793 | Cadm4 | cell adhesion molecule 4 | 2.14 |
| ENSMUSG00000105681 | Gm43428 | predicted gene 43428 | 2.14 |
| ENSMUSG00000044043 | Pcdhb14 | protocadherin beta 14 | 2.14 |
| ENSMUSG00000039457 | Ppl | periplakin | 2.13 |
| ENSMUSG00000041930 | Fam222a | family with sequence similarity 222, member A | 2.13 |
| ENSMUSG00000107796 | Gm44068 | predicted gene, 44068 | 2.13 |
| ENSMUSG00000001473 | Tubb6 | tubulin, beta 6 class V | 2.12 |
| ENSMUSG00000045319 | Proser2 | proline and serine rich 2 | 2.12 |
| ENSMUSG00000048058 | Ldlrad3 | low density lipoprotein receptor class A domain containing 3 | 2.12 |
| ENSMUSG00000060671 | Atp8b2 | ATPase, class I, type 8B, member 2 | 2.12 |
| ENSMUSG00000033967 | Rnf225 | ring finger protein 225 | 2.12 |
| ENSMUSG00000042524 | Sun2 | Sad1 and UNC84 domain containing 2 | 2.11 |
| ENSMUSG00000020882 | Cacnb1 | calcium channel, voltage-dependent, beta 1 subunit | 2.11 |
| ENSMUSG00000032013 | Trim29 | tripartite motif-containing 29 | 2.11 |
| ENSMUSG00000028780 | Sema3c | sema domain, immunoglobulin domain (Ig), short basic domain, secreted, (semaphorin) 3C | 2.10 |
| ENSMUSG00000000320 | Alox12 | arachidonate 12-lipoxygenase | 2.10 |
| ENSMUSG00000029094 | Afap1 | actin filament associated protein 1 | 2.09 |
| ENSMUSG00000032643 | Fhl3 | four and a half LIM domains 3 | 2.09 |
| ENSMUSG00000023885 | Thbs2 | thrombospondin 2 | 2.08 |
| ENSMUSG00000025582 | Nptx1 | neuronal pentraxin 1 | 2.08 |

| | | | |
|---|---|---|---|
| ENSMUSG00000022754 | Tmem45a | transmembrane protein 45a | 2.08 |
| ENSMUSG00000037239 | Spred3 | sprouty-related, EVH1 domain containing 3 | 2.08 |
| ENSMUSG00000086537 | Nespas | neuroendocrine secretory protein antisense | 2.08 |
| ENSMUSG00000039328 | Rnf122 | ring finger protein 122 | 2.07 |
| ENSMUSG00000017723 | Wfdc2 | WAP four-disulfide core domain 2 | 2.07 |
| ENSMUSG00000009378 | Slc16a12 | solute carrier family 16 (monocarboxylic acid transporters), member 12 | 2.06 |
| ENSMUSG00000002058 | Unc119 | unc-119 lipid binding chaperone | 2.06 |
| ENSMUSG00000044034 | Npb | neuropeptide B | 2.06 |
| ENSMUSG00000085514 | Bcas3os2 | breast carcinoma amplified sequence 3 opposite strand 2 | 2.06 |
| ENSMUSG00000102758 | Naaladl2 | N-acetylated alpha-linked acidic dipeptidase-like 2 | 2.05 |
| ENSMUSG00000023905 | Tnfrsf12a | tumor necrosis factor receptor superfamily, member 12a | 2.05 |
| ENSMUSG00000017667 | Zfp334 | zinc finger protein 334 | 2.05 |
| ENSMUSG00000026405 | C4bp | complement component 4 binding protein | 2.05 |
| ENSMUSG00000042268 | Slc26a9 | solute carrier family 26, member 9 | 2.05 |
| ENSMUSG00000059493 | Nhs | NHS actin remodeling regulator | 2.04 |
| ENSMUSG00000036412 | Arsi | arylsulfatase i | 2.04 |
| ENSMUSG00000021756 | Il6st | interleukin 6 signal transducer | 2.03 |
| ENSMUSG00000025608 | Podxl | podocalyxin-like | 2.03 |
| ENSMUSG00000110218 | Gm20219 | predicted gene, 20219 | 2.03 |
| ENSMUSG00000004709 | Cd244a | CD244 molecule A | 2.03 |
| ENSMUSG00000068270 | Shroom4 | shroom family member 4 | 2.03 |
| ENSMUSG00000037946 | Fgd3 | FYVE, RhoGEF and PH domain containing 3 | 2.02 |
| ENSMUSG00000031074 | Fgf3 | fibroblast growth factor 3 | 2.02 |
| ENSMUSG00000068617 | Efcab1 | EF-hand calcium binding domain 1 | 2.02 |
| ENSMUSG00000037736 | Limch1 | LIM and calponin homology domains 1 | 2.01 |
| ENSMUSG00000028600 | Podn | podocan | 2.01 |
| ENSMUSG00000003541 | Ier3 | immediate early response 3 | 2.01 |
| ENSMUSG00000019817 | Plagl1 | pleiomorphic adenoma gene-like 1 | 2.01 |
| ENSMUSG00000073409 | H2-Q6 | histocompatibility 2, Q region locus 6 | 2.01 |
| ENSMUSG00000037892 | Pcdh18 | protocadherin 18 | 2.01 |
| ENSMUSG00000046470 | Sox18 | SRY (sex determining region Y)-box 18 | 2.01 |
| ENSMUSG00000071855 | Ccdc112 | coiled-coil domain containing 112 | 2.00 |
| ENSMUSG00000022425 | Enpp2 | ectonucleotide pyrophosphatase/phosphodiesterase 2 | 2.00 |
| ENSMUSG00000044022 | Pcdhb21 | protocadherin beta 21 | 2.00 |
| ENSMUSG00000051790 | Nlgn2 | neuroligin 2 | 1.99 |
| ENSMUSG00000024270 | Slc39a6 | solute carrier family 39 (metal ion transporter), member 6 | 1.99 |
| ENSMUSG00000007207 | Stx1a | syntaxin 1A (brain) | 1.99 |
| ENSMUSG00000089762 | Ier5l | immediate early response 5-like | 1.99 |
| ENSMUSG00000039239 | Tgfb2 | transforming growth factor, beta 2 | 1.99 |
| ENSMUSG00000028949 | Smarcd3 | SWI/SNF related, matrix associated, actin dependent regulator of chromatin, subfamily d, member 3 | 1.99 |
| ENSMUSG00000021477 | Ctsl | cathepsin L | 1.98 |
| ENSMUSG00000035901 | Dennd5a | DENN/MADD domain containing 5A | 1.98 |
| ENSMUSG00000056758 | Hmga2 | high mobility group AT-hook 2 | 1.98 |
| ENSMUSG00000029312 | Klhl8 | kelch-like 8 | 1.98 |
| ENSMUSG00000044350 | Lacc1 | laccase domain containing 1 | 1.98 |
| ENSMUSG00000059336 | Slc14a1 | solute carrier family 14 (urea transporter), member 1 | 1.98 |
| ENSMUSG00000037060 | Cavin3 | caveolae associated 3 | 1.98 |
| ENSMUSG00000027381 | Bcl2l11 | BCL2-like 11 (apoptosis facilitator) | 1.97 |
| ENSMUSG00000000555 | Itga5 | integrin alpha 5 (fibronectin receptor alpha) | 1.97 |
| ENSMUSG00000026411 | Tmem9 | transmembrane protein 9 | 1.97 |
| ENSMUSG00000046546 | Fam43a | family with sequence similarity 43, member A | 1.97 |
| ENSMUSG00000043895 | S1pr2 | sphingosine-1-phosphate receptor 2 | 1.97 |
| ENSMUSG00000038146 | Notch3 | notch 3 | 1.97 |
| ENSMUSG00000026121 | Sema4c | sema domain, immunoglobulin domain (Ig), transmembrane domain (TM) and short cytoplasmic dom | 1.96 |
| ENSMUSG00000091183 | Gm5141 | predicted gene 5141 | 1.96 |
| ENSMUSG00000037846 | Rtkn2 | rhotekin 2 | 1.95 |
| ENSMUSG00000024558 | Mapk4 | mitogen-activated protein kinase 4 | 1.94 |
| ENSMUSG00000046329 | Slc25a23 | solute carrier family 25 (mitochondrial carrier; phosphate carrier), member 23 | 1.94 |
| ENSMUSG00000038775 | Vill | villin-like | 1.94 |
| ENSMUSG00000002900 | Lamb1 | laminin B1 | 1.94 |
| ENSMUSG00000031066 | Usp11 | ubiquitin specific peptidase 11 | 1.94 |
| ENSMUSG00000030352 | Tspan9 | tetraspanin 9 | 1.94 |
| ENSMUSG00000072964 | Bhlhb9 | basic helix-loop-helix domain containing, class B9 | 1.93 |
| ENSMUSG00000037405 | Icam1 | intercellular adhesion molecule 1 | 1.93 |
| ENSMUSG00000055745 | Rtl6 | retrotransposon Gag like 6 | 1.93 |
| ENSMUSG00000021253 | Tgfb3 | transforming growth factor, beta 3 | 1.93 |
| ENSMUSG00000057895 | Zfp105 | zinc finger protein 105 | 1.93 |
| ENSMUSG00000044548 | Dact1 | dishevelled-binding antagonist of beta-catenin 1 | 1.93 |
| ENSMUSG00000040990 | Sh3kbp1 | SH3-domain kinase binding protein 1 | 1.92 |
| ENSMUSG00000071637 | Cebpd | CCAAT/enhancer binding protein (C/EBP), delta | 1.92 |
| ENSMUSG00000037447 | Arid5a | AT rich interactive domain 5A (MRF1-like) | 1.92 |

| | | | |
|---|---|---|---|
| ENSMUSG00000007039 | Ddah2 | dimethylarginine dimethylaminohydrolase 2 | 1.92 |
| ENSMUSG00000062012 | Zfp13 | zinc finger protein 13 | 1.92 |
| ENSMUSG00000060548 | Tnfrsf19 | tumor necrosis factor receptor superfamily, member 19 | 1.92 |
| ENSMUSG00000043448 | Gjc2 | gap junction protein, gamma 2 | 1.92 |
| ENSMUSG00000002059 | Rab34 | RAB34, member RAS oncogene family | 1.92 |
| ENSMUSG00000045287 | Rtn4rl1 | reticulon 4 receptor-like 1 | 1.91 |
| ENSMUSG00000015243 | Abca1 | ATP-binding cassette, sub-family A (ABC1), member 1 | 1.90 |
| ENSMUSG00000026223 | Itm2c | integral membrane protein 2C | 1.90 |
| ENSMUSG00000031530 | Dusp4 | dual specificity phosphatase 4 | 1.90 |
| ENSMUSG00000003363 | Pld3 | phospholipase D family, member 3 | 1.90 |
| ENSMUSG00000019732 | Calr3 | calreticulin 3 | 1.90 |
| ENSMUSG00000058908 | Pla2g2a | phospholipase A2, group IIA (platelets, synovial fluid) | 1.89 |
| ENSMUSG00000039103 | Nexn | nexilin | 1.89 |
| ENSMUSG00000031734 | Irx3 | Iroquois related homeobox 3 | 1.89 |
| ENSMUSG00000028358 | Zfp618 | zinc finger protein 618 | 1.88 |
| ENSMUSG00000041202 | Pla2g2d | phospholipase A2, group IID | 1.88 |
| ENSMUSG00000113603 | Gm47004 | predicted gene, 47004 | 1.88 |
| ENSMUSG00000026478 | Lamc1 | laminin, gamma 1 | 1.87 |
| ENSMUSG00000022178 | Ajuba | ajuba LIM protein | 1.87 |
| ENSMUSG00000090386 | Mir99ahg | Mir99a and Mirlet7c-1 host gene (non-protein coding) | 1.87 |
| ENSMUSG00000044562 | Rasip1 | Ras interacting protein 1 | 1.87 |
| ENSMUSG00000015709 | Arnt2 | aryl hydrocarbon receptor nuclear translocator 2 | 1.86 |
| ENSMUSG00000020620 | Abca8b | ATP-binding cassette, sub-family A (ABC1), member 8b | 1.86 |
| ENSMUSG00000053141 | Ptprt | protein tyrosine phosphatase, receptor type, T | 1.85 |
| ENSMUSG00000031596 | Slc7a2 | solute carrier family 7 (cationic amino acid transporter, y+ system), member 2 | 1.85 |
| ENSMUSG00000007989 | Fzd3 | frizzled class receptor 3 | 1.85 |
| ENSMUSG00000019235 | Rps6kl1 | ribosomal protein S6 kinase-like 1 | 1.85 |
| ENSMUSG00000028108 | Ecm1 | extracellular matrix protein 1 | 1.85 |
| ENSMUSG00000030409 | Dmpk | dystrophia myotonica-protein kinase | 1.84 |
| ENSMUSG00000026944 | Abca2 | ATP-binding cassette, sub-family A (ABC1), member 2 | 1.84 |
| ENSMUSG00000073987 | Ggh | gamma-glutamyl hydrolase | 1.84 |
| ENSMUSG00000039481 | Nrtn | neurturin | 1.84 |
| ENSMUSG00000069170 | Adgrv1 | adhesion G protein-coupled receptor V1 | 1.84 |
| ENSMUSG00000052212 | Cd177 | CD177 antigen | 1.83 |
| ENSMUSG00000025921 | Rdh10 | retinol dehydrogenase 10 (all-trans) | 1.83 |
| ENSMUSG00000028763 | Hspg2 | perlecan (heparan sulfate proteoglycan 2) | 1.83 |
| ENSMUSG00000001521 | Tulp3 | tubby-like protein 3 | 1.83 |
| ENSMUSG00000022415 | Syngr1 | synaptogyrin 1 | 1.83 |
| ENSMUSG00000019986 | Ahi1 | Abelson helper integration site 1 | 1.83 |
| ENSMUSG00000104063 | Pcdhgb7 | protocadherin gamma subfamily B, 7 | 1.83 |
| ENSMUSG00000038545 | Cul7 | cullin 7 | 1.82 |
| ENSMUSG00000037071 | Scd1 | stearoyl-Coenzyme A desaturase 1 | 1.82 |
| ENSMUSG00000062661 | Ncs1 | neuronal calcium sensor 1 | 1.82 |
| ENSMUSG00000091941 | Gm7399 | predicted gene 7399 | 1.82 |
| ENSMUSG00000028173 | Wls | wntless WNT ligand secretion mediator | 1.80 |
| ENSMUSG00000053897 | Slc39a8 | solute carrier family 39 (metal ion transporter), member 8 | 1.80 |
| ENSMUSG00000046572 | Zfp518b | zinc finger protein 518B | 1.80 |
| ENSMUSG00000020646 | Mboat2 | membrane bound O-acyltransferase domain containing 2 | 1.80 |
| ENSMUSG00000032698 | Lmo2 | LIM domain only 2 | 1.80 |
| ENSMUSG00000026637 | Traf5 | TNF receptor-associated factor 5 | 1.80 |
| ENSMUSG00000027712 | Anxa5 | annexin A5 | 1.79 |
| ENSMUSG00000031875 | Cmtm3 | CKLF-like MARVEL transmembrane domain containing 3 | 1.79 |
| ENSMUSG00000046589 | Lrrc8e | leucine rich repeat containing 8 family, member E | 1.79 |
| ENSMUSG00000026065 | Slc9a4 | solute carrier family 9 (sodium/hydrogen exchanger), member 4 | 1.79 |
| ENSMUSG00000040723 | Rcsd1 | RCSD domain containing 1 | 1.79 |
| ENSMUSG00000076431 | Sox4 | SRY (sex determining region Y)-box 4 | 1.78 |
| ENSMUSG00000028357 | Kif12 | kinesin family member 12 | 1.78 |
| ENSMUSG00000021373 | Cap2 | CAP, adenylate cyclase-associated protein, 2 (yeast) | 1.78 |
| ENSMUSG00000066607 | Insyn1 | inhibitory synaptic factor 1 | 1.78 |
| ENSMUSG00000039238 | Zfp750 | zinc finger protein 750 | 1.78 |
| ENSMUSG00000063018 | 2010204K13Rik | RIKEN cDNA 2010204K13 gene | 1.78 |
| ENSMUSG00000027858 | Tspan2 | tetraspanin 2 | 1.78 |
| ENSMUSG00000027860 | Vangl1 | VANGL planar cell polarity 1 | 1.77 |
| ENSMUSG00000033730 | Egr3 | early growth response 3 | 1.77 |
| ENSMUSG00000112014 | Gm48435 | predicted gene, 48435 | 1.77 |
| ENSMUSG00000021340 | Gpld1 | glycosylphosphatidylinositol specific phospholipase D1 | 1.76 |
| ENSMUSG00000025372 | Baiap2 | brain-specific angiogenesis inhibitor 1-associated protein 2 | 1.76 |
| ENSMUSG00000053080 | 2700081O15Rik | RIKEN cDNA 2700081O15 gene | 1.76 |
| ENSMUSG00000012428 | Steap4 | STEAP family member 4 | 1.76 |
| ENSMUSG00000050201 | Otop2 | otopetrin 2 | 1.76 |
| ENSMUSG00000031996 | Aplp2 | amyloid beta (A4) precursor-like protein 2 | 1.75 |

| | | | |
|---|---|---|---|
| ENSMUSG00000016382 | Pls3 | plastin 3 (T-isoform) | 1.75 |
| ENSMUSG00000038578 | Susd1 | sushi domain containing 1 | 1.75 |
| ENSMUSG00000117123 | AL805899.1 | novel transcript | 1.75 |
| ENSMUSG00000073158 | 9030624G23Rik | RIKEN cDNA 9030624G23 gene | 1.75 |
| ENSMUSG00000015053 | Gata2 | GATA binding protein 2 | 1.75 |
| ENSMUSG00000022996 | Wnt10b | wingless-type MMTV integration site family, member 10B | 1.75 |
| ENSMUSG00000030223 | Ptpro | protein tyrosine phosphatase, receptor type, O | 1.74 |
| ENSMUSG00000028347 | Tmeff1 | transmembrane protein with EGF-like and two follistatin-like domains 1 | 1.74 |
| ENSMUSG00000024070 | Prkd3 | protein kinase D3 | 1.74 |
| ENSMUSG00000024177 | Nme4 | NME/NM23 nucleoside diphosphate kinase 4 | 1.74 |
| ENSMUSG00000020037 | Rfx4 | regulatory factor X, 4 (influences HLA class II expression) | 1.74 |
| ENSMUSG00000098068 | Gm7909 | predicted gene 7909 | 1.74 |
| ENSMUSG00000051510 | Mafg | v-maf musculoaponeurotic fibrosarcoma oncogene family, protein G (avian) | 1.73 |
| ENSMUSG00000057092 | Fxyd3 | FXYD domain-containing ion transport regulator 3 | 1.73 |
| ENSMUSG00000038980 | Rbbp8nl | RBBP8 N-terminal like | 1.73 |
| ENSMUSG00000032607 | Amt | aminomethyltransferase | 1.73 |
| ENSMUSG00000024968 | Rcor2 | REST corepressor 2 | 1.73 |
| ENSMUSG00000006931 | P3h4 | prolyl 3-hydroxylase family member 4 (non-enzymatic) | 1.73 |
| ENSMUSG00000031736 | Crnde | colorectal neoplasia differentially expressed (non-protein coding) | 1.73 |
| ENSMUSG00000028541 | B4galt2 | UDP-Gal:betaGlcNAc beta 1,4- galactosyltransferase, polypeptide 2 | 1.73 |
| ENSMUSG00000034177 | Rnf43 | ring finger protein 43 | 1.72 |
| ENSMUSG00000047501 | Cldn4 | claudin 4 | 1.71 |
| ENSMUSG00000028464 | Tpm2 | tropomyosin 2, beta | 1.71 |
| ENSMUSG00000033032 | Afap1l1 | actin filament associated protein 1-like 1 | 1.70 |
| ENSMUSG00000042099 | Kank3 | KN motif and ankyrin repeat domains 3 | 1.70 |
| ENSMUSG00000105990 | Gm43307 | predicted gene 43307 | 1.70 |
| ENSMUSG00000031430 | Vsig1 | V-set and immunoglobulin domain containing 1 | 1.70 |
| ENSMUSG00000022505 | Emp2 | epithelial membrane protein 2 | 1.69 |
| ENSMUSG00000014592 | Camta1 | calmodulin binding transcription activator 1 | 1.69 |
| ENSMUSG00000024535 | Snx24 | sorting nexing 24 | 1.69 |
| ENSMUSG00000038594 | Cep85l | centrosomal protein 85-like | 1.69 |
| ENSMUSG00000112084 | Gm10773 | predicted gene 10773 | 1.69 |
| ENSMUSG00000004110 | Cacna1e | calcium channel, voltage-dependent, R type, alpha 1E subunit | 1.69 |
| ENSMUSG00000060261 | Gtf2i | general transcription factor II I | 1.68 |
| ENSMUSG00000000142 | Axin2 | axin 2 | 1.68 |
| ENSMUSG00000025017 | Pik3ap1 | phosphoinositide-3-kinase adaptor protein 1 | 1.68 |
| ENSMUSG00000048285 | Frmd6 | FERM domain containing 6 | 1.68 |
| ENSMUSG00000037972 | Snn | stannin | 1.68 |
| ENSMUSG00000027737 | Slc7a11 | solute carrier family 7 (cationic amino acid transporter, y+ system), member 11 | 1.68 |
| ENSMUSG00000027346 | Gpcpd1 | glycerophosphocholine phosphodiesterase 1 | 1.67 |
| ENSMUSG00000019843 | Fyn | Fyn proto-oncogene | 1.67 |
| ENSMUSG00000034161 | Scx | scleraxis | 1.67 |
| ENSMUSG00000050821 | Fam131a | family with sequence similarity 131, member A | 1.67 |
| ENSMUSG00000029669 | Tspan12 | tetraspanin 12 | 1.66 |
| ENSMUSG00000038295 | Atg9b | autophagy related 9B | 1.66 |
| ENSMUSG00000054720 | Lrrc8c | leucine rich repeat containing 8 family, member C | 1.66 |
| ENSMUSG00000059540 | Tcea2 | transcription elongation factor A (SII), 2 | 1.65 |
| ENSMUSG00000030519 | Apba2 | amyloid beta (A4) precursor protein-binding, family A, member 2 | 1.65 |
| ENSMUSG00000080198 | Gm13573 | predicted gene 13573 | 1.65 |
| ENSMUSG00000087589 | D430040D24Rik | RIKEN cDNA D430040D24 gene | 1.65 |
| ENSMUSG00000000957 | Mmp14 | matrix metallopeptidase 14 (membrane-inserted) | 1.64 |
| ENSMUSG00000020205 | Phlda1 | pleckstrin homology like domain, family A, member 1 | 1.64 |
| ENSMUSG00000022475 | Hdac7 | histone deacetylase 7 | 1.64 |
| ENSMUSG00000017390 | Aldoc | aldolase C, fructose-bisphosphate | 1.64 |
| ENSMUSG00000076441 | Ass1 | argininosuccinate synthetase 1 | 1.64 |
| ENSMUSG00000055407 | Map6 | microtubule-associated protein 6 | 1.64 |
| ENSMUSG00000021539 | Lect2 | leukocyte cell-derived chemotaxin 2 | 1.64 |
| ENSMUSG00000031633 | Slc25a4 | solute carrier family 25 (mitochondrial carrier, adenine nucleotide translocator), member 4 | 1.63 |
| ENSMUSG00000031586 | Rbpms | RNA binding protein gene with multiple splicing | 1.63 |
| ENSMUSG00000032606 | Nicn1 | nicolin 1 | 1.63 |
| ENSMUSG00000049321 | Zfp2 | zinc finger protein 2 | 1.63 |
| ENSMUSG00000026458 | Ppfia4 | protein tyrosine phosphatase, receptor type, f polypeptide (PTPRF), interacting protein (liprin), alpha | 1.63 |
| ENSMUSG00000006777 | Krt23 | keratin 23 | 1.62 |
| ENSMUSG00000062937 | Mtap | methylthioadenosine phosphorylase | 1.62 |
| ENSMUSG00000024524 | Gnal | guanine nucleotide binding protein, alpha stimulating, olfactory type | 1.62 |
| ENSMUSG00000069874 | Irgm2 | immunity-related GTPase family M member 2 | 1.62 |
| ENSMUSG00000090965 | Gm17203 | predicted gene 17203 | 1.62 |
| ENSMUSG00000032232 | Cgnl1 | cingulin-like 1 | 1.61 |
| ENSMUSG00000109727 | Gm45464 | predicted gene 45464 | 1.61 |
| ENSMUSG00000001870 | Ltbp1 | latent transforming growth factor beta binding protein 1 | 1.61 |
| ENSMUSG00000079330 | Lemd1 | LEM domain containing 1 | 1.60 |

| | | | |
|---|---|---|---|
| ENSMUSG00000046743 | Fat4 | FAT atypical cadherin 4 | 1.60 |
| ENSMUSG00000090399 | Gm38399 | predicted gene, 38399 | 1.60 |
| ENSMUSG00000024109 | Nrxn1 | neurexin I | 1.60 |
| ENSMUSG00000091055 | Siglec15 | sialic acid binding Ig-like lectin 15 | 1.60 |
| ENSMUSG00000032849 | Abcc4 | ATP-binding cassette, sub-family C (CFTR/MRP), member 4 | 1.59 |
| ENSMUSG00000020453 | Patz1 | POZ (BTB) and AT hook containing zinc finger 1 | 1.59 |
| ENSMUSG00000022433 | Csnk1e | casein kinase 1, epsilon | 1.58 |
| ENSMUSG00000013033 | Adgrl1 | adhesion G protein-coupled receptor L1 | 1.58 |
| ENSMUSG00000046718 | Bst2 | bone marrow stromal cell antigen 2 | 1.58 |
| ENSMUSG00000000486 | Sept1 | septin 1 | 1.58 |
| ENSMUSG00000038738 | Shank1 | SH3 and multiple ankyrin repeat domains 1 | 1.58 |
| ENSMUSG00000058046 | 4933430I17Rik | RIKEN cDNA 4933430I17 gene | 1.58 |
| ENSMUSG00000044349 | Snhg11 | small nucleolar RNA host gene 11 | 1.58 |
| ENSMUSG00000027570 | Col9a3 | collagen, type IX, alpha 3 | 1.57 |
| ENSMUSG00000051735 | Rinl | Ras and Rab interactor-like | 1.57 |
| ENSMUSG00000027485 | Bpifb1 | BPI fold containing family B, member 1 | 1.57 |
| ENSMUSG00000036564 | Ndrg4 | N-myc downstream regulated gene 4 | 1.57 |
| ENSMUSG00000019256 | Ahr | aryl-hydrocarbon receptor | 1.56 |
| ENSMUSG00000082508 | Rpl15-ps4 | ribosomal protein L15, pseudogene 4 | 1.56 |
| ENSMUSG00000049521 | Cdc42ep1 | CDC42 effector protein (Rho GTPase binding) 1 | 1.55 |
| ENSMUSG00000006435 | Neurl1a | neuralized E3 ubiquitin protein ligase 1A | 1.55 |
| ENSMUSG00000020032 | Nuak1 | NUAK family, SNF1-like kinase, 1 | 1.55 |
| ENSMUSG00000103220 | Gm37728 | predicted gene, 37728 | 1.55 |
| ENSMUSG00000019841 | Rev3l | REV3 like, DNA directed polymerase zeta catalytic subunit | 1.54 |
| ENSMUSG00000035914 | Cd276 | CD276 antigen | 1.54 |
| ENSMUSG00000027784 | Ppm1l | protein phosphatase 1 (formerly 2C)-like | 1.54 |
| ENSMUSG00000008450 | Nutf2 | nuclear transport factor 2 | 1.54 |
| ENSMUSG00000018906 | P4ha2 | procollagen-proline, 2-oxoglutarate 4-dioxygenase (proline 4-hydroxylase), alpha II polypeptide | 1.54 |
| ENSMUSG00000110662 | Gm45722 | predicted gene 45722 | 1.54 |
| ENSMUSG00000022995 | Enah | ENAH actin regulator | 1.54 |
| ENSMUSG00000017493 | Igfbp4 | insulin-like growth factor binding protein 4 | 1.53 |
| ENSMUSG00000032300 | 1700017B05Rik | RIKEN cDNA 1700017B05 gene | 1.53 |
| ENSMUSG00000054169 | Ceacam10 | carcinoembryonic antigen-related cell adhesion molecule 10 | 1.53 |
| ENSMUSG00000072694 | 1500011B03Rik | RIKEN cDNA 1500011B03 gene | 1.53 |
| ENSMUSG00000050192 | Eif5a2 | eukaryotic translation initiation factor 5A2 | 1.53 |
| ENSMUSG00000001025 | S100a6 | S100 calcium binding protein A6 (calcyclin) | 1.52 |
| ENSMUSG00000016534 | Lamp2 | lysosomal-associated membrane protein 2 | 1.52 |
| ENSMUSG00000026383 | Epb41l5 | erythrocyte membrane protein band 4.1 like 5 | 1.52 |
| ENSMUSG00000051359 | Ncald | neurocalcin delta | 1.52 |
| ENSMUSG00000043068 | Fam89a | family with sequence similarity 89, member A | 1.52 |
| ENSMUSG00000041592 | Sdk2 | sidekick cell adhesion molecule 2 | 1.52 |
| ENSMUSG00000102278 | Gm37145 | predicted gene, 37145 | 1.52 |
| ENSMUSG00000038872 | Zfhx3 | zinc finger homeobox 3 | 1.51 |
| ENSMUSG00000016541 | Atxn10 | ataxin 10 | 1.51 |
| ENSMUSG00000017376 | Nlk | nemo like kinase | 1.51 |
| ENSMUSG00000090100 | Ttbk2 | tau tubulin kinase 2 | 1.51 |
| ENSMUSG00000028601 | Echdc2 | enoyl Coenzyme A hydratase domain containing 2 | 1.51 |
| ENSMUSG00000017897 | Eya2 | EYA transcriptional coactivator and phosphatase 2 | 1.51 |
| ENSMUSG00000031292 | Cdkl5 | cyclin-dependent kinase-like 5 | 1.51 |
| ENSMUSG00000047910 | Pcdhb16 | protocadherin beta 16 | 1.51 |
| ENSMUSG00000047407 | Tgif1 | TGFB-induced factor homeobox 1 | 1.50 |
| ENSMUSG00000039081 | Zfp503 | zinc finger protein 503 | 1.50 |
| ENSMUSG00000001248 | Gramd1a | GRAM domain containing 1A | 1.50 |
| ENSMUSG00000022780 | Meltf | melanotransferrin | 1.50 |
| ENSMUSG00000027676 | Ccdc39 | coiled-coil domain containing 39 | 1.50 |
| ENSMUSG00000013275 | Slc41a1 | solute carrier family 41, member 1 | 1.49 |
| ENSMUSG00000048696 | Mex3d | mex3 RNA binding family member D | 1.49 |
| ENSMUSG00000041025 | Iffo2 | intermediate filament family orphan 2 | 1.49 |
| ENSMUSG00000069114 | Zbtb10 | zinc finger and BTB domain containing 10 | 1.49 |
| ENSMUSG00000020546 | Stxbp4 | syntaxin binding protein 4 | 1.49 |
| ENSMUSG00000020253 | Ppm1m | protein phosphatase 1M | 1.49 |
| ENSMUSG00000096687 | Mfsd4b4 | major facilitator superfamily domain containing 4B4 | 1.49 |
| ENSMUSG00000031660 | Brd7 | bromodomain containing 7 | 1.48 |
| ENSMUSG00000045991 | Onecut2 | one cut domain, family member 2 | 1.48 |
| ENSMUSG00000059895 | Ptp4a3 | protein tyrosine phosphatase 4a3 | 1.48 |
| ENSMUSG00000022604 | Cep97 | centrosomal protein 97 | 1.48 |
| ENSMUSG00000039683 | Sdk1 | sidekick cell adhesion molecule 1 | 1.48 |
| ENSMUSG00000029093 | Sorcs2 | sortilin-related VPS10 domain containing receptor 2 | 1.47 |
| ENSMUSG00000027907 | S100a11 | S100 calcium binding protein A11 | 1.47 |
| ENSMUSG00000025577 | Cbx2 | chromobox 2 | 1.47 |
| ENSMUSG00000028621 | Cyb5rl | cytochrome b5 reductase-like | 1.47 |

| | | | |
|---|---|---|---|
| ENSMUSG00000020773 | Trim47 | tripartite motif-containing 47 | 1.47 |
| ENSMUSG00000010751 | Tnfrsf22 | tumor necrosis factor receptor superfamily, member 22 | 1.47 |
| ENSMUSG00000062151 | Unc13c | unc-13 homolog C | 1.47 |
| ENSMUSG00000043065 | Spice1 | spindle and centriole associated protein 1 | 1.46 |
| ENSMUSG00000020648 | Dus4l | dihydrouridine synthase 4-like (S. cerevisiae) | 1.46 |
| ENSMUSG00000021108 | Prkch | protein kinase C, eta | 1.46 |
| ENSMUSG00000071042 | Rasgrp3 | RAS, guanyl releasing protein 3 | 1.46 |
| ENSMUSG00000007891 | Ctsd | cathepsin D | 1.45 |
| ENSMUSG00000025268 | Maged2 | melanoma antigen, family D, 2 | 1.45 |
| ENSMUSG00000058486 | Wdr91 | WD repeat domain 91 | 1.45 |
| ENSMUSG00000002257 | Def6 | differentially expressed in FDCP 6 | 1.45 |
| ENSMUSG00000030643 | Rab30 | RAB30, member RAS oncogene family | 1.45 |
| ENSMUSG00000038132 | Rbm24 | RNA binding motif protein 24 | 1.45 |
| ENSMUSG00000052534 | Pbx1 | pre B cell leukemia homeobox 1 | 1.44 |
| ENSMUSG00000028184 | Adgrl2 | adhesion G protein-coupled receptor L2 | 1.44 |
| ENSMUSG00000030759 | Far1 | fatty acyl CoA reductase 1 | 1.44 |
| ENSMUSG00000039384 | Dusp10 | dual specificity phosphatase 10 | 1.44 |
| ENSMUSG00000032359 | Ctsh | cathepsin H | 1.44 |
| ENSMUSG00000055639 | Dach1 | dachshund family transcription factor 1 | 1.43 |
| ENSMUSG00000004032 | Gstm5 | glutathione S-transferase, mu 5 | 1.43 |
| ENSMUSG00000095098 | Ccdc85b | coiled-coil domain containing 85B | 1.43 |
| ENSMUSG00000024736 | Tmem132a | transmembrane protein 132A | 1.43 |
| ENSMUSG00000103088 | Pcdhgb6 | protocadherin gamma subfamily B, 6 | 1.43 |
| ENSMUSG00000028152 | Tspan5 | tetraspanin 5 | 1.42 |
| ENSMUSG00000039801 | Cplane1 | ciliogenesis and planar polarity effector 1 | 1.42 |
| ENSMUSG00000028542 | Slc6a9 | solute carrier family 6 (neurotransmitter transporter, glycine), member 9 | 1.42 |
| ENSMUSG00000023348 | Trip6 | thyroid hormone receptor interactor 6 | 1.42 |
| ENSMUSG00000030725 | Lipt2 | lipoyl(octanoyl) transferase 2 (putative) | 1.42 |
| ENSMUSG00000050295 | Foxc1 | forkhead box C1 | 1.41 |
| ENSMUSG00000042694 | Stn1 | STN1, CST complex subunit | 1.41 |
| ENSMUSG00000024774 | Ankrd22 | ankyrin repeat domain 22 | 1.41 |
| ENSMUSG00000031626 | Sorbs2 | sorbin and SH3 domain containing 2 | 1.40 |
| ENSMUSG00000037936 | Scarb1 | scavenger receptor class B, member 1 | 1.40 |
| ENSMUSG00000044927 | H1fx | H1 histone family, member X | 1.40 |
| ENSMUSG00000047146 | Tet1 | tet methylcytosine dioxygenase 1 | 1.40 |
| ENSMUSG00000034156 | Tspoap1 | TSPO associated protein 1 | 1.40 |
| ENSMUSG00000029392 | Rilpl1 | Rab interacting lysosomal protein-like 1 | 1.40 |
| ENSMUSG00000025145 | Lrrc45 | leucine rich repeat containing 45 | 1.39 |
| ENSMUSG00000037664 | Cdkn1c | cyclin-dependent kinase inhibitor 1C (P57) | 1.39 |
| ENSMUSG00000041886 | Macc1 | metastasis associated in colon cancer 1 | 1.39 |
| ENSMUSG00000034854 | Mfsd12 | major facilitator superfamily domain containing 12 | 1.39 |
| ENSMUSG00000024013 | Fgd2 | FYVE, RhoGEF and PH domain containing 2 | 1.39 |
| ENSMUSG00000022055 | Nefl | neurofilament, light polypeptide | 1.39 |
| ENSMUSG00000008090 | Fgfrl1 | fibroblast growth factor receptor-like 1 | 1.38 |
| ENSMUSG00000006519 | Cyba | cytochrome b-245, alpha polypeptide | 1.38 |
| ENSMUSG00000009418 | Nav1 | neuron navigator 1 | 1.38 |
| ENSMUSG00000020650 | Bcap29 | B cell receptor associated protein 29 | 1.38 |
| ENSMUSG00000033149 | Phldb2 | pleckstrin homology like domain, family B, member 2 | 1.38 |
| ENSMUSG00000021792 | Fam213a | family with sequence similarity 213, member A | 1.38 |
| ENSMUSG00000038797 | Zscan2 | zinc finger and SCAN domain containing 2 | 1.38 |
| ENSMUSG00000027378 | Nphp1 | nephronophthisis 1 (juvenile) homolog (human) | 1.38 |
| ENSMUSG00000068758 | Il3ra | interleukin 3 receptor, alpha chain | 1.38 |
| ENSMUSG00000000202 | Btbd17 | BTB (POZ) domain containing 17 | 1.38 |
| ENSMUSG00000097787 | 2700046G09Rik | RIKEN cDNA 2700046G09 gene | 1.38 |
| ENSMUSG00000027997 | Casp6 | caspase 6 | 1.37 |
| ENSMUSG00000037656 | Slc20a2 | solute carrier family 20, member 2 | 1.37 |
| ENSMUSG00000040446 | Rprd1a | regulation of nuclear pre-mRNA domain containing 1A | 1.37 |
| ENSMUSG00000032470 | Mras | muscle and microspikes RAS | 1.37 |
| ENSMUSG00000038776 | Ephx1 | epoxide hydrolase 1, microsomal | 1.37 |
| ENSMUSG00000028573 | Fggy | FGGY carbohydrate kinase domain containing | 1.37 |
| ENSMUSG00000028626 | Col9a2 | collagen, type IX, alpha 2 | 1.37 |
| ENSMUSG00000040327 | Cul9 | cullin 9 | 1.36 |
| ENSMUSG00000032609 | Klhdc8b | kelch domain containing 8B | 1.36 |
| ENSMUSG00000025223 | Ldb1 | LIM domain binding 1 | 1.35 |
| ENSMUSG00000021482 | Aaed1 | AhpC/TSA antioxidant enzyme domain containing 1 | 1.35 |
| ENSMUSG00000033545 | Znrf1 | zinc and ring finger 1 | 1.35 |
| ENSMUSG00000085972 | 1110028F11Rik | RIKEN cDNA 1110028F11 gene | 1.35 |
| ENSMUSG00000020388 | Pdlim4 | PDZ and LIM domain 4 | 1.35 |
| ENSMUSG00000035615 | Frmpd1 | FERM and PDZ domain containing 1 | 1.35 |
| ENSMUSG00000079414 | Gm11110 | predicted gene 11110 | 1.35 |
| ENSMUSG00000029763 | Exoc4 | exocyst complex component 4 | 1.34 |

| | | | |
|---|---|---|---|
| ENSMUSG00000023088 | Abcc1 | ATP-binding cassette, sub-family C (CFTR/MRP), member 1 | 1.34 |
| ENSMUSG00000055485 | Soga1 | suppressor of glucose, autophagy associated 1 | 1.34 |
| ENSMUSG00000036097 | Slf2 | SMC5-SMC6 complex localization factor 2 | 1.33 |
| ENSMUSG00000025091 | Pnliprp2 | pancreatic lipase-related protein 2 | 1.33 |
| ENSMUSG00000044468 | Tent5c | terminal nucleotidyltransferase 5C | 1.33 |
| ENSMUSG00000048450 | Msx1 | msh homeobox 1 | 1.33 |
| ENSMUSG00000059970 | Hspa2 | heat shock protein 2 | 1.33 |
| ENSMUSG00000030284 | Creld1 | cysteine-rich with EGF-like domains 1 | 1.33 |
| ENSMUSG00000019831 | Wasf1 | WAS protein family, member 1 | 1.33 |
| ENSMUSG00000021013 | Ttc8 | tetratricopeptide repeat domain 8 | 1.33 |
| ENSMUSG00000038663 | Fsd2 | fibronectin type III and SPRY domain containing 2 | 1.33 |
| ENSMUSG00000058799 | Nap1l1 | nucleosome assembly protein 1-like 1 | 1.32 |
| ENSMUSG00000009585 | Apobec3 | apolipoprotein B mRNA editing enzyme, catalytic polypeptide 3 | 1.32 |
| ENSMUSG00000028838 | Extl1 | exostoses (multiple)-like 1 | 1.32 |
| ENSMUSG00000028318 | Polr1e | polymerase (RNA) I polypeptide E | 1.32 |
| ENSMUSG00000027189 | Trim44 | tripartite motif-containing 44 | 1.31 |
| ENSMUSG00000028266 | Lmo4 | LIM domain only 4 | 1.31 |
| ENSMUSG00000043439 | Epop | elongin BC and polycomb repressive complex 2 associated protein | 1.31 |
| ENSMUSG00000014907 | Naf1 | nuclear assembly factor 1 ribonucleoprotein | 1.31 |
| ENSMUSG00000045288 | Ush1g | USH1 protein network component sans | 1.31 |
| ENSMUSG00000047473 | Zfp30 | zinc finger protein 30 | 1.31 |
| ENSMUSG00000027333 | Smox | spermine oxidase | 1.30 |
| ENSMUSG00000026421 | Csrp1 | cysteine and glycine-rich protein 1 | 1.30 |
| ENSMUSG00000032348 | Gsta4 | glutathione S-transferase, alpha 4 | 1.30 |
| ENSMUSG00000035776 | Cd99l2 | CD99 antigen-like 2 | 1.30 |
| ENSMUSG00000049871 | Nlrc3 | NLR family, CARD domain containing 3 | 1.30 |
| ENSMUSG00000053931 | Cnn3 | calponin 3, acidic | 1.29 |
| ENSMUSG00000024896 | Minpp1 | multiple inositol polyphosphate histidine phosphatase 1 | 1.29 |
| ENSMUSG00000032440 | Tgfbr2 | transforming growth factor, beta receptor II | 1.29 |
| ENSMUSG00000022096 | Hr | hairless | 1.29 |
| ENSMUSG00000060591 | Ifitm2 | interferon induced transmembrane protein 2 | 1.29 |
| ENSMUSG00000020376 | Rnf130 | ring finger protein 130 | 1.29 |
| ENSMUSG00000025969 | Nrp2 | neuropilin 2 | 1.29 |
| ENSMUSG00000034771 | Tle2 | transducin-like enhancer of split 2 | 1.29 |
| ENSMUSG00000003190 | Bcl2l12 | BCL2-like 12 (proline rich) | 1.29 |
| ENSMUSG00000008734 | Gprc5b | G protein-coupled receptor, family C, group 5, member B | 1.29 |
| ENSMUSG00000021720 | Rnf180 | ring finger protein 180 | 1.28 |
| ENSMUSG00000004665 | Cnn2 | calponin 2 | 1.28 |
| ENSMUSG00000068245 | Phf11d | PHD finger protein 11D | 1.28 |
| ENSMUSG00000056486 | Chn1 | chimerin 1 | 1.28 |
| ENSMUSG00000031342 | Gpm6b | glycoprotein m6b | 1.28 |
| ENSMUSG00000059248 | Sept9 | septin 9 | 1.27 |
| ENSMUSG00000018378 | Cuedc1 | CUE domain containing 1 | 1.27 |
| ENSMUSG00000052852 | Reep1 | receptor accessory protein 1 | 1.27 |
| ENSMUSG00000017314 | Mpp2 | membrane protein, palmitoylated 2 (MAGUK p55 subfamily member 2) | 1.27 |
| ENSMUSG00000097062 | Gm17586 | predicted gene, 17586 | 1.27 |
| ENSMUSG00000084941 | Gm11944 | predicted gene 11944 | 1.27 |
| ENSMUSG00000032238 | Rora | RAR-related orphan receptor alpha | 1.27 |
| ENSMUSG00000034118 | Tpst1 | protein-tyrosine sulfotransferase 1 | 1.27 |
| ENSMUSG00000035547 | Capn5 | calpain 5 | 1.26 |
| ENSMUSG00000022390 | Zc3h7b | zinc finger CCCH type containing 7B | 1.26 |
| ENSMUSG00000028832 | Stmn1 | stathmin 1 | 1.26 |
| ENSMUSG00000034586 | Hid1 | HID1 domain containing | 1.26 |
| ENSMUSG00000026335 | Pam | peptidylglycine alpha-amidating monooxygenase | 1.26 |
| ENSMUSG00000033319 | Fem1c | fem-1 homolog c (C.elegans) | 1.26 |
| ENSMUSG00000068740 | Celsr2 | cadherin, EGF LAG seven-pass G-type receptor 2 | 1.26 |
| ENSMUSG00000061981 | Flot2 | flotillin 2 | 1.26 |
| ENSMUSG00000025212 | Sfxn3 | sideroflexin 3 | 1.26 |
| ENSMUSG00000023495 | Pcbp4 | poly(rC) binding protein 4 | 1.26 |
| ENSMUSG00000041836 | Ptpre | protein tyrosine phosphatase, receptor type, E | 1.25 |
| ENSMUSG00000037287 | Tbcel | tubulin folding cofactor E-like | 1.25 |
| ENSMUSG00000020776 | Fbf1 | Fas (TNFRSF6) binding factor 1 | 1.25 |
| ENSMUSG00000037499 | Nenf | neuron derived neurotrophic factor | 1.25 |
| ENSMUSG00000033039 | Micall1 | microtubule associated monooxygenase, calponin and LIM domain containing -like 1 | 1.24 |
| ENSMUSG00000071662 | Polr2g | polymerase (RNA) II (DNA directed) polypeptide G | 1.24 |
| ENSMUSG00000041135 | Ripk2 | receptor (TNFRSF)-interacting serine-threonine kinase 2 | 1.24 |
| ENSMUSG00000021611 | Tert | telomerase reverse transcriptase | 1.24 |
| ENSMUSG00000025887 | Casp12 | caspase 12 | 1.24 |
| ENSMUSG00000044716 | Dok7 | docking protein 7 | 1.24 |
| ENSMUSG00000013698 | Pea15a | phosphoprotein enriched in astrocytes 15A | 1.23 |
| ENSMUSG00000020536 | Llgl1 | LLGL1 scribble cell polarity complex component | 1.23 |

| | | | |
|---|---|---|---|
| ENSMUSG00000025144 | Cenpx | centromere protein X | 1.23 |
| ENSMUSG00000033900 | Map9 | microtubule-associated protein 9 | 1.23 |
| ENSMUSG00000037890 | Wdr19 | WD repeat domain 19 | 1.23 |
| ENSMUSG00000040187 | Arntl2 | aryl hydrocarbon receptor nuclear translocator-like 2 | 1.23 |
| ENSMUSG00000009406 | Elk1 | ELK1, member of ETS oncogene family | 1.23 |
| ENSMUSG00000038463 | Olfml2b | olfactomedin-like 2B | 1.23 |
| ENSMUSG00000030413 | Pglyrp1 | peptidoglycan recognition protein 1 | 1.22 |
| ENSMUSG00000028378 | Ptgr1 | prostaglandin reductase 1 | 1.22 |
| ENSMUSG00000020015 | Cdk17 | cyclin-dependent kinase 17 | 1.22 |
| ENSMUSG00000032184 | Lysmd2 | LysM, putative peptidoglycan-binding, domain containing 2 | 1.22 |
| ENSMUSG00000041540 | Sox5 | SRY (sex determining region Y)-box 5 | 1.22 |
| ENSMUSG00000027878 | Notch2 | notch 2 | 1.21 |
| ENSMUSG00000021506 | Pitx1 | paired-like homeodomain transcription factor 1 | 1.21 |
| ENSMUSG00000038007 | Acer2 | alkaline ceramidase 2 | 1.21 |
| ENSMUSG00000081683 | Fzd10 | frizzled class receptor 10 | 1.21 |
| ENSMUSG00000083012 | Fam220a | family with sequence similarity 220, member A | 1.21 |
| ENSMUSG00000020592 | Sdc1 | syndecan 1 | 1.20 |
| ENSMUSG00000029229 | Chic2 | cysteine-rich hydrophobic domain 2 | 1.20 |
| ENSMUSG00000046432 | Bex3 | brain expressed X-linked 3 | 1.20 |
| ENSMUSG00000023031 | Cela1 | chymotrypsin-like elastase family, member 1 | 1.20 |
| ENSMUSG00000034848 | Ttc21b | tetratricopeptide repeat domain 21B | 1.20 |
| ENSMUSG00000046058 | Eid2 | EP300 interacting inhibitor of differentiation 2 | 1.20 |
| ENSMUSG00000025153 | Fasn | fatty acid synthase | 1.19 |
| ENSMUSG00000045211 | Nudt18 | nudix (nucleoside diphosphate linked moiety X)-type motif 18 | 1.19 |
| ENSMUSG00000095325 | Zfp870 | zinc finger protein 870 | 1.19 |
| ENSMUSG00000047749 | Zc3hav1l | zinc finger CCCH-type, antiviral 1-like | 1.19 |
| ENSMUSG00000032657 | Fam189b | family with sequence similarity 189, member B | 1.19 |
| ENSMUSG00000028836 | Slc30a2 | solute carrier family 30 (zinc transporter), member 2 | 1.18 |
| ENSMUSG00000060510 | Zfp266 | zinc finger protein 266 | 1.18 |
| ENSMUSG00000023055 | Calcoco1 | calcium binding and coiled coil domain 1 | 1.18 |
| ENSMUSG00000049421 | Zfp260 | zinc finger protein 260 | 1.18 |
| ENSMUSG00000022678 | Nde1 | nudE neurodevelopment protein 1 | 1.18 |
| ENSMUSG00000020654 | Adcy3 | adenylate cyclase 3 | 1.18 |
| ENSMUSG00000072621 | Slfn10-ps | schlafen 10, pseudogene | 1.18 |
| ENSMUSG00000030315 | Vgll4 | vestigial like family member 4 | 1.17 |
| ENSMUSG00000109324 | Prmt1 | protein arginine N-methyltransferase 1 | 1.17 |
| ENSMUSG00000056648 | Hoxb8 | homeobox B8 | 1.17 |
| ENSMUSG00000021466 | Ptch1 | patched 1 | 1.17 |
| ENSMUSG00000000861 | Bcl11a | B cell CLL/lymphoma 11A (zinc finger protein) | 1.17 |
| ENSMUSG00000040711 | Sh3pxd2b | SH3 and PX domains 2B | 1.17 |
| ENSMUSG00000041096 | Tspyl2 | TSPY-like 2 | 1.17 |
| ENSMUSG00000039831 | Arhgap29 | Rho GTPase activating protein 29 | 1.17 |
| ENSMUSG00000050910 | Cdr2l | cerebellar degeneration-related protein 2-like | 1.17 |
| ENSMUSG00000039236 | Isg20 | interferon-stimulated protein | 1.17 |
| ENSMUSG00000039262 | Prrc2b | proline-rich coiled-coil 2B | 1.16 |
| ENSMUSG00000030729 | Pgm2l1 | phosphoglucomutase 2-like 1 | 1.16 |
| ENSMUSG00000026425 | Srgap2 | SLIT-ROBO Rho GTPase activating protein 2 | 1.16 |
| ENSMUSG00000025534 | Gusb | glucuronidase, beta | 1.16 |
| ENSMUSG00000045045 | Lrfn4 | leucine rich repeat and fibronectin type III domain containing 4 | 1.16 |
| ENSMUSG00000039976 | Tbc1d16 | TBC1 domain family, member 16 | 1.16 |
| ENSMUSG00000010080 | Epn3 | epsin 3 | 1.16 |
| ENSMUSG00000040543 | Pitpnm3 | PITPNM family member 3 | 1.16 |
| ENSMUSG00000035164 | Zc3h12c | zinc finger CCCH type containing 12C | 1.16 |
| ENSMUSG00000031207 | Msn | moesin | 1.16 |
| ENSMUSG00000053110 | Yap1 | yes-associated protein 1 | 1.15 |
| ENSMUSG00000049577 | Zfpm1 | zinc finger protein, multitype 1 | 1.15 |
| ENSMUSG00000034981 | Parm1 | prostate androgen-regulated mucin-like protein 1 | 1.15 |
| ENSMUSG00000025323 | Sp4 | trans-acting transcription factor 4 | 1.15 |
| ENSMUSG00000031502 | Col4a1 | collagen, type IV, alpha 1 | 1.15 |
| ENSMUSG00000033906 | Zdhhc15 | zinc finger, DHHC domain containing 15 | 1.14 |
| ENSMUSG00000034723 | Tmx4 | thioredoxin-related transmembrane protein 4 | 1.14 |
| ENSMUSG00000036006 | Ripor2 | RHO family interacting cell polarization regulator 2 | 1.14 |
| ENSMUSG00000031919 | Tmed6 | transmembrane p24 trafficking protein 6 | 1.14 |
| ENSMUSG00000022912 | Pros1 | protein S (alpha) | 1.14 |
| ENSMUSG00000064326 | Siva1 | SIVA1, apoptosis-inducing factor | 1.14 |
| ENSMUSG00000022895 | Ets2 | E26 avian leukemia oncogene 2, 3' domain | 1.13 |
| ENSMUSG00000019966 | Kitl | kit ligand | 1.13 |
| ENSMUSG00000048371 | Pdp2 | pyruvate dehyrogenase phosphatase catalytic subunit 2 | 1.13 |
| ENSMUSG00000040669 | Phc1 | polyhomeotic 1 | 1.13 |
| ENSMUSG00000019461 | Plscr3 | phospholipid scramblase 3 | 1.13 |
| ENSMUSG00000026727 | Rsu1 | Ras suppressor protein 1 | 1.13 |

| | | | |
|---|---|---|---|
| ENSMUSG00000027684 | Mecom | MDS1 and EVI1 complex locus | 1.12 |
| ENSMUSG00000084939 | Gm830 | predicted gene 830 | 1.12 |
| ENSMUSG00000018199 | Trove2 | TROVE domain family, member 2 | 1.12 |
| ENSMUSG00000057156 | Homez | homeodomain leucine zipper-encoding gene | 1.12 |
| ENSMUSG00000069631 | Strada | STE20-related kinase adaptor alpha | 1.12 |
| ENSMUSG00000038387 | Rras | related RAS viral (r-ras) oncogene | 1.12 |
| ENSMUSG00000049791 | Fzd4 | frizzled class receptor 4 | 1.12 |
| ENSMUSG00000002489 | Tiam1 | T cell lymphoma invasion and metastasis 1 | 1.11 |
| ENSMUSG00000022708 | Zbtb20 | zinc finger and BTB domain containing 20 | 1.11 |
| ENSMUSG00000059552 | Trp53 | transformation related protein 53 | 1.11 |
| ENSMUSG00000021262 | Evl | Ena-vasodilator stimulated phosphoprotein | 1.11 |
| ENSMUSG00000025986 | Slc39a10 | solute carrier family 39 (zinc transporter), member 10 | 1.11 |
| ENSMUSG00000000325 | Arvcf | armadillo repeat gene deleted in velocardiofacial syndrome | 1.11 |
| ENSMUSG00000033282 | Rpgrip1l | Rpgrip1-like | 1.11 |
| ENSMUSG00000021589 | Rhobtb3 | Rho-related BTB domain containing 3 | 1.11 |
| ENSMUSG00000021003 | Galc | galactosylceramidase | 1.11 |
| ENSMUSG00000027995 | Tlr2 | toll-like receptor 2 | 1.11 |
| ENSMUSG00000023947 | Nfkbie | nuclear factor of kappa light polypeptide gene enhancer in B cells inhibitor, epsilon | 1.11 |
| ENSMUSG00000115902 | AC113595.1 | novel transcript, antisense to Apobec3 | 1.11 |
| ENSMUSG00000020167 | Tcf3 | transcription factor 3 | 1.10 |
| ENSMUSG00000029246 | Ppat | phosphoribosyl pyrophosphate amidotransferase | 1.10 |
| ENSMUSG00000028245 | Nsmaf | neutral sphingomyelinase (N-SMase) activation associated factor | 1.09 |
| ENSMUSG00000040466 | Blvrb | biliverdin reductase B (flavin reductase (NADPH)) | 1.09 |
| ENSMUSG00000058672 | Tubb2a | tubulin, beta 2A class IIA | 1.09 |
| ENSMUSG00000044469 | Tnfaip8l1 | tumor necrosis factor, alpha-induced protein 8-like 1 | 1.09 |
| ENSMUSG00000089809 | Rasgef1b | RasGEF domain family, member 1B | 1.09 |
| ENSMUSG00000036052 | Dnajb5 | DnaJ heat shock protein family (Hsp40) member B5 | 1.09 |
| ENSMUSG00000028576 | Ift74 | intraflagellar transport 74 | 1.09 |
| ENSMUSG00000027490 | E2f1 | E2F transcription factor 1 | 1.09 |
| ENSMUSG00000040681 | Hmgn1 | high mobility group nucleosomal binding domain 1 | 1.08 |
| ENSMUSG00000041238 | Rbbp8 | retinoblastoma binding protein 8, endonuclease | 1.08 |
| ENSMUSG00000015217 | Hmgb3 | high mobility group box 3 | 1.08 |
| ENSMUSG00000020393 | Kremen1 | kringle containing transmembrane protein 1 | 1.08 |
| ENSMUSG00000046668 | Cxxc5 | CXXC finger 5 | 1.08 |
| ENSMUSG00000049939 | Lrrc4 | leucine rich repeat containing 4 | 1.08 |
| ENSMUSG00000032068 | Plet1 | placenta expressed transcript 1 | 1.08 |
| ENSMUSG00000049300 | Prmt6 | protein arginine N-methyltransferase 6 | 1.08 |
| ENSMUSG00000097431 | Gm26782 | predicted gene, 26782 | 1.08 |
| ENSMUSG00000048232 | Fbxo10 | F-box protein 10 | 1.08 |
| ENSMUSG00000035451 | Foxa1 | forkhead box A1 | 1.07 |
| ENSMUSG00000021068 | Nin | ninein | 1.07 |
| ENSMUSG00000041308 | Sntb2 | syntrophin, basic 2 | 1.07 |
| ENSMUSG00000070305 | Mpzl3 | myelin protein zero-like 3 | 1.07 |
| ENSMUSG00000048924 | Ccdc125 | coiled-coil domain containing 125 | 1.07 |
| ENSMUSG00000022442 | Ttll1 | tubulin tyrosine ligase-like 1 | 1.07 |
| ENSMUSG00000031799 | Tpm4 | tropomyosin 4 | 1.06 |
| ENSMUSG00000026796 | Fam129b | family with sequence similarity 129, member B | 1.06 |
| ENSMUSG00000022231 | Sema5a | sema domain, seven thrombospondin repeats (type 1 and type 1-like), transmembrane domain (TM) a | 1.06 |
| ENSMUSG00000052117 | D630039A03Rik | RIKEN cDNA D630039A03 gene | 1.06 |
| ENSMUSG00000006369 | Fbln1 | fibulin 1 | 1.06 |
| ENSMUSG00000029999 | Tgfa | transforming growth factor alpha | 1.06 |
| ENSMUSG00000040596 | Pogk | pogo transposable element with KRAB domain | 1.06 |
| ENSMUSG00000040717 | Il17rd | interleukin 17 receptor D | 1.06 |
| ENSMUSG00000018547 | Pip4k2b | phosphatidylinositol-5-phosphate 4-kinase, type II, beta | 1.06 |
| ENSMUSG00000055945 | Prr18 | proline rich 18 | 1.06 |
| ENSMUSG00000034064 | Poglut1 | protein O-glucosyltransferase 1 | 1.06 |
| ENSMUSG00000038205 | Prkab2 | protein kinase, AMP-activated, beta 2 non-catalytic subunit | 1.06 |
| ENSMUSG00000032481 | Smarcc1 | SWI/SNF related, matrix associated, actin dependent regulator of chromatin, subfamily c, member 1 | 1.05 |
| ENSMUSG00000042506 | Usp22 | ubiquitin specific peptidase 22 | 1.05 |
| ENSMUSG00000026509 | Capn2 | calpain 2 | 1.05 |
| ENSMUSG00000029055 | Plch2 | phospholipase C, eta 2 | 1.05 |
| ENSMUSG00000019433 | Gipc1 | GIPC PDZ domain containing family, member 1 | 1.05 |
| ENSMUSG00000058070 | Eml1 | echinoderm microtubule associated protein like 1 | 1.05 |
| ENSMUSG00000002308 | Cd320 | CD320 antigen | 1.05 |
| ENSMUSG00000074876 | Spata5l1 | spermatogenesis associated 5-like 1 | 1.05 |
| ENSMUSG00000009614 | Sardh | sarcosine dehydrogenase | 1.05 |
| ENSMUSG00000033965 | Slc16a2 | solute carrier family 16 (monocarboxylic acid transporters), member 2 | 1.05 |
| ENSMUSG00000057133 | Chd6 | chromodomain helicase DNA binding protein 6 | 1.04 |
| ENSMUSG00000030342 | Cd9 | CD9 antigen | 1.04 |
| ENSMUSG00000053580 | Tanc2 | tetratricopeptide repeat, ankyrin repeat and coiled-coil containing 2 | 1.04 |
| ENSMUSG00000025151 | Maged1 | melanoma antigen, family D, 1 | 1.04 |

| | | | |
|---|---|---|---|
| ENSMUSG00000005034 | Prkacb | protein kinase, cAMP dependent, catalytic, beta | 1.04 |
| ENSMUSG00000031360 | Ctps2 | cytidine 5'-triphosphate synthase 2 | 1.04 |
| ENSMUSG00000024501 | Dpysl3 | dihydropyrimidinase-like 3 | 1.04 |
| ENSMUSG00000046380 | Jrk | jerky | 1.04 |
| ENSMUSG00000036368 | Rmdn2 | regulator of microtubule dynamics 2 | 1.04 |
| ENSMUSG00000037706 | Cd81 | CD81 antigen | 1.03 |
| ENSMUSG00000022828 | Gtf2e1 | general transcription factor II E, polypeptide 1 (alpha subunit) | 1.03 |
| ENSMUSG00000040785 | Ttc3 | tetratricopeptide repeat domain 3 | 1.02 |
| ENSMUSG00000040274 | Cdk6 | cyclin-dependent kinase 6 | 1.02 |
| ENSMUSG00000028849 | Map7d1 | MAP7 domain containing 1 | 1.02 |
| ENSMUSG00000028044 | Cks1b | CDC28 protein kinase 1b | 1.02 |
| ENSMUSG00000032228 | Tcf12 | transcription factor 12 | 1.01 |
| ENSMUSG00000051391 | Ywhag | tyrosine 3-monooxygenase/tryptophan 5-monooxygenase activation protein, gamma polypeptide | 1.01 |
| ENSMUSG00000041688 | Amot | angiomotin | 1.01 |
| ENSMUSG00000050912 | Tmem123 | transmembrane protein 123 | 1.01 |
| ENSMUSG00000054520 | Sh3bp2 | SH3-domain binding protein 2 | 1.01 |
| ENSMUSG00000049532 | Sall2 | spalt like transcription factor 2 | 1.01 |
| ENSMUSG00000030980 | Knop1 | lysine rich nucleolar protein 1 | 1.00 |
| ENSMUSG00000002107 | Celf2 | CUGBP, Elav-like family member 2 | 1.00 |
| ENSMUSG00000018569 | Cldn7 | claudin 7 | -1.00 |
| ENSMUSG00000001089 | Luzp1 | leucine zipper protein 1 | -1.00 |
| ENSMUSG00000054277 | Arfgap3 | ADP-ribosylation factor GTPase activating protein 3 | -1.00 |
| ENSMUSG00000021948 | Prkcd | protein kinase C, delta | -1.00 |
| ENSMUSG00000019969 | Psen1 | presenilin 1 | -1.00 |
| ENSMUSG00000040188 | Scamp2 | secretory carrier membrane protein 2 | -1.00 |
| ENSMUSG00000041577 | Prelp | proline arginine-rich end leucine-rich repeat | -1.00 |
| ENSMUSG00000030499 | Kctd15 | potassium channel tetramerisation domain containing 15 | -1.00 |
| ENSMUSG00000037400 | Atp11b | ATPase, class VI, type 11B | -1.01 |
| ENSMUSG00000027957 | Slc35a3 | solute carrier family 35 (UDP-N-acetylglucosamine (UDP-GlcNAc) transporter), member 3 | -1.01 |
| ENSMUSG00000034435 | Tmem30b | transmembrane protein 30B | -1.01 |
| ENSMUSG00000024036 | Slc37a1 | solute carrier family 37 (glycerol-3-phosphate transporter), member 1 | -1.01 |
| ENSMUSG00000035064 | Eef2k | eukaryotic elongation factor-2 kinase | -1.01 |
| ENSMUSG00000053898 | Ech1 | enoyl coenzyme A hydratase 1, peroxisomal | -1.01 |
| ENSMUSG00000053886 | Sh2d4a | SH2 domain containing 4A | -1.01 |
| ENSMUSG00000033318 | Gstt2 | glutathione S-transferase, theta 2 | -1.01 |
| ENSMUSG00000041845 | Rhod | ras homolog family member D | -1.01 |
| ENSMUSG00000057137 | Tmem140 | transmembrane protein 140 | -1.01 |
| ENSMUSG00000023328 | Ache | acetylcholinesterase | -1.01 |
| ENSMUSG00000070780 | Rbm47 | RNA binding motif protein 47 | -1.02 |
| ENSMUSG00000040659 | Efhd2 | EF hand domain containing 2 | -1.02 |
| ENSMUSG00000019302 | Atp6v0a1 | ATPase, H+ transporting, lysosomal V0 subunit A1 | -1.02 |
| ENSMUSG00000074305 | Peak1 | pseudopodium-enriched atypical kinase 1 | -1.02 |
| ENSMUSG00000061313 | Ddhd2 | DDHD domain containing 2 | -1.02 |
| ENSMUSG00000039349 | C130074G19Rik | RIKEN cDNA C130074G19 gene | -1.02 |
| ENSMUSG00000015536 | Mocs2 | molybdenum cofactor synthesis 2 | -1.02 |
| ENSMUSG00000037709 | Fam13a | family with sequence similarity 13, member A | -1.02 |
| ENSMUSG00000028885 | Smpdl3b | sphingomyelin phosphodiesterase, acid-like 3B | -1.02 |
| ENSMUSG00000043085 | Tmem82 | transmembrane protein 82 | -1.02 |
| ENSMUSG00000000386 | Mx1 | MX dynamin-like GTPase 1 | -1.02 |
| ENSMUSG00000000631 | Myo18a | myosin XVIIIA | -1.03 |
| ENSMUSG00000024052 | Lpin2 | lipin 2 | -1.03 |
| ENSMUSG00000020451 | Limk2 | LIM motif-containing protein kinase 2 | -1.03 |
| ENSMUSG00000022973 | Synj1 | synaptojanin 1 | -1.03 |
| ENSMUSG00000020803 | Txndc17 | thioredoxin domain containing 17 | -1.03 |
| ENSMUSG00000055116 | Arntl | aryl hydrocarbon receptor nuclear translocator-like | -1.03 |
| ENSMUSG00000074653 | Lrrc31 | leucine rich repeat containing 31 | -1.03 |
| ENSMUSG00000002346 | Slc25a42 | solute carrier family 25, member 42 | -1.03 |
| ENSMUSG00000027422 | Rrbp1 | ribosome binding protein 1 | -1.04 |
| ENSMUSG00000046447 | Camk2n1 | calcium/calmodulin-dependent protein kinase II inhibitor 1 | -1.04 |
| ENSMUSG00000053965 | Pde5a | phosphodiesterase 5A, cGMP-specific | -1.04 |
| ENSMUSG00000027215 | Cd82 | CD82 antigen | -1.04 |
| ENSMUSG00000028413 | B4galt1 | UDP-Gal:betaGlcNAc beta 1,4- galactosyltransferase, polypeptide 1 | -1.04 |
| ENSMUSG00000032741 | Tpcn1 | two pore channel 1 | -1.04 |
| ENSMUSG00000074447 | Defa21 | defensin, alpha, 21 | -1.04 |
| ENSMUSG00000063206 | Defa34 | defensin, alpha, 34 | -1.04 |
| ENSMUSG00000026784 | Pdss1 | prenyl (solanesyl) diphosphate synthase, subunit 1 | -1.04 |
| ENSMUSG00000046794 | Ppp1r3b | protein phosphatase 1, regulatory subunit 3B | -1.04 |
| ENSMUSG00000039958 | Etfbkmt | electron transfer flavoprotein beta subunit lysine methyltransferase | -1.04 |
| ENSMUSG00000001300 | Efnb2 | ephrin B2 | -1.05 |
| ENSMUSG00000044786 | Zfp36 | zinc finger protein 36 | -1.05 |
| ENSMUSG00000046876 | Atxn1 | ataxin 1 | -1.05 |

| | | | |
|---|---|---|---|
| ENSMUSG00000034422 | Parp14 | poly (ADP-ribose) polymerase family, member 14 | -1.05 |
| ENSMUSG00000034330 | Plcg2 | phospholipase C, gamma 2 | -1.05 |
| ENSMUSG00000031853 | Map3k21 | mitogen-activated protein kinase kinase kinase 21 | -1.05 |
| ENSMUSG00000028480 | Glipr2 | GLI pathogenesis-related 2 | -1.05 |
| ENSMUSG00000032369 | Plscr1 | phospholipid scramblase 1 | -1.05 |
| ENSMUSG00000031639 | Tlr3 | toll-like receptor 3 | -1.05 |
| ENSMUSG00000056313 | Tcim | transcriptional and immune response regulator | -1.05 |
| ENSMUSG00000046352 | Gjb2 | gap junction protein, beta 2 | -1.05 |
| ENSMUSG00000038459 | Abhd17c | abhydrolase domain containing 17C | -1.06 |
| ENSMUSG00000035382 | Pcsk7 | proprotein convertase subtilisin/kexin type 7 | -1.06 |
| ENSMUSG00000041625 | Ggact | gamma-glutamylamine cyclotransferase | -1.06 |
| ENSMUSG00000027605 | Acss2 | acyl-CoA synthetase short-chain family member 2 | -1.06 |
| ENSMUSG00000022768 | Ccdc116 | coiled-coil domain containing 116 | -1.06 |
| ENSMUSG00000034271 | Jdp2 | Jun dimerization protein 2 | -1.06 |
| ENSMUSG00000042647 | Acad12 | acyl-Coenzyme A dehydrogenase family, member 12 | -1.06 |
| ENSMUSG00000038393 | Txnip | thioredoxin interacting protein | -1.07 |
| ENSMUSG00000021866 | Anxa11 | annexin A11 | -1.07 |
| ENSMUSG00000020134 | Peli1 | pellino 1 | -1.07 |
| ENSMUSG00000003545 | Fosb | FBJ osteosarcoma oncogene B | -1.07 |
| ENSMUSG00000034613 | Ppm1h | protein phosphatase 1H (PP2C domain containing) | -1.07 |
| ENSMUSG00000026360 | Rgs2 | regulator of G-protein signaling 2 | -1.07 |
| ENSMUSG00000034640 | Tiparp | TCDD-inducible poly(ADP-ribose) polymerase | -1.07 |
| ENSMUSG00000052776 | Oas1a | 2'-5' oligoadenylate synthetase 1A | -1.07 |
| ENSMUSG00000067199 | Frat1 | frequently rearranged in advanced T cell lymphomas | -1.07 |
| ENSMUSG00000046733 | Gprc5a | G protein-coupled receptor, family C, group 5, member A | -1.07 |
| ENSMUSG00000042750 | Bex2 | brain expressed X-linked 2 | -1.07 |
| ENSMUSG00000069833 | Ahnak | AHNAK nucleoprotein (desmoyokin) | -1.08 |
| ENSMUSG00000041870 | Ankrd13a | ankyrin repeat domain 13a | -1.08 |
| ENSMUSG00000038023 | Atp6v0a2 | ATPase, H+ transporting, lysosomal V0 subunit A2 | -1.08 |
| ENSMUSG00000026110 | Mgat4a | mannoside acetylglucosaminyltransferase 4, isoenzyme A | -1.08 |
| ENSMUSG00000022949 | Clic6 | chloride intracellular channel 6 | -1.08 |
| ENSMUSG00000090946 | Ccdc71l | coiled-coil domain containing 71 like | -1.08 |
| ENSMUSG00000066258 | Trim12a | tripartite motif-containing 12A | -1.08 |
| ENSMUSG00000031712 | Il15 | interleukin 15 | -1.08 |
| ENSMUSG00000057778 | Cyb5d2 | cytochrome b5 domain containing 2 | -1.08 |
| ENSMUSG00000103696 | Gm37531 | predicted gene, 37531 | -1.08 |
| ENSMUSG00000056185 | Snx32 | sorting nexin 32 | -1.08 |
| ENSMUSG00000032601 | Prkar2a | protein kinase, cAMP dependent regulatory, type II alpha | -1.09 |
| ENSMUSG00000030313 | Dennd5b | DENN/MADD domain containing 5B | -1.09 |
| ENSMUSG00000038967 | Pdk2 | pyruvate dehydrogenase kinase, isoenzyme 2 | -1.09 |
| ENSMUSG00000031156 | Slc35a2 | solute carrier family 35 (UDP-galactose transporter), member A2 | -1.09 |
| ENSMUSG00000021185 | Dglucy | D-glutamate cyclase | -1.09 |
| ENSMUSG00000024193 | Phf1 | PHD finger protein 1 | -1.09 |
| ENSMUSG00000026189 | Pecr | peroxisomal trans-2-enoyl-CoA reductase | -1.09 |
| ENSMUSG00000103711 | Tstd1 | thiosulfate sulfurtransferase (rhodanese)-like domain containing 1 | -1.09 |
| ENSMUSG00000109438 | Gm45073 | predicted gene 45073 | -1.09 |
| ENSMUSG00000056429 | Tgoln1 | trans-golgi network protein | -1.10 |
| ENSMUSG00000038859 | Baiap2l1 | BAI1-associated protein 2-like 1 | -1.10 |
| ENSMUSG00000023087 | Noct | nocturnin | -1.10 |
| ENSMUSG00000032306 | Mpi | mannose phosphate isomerase | -1.10 |
| ENSMUSG00000049422 | Chchd10 | coiled-coil-helix-coiled-coil-helix domain containing 10 | -1.10 |
| ENSMUSG00000048970 | C1galt1c1 | C1GALT1-specific chaperone 1 | -1.10 |
| ENSMUSG00000028538 | St3gal3 | ST3 beta-galactoside alpha-2,3-sialyltransferase 3 | -1.10 |
| ENSMUSG00000034265 | Zdhhc14 | zinc finger, DHHC domain containing 14 | -1.10 |
| ENSMUSG00000038422 | Hdhd3 | haloacid dehalogenase-like hydrolase domain containing 3 | -1.10 |
| ENSMUSG00000019370 | Calm3 | calmodulin 3 | -1.11 |
| ENSMUSG00000028830 | AU040320 | expressed sequence AU040320 | -1.11 |
| ENSMUSG00000030494 | Rhpn2 | rhophilin, Rho GTPase binding protein 2 | -1.11 |
| ENSMUSG00000031090 | Nadsyn1 | NAD synthetase 1 | -1.11 |
| ENSMUSG00000028521 | Slc35d1 | solute carrier family 35 (UDP-glucuronic acid/UDP-N-acetylgalactosamine dual transporter), member | -1.11 |
| ENSMUSG00000040613 | Apobec1 | apolipoprotein B mRNA editing enzyme, catalytic polypeptide 1 | -1.11 |
| ENSMUSG00000042251 | Pm20d1 | peptidase M20 domain containing 1 | -1.11 |
| ENSMUSG00000040957 | Cables1 | CDK5 and Abl enzyme substrate 1 | -1.11 |
| ENSMUSG00000026271 | Gpr35 | G protein-coupled receptor 35 | -1.11 |
| ENSMUSG00000068874 | Selenbp1 | selenium binding protein 1 | -1.11 |
| ENSMUSG00000050854 | Tmem125 | transmembrane protein 125 | -1.11 |
| ENSMUSG00000007817 | Zmiz1 | zinc finger, MIZ-type containing 1 | -1.12 |
| ENSMUSG00000020827 | Mink1 | misshapen-like kinase 1 (zebrafish) | -1.12 |
| ENSMUSG00000070002 | Ell | elongation factor RNA polymerase II | -1.12 |
| ENSMUSG00000042363 | Lgalsl | lectin, galactoside binding-like | -1.12 |
| ENSMUSG00000054252 | Fgfr3 | fibroblast growth factor receptor 3 | -1.12 |

| | | | |
|---|---|---|---|
| ENSMUSG00000042073 | Abhd14b | abhydrolase domain containing 14b | -1.12 |
| ENSMUSG00000029735 | Tpk1 | thiamine pyrophosphokinase | -1.12 |
| ENSMUSG00000021709 | Erbin | Erbb2 interacting protein | -1.13 |
| ENSMUSG00000024378 | Stard4 | StAR-related lipid transfer (START) domain containing 4 | -1.13 |
| ENSMUSG00000005580 | Adcy9 | adenylate cyclase 9 | -1.13 |
| ENSMUSG00000026956 | Uap1l1 | UDP-N-acteylglucosamine pyrophosphorylase 1-like 1 | -1.13 |
| ENSMUSG00000106375 | Gm43361 | predicted gene 43361 | -1.13 |
| ENSMUSG00000030545 | Pex11a | peroxisomal biogenesis factor 11 alpha | -1.13 |
| ENSMUSG00000029154 | Cwh43 | cell wall biogenesis 43 C-terminal homolog | -1.13 |
| ENSMUSG00000020777 | Acox1 | acyl-Coenzyme A oxidase 1, palmitoyl | -1.14 |
| ENSMUSG00000025068 | Gsto1 | glutathione S-transferase omega 1 | -1.14 |
| ENSMUSG00000092274 | Neat1 | nuclear paraspeckle assembly transcript 1 (non-protein coding) | -1.14 |
| ENSMUSG00000024431 | Nr3c1 | nuclear receptor subfamily 3, group C, member 1 | -1.14 |
| ENSMUSG00000009828 | Ick | intestinal cell kinase | -1.14 |
| ENSMUSG00000048707 | Tprn | taperin | -1.14 |
| ENSMUSG00000036918 | Ttc7 | tetratricopeptide repeat domain 7 | -1.14 |
| ENSMUSG00000029102 | Hgfac | hepatocyte growth factor activator | -1.14 |
| ENSMUSG00000040631 | Dok4 | docking protein 4 | -1.14 |
| ENSMUSG00000028223 | Decr1 | 2,4-dienoyl CoA reductase 1, mitochondrial | -1.14 |
| ENSMUSG00000041143 | Tmco4 | transmembrane and coiled-coil domains 4 | -1.14 |
| ENSMUSG00000105071 | Gm43336 | predicted gene 43336 | -1.14 |
| ENSMUSG00000048442 | Smim5 | small integral membrane protein 5 | -1.14 |
| ENSMUSG00000034579 | Pla2g3 | phospholipase A2, group III | -1.14 |
| ENSMUSG00000061718 | Ppp1r1b | protein phosphatase 1, regulatory inhibitor subunit 1B | -1.15 |
| ENSMUSG00000037434 | Slc30a1 | solute carrier family 30 (zinc transporter), member 1 | -1.15 |
| ENSMUSG00000022750 | Klhl22 | kelch-like 22 | -1.15 |
| ENSMUSG00000038291 | Snx25 | sorting nexin 25 | -1.15 |
| ENSMUSG00000015222 | Map2 | microtubule-associated protein 2 | -1.15 |
| ENSMUSG00000006641 | Slc5a6 | solute carrier family 5 (sodium-dependent vitamin transporter), member 6 | -1.15 |
| ENSMUSG00000050914 | Ankrd37 | ankyrin repeat domain 37 | -1.15 |
| ENSMUSG00000103948 | 4930594C11Rik | RIKEN cDNA 4930594C11 gene | -1.15 |
| ENSMUSG00000041757 | Plekha6 | pleckstrin homology domain containing, family A member 6 | -1.16 |
| ENSMUSG00000020986 | Sec23a | SEC23 homolog A, COPII coat complex component | -1.16 |
| ENSMUSG00000021550 | 2210016F16Rik | RIKEN cDNA 2210016F16 gene | -1.16 |
| ENSMUSG00000022221 | Ripk3 | receptor-interacting serine-threonine kinase 3 | -1.16 |
| ENSMUSG00000022090 | Pdlim2 | PDZ and LIM domain 2 | -1.16 |
| ENSMUSG00000029449 | Rhof | ras homolog family member F (in filopodia) | -1.16 |
| ENSMUSG00000022469 | Rapgef3 | Rap guanine nucleotide exchange factor (GEF) 3 | -1.16 |
| ENSMUSG00000017453 | Pipox | pipecolic acid oxidase | -1.16 |
| ENSMUSG00000021751 | Acox2 | acyl-Coenzyme A oxidase 2, branched chain | -1.16 |
| ENSMUSG00000026131 | Dst | dystonin | -1.17 |
| ENSMUSG00000019866 | Crybg1 | crystallin beta-gamma domain containing 1 | -1.17 |
| ENSMUSG00000029032 | Arhgef16 | Rho guanine nucleotide exchange factor (GEF) 16 | -1.17 |
| ENSMUSG00000038781 | Stap2 | signal transducing adaptor family member 2 | -1.17 |
| ENSMUSG00000005802 | Slc30a4 | solute carrier family 30 (zinc transporter), member 4 | -1.17 |
| ENSMUSG00000025477 | Inpp5a | inositol polyphosphate-5-phosphatase A | -1.17 |
| ENSMUSG00000070730 | Rmdn3 | regulator of microtubule dynamics 3 | -1.17 |
| ENSMUSG00000016495 | Plgrkt | plasminogen receptor, C-terminal lysine transmembrane protein | -1.17 |
| ENSMUSG00000021587 | Pcsk1 | proprotein convertase subtilisin/kexin type 1 | -1.17 |
| ENSMUSG00000045679 | Pqlc3 | PQ loop repeat containing | -1.17 |
| ENSMUSG00000043760 | Pkhd1 | polycystic kidney and hepatic disease 1 | -1.17 |
| ENSMUSG00000022565 | Plec | plectin | -1.18 |
| ENSMUSG00000024388 | Myo7b | myosin VIIB | -1.18 |
| ENSMUSG00000041959 | S100a10 | S100 calcium binding protein A10 (calpactin) | -1.18 |
| ENSMUSG00000063952 | Brpf3 | bromodomain and PHD finger containing, 3 | -1.18 |
| ENSMUSG00000020173 | Cobl | cordon-bleu WH2 repeat | -1.18 |
| ENSMUSG00000039145 | Camk1d | calcium/calmodulin-dependent protein kinase ID | -1.18 |
| ENSMUSG00000031391 | L1cam | L1 cell adhesion molecule | -1.18 |
| ENSMUSG00000022683 | Pla2g10 | phospholipase A2, group X | -1.18 |
| ENSMUSG00000027984 | Hadh | hydroxyacyl-Coenzyme A dehydrogenase | -1.19 |
| ENSMUSG00000038244 | Mical2 | microtubule associated monooxygenase, calponin and LIM domain containing 2 | -1.19 |
| ENSMUSG00000047123 | Ticam1 | toll-like receptor adaptor molecule 1 | -1.19 |
| ENSMUSG00000024112 | Cacna1h | calcium channel, voltage-dependent, T type, alpha 1H subunit | -1.19 |
| ENSMUSG00000033902 | Mapkbp1 | mitogen-activated protein kinase binding protein 1 | -1.19 |
| ENSMUSG00000069804 | Gm10277 | predicted gene 10277 | -1.19 |
| ENSMUSG00000069227 | Gprin1 | G protein-regulated inducer of neurite outgrowth 1 | -1.19 |
| ENSMUSG00000013418 | B4galnt2 | beta-1,4-N-acetyl-galactosaminyl transferase 2 | -1.20 |
| ENSMUSG00000064210 | Ano6 | anoctamin 6 | -1.20 |
| ENSMUSG00000032737 | Inppl1 | inositol polyphosphate phosphatase-like 1 | -1.20 |
| ENSMUSG00000017801 | Mlx | MAX-like protein X | -1.20 |
| ENSMUSG00000054733 | Msra | methionine sulfoxide reductase A | -1.20 |

| | | | |
|---|---|---|---|
| ENSMUSG00000042797 | Aqp11 | aquaporin 11 | -1.20 |
| ENSMUSG00000039747 | Orai2 | ORAI calcium release-activated calcium modulator 2 | -1.20 |
| ENSMUSG00000022680 | Pdxdc1 | pyridoxal-dependent decarboxylase domain containing 1 | -1.21 |
| ENSMUSG00000022946 | Dopey2 | dopey family member 2 | -1.21 |
| ENSMUSG00000039193 | Nlrc4 | NLR family, CARD domain containing 4 | -1.21 |
| ENSMUSG00000027938 | Creb3l4 | cAMP responsive element binding protein 3-like 4 | -1.21 |
| ENSMUSG00000074218 | Cox7a1 | cytochrome c oxidase subunit 7A1 | -1.21 |
| ENSMUSG00000020377 | Ltc4s | leukotriene C4 synthase | -1.21 |
| ENSMUSG00000033161 | Atp1a1 | ATPase, Na+/K+ transporting, alpha 1 polypeptide | -1.22 |
| ENSMUSG00000021876 | Rnase4 | ribonuclease, RNase A family 4 | -1.22 |
| ENSMUSG00000039682 | Lap3 | leucine aminopeptidase 3 | -1.22 |
| ENSMUSG00000050931 | Sgms2 | sphingomyelin synthase 2 | -1.22 |
| ENSMUSG00000022265 | Ank | progressive ankylosis | -1.22 |
| ENSMUSG00000039166 | Akap7 | A kinase (PRKA) anchor protein 7 | -1.22 |
| ENSMUSG00000029167 | Ppargc1a | peroxisome proliferative activated receptor, gamma, coactivator 1 alpha | -1.22 |
| ENSMUSG00000059323 | Tonsl | tonsoku-like, DNA repair protein | -1.22 |
| ENSMUSG00000031378 | Abcd1 | ATP-binding cassette, sub-family D (ALD), member 1 | -1.22 |
| ENSMUSG00000029471 | Camkk2 | calcium/calmodulin-dependent protein kinase kinase 2, beta | -1.22 |
| ENSMUSG00000054200 | Ffar4 | free fatty acid receptor 4 | -1.22 |
| ENSMUSG00000001227 | Sema6b | sema domain, transmembrane domain (TM), and cytoplasmic domain, (semaphorin) 6B | -1.22 |
| ENSMUSG00000035441 | Myo1d | myosin ID | -1.23 |
| ENSMUSG00000020841 | Cpd | carboxypeptidase D | -1.23 |
| ENSMUSG00000027007 | Ssfa2 | sperm specific antigen 2 | -1.23 |
| ENSMUSG00000038260 | Trpm4 | transient receptor potential cation channel, subfamily M, member 4 | -1.23 |
| ENSMUSG00000036499 | Eea1 | early endosome antigen 1 | -1.23 |
| ENSMUSG00000028894 | Inpp5b | inositol polyphosphate-5-phosphatase B | -1.23 |
| ENSMUSG00000015342 | Xk | X-linked Kx blood group | -1.23 |
| ENSMUSG00000021379 | Id4 | inhibitor of DNA binding 4 | -1.23 |
| ENSMUSG00000097006 | 9530082P21Rik | RIKEN cDNA 9530082P21 gene | -1.23 |
| ENSMUSG00000102317 | Gm37628 | predicted gene, 37628 | -1.23 |
| ENSMUSG00000033577 | Myo6 | myosin VI | -1.24 |
| ENSMUSG00000033885 | Pxk | PX domain containing serine/threonine kinase | -1.24 |
| ENSMUSG00000027634 | Ndrg3 | N-myc downstream regulated gene 3 | -1.24 |
| ENSMUSG00000039197 | Adk | adenosine kinase | -1.24 |
| ENSMUSG00000028862 | Map3k6 | mitogen-activated protein kinase kinase kinase 6 | -1.24 |
| ENSMUSG00000034731 | Dgkh | diacylglycerol kinase, eta | -1.24 |
| ENSMUSG00000030590 | Fam98c | family with sequence similarity 98, member C | -1.24 |
| ENSMUSG00000110289 | 4930412F12Rik | RIKEN cDNA 4930412F12 gene | -1.24 |
| ENSMUSG00000020009 | Ifngr1 | interferon gamma receptor 1 | -1.25 |
| ENSMUSG00000033416 | Gucd1 | guanylyl cyclase domain containing 1 | -1.25 |
| ENSMUSG00000023019 | Gpd1 | glycerol-3-phosphate dehydrogenase 1 (soluble) | -1.25 |
| ENSMUSG00000024063 | Lbh | limb-bud and heart | -1.25 |
| ENSMUSG00000021236 | Entpd5 | ectonucleoside triphosphate diphosphohydrolase 5 | -1.25 |
| ENSMUSG00000025190 | Got1 | glutamic-oxaloacetic transaminase 1, soluble | -1.25 |
| ENSMUSG00000029413 | Naaa | N-acylethanolamine acid amidase | -1.25 |
| ENSMUSG00000058022 | Adtrp | androgen dependent TFPI regulating protein | -1.25 |
| ENSMUSG00000050777 | Tmem37 | transmembrane protein 37 | -1.25 |
| ENSMUSG00000049598 | Vsig8 | V-set and immunoglobulin domain containing 8 | -1.25 |
| ENSMUSG00000044042 | Fmn1 | formin 1 | -1.26 |
| ENSMUSG00000042808 | Gpx2 | glutathione peroxidase 2 | -1.26 |
| ENSMUSG00000084883 | Ccdc85c | coiled-coil domain containing 85C | -1.26 |
| ENSMUSG00000061845 | Defa35 | defensin, alpha, 35 | -1.26 |
| ENSMUSG00000029470 | P2rx4 | purinergic receptor P2X, ligand-gated ion channel 4 | -1.26 |
| ENSMUSG00000026259 | Ngef | neuronal guanine nucleotide exchange factor | -1.26 |
| ENSMUSG00000006567 | Atp7b | ATPase, Cu++ transporting, beta polypeptide | -1.26 |
| ENSMUSG00000093394 | Gm20621 | predicted gene 20621 | -1.26 |
| ENSMUSG00000022711 | Pmm2 | phosphomannomutase 2 | -1.27 |
| ENSMUSG00000038507 | Parp12 | poly (ADP-ribose) polymerase family, member 12 | -1.27 |
| ENSMUSG00000027457 | Snph | syntaphilin | -1.27 |
| ENSMUSG00000090115 | Usp49 | ubiquitin specific peptidase 49 | -1.27 |
| ENSMUSG00000042097 | Zfp239 | zinc finger protein 239 | -1.27 |
| ENSMUSG00000025584 | Pde8a | phosphodiesterase 8A | -1.27 |
| ENSMUSG00000010830 | Kdelr3 | KDEL (Lys-Asp-Glu-Leu) endoplasmic reticulum protein retention receptor 3 | -1.27 |
| ENSMUSG00000034509 | Mad2l1bp | MAD2L1 binding protein | -1.27 |
| ENSMUSG00000056492 | Adgrf5 | adhesion G protein-coupled receptor F5 | -1.27 |
| ENSMUSG00000099582 | Gm29014 | predicted gene 29014 | -1.27 |
| ENSMUSG00000032324 | Tspan3 | tetraspanin 3 | -1.28 |
| ENSMUSG00000005125 | Ndrg1 | N-myc downstream regulated gene 1 | -1.28 |
| ENSMUSG00000063931 | Pepd | peptidase D | -1.28 |
| ENSMUSG00000046519 | Golph3l | golgi phosphoprotein 3-like | -1.28 |
| ENSMUSG00000034066 | Farp2 | FERM, RhoGEF and pleckstrin domain protein 2 | -1.28 |

| | | | |
|---|---|---|---|
| ENSMUSG00000047409 | Ctdspl | CTD (carboxy-terminal domain, RNA polymerase II, polypeptide A) small phosphatase-like | -1.28 |
| ENSMUSG00000032842 | Abcc10 | ATP-binding cassette, sub-family C (CFTR/MRP), member 10 | -1.28 |
| ENSMUSG00000027314 | Dll4 | delta like canonical Notch ligand 4 | -1.28 |
| ENSMUSG00000072115 | Ang | angiogenin, ribonuclease, RNase A family, 5 | -1.28 |
| ENSMUSG00000104576 | F830115B05Rik | RIKEN cDNA F830115B05 gene | -1.28 |
| ENSMUSG00000035852 | Misp | mitotic spindle positioning | -1.29 |
| ENSMUSG00000029859 | Epha1 | Eph receptor A1 | -1.29 |
| ENSMUSG00000039304 | Tnfsf10 | tumor necrosis factor (ligand) superfamily, member 10 | -1.29 |
| ENSMUSG00000060147 | Serpinb6a | serine (or cysteine) peptidase inhibitor, clade B, member 6a | -1.30 |
| ENSMUSG00000033684 | Qsox1 | quiescin Q6 sulfhydryl oxidase 1 | -1.30 |
| ENSMUSG00000035722 | Abca7 | ATP-binding cassette, sub-family A (ABC1), member 7 | -1.30 |
| ENSMUSG00000024955 | Esrra | estrogen related receptor, alpha | -1.30 |
| ENSMUSG00000038332 | Sesn1 | sestrin 1 | -1.30 |
| ENSMUSG00000029166 | Mapre3 | microtubule-associated protein, RP/EB family, member 3 | -1.30 |
| ENSMUSG00000054474 | Thnsl2 | threonine synthase-like 2 (bacterial) | -1.30 |
| ENSMUSG00000050944 | Efcab5 | EF-hand calcium binding domain 5 | -1.30 |
| ENSMUSG00000063689 | Hist2h2ab | histone cluster 2, H2ab | -1.30 |
| ENSMUSG00000029452 | Tmem116 | transmembrane protein 116 | -1.30 |
| ENSMUSG00000102837 | Gm37383 | predicted gene, 37383 | -1.30 |
| ENSMUSG00000024597 | Slc12a2 | solute carrier family 12, member 2 | -1.31 |
| ENSMUSG00000020432 | Tcn2 | transcobalamin 2 | -1.31 |
| ENSMUSG00000042659 | Arrdc4 | arrestin domain containing 4 | -1.31 |
| ENSMUSG00000021665 | Hexb | hexosaminidase B | -1.31 |
| ENSMUSG00000031453 | Rasa3 | RAS p21 protein activator 3 | -1.31 |
| ENSMUSG00000031788 | Kifc3 | kinesin family member C3 | -1.31 |
| ENSMUSG00000032860 | P2ry2 | purinergic receptor P2Y, G-protein coupled 2 | -1.31 |
| ENSMUSG00000086674 | Zfp286os | zinc finger protein 286, opposite strand | -1.31 |
| ENSMUSG00000059412 | Fxyd2 | FXYD domain-containing ion transport regulator 2 | -1.31 |
| ENSMUSG00000027800 | Tm4sf1 | transmembrane 4 superfamily member 1 | -1.31 |
| ENSMUSG00000109852 | Gm45360 | predicted gene 45360 | -1.31 |
| ENSMUSG00000028399 | Ptprd | protein tyrosine phosphatase, receptor type, D | -1.32 |
| ENSMUSG00000036402 | Gng12 | guanine nucleotide binding protein (G protein), gamma 12 | -1.32 |
| ENSMUSG00000039131 | Gipc2 | GIPC PDZ domain containing family, member 2 | -1.32 |
| ENSMUSG00000029053 | Prkcz | protein kinase C, zeta | -1.32 |
| ENSMUSG00000034248 | Slc25a37 | solute carrier family 25, member 37 | -1.32 |
| ENSMUSG00000020573 | Pik3cg | phosphatidylinositol-4,5-bisphosphate 3-kinase catalytic subunit gamma | -1.32 |
| ENSMUSG00000038365 | Fbxo25 | F-box protein 25 | -1.32 |
| ENSMUSG00000034371 | Tkfc | triokinase, FMN cyclase | -1.32 |
| ENSMUSG00000022900 | Ildr1 | immunoglobulin-like domain containing receptor 1 | -1.32 |
| ENSMUSG00000058586 | Serhl | serine hydrolase-like | -1.32 |
| ENSMUSG00000023150 | Ivns1abp | influenza virus NS1A binding protein | -1.33 |
| ENSMUSG00000025504 | Eps8l2 | EPS8-like 2 | -1.33 |
| ENSMUSG00000020520 | Galnt10 | polypeptide N-acetylgalactosaminyltransferase 10 | -1.33 |
| ENSMUSG00000022330 | Osr2 | odd-skipped related 2 | -1.33 |
| ENSMUSG00000056602 | Fry | FRY microtubule binding protein | -1.33 |
| ENSMUSG00000055866 | Per2 | period circadian clock 2 | -1.33 |
| ENSMUSG00000032380 | Dapk2 | death-associated protein kinase 2 | -1.33 |
| ENSMUSG00000086549 | Gm13648 | predicted gene 13648 | -1.33 |
| ENSMUSG00000116594 | AC133488.1 | novel protein | -1.33 |
| ENSMUSG00000045664 | Cdc42ep2 | CDC42 effector protein (Rho GTPase binding) 2 | -1.33 |
| ENSMUSG00000021223 | Papln | papilin, proteoglycan-like sulfated glycoprotein | -1.33 |
| ENSMUSG00000050270 | Tmem220 | transmembrane protein 220 | -1.33 |
| ENSMUSG00000026103 | Gls | glutaminase | -1.34 |
| ENSMUSG00000021411 | Pxdc1 | PX domain containing 1 | -1.34 |
| ENSMUSG00000115062 | 4930544F09Rik | RIKEN cDNA 4930544F09 gene | -1.34 |
| ENSMUSG00000021803 | Cdhr1 | cadherin-related family member 1 | -1.34 |
| ENSMUSG00000114612 | Gm47550 | predicted gene, 47550 | -1.34 |
| ENSMUSG00000111176 | Gm47233 | predicted gene, 47233 | -1.34 |
| ENSMUSG00000028719 | Cmpk1 | cytidine monophosphate (UMP-CMP) kinase 1 | -1.35 |
| ENSMUSG00000060012 | Kif13b | kinesin family member 13B | -1.35 |
| ENSMUSG00000014418 | Hps5 | HPS5, biogenesis of lysosomal organelles complex 2 subunit 2 | -1.35 |
| ENSMUSG00000053702 | Nebl | nebulette | -1.35 |
| ENSMUSG00000035473 | Galm | galactose mutarotase | -1.35 |
| ENSMUSG00000029385 | Ccng2 | cyclin G2 | -1.35 |
| ENSMUSG00000045314 | Sowahb | sosondowah ankyrin repeat domain family member B | -1.35 |
| ENSMUSG00000031897 | Psmb10 | proteasome (prosome, macropain) subunit, beta type 10 | -1.35 |
| ENSMUSG00000063838 | Cdc42ep5 | CDC42 effector protein (Rho GTPase binding) 5 | -1.35 |
| ENSMUSG00000026480 | Ncf2 | neutrophil cytosolic factor 2 | -1.35 |
| ENSMUSG00000112944 | Gm48885 | predicted gene, 48885 | -1.35 |
| ENSMUSG00000067203 | H2-K2 | histocompatibility 2, K region locus 2 | -1.35 |
| ENSMUSG00000078954 | Arhgap8 | Rho GTPase activating protein 8 | -1.35 |

| | | | |
|---|---|---|---|
| ENSMUSG00000102953 | Gm37019 | predicted gene, 37019 | -1.35 |
| ENSMUSG00000106659 | Gm42161 | predicted gene, 42161 | -1.35 |
| ENSMUSG00000030739 | Myh14 | myosin, heavy polypeptide 14 | -1.36 |
| ENSMUSG00000020029 | Nudt4 | nudix (nucleoside diphosphate linked moiety X)-type motif 4 | -1.36 |
| ENSMUSG00000024981 | Acsl5 | acyl-CoA synthetase long-chain family member 5 | -1.36 |
| ENSMUSG00000061666 | Gdpd1 | glycerophosphodiester phosphodiesterase domain containing 1 | -1.36 |
| ENSMUSG00000020752 | Recql5 | RecQ protein-like 5 | -1.36 |
| ENSMUSG00000037579 | Kcnh3 | potassium voltage-gated channel, subfamily H (eag-related), member 3 | -1.36 |
| ENSMUSG00000032454 | Rbp2 | retinol binding protein 2, cellular | -1.36 |
| ENSMUSG00000030921 | Trim30a | tripartite motif-containing 30A | -1.36 |
| ENSMUSG00000024187 | Fam234a | family with sequence similarity 234, member A | -1.37 |
| ENSMUSG00000022843 | Clcn2 | chloride channel, voltage-sensitive 2 | -1.37 |
| ENSMUSG00000036452 | Arhgap26 | Rho GTPase activating protein 26 | -1.37 |
| ENSMUSG00000043445 | Pgp | phosphoglycolate phosphatase | -1.37 |
| ENSMUSG00000026102 | Inpp1 | inositol polyphosphate-1-phosphatase | -1.37 |
| ENSMUSG00000044681 | Cnpy1 | canopy FGF signaling regulator 1 | -1.37 |
| ENSMUSG00000070034 | Sp110 | Sp110 nuclear body protein | -1.37 |
| ENSMUSG00000108526 | Gm45828 | predicted gene 45828 | -1.37 |
| ENSMUSG00000028217 | Cdh17 | cadherin 17 | -1.38 |
| ENSMUSG00000000915 | Hip1r | huntingtin interacting protein 1 related | -1.38 |
| ENSMUSG00000031482 | Slc25a15 | solute carrier family 25 (mitochondrial carrier ornithine transporter), member 15 | -1.38 |
| ENSMUSG00000031887 | Tradd | TNFRSF1A-associated via death domain | -1.38 |
| ENSMUSG00000075010 | AW112010 | expressed sequence AW112010 | -1.38 |
| ENSMUSG00000029275 | Gfi1 | growth factor independent 1 | -1.38 |
| ENSMUSG00000025466 | Fuom | fucose mutarotase | -1.38 |
| ENSMUSG00000029923 | Rab19 | RAB19, member RAS oncogene family | -1.38 |
| ENSMUSG00000074882 | Cyp2c68 | cytochrome P450, family 2, subfamily c, polypeptide 68 | -1.38 |
| ENSMUSG00000033721 | Vav3 | vav 3 oncogene | -1.38 |
| ENSMUSG00000104693 | Gm42941 | predicted gene 42941 | -1.38 |
| ENSMUSG00000038239 | Hrc | histidine rich calcium binding protein | -1.38 |
| ENSMUSG00000036813 | Entpd8 | ectonucleoside triphosphate diphosphohydrolase 8 | -1.39 |
| ENSMUSG00000041827 | Oasl1 | 2'-5' oligoadenylate synthetase-like 1 | -1.39 |
| ENSMUSG00000056938 | Acbd4 | acyl-Coenzyme A binding domain containing 4 | -1.39 |
| ENSMUSG00000021553 | Slc28a3 | solute carrier family 28 (sodium-coupled nucleoside transporter), member 3 | -1.39 |
| ENSMUSG00000002289 | Angptl4 | angiopoietin-like 4 | -1.39 |
| ENSMUSG00000113516 | Gm47368 | predicted gene, 47368 | -1.39 |
| ENSMUSG00000109446 | Gm9195 | predicted gene 9195 | -1.39 |
| ENSMUSG00000040345 | Arhgap9 | Rho GTPase activating protein 9 | -1.39 |
| ENSMUSG00000106206 | Gm43094 | predicted gene 43094 | -1.39 |
| ENSMUSG00000104094 | Gm37314 | predicted gene, 37314 | -1.39 |
| ENSMUSG00000018166 | Erbb3 | erb-b2 receptor tyrosine kinase 3 | -1.40 |
| ENSMUSG00000031955 | Bcar1 | breast cancer anti-estrogen resistance 1 | -1.40 |
| ENSMUSG00000046861 | Hectd3 | HECT domain E3 ubiquitin protein ligase 3 | -1.40 |
| ENSMUSG00000039234 | Sec24d | Sec24 related gene family, member D (S. cerevisiae) | -1.40 |
| ENSMUSG00000031823 | Zdhhc7 | zinc finger, DHHC domain containing 7 | -1.40 |
| ENSMUSG00000030761 | Myo7a | myosin VIIA | -1.40 |
| ENSMUSG00000023915 | Tnfrsf21 | tumor necrosis factor receptor superfamily, member 21 | -1.40 |
| ENSMUSG00000018983 | E2f2 | E2F transcription factor 2 | -1.40 |
| ENSMUSG00000020182 | Ddc | dopa decarboxylase | -1.40 |
| ENSMUSG00000028795 | Ccdc28b | coiled coil domain containing 28B | -1.40 |
| ENSMUSG00000046456 | Tmem150b | transmembrane protein 150B | -1.40 |
| ENSMUSG00000022150 | Dab2 | disabled 2, mitogen-responsive phosphoprotein | -1.40 |
| ENSMUSG00000030934 | Oat | ornithine aminotransferase | -1.41 |
| ENSMUSG00000026687 | Aldh9a1 | aldehyde dehydrogenase 9, subfamily A1 | -1.41 |
| ENSMUSG00000021266 | Wars | tryptophanyl-tRNA synthetase | -1.41 |
| ENSMUSG00000042228 | Lyn | LYN proto-oncogene, Src family tyrosine kinase | -1.41 |
| ENSMUSG00000070661 | Rnf186 | ring finger protein 186 | -1.41 |
| ENSMUSG00000024247 | Pkdcc | protein kinase domain containing, cytoplasmic | -1.41 |
| ENSMUSG00000006395 | Hyi | hydroxypyruvate isomerase (putative) | -1.41 |
| ENSMUSG00000020847 | Rph3al | rabphilin 3A-like (without C2 domains) | -1.41 |
| ENSMUSG00000023393 | Slc17a9 | solute carrier family 17, member 9 | -1.41 |
| ENSMUSG00000106969 | Gm42882 | predicted gene 42882 | -1.41 |
| ENSMUSG00000098678 | Mroh6 | maestro heat-like repeat family member 6 | -1.41 |
| ENSMUSG00000074440 | Defa3 | defensin, alpha, 3 | -1.41 |
| ENSMUSG00000022419 | Deptor | DEP domain containing MTOR-interacting protein | -1.42 |
| ENSMUSG00000007038 | Neu1 | neuraminidase 1 | -1.42 |
| ENSMUSG00000038331 | Satb2 | special AT-rich sequence binding protein 2 | -1.42 |
| ENSMUSG00000039813 | Tbc1d2 | TBC1 domain family, member 2 | -1.42 |
| ENSMUSG00000026976 | Pax8 | paired box 8 | -1.42 |
| ENSMUSG00000111468 | 5033425B01Rik | RIKEN cDNA 5033425B01 gene | -1.42 |
| ENSMUSG00000086067 | Gm16183 | predicted gene 16183 | -1.42 |

| | | | |
|---|---|---|---|
| ENSMUSG00000022186 | Oxct1 | 3-oxoacid CoA transferase 1 | -1.43 |
| ENSMUSG00000047153 | Khnyn | KH and NYN domain containing | -1.43 |
| ENSMUSG00000023262 | Acy1 | aminoacylase 1 | -1.43 |
| ENSMUSG00000035509 | Fbxl21 | F-box and leucine-rich repeat protein 21 | -1.43 |
| ENSMUSG00000029381 | Shroom3 | shroom family member 3 | -1.44 |
| ENSMUSG00000026879 | Gsn | gelsolin | -1.44 |
| ENSMUSG00000020111 | Micu1 | mitochondrial calcium uptake 1 | -1.44 |
| ENSMUSG00000032602 | Slc25a20 | solute carrier family 25 (mitochondrial carnitine/acylcarnitine translocase), member 20 | -1.44 |
| ENSMUSG00000018334 | Ksr1 | kinase suppressor of ras 1 | -1.44 |
| ENSMUSG00000034656 | Cacna1a | calcium channel, voltage-dependent, P/Q type, alpha 1A subunit | -1.44 |
| ENSMUSG00000013539 | Tango2 | transport and golgi organization 2 | -1.44 |
| ENSMUSG00000029401 | Rilpl2 | Rab interacting lysosomal protein-like 2 | -1.44 |
| ENSMUSG00000003585 | Sec14l2 | SEC14-like lipid binding 2 | -1.44 |
| ENSMUSG00000085088 | 4931413K12Rik | RIKEN cDNA 4931413K12 gene | -1.44 |
| ENSMUSG00000019122 | Ccl9 | chemokine (C-C motif) ligand 9 | -1.44 |
| ENSMUSG00000071550 | Cfap44 | cilia and flagella associated protein 44 | -1.44 |
| ENSMUSG00000018776 | Slc35g3 | solute carrier family 35, member G3 | -1.44 |
| ENSMUSG00000111412 | Gm47047 | predicted gene, 47047 | -1.44 |
| ENSMUSG00000102540 | Gm38134 | predicted gene, 38134 | -1.44 |
| ENSMUSG00000009647 | Mcu | mitochondrial calcium uniporter | -1.45 |
| ENSMUSG00000041605 | Inava | innate immunity activator | -1.45 |
| ENSMUSG00000003746 | Man1a | mannosidase 1, alpha | -1.45 |
| ENSMUSG00000000538 | Tom1l2 | target of myb1-like 2 (chicken) | -1.45 |
| ENSMUSG00000031732 | Phlpp2 | PH domain and leucine rich repeat protein phosphatase 2 | -1.45 |
| ENSMUSG00000026922 | Agpat2 | 1-acylglycerol-3-phosphate O-acyltransferase 2 (lysophosphatidic acid acyltransferase, beta) | -1.45 |
| ENSMUSG00000037921 | Ddx60 | DEAD (Asp-Glu-Ala-Asp) box polypeptide 60 | -1.45 |
| ENSMUSG00000047250 | Ptgs1 | prostaglandin-endoperoxide synthase 1 | -1.45 |
| ENSMUSG00000078612 | Fyb2 | FYN binding protein 2 | -1.45 |
| ENSMUSG00000091890 | A830073O21Rik | RIKEN cDNA A830073O21 gene | -1.45 |
| ENSMUSG00000092074 | Dynlt1a | dynein light chain Tctex-type 1A | -1.45 |
| ENSMUSG00000113119 | Gm48883 | predicted gene, 48883 | -1.45 |
| ENSMUSG00000037033 | Clca3b | chloride channel accessory 3B | -1.46 |
| ENSMUSG00000021556 | Golm1 | golgi membrane protein 1 | -1.46 |
| ENSMUSG00000033124 | Atg9a | autophagy related 9A | -1.46 |
| ENSMUSG00000068417 | Pnp2 | purine-nucleoside phosphorylase 2 | -1.46 |
| ENSMUSG00000038811 | Gngt2 | guanine nucleotide binding protein (G protein), gamma transducing activity polypeptide 2 | -1.46 |
| ENSMUSG00000116220 | A430088P11Rik | RIKEN cDNA A430088P11 gene | -1.46 |
| ENSMUSG00000107310 | Gm7452 | predicted pseudogene 7452 | -1.46 |
| ENSMUSG00000030681 | Mvp | major vault protein | -1.47 |
| ENSMUSG00000024556 | Me2 | malic enzyme 2, NAD(+)-dependent, mitochondrial | -1.47 |
| ENSMUSG00000007036 | Abhd16a | abhydrolase domain containing 16A | -1.47 |
| ENSMUSG00000039637 | Coro7 | coronin 7 | -1.47 |
| ENSMUSG00000036106 | Prr5 | proline rich 5 (renal) | -1.47 |
| ENSMUSG00000035878 | Hykk | hydroxylysine kinase 1 | -1.47 |
| ENSMUSG00000079174 | Gm3054 | predicted gene 3054 | -1.47 |
| ENSMUSG00000090293 | Gm17034 | predicted gene 17034 | -1.47 |
| ENSMUSG00000113935 | Gm35732 | predicted gene, 35732 | -1.47 |
| ENSMUSG00000048992 | Prss32 | protease, serine 32 | -1.48 |
| ENSMUSG00000039774 | Galnt12 | polypeptide N-acetylgalactosaminyltransferase 12 | -1.48 |
| ENSMUSG00000020099 | Unc5b | unc-5 netrin receptor B | -1.48 |
| ENSMUSG00000038497 | Tmco3 | transmembrane and coiled-coil domains 3 | -1.48 |
| ENSMUSG00000026094 | Stk17b | serine/threonine kinase 17b (apoptosis-inducing) | -1.48 |
| ENSMUSG00000022560 | Slc52a2 | solute carrier protein 52, member 2 | -1.48 |
| ENSMUSG00000087535 | Zmiz1os1 | Zmiz1 opposite strand 1 | -1.48 |
| ENSMUSG00000084978 | Gm11655 | predicted gene 11655 | -1.48 |
| ENSMUSG00000013707 | Tnfaip8l2 | tumor necrosis factor, alpha-induced protein 8-like 2 | -1.48 |
| ENSMUSG00000043004 | Gng2 | guanine nucleotide binding protein (G protein), gamma 2 | -1.48 |
| ENSMUSG00000030616 | Sytl2 | synaptotagmin-like 2 | -1.49 |
| ENSMUSG00000028064 | Sema4a | sema domain, immunoglobulin domain (Ig), transmembrane domain (TM) and short cytoplasmic dom | -1.49 |
| ENSMUSG00000023909 | Paqr4 | progestin and adipoQ receptor family member IV | -1.49 |
| ENSMUSG00000031021 | Tmem9b | TMEM9 domain family, member B | -1.49 |
| ENSMUSG00000036553 | Sh3tc1 | SH3 domain and tetratricopeptide repeats 1 | -1.49 |
| ENSMUSG00000087691 | Gm15674 | predicted gene 15674 | -1.49 |
| ENSMUSG00000113776 | Gm19327 | predicted gene, 19327 | -1.49 |
| ENSMUSG00000092212 | Slc22a13b-ps | solute carrier family 22 (organic cation transporter), member 13b, pseudogene | -1.49 |
| ENSMUSG00000081245 | Gm6587 | predicted gene 6587 | -1.49 |
| ENSMUSG00000038298 | Pdzk1 | PDZ domain containing 1 | -1.49 |
| ENSMUSG00000097862 | Gm26859 | predicted gene, 26859 | -1.49 |
| ENSMUSG00000085944 | 1700003D09Rik | RIKEN cDNA 1700003D09 gene | -1.49 |
| ENSMUSG00000021591 | Glrx | glutaredoxin | -1.50 |
| ENSMUSG00000033917 | Gde1 | glycerophosphodiester phosphodiesterase 1 | -1.50 |

| | | | |
|---|---|---|---|
| ENSMUSG00000021913 | Ogdhl | oxoglutarate dehydrogenase-like | -1.50 |
| ENSMUSG00000085333 | 1700030A11Rik | RIKEN cDNA 1700030A11 gene | -1.50 |
| ENSMUSG00000096145 | Vkorc1 | vitamin K epoxide reductase complex, subunit 1 | -1.50 |
| ENSMUSG00000083020 | Gm15331 | predicted gene 15331 | -1.50 |
| ENSMUSG00000026546 | Cfap45 | cilia and flagella associated protein 45 | -1.50 |
| ENSMUSG00000027695 | Pld1 | phospholipase D1 | -1.51 |
| ENSMUSG00000076435 | Acsf2 | acyl-CoA synthetase family member 2 | -1.51 |
| ENSMUSG00000032047 | Acat1 | acetyl-Coenzyme A acetyltransferase 1 | -1.51 |
| ENSMUSG00000027765 | P2ry1 | purinergic receptor P2Y, G-protein coupled 1 | -1.51 |
| ENSMUSG00000036136 | Fam110c | family with sequence similarity 110, member C | -1.51 |
| ENSMUSG00000001665 | Gstt3 | glutathione S-transferase, theta 3 | -1.51 |
| ENSMUSG00000104132 | Gm36952 | predicted gene, 36952 | -1.51 |
| ENSMUSG00000085152 | Gm11496 | predicted gene 11496 | -1.51 |
| ENSMUSG00000109715 | Gm45606 | predicted gene 45606 | -1.51 |
| ENSMUSG00000038209 | Itln1 | intelectin 1 (galactofuranose binding) | -1.52 |
| ENSMUSG00000024074 | Crim1 | cysteine rich transmembrane BMP regulator 1 (chordin like) | -1.52 |
| ENSMUSG00000031327 | Chic1 | cysteine-rich hydrophobic domain 1 | -1.52 |
| ENSMUSG00000032080 | Apoa4 | apolipoprotein A-IV | -1.52 |
| ENSMUSG00000020334 | Slc22a4 | solute carrier family 22 (organic cation transporter), member 4 | -1.52 |
| ENSMUSG00000096954 | Gdap10 | ganglioside-induced differentiation-associated-protein 10 | -1.52 |
| ENSMUSG00000019762 | Iyd | iodotyrosine deiodinase | -1.52 |
| ENSMUSG00000024386 | Proc | protein C | -1.52 |
| ENSMUSG00000042942 | Greb1l | growth regulation by estrogen in breast cancer-like | -1.52 |
| ENSMUSG00000066178 | 6030445D17Rik | RIKEN cDNA 6030445D17 gene | -1.52 |
| ENSMUSG00000020154 | Ptprb | protein tyrosine phosphatase, receptor type, B | -1.52 |
| ENSMUSG00000073608 | Gal3st2c | galactose-3-O-sulfotransferase 2C | -1.52 |
| ENSMUSG00000020226 | Slc5a4b | solute carrier family 5 (neutral amino acid transporters, system A), member 4b | -1.52 |
| ENSMUSG00000111028 | Gm5922 | predicted gene 5922 | -1.52 |
| ENSMUSG00000022048 | Dpysl2 | dihydropyrimidinase-like 2 | -1.53 |
| ENSMUSG00000045934 | Mtmr11 | myotubularin related protein 11 | -1.53 |
| ENSMUSG00000029847 | Slc23a4 | solute carrier family 23 member 4 | -1.53 |
| ENSMUSG00000039754 | Alkbh4 | alkB homolog 4, lysine demethylase | -1.53 |
| ENSMUSG00000078664 | Sprr2a1 | small proline-rich protein 2A1 | -1.53 |
| ENSMUSG00000046005 | D830044D21Rik | RIKEN cDNA D830044D21 gene | -1.53 |
| ENSMUSG00000041237 | Pklr | pyruvate kinase liver and red blood cell | -1.54 |
| ENSMUSG00000022105 | Rb1 | RB transcriptional corepressor 1 | -1.54 |
| ENSMUSG00000041992 | Rapgef5 | Rap guanine nucleotide exchange factor (GEF) 5 | -1.54 |
| ENSMUSG00000026304 | Rab17 | RAB17, member RAS oncogene family | -1.54 |
| ENSMUSG00000070563 | Spaca4 | sperm acrosome associated 4 | -1.54 |
| ENSMUSG00000029352 | Crybb3 | crystallin, beta B3 | -1.54 |
| ENSMUSG00000104776 | Gm43691 | predicted gene 43691 | -1.54 |
| ENSMUSG00000078307 | AI593442 | expressed sequence AI593442 | -1.54 |
| ENSMUSG00000074217 | 2210011C24Rik | RIKEN cDNA 2210011C24 gene | -1.54 |
| ENSMUSG00000103845 | Gm19026 | predicted gene, 19026 | -1.54 |
| ENSMUSG00000091475 | 2810468N07Rik | RIKEN cDNA 2810468N07 gene | -1.54 |
| ENSMUSG00000038541 | Srd5a2 | steroid 5 alpha-reductase 2 | -1.54 |
| ENSMUSG00000065581 | Mir194-1 | microRNA 194-1 | -1.54 |
| ENSMUSG00000029646 | Cdx2 | caudal type homeobox 2 | -1.55 |
| ENSMUSG00000028519 | Dab1 | disabled 1 | -1.55 |
| ENSMUSG00000027524 | Edn3 | endothelin 3 | -1.55 |
| ENSMUSG00000042333 | Tnfrsf14 | tumor necrosis factor receptor superfamily, member 14 (herpesvirus entry mediator) | -1.55 |
| ENSMUSG00000071604 | Fam189a2 | family with sequence similarity 189, member A2 | -1.55 |
| ENSMUSG00000040147 | Maob | monoamine oxidase B | -1.55 |
| ENSMUSG00000104950 | 4833413G10Rik | RIKEN cDNA 4833413G10 gene | -1.55 |
| ENSMUSG00000052767 | Gm12703 | predicted gene 12703 | -1.55 |
| ENSMUSG00000064036 | Mro | maestro | -1.55 |
| ENSMUSG00000040528 | Milr1 | mast cell immunoglobulin like receptor 1 | -1.55 |
| ENSMUSG00000067049 | Unc93a | unc-93 homolog A | -1.55 |
| ENSMUSG00000057914 | Cacnb2 | calcium channel, voltage-dependent, beta 2 subunit | -1.55 |
| ENSMUSG00000024610 | Cd74 | CD74 antigen (invariant polypeptide of major histocompatibility complex, class II antigen-associated) | -1.56 |
| ENSMUSG00000024511 | Rab27b | RAB27B, member RAS oncogene family | -1.56 |
| ENSMUSG00000024887 | Asah2 | N-acylsphingosine amidohydrolase 2 | -1.56 |
| ENSMUSG00000021779 | Thrb | thyroid hormone receptor beta | -1.56 |
| ENSMUSG00000109314 | Gm44699 | predicted gene 44699 | -1.56 |
| ENSMUSG00000026069 | Il1rl1 | interleukin 1 receptor-like 1 | -1.56 |
| ENSMUSG00000053819 | Camk2d | calcium/calmodulin-dependent protein kinase II, delta | -1.57 |
| ENSMUSG00000027901 | Dennd2d | DENN/MADD domain containing 2D | -1.57 |
| ENSMUSG00000052396 | Mogat2 | monoacylglycerol O-acyltransferase 2 | -1.57 |
| ENSMUSG00000025006 | Sorbs1 | sorbin and SH3 domain containing 1 | -1.57 |
| ENSMUSG00000034591 | Slc41a2 | solute carrier family 41, member 2 | -1.57 |
| ENSMUSG00000031337 | Mtm1 | X-linked myotubular myopathy gene 1 | -1.57 |

| | | | |
|---|---|---|---|
| ENSMUSG00000022799 | Arhgap31 | Rho GTPase activating protein 31 | -1.57 |
| ENSMUSG00000063652 | Slc22a21 | solute carrier family 22 (organic cation transporter), member 21 | -1.57 |
| ENSMUSG00000026969 | Fam166a | family with sequence similarity 166, member A | -1.57 |
| ENSMUSG00000051339 | 2900026A02Rik | RIKEN cDNA 2900026A02 gene | -1.58 |
| ENSMUSG00000005373 | Mlxipl | MLX interacting protein-like | -1.58 |
| ENSMUSG00000024867 | Pip5k1b | phosphatidylinositol-4-phosphate 5-kinase, type 1 beta | -1.58 |
| ENSMUSG00000005686 | Ampd3 | adenosine monophosphate deaminase 3 | -1.58 |
| ENSMUSG00000022574 | Naprt | nicotinate phosphoribosyltransferase | -1.58 |
| ENSMUSG00000093695 | Gm20717 | predicted gene 20717 | -1.58 |
| ENSMUSG00000112129 | Pbld1 | phenazine biosynthesis-like protein domain containing 1 | -1.58 |
| ENSMUSG00000003546 | Klc4 | kinesin light chain 4 | -1.59 |
| ENSMUSG00000005615 | Pcyt1a | phosphate cytidylyltransferase 1, choline, alpha isoform | -1.59 |
| ENSMUSG00000025648 | Pfkfb4 | 6-phosphofructo-2-kinase/fructose-2,6-biphosphatase 4 | -1.59 |
| ENSMUSG00000087230 | Mroh3 | maestro heat-like repeat family member 3 | -1.59 |
| ENSMUSG00000112513 | 4930422I22Rik | RIKEN cDNA 4930422I22 gene | -1.59 |
| ENSMUSG00000107760 | Gm44401 | predicted gene, 44401 | -1.59 |
| ENSMUSG00000040134 | Rdh7 | retinol dehydrogenase 7 | -1.59 |
| ENSMUSG00000102283 | Gm37000 | predicted gene, 37000 | -1.59 |
| ENSMUSG00000063730 | Hsd3b2 | hydroxy-delta-5-steroid dehydrogenase, 3 beta- and steroid delta-isomerase 2 | -1.59 |
| ENSMUSG00000084325 | Gm15550 | predicted gene 15550 | -1.59 |
| ENSMUSG00000056973 | Ces1d | carboxylesterase 1D | -1.59 |
| ENSMUSG00000101814 | Gm17807 | predicted gene, 17807 | -1.59 |
| ENSMUSG00000032661 | Oas3 | 2'-5' oligoadenylate synthetase 3 | -1.60 |
| ENSMUSG00000062410 | Hsd3b3 | hydroxy-delta-5-steroid dehydrogenase, 3 beta- and steroid delta-isomerase 3 | -1.60 |
| ENSMUSG00000004151 | Etv1 | ets variant 1 | -1.60 |
| ENSMUSG00000022575 | Gsdmd | gasdermin D | -1.61 |
| ENSMUSG00000026670 | Uap1 | UDP-N-acetylglucosamine pyrophosphorylase 1 | -1.61 |
| ENSMUSG00000028121 | Bcar3 | breast cancer anti-estrogen resistance 3 | -1.61 |
| ENSMUSG00000003271 | Sult2b1 | sulfotransferase family, cytosolic, 2B, member 1 | -1.61 |
| ENSMUSG00000074063 | Osgin1 | oxidative stress induced growth inhibitor 1 | -1.61 |
| ENSMUSG00000044080 | S100a1 | S100 calcium binding protein A1 | -1.61 |
| ENSMUSG00000041372 | B4galnt3 | beta-1,4-N-acetyl-galactosaminyl transferase 3 | -1.61 |
| ENSMUSG00000104571 | Gm43010 | predicted gene 43010 | -1.61 |
| ENSMUSG00000104114 | Gm37297 | predicted gene, 37297 | -1.61 |
| ENSMUSG00000036960 | Clca2 | chloride channel accessory 2 | -1.61 |
| ENSMUSG00000034528 | Hsd17b13 | hydroxysteroid (17-beta) dehydrogenase 13 | -1.62 |
| ENSMUSG00000092474 | Gm20478 | predicted gene 20478 | -1.62 |
| ENSMUSG00000090582 | Gm17024 | predicted gene 17024 | -1.62 |
| ENSMUSG00000032536 | Trak1 | trafficking protein, kinesin binding 1 | -1.63 |
| ENSMUSG00000026896 | Ifih1 | interferon induced with helicase C domain 1 | -1.63 |
| ENSMUSG00000052102 | Gnpda1 | glucosamine-6-phosphate deaminase 1 | -1.63 |
| ENSMUSG00000037649 | H2-DMa | histocompatibility 2, class II, locus DMa | -1.63 |
| ENSMUSG00000112346 | Gm48768 | predicted gene, 48768 | -1.63 |
| ENSMUSG00000048473 | Sult6b2 | sulfotransferase family 6B, member 2 | -1.63 |
| ENSMUSG00000024379 | Tslp | thymic stromal lymphopoietin | -1.63 |
| ENSMUSG00000040712 | Camta2 | calmodulin binding transcription activator 2 | -1.64 |
| ENSMUSG00000028088 | Fmo5 | flavin containing monooxygenase 5 | -1.64 |
| ENSMUSG00000025198 | Erlin1 | ER lipid raft associated 1 | -1.64 |
| ENSMUSG00000064254 | Ethe1 | ethylmalonic encephalopathy 1 | -1.64 |
| ENSMUSG00000033538 | Casp4 | caspase 4, apoptosis-related cysteine peptidase | -1.64 |
| ENSMUSG00000015085 | Entpd2 | ectonucleoside triphosphate diphosphohydrolase 2 | -1.64 |
| ENSMUSG00000006711 | D130043K22Rik | RIKEN cDNA D130043K22 gene | -1.64 |
| ENSMUSG00000082677 | Gm15371 | predicted gene 15371 | -1.64 |
| ENSMUSG00000038188 | Scarf1 | scavenger receptor class F, member 1 | -1.64 |
| ENSMUSG00000071265 | 1700086L19Rik | RIKEN cDNA 1700086L19 gene | -1.64 |
| ENSMUSG00000103391 | Gm38302 | predicted gene, 38302 | -1.64 |
| ENSMUSG00000043501 | Lgals2 | lectin, galactose-binding, soluble 2 | -1.65 |
| ENSMUSG00000029455 | Aldh2 | aldehyde dehydrogenase 2, mitochondrial | -1.65 |
| ENSMUSG00000017057 | Il13ra1 | interleukin 13 receptor, alpha 1 | -1.65 |
| ENSMUSG00000020889 | Nr1d1 | nuclear receptor subfamily 1, group D, member 1 | -1.65 |
| ENSMUSG00000055675 | Kbtbd11 | kelch repeat and BTB (POZ) domain containing 11 | -1.65 |
| ENSMUSG00000051065 | Mb21d2 | Mab-21 domain containing 2 | -1.65 |
| ENSMUSG00000025759 | Mfsd8 | major facilitator superfamily domain containing 8 | -1.65 |
| ENSMUSG00000036473 | Tbc1d24 | TBC1 domain family, member 24 | -1.65 |
| ENSMUSG00000111379 | Gm48346 | predicted gene, 48346 | -1.65 |
| ENSMUSG00000102626 | Gm18553 | predicted gene, 18553 | -1.65 |
| ENSMUSG00000106583 | 4930447N08Rik | RIKEN cDNA 4930447N08 gene | -1.65 |
| ENSMUSG00000034127 | Tspan8 | tetraspanin 8 | -1.66 |
| ENSMUSG00000108291 | Gm44292 | predicted gene, 44292 | -1.66 |
| ENSMUSG00000030790 | Adm | adrenomedullin | -1.66 |
| ENSMUSG00000106541 | Gm43246 | predicted gene 43246 | -1.66 |

| | | | |
|---|---|---|---|
| ENSMUSG00000028017 | Egf | epidermal growth factor | -1.66 |
| ENSMUSG00000027669 | Gnb4 | guanine nucleotide binding protein (G protein), beta 4 | -1.66 |
| ENSMUSG00000049037 | Clec4a1 | C-type lectin domain family 4, member a1 | -1.66 |
| ENSMUSG00000106466 | Gm40348 | predicted gene, 40348 | -1.66 |
| ENSMUSG00000108211 | Gm44130 | predicted gene, 44130 | -1.66 |
| ENSMUSG00000102752 | Gm7694 | predicted gene 7694 | -1.67 |
| ENSMUSG00000053310 | Nrgn | neurogranin | -1.67 |
| ENSMUSG00000001348 | Acp5 | acid phosphatase 5, tartrate resistant | -1.67 |
| ENSMUSG00000037818 | Abhd18 | abhydrolase domain containing 18 | -1.67 |
| ENSMUSG00000090174 | Gm10612 | predicted gene 10612 | -1.67 |
| ENSMUSG00000084329 | Gm6733 | predicted gene 6733 | -1.67 |
| ENSMUSG00000106178 | Gm42987 | predicted gene 42987 | -1.67 |
| ENSMUSG00000049176 | Frmpd4 | FERM and PDZ domain containing 4 | -1.67 |
| ENSMUSG00000021097 | Clmn | calmin | -1.68 |
| ENSMUSG00000032802 | Srxn1 | sulfiredoxin 1 homolog (S. cerevisiae) | -1.68 |
| ENSMUSG00000006651 | Aplp1 | amyloid beta (A4) precursor-like protein 1 | -1.68 |
| ENSMUSG00000045312 | Lhfpl2 | lipoma HMGIC fusion partner-like 2 | -1.68 |
| ENSMUSG00000072214 | Sept5 | septin 5 | -1.68 |
| ENSMUSG00000022853 | Ehhadh | enoyl-Coenzyme A, hydratase/3-hydroxyacyl Coenzyme A dehydrogenase | -1.68 |
| ENSMUSG00000105456 | Gm43745 | predicted gene 43745 | -1.68 |
| ENSMUSG00000071551 | Akr1c19 | aldo-keto reductase family 1, member C19 | -1.68 |
| ENSMUSG00000005360 | Slc1a3 | solute carrier family 1 (glial high affinity glutamate transporter), member 3 | -1.68 |
| ENSMUSG00000038473 | Nos1ap | nitric oxide synthase 1 (neuronal) adaptor protein | -1.69 |
| ENSMUSG00000000958 | Slc7a7 | solute carrier family 7 (cationic amino acid transporter, y+ system), member 7 | -1.69 |
| ENSMUSG00000106475 | Gm43011 | predicted gene 43011 | -1.69 |
| ENSMUSG00000020600 | Slc7a15 | solute carrier family 7 (cationic amino acid transporter, y+ system), member 15 | -1.69 |
| ENSMUSG00000027834 | Serpini1 | serine (or cysteine) peptidase inhibitor, clade I, member 1 | -1.69 |
| ENSMUSG00000041323 | Ak7 | adenylate kinase 7 | -1.69 |
| ENSMUSG00000085095 | Gm15635 | predicted gene 15635 | -1.69 |
| ENSMUSG00000068323 | Slc4a5 | solute carrier family 4, sodium bicarbonate cotransporter, member 5 | -1.69 |
| ENSMUSG00000021250 | Fos | FBJ osteosarcoma oncogene | -1.70 |
| ENSMUSG00000044626 | Liph | lipase, member H | -1.70 |
| ENSMUSG00000024222 | Fkbp5 | FK506 binding protein 5 | -1.70 |
| ENSMUSG00000101517 | 4732465J04Rik | RIKEN cDNA 4732465J04 gene | -1.70 |
| ENSMUSG00000105460 | Gm42671 | predicted gene 42671 | -1.70 |
| ENSMUSG00000049493 | Pls1 | plastin 1 (I-isoform) | -1.71 |
| ENSMUSG00000032120 | C2cd2l | C2 calcium-dependent domain containing 2-like | -1.71 |
| ENSMUSG00000024712 | Rfk | riboflavin kinase | -1.71 |
| ENSMUSG00000025876 | Unc5a | unc-5 netrin receptor A | -1.71 |
| ENSMUSG00000100183 | Gm28512 | predicted gene 28512 | -1.71 |
| ENSMUSG00000058740 | Kcnt1 | potassium channel, subfamily T, member 1 | -1.71 |
| ENSMUSG00000081236 | Gm13574 | predicted gene 13574 | -1.71 |
| ENSMUSG00000047728 | BC025446 | cDNA sequence BC025446 | -1.72 |
| ENSMUSG00000019699 | Akt3 | thymoma viral proto-oncogene 3 | -1.72 |
| ENSMUSG00000026204 | Ptprn | protein tyrosine phosphatase, receptor type, N | -1.72 |
| ENSMUSG00000030228 | Pik3c2g | phosphatidylinositol-4-phosphate 3-kinase catalytic subunit type 2 gamma | -1.72 |
| ENSMUSG00000006281 | Tep1 | telomerase associated protein 1 | -1.73 |
| ENSMUSG00000069515 | Lyz1 | lysozyme 1 | -1.73 |
| ENSMUSG00000020733 | Slc9a3r1 | solute carrier family 9 (sodium/hydrogen exchanger), member 3 regulator 1 | -1.73 |
| ENSMUSG00000054662 | Ano9 | anoctamin 9 | -1.73 |
| ENSMUSG00000035642 | Aamdc | adipogenesis associated Mth938 domain containing | -1.73 |
| ENSMUSG00000039208 | Metrnl | meteorin, glial cell differentiation regulator-like | -1.73 |
| ENSMUSG00000043333 | Rhbdl2 | rhomboid like 2 | -1.73 |
| ENSMUSG00000037455 | Slc18b1 | solute carrier family 18, subfamily B, member 1 | -1.73 |
| ENSMUSG00000001985 | Grik3 | glutamate receptor, ionotropic, kainate 3 | -1.73 |
| ENSMUSG00000024960 | Plcb3 | phospholipase C, beta 3 | -1.74 |
| ENSMUSG00000034781 | Gna11 | guanine nucleotide binding protein, alpha 11 | -1.74 |
| ENSMUSG00000041737 | Tmem45b | transmembrane protein 45b | -1.74 |
| ENSMUSG00000041515 | Irf8 | interferon regulatory factor 8 | -1.74 |
| ENSMUSG00000020407 | Upp1 | uridine phosphorylase 1 | -1.74 |
| ENSMUSG00000043432 | Leng9 | leukocyte receptor cluster (LRC) member 9 | -1.74 |
| ENSMUSG00000037661 | Gpr160 | G protein-coupled receptor 160 | -1.74 |
| ENSMUSG00000052392 | Acot4 | acyl-CoA thioesterase 4 | -1.74 |
| ENSMUSG00000056671 | Prelid2 | PRELI domain containing 2 | -1.74 |
| ENSMUSG00000105872 | Gm9515 | predicted gene 9515 | -1.74 |
| ENSMUSG00000026062 | Slc9a2 | solute carrier family 9 (sodium/hydrogen exchanger), member 2 | -1.75 |
| ENSMUSG00000028518 | Prkaa2 | protein kinase, AMP-activated, alpha 2 catalytic subunit | -1.75 |
| ENSMUSG00000027412 | Lpin3 | lipin 3 | -1.75 |
| ENSMUSG00000074604 | Mgst2 | microsomal glutathione S-transferase 2 | -1.75 |
| ENSMUSG00000048440 | Cyp4f16 | cytochrome P450, family 4, subfamily f, polypeptide 16 | -1.75 |
| ENSMUSG00000004815 | Dgkq | diacylglycerol kinase, theta | -1.75 |

| | | | |
|---|---|---|---|
| ENSMUSG00000028179 | Cth | cystathionase (cystathionine gamma-lyase) | -1.75 |
| ENSMUSG00000002108 | Nr1h3 | nuclear receptor subfamily 1, group H, member 3 | -1.75 |
| ENSMUSG00000031255 | Sytl4 | synaptotagmin-like 4 | -1.75 |
| ENSMUSG00000024471 | Myot | myotilin | -1.75 |
| ENSMUSG00000037605 | Adgrl3 | adhesion G protein-coupled receptor L3 | -1.75 |
| ENSMUSG00000102425 | Gm26616 | predicted gene, 26616 | -1.75 |
| ENSMUSG00000029772 | Ahcyl2 | S-adenosylhomocysteine hydrolase-like 2 | -1.76 |
| ENSMUSG00000061751 | Kalrn | kalirin, RhoGEF kinase | -1.76 |
| ENSMUSG00000026688 | Mgst3 | microsomal glutathione S-transferase 3 | -1.76 |
| ENSMUSG00000046688 | Tifa | TRAF-interacting protein with forkhead-associated domain | -1.76 |
| ENSMUSG00000042428 | Mgat3 | mannoside acetylglucosaminyltransferase 3 | -1.76 |
| ENSMUSG00000027510 | Rbm38 | RNA binding motif protein 38 | -1.76 |
| ENSMUSG00000029919 | Hpgds | hematopoietic prostaglandin D synthase | -1.76 |
| ENSMUSG00000030278 | Cidec | cell death-inducing DFFA-like effector c | -1.76 |
| ENSMUSG00000067813 | Xkr9 | X-linked Kx blood group related 9 | -1.76 |
| ENSMUSG00000038917 | 3930402G23Rik | RIKEN cDNA 3930402G23 gene | -1.76 |
| ENSMUSG00000086479 | Gm16014 | predicted gene 16014 | -1.76 |
| ENSMUSG00000020017 | Hal | histidine ammonia lyase | -1.76 |
| ENSMUSG00000027870 | Hao2 | hydroxyacid oxidase 2 | -1.76 |
| ENSMUSG00000038807 | Rap1gap2 | RAP1 GTPase activating protein 2 | -1.77 |
| ENSMUSG00000018909 | Arrb1 | arrestin, beta 1 | -1.77 |
| ENSMUSG00000059316 | Slc27a4 | solute carrier family 27 (fatty acid transporter), member 4 | -1.77 |
| ENSMUSG00000054469 | Lclat1 | lysocardiolipin acyltransferase 1 | -1.77 |
| ENSMUSG00000021830 | Txndc16 | thioredoxin domain containing 16 | -1.77 |
| ENSMUSG00000022504 | Ciita | class II transactivator | -1.77 |
| ENSMUSG00000067219 | Nipal1 | NIPA-like domain containing 1 | -1.77 |
| ENSMUSG00000034957 | Cebpa | CCAAT/enhancer binding protein (C/EBP), alpha | -1.77 |
| ENSMUSG00000005089 | Slc1a2 | solute carrier family 1 (glial high affinity glutamate transporter), member 2 | -1.77 |
| ENSMUSG00000081219 | Bambi-ps1 | BMP and activin membrane-bound inhibitor, pseudogene (Xenopus laevis) | -1.77 |
| ENSMUSG00000021566 | Slc6a19os | solute carrier family 6 (neurotransmitter transporter), member 19, opposite strand | -1.77 |
| ENSMUSG00000055670 | Zzef1 | zinc finger, ZZ-type with EF hand domain 1 | -1.78 |
| ENSMUSG00000032246 | Calml4 | calmodulin-like 4 | -1.78 |
| ENSMUSG00000025888 | Casp1 | caspase 1 | -1.78 |
| ENSMUSG00000047496 | Rnf152 | ring finger protein 152 | -1.78 |
| ENSMUSG00000063142 | Kcnma1 | potassium large conductance calcium-activated channel, subfamily M, alpha member 1 | -1.78 |
| ENSMUSG00000025726 | Slc28a1 | solute carrier family 28 (sodium-coupled nucleoside transporter), member 1 | -1.78 |
| ENSMUSG00000031444 | F10 | coagulation factor X | -1.78 |
| ENSMUSG00000104291 | A130071D04Rik | RIKEN cDNA A130071D04 gene | -1.78 |
| ENSMUSG00000109244 | Gm44751 | predicted gene 44751 | -1.78 |
| ENSMUSG00000037031 | Tspan15 | tetraspanin 15 | -1.79 |
| ENSMUSG00000106078 | Gm43697 | predicted gene 43697 | -1.79 |
| ENSMUSG00000060208 | Defa17 | defensin, alpha, 17 | -1.79 |
| ENSMUSG00000034115 | Scn11a | sodium channel, voltage-gated, type XI, alpha | -1.79 |
| ENSMUSG00000040600 | Eps8l3 | EPS8-like 3 | -1.80 |
| ENSMUSG00000067297 | Ifit1bl2 | interferon induced protein with tetratricopeptide repeats 1B like 2 | -1.80 |
| ENSMUSG00000064262 | Gimap8 | GTPase, IMAP family member 8 | -1.80 |
| ENSMUSG00000106933 | Gm43621 | predicted gene 43621 | -1.80 |
| ENSMUSG00000031364 | Grpr | gastrin releasing peptide receptor | -1.80 |
| ENSMUSG00000061171 | Slc38a11 | solute carrier family 38, member 11 | -1.80 |
| ENSMUSG00000107624 | Gm44005 | predicted gene, 44005 | -1.80 |
| ENSMUSG00000005640 | Insrr | insulin receptor-related receptor | -1.80 |
| ENSMUSG00000025076 | Casp7 | caspase 7 | -1.81 |
| ENSMUSG00000021062 | Rab15 | RAB15, member RAS oncogene family | -1.81 |
| ENSMUSG00000017868 | Sgk2 | serum/glucocorticoid regulated kinase 2 | -1.81 |
| ENSMUSG00000089900 | Tbc1d22bos | TBC1 domain family, member 22B, opposite strand | -1.81 |
| ENSMUSG00000106186 | Gm43627 | predicted gene 43627 | -1.81 |
| ENSMUSG00000009772 | Nuak2 | NUAK family, SNF1-like kinase, 2 | -1.82 |
| ENSMUSG00000034714 | Ttyh2 | tweety family member 2 | -1.82 |
| ENSMUSG00000021280 | Exoc3l4 | exocyst complex component 3-like 4 | -1.82 |
| ENSMUSG00000052131 | Akr1b7 | aldo-keto reductase family 1, member B7 | -1.82 |
| ENSMUSG00000021221 | Dpf3 | D4, zinc and double PHD fingers, family 3 | -1.82 |
| ENSMUSG00000074628 | Tldc2 | TBC/LysM associated domain containing 2 | -1.82 |
| ENSMUSG00000099025 | Gm27162 | predicted gene 27162 | -1.82 |
| ENSMUSG00000035561 | Aldh1b1 | aldehyde dehydrogenase 1 family, member B1 | -1.83 |
| ENSMUSG00000024644 | Cndp2 | CNDP dipeptidase 2 (metallopeptidase M20 family) | -1.83 |
| ENSMUSG00000061411 | Nol4l | nucleolar protein 4-like | -1.83 |
| ENSMUSG00000025993 | Slc40a1 | solute carrier family 40 (iron-regulated transporter), member 1 | -1.83 |
| ENSMUSG00000026880 | Stom | stomatin | -1.83 |
| ENSMUSG00000052085 | Dock8 | dedicator of cytokinesis 8 | -1.83 |
| ENSMUSG00000048572 | Tmem252 | transmembrane protein 252 | -1.83 |
| ENSMUSG00000051452 | Gm11437 | predicted gene 11437 | -1.83 |

| | | | |
|---|---|---|---|
| ENSMUSG00000011034 | Slc5a1 | solute carrier family 5 (sodium/glucose cotransporter), member 1 | -1.83 |
| ENSMUSG00000044017 | Adgrd1 | adhesion G protein-coupled receptor D1 | -1.83 |
| ENSMUSG00000101624 | Gm28882 | predicted gene 28882 | -1.83 |
| ENSMUSG00000050762 | Prss27 | protease, serine 27 | -1.83 |
| ENSMUSG00000108850 | Gm44914 | predicted gene 44914 | -1.83 |
| ENSMUSG00000021596 | Mctp1 | multiple C2 domains, transmembrane 1 | -1.83 |
| ENSMUSG00000072244 | Trim6 | tripartite motif-containing 6 | -1.83 |
| ENSMUSG00000001156 | Mxd1 | MAX dimerization protein 1 | -1.84 |
| ENSMUSG00000028943 | Espn | espin | -1.84 |
| ENSMUSG00000057069 | Ero1lb | ERO1-like beta (S. cerevisiae) | -1.84 |
| ENSMUSG00000045871 | Slitrk6 | SLIT and NTRK-like family, member 6 | -1.84 |
| ENSMUSG00000055963 | Triqk | triple QxxK/R motif containing | -1.84 |
| ENSMUSG00000028427 | Aqp7 | aquaporin 7 | -1.84 |
| ENSMUSG00000073405 | H2-T-ps | histocompatibility 2, T region locus, pseudogene | -1.84 |
| ENSMUSG00000038372 | Gmds | GDP-mannose 4, 6-dehydratase | -1.85 |
| ENSMUSG00000032899 | Styk1 | serine/threonine/tyrosine kinase 1 | -1.85 |
| ENSMUSG00000030281 | Il17rc | interleukin 17 receptor C | -1.85 |
| ENSMUSG00000060002 | Chpt1 | choline phosphotransferase 1 | -1.85 |
| ENSMUSG00000027358 | Bmp2 | bone morphogenetic protein 2 | -1.85 |
| ENSMUSG00000027219 | Slc28a2 | solute carrier family 28 (sodium-coupled nucleoside transporter), member 2 | -1.85 |
| ENSMUSG00000028262 | Clca3a2 | chloride channel accessory 3A2 | -1.85 |
| ENSMUSG00000097099 | Gm9917 | predicted gene 9917 | -1.85 |
| ENSMUSG00000028544 | Slc5a9 | solute carrier family 5 (sodium/glucose cotransporter), member 9 | -1.85 |
| ENSMUSG00000110266 | Gm32742 | predicted gene, 32742 | -1.85 |
| ENSMUSG00000030621 | Me3 | malic enzyme 3, NADP(+)-dependent, mitochondrial | -1.85 |
| ENSMUSG00000041617 | Ccdc74a | coiled-coil domain containing 74A | -1.85 |
| ENSMUSG00000021337 | Scgn | secretagogin, EF-hand calcium binding protein | -1.85 |
| ENSMUSG00000017950 | Hnf4a | hepatic nuclear factor 4, alpha | -1.86 |
| ENSMUSG00000048905 | 4930539E08Rik | RIKEN cDNA 4930539E08 gene | -1.86 |
| ENSMUSG00000063522 | 2010109I03Rik | RIKEN cDNA 2010109I03 gene | -1.86 |
| ENSMUSG00000090214 | Gm15657 | predicted gene 15657 | -1.86 |
| ENSMUSG00000034765 | Dusp5 | dual specificity phosphatase 5 | -1.86 |
| ENSMUSG00000074892 | B3galt5 | UDP-Gal:betaGlcNAc beta 1,3-galactosyltransferase, polypeptide 5 | -1.86 |
| ENSMUSG00000054150 | Syne3 | spectrin repeat containing, nuclear envelope family member 3 | -1.86 |
| ENSMUSG00000110520 | Gm45776 | predicted gene 45776 | -1.86 |
| ENSMUSG00000110735 | Gm47173 | predicted gene, 47173 | -1.86 |
| ENSMUSG00000105986 | Gm43065 | predicted gene 43065 | -1.86 |
| ENSMUSG00000030838 | Ush1c | USH1 protein network component harmonin | -1.87 |
| ENSMUSG00000031442 | Mcf2l | mcf.2 transforming sequence-like | -1.87 |
| ENSMUSG00000024066 | Xdh | xanthine dehydrogenase | -1.87 |
| ENSMUSG00000030208 | Emp1 | epithelial membrane protein 1 | -1.87 |
| ENSMUSG00000028536 | 2610528J11Rik | RIKEN cDNA 2610528J11 gene | -1.87 |
| ENSMUSG00000020709 | Adap2 | ArfGAP with dual PH domains 2 | -1.87 |
| ENSMUSG00000104316 | Gm37909 | predicted gene, 37909 | -1.87 |
| ENSMUSG00000031129 | Slc9a9 | solute carrier family 9 (sodium/hydrogen exchanger), member 9 | -1.87 |
| ENSMUSG00000035186 | Ubd | ubiquitin D | -1.87 |
| ENSMUSG00000114705 | Gm18760 | predicted gene, 18760 | -1.87 |
| ENSMUSG00000031608 | Galnt7 | polypeptide N-acetylgalactosaminyltransferase 7 | -1.88 |
| ENSMUSG00000026175 | Vil1 | villin 1 | -1.88 |
| ENSMUSG00000064213 | Defa24 | defensin, alpha, 24 | -1.88 |
| ENSMUSG00000039910 | Cited2 | Cbp/p300-interacting transactivator, with Glu/Asp-rich carboxy-terminal domain, 2 | -1.88 |
| ENSMUSG00000029727 | Cyp3a13 | cytochrome P450, family 3, subfamily a, polypeptide 13 | -1.88 |
| ENSMUSG00000020272 | Stk10 | serine/threonine kinase 10 | -1.88 |
| ENSMUSG00000032776 | Mctp2 | multiple C2 domains, transmembrane 2 | -1.88 |
| ENSMUSG00000022040 | Ephx2 | epoxide hydrolase 2, cytoplasmic | -1.88 |
| ENSMUSG00000024225 | Clps | colipase, pancreatic | -1.88 |
| ENSMUSG00000089960 | Ugt1a1 | UDP glucuronosyltransferase 1 family, polypeptide A1 | -1.88 |
| ENSMUSG00000003032 | Klf4 | Kruppel-like factor 4 (gut) | -1.89 |
| ENSMUSG00000029561 | Oasl2 | 2'-5' oligoadenylate synthetase-like 2 | -1.89 |
| ENSMUSG00000048373 | Fgfbp1 | fibroblast growth factor binding protein 1 | -1.89 |
| ENSMUSG00000022018 | Rgcc | regulator of cell cycle | -1.89 |
| ENSMUSG00000029490 | Mfsd7a | major facilitator superfamily domain containing 7A | -1.89 |
| ENSMUSG00000029059 | Fam213b | family with sequence similarity 213, member B | -1.90 |
| ENSMUSG00000037003 | Tns2 | tensin 2 | -1.90 |
| ENSMUSG00000052135 | Foxo6 | forkhead box O6 | -1.90 |
| ENSMUSG00000047878 | A4galt | alpha 1,4-galactosyltransferase | -1.90 |
| ENSMUSG00000029802 | Abcg2 | ATP binding cassette subfamily G member 2 (Junior blood group) | -1.90 |
| ENSMUSG00000037451 | Slc22a20 | solute carrier family 22 (organic anion transporter), member 20 | -1.90 |
| ENSMUSG00000045281 | Gpr20 | G protein-coupled receptor 20 | -1.90 |
| ENSMUSG00000029992 | Gfpt1 | glutamine fructose-6-phosphate transaminase 1 | -1.91 |
| ENSMUSG00000054161 | Fam83e | family with sequence similarity 83, member E | -1.91 |

| | | | |
|---|---|---|---|
| ENSMUSG00000024180 | Tmem8 | transmembrane protein 8 (five membrane-spanning domains) | -1.91 |
| ENSMUSG00000022836 | Mylk | myosin, light polypeptide kinase | -1.91 |
| ENSMUSG00000043079 | Synpo | synaptopodin | -1.91 |
| ENSMUSG00000028194 | Ddah1 | dimethylarginine dimethylaminohydrolase 1 | -1.91 |
| ENSMUSG00000067206 | Lrrc66 | leucine rich repeat containing 66 | -1.91 |
| ENSMUSG00000057596 | Trim30d | tripartite motif-containing 30D | -1.91 |
| ENSMUSG00000022340 | Sybu | syntabulin (syntaxin-interacting) | -1.91 |
| ENSMUSG00000022445 | Cyp2d26 | cytochrome P450, family 2, subfamily d, polypeptide 26 | -1.91 |
| ENSMUSG00000028699 | Tspan1 | tetraspanin 1 | -1.92 |
| ENSMUSG00000021336 | Slc17a4 | solute carrier family 17 (sodium phosphate), member 4 | -1.92 |
| ENSMUSG00000041120 | Nbl1 | neuroblastoma, suppression of tumorigenicity 1 | -1.92 |
| ENSMUSG00000030214 | Plbd1 | phospholipase B domain containing 1 | -1.92 |
| ENSMUSG00000028364 | Tnc | tenascin C | -1.92 |
| ENSMUSG00000054453 | Sytl5 | synaptotagmin-like 5 | -1.92 |
| ENSMUSG00000022947 | Cbr3 | carbonyl reductase 3 | -1.92 |
| ENSMUSG00000015401 | Cltrn | collectrin, amino acid transport regulator | -1.92 |
| ENSMUSG00000034320 | Slc26a2 | solute carrier family 26 (sulfate transporter), member 2 | -1.93 |
| ENSMUSG00000031838 | Ifi30 | interferon gamma inducible protein 30 | -1.93 |
| ENSMUSG00000025429 | Pstpip2 | proline-serine-threonine phosphatase-interacting protein 2 | -1.93 |
| ENSMUSG00000035258 | Abi3bp | ABI gene family, member 3 (NESH) binding protein | -1.93 |
| ENSMUSG00000032690 | Oas2 | 2'-5' oligoadenylate synthetase 2 | -1.93 |
| ENSMUSG00000040537 | Adam22 | a disintegrin and metallopeptidase domain 22 | -1.93 |
| ENSMUSG00000069456 | Rdh16 | retinol dehydrogenase 16 | -1.93 |
| ENSMUSG00000061762 | Tac1 | tachykinin 1 | -1.93 |
| ENSMUSG00000005045 | Chd5 | chromodomain helicase DNA binding protein 5 | -1.93 |
| ENSMUSG00000019359 | Gdpd2 | glycerophosphodiester phosphodiesterase domain containing 2 | -1.93 |
| ENSMUSG00000004864 | Mapk13 | mitogen-activated protein kinase 13 | -1.94 |
| ENSMUSG00000057963 | Itpk1 | inositol 1,3,4-triphosphate 5/6 kinase | -1.94 |
| ENSMUSG00000024642 | Tle4 | transducin-like enhancer of split 4 | -1.94 |
| ENSMUSG00000005107 | Slc2a9 | solute carrier family 2 (facilitated glucose transporter), member 9 | -1.94 |
| ENSMUSG00000023827 | Agpat4 | 1-acylglycerol-3-phosphate O-acyltransferase 4 (lysophosphatidic acid acyltransferase, delta) | -1.94 |
| ENSMUSG00000040298 | Btbd16 | BTB (POZ) domain containing 16 | -1.94 |
| ENSMUSG00000084948 | 1700061H18Rik | RIKEN cDNA 1700061H18 gene | -1.94 |
| ENSMUSG00000029322 | Plac8 | placenta-specific 8 | -1.95 |
| ENSMUSG00000025498 | Irf7 | interferon regulatory factor 7 | -1.95 |
| ENSMUSG00000020681 | Ace | angiotensin I converting enzyme (peptidyl-dipeptidase A) 1 | -1.95 |
| ENSMUSG00000038534 | Osbpl7 | oxysterol binding protein-like 7 | -1.95 |
| ENSMUSG00000103527 | Gm37261 | predicted gene, 37261 | -1.95 |
| ENSMUSG00000019989 | Enpp3 | ectonucleotide pyrophosphatase/phosphodiesterase 3 | -1.95 |
| ENSMUSG00000086844 | B230206H07Rik | RIKEN cDNA B230206H07 gene | -1.95 |
| ENSMUSG00000106108 | Gm43221 | predicted gene 43221 | -1.95 |
| ENSMUSG00000115388 | Eppk1 | epiplakin 1 | -1.96 |
| ENSMUSG00000038178 | Slc43a2 | solute carrier family 43, member 2 | -1.96 |
| ENSMUSG00000053279 | Aldh1a1 | aldehyde dehydrogenase family 1, subfamily A1 | -1.96 |
| ENSMUSG00000000204 | Slfn4 | schlafen 4 | -1.96 |
| ENSMUSG00000052271 | Bhlha15 | basic helix-loop-helix family, member a15 | -1.96 |
| ENSMUSG00000023206 | Il15ra | interleukin 15 receptor, alpha chain | -1.96 |
| ENSMUSG00000090608 | Gm17200 | predicted gene 17200 | -1.96 |
| ENSMUSG00000110597 | Gm8798 | predicted gene 8798 | -1.96 |
| ENSMUSG00000017978 | Cadps2 | Ca2+-dependent activator protein for secretion 2 | -1.97 |
| ENSMUSG00000018217 | Pmp22 | peripheral myelin protein 22 | -1.97 |
| ENSMUSG00000057897 | Camk2b | calcium/calmodulin-dependent protein kinase II, beta | -1.97 |
| ENSMUSG00000051379 | Flrt3 | fibronectin leucine rich transmembrane protein 3 | -1.97 |
| ENSMUSG00000025002 | Cyp2c55 | cytochrome P450, family 2, subfamily c, polypeptide 55 | -1.97 |
| ENSMUSG00000010122 | Slc47a1 | solute carrier family 47, member 1 | -1.97 |
| ENSMUSG00000037157 | Il22ra1 | interleukin 22 receptor, alpha 1 | -1.98 |
| ENSMUSG00000034687 | Fras1 | Fraser extracellular matrix complex subunit 1 | -1.98 |
| ENSMUSG00000041261 | Car8 | carbonic anhydrase 8 | -1.98 |
| ENSMUSG00000085747 | Slc13a2os | solute carrier family 13 (sodium-dependent dicarboxylate transporter), member 2, opposite strand | -1.98 |
| ENSMUSG00000017718 | Afmid | arylformamidase | -1.98 |
| ENSMUSG00000056032 | BC018473 | cDNA sequence BC018473 | -1.98 |
| ENSMUSG00000032377 | Plscr4 | phospholipid scramblase 4 | -1.98 |
| ENSMUSG00000116010 | Gm36026 | predicted gene, 36026 | -1.98 |
| ENSMUSG00000103553 | Gm38218 | predicted gene, 38218 | -1.98 |
| ENSMUSG00000030909 | Anks4b | ankyrin repeat and sterile alpha motif domain containing 4B | -1.99 |
| ENSMUSG00000074445 | Sprr2a3 | small proline-rich protein 2A3 | -1.99 |
| ENSMUSG00000046410 | Kcnk6 | potassium inwardly-rectifying channel, subfamily K, member 6 | -1.99 |
| ENSMUSG00000038963 | Slco4a1 | solute carrier organic anion transporter family, member 4a1 | -1.99 |
| ENSMUSG00000039981 | Zc3h12d | zinc finger CCCH type containing 12D | -1.99 |
| ENSMUSG00000104966 | Gm43273 | predicted gene 43273 | -1.99 |
| ENSMUSG00000038037 | Socs1 | suppressor of cytokine signaling 1 | -1.99 |

| | | | |
|---|---|---|---|
| ENSMUSG00000090145 | Ugt1a6b | UDP glucuronosyltransferase 1 family, polypeptide A6B | -1.99 |
| ENSMUSG00000078964 | Ces1b | carboxylesterase 1B | -1.99 |
| ENSMUSG00000018126 | Baiap2l2 | BAI1-associated protein 2-like 2 | -2.00 |
| ENSMUSG00000060733 | Ipmk | inositol polyphosphate multikinase | -2.00 |
| ENSMUSG00000025357 | Dgka | diacylglycerol kinase, alpha | -2.00 |
| ENSMUSG00000045038 | Prkce | protein kinase C, epsilon | -2.00 |
| ENSMUSG00000022555 | Dgat1 | diacylglycerol O-acyltransferase 1 | -2.00 |
| ENSMUSG00000028635 | Edn2 | endothelin 2 | -2.00 |
| ENSMUSG00000085941 | Gm11201 | predicted gene 11201 | -2.00 |
| ENSMUSG00000006731 | B4galnt1 | beta-1,4-N-acetyl-galactosaminyl transferase 1 | -2.01 |
| ENSMUSG00000024937 | Ehbp1l1 | EH domain binding protein 1-like 1 | -2.01 |
| ENSMUSG00000023243 | Kcnk5 | potassium channel, subfamily K, member 5 | -2.01 |
| ENSMUSG00000079547 | H2-DMb1 | histocompatibility 2, class II, locus Mb1 | -2.01 |
| ENSMUSG00000090066 | 1110002E22Rik | RIKEN cDNA 1110002E22 gene | -2.01 |
| ENSMUSG00000055027 | Smyd1 | SET and MYND domain containing 1 | -2.01 |
| ENSMUSG00000021624 | Cd180 | CD180 antigen | -2.01 |
| ENSMUSG00000068587 | Mgam | maltase-glucoamylase | -2.02 |
| ENSMUSG00000007034 | Slc44a4 | solute carrier family 44, member 4 | -2.02 |
| ENSMUSG00000019970 | Sgk1 | serum/glucocorticoid regulated kinase 1 | -2.02 |
| ENSMUSG00000074622 | Mafb | v-maf musculoaponeurotic fibrosarcoma oncogene family, protein B (avian) | -2.02 |
| ENSMUSG00000043629 | 1700019D03Rik | RIKEN cDNA 1700019D03 gene | -2.02 |
| ENSMUSG00000103854 | Gm37250 | predicted gene, 37250 | -2.02 |
| ENSMUSG00000079242 | C730034F03Rik | RIKEN cDNA C730034F03 gene | -2.02 |
| ENSMUSG00000021456 | Fbp2 | fructose bisphosphatase 2 | -2.03 |
| ENSMUSG00000034570 | Inpp5j | inositol polyphosphate 5-phosphatase J | -2.03 |
| ENSMUSG00000024743 | Syt7 | synaptotagmin VII | -2.03 |
| ENSMUSG00000078945 | Naip2 | NLR family, apoptosis inhibitory protein 2 | -2.03 |
| ENSMUSG00000042041 | 2010003K11Rik | RIKEN cDNA 2010003K11 gene | -2.03 |
| ENSMUSG00000024697 | Gna14 | guanine nucleotide binding protein, alpha 14 | -2.03 |
| ENSMUSG00000023057 | Fabp2 | fatty acid binding protein 2, intestinal | -2.03 |
| ENSMUSG00000082148 | Gm12266 | predicted gene 12266 | -2.03 |
| ENSMUSG00000075517 | Cyp2d37-ps | cytochrome P450, family 2, subfamily d, polypeptide 37, pseudogene | -2.03 |
| ENSMUSG00000028337 | Coro2a | coronin, actin binding protein 2A | -2.04 |
| ENSMUSG00000034116 | Vav1 | vav 1 oncogene | -2.04 |
| ENSMUSG00000027452 | Acss1 | acyl-CoA synthetase short-chain family member 1 | -2.04 |
| ENSMUSG00000020614 | Fam20a | family with sequence similarity 20, member A | -2.04 |
| ENSMUSG00000020774 | Aspa | aspartoacylase | -2.04 |
| ENSMUSG00000020105 | Lrig3 | leucine-rich repeats and immunoglobulin-like domains 3 | -2.04 |
| ENSMUSG00000038020 | Rapgefl1 | Rap guanine nucleotide exchange factor (GEF)-like 1 | -2.04 |
| ENSMUSG00000049799 | Lrrc19 | leucine rich repeat containing 19 | -2.04 |
| ENSMUSG00000072571 | Tmem253 | transmembrane protein 253 | -2.04 |
| ENSMUSG00000030683 | Sez6l2 | seizure related 6 homolog like 2 | -2.04 |
| ENSMUSG00000085196 | Gm14963 | predicted gene 14963 | -2.04 |
| ENSMUSG00000029811 | Aoc1 | amine oxidase, copper-containing 1 | -2.05 |
| ENSMUSG00000032033 | Barx2 | BarH-like homeobox 2 | -2.05 |
| ENSMUSG00000035041 | Creb3l3 | cAMP responsive element binding protein 3-like 3 | -2.05 |
| ENSMUSG00000038349 | Plcl1 | phospholipase C-like 1 | -2.05 |
| ENSMUSG00000101581 | C430002N11Rik | RIKEN cDNA C430002N11 gene | -2.05 |
| ENSMUSG00000029821 | Gsdme | gasdermin E | -2.06 |
| ENSMUSG00000027227 | Sord | sorbitol dehydrogenase | -2.06 |
| ENSMUSG00000020826 | Nos2 | nitric oxide synthase 2, inducible | -2.06 |
| ENSMUSG00000020151 | Ptprr | protein tyrosine phosphatase, receptor type, R | -2.06 |
| ENSMUSG00000025467 | Prap1 | proline-rich acidic protein 1 | -2.06 |
| ENSMUSG00000035967 | Ints6l | integrator complex subunit 6 like | -2.06 |
| ENSMUSG00000015970 | Chdh | choline dehydrogenase | -2.08 |
| ENSMUSG00000066800 | Rnasel | ribonuclease L (2', 5'-oligoisoadenylate synthetase-dependent) | -2.08 |
| ENSMUSG00000026726 | Cubn | cubilin (intrinsic factor-cobalamin receptor) | -2.08 |
| ENSMUSG00000022938 | Fam3b | family with sequence similarity 3, member B | -2.08 |
| ENSMUSG00000079339 | Ifit1bl1 | interferon induced protein with tetratricpeptide repeats 1B like 1 | -2.08 |
| ENSMUSG00000026489 | Coq8a | coenzyme Q8A | -2.08 |
| ENSMUSG00000050106 | Tmc8 | transmembrane channel-like gene family 8 | -2.08 |
| ENSMUSG00000026692 | Fmo4 | flavin containing monooxygenase 4 | -2.08 |
| ENSMUSG00000092517 | Art2a-ps | ADP-ribosyltransferase 2a, pseudogene | -2.08 |
| ENSMUSG00000043673 | Kcns3 | potassium voltage-gated channel, delayed-rectifier, subfamily S, member 3 | -2.08 |
| ENSMUSG00000082485 | Gm15693 | predicted gene 15693 | -2.08 |
| ENSMUSG00000041268 | Dmxl2 | Dmx-like 2 | -2.09 |
| ENSMUSG00000036594 | H2-Aa | histocompatibility 2, class II antigen A, alpha | -2.09 |
| ENSMUSG00000032373 | Car12 | carbonic anhydrase 12 | -2.09 |
| ENSMUSG00000020447 | Npc1l1 | NPC1 like intracellular cholesterol transporter 1 | -2.09 |
| ENSMUSG00000032083 | Apoa1 | apolipoprotein A-I | -2.09 |
| ENSMUSG00000075014 | Gm10800 | predicted gene 10800 | -2.09 |

| | | | |
|---|---|---|---|
| ENSMUSG00000040584 | Abcb1a | ATP-binding cassette, sub-family B (MDR/TAP), member 1A | -2.10 |
| ENSMUSG00000024619 | Cdx1 | caudal type homeobox 1 | -2.10 |
| ENSMUSG00000013523 | Bcas1 | breast carcinoma amplified sequence 1 | -2.10 |
| ENSMUSG00000034413 | Neurl1b | neuralized E3 ubiquitin protein ligase 1B | -2.10 |
| ENSMUSG00000026077 | Npas2 | neuronal PAS domain protein 2 | -2.10 |
| ENSMUSG00000041741 | Pde3a | phosphodiesterase 3A, cGMP inhibited | -2.10 |
| ENSMUSG00000040767 | Snrnp25 | small nuclear ribonucleoprotein 25 (U11/U12) | -2.10 |
| ENSMUSG00000027508 | Pag1 | phosphoprotein associated with glycosphingolipid microdomains 1 | -2.10 |
| ENSMUSG00000042312 | S100a13 | S100 calcium binding protein A13 | -2.10 |
| ENSMUSG00000052415 | Tchh | trichohyalin | -2.10 |
| ENSMUSG00000039699 | Batf2 | basic leucine zipper transcription factor, ATF-like 2 | -2.10 |
| ENSMUSG00000000805 | Car4 | carbonic anhydrase 4 | -2.10 |
| ENSMUSG00000004113 | Cacna1b | calcium channel, voltage-dependent, N type, alpha 1B subunit | -2.10 |
| ENSMUSG00000020405 | Fabp6 | fatty acid binding protein 6, ileal (gastrotropin) | -2.10 |
| ENSMUSG00000079588 | Tmem182 | transmembrane protein 182 | -2.10 |
| ENSMUSG00000030650 | Tmc5 | transmembrane channel-like gene family 5 | -2.11 |
| ENSMUSG00000021196 | Pfkp | phosphofructokinase, platelet | -2.11 |
| ENSMUSG00000017639 | Rab11fip4 | RAB11 family interacting protein 4 (class II) | -2.11 |
| ENSMUSG00000022747 | St3gal6 | ST3 beta-galactoside alpha-2,3-sialyltransferase 6 | -2.11 |
| ENSMUSG00000039578 | Ccser1 | coiled-coil serine rich 1 | -2.11 |
| ENSMUSG00000104350 | Gm38244 | predicted gene, 38244 | -2.11 |
| ENSMUSG00000029082 | Bst1 | bone marrow stromal cell antigen 1 | -2.11 |
| ENSMUSG00000025037 | Maoa | monoamine oxidase A | -2.12 |
| ENSMUSG00000052595 | A1cf | APOBEC1 complementation factor | -2.12 |
| ENSMUSG00000068547 | Clca4a | chloride channel accessory 4A | -2.12 |
| ENSMUSG00000096001 | 2610528A11Rik | RIKEN cDNA 2610528A11 gene | -2.12 |
| ENSMUSG00000028158 | Mttp | microsomal triglyceride transfer protein | -2.12 |
| ENSMUSG00000027230 | Creb3l1 | cAMP responsive element binding protein 3-like 1 | -2.13 |
| ENSMUSG00000041642 | Kif21b | kinesin family member 21B | -2.13 |
| ENSMUSG00000050747 | Trim15 | tripartite motif-containing 15 | -2.13 |
| ENSMUSG00000003420 | Fcgrt | Fc receptor, IgG, alpha chain transporter | -2.13 |
| ENSMUSG00000023045 | Soat2 | sterol O-acyltransferase 2 | -2.13 |
| ENSMUSG00000032105 | Pdzd3 | PDZ domain containing 3 | -2.13 |
| ENSMUSG00000009646 | Pla2g12b | phospholipase A2, group XIIB | -2.13 |
| ENSMUSG00000000125 | Wnt3 | wingless-type MMTV integration site family, member 3 | -2.13 |
| ENSMUSG00000097254 | C430042M11Rik | RIKEN cDNA C430042M11 gene | -2.13 |
| ENSMUSG00000041644 | Slc5a12 | solute carrier family 5 (sodium/glucose cotransporter), member 12 | -2.13 |
| ENSMUSG00000026417 | Pigr | polymeric immunoglobulin receptor | -2.14 |
| ENSMUSG00000024747 | Aldh1a7 | aldehyde dehydrogenase family 1, subfamily A7 | -2.14 |
| ENSMUSG00000019935 | Slc17a8 | solute carrier family 17 (sodium-dependent inorganic phosphate cotransporter), member 8 | -2.14 |
| ENSMUSG00000030587 | 2200002D01Rik | RIKEN cDNA 2200002D01 gene | -2.15 |
| ENSMUSG00000022824 | Muc13 | mucin 13, epithelial transmembrane | -2.15 |
| ENSMUSG00000024254 | Abcg8 | ATP binding cassette subfamily G member 8 | -2.15 |
| ENSMUSG00000056413 | Adap1 | ArfGAP with dual PH domains 1 | -2.15 |
| ENSMUSG00000108688 | Gm44985 | predicted gene 44985 | -2.15 |
| ENSMUSG00000015405 | Ace2 | angiotensin I converting enzyme (peptidyl-dipeptidase A) 2 | -2.15 |
| ENSMUSG00000060586 | H2-Eb1 | histocompatibility 2, class II antigen E beta | -2.16 |
| ENSMUSG00000073802 | Cdkn2b | cyclin dependent kinase inhibitor 2B | -2.16 |
| ENSMUSG00000034919 | Ttc22 | tetratricopeptide repeat domain 22 | -2.16 |
| ENSMUSG00000039405 | Prss23 | protease, serine 23 | -2.16 |
| ENSMUSG00000031089 | Slc6a14 | solute carrier family 6 (neurotransmitter transporter), member 14 | -2.16 |
| ENSMUSG00000079445 | B3gnt7 | UDP-GlcNAc:betaGal beta-1,3-N-acetylglucosaminyltransferase 7 | -2.16 |
| ENSMUSG00000075611 | Gm11545 | predicted gene 11545 | -2.16 |
| ENSMUSG00000097779 | 4833407H14Rik | RIKEN cDNA 4833407H14 gene | -2.16 |
| ENSMUSG00000030786 | Itgam | integrin alpha M | -2.16 |
| ENSMUSG00000096914 | Galntl6 | UDP-N-acetyl-alpha-D-galactosamine:polypeptide N-acetylgalactosaminyltransferase-like 6 | -2.16 |
| ENSMUSG00000004266 | Ptpn6 | protein tyrosine phosphatase, non-receptor type 6 | -2.17 |
| ENSMUSG00000037408 | Cnnm4 | cyclin M4 | -2.17 |
| ENSMUSG00000073421 | H2-Ab1 | histocompatibility 2, class II antigen A, beta 1 | -2.17 |
| ENSMUSG00000043592 | Unc5cl | unc-5 family C-terminal like | -2.17 |
| ENSMUSG00000037686 | Aspg | asparaginase | -2.17 |
| ENSMUSG00000100927 | Gm28536 | predicted gene 28536 | -2.17 |
| ENSMUSG00000106191 | 4930557B06Rik | RIKEN cDNA 4930557B06 gene | -2.17 |
| ENSMUSG00000072875 | Gpr27 | G protein-coupled receptor 27 | -2.17 |
| ENSMUSG00000050965 | Prkca | protein kinase C, alpha | -2.18 |
| ENSMUSG00000030866 | Ern2 | endoplasmic reticulum (ER) to nucleus signalling 2 | -2.18 |
| ENSMUSG00000023073 | Slc10a2 | solute carrier family 10, member 2 | -2.18 |
| ENSMUSG00000055435 | Maf | avian musculoaponeurotic fibrosarcoma oncogene homolog | -2.18 |
| ENSMUSG00000012123 | Crybg2 | crystallin beta-gamma domain containing 2 | -2.18 |
| ENSMUSG00000031209 | Heph | hephaestin | -2.18 |
| ENSMUSG00000027463 | Slc52a3 | solute carrier protein family 52, member 3 | -2.18 |

| | | | |
|---|---|---|---|
| ENSMUSG00000028278 | Rragd | Ras-related GTP binding D | -2.18 |
| ENSMUSG00000040249 | Lrp1 | low density lipoprotein receptor-related protein 1 | -2.19 |
| ENSMUSG00000045316 | Fahd1 | fumarylacetoacetate hydrolase domain containing 1 | -2.19 |
| ENSMUSG00000025557 | Slc15a1 | solute carrier family 15 (oligopeptide transporter), member 1 | -2.19 |
| ENSMUSG00000091956 | C2cd4b | C2 calcium-dependent domain containing 4B | -2.19 |
| ENSMUSG00000029409 | U90926 | cDNA sequence U90926 | -2.19 |
| ENSMUSG00000107425 | Gm44216 | predicted gene, 44216 | -2.19 |
| ENSMUSG00000066687 | Zbtb16 | zinc finger and BTB domain containing 16 | -2.19 |
| ENSMUSG00000102802 | Mgam2-ps | maltase-glucoamylase 2, pseudogene | -2.20 |
| ENSMUSG00000030793 | Pycard | PYD and CARD domain containing | -2.20 |
| ENSMUSG00000024277 | Mapre2 | microtubule-associated protein, RP/EB family, member 2 | -2.20 |
| ENSMUSG00000024727 | Trpm6 | transient receptor potential cation channel, subfamily M, member 6 | -2.20 |
| ENSMUSG00000033871 | Ppargc1b | peroxisome proliferative activated receptor, gamma, coactivator 1 beta | -2.20 |
| ENSMUSG00000061825 | Ces2c | carboxylesterase 2C | -2.20 |
| ENSMUSG00000026222 | Sp100 | nuclear antigen Sp100 | -2.20 |
| ENSMUSG00000093867 | Gsdmcl2 | gasdermin C-like 2 | -2.20 |
| ENSMUSG00000047115 | Fam221a | family with sequence similarity 221, member A | -2.20 |
| ENSMUSG00000022219 | Cideb | cell death-inducing DNA fragmentation factor, alpha subunit-like effector B | -2.20 |
| ENSMUSG00000047414 | Flrt2 | fibronectin leucine rich transmembrane protein 2 | -2.20 |
| ENSMUSG00000109295 | Gm5739 | predicted gene 5739 | -2.20 |
| ENSMUSG00000044734 | Serpinb1a | serine (or cysteine) peptidase inhibitor, clade B, member 1a | -2.21 |
| ENSMUSG00000035000 | Dpp4 | dipeptidylpeptidase 4 | -2.21 |
| ENSMUSG00000021416 | Eci3 | enoyl-Coenzyme A delta isomerase 3 | -2.21 |
| ENSMUSG00000019558 | Slc6a8 | solute carrier family 6 (neurotransmitter transporter, creatine), member 8 | -2.22 |
| ENSMUSG00000031803 | B3gnt3 | UDP-GlcNAc:betaGal beta-1,3-N-acetylglucosaminyltransferase 3 | -2.22 |
| ENSMUSG00000031618 | Nr3c2 | nuclear receptor subfamily 3, group C, member 2 | -2.22 |
| ENSMUSG00000038305 | Spats2l | spermatogenesis associated, serine-rich 2-like | -2.22 |
| ENSMUSG00000044165 | Bcl2l15 | BCLl2-like 15 | -2.22 |
| ENSMUSG00000110573 | Gm5485 | predicted gene 5485 | -2.22 |
| ENSMUSG00000039062 | Anpep | alanyl (membrane) aminopeptidase | -2.22 |
| ENSMUSG00000074345 | Tnfaip8l3 | tumor necrosis factor, alpha-induced protein 8-like 3 | -2.22 |
| ENSMUSG00000060703 | Cd302 | CD302 antigen | -2.22 |
| ENSMUSG00000110498 | A630001O12Rik | RIKEN cDNA A630001O12 gene | -2.22 |
| ENSMUSG00000050158 | Olfr165 | olfactory receptor 165 | -2.22 |
| ENSMUSG00000072941 | Sod3 | superoxide dismutase 3, extracellular | -2.22 |
| ENSMUSG00000061958 | Gm14851 | predicted gene 14851 | -2.23 |
| ENSMUSG00000020609 | Apob | apolipoprotein B | -2.23 |
| ENSMUSG00000029377 | Ereg | epiregulin | -2.23 |
| ENSMUSG00000004655 | Aqp1 | aquaporin 1 | -2.24 |
| ENSMUSG00000022809 | Nr1i2 | nuclear receptor subfamily 1, group I, member 2 | -2.24 |
| ENSMUSG00000110060 | Gm9860 | predicted gene 9860 | -2.24 |
| ENSMUSG00000029188 | Slc34a2 | solute carrier family 34 (sodium phosphate), member 2 | -2.24 |
| ENSMUSG00000097316 | Gm10516 | predicted gene 10516 | -2.24 |
| ENSMUSG00000027474 | Ccm2l | cerebral cavernous malformation 2-like | -2.24 |
| ENSMUSG00000025738 | Fbxl16 | F-box and leucine-rich repeat protein 16 | -2.24 |
| ENSMUSG00000037280 | Galnt6 | polypeptide N-acetylgalactosaminyltransferase 6 | -2.25 |
| ENSMUSG00000017688 | Hnf4g | hepatocyte nuclear factor 4, gamma | -2.25 |
| ENSMUSG00000085683 | Tmem238l | transmembrane protein 238 like | -2.25 |
| ENSMUSG00000034449 | Dhrs11 | dehydrogenase/reductase (SDR family) member 11 | -2.25 |
| ENSMUSG00000049690 | Nckap5 | NCK-associated protein 5 | -2.25 |
| ENSMUSG00000024411 | Aqp4 | aquaporin 4 | -2.25 |
| ENSMUSG00000028976 | Slc2a5 | solute carrier family 2 (facilitated glucose transporter), member 5 | -2.25 |
| ENSMUSG00000042388 | Dlgap3 | DLG associated protein 3 | -2.25 |
| ENSMUSG00000071335 | Mfsd4b3 | major facilitator superfamily domain containing 4B3 | -2.25 |
| ENSMUSG00000032087 | Dscaml1 | DS cell adhesion molecule like 1 | -2.25 |
| ENSMUSG00000053695 | Defb37 | defensin beta 37 | -2.25 |
| ENSMUSG00000036912 | Piwil4 | piwi-like RNA-mediated gene silencing 4 | -2.25 |
| ENSMUSG00000038642 | Ctss | cathepsin S | -2.26 |
| ENSMUSG00000040505 | Abcg5 | ATP binding cassette subfamily G member 5 | -2.26 |
| ENSMUSG00000112592 | Gm19972 | predicted gene, 19972 | -2.26 |
| ENSMUSG00000061414 | Cracr2a | calcium release activated channel regulator 2A | -2.27 |
| ENSMUSG00000026399 | Cd55 | CD55 molecule, decay accelerating factor for complement | -2.27 |
| ENSMUSG00000038903 | Ccdc68 | coiled-coil domain containing 68 | -2.27 |
| ENSMUSG00000014773 | Dll1 | delta like canonical Notch ligand 1 | -2.27 |
| ENSMUSG00000075702 | Selenom | selenoprotein M | -2.27 |
| ENSMUSG00000027068 | Dhrs9 | dehydrogenase/reductase (SDR family) member 9 | -2.27 |
| ENSMUSG00000048022 | Tmem229a | transmembrane protein 229A | -2.27 |
| ENSMUSG00000071506 | Tmem139 | transmembrane protein 139 | -2.27 |
| ENSMUSG00000102759 | Gm10463 | predicted gene 10463 | -2.27 |
| ENSMUSG00000025207 | Sema4g | sema domain, immunoglobulin domain (Ig), transmembrane domain (TM) and short cytoplasmic dom | -2.28 |
| ENSMUSG00000031402 | Mpp1 | membrane protein, palmitoylated | -2.28 |

| | | | |
|---|---|---|---|
| ENSMUSG00000019872 | Smpdl3a | sphingomyelin phosphodiesterase, acid-like 3A | -2.28 |
| ENSMUSG00000006362 | Cbfa2t3 | core-binding factor, runt domain, alpha subunit 2, translocated to, 3 (human) | -2.28 |
| ENSMUSG00000031844 | Hsd17b2 | hydroxysteroid (17-beta) dehydrogenase 2 | -2.28 |
| ENSMUSG00000057880 | Abat | 4-aminobutyrate aminotransferase | -2.28 |
| ENSMUSG00000028150 | Rorc | RAR-related orphan receptor gamma | -2.28 |
| ENSMUSG00000040471 | Ggt6 | gamma-glutamyltransferase 6 | -2.28 |
| ENSMUSG00000031937 | Vstm5 | V-set and transmembrane domain containing 5 | -2.28 |
| ENSMUSG00000033453 | Adamts15 | a disintegrin-like and metallopeptidase (reprolysin type) with thrombospondin type 1 motif, 15 | -2.28 |
| ENSMUSG00000042096 | Dao | D-amino acid oxidase | -2.28 |
| ENSMUSG00000058624 | Gda | guanine deaminase | -2.29 |
| ENSMUSG00000024713 | Pcsk5 | proprotein convertase subtilisin/kexin type 5 | -2.29 |
| ENSMUSG00000025528 | 2010106E10Rik | RIKEN cDNA 2010106E10 gene | -2.29 |
| ENSMUSG00000104535 | Gm42686 | predicted gene 42686 | -2.29 |
| ENSMUSG00000031389 | Arhgap4 | Rho GTPase activating protein 4 | -2.29 |
| ENSMUSG00000055833 | 1700034H15Rik | RIKEN cDNA 1700034H15 gene | -2.29 |
| ENSMUSG00000032034 | Kcnj5 | potassium inwardly-rectifying channel, subfamily J, member 5 | -2.29 |
| ENSMUSG00000006313 | Upk1a | uroplakin 1A | -2.29 |
| ENSMUSG00000035493 | Tgfbi | transforming growth factor, beta induced | -2.30 |
| ENSMUSG00000030157 | Clec2d | C-type lectin domain family 2, member d | -2.30 |
| ENSMUSG00000019478 | Rab4a | RAB4A, member RAS oncogene family | -2.30 |
| ENSMUSG00000025790 | Slco3a1 | solute carrier organic anion transporter family, member 3a1 | -2.30 |
| ENSMUSG00000033213 | AA467197 | expressed sequence AA467197 | -2.30 |
| ENSMUSG00000019278 | Dpep1 | dipeptidase 1 (renal) | -2.30 |
| ENSMUSG00000050982 | Apol10a | apolipoprotein L 10A | -2.30 |
| ENSMUSG00000090125 | Pou3f1 | POU domain, class 3, transcription factor 1 | -2.30 |
| ENSMUSG00000081303 | Gm16011 | predicted gene 16011 | -2.31 |
| ENSMUSG00000049307 | Fut4 | fucosyltransferase 4 | -2.31 |
| ENSMUSG00000104737 | Gm42937 | predicted gene 42937 | -2.31 |
| ENSMUSG00000109936 | Gm45889 | predicted gene 45889 | -2.31 |
| ENSMUSG00000094559 | Cyp2d34 | cytochrome P450, family 2, subfamily d, polypeptide 34 | -2.31 |
| ENSMUSG00000075184 | F930017D23Rik | RIKEN cDNA F930017D23 gene | -2.31 |
| ENSMUSG00000117084 | AC154912.1 | novel transcript, antisense to Fkbp5 | -2.31 |
| ENSMUSG00000026786 | Apbb1ip | amyloid beta (A4) precursor protein-binding, family B, member 1 interacting protein | -2.31 |
| ENSMUSG00000045005 | Fzd5 | frizzled class receptor 5 | -2.32 |
| ENSMUSG00000028307 | Aldob | aldolase B, fructose-bisphosphate | -2.32 |
| ENSMUSG00000025059 | Gk | glycerol kinase | -2.32 |
| ENSMUSG00000030155 | Clec2e | C-type lectin domain family 2, member e | -2.32 |
| ENSMUSG00000064373 | Selenop | selenoprotein P | -2.32 |
| ENSMUSG00000086825 | Gm15675 | predicted gene 15675 | -2.32 |
| ENSMUSG00000032098 | Treh | trehalase (brush-border membrane glycoprotein) | -2.32 |
| ENSMUSG00000054999 | Naaladl1 | N-acetylated alpha-linked acidic dipeptidase-like 1 | -2.32 |
| ENSMUSG00000079559 | Colca2 | COLCA2 homolog | -2.32 |
| ENSMUSG00000082154 | Gm16464 | predicted gene 16464 | -2.32 |
| ENSMUSG00000038528 | Mfsd4b5 | major facilitator superfamily domain containing 4B5 | -2.32 |
| ENSMUSG00000053168 | 9030619P08Rik | RIKEN cDNA 9030619P08 gene | -2.32 |
| ENSMUSG00000017943 | Gdap1l1 | ganglioside-induced differentiation-associated protein 1-like 1 | -2.32 |
| ENSMUSG00000030303 | Far2 | fatty acyl CoA reductase 2 | -2.33 |
| ENSMUSG00000024731 | Ms4a10 | membrane-spanning 4-domains, subfamily A, member 10 | -2.33 |
| ENSMUSG00000074195 | Clca4b | chloride channel accessory 4B | -2.33 |
| ENSMUSG00000073403 | Gm10499 | predicted gene 10499 | -2.33 |
| ENSMUSG00000113800 | 2210039B01Rik | RIKEN cDNA 2210039B01 gene | -2.33 |
| ENSMUSG00000114150 | Gm46367 | predicted gene, 46367 | -2.33 |
| ENSMUSG00000050866 | Clrn3 | clarin 3 | -2.34 |
| ENSMUSG00000023914 | Mep1a | meprin 1 alpha | -2.34 |
| ENSMUSG00000078670 | Fam174b | family with sequence similarity 174, member B | -2.34 |
| ENSMUSG00000073399 | Trim40 | tripartite motif-containing 40 | -2.34 |
| ENSMUSG00000001670 | Tat | tyrosine aminotransferase | -2.34 |
| ENSMUSG00000027555 | Car13 | carbonic anhydrase 13 | -2.34 |
| ENSMUSG00000028525 | Pde4b | phosphodiesterase 4B, cAMP specific | -2.34 |
| ENSMUSG00000058260 | Serpina9 | serine (or cysteine) peptidase inhibitor, clade A (alpha-1 antiproteinase, antitrypsin), member 9 | -2.34 |
| ENSMUSG00000050505 | Pcdh20 | protocadherin 20 | -2.34 |
| ENSMUSG00000041301 | Cftr | cystic fibrosis transmembrane conductance regulator | -2.35 |
| ENSMUSG00000000308 | Ckmt1 | creatine kinase, mitochondrial 1, ubiquitous | -2.35 |
| ENSMUSG00000040412 | 5330417C22Rik | RIKEN cDNA 5330417C22 gene | -2.35 |
| ENSMUSG00000023259 | Slc26a6 | solute carrier family 26, member 6 | -2.35 |
| ENSMUSG00000094651 | Gal3st2 | galactose-3-O-sulfotransferase 2 | -2.35 |
| ENSMUSG00000030834 | Abcc6 | ATP-binding cassette, sub-family C (CFTR/MRP), member 6 | -2.35 |
| ENSMUSG00000024885 | Aldh3b1 | aldehyde dehydrogenase 3 family, member B1 | -2.35 |
| ENSMUSG00000044827 | Tlr1 | toll-like receptor 1 | -2.35 |
| ENSMUSG00000027297 | Ltk | leukocyte tyrosine kinase | -2.35 |
| ENSMUSG00000078439 | Smim24 | small integral membrane protein 24 | -2.36 |

| | | | |
|---|---|---|---|
| ENSMUSG00000029314 | Gpat3 | glycerol-3-phosphate acyltransferase 3 | -2.36 |
| ENSMUSG00000074300 | BC030870 | cDNA sequence BC030870 | -2.36 |
| ENSMUSG00000036242 | Armh4 | armadillo-like helical domain containing 4 | -2.36 |
| ENSMUSG00000074183 | Gsta1 | glutathione S-transferase, alpha 1 (Ya) | -2.36 |
| ENSMUSG00000053964 | Lgals4 | lectin, galactose binding, soluble 4 | -2.37 |
| ENSMUSG00000032528 | Vipr1 | vasoactive intestinal peptide receptor 1 | -2.37 |
| ENSMUSG00000068452 | Duox2 | dual oxidase 2 | -2.37 |
| ENSMUSG00000035506 | Slc12a8 | solute carrier family 12 (potassium/chloride transporters), member 8 | -2.37 |
| ENSMUSG00000027225 | Duoxa2 | dual oxidase maturation factor 2 | -2.37 |
| ENSMUSG00000038515 | Grtp1 | GH regulated TBC protein 1 | -2.37 |
| ENSMUSG00000006345 | Ggt1 | gamma-glutamyltransferase 1 | -2.37 |
| ENSMUSG00000073402 | Gm8909 | predicted gene 8909 | -2.37 |
| ENSMUSG00000037390 | Muc3 | mucin 3, intestinal | -2.37 |
| ENSMUSG00000024503 | Spink1 | serine peptidase inhibitor, Kazal type 1 | -2.37 |
| ENSMUSG00000038745 | Nlrp6 | NLR family, pyrin domain containing 6 | -2.38 |
| ENSMUSG00000031451 | Gas6 | growth arrest specific 6 | -2.38 |
| ENSMUSG00000006360 | Crip1 | cysteine-rich protein 1 (intestinal) | -2.38 |
| ENSMUSG00000001739 | Cldn15 | claudin 15 | -2.38 |
| ENSMUSG00000030865 | Chp2 | calcineurin-like EF hand protein 2 | -2.38 |
| ENSMUSG00000034947 | Tmem106a | transmembrane protein 106A | -2.38 |
| ENSMUSG00000038843 | Gcnt1 | glucosaminyl (N-acetyl) transferase 1, core 2 | -2.38 |
| ENSMUSG00000074261 | Erich4 | glutamate rich 4 | -2.38 |
| ENSMUSG00000084174 | Sycn | syncollin | -2.38 |
| ENSMUSG00000039438 | Ttc36 | tetratricopeptide repeat domain 36 | -2.38 |
| ENSMUSG00000030492 | Slc7a9 | solute carrier family 7 (cationic amino acid transporter, y+ system), member 9 | -2.39 |
| ENSMUSG00000091705 | H2-Q2 | histocompatibility 2, Q region locus 2 | -2.39 |
| ENSMUSG00000001095 | Slc13a2 | solute carrier family 13 (sodium-dependent dicarboxylate transporter), member 2 | -2.39 |
| ENSMUSG00000043822 | Adamtsl5 | ADAMTS-like 5 | -2.39 |
| ENSMUSG00000019888 | Mgat4c | MGAT4 family, member C | -2.40 |
| ENSMUSG00000051111 | Sv2c | synaptic vesicle glycoprotein 2c | -2.40 |
| ENSMUSG00000072849 | Serpina1e | serine (or cysteine) peptidase inhibitor, clade A, member 1E | -2.40 |
| ENSMUSG00000104906 | Gm43005 | predicted gene 43005 | -2.40 |
| ENSMUSG00000031621 | Isx | intestine specific homeobox | -2.41 |
| ENSMUSG00000026126 | Ptpn18 | protein tyrosine phosphatase, non-receptor type 18 | -2.41 |
| ENSMUSG00000020062 | Slc5a8 | solute carrier family 5 (iodide transporter), member 8 | -2.41 |
| ENSMUSG00000020839 | Tmigd1 | transmembrane and immunoglobulin domain containing 1 | -2.41 |
| ENSMUSG00000032978 | Guca2b | guanylate cyclase activator 2b (retina) | -2.42 |
| ENSMUSG00000039899 | Fgl2 | fibrinogen-like protein 2 | -2.42 |
| ENSMUSG00000010660 | Plcd1 | phospholipase C, delta 1 | -2.42 |
| ENSMUSG00000020072 | Pbld2 | phenazine biosynthesis-like protein domain containing 2 | -2.42 |
| ENSMUSG00000032357 | Tinag | tubulointerstitial nephritis antigen | -2.42 |
| ENSMUSG00000035200 | Chrnb4 | cholinergic receptor, nicotinic, beta polypeptide 4 | -2.42 |
| ENSMUSG00000044453 | Ffar1 | free fatty acid receptor 1 | -2.42 |
| ENSMUSG00000059654 | Reg1 | regenerating islet-derived 1 | -2.42 |
| ENSMUSG00000005672 | Kit | KIT proto-oncogene receptor tyrosine kinase | -2.43 |
| ENSMUSG00000021676 | Iqgap2 | IQ motif containing GTPase activating protein 2 | -2.43 |
| ENSMUSG00000031886 | Ces2e | carboxylesterase 2E | -2.43 |
| ENSMUSG00000043705 | Capn13 | calpain 13 | -2.43 |
| ENSMUSG00000079440 | Alpi | alkaline phosphatase, intestinal | -2.43 |
| ENSMUSG00000055730 | Ces2a | carboxylesterase 2A | -2.43 |
| ENSMUSG00000057719 | Sh3rf2 | SH3 domain containing ring finger 2 | -2.43 |
| ENSMUSG00000027420 | Bfsp1 | beaded filament structural protein 1, in lens-CP94 | -2.43 |
| ENSMUSG00000040740 | Slc25a34 | solute carrier family 25, member 34 | -2.43 |
| ENSMUSG00000028940 | Hes2 | hes family bHLH transcription factor 2 | -2.43 |
| ENSMUSG00000049122 | Frmd3 | FERM domain containing 3 | -2.43 |
| ENSMUSG00000023959 | Clic5 | chloride intracellular channel 5 | -2.44 |
| ENSMUSG00000026398 | Nr5a2 | nuclear receptor subfamily 5, group A, member 2 | -2.44 |
| ENSMUSG00000072812 | Ahnak2 | AHNAK nucleoprotein 2 | -2.44 |
| ENSMUSG00000028024 | Enpep | glutamyl aminopeptidase | -2.44 |
| ENSMUSG00000010066 | Cacna2d2 | calcium channel, voltage-dependent, alpha 2/delta subunit 2 | -2.44 |
| ENSMUSG00000028186 | Uox | urate oxidase | -2.44 |
| ENSMUSG00000041119 | Pde9a | phosphodiesterase 9A | -2.45 |
| ENSMUSG00000074272 | Ceacam1 | carcinoembryonic antigen-related cell adhesion molecule 1 | -2.45 |
| ENSMUSG00000027317 | Ppp1r14d | protein phosphatase 1, regulatory inhibitor subunit 14D | -2.45 |
| ENSMUSG00000024131 | Slc3a1 | solute carrier family 3, member 1 | -2.45 |
| ENSMUSG00000087651 | 1500009L16Rik | RIKEN cDNA 1500009L16 gene | -2.45 |
| ENSMUSG00000025189 | Cnnm1 | cyclin M1 | -2.45 |
| ENSMUSG00000053182 | Gm609 | predicted gene 609 | -2.46 |
| ENSMUSG00000025347 | Mettl7b | methyltransferase like 7B | -2.46 |
| ENSMUSG00000090700 | Cyp4f40 | cytochrome P450, family 4, subfamily f, polypeptide 40 | -2.46 |
| ENSMUSG00000034785 | Dio1 | deiodinase, iodothyronine, type I | -2.46 |

| | | | |
|---|---|---|---|
| ENSMUSG00000079507 | H2-Q1 | histocompatibility 2, Q region locus 1 | -2.46 |
| ENSMUSG00000027843 | Ptpn22 | protein tyrosine phosphatase, non-receptor type 22 (lymphoid) | -2.46 |
| ENSMUSG00000028003 | Lrat | lecithin-retinol acyltransferase (phosphatidylcholine-retinol-O-acyltransferase) | -2.46 |
| ENSMUSG00000028874 | Fgr | FGR proto-oncogene, Src family tyrosine kinase | -2.46 |
| ENSMUSG00000030340 | Scnn1a | sodium channel, nonvoltage-gated 1 alpha | -2.47 |
| ENSMUSG00000062638 | Btnl1 | butyrophilin-like 1 | -2.47 |
| ENSMUSG00000029700 | Slc13a1 | solute carrier family 13 (sodium/sulfate symporters), member 1 | -2.47 |
| ENSMUSG00000053862 | Slc51b | solute carrier family 51, beta subunit | -2.47 |
| ENSMUSG00000026614 | Slc30a10 | solute carrier family 30, member 10 | -2.47 |
| ENSMUSG00000025401 | Myo1a | myosin IA | -2.48 |
| ENSMUSG00000026825 | Dnm1 | dynamin 1 | -2.48 |
| ENSMUSG00000029630 | Cyp3a25 | cytochrome P450, family 3, subfamily a, polypeptide 25 | -2.48 |
| ENSMUSG00000034918 | Cdhr2 | cadherin-related family member 2 | -2.48 |
| ENSMUSG00000029095 | Ablim2 | actin-binding LIM protein 2 | -2.48 |
| ENSMUSG00000045930 | Clec14a | C-type lectin domain family 14, member a | -2.48 |
| ENSMUSG00000038910 | Plcl2 | phospholipase C-like 2 | -2.49 |
| ENSMUSG00000027200 | Sema6d | sema domain, transmembrane domain (TM), and cytoplasmic domain, (semaphorin) 6D | -2.49 |
| ENSMUSG00000049858 | Suox | sulfite oxidase | -2.49 |
| ENSMUSG00000021565 | Slc6a19 | solute carrier family 6 (neurotransmitter transporter), member 19 | -2.49 |
| ENSMUSG00000029162 | Khk | ketohexokinase | -2.49 |
| ENSMUSG00000042770 | Hebp1 | heme binding protein 1 | -2.49 |
| ENSMUSG00000060615 | Ang4 | angiogenin, ribonuclease A family, member 4 | -2.50 |
| ENSMUSG00000032548 | Slco2a1 | solute carrier organic anion transporter family, member 2a1 | -2.50 |
| ENSMUSG00000033308 | Dpyd | dihydropyrimidine dehydrogenase | -2.50 |
| ENSMUSG00000028327 | Stra6l | STRA6-like | -2.50 |
| ENSMUSG00000040703 | Cyp2s1 | cytochrome P450, family 2, subfamily s, polypeptide 1 | -2.50 |
| ENSMUSG00000021947 | Cryl1 | crystallin, lambda 1 | -2.50 |
| ENSMUSG00000021798 | Ldb3 | LIM domain binding 3 | -2.50 |
| ENSMUSG00000023032 | Slc4a8 | solute carrier family 4 (anion exchanger), member 8 | -2.50 |
| ENSMUSG00000021278 | Amn | amnionless | -2.50 |
| ENSMUSG00000089948 | Far2os1 | fatty acyl CoA reductase 2, opposite strand 1 | -2.50 |
| ENSMUSG00000026380 | Tfcp2l1 | transcription factor CP2-like 1 | -2.51 |
| ENSMUSG00000024105 | Themis3 | thymocyte selection associated family member 3 | -2.51 |
| ENSMUSG00000022235 | Cmbl | carboxymethylenebutenolidase-like (Pseudomonas) | -2.51 |
| ENSMUSG00000029420 | Rimbp2 | RIMS binding protein 2 | -2.51 |
| ENSMUSG00000020805 | Slc13a5 | solute carrier family 13 (sodium-dependent citrate transporter), member 5 | -2.51 |
| ENSMUSG00000029084 | Cd38 | CD38 antigen | -2.52 |
| ENSMUSG00000090124 | Ugt1a7c | UDP glucuronosyltransferase 1 family, polypeptide A7C | -2.52 |
| ENSMUSG00000108476 | Gm44974 | predicted gene 44974 | -2.52 |
| ENSMUSG00000021125 | Arg2 | arginase type II | -2.52 |
| ENSMUSG00000057614 | Gnai1 | guanine nucleotide binding protein (G protein), alpha inhibiting 1 | -2.52 |
| ENSMUSG00000093580 | Gm20706 | predicted gene 20706 | -2.52 |
| ENSMUSG00000026544 | Dusp23 | dual specificity phosphatase 23 | -2.52 |
| ENSMUSG00000024292 | Cyp4f14 | cytochrome P450, family 4, subfamily f, polypeptide 14 | -2.53 |
| ENSMUSG00000027377 | Mall | mal, T cell differentiation protein-like | -2.53 |
| ENSMUSG00000022546 | Gpt | glutamic pyruvic transaminase, soluble | -2.53 |
| ENSMUSG00000021509 | Slc25a48 | solute carrier family 25, member 48 | -2.53 |
| ENSMUSG00000107456 | Gm10400 | predicted gene 10400 | -2.53 |
| ENSMUSG00000032226 | Gcnt3 | glucosaminyl (N-acetyl) transferase 3, mucin type | -2.54 |
| ENSMUSG00000033847 | Pla2g4c | phospholipase A2, group IVC (cytosolic, calcium-independent) | -2.54 |
| ENSMUSG00000044352 | Sowaha | sosondowah ankyrin repeat domain family member A | -2.54 |
| ENSMUSG00000037440 | Vnn1 | vanin 1 | -2.54 |
| ENSMUSG00000027359 | Slc27a2 | solute carrier family 27 (fatty acid transporter), member 2 | -2.54 |
| ENSMUSG00000031163 | Glod5 | glyoxalase domain containing 5 | -2.54 |
| ENSMUSG00000023122 | Sult1c2 | sulfotransferase family, cytosolic, 1C, member 2 | -2.54 |
| ENSMUSG00000025497 | Cdhr5 | cadherin-related family member 5 | -2.55 |
| ENSMUSG00000024313 | Mep1b | meprin 1 beta | -2.55 |
| ENSMUSG00000055114 | Anxa13 | annexin A13 | -2.55 |
| ENSMUSG00000024124 | Prss30 | protease, serine 30 | -2.55 |
| ENSMUSG00000036151 | Tm6sf2 | transmembrane 6 superfamily member 2 | -2.55 |
| ENSMUSG00000042284 | Itga1 | integrin alpha 1 | -2.55 |
| ENSMUSG00000026928 | Card9 | caspase recruitment domain family, member 9 | -2.55 |
| ENSMUSG00000032487 | Ptgs2 | prostaglandin-endoperoxide synthase 2 | -2.55 |
| ENSMUSG00000106417 | Gm42628 | predicted gene 42628 | -2.55 |
| ENSMUSG00000088609 | Gm24187 | predicted gene, 24187 | -2.56 |
| ENSMUSG00000010601 | Apol7a | apolipoprotein L 7a | -2.56 |
| ENSMUSG00000091387 | Gcnt4 | glucosaminyl (N-acetyl) transferase 4, core 2 (beta-1,6-N-acetylglucosaminyltransferase) | -2.56 |
| ENSMUSG00000033174 | Mgll | monoglyceride lipase | -2.56 |
| ENSMUSG00000061808 | Ttr | transthyretin | -2.56 |
| ENSMUSG00000031980 | Agt | angiotensinogen (serpin peptidase inhibitor, clade A, member 8) | -2.57 |
| ENSMUSG00000056162 | Cndp1 | carnosine dipeptidase 1 (metallopeptidase M20 family) | -2.57 |

| | | | |
|---|---|---|---|
| ENSMUSG00000036422 | Pcdh8 | protocadherin 8 | -2.57 |
| ENSMUSG00000074441 | Gm15292 | predicted gene 15292 | -2.57 |
| ENSMUSG00000075217 | 4833423E24Rik | RIKEN cDNA 4833423E24 gene | -2.57 |
| ENSMUSG00000094840 | Muc3a | mucin 3A, cell surface associated | -2.58 |
| ENSMUSG00000038751 | Ptk6 | PTK6 protein tyrosine kinase 6 | -2.58 |
| ENSMUSG00000109644 | 0610005C13Rik | RIKEN cDNA 0610005C13 gene | -2.58 |
| ENSMUSG00000061959 | Ces1e | carboxylesterase 1E | -2.58 |
| ENSMUSG00000027790 | Sis | sucrase isomaltase (alpha-glucosidase) | -2.59 |
| ENSMUSG00000036377 | C530008M17Rik | RIKEN cDNA C530008M17 gene | -2.59 |
| ENSMUSG00000038167 | Plekhg6 | pleckstrin homology domain containing, family G (with RhoGef domain) member 6 | -2.59 |
| ENSMUSG00000027983 | Cyp2u1 | cytochrome P450, family 2, subfamily u, polypeptide 1 | -2.59 |
| ENSMUSG00000020838 | Slc6a4 | solute carrier family 6 (neurotransmitter transporter, serotonin), member 4 | -2.59 |
| ENSMUSG00000035429 | Ptprh | protein tyrosine phosphatase, receptor type, H | -2.60 |
| ENSMUSG00000041794 | Myrip | myosin VIIA and Rab interacting protein | -2.60 |
| ENSMUSG00000049134 | Nrap | nebulin-related anchoring protein | -2.60 |
| ENSMUSG00000049154 | Fam183b | family with sequence similarity 183, member B | -2.60 |
| ENSMUSG00000097266 | Gm26802 | predicted gene, 26802 | -2.60 |
| ENSMUSG00000035775 | Krt20 | keratin 20 | -2.61 |
| ENSMUSG00000058435 | Btnl4 | butyrophilin-like 4 | -2.61 |
| ENSMUSG00000036814 | Slc6a20a | solute carrier family 6 (neurotransmitter transporter), member 20A | -2.61 |
| ENSMUSG00000085042 | Abhd11os | abhydrolase domain containing 11, opposite strand | -2.61 |
| ENSMUSG00000025092 | Hspa12a | heat shock protein 12A | -2.61 |
| ENSMUSG00000027350 | Chgb | chromogranin B | -2.61 |
| ENSMUSG00000000049 | Apoh | apolipoprotein H | -2.61 |
| ENSMUSG00000015468 | Notch4 | notch 4 | -2.62 |
| ENSMUSG00000096546 | Smlr1 | small leucine-rich protein 1 | -2.62 |
| ENSMUSG00000040770 | Il25 | interleukin 25 | -2.62 |
| ENSMUSG00000030364 | Clec2h | C-type lectin domain family 2, member h | -2.63 |
| ENSMUSG00000001225 | Slc26a3 | solute carrier family 26, member 3 | -2.63 |
| ENSMUSG00000022408 | Fam83f | family with sequence similarity 83, member F | -2.63 |
| ENSMUSG00000024548 | Setbp1 | SET binding protein 1 | -2.63 |
| ENSMUSG00000052026 | Slc6a7 | solute carrier family 6 (neurotransmitter transporter, L-proline), member 7 | -2.63 |
| ENSMUSG00000021640 | Naip1 | NLR family, apoptosis inhibitory protein 1 | -2.64 |
| ENSMUSG00000045775 | Slc16a5 | solute carrier family 16 (monocarboxylic acid transporters), member 5 | -2.64 |
| ENSMUSG00000046697 | Enpp7 | ectonucleotide pyrophosphatase/phosphodiesterase 7 | -2.64 |
| ENSMUSG00000021675 | F2rl2 | coagulation factor II (thrombin) receptor-like 2 | -2.64 |
| ENSMUSG00000066176 | Gm12511 | predicted gene 12511 | -2.64 |
| ENSMUSG00000034427 | Myo15b | myosin XVB | -2.65 |
| ENSMUSG00000092618 | Btnl6 | butyrophilin-like 6 | -2.65 |
| ENSMUSG00000021208 | Ifi27l2b | interferon, alpha-inducible protein 27 like 2B | -2.65 |
| ENSMUSG00000027792 | Bche | butyrylcholinesterase | -2.65 |
| ENSMUSG00000070315 | 4930581F22Rik | RIKEN cDNA 4930581F22 gene | -2.65 |
| ENSMUSG00000049892 | Rasd1 | RAS, dexamethasone-induced 1 | -2.65 |
| ENSMUSG00000024371 | C2 | complement component 2 (within H-2S) | -2.65 |
| ENSMUSG00000078650 | G6pc | glucose-6-phosphatase, catalytic | -2.65 |
| ENSMUSG00000071356 | Reg3b | regenerating islet-derived 3 beta | -2.66 |
| ENSMUSG00000037005 | Xpnpep2 | X-prolyl aminopeptidase (aminopeptidase P) 2, membrane-bound | -2.66 |
| ENSMUSG00000050592 | Fam78a | family with sequence similarity 78, member A | -2.66 |
| ENSMUSG00000058163 | Gm5431 | predicted gene 5431 | -2.66 |
| ENSMUSG00000031445 | Proz | protein Z, vitamin K-dependent plasma glycoprotein | -2.66 |
| ENSMUSG00000096299 | Gm21814 | predicted gene, 21814 | -2.66 |
| ENSMUSG00000054128 | H2-T3 | histocompatibility 2, T region locus 3 | -2.67 |
| ENSMUSG00000027876 | Reg4 | regenerating islet-derived family, member 4 | -2.67 |
| ENSMUSG00000036298 | Slc2a13 | solute carrier family 2 (facilitated glucose transporter), member 13 | -2.67 |
| ENSMUSG00000033715 | Akr1c14 | aldo-keto reductase family 1, member C14 | -2.67 |
| ENSMUSG00000046613 | Vwa5b2 | von Willebrand factor A domain containing 5B2 | -2.67 |
| ENSMUSG00000021194 | Chga | chromogranin A | -2.67 |
| ENSMUSG00000044505 | Lingo4 | leucine rich repeat and Ig domain containing 4 | -2.67 |
| ENSMUSG00000023235 | Ccl25 | chemokine (C-C motif) ligand 25 | -2.68 |
| ENSMUSG00000074437 | Defa29 | defensin, alpha, 29 | -2.68 |
| ENSMUSG00000038195 | Rilp | Rab interacting lysosomal protein | -2.68 |
| ENSMUSG00000067231 | Cyp2c65 | cytochrome P450, family 2, subfamily c, polypeptide 65 | -2.69 |
| ENSMUSG00000002100 | Mybpc3 | myosin binding protein C, cardiac | -2.69 |
| ENSMUSG00000027513 | Pck1 | phosphoenolpyruvate carboxykinase 1, cytosolic | -2.70 |
| ENSMUSG00000091575 | 2010016I18Rik | RIKEN cDNA 2010016I18 gene | -2.70 |
| ENSMUSG00000018919 | Tm4sf5 | transmembrane 4 superfamily member 5 | -2.70 |
| ENSMUSG00000039339 | Mfsd4b2 | major facilitator superfamily domain containing 4B2 | -2.70 |
| ENSMUSG00000075520 | Malrd1 | MAM and LDL receptor class A domain containing 1 | -2.71 |
| ENSMUSG00000035699 | Slc51a | solute carrier family 51, alpha subunit | -2.71 |
| ENSMUSG00000029161 | Cgref1 | cell growth regulator with EF hand domain 1 | -2.71 |
| ENSMUSG00000014361 | Mertk | c-mer proto-oncogene tyrosine kinase | -2.71 |

| | | | |
|---|---|---|---|
| ENSMUSG00000021850 | ccdc198 | coiled-coil domain containing 198 | -2.71 |
| ENSMUSG00000031073 | Fgf15 | fibroblast growth factor 15 | -2.71 |
| ENSMUSG00000031497 | Tnfsf13b | tumor necrosis factor (ligand) superfamily, member 13b | -2.71 |
| ENSMUSG00000028251 | Tstd3 | thiosulfate sulfurtransferase (rhodanese)-like domain containing 3 | -2.71 |
| ENSMUSG00000029134 | Plb1 | phospholipase B1 | -2.71 |
| ENSMUSG00000029330 | Cds1 | CDP-diacylglycerol synthase 1 | -2.72 |
| ENSMUSG00000045441 | Gprin3 | GPRIN family member 3 | -2.72 |
| ENSMUSG00000060070 | Defa26 | defensin, alpha, 26 | -2.72 |
| ENSMUSG00000073413 | Ly6g6d | lymphocyte antigen 6 complex, locus G6D | -2.72 |
| ENSMUSG00000030017 | Reg3g | regenerating islet-derived 3 gamma | -2.73 |
| ENSMUSG00000071203 | Naip5 | NLR family, apoptosis inhibitory protein 5 | -2.73 |
| ENSMUSG00000029269 | Sult1b1 | sulfotransferase family 1B, member 1 | -2.73 |
| ENSMUSG00000024935 | Slc1a1 | solute carrier family 1 (neuronal/epithelial high affinity glutamate transporter, system Xag), member [illegible] | -2.73 |
| ENSMUSG00000052229 | Gpr17 | G protein-coupled receptor 17 | -2.73 |
| ENSMUSG00000026610 | Esrrg | estrogen-related receptor gamma | -2.73 |
| ENSMUSG00000034112 | Atp2c2 | ATPase, Ca++ transporting, type 2C, member 2 | -2.74 |
| ENSMUSG00000021749 | Oit1 | oncoprotein induced transcript 1 | -2.74 |
| ENSMUSG00000021806 | Nid2 | nidogen 2 | -2.74 |
| ENSMUSG00000106542 | Gm43410 | predicted gene 43410 | -2.74 |
| ENSMUSG00000024140 | Epas1 | endothelial PAS domain protein 1 | -2.75 |
| ENSMUSG00000020080 | Hkdc1 | hexokinase domain containing 1 | -2.75 |
| ENSMUSG00000005268 | Prlr | prolactin receptor | -2.75 |
| ENSMUSG00000027173 | Depdc7 | DEP domain containing 7 | -2.75 |
| ENSMUSG00000094091 | Gm21885 | predicted gene, 21885 | -2.75 |
| ENSMUSG00000022479 | Vdr | vitamin D (1,25-dihydroxyvitamin D3) receptor | -2.76 |
| ENSMUSG00000032038 | St3gal4 | ST3 beta-galactoside alpha-2,3-sialyltransferase 4 | -2.76 |
| ENSMUSG00000020865 | Abcc3 | ATP-binding cassette, sub-family C (CFTR/MRP), member 3 | -2.76 |
| ENSMUSG00000073420 | Btnl5-ps | butyrophilin-like 5, pseudogene | -2.76 |
| ENSMUSG00000047638 | Nr1h4 | nuclear receptor subfamily 1, group H, member 4 | -2.76 |
| ENSMUSG00000058921 | Slc10a5 | solute carrier family 10 (sodium/bile acid cotransporter family), member 5 | -2.77 |
| ENSMUSG00000096146 | Kcnj11 | potassium inwardly rectifying channel, subfamily J, member 11 | -2.77 |
| ENSMUSG00000074115 | Saa1 | serum amyloid A 1 | -2.78 |
| ENSMUSG00000031725 | Ces1f | carboxylesterase 1F | -2.78 |
| ENSMUSG00000034762 | Glis1 | GLIS family zinc finger 1 | -2.78 |
| ENSMUSG00000056293 | Gsdmc2 | gasdermin C2 | -2.79 |
| ENSMUSG00000074715 | Ccl28 | chemokine (C-C motif) ligand 28 | -2.79 |
| ENSMUSG00000061728 | Btnl7-ps | butyrophilin-like 7, pseudogene | -2.79 |
| ENSMUSG00000026163 | Sphkap | SPHK1 interactor, AKAP domain containing | -2.79 |
| ENSMUSG00000063296 | Tmem117 | transmembrane protein 117 | -2.80 |
| ENSMUSG00000073406 | H2-Bl | histocompatibility 2, blastocyst | -2.80 |
| ENSMUSG00000020732 | Rab37 | RAB37, member RAS oncogene family | -2.81 |
| ENSMUSG00000058063 | Trim31 | tripartite motif-containing 31 | -2.82 |
| ENSMUSG00000070777 | Ceacam20 | carcinoembryonic antigen-related cell adhesion molecule 20 | -2.82 |
| ENSMUSG00000024846 | Cst6 | cystatin E/M | -2.82 |
| ENSMUSG00000031410 | Nxf7 | nuclear RNA export factor 7 | -2.82 |
| ENSMUSG00000024620 | Pdgfrb | platelet derived growth factor receptor, beta polypeptide | -2.82 |
| ENSMUSG00000103356 | Gm37595 | predicted gene, 37595 | -2.82 |
| ENSMUSG00000031551 | Ido1 | indoleamine 2,3-dioxygenase 1 | -2.83 |
| ENSMUSG00000030560 | Ctsc | cathepsin C | -2.85 |
| ENSMUSG00000036492 | Rnf39 | ring finger protein 39 | -2.85 |
| ENSMUSG00000059003 | Grin2a | glutamate receptor, ionotropic, NMDA2A (epsilon 1) | -2.85 |
| ENSMUSG00000035277 | Arx | aristaless related homeobox | -2.85 |
| ENSMUSG00000031613 | Hpgd | hydroxyprostaglandin dehydrogenase 15 (NAD) | -2.86 |
| ENSMUSG00000024730 | Ms4a8a | membrane-spanning 4-domains, subfamily A, member 8A | -2.86 |
| ENSMUSG00000022383 | Ppara | peroxisome proliferator activated receptor alpha | -2.86 |
| ENSMUSG00000034645 | Zyg11a | zyg-11 family member A, cell cycle regulator | -2.86 |
| ENSMUSG00000052955 | Cpvl | carboxypeptidase, vitellogenic-like | -2.86 |
| ENSMUSG00000011008 | Mcoln2 | mucolipin 2 | -2.87 |
| ENSMUSG00000108155 | Gm44443 | predicted gene, 44443 | -2.87 |
| ENSMUSG00000001604 | Tcea3 | transcription elongation factor A (SII), 3 | -2.88 |
| ENSMUSG00000022661 | Cd200 | CD200 antigen | -2.88 |
| ENSMUSG00000079355 | Ackr4 | atypical chemokine receptor 4 | -2.88 |
| ENSMUSG00000064177 | Ghrl | ghrelin | -2.88 |
| ENSMUSG00000022489 | Pde1b | phosphodiesterase 1B, Ca2+-calmodulin dependent | -2.88 |
| ENSMUSG00000021070 | Bdkrb2 | bradykinin receptor, beta 2 | -2.88 |
| ENSMUSG00000089635 | Gm16559 | predicted gene 16559 | -2.88 |
| ENSMUSG00000044860 | Gm1123 | predicted gene 1123 | -2.89 |
| ENSMUSG00000098557 | Kctd12 | potassium channel tetramerisation domain containing 12 | -2.89 |
| ENSMUSG00000027661 | Slc2a10 | solute carrier family 2 (facilitated glucose transporter), member 10 | -2.89 |
| ENSMUSG00000116383 | Apol10c-ps | apolipoprotein L 10C, pseudogene | -2.89 |
| ENSMUSG00000002565 | Scin | scinderin | -2.90 |

| | | | |
|---|---|---|---|
| ENSMUSG00000090062 | Galnt6os | polypeptide N-acetylgalactosaminyltransferase 6, opposite strand | -2.90 |
| ENSMUSG00000022755 | Adgrg7 | adhesion G protein-coupled receptor G7 | -2.91 |
| ENSMUSG00000061531 | Tmem236 | transmembrane protein 236 | -2.91 |
| ENSMUSG00000074207 | Adh1 | alcohol dehydrogenase 1 (class I) | -2.91 |
| ENSMUSG00000039878 | Slc39a5 | solute carrier family 39 (metal ion transporter), member 5 | -2.91 |
| ENSMUSG00000030472 | Ceacam18 | carcinoembryonic antigen-related cell adhesion molecule 18 | -2.91 |
| ENSMUSG00000006642 | Tcf23 | transcription factor 23 | -2.92 |
| ENSMUSG00000072623 | Zfp9 | zinc finger protein 9 | -2.92 |
| ENSMUSG00000033200 | Tpsg1 | tryptase gamma 1 | -2.93 |
| ENSMUSG00000030737 | Slco2b1 | solute carrier organic anion transporter family, member 2b1 | -2.93 |
| ENSMUSG00000064272 | Gpbar1 | G protein-coupled bile acid receptor 1 | -2.93 |
| ENSMUSG00000037953 | A4gnt | alpha-1,4-N-acetylglucosaminyltransferase | -2.94 |
| ENSMUSG00000027224 | Duoxa1 | dual oxidase maturation factor 1 | -2.94 |
| ENSMUSG00000067377 | Tspan6 | tetraspanin 6 | -2.95 |
| ENSMUSG00000041696 | Rasl12 | RAS-like, family 12 | -2.95 |
| ENSMUSG00000040653 | Ppp1r14c | protein phosphatase 1, regulatory inhibitor subunit 14C | -2.95 |
| ENSMUSG00000032181 | Scg3 | secretogranin III | -2.96 |
| ENSMUSG00000087518 | Gm13561 | predicted gene 13561 | -2.96 |
| ENSMUSG00000050097 | Ces2b | carboxyesterase 2B | -2.97 |
| ENSMUSG00000025010 | Ccnj | cyclin J | -2.98 |
| ENSMUSG00000023247 | Guca2a | guanylate cyclase activator 2a (guanylin) | -2.98 |
| ENSMUSG00000078776 | 9530053A07Rik | RIKEN cDNA 9530053A07 gene | -2.98 |
| ENSMUSG00000046804 | Phgr1 | proline/histidine/glycine-rich 1 | -2.99 |
| ENSMUSG00000021216 | Tubal3 | tubulin, alpha-like 3 | -2.99 |
| ENSMUSG00000055748 | Gsdmc4 | gasdermin C4 | -3.00 |
| ENSMUSG00000029273 | Sult1d1 | sulfotransferase family 1D, member 1 | -3.00 |
| ENSMUSG00000026870 | Cutal | cutA divalent cation tolerance homolog-like | -3.00 |
| ENSMUSG00000020120 | Plek | pleckstrin | -3.00 |
| ENSMUSG00000013643 | Lypd8 | LY6/PLAUR domain containing 8 | -3.01 |
| ENSMUSG00000094584 | Ms4a18 | membrane-spanning 4-domains, subfamily A, member 18 | -3.02 |
| ENSMUSG00000110405 | Gm45534 | predicted gene 45534 | -3.02 |
| ENSMUSG00000024340 | Btnl2 | butyrophilin-like 2 | -3.03 |
| ENSMUSG00000079180 | Mptx2 | mucosal pentraxin 2 | -3.04 |
| ENSMUSG00000072723 | Gm10044 | predicted gene 10044 | -3.04 |
| ENSMUSG00000047564 | Krtap3-1 | keratin associated protein 3-1 | -3.04 |
| ENSMUSG00000035849 | Krt222 | keratin 222 | -3.05 |
| ENSMUSG00000027220 | Syt13 | synaptotagmin XIII | -3.05 |
| ENSMUSG00000034472 | Rasd2 | RASD family, member 2 | -3.07 |
| ENSMUSG00000041476 | Smpx | small muscle protein, X-linked | -3.07 |
| ENSMUSG00000024553 | Galr1 | galanin receptor 1 | -3.07 |
| ENSMUSG00000029335 | Bmp3 | bone morphogenetic protein 3 | -3.08 |
| ENSMUSG00000078137 | Ankrd63 | ankyrin repeat domain 63 | -3.09 |
| ENSMUSG00000034057 | Myrfl | myelin regulatory factor-like | -3.10 |
| ENSMUSG00000054640 | Slc8a1 | solute carrier family 8 (sodium/calcium exchanger), member 1 | -3.10 |
| ENSMUSG00000024215 | Spdef | SAM pointed domain containing ets transcription factor | -3.11 |
| ENSMUSG00000032202 | Rab27a | RAB27A, member RAS oncogene family | -3.12 |
| ENSMUSG00000060508 | Nlrp9b | NLR family, pyrin domain containing 9B | -3.12 |
| ENSMUSG00000050296 | Abca12 | ATP-binding cassette, sub-family A (ABC1), member 12 | -3.12 |
| ENSMUSG00000038599 | Capn8 | calpain 8 | -3.13 |
| ENSMUSG00000060224 | Pyroxd2 | pyridine nucleotide-disulphide oxidoreductase domain 2 | -3.13 |
| ENSMUSG00000027801 | Tm4sf4 | transmembrane 4 superfamily member 4 | -3.14 |
| ENSMUSG00000030207 | Fam234b | family with sequence similarity 234, member B | -3.14 |
| ENSMUSG00000055235 | Wdr86 | WD repeat domain 86 | -3.14 |
| ENSMUSG00000049036 | Tmem121 | transmembrane protein 121 | -3.14 |
| ENSMUSG00000055827 | Gsdmc3 | gasdermin C3 | -3.15 |
| ENSMUSG00000040121 | Rep15 | RAB15 effector protein | -3.15 |
| ENSMUSG00000038580 | Sct | secretin | -3.16 |
| ENSMUSG00000027254 | Map1a | microtubule-associated protein 1 A | -3.16 |
| ENSMUSG00000040432 | Ltb4r2 | leukotriene B4 receptor 2 | -3.16 |
| ENSMUSG00000078942 | Naip6 | NLR family, apoptosis inhibitory protein 6 | -3.17 |
| ENSMUSG00000031891 | Hsd11b2 | hydroxysteroid 11-beta dehydrogenase 2 | -3.17 |
| ENSMUSG00000054385 | Ceacam2 | carcinoembryonic antigen-related cell adhesion molecule 2 | -3.17 |
| ENSMUSG00000023092 | Fhl1 | four and a half LIM domains 1 | -3.17 |
| ENSMUSG00000089669 | Tnfsf13 | tumor necrosis factor (ligand) superfamily, member 13 | -3.17 |
| ENSMUSG00000110195 | Pde2a | phosphodiesterase 2A, cGMP-stimulated | -3.17 |
| ENSMUSG00000026834 | Acvr1c | activin A receptor, type IC | -3.18 |
| ENSMUSG00000096852 | Cyp2d12 | cytochrome P450, family 2, subfamily d, polypeptide 12 | -3.19 |
| ENSMUSG00000029445 | Hpd | 4-hydroxyphenylpyruvic acid dioxygenase | -3.20 |
| ENSMUSG00000017311 | Pyy | peptide YY | -3.20 |
| ENSMUSG00000053825 | Ppfia2 | protein tyrosine phosphatase, receptor type, f polypeptide (PTPRF), interacting protein (liprin), alpha | -3.21 |
| ENSMUSG00000070704 | Ugt2b36 | UDP glucuronosyltransferase 2 family, polypeptide B36 | -3.21 |

| | | | |
|---|---|---|---|
| ENSMUSG00000020589 | Fam49a | family with sequence similarity 49, member A | -3.22 |
| ENSMUSG00000034107 | Ano7 | anoctamin 7 | -3.23 |
| ENSMUSG00000056220 | Pla2g4a | phospholipase A2, group IVA (cytosolic, calcium-dependent) | -3.24 |
| ENSMUSG00000068154 | Insm1 | insulinoma-associated 1 | -3.24 |
| ENSMUSG00000047528 | Als2cr12 | amyotrophic lateral sclerosis 2 chromosome region 12 | -3.24 |
| ENSMUSG00000108950 | 9130015G15Rik | RIKEN cDNA 9130015G15 gene | -3.24 |
| ENSMUSG00000026832 | Cytip | cytohesin 1 interacting protein | -3.26 |
| ENSMUSG00000085651 | Gm11695 | predicted gene 11695 | -3.27 |
| ENSMUSG00000040896 | Kcnd3 | potassium voltage-gated channel, Shal-related family, member 3 | -3.28 |
| ENSMUSG00000052861 | Dnah6 | dynein, axonemal, heavy chain 6 | -3.28 |
| ENSMUSG00000107296 | Gm43500 | predicted gene 43500 | -3.28 |
| ENSMUSG00000095649 | Gm8979 | predicted gene 8979 | -3.29 |
| ENSMUSG00000047517 | Dmbt1 | deleted in malignant brain tumors 1 | -3.30 |
| ENSMUSG00000048217 | Nags | N-acetylglutamate synthase | -3.30 |
| ENSMUSG00000028415 | Spink4 | serine peptidase inhibitor, Kazal type 4 | -3.30 |
| ENSMUSG00000073043 | Atoh1 | atonal bHLH transcription factor 1 | -3.31 |
| ENSMUSG00000030911 | Zp2 | zona pellucida glycoprotein 2 | -3.31 |
| ENSMUSG00000037106 | Fer1l6 | fer-1-like 6 (C. elegans) | -3.32 |
| ENSMUSG00000042115 | Klhdc8a | kelch domain containing 8A | -3.32 |
| ENSMUSG00000015966 | Il17rb | interleukin 17 receptor B | -3.33 |
| ENSMUSG00000058618 | AY761184 | cDNA sequence AY761184 | -3.33 |
| ENSMUSG00000074196 | Clca4c-ps | chloride channel accessory 4C, pseudogene | -3.33 |
| ENSMUSG00000111377 | Gm40634 | predicted gene, 40634 | -3.33 |
| ENSMUSG00000031173 | Otc | ornithine transcarbamylase | -3.34 |
| ENSMUSG00000033740 | St18 | suppression of tumorigenicity 18 | -3.34 |
| ENSMUSG00000053004 | Hrh1 | histamine receptor H1 | -3.35 |
| ENSMUSG00000026815 | Gfi1b | growth factor independent 1B | -3.35 |
| ENSMUSG00000036027 | 1810046K07Rik | RIKEN cDNA 1810046K07 gene | -3.36 |
| ENSMUSG00000063903 | Klk1 | kallikrein 1 | -3.37 |
| ENSMUSG00000109685 | Gm45912 | predicted gene 45912 | -3.37 |
| ENSMUSG00000095026 | Gm3336 | predicted gene 3336 | -3.37 |
| ENSMUSG00000060176 | Kif27 | kinesin family member 27 | -3.37 |
| ENSMUSG00000101031 | Ms4a12 | membrane-spanning 4-domains, subfamily A, member 12 | -3.39 |
| ENSMUSG00000055567 | Unc80 | unc-80, NALCN activator | -3.39 |
| ENSMUSG00000111340 | Gm47171 | predicted gene, 47171 | -3.40 |
| ENSMUSG00000042351 | Grap2 | GRB2-related adaptor protein 2 | -3.40 |
| ENSMUSG00000019890 | Nts | neurotensin | -3.41 |
| ENSMUSG00000050556 | Kcnb1 | potassium voltage gated channel, Shab-related subfamily, member 1 | -3.41 |
| ENSMUSG00000020581 | Agr2 | anterior gradient 2 | -3.43 |
| ENSMUSG00000020279 | Il9r | interleukin 9 receptor | -3.43 |
| ENSMUSG00000008193 | Spib | Spi-B transcription factor (Spi-1/PU.1 related) | -3.44 |
| ENSMUSG00000050824 | Sstr5 | somatostatin receptor 5 | -3.44 |
| ENSMUSG00000034353 | Ramp1 | receptor (calcitonin) activity modifying protein 1 | -3.45 |
| ENSMUSG00000032015 | Pou2f3 | POU domain, class 2, transcription factor 3 | -3.45 |
| ENSMUSG00000096751 | Gm28373 | predicted gene 28373 | -3.46 |
| ENSMUSG00000050100 | Hmx2 | H6 homeobox 2 | -3.46 |
| ENSMUSG00000040680 | Kremen2 | kringle containing transmembrane protein 2 | -3.47 |
| ENSMUSG00000025991 | Cps1 | carbamoyl-phosphate synthetase 1 | -3.48 |
| ENSMUSG00000102615 | Gm37844 | predicted gene, 37844 | -3.48 |
| ENSMUSG00000087801 | Gm25980 | predicted gene, 25980 | -3.49 |
| ENSMUSG00000038665 | Dgki | diacylglycerol kinase, iota | -3.50 |
| ENSMUSG00000040136 | Abcc8 | ATP-binding cassette, sub-family C (CFTR/MRP), member 8 | -3.50 |
| ENSMUSG00000114576 | Naip3-ps1 | NLR family, apoptosis inhibitory protein 3, pseudogene 1 | -3.51 |
| ENSMUSG00000034701 | Neurod1 | neurogenic differentiation 1 | -3.51 |
| ENSMUSG00000004952 | Rasa4 | RAS p21 protein activator 4 | -3.52 |
| ENSMUSG00000026828 | Galnt5 | polypeptide N-acetylgalactosaminyltransferase 5 | -3.52 |
| ENSMUSG00000056553 | Ptprn2 | protein tyrosine phosphatase, receptor type, N polypeptide 2 | -3.52 |
| ENSMUSG00000027296 | Itpka | inositol 1,4,5-trisphosphate 3-kinase A | -3.52 |
| ENSMUSG00000022840 | Adcy5 | adenylate cyclase 5 | -3.55 |
| ENSMUSG00000026811 | St6galnac6 | ST6 (alpha-N-acetyl-neuraminyl-2,3-beta-galactosyl-1,3)-N-acetylgalactosaminide alpha-2,6-sialyltran | -3.56 |
| ENSMUSG00000046215 | Rprml | reprimo-like | -3.56 |
| ENSMUSG00000049350 | Zg16 | zymogen granule protein 16 | -3.58 |
| ENSMUSG00000047730 | Fcgbp | Fc fragment of IgG binding protein | -3.59 |
| ENSMUSG00000027762 | Sucnr1 | succinate receptor 1 | -3.59 |
| ENSMUSG00000110397 | Gm45540 | predicted gene 45540 | -3.59 |
| ENSMUSG00000012819 | Cdh23 | cadherin 23 (otocadherin) | -3.59 |
| ENSMUSG00000027797 | Dclk1 | doublecortin-like kinase 1 | -3.60 |
| ENSMUSG00000044156 | Hepacam2 | HEPACAM family member 2 | -3.61 |
| ENSMUSG00000051606 | 2010001K21Rik | RIKEN cDNA 2010001K21 gene | -3.61 |
| ENSMUSG00000086513 | 9130208D14Rik | RIKEN cDNA 9130208D14 gene | -3.63 |
| ENSMUSG00000022262 | Dnah5 | dynein, axonemal, heavy chain 5 | -3.63 |

| | | | |
|---|---|---|---|
| ENSMUSG00000020901 | Pik3r5 | phosphoinositide-3-kinase regulatory subunit 5 | -3.64 |
| ENSMUSG00000032561 | Acpp | acid phosphatase, prostate | -3.64 |
| ENSMUSG00000106651 | Gm42608 | predicted gene 42608 | -3.65 |
| ENSMUSG00000039264 | Gimap3 | GTPase, IMAP family member 3 | -3.66 |
| ENSMUSG00000035930 | Chst4 | carbohydrate (chondroitin 6/keratan) sulfotransferase 4 | -3.67 |
| ENSMUSG00000024029 | Tff3 | trefoil factor 3, intestinal | -3.68 |
| ENSMUSG00000046314 | Stxbp6 | syntaxin binding protein 6 (amisyn) | -3.68 |
| ENSMUSG00000028255 | Clca1 | chloride channel accessory 1 | -3.69 |
| ENSMUSG00000094219 | Calhm3 | calcium homeostasis modulator 3 | -3.69 |
| ENSMUSG00000020788 | Atp2a3 | ATPase, Ca++ transporting, ubiquitous | -3.71 |
| ENSMUSG00000040061 | Plcb2 | phospholipase C, beta 2 | -3.71 |
| ENSMUSG00000022148 | Fyb | FYN binding protein | -3.72 |
| ENSMUSG00000060063 | Alox5ap | arachidonate 5-lipoxygenase activating protein | -3.72 |
| ENSMUSG00000022686 | B3gnt5 | UDP-GlcNAc:betaGal beta-1,3-N-acetylglucosaminyltransferase 5 | -3.76 |
| ENSMUSG00000032036 | Kirrel3 | kirre like nephrin family adhesion molecule 3 | -3.77 |
| ENSMUSG00000044359 | P2ry4 | pyrimidinergic receptor P2Y, G-protein coupled, 4 | -3.78 |
| ENSMUSG00000050321 | Neto1 | neuropilin (NRP) and tolloid (TLL)-like 1 | -3.78 |
| ENSMUSG00000021730 | Hcn1 | hyperpolarization-activated, cyclic nucleotide-gated K+ 1 | -3.79 |
| ENSMUSG00000029865 | Sval1 | seminal vesicle antigen-like 1 | -3.79 |
| ENSMUSG00000009246 | Trpm5 | transient receptor potential cation channel, subfamily M, member 5 | -3.81 |
| ENSMUSG00000071178 | Serpina1b | serine (or cysteine) preptidase inhibitor, clade A, member 1B | -3.81 |
| ENSMUSG00000021700 | Rab3c | RAB3C, member RAS oncogene family | -3.81 |
| ENSMUSG00000054422 | Fabp1 | fatty acid binding protein 1, liver | -3.82 |
| ENSMUSG00000025515 | Muc2 | mucin 2 | -3.83 |
| ENSMUSG00000046460 | Sh2d7 | SH2 domain containing 7 | -3.86 |
| ENSMUSG00000028996 | Rbp7 | retinol binding protein 7, cellular | -3.87 |
| ENSMUSG00000024366 | Gfra3 | glial cell line derived neurotrophic factor family receptor alpha 3 | -3.88 |
| ENSMUSG00000004933 | Matk | megakaryocyte-associated tyrosine kinase | -3.89 |
| ENSMUSG00000026288 | Inpp5d | inositol polyphosphate-5-phosphatase D | -3.90 |
| ENSMUSG00000109237 | 9130214F15Rik | RIKEN cDNA 9130214F15 gene | -3.90 |
| ENSMUSG00000082956 | Naip3 | NLR family, apoptosis inhibitory protein 3 | -3.91 |
| ENSMUSG00000116929 | AC164088.2 | TEC | -3.91 |
| ENSMUSG00000114443 | Gm19241 | predicted gene, 19241 | -3.93 |
| ENSMUSG00000001827 | Folr1 | folate receptor 1 (adult) | -3.95 |
| ENSMUSG00000084384 | Gm12251 | predicted gene 12251 | -3.95 |
| ENSMUSG00000026395 | Ptprc | protein tyrosine phosphatase, receptor type, C | -3.96 |
| ENSMUSG00000030302 | Atp2b2 | ATPase, Ca++ transporting, plasma membrane 2 | -3.96 |
| ENSMUSG00000031377 | Bmx | BMX non-receptor tyrosine kinase | -3.97 |
| ENSMUSG00000029797 | Sspo | SCO-spondin | -3.97 |
| ENSMUSG00000025701 | Alox5 | arachidonate 5-lipoxygenase | -3.98 |
| ENSMUSG00000030263 | Lrmp | lymphoid-restricted membrane protein | -4.00 |
| ENSMUSG00000018927 | Ccl6 | chemokine (C-C motif) ligand 6 | -4.01 |
| ENSMUSG00000031981 | Capn9 | calpain 9 | -4.02 |
| ENSMUSG00000027168 | Pax6 | paired box 6 | -4.04 |
| ENSMUSG00000051079 | Rgs13 | regulator of G-protein signaling 13 | -4.05 |
| ENSMUSG00000050711 | Scg2 | secretogranin II | -4.06 |
| ENSMUSG00000090254 | Gm1965 | predicted gene 1965 | -4.08 |
| ENSMUSG00000058216 | Gstp3 | glutathione S-transferase pi 3 | -4.09 |
| ENSMUSG00000057400 | Ces1c | carboxylesterase 1C | -4.09 |
| ENSMUSG00000022026 | Olfm4 | olfactomedin 4 | -4.12 |
| ENSMUSG00000021069 | Pygl | liver glycogen phosphorylase | -4.12 |
| ENSMUSG00000051497 | Kcnj16 | potassium inwardly-rectifying channel, subfamily J, member 16 | -4.14 |
| ENSMUSG00000105302 | Gm19817 | predicted gene, 19817 | -4.14 |
| ENSMUSG00000039629 | Strip2 | striatin interacting protein 2 | -4.16 |
| ENSMUSG00000033220 | Rac2 | Rac family small GTPase 2 | -4.17 |
| ENSMUSG00000042010 | Acacb | acetyl-Coenzyme A carboxylase beta | -4.26 |
| ENSMUSG00000026531 | Mptx1 | mucosal pentraxin 1 | -4.26 |
| ENSMUSG00000040035 | Disp2 | dispatched RND tramsporter family member 2 | -4.28 |
| ENSMUSG00000040148 | Hmx3 | H6 homeobox 3 | -4.31 |
| ENSMUSG00000105975 | Gm42609 | predicted gene 42609 | -4.31 |
| ENSMUSG00000003283 | Hck | hemopoietic cell kinase | -4.32 |
| ENSMUSG00000029878 | Dbpht2 | DNA binding protein with his-thr domain | -4.32 |
| ENSMUSG00000052631 | Sh2d6 | SH2 domain containing 6 | -4.34 |
| ENSMUSG00000064140 | Trim38 | tripartite motif-containing 38 | -4.35 |
| ENSMUSG00000019429 | Ffar3 | free fatty acid receptor 3 | -4.38 |
| ENSMUSG00000028339 | Col15a1 | collagen, type XV, alpha 1 | -4.42 |
| ENSMUSG00000040380 | Cbln3 | cerebellin 3 precursor protein | -4.43 |
| ENSMUSG00000032826 | Ank2 | ankyrin 2, brain | -4.53 |
| ENSMUSG00000037973 | Ccdc129 | coiled-coil domain containing 129 | -4.54 |
| ENSMUSG00000047798 | Cd300lf | CD300 molecule like family member F | -4.57 |
| ENSMUSG00000020787 | P2rx1 | purinergic receptor P2X, ligand-gated ion channel, 1 | -4.64 |

| | | | |
|---|---|---|---|
| ENSMUSG00000039013 | Siglecf | sialic acid binding Ig-like lectin F | -4.65 |
| ENSMUSG00000096630 | Vmn2r26 | vomeronasal 2, receptor 26 | -4.68 |
| ENSMUSG00000033569 | Adgrb3 | adhesion G protein-coupled receptor B3 | -4.68 |
| ENSMUSG00000047497 | Adamts12 | a disintegrin-like and metallopeptidase (reprolysin type) with thrombospondin type 1 motif, 12 | -4.81 |
| ENSMUSG00000034923 | Ly6g6f | lymphocyte antigen 6 complex, locus G6F | -4.82 |
| ENSMUSG00000106354 | Gm42607 | predicted gene 42607 | -4.83 |
| ENSMUSG00000115816 | Gm34589 | predicted gene, 34589 | -4.95 |
| ENSMUSG00000037627 | Rgs22 | regulator of G-protein signalling 22 | -5.06 |
| ENSMUSG00000028777 | Gnat3 | guanine nucleotide binding protein, alpha transducing 3 | -5.06 |
| ENSMUSG00000022650 | Retnlb | resistin like beta | -5.10 |
| ENSMUSG00000021919 | Chat | choline acetyltransferase | -5.28 |
| ENSMUSG00000046318 | Ccbe1 | collagen and calcium binding EGF domains 1 | -5.28 |
| ENSMUSG00000032532 | Cck | cholecystokinin | -5.41 |
| ENSMUSG00000021536 | Adcy2 | adenylate cyclase 2 | -5.55 |
| ENSMUSG00000026938 | Fcna | ficolin A | -6.75 |
| ENSMUSG00000014351 | Gip | gastric inhibitory polypeptide | -6.98 |
| ENSMUSG00000000394 | Gcg | glucagon | -7.26 |
| ENSMUSG00000053852 | Adgrg4 | adhesion G protein-coupled receptor G4 | -7.27 |
| ENSMUSG00000026532 | Spta1 | spectrin alpha, erythrocytic 1 | -7.38 |

**Tom+Trop2+ vs. differentiated cells**

| Ensemble gene code | Symbol | Gene name | logFC |
|---|---|---|---|
| ENSMUSG00000031841 | Cdh13 | cadherin 13 | 11.80 |
| ENSMUSG00000029838 | Ptn | pleiotrophin | 10.90 |
| ENSMUSG00000028031 | Dkk2 | dickkopf WNT signaling pathway inhibitor 2 | 10.70 |
| ENSMUSG00000036480 | Prss56 | protease, serine 56 | 10.30 |
| ENSMUSG00000053613 | Notumos | notum palmitoleoyl-protein carboxylesterase, opposite strand | 10.20 |
| ENSMUSG00000037035 | Inhbb | inhibin beta-B | 10.20 |
| ENSMUSG00000025330 | Padi4 | peptidyl arginine deiminase, type IV | 10.10 |
| ENSMUSG00000029368 | Alb | albumin | 10.00 |
| ENSMUSG00000026167 | Wnt10a | wingless-type MMTV integration site family, member 10A | 9.81 |
| ENSMUSG00000017491 | Rarb | retinoic acid receptor, beta | 9.68 |
| ENSMUSG00000063632 | Sox11 | SRY (sex determining region Y)-box 11 | 9.52 |
| ENSMUSG00000040852 | Plekhh2 | pleckstrin homology domain containing, family H (with MyTH4 domain) member 2 | 9.38 |
| ENSMUSG00000029648 | Flt1 | FMS-like tyrosine kinase 1 | 9.29 |
| ENSMUSG00000028360 | Slc44a5 | solute carrier family 44, member 5 | 9.24 |
| ENSMUSG00000031871 | Cdh5 | cadherin 5 | 9.22 |
| ENSMUSG00000022883 | Robo1 | roundabout guidance receptor 1 | 9.05 |
| ENSMUSG00000035615 | Frmpd1 | FERM and PDZ domain containing 1 | 8.94 |
| ENSMUSG00000042268 | Slc26a9 | solute carrier family 26, member 9 | 8.87 |
| ENSMUSG00000110290 | Gm45336 | predicted gene 45336 | 8.87 |
| ENSMUSG00000032965 | Ift57 | intraflagellar transport 57 | 8.85 |
| ENSMUSG00000048960 | Prex2 | phosphatidylinositol-3,4,5-trisphosphate-dependent Rac exchange factor 2 | 8.82 |
| ENSMUSG00000051397 | Tacstd2 | tumor-associated calcium signal transducer 2 | 8.81 |
| ENSMUSG00000020181 | Nav3 | neuron navigator 3 | 8.81 |
| ENSMUSG00000068699 | Flnc | filamin C, gamma | 8.79 |
| ENSMUSG00000020388 | Pdlim4 | PDZ and LIM domain 4 | 8.70 |
| ENSMUSG00000005503 | Evx1 | even-skipped homeobox 1 | 8.70 |
| ENSMUSG00000054263 | Lifr | leukemia inhibitory factor receptor | 8.69 |
| ENSMUSG00000039232 | Stx11 | syntaxin 11 | 8.66 |
| ENSMUSG00000055322 | Tns1 | tensin 1 | 8.50 |
| ENSMUSG00000041046 | Ramp3 | receptor (calcitonin) activity modifying protein 3 | 8.46 |
| ENSMUSG00000013584 | Aldh1a2 | aldehyde dehydrogenase family 1, subfamily A2 | 8.46 |
| ENSMUSG00000079330 | Lemd1 | LEM domain containing 1 | 8.43 |
| ENSMUSG00000000120 | Ngfr | nerve growth factor receptor (TNFR superfamily, member 16) | 8.42 |
| ENSMUSG00000032936 | Camkv | CaM kinase-like vesicle-associated | 8.37 |
| ENSMUSG00000026117 | Zap70 | zeta-chain (TCR) associated protein kinase | 8.30 |
| ENSMUSG00000016763 | Scube1 | signal peptide, CUB domain, EGF-like 1 | 8.28 |
| ENSMUSG00000023885 | Thbs2 | thrombospondin 2 | 8.21 |
| ENSMUSG00000059854 | Hydin | HYDIN, axonemal central pair apparatus protein | 8.21 |
| ENSMUSG00000071531 | Gprin2 | G protein regulated inducer of neurite outgrowth 2 | 8.19 |
| ENSMUSG00000110187 | Gm45496 | predicted gene 45496 | 8.17 |
| ENSMUSG00000022754 | Tmem45a | transmembrane protein 45a | 8.17 |
| ENSMUSG00000054555 | Adam12 | a disintegrin and metallopeptidase domain 12 (meltrin alpha) | 8.15 |
| ENSMUSG00000034460 | Six4 | sine oculis-related homeobox 4 | 8.11 |
| ENSMUSG00000004552 | Ctse | cathepsin E | 8.05 |
| ENSMUSG00000084822 | Myadml2os | myeloid-associated differentiation marker-like 2, opposite strand | 8.03 |
| ENSMUSG00000034634 | Ly6d | lymphocyte antigen 6 complex, locus D | 7.98 |
| ENSMUSG00000029869 | Ephb6 | Eph receptor B6 | 7.95 |
| ENSMUSG00000063531 | Sema3e | sema domain, immunoglobulin domain (Ig), short basic domain, secreted, (semaphorin) 3E | 7.87 |
| ENSMUSG00000012428 | Steap4 | STEAP family member 4 | 7.86 |
| ENSMUSG00000022037 | Clu | clusterin | 7.85 |
| ENSMUSG00000044313 | Mab21l3 | mab-21-like 3 (C. elegans) | 7.74 |
| ENSMUSG00000020218 | Wif1 | Wnt inhibitory factor 1 | 7.67 |
| ENSMUSG00000070867 | Trabd2b | TraB domain containing 2B | 7.67 |
| ENSMUSG00000033377 | Palmd | palmdelphin | 7.65 |
| ENSMUSG00000013415 | Igf2bp1 | insulin-like growth factor 2 mRNA binding protein 1 | 7.61 |
| ENSMUSG00000062591 | Tubb4a | tubulin, beta 4A class IVA | 7.60 |
| ENSMUSG00000032719 | Sbspon | somatomedin B and thrombospondin, type 1 domain containing | 7.48 |
| ENSMUSG00000043903 | Zfp469 | zinc finger protein 469 | 7.47 |
| ENSMUSG00000030772 | Dkk3 | dickkopf WNT signaling pathway inhibitor 3 | 7.45 |
| ENSMUSG00000049848 | Ceacam19 | carcinoembryonic antigen-related cell adhesion molecule 19 | 7.40 |
| ENSMUSG00000041134 | Cyyr1 | cysteine and tyrosine-rich protein 1 | 7.37 |
| ENSMUSG00000025491 | Ifitm1 | interferon induced transmembrane protein 1 | 7.37 |
| ENSMUSG00000018211 | Wfdc15b | WAP four-disulfide core domain 15B | 7.37 |
| ENSMUSG00000038415 | Foxq1 | forkhead box Q1 | 7.34 |
| ENSMUSG00000029223 | Uchl1 | ubiquitin carboxy-terminal hydrolase L1 | 7.29 |
| ENSMUSG00000022146 | Osmr | oncostatin M receptor | 7.29 |
| ENSMUSG00000042988 | Notum | notum palmitoleoyl-protein carboxylesterase | 7.28 |
| ENSMUSG00000031074 | Fgf3 | fibroblast growth factor 3 | 7.24 |
| ENSMUSG00000032502 | Stac | src homology three (SH3) and cysteine rich domain | 7.23 |
| ENSMUSG00000038572 | Bpifb5 | BPI fold containing family B, member 5 | 7.22 |
| ENSMUSG00000031284 | Pak3 | p21 (RAC1) activated kinase 3 | 7.17 |
| ENSMUSG00000031503 | Col4a2 | collagen, type IV, alpha 2 | 7.17 |
| ENSMUSG00000022464 | Slc38a4 | solute carrier family 38, member 4 | 7.16 |

| | | | |
|---|---|---|---|
| ENSMUSG00000034463 | Scara3 | scavenger receptor class A, member 3 | 7.13 |
| ENSMUSG00000036599 | Chst12 | carbohydrate sulfotransferase 12 | 7.08 |
| ENSMUSG00000026822 | Lcn2 | lipocalin 2 | 7.05 |
| ENSMUSG00000021091 | Serpina3n | serine (or cysteine) peptidase inhibitor, clade A, member 3N | 7.01 |
| ENSMUSG00000037712 | Fermt2 | fermitin family member 2 | 6.97 |
| ENSMUSG00000059898 | Dsc3 | desmocollin 3 | 6.97 |
| ENSMUSG00000014813 | Stc1 | stanniocalcin 1 | 6.97 |
| ENSMUSG00000020363 | Gfpt2 | glutamine fructose-6-phosphate transaminase 2 | 6.96 |
| ENSMUSG00000056856 | Jakmip3 | janus kinase and microtubule interacting protein 3 | 6.96 |
| ENSMUSG00000048078 | Tenm4 | teneurin transmembrane protein 4 | 6.94 |
| ENSMUSG00000029371 | Cxcl5 | chemokine (C-X-C motif) ligand 5 | 6.94 |
| ENSMUSG00000020037 | Rfx4 | regulatory factor X, 4 (influences HLA class II expression) | 6.94 |
| ENSMUSG00000108218 | Olfr1372-ps1 | olfactory receptor 1372, pseudogene 1 | 6.90 |
| ENSMUSG00000031661 | Nkd1 | naked cuticle 1 | 6.89 |
| ENSMUSG00000053007 | Creb5 | cAMP responsive element binding protein 5 | 6.86 |
| ENSMUSG00000032717 | Mdfi | MyoD family inhibitor | 6.85 |
| ENSMUSG00000054932 | Afp | alpha fetoprotein | 6.85 |
| ENSMUSG00000085811 | Cep112it | centrosomal protein 112, intronic transcript | 6.77 |
| ENSMUSG00000015134 | Aldh1a3 | aldehyde dehydrogenase family 1, subfamily A3 | 6.76 |
| ENSMUSG00000032327 | Stra6 | stimulated by retinoic acid gene 6 | 6.74 |
| ENSMUSG00000044071 | Fam19a2 | family with sequence similarity 19, member A2 | 6.73 |
| ENSMUSG00000061048 | Cdh3 | cadherin 3 | 6.71 |
| ENSMUSG00000014599 | Csf1 | colony stimulating factor 1 (macrophage) | 6.70 |
| ENSMUSG00000110666 | Gm9172 | predicted gene 9172 | 6.69 |
| ENSMUSG00000004951 | Hspb1 | heat shock protein 1 | 6.68 |
| ENSMUSG00000032186 | Tmod2 | tropomodulin 2 | 6.67 |
| ENSMUSG00000031736 | Crnde | colorectal neoplasia differentially expressed (non-protein coding) | 6.66 |
| ENSMUSG00000006205 | Htra1 | HtrA serine peptidase 1 | 6.64 |
| ENSMUSG00000024810 | Il33 | interleukin 33 | 6.59 |
| ENSMUSG00000001864 | Aif1l | allograft inflammatory factor 1-like | 6.59 |
| ENSMUSG00000049866 | Arl4c | ADP-ribosylation factor-like 4C | 6.58 |
| ENSMUSG00000042734 | Ttc9 | tetratricopeptide repeat domain 9 | 6.58 |
| ENSMUSG00000046623 | Gjb4 | gap junction protein, beta 4 | 6.57 |
| ENSMUSG00000027376 | Prom2 | prominin 2 | 6.56 |
| ENSMUSG00000000901 | Mmp11 | matrix metallopeptidase 11 | 6.55 |
| ENSMUSG00000040488 | Ltbp4 | latent transforming growth factor beta binding protein 4 | 6.54 |
| ENSMUSG00000031595 | Pdgfrl | platelet-derived growth factor receptor-like | 6.54 |
| ENSMUSG00000022176 | Rem2 | rad and gem related GTP binding protein 2 | 6.54 |
| ENSMUSG00000000983 | Wfdc18 | WAP four-disulfide core domain 18 | 6.53 |
| ENSMUSG00000019577 | Pdk4 | pyruvate dehydrogenase kinase, isoenzyme 4 | 6.53 |
| ENSMUSG00000037016 | Frem2 | Fras1 related extracellular matrix protein 2 | 6.51 |
| ENSMUSG00000029769 | Ccdc136 | coiled-coil domain containing 136 | 6.48 |
| ENSMUSG00000029108 | Pcdh7 | protocadherin 7 | 6.47 |
| ENSMUSG00000046169 | Adamts6 | a disintegrin-like and metallopeptidase (reprolysin type) with thrombospondin type 1 motif, 6 | 6.47 |
| ENSMUSG00000001131 | Timp1 | tissue inhibitor of metalloproteinase 1 | 6.45 |
| ENSMUSG00000042357 | Gjb5 | gap junction protein, beta 5 | 6.44 |
| ENSMUSG00000025352 | Gdf11 | growth differentiation factor 11 | 6.42 |
| ENSMUSG00000032643 | Fhl3 | four and a half LIM domains 3 | 6.39 |
| ENSMUSG00000021253 | Tgfb3 | transforming growth factor, beta 3 | 6.37 |
| ENSMUSG00000051159 | Cited1 | Cbp/p300-interacting transactivator with Glu/Asp-rich carboxy-terminal domain 1 | 6.36 |
| ENSMUSG00000022995 | Enah | ENAH actin regulator | 6.36 |
| ENSMUSG00000025202 | Scd3 | stearoyl-coenzyme A desaturase 3 | 6.36 |
| ENSMUSG00000051236 | Msrb3 | methionine sulfoxide reductase B3 | 6.35 |
| ENSMUSG00000097651 | 4930461G14Rik | RIKEN cDNA 4930461G14 gene | 6.35 |
| ENSMUSG00000058297 | Spock2 | sparc/osteonectin, cwcv and kazal-like domains proteoglycan 2 | 6.33 |
| ENSMUSG00000024268 | Celf4 | CUGBP, Elav-like family member 4 | 6.32 |
| ENSMUSG00000042179 | Pnliprp1 | pancreatic lipase related protein 1 | 6.31 |
| ENSMUSG00000021822 | Plau | plasminogen activator, urokinase | 6.31 |
| ENSMUSG00000040836 | Gpr161 | G protein-coupled receptor 161 | 6.29 |
| ENSMUSG00000025473 | Adam8 | a disintegrin and metallopeptidase domain 8 | 6.28 |
| ENSMUSG00000037129 | Tmprss13 | transmembrane protease, serine 13 | 6.22 |
| ENSMUSG00000022425 | Enpp2 | ectonucleotide pyrophosphatase/phosphodiesterase 2 | 6.22 |
| ENSMUSG00000052957 | Gas1 | growth arrest specific 1 | 6.20 |
| ENSMUSG00000062327 | T | brachyury, T-box transcription factor T | 6.19 |
| ENSMUSG00000038860 | Garnl3 | GTPase activating RANGAP domain-like 3 | 6.18 |
| ENSMUSG00000034220 | Gpc1 | glypican 1 | 6.16 |
| ENSMUSG00000049807 | Arhgap23 | Rho GTPase activating protein 23 | 6.15 |
| ENSMUSG00000034450 | Gulo | gulonolactone (L-) oxidase | 6.15 |
| ENSMUSG00000097365 | C030034L19Rik | RIKEN cDNA C030034L19 gene | 6.15 |
| ENSMUSG00000001870 | Ltbp1 | latent transforming growth factor beta binding protein 1 | 6.12 |
| ENSMUSG00000063727 | Tnfrsf11b | tumor necrosis factor receptor superfamily, member 11b (osteoprotegerin) | 6.08 |
| ENSMUSG00000026764 | Kif5c | kinesin family member 5C | 6.08 |
| ENSMUSG00000015053 | Gata2 | GATA binding protein 2 | 6.08 |
| ENSMUSG00000103738 | Gm37652 | predicted gene, 37652 | 6.06 |
| ENSMUSG00000066607 | Insyn1 | inhibitory synaptic factor 1 | 6.01 |
| ENSMUSG00000050052 | Tdrp | testis development related protein | 6.00 |

| | | | |
|---|---|---|---|
| ENSMUSG00000050201 | Otop2 | otopetrin 2 | 6.00 |
| ENSMUSG00000071847 | Apcdd1 | adenomatosis polyposis coli down-regulated 1 | 5.99 |
| ENSMUSG00000025938 | Slco5a1 | solute carrier organic anion transporter family, member 5A1 | 5.96 |
| ENSMUSG00000078161 | Erich3 | glutamate rich 3 | 5.95 |
| ENSMUSG00000025324 | Atp10a | ATPase, class V, type 10A | 5.94 |
| ENSMUSG00000034684 | Sema3f | sema domain, immunoglobulin domain (Ig), short basic domain, secreted, (semaphorin) 3F | 5.93 |
| ENSMUSG00000042804 | Gpr153 | G protein-coupled receptor 153 | 5.89 |
| ENSMUSG00000027985 | Lef1 | lymphoid enhancer binding factor 1 | 5.88 |
| ENSMUSG00000014329 | Bicc1 | BicC family RNA binding protein 1 | 5.88 |
| ENSMUSG00000010751 | Tnfrsf22 | tumor necrosis factor receptor superfamily, member 22 | 5.86 |
| ENSMUSG00000040998 | Npnt | nephronectin | 5.85 |
| ENSMUSG00000038146 | Notch3 | notch 3 | 5.84 |
| ENSMUSG00000028497 | Hacd4 | 3-hydroxyacyl-CoA dehydratase 4 | 5.83 |
| ENSMUSG00000110802 | Gm47141 | predicted gene, 47141 | 5.83 |
| ENSMUSG00000020086 | H2afy2 | H2A histone family, member Y2 | 5.82 |
| ENSMUSG00000085412 | Halr1 | Hoxa adjacent long noncoding RNA 1 | 5.81 |
| ENSMUSG00000042514 | Klhl14 | kelch-like 14 | 5.79 |
| ENSMUSG00000007097 | Atp1a2 | ATPase, Na+/K+ transporting, alpha 2 polypeptide | 5.77 |
| ENSMUSG00000013338 | Fer1l4 | fer-1-like 4 (C. elegans) | 5.77 |
| ENSMUSG00000022665 | Ccdc80 | coiled-coil domain containing 80 | 5.76 |
| ENSMUSG00000023008 | Fmnl3 | formin-like 3 | 5.75 |
| ENSMUSG00000005883 | Spo11 | SPO11 meiotic protein covalently bound to DSB | 5.75 |
| ENSMUSG00000078235 | Fam43b | family with sequence similarity 43, member B | 5.74 |
| ENSMUSG00000027547 | Sall4 | spalt like transcription factor 4 | 5.70 |
| ENSMUSG00000052516 | Robo2 | roundabout guidance receptor 2 | 5.69 |
| ENSMUSG00000108145 | Gm38811 | predicted gene, 38811 | 5.69 |
| ENSMUSG00000051596 | Otop1 | otopetrin 1 | 5.69 |
| ENSMUSG00000026193 | Fn1 | fibronectin 1 | 5.68 |
| ENSMUSG00000027210 | Meis2 | Meis homeobox 2 | 5.64 |
| ENSMUSG00000017466 | Timp2 | tissue inhibitor of metalloproteinase 2 | 5.58 |
| ENSMUSG00000026579 | F5 | coagulation factor V | 5.58 |
| ENSMUSG00000040289 | Hey1 | hairy/enhancer-of-split related with YRPW motif 1 | 5.58 |
| ENSMUSG00000033508 | Asprv1 | aspartic peptidase, retroviral-like 1 | 5.58 |
| ENSMUSG00000048191 | Muc6 | mucin 6, gastric | 5.56 |
| ENSMUSG00000041202 | Pla2g2d | phospholipase A2, group IID | 5.56 |
| ENSMUSG00000113642 | Gm49329 | predicted gene, 49329 | 5.55 |
| ENSMUSG00000097767 | Miat | myocardial infarction associated transcript (non-protein coding) | 5.52 |
| ENSMUSG00000045667 | Smtnl2 | smoothelin-like 2 | 5.50 |
| ENSMUSG00000026728 | Vim | vimentin | 5.45 |
| ENSMUSG00000029757 | Dync1i1 | dynein cytoplasmic 1 intermediate chain 1 | 5.45 |
| ENSMUSG00000025089 | Gfra1 | glial cell line derived neurotrophic factor family receptor alpha 1 | 5.45 |
| ENSMUSG00000039239 | Tgfb2 | transforming growth factor, beta 2 | 5.44 |
| ENSMUSG00000022548 | Apod | apolipoprotein D | 5.44 |
| ENSMUSG00000053049 | Gm15413 | predicted gene 15413 | 5.42 |
| ENSMUSG00000029868 | Trpv6 | transient receptor potential cation channel, subfamily V, member 6 | 5.41 |
| ENSMUSG00000022456 | Sept3 | septin 3 | 5.38 |
| ENSMUSG00000075033 | Nxpe3 | neurexophilin and PC-esterase domain family, member 3 | 5.37 |
| ENSMUSG00000085903 | Gm15340 | predicted gene 15340 | 5.36 |
| ENSMUSG00000028246 | Faxc | failed axon connections homolog | 5.35 |
| ENSMUSG00000033227 | Wnt6 | wingless-type MMTV integration site family, member 6 | 5.35 |
| ENSMUSG00000091055 | Siglec15 | sialic acid binding Ig-like lectin 15 | 5.34 |
| ENSMUSG00000106073 | Gm42892 | predicted gene 42892 | 5.34 |
| ENSMUSG00000027858 | Tspan2 | tetraspanin 2 | 5.33 |
| ENSMUSG00000035678 | Tnfsf9 | tumor necrosis factor (ligand) superfamily, member 9 | 5.32 |
| ENSMUSG00000027931 | Npr1 | natriuretic peptide receptor 1 | 5.32 |
| ENSMUSG00000044562 | Rasip1 | Ras interacting protein 1 | 5.31 |
| ENSMUSG00000041801 | Phlda3 | pleckstrin homology like domain, family A, member 3 | 5.30 |
| ENSMUSG00000026620 | Mark1 | MAP/microtubule affinity regulating kinase 1 | 5.29 |
| ENSMUSG00000052221 | Ppp1r36 | protein phosphatase 1, regulatory subunit 36 | 5.29 |
| ENSMUSG00000040663 | Clcf1 | cardiotrophin-like cytokine factor 1 | 5.28 |
| ENSMUSG00000045991 | Onecut2 | one cut domain, family member 2 | 5.26 |
| ENSMUSG00000060548 | Tnfrsf19 | tumor necrosis factor receptor superfamily, member 19 | 5.25 |
| ENSMUSG00000036412 | Arsi | arylsulfatase i | 5.25 |
| ENSMUSG00000035164 | Zc3h12c | zinc finger CCCH type containing 12C | 5.23 |
| ENSMUSG00000067818 | Myl9 | myosin, light polypeptide 9, regulatory | 5.21 |
| ENSMUSG00000032496 | Ltf | lactotransferrin | 5.21 |
| ENSMUSG00000068923 | Syt11 | synaptotagmin XI | 5.20 |
| ENSMUSG00000063011 | Msln | mesothelin | 5.20 |
| ENSMUSG00000014602 | Kif1a | kinesin family member 1A | 5.18 |
| ENSMUSG00000024659 | Anxa1 | annexin A1 | 5.18 |
| ENSMUSG00000022371 | Col14a1 | collagen, type XIV, alpha 1 | 5.18 |
| ENSMUSG00000029417 | Cxcl9 | chemokine (C-X-C motif) ligand 9 | 5.18 |
| ENSMUSG00000040415 | Dtx3 | deltex 3, E3 ubiquitin ligase | 5.15 |
| ENSMUSG00000046470 | Sox18 | SRY (sex determining region Y)-box 18 | 5.15 |
| ENSMUSG00000031538 | Plat | plasminogen activator, tissue | 5.13 |
| ENSMUSG00000050382 | Kif7 | kinesin family member 7 | 5.12 |

| | | | |
|---|---|---|---|
| ENSMUSG00000074813 | Gm14005 | predicted gene 14005 | 5.12 |
| ENSMUSG00000055333 | Fat2 | FAT atypical cadherin 2 | 5.11 |
| ENSMUSG00000020431 | Adcy1 | adenylate cyclase 1 | 5.10 |
| ENSMUSG00000018340 | Anxa6 | annexin A6 | 5.09 |
| ENSMUSG00000081016 | Olfr1397-ps1 | olfactory receptor 1397, pseudogene 1 | 5.07 |
| ENSMUSG00000000031 | H19 | H19, imprinted maternally expressed transcript | 5.06 |
| ENSMUSG00000031737 | Irx5 | Iroquois homeobox 5 | 5.06 |
| ENSMUSG00000009097 | Tbx1 | T-box 1 | 5.05 |
| ENSMUSG00000025666 | Tmem47 | transmembrane protein 47 | 5.05 |
| ENSMUSG00000070337 | Gpr179 | G protein-coupled receptor 179 | 5.04 |
| ENSMUSG00000036502 | Tmem255a | transmembrane protein 255A | 5.03 |
| ENSMUSG00000022594 | Lynx1 | Ly6/neurotoxin 1 | 5.00 |
| ENSMUSG00000006576 | Slc4a3 | solute carrier family 4 (anion exchanger), member 3 | 5.00 |
| ENSMUSG00000020427 | Igfbp3 | insulin-like growth factor binding protein 3 | 4.99 |
| ENSMUSG00000039323 | Igfbp2 | insulin-like growth factor binding protein 2 | 4.98 |
| ENSMUSG00000100733 | 4932411K12Rik | RIKEN cDNA 4932411K12 gene | 4.98 |
| ENSMUSG00000081392 | Gm11668 | predicted gene 11668 | 4.97 |
| ENSMUSG00000030844 | Rgs10 | regulator of G-protein signalling 10 | 4.97 |
| ENSMUSG00000035062 | Zc4h2 | zinc finger, C4H2 domain containing | 4.97 |
| ENSMUSG00000031734 | Irx3 | Iroquois related homeobox 3 | 4.95 |
| ENSMUSG00000050640 | Tmem150c | transmembrane protein 150C | 4.94 |
| ENSMUSG00000040430 | Pitpnc1 | phosphatidylinositol transfer protein, cytoplasmic 1 | 4.91 |
| ENSMUSG00000000552 | Zfp385a | zinc finger protein 385A | 4.91 |
| ENSMUSG00000034675 | Dbn1 | drebrin 1 | 4.91 |
| ENSMUSG00000038496 | Slc19a3 | solute carrier family 19, member 3 | 4.91 |
| ENSMUSG00000034685 | Fam171a2 | family with sequence similarity 171, member A2 | 4.91 |
| ENSMUSG00000030873 | Scnn1b | sodium channel, nonvoltage-gated 1 beta | 4.86 |
| ENSMUSG00000029380 | Cxcl1 | chemokine (C-X-C motif) ligand 1 | 4.84 |
| ENSMUSG00000068748 | Ptprz1 | protein tyrosine phosphatase, receptor type Z, polypeptide 1 | 4.84 |
| ENSMUSG00000032875 | Arhgef17 | Rho guanine nucleotide exchange factor (GEF) 17 | 4.83 |
| ENSMUSG00000043430 | Psapl1 | prosaposin-like 1 | 4.83 |
| ENSMUSG00000032625 | Thsd7a | thrombospondin, type I, domain containing 7A | 4.82 |
| ENSMUSG00000021950 | Anxa8 | annexin A8 | 4.82 |
| ENSMUSG00000022435 | Upk3a | uroplakin 3A | 4.82 |
| ENSMUSG00000035783 | Acta2 | actin, alpha 2, smooth muscle, aorta | 4.82 |
| ENSMUSG00000029359 | Tesc | tescalcin | 4.81 |
| ENSMUSG00000058952 | Cfi | complement component factor i | 4.79 |
| ENSMUSG00000017204 | Gsdma | gasdermin A | 4.79 |
| ENSMUSG00000020937 | Plcd3 | phospholipase C, delta 3 | 4.79 |
| ENSMUSG00000010175 | Prox1 | prospero homeobox 1 | 4.78 |
| ENSMUSG00000000093 | Tbx2 | T-box 2 | 4.78 |
| ENSMUSG00000030351 | Tspan11 | tetraspanin 11 | 4.78 |
| ENSMUSG00000027485 | Bpifb1 | BPI fold containing family B, member 1 | 4.78 |
| ENSMUSG00000038775 | Vill | villin-like | 4.77 |
| ENSMUSG00000032076 | Cadm1 | cell adhesion molecule 1 | 4.76 |
| ENSMUSG00000045062 | Pcdhb7 | protocadherin beta 7 | 4.76 |
| ENSMUSG00000027570 | Col9a3 | collagen, type IX, alpha 3 | 4.75 |
| ENSMUSG00000052105 | Mtcl1 | microtubule crosslinking factor 1 | 4.73 |
| ENSMUSG00000006931 | P3h4 | prolyl 3-hydroxylase family member 4 (non-enzymatic) | 4.73 |
| ENSMUSG00000069170 | Adgrv1 | adhesion G protein-coupled receptor V1 | 4.73 |
| ENSMUSG00000015647 | Lama5 | laminin, alpha 5 | 4.72 |
| ENSMUSG00000020844 | Nxn | nucleoredoxin | 4.72 |
| ENSMUSG00000046618 | Olfml2a | olfactomedin-like 2A | 4.72 |
| ENSMUSG00000072437 | Nanos1 | nanos C2HC-type zinc finger 1 | 4.71 |
| ENSMUSG00000038456 | Dennd2a | DENN/MADD domain containing 2A | 4.70 |
| ENSMUSG00000062151 | Unc13c | unc-13 homolog C | 4.69 |
| ENSMUSG00000020814 | Mxra7 | matrix-remodelling associated 7 | 4.68 |
| ENSMUSG00000034295 | Fhod3 | formin homology 2 domain containing 3 | 4.68 |
| ENSMUSG00000025492 | Ifitm3 | interferon induced transmembrane protein 3 | 4.67 |
| ENSMUSG00000039607 | Rbms3 | RNA binding motif, single stranded interacting protein | 4.67 |
| ENSMUSG00000028626 | Col9a2 | collagen, type IX, alpha 2 | 4.67 |
| ENSMUSG00000037188 | Grhl3 | grainyhead like transcription factor 3 | 4.66 |
| ENSMUSG00000004044 | Cavin1 | caveolae associated 1 | 4.66 |
| ENSMUSG00000106709 | Gm30270 | predicted gene, 30270 | 4.66 |
| ENSMUSG00000032492 | Pth1r | parathyroid hormone 1 receptor | 4.64 |
| ENSMUSG00000039959 | Hip1 | huntingtin interacting protein 1 | 4.63 |
| ENSMUSG00000026065 | Slc9a4 | solute carrier family 9 (sodium/hydrogen exchanger), member 4 | 4.62 |
| ENSMUSG00000038295 | Atg9b | autophagy related 9B | 4.61 |
| ENSMUSG00000059401 | Mamld1 | mastermind-like domain containing 1 | 4.61 |
| ENSMUSG00000048450 | Msx1 | msh homeobox 1 | 4.60 |
| ENSMUSG00000050989 | Selenon | selenoprotein N | 4.60 |
| ENSMUSG00000024164 | C3 | complement component 3 | 4.56 |
| ENSMUSG00000044034 | Npb | neuropeptide B | 4.56 |
| ENSMUSG00000030796 | Tead2 | TEA domain family member 2 | 4.55 |
| ENSMUSG00000037813 | D630003M21Rik | RIKEN cDNA D630003M21 gene | 4.55 |
| ENSMUSG00000021539 | Lect2 | leukocyte cell-derived chemotaxin 2 | 4.55 |

| | | | |
|---|---|---|---|
| ENSMUSG00000021765 | Fst | follistatin | 4.55 |
| ENSMUSG00000112871 | Gm35404 | predicted gene, 35404 | 4.55 |
| ENSMUSG00000021763 | BC067074 | cDNA sequence BC067074 | 4.54 |
| ENSMUSG00000050192 | Eif5a2 | eukaryotic translation initiation factor 5A2 | 4.53 |
| ENSMUSG00000025328 | Padi3 | peptidyl arginine deiminase, type III | 4.53 |
| ENSMUSG00000031328 | Flna | filamin, alpha | 4.52 |
| ENSMUSG00000027803 | Wwtr1 | WW domain containing transcription regulator 1 | 4.52 |
| ENSMUSG00000073599 | Ecscr | endothelial cell surface expressed chemotaxis and apoptosis regulator | 4.52 |
| ENSMUSG00000059336 | Slc14a1 | solute carrier family 14 (urea transporter), member 1 | 4.50 |
| ENSMUSG00000038132 | Rbm24 | RNA binding motif protein 24 | 4.50 |
| ENSMUSG00000070436 | Serpinh1 | serine (or cysteine) peptidase inhibitor, clade H, member 1 | 4.50 |
| ENSMUSG00000060429 | Sntb1 | syntrophin, basic 1 | 4.49 |
| ENSMUSG00000030246 | Ldhb | lactate dehydrogenase B | 4.49 |
| ENSMUSG00000044033 | Ccdc141 | coiled-coil domain containing 141 | 4.48 |
| ENSMUSG00000030048 | Gkn3 | gastrokine 3 | 4.48 |
| ENSMUSG00000021986 | Amer2 | APC membrane recruitment 2 | 4.47 |
| ENSMUSG00000028883 | Sema3a | sema domain, immunoglobulin domain (Ig), short basic domain, secreted, (semaphorin) 3A | 4.43 |
| ENSMUSG00000004668 | Abca13 | ATP-binding cassette, sub-family A (ABC1), member 13 | 4.43 |
| ENSMUSG00000067889 | Sptbn2 | spectrin beta, non-erythrocytic 2 | 4.42 |
| ENSMUSG00000040152 | Thbs1 | thrombospondin 1 | 4.42 |
| ENSMUSG00000026888 | Grb14 | growth factor receptor bound protein 14 | 4.42 |
| ENSMUSG00000086484 | Nron | non-protein coding RNA, repressor of NFAT | 4.42 |
| ENSMUSG00000103636 | Gm38331 | predicted gene, 38331 | 4.39 |
| ENSMUSG00000038180 | Spag4 | sperm associated antigen 4 | 4.36 |
| ENSMUSG00000008398 | Elk3 | ELK3, member of ETS oncogene family | 4.34 |
| ENSMUSG00000031355 | Arhgap6 | Rho GTPase activating protein 6 | 4.34 |
| ENSMUSG00000072591 | 5930412G12Rik | RIKEN cDNA 5930412G12 gene | 4.34 |
| ENSMUSG00000026676 | Ccdc3 | coiled-coil domain containing 3 | 4.33 |
| ENSMUSG00000024502 | Jakmip2 | janus kinase and microtubule interacting protein 2 | 4.33 |
| ENSMUSG00000017897 | Eya2 | EYA transcriptional coactivator and phosphatase 2 | 4.32 |
| ENSMUSG00000068327 | Tlx2 | T cell leukemia, homeobox 2 | 4.32 |
| ENSMUSG00000013089 | Etv5 | ets variant 5 | 4.29 |
| ENSMUSG00000033763 | Mtss1l | metastasis suppressor 1-like | 4.29 |
| ENSMUSG00000028328 | Tmod1 | tropomodulin 1 | 4.29 |
| ENSMUSG00000042759 | Apobr | apolipoprotein B receptor | 4.29 |
| ENSMUSG00000019997 | Ctgf | connective tissue growth factor | 4.29 |
| ENSMUSG00000020092 | Pald1 | phosphatase domain containing, paladin 1 | 4.27 |
| ENSMUSG00000063605 | Ccdc102a | coiled-coil domain containing 102A | 4.26 |
| ENSMUSG00000060962 | Dmkn | dermokine | 4.26 |
| ENSMUSG00000098132 | Rassf10 | Ras association (RalGDS/AF-6) domain family (N-terminal) member 10 | 4.25 |
| ENSMUSG00000009628 | Tex15 | testis expressed gene 15 | 4.25 |
| ENSMUSG00000080316 | Spaca6 | sperm acrosome associated 6 | 4.25 |
| ENSMUSG00000058070 | Eml1 | echinoderm microtubule associated protein like 1 | 4.22 |
| ENSMUSG00000024940 | Ltbp3 | latent transforming growth factor beta binding protein 3 | 4.20 |
| ENSMUSG00000020926 | Adam11 | a disintegrin and metallopeptidase domain 11 | 4.19 |
| ENSMUSG00000025656 | Arhgef9 | CDC42 guanine nucleotide exchange factor (GEF) 9 | 4.18 |
| ENSMUSG00000074796 | Slc4a11 | solute carrier family 4, sodium bicarbonate transporter-like, member 11 | 4.17 |
| ENSMUSG00000039601 | Rcan2 | regulator of calcineurin 2 | 4.16 |
| ENSMUSG00000053297 | AI854703 | expressed sequence AI854703 | 4.16 |
| ENSMUSG00000027257 | Pacsin3 | protein kinase C and casein kinase substrate in neurons 3 | 4.14 |
| ENSMUSG00000020900 | Myh10 | myosin, heavy polypeptide 10, non-muscle | 4.13 |
| ENSMUSG00000036944 | Tmem71 | transmembrane protein 71 | 4.13 |
| ENSMUSG00000075304 | Sp5 | trans-acting transcription factor 5 | 4.12 |
| ENSMUSG00000050212 | Eva1b | eva-1 homolog B (C. elegans) | 4.12 |
| ENSMUSG00000022871 | Fetub | fetuin beta | 4.12 |
| ENSMUSG00000104612 | Gm42449 | predicted gene 42449 | 4.07 |
| ENSMUSG00000004791 | Pgf | placental growth factor | 4.07 |
| ENSMUSG00000025140 | Pycr1 | pyrroline-5-carboxylate reductase 1 | 4.06 |
| ENSMUSG00000078922 | Tgtp1 | T cell specific GTPase 1 | 4.06 |
| ENSMUSG00000013076 | Amotl1 | angiomotin-like 1 | 4.05 |
| ENSMUSG00000026950 | Neb | nebulin | 4.05 |
| ENSMUSG00000031502 | Col4a1 | collagen, type IV, alpha 1 | 4.05 |
| ENSMUSG00000025855 | Prkar1b | protein kinase, cAMP dependent regulatory, type I beta | 4.05 |
| ENSMUSG00000044548 | Dact1 | dishevelled-binding antagonist of beta-catenin 1 | 4.04 |
| ENSMUSG00000022996 | Wnt10b | wingless-type MMTV integration site family, member 10B | 4.03 |
| ENSMUSG00000002980 | Bcam | basal cell adhesion molecule | 4.02 |
| ENSMUSG00000039153 | Runx2 | runt related transcription factor 2 | 4.01 |
| ENSMUSG00000032698 | Lmo2 | LIM domain only 2 | 4.01 |
| ENSMUSG00000041608 | Entpd3 | ectonucleoside triphosphate diphosphohydrolase 3 | 4.00 |
| ENSMUSG00000009687 | Fxyd5 | FXYD domain-containing ion transport regulator 5 | 3.99 |
| ENSMUSG00000031760 | Mt3 | metallothionein 3 | 3.99 |
| ENSMUSG00000025902 | Sox17 | SRY (sex determining region Y)-box 17 | 3.98 |
| ENSMUSG00000021256 | Vash1 | vasohibin 1 | 3.98 |
| ENSMUSG00000032452 | Clstn2 | calsyntenin 2 | 3.97 |
| ENSMUSG00000055980 | Irs1 | insulin receptor substrate 1 | 3.97 |
| ENSMUSG00000044022 | Pcdhb21 | protocadherin beta 21 | 3.97 |

| | | | |
|---|---|---|---|
| ENSMUSG00000062661 | Ncs1 | neuronal calcium sensor 1 | 3.96 |
| ENSMUSG00000000320 | Alox12 | arachidonate 12-lipoxygenase | 3.96 |
| ENSMUSG00000087179 | Gm14230 | predicted gene 14230 | 3.96 |
| ENSMUSG00000079363 | Gbp4 | guanylate binding protein 4 | 3.95 |
| ENSMUSG00000055413 | H2-Q5 | histocompatibility 2, Q region locus 5 | 3.95 |
| ENSMUSG00000043068 | Fam89a | family with sequence similarity 89, member A | 3.93 |
| ENSMUSG00000103749 | Pcdhgb5 | protocadherin gamma subfamily B, 5 | 3.92 |
| ENSMUSG00000037095 | Lrg1 | leucine-rich alpha-2-glycoprotein 1 | 3.90 |
| ENSMUSG00000029287 | Tgfbr3 | transforming growth factor, beta receptor III | 3.90 |
| ENSMUSG00000049556 | Lingo1 | leucine rich repeat and Ig domain containing 1 | 3.90 |
| ENSMUSG00000019817 | Plagl1 | pleiomorphic adenoma gene-like 1 | 3.89 |
| ENSMUSG00000113211 | 4921525O09Rik | RIKEN cDNA 4921525O09 gene | 3.89 |
| ENSMUSG00000040187 | Arntl2 | aryl hydrocarbon receptor nuclear translocator-like 2 | 3.87 |
| ENSMUSG00000105096 | Gbp10 | guanylate-binding protein 10 | 3.87 |
| ENSMUSG00000092586 | Ly6g6c | lymphocyte antigen 6 complex, locus G6C | 3.86 |
| ENSMUSG00000035640 | Cbarp | calcium channel, voltage-dependent, beta subunit associated regulatory protein | 3.85 |
| ENSMUSG00000090399 | Gm38399 | predicted gene, 38399 | 3.85 |
| ENSMUSG00000029298 | Gbp9 | guanylate-binding protein 9 | 3.85 |
| ENSMUSG00000021493 | Pdlim7 | PDZ and LIM domain 7 | 3.84 |
| ENSMUSG00000000782 | Tcf7 | transcription factor 7, T cell specific | 3.83 |
| ENSMUSG00000074766 | Ism1 | isthmin 1, angiogenesis inhibitor | 3.83 |
| ENSMUSG00000041189 | Chrnb1 | cholinergic receptor, nicotinic, beta polypeptide 1 (muscle) | 3.81 |
| ENSMUSG00000038578 | Susd1 | sushi domain containing 1 | 3.81 |
| ENSMUSG00000046058 | Eid2 | EP300 interacting inhibitor of differentiation 2 | 3.81 |
| ENSMUSG00000074916 | Chst14 | carbohydrate (N-acetylgalactosamine 4-0) sulfotransferase 14 | 3.81 |
| ENSMUSG00000023828 | Slc22a3 | solute carrier family 22 (organic cation transporter), member 3 | 3.80 |
| ENSMUSG00000023009 | Nckap5l | NCK-associated protein 5-like | 3.80 |
| ENSMUSG00000107796 | Gm44068 | predicted gene, 44068 | 3.80 |
| ENSMUSG00000030223 | Ptpro | protein tyrosine phosphatase, receptor type, O | 3.79 |
| ENSMUSG00000047363 | Cstad | CSA-conditional, T cell activation-dependent protein | 3.79 |
| ENSMUSG00000061517 | Sox21 | SRY (sex determining region Y)-box 21 | 3.78 |
| ENSMUSG00000045288 | Ush1g | USH1 protein network component sans | 3.77 |
| ENSMUSG00000115602 | Gm7908 | predicted gene 7908 | 3.77 |
| ENSMUSG00000020646 | Mboat2 | membrane bound O-acyltransferase domain containing 2 | 3.76 |
| ENSMUSG00000047910 | Pcdhb16 | protocadherin beta 16 | 3.76 |
| ENSMUSG00000034156 | Tspoap1 | TSPO associated protein 1 | 3.76 |
| ENSMUSG00000041324 | Inhba | inhibin beta-A | 3.76 |
| ENSMUSG00000112084 | Gm10773 | predicted gene 10773 | 3.76 |
| ENSMUSG00000055945 | Prr18 | proline rich 18 | 3.75 |
| ENSMUSG00000036062 | Phf24 | PHD finger protein 24 | 3.75 |
| ENSMUSG00000015957 | Wnt11 | wingless-type MMTV integration site family, member 11 | 3.75 |
| ENSMUSG00000048058 | Ldlrad3 | low density lipoprotein receptor class A domain containing 3 | 3.74 |
| ENSMUSG00000055805 | Fmnl1 | formin-like 1 | 3.74 |
| ENSMUSG00000021306 | Gpr137b | G protein-coupled receptor 137B | 3.74 |
| ENSMUSG00000029832 | Nfe2l3 | nuclear factor, erythroid derived 2, like 3 | 3.72 |
| ENSMUSG00000001521 | Tulp3 | tubby-like protein 3 | 3.71 |
| ENSMUSG00000054793 | Cadm4 | cell adhesion molecule 4 | 3.70 |
| ENSMUSG00000103189 | Gm37092 | predicted gene, 37092 | 3.70 |
| ENSMUSG00000030352 | Tspan9 | tetraspanin 9 | 3.69 |
| ENSMUSG00000059668 | Krt4 | keratin 4 | 3.68 |
| ENSMUSG00000029361 | Nos1 | nitric oxide synthase 1, neuronal | 3.68 |
| ENSMUSG00000046818 | Ddit4l | DNA-damage-inducible transcript 4-like | 3.68 |
| ENSMUSG00000049907 | Rasl11b | RAS-like, family 11, member B | 3.67 |
| ENSMUSG00000003541 | Ier3 | immediate early response 3 | 3.65 |
| ENSMUSG00000028199 | Cryz | crystallin, zeta | 3.65 |
| ENSMUSG00000036500 | Akp3 | alkaline phosphatase 3, intestine, not Mn requiring | 3.64 |
| ENSMUSG00000046768 | Rhoj | ras homolog family member J | 3.64 |
| ENSMUSG00000034612 | Chst11 | carbohydrate sulfotransferase 11 | 3.63 |
| ENSMUSG00000048644 | Ctxn1 | cortexin 1 | 3.63 |
| ENSMUSG00000078921 | Tgtp2 | T cell specific GTPase 2 | 3.62 |
| ENSMUSG00000085111 | Ascl4 | achaete-scute family bHLH transcription factor 4 | 3.62 |
| ENSMUSG00000043924 | Ncmap | noncompact myelin associated protein | 3.62 |
| ENSMUSG00000013419 | Zfp651 | zinc finger protein 651 | 3.62 |
| ENSMUSG00000111293 | Gm34006 | predicted gene, 34006 | 3.62 |
| ENSMUSG00000008932 | Slc1a7 | solute carrier family 1 (glutamate transporter), member 7 | 3.62 |
| ENSMUSG00000045136 | Tubb2b | tubulin, beta 2B class IIB | 3.61 |
| ENSMUSG00000090394 | 4930523C07Rik | RIKEN cDNA 4930523C07 gene | 3.61 |
| ENSMUSG00000054720 | Lrrc8c | leucine rich repeat containing 8 family, member C | 3.61 |
| ENSMUSG00000020651 | Slc26a4 | solute carrier family 26, member 4 | 3.60 |
| ENSMUSG00000026981 | Il1rn | interleukin 1 receptor antagonist | 3.58 |
| ENSMUSG00000022816 | Fstl1 | follistatin-like 1 | 3.56 |
| ENSMUSG00000019990 | Pde7b | phosphodiesterase 7B | 3.56 |
| ENSMUSG00000108897 | Gm44861 | predicted gene 44861 | 3.56 |
| ENSMUSG00000027238 | Frmd5 | FERM domain containing 5 | 3.55 |
| ENSMUSG00000037071 | Scd1 | stearoyl-Coenzyme A desaturase 1 | 3.54 |
| ENSMUSG00000037379 | Spon2 | spondin 2, extracellular matrix protein | 3.54 |

| | | | |
|---|---|---|---|
| ENSMUSG00000001473 | Tubb6 | tubulin, beta 6 class V | 3.53 |
| ENSMUSG00000001288 | Rarg | retinoic acid receptor, gamma | 3.53 |
| ENSMUSG00000029674 | Limk1 | LIM-domain containing, protein kinase | 3.53 |
| ENSMUSG00000008734 | Gprc5b | G protein-coupled receptor, family C, group 5, member B | 3.53 |
| ENSMUSG00000035246 | Pcyt1b | phosphate cytidylyltransferase 1, choline, beta isoform | 3.52 |
| ENSMUSG00000048776 | Pthlh | parathyroid hormone-like peptide | 3.52 |
| ENSMUSG00000024401 | Tnf | tumor necrosis factor | 3.51 |
| ENSMUSG00000055407 | Map6 | microtubule-associated protein 6 | 3.50 |
| ENSMUSG00000022893 | Adamts1 | a disintegrin-like and metallopeptidase (reprolysin type) with thrombospondin type 1 motif, 1 | 3.50 |
| ENSMUSG00000006411 | Nectin4 | nectin cell adhesion molecule 4 | 3.48 |
| ENSMUSG00000057193 | Slc44a2 | solute carrier family 44, member 2 | 3.47 |
| ENSMUSG00000063455 | D630045J12Rik | RIKEN cDNA D630045J12 gene | 3.47 |
| ENSMUSG00000025582 | Nptx1 | neuronal pentraxin 1 | 3.47 |
| ENSMUSG00000028600 | Podn | podocan | 3.46 |
| ENSMUSG00000016526 | Dyrk3 | dual-specificity tyrosine-(Y)-phosphorylation regulated kinase 3 | 3.46 |
| ENSMUSG00000029761 | Cald1 | caldesmon 1 | 3.45 |
| ENSMUSG00000020325 | Fstl3 | follistatin-like 3 | 3.45 |
| ENSMUSG00000014592 | Camta1 | calmodulin binding transcription activator 1 | 3.43 |
| ENSMUSG00000019235 | Rps6kl1 | ribosomal protein S6 kinase-like 1 | 3.43 |
| ENSMUSG00000081683 | Fzd10 | frizzled class receptor 10 | 3.43 |
| ENSMUSG00000055653 | Gpc3 | glypican 3 | 3.43 |
| ENSMUSG00000028108 | Ecm1 | extracellular matrix protein 1 | 3.42 |
| ENSMUSG00000039004 | Bmp6 | bone morphogenetic protein 6 | 3.41 |
| ENSMUSG00000074384 | AI429214 | expressed sequence AI429214 | 3.41 |
| ENSMUSG00000030510 | Cers3 | ceramide synthase 3 | 3.41 |
| ENSMUSG00000032387 | Rbpms2 | RNA binding protein with multiple splicing 2 | 3.41 |
| ENSMUSG00000023905 | Tnfrsf12a | tumor necrosis factor receptor superfamily, member 12a | 3.40 |
| ENSMUSG00000059991 | Nptx2 | neuronal pentraxin 2 | 3.40 |
| ENSMUSG00000025608 | Podxl | podocalyxin-like | 3.39 |
| ENSMUSG00000058145 | Adamts17 | a disintegrin-like and metallopeptidase (reprolysin type) with thrombospondin type 1 motif, 17 | 3.38 |
| ENSMUSG00000030409 | Dmpk | dystrophia myotonica-protein kinase | 3.38 |
| ENSMUSG00000058908 | Pla2g2a | phospholipase A2, group IIA (platelets, synovial fluid) | 3.37 |
| ENSMUSG00000036158 | Prickle1 | prickle planar cell polarity protein 1 | 3.37 |
| ENSMUSG00000024349 | Tmem173 | transmembrane protein 173 | 3.36 |
| ENSMUSG00000043439 | Epop | elongin BC and polycomb repressive complex 2 associated protein | 3.36 |
| ENSMUSG00000037681 | Esyt3 | extended synaptotagmin-like protein 3 | 3.35 |
| ENSMUSG00000053646 | Plxnb1 | plexin B1 | 3.34 |
| ENSMUSG00000052911 | Lamb2 | laminin, beta 2 | 3.34 |
| ENSMUSG00000035954 | Dock4 | dedicator of cytokinesis 4 | 3.34 |
| ENSMUSG00000071723 | Gspt2 | G1 to S phase transition 2 | 3.34 |
| ENSMUSG00000034595 | Ppp1r18 | protein phosphatase 1, regulatory subunit 18 | 3.34 |
| ENSMUSG00000057895 | Zfp105 | zinc finger protein 105 | 3.34 |
| ENSMUSG00000006403 | Adamts4 | a disintegrin-like and metallopeptidase (reprolysin type) with thrombospondin type 1 motif, 4 | 3.34 |
| ENSMUSG00000032068 | Plet1 | placenta expressed transcript 1 | 3.32 |
| ENSMUSG00000026556 | Vangl2 | VANGL planar cell polarity 2 | 3.31 |
| ENSMUSG00000070304 | Scn2b | sodium channel, voltage-gated, type II, beta | 3.31 |
| ENSMUSG00000030600 | Lrfn1 | leucine rich repeat and fibronectin type III domain containing 1 | 3.31 |
| ENSMUSG00000019853 | Hebp2 | heme binding protein 2 | 3.30 |
| ENSMUSG00000044350 | Lacc1 | laccase domain containing 1 | 3.30 |
| ENSMUSG00000037405 | Icam1 | intercellular adhesion molecule 1 | 3.30 |
| ENSMUSG00000014303 | Glis2 | GLIS family zinc finger 2 | 3.30 |
| ENSMUSG00000032940 | Rbm11 | RNA binding motif protein 11 | 3.28 |
| ENSMUSG00000045545 | Krt14 | keratin 14 | 3.28 |
| ENSMUSG00000001435 | Col18a1 | collagen, type XVIII, alpha 1 | 3.27 |
| ENSMUSG00000043391 | 2510009E07Rik | RIKEN cDNA 2510009E07 gene | 3.26 |
| ENSMUSG00000030259 | Rassf8 | Ras association (RalGDS/AF-6) domain family (N-terminal) member 8 | 3.25 |
| ENSMUSG00000046550 | Spin2c | spindlin family, member 2C | 3.25 |
| ENSMUSG00000017493 | Igfbp4 | insulin-like growth factor binding protein 4 | 3.24 |
| ENSMUSG00000035357 | Pdzrn3 | PDZ domain containing RING finger 3 | 3.24 |
| ENSMUSG00000073164 | 2410018L13Rik | RIKEN cDNA 2410018L13 gene | 3.24 |
| ENSMUSG00000020044 | Timp3 | tissue inhibitor of metalloproteinase 3 | 3.23 |
| ENSMUSG00000062488 | Ifit3b | interferon-induced protein with tetratricopeptide repeats 3B | 3.23 |
| ENSMUSG00000022780 | Meltf | melanotransferrin | 3.22 |
| ENSMUSG00000063919 | Srrm4 | serine/arginine repetitive matrix 4 | 3.22 |
| ENSMUSG00000090386 | Mir99ahg | Mir99a and Mirlet7c-1 host gene (non-protein coding) | 3.22 |
| ENSMUSG00000087589 | D430040D24Rik | RIKEN cDNA D430040D24 gene | 3.22 |
| ENSMUSG00000062175 | Tgif2 | TGFB-induced factor homeobox 2 | 3.20 |
| ENSMUSG00000002900 | Lamb1 | laminin B1 | 3.18 |
| ENSMUSG00000040093 | Bmf | BCL2 modifying factor | 3.18 |
| ENSMUSG00000055745 | Rtl6 | retrotransposon Gag like 6 | 3.18 |
| ENSMUSG00000022178 | Ajuba | ajuba LIM protein | 3.17 |
| ENSMUSG00000024109 | Nrxn1 | neurexin I | 3.16 |
| ENSMUSG00000019831 | Wasf1 | WAS protein family, member 1 | 3.15 |
| ENSMUSG00000056947 | Mab21l1 | mab-21-like 1 (C. elegans) | 3.15 |
| ENSMUSG00000041592 | Sdk2 | sidekick cell adhesion molecule 2 | 3.15 |
| ENSMUSG00000030605 | Mfge8 | milk fat globule-EGF factor 8 protein | 3.14 |

| | | | |
|---|---|---|---|
| ENSMUSG00000051343 | Rab11fip5 | RAB11 family interacting protein 5 (class I) | 3.14 |
| ENSMUSG00000027978 | Prss12 | protease, serine 12 neurotrypsin (motopsin) | 3.14 |
| ENSMUSG00000015709 | Arnt2 | aryl hydrocarbon receptor nuclear translocator 2 | 3.13 |
| ENSMUSG00000000202 | Btbd17 | BTB (POZ) domain containing 17 | 3.13 |
| ENSMUSG00000033436 | Armcx2 | armadillo repeat containing, X-linked 2 | 3.13 |
| ENSMUSG00000046402 | Rbp1 | retinol binding protein 1, cellular | 3.12 |
| ENSMUSG00000073409 | H2-Q6 | histocompatibility 2, Q region locus 6 | 3.12 |
| ENSMUSG00000031441 | Atp11a | ATPase, class VI, type 11A | 3.11 |
| ENSMUSG00000037736 | Limch1 | LIM and calponin homology domains 1 | 3.11 |
| ENSMUSG00000046546 | Fam43a | family with sequence similarity 43, member A | 3.11 |
| ENSMUSG00000004709 | Cd244a | CD244 molecule A | 3.11 |
| ENSMUSG00000060181 | Slc35e3 | solute carrier family 35, member E3 | 3.11 |
| ENSMUSG00000023972 | Ptk7 | PTK7 protein tyrosine kinase 7 | 3.10 |
| ENSMUSG00000027375 | Mal | myelin and lymphocyte protein, T cell differentiation protein | 3.10 |
| ENSMUSG00000044043 | Pcdhb14 | protocadherin beta 14 | 3.10 |
| ENSMUSG00000037053 | Azgp1 | alpha-2-glycoprotein 1, zinc | 3.10 |
| ENSMUSG00000037664 | Cdkn1c | cyclin-dependent kinase inhibitor 1C (P57) | 3.09 |
| ENSMUSG00000042523 | Dnal1 | dynein, axonemal, light chain 1 | 3.09 |
| ENSMUSG00000038651 | Sycp2l | synaptonemal complex protein 2-like | 3.09 |
| ENSMUSG00000048347 | Pcdhb18 | protocadherin beta 18 | 3.09 |
| ENSMUSG00000089762 | Ier5l | immediate early response 5-like | 3.08 |
| ENSMUSG00000050295 | Foxc1 | forkhead box C1 | 3.08 |
| ENSMUSG00000026072 | Il1r1 | interleukin 1 receptor, type I | 3.08 |
| ENSMUSG00000036036 | Zfp57 | zinc finger protein 57 | 3.08 |
| ENSMUSG00000041390 | Mdfic | MyoD family inhibitor domain containing | 3.07 |
| ENSMUSG00000038738 | Shank1 | SH3 and multiple ankyrin repeat domains 1 | 3.07 |
| ENSMUSG00000062078 | Qk | quaking | 3.07 |
| ENSMUSG00000027015 | Cybrd1 | cytochrome b reductase 1 | 3.05 |
| ENSMUSG00000029101 | Rgs12 | regulator of G-protein signaling 12 | 3.03 |
| ENSMUSG00000072964 | Bhlhb9 | basic helix-loop-helix domain containing, class B9 | 3.03 |
| ENSMUSG00000020728 | Cep112 | centrosomal protein 112 | 3.02 |
| ENSMUSG00000046387 | Pcdhb17 | protocadherin beta 17 | 3.02 |
| ENSMUSG00000104379 | Gm37509 | predicted gene, 37509 | 3.02 |
| ENSMUSG00000013236 | Ptprs | protein tyrosine phosphatase, receptor type, S | 3.01 |
| ENSMUSG00000050447 | Lypd6 | LY6/PLAUR domain containing 6 | 3.01 |
| ENSMUSG00000053477 | Tcf4 | transcription factor 4 | 3.00 |
| ENSMUSG00000020674 | Pxdn | peroxidasin | 3.00 |
| ENSMUSG00000038980 | Rbbp8nl | RBBP8 N-terminal like | 3.00 |
| ENSMUSG00000048285 | Frmd6 | FERM domain containing 6 | 3.00 |
| ENSMUSG00000022790 | Igsf11 | immunoglobulin superfamily, member 11 | 3.00 |
| ENSMUSG00000038463 | Olfml2b | olfactomedin-like 2B | 3.00 |
| ENSMUSG00000022587 | Ly6e | lymphocyte antigen 6 complex, locus E | 2.99 |
| ENSMUSG00000063623 | C230062I16Rik | RIKEN cDNA C230062I16 gene | 2.99 |
| ENSMUSG00000064263 | Platr26 | pluripotency associated transcript 26 | 2.98 |
| ENSMUSG00000032554 | Trf | transferrin | 2.98 |
| ENSMUSG00000032232 | Cgnl1 | cingulin-like 1 | 2.96 |
| ENSMUSG00000074480 | Mex3a | mex3 RNA binding family member A | 2.96 |
| ENSMUSG00000033149 | Phldb2 | pleckstrin homology like domain, family B, member 2 | 2.96 |
| ENSMUSG00000063851 | Rnf183 | ring finger protein 183 | 2.96 |
| ENSMUSG00000021477 | Ctsl | cathepsin L | 2.95 |
| ENSMUSG00000091243 | Vgll3 | vestigial like family member 3 | 2.95 |
| ENSMUSG00000080937 | Gm13510 | predicted gene 13510 | 2.95 |
| ENSMUSG00000007039 | Ddah2 | dimethylarginine dimethylaminohydrolase 2 | 2.94 |
| ENSMUSG00000053080 | 2700081O15Rik | RIKEN cDNA 2700081O15 gene | 2.93 |
| ENSMUSG00000041570 | Camsap2 | calmodulin regulated spectrin-associated protein family, member 2 | 2.92 |
| ENSMUSG00000107076 | Gm43480 | predicted gene 43480 | 2.92 |
| ENSMUSG00000028698 | Pik3r3 | phosphoinositide-3-kinase regulatory subunit 3 | 2.91 |
| ENSMUSG00000108833 | Gm17909 | predicted gene, 17909 | 2.91 |
| ENSMUSG00000031849 | Comp | cartilage oligomeric matrix protein | 2.91 |
| ENSMUSG00000066705 | Fxyd6 | FXYD domain-containing ion transport regulator 6 | 2.91 |
| ENSMUSG00000016382 | Pls3 | plastin 3 (T-isoform) | 2.90 |
| ENSMUSG00000040690 | Col16a1 | collagen, type XVI, alpha 1 | 2.90 |
| ENSMUSG00000019846 | Lama4 | laminin, alpha 4 | 2.89 |
| ENSMUSG00000026475 | Rgs16 | regulator of G-protein signaling 16 | 2.88 |
| ENSMUSG00000017314 | Mpp2 | membrane protein, palmitoylated 2 (MAGUK p55 subfamily member 2) | 2.88 |
| ENSMUSG00000033276 | Stk36 | serine/threonine kinase 36 | 2.88 |
| ENSMUSG00000007207 | Stx1a | syntaxin 1A (brain) | 2.86 |
| ENSMUSG00000026970 | Rbms1 | RNA binding motif, single stranded interacting protein 1 | 2.86 |
| ENSMUSG00000079669 | Gm17396 | predicted gene, 17396 | 2.86 |
| ENSMUSG00000026121 | Sema4c | sema domain, immunoglobulin domain (Ig), transmembrane domain (TM) and short cytoplasmic do | 2.84 |
| ENSMUSG00000039693 | Msantd3 | Myb/SANT-like DNA-binding domain containing 3 | 2.84 |
| ENSMUSG00000089715 | Cbx6 | chromobox 6 | 2.83 |
| ENSMUSG00000056214 | Pard6g | par-6 family cell polarity regulator gamma | 2.83 |
| ENSMUSG00000055633 | Zfp580 | zinc finger protein 580 | 2.83 |
| ENSMUSG00000085440 | Sorbs2os | sorbin and SH3 domain containing 2, opposite strand | 2.83 |
| ENSMUSG00000036123 | Slc9a3 | solute carrier family 9 (sodium/hydrogen exchanger), member 3 | 2.82 |

| | | | |
|---|---|---|---|
| ENSMUSG00000010021 | Kif19a | kinesin family member 19A | 2.82 |
| ENSMUSG00000026509 | Capn2 | calpain 2 | 2.81 |
| ENSMUSG00000044250 | Pced1b | PC-esterase domain containing 1B | 2.81 |
| ENSMUSG00000022864 | D16Ertd472e | DNA segment, Chr 16, ERATO Doi 472, expressed | 2.81 |
| ENSMUSG00000026185 | Igfbp5 | insulin-like growth factor binding protein 5 | 2.81 |
| ENSMUSG00000028780 | Sema3c | sema domain, immunoglobulin domain (Ig), short basic domain, secreted, (semaphorin) 3C | 2.80 |
| ENSMUSG00000036053 | Fmnl2 | formin-like 2 | 2.80 |
| ENSMUSG00000028347 | Tmeff1 | transmembrane protein with EGF-like and two follistatin-like domains 1 | 2.80 |
| ENSMUSG00000017667 | Zfp334 | zinc finger protein 334 | 2.80 |
| ENSMUSG00000097857 | Gm26603 | predicted gene, 26603 | 2.80 |
| ENSMUSG00000052921 | Arhgef15 | Rho guanine nucleotide exchange factor (GEF) 15 | 2.80 |
| ENSMUSG00000041961 | Znrf3 | zinc and ring finger 3 | 2.79 |
| ENSMUSG00000031586 | Rbpms | RNA binding protein gene with multiple splicing | 2.79 |
| ENSMUSG00000023912 | Slc25a27 | solute carrier family 25, member 27 | 2.79 |
| ENSMUSG00000030630 | Fah | fumarylacetoacetate hydrolase | 2.79 |
| ENSMUSG00000040714 | Klc3 | kinesin light chain 3 | 2.78 |
| ENSMUSG00000084087 | Gm13650 | predicted gene 13650 | 2.78 |
| ENSMUSG00000013275 | Slc41a1 | solute carrier family 41, member 1 | 2.77 |
| ENSMUSG00000043639 | Rbm20 | RNA binding motif protein 20 | 2.77 |
| ENSMUSG00000009614 | Sardh | sarcosine dehydrogenase | 2.77 |
| ENSMUSG00000034774 | Dsg1c | desmoglein 1 gamma | 2.77 |
| ENSMUSG00000026365 | Cfh | complement component factor h | 2.76 |
| ENSMUSG00000026494 | Kif26b | kinesin family member 26B | 2.75 |
| ENSMUSG00000035901 | Dennd5a | DENN/MADD domain containing 5A | 2.75 |
| ENSMUSG00000051790 | Nlgn2 | neuroligin 2 | 2.75 |
| ENSMUSG00000030513 | Pcsk6 | proprotein convertase subtilisin/kexin type 6 | 2.75 |
| ENSMUSG00000024901 | Peli3 | pellino 3 | 2.75 |
| ENSMUSG00000003665 | Has1 | hyaluronan synthase 1 | 2.75 |
| ENSMUSG00000046449 | Nexmif | neurite extension and migration factor | 2.74 |
| ENSMUSG00000028357 | Kif12 | kinesin family member 12 | 2.73 |
| ENSMUSG00000049871 | Nlrc3 | NLR family, CARD domain containing 3 | 2.73 |
| ENSMUSG00000027381 | Bcl2l11 | BCL2-like 11 (apoptosis facilitator) | 2.72 |
| ENSMUSG00000050910 | Cdr2l | cerebellar degeneration-related protein 2-like | 2.72 |
| ENSMUSG00000073433 | Arhgdig | Rho GDP dissociation inhibitor (GDI) gamma | 2.72 |
| ENSMUSG00000043313 | Pcdhb19 | protocadherin beta 19 | 2.72 |
| ENSMUSG00000020205 | Phlda1 | pleckstrin homology like domain, family A, member 1 | 2.71 |
| ENSMUSG00000037972 | Snn | stannin | 2.71 |
| ENSMUSG00000028358 | Zfp618 | zinc finger protein 618 | 2.70 |
| ENSMUSG00000019843 | Fyn | Fyn proto-oncogene | 2.70 |
| ENSMUSG00000060166 | Zdhhc8 | zinc finger, DHHC domain containing 8 | 2.70 |
| ENSMUSG00000003573 | Homer3 | homer scaffolding protein 3 | 2.70 |
| ENSMUSG00000060314 | Zfp941 | zinc finger protein 941 | 2.69 |
| ENSMUSG00000105681 | Gm43428 | predicted gene 43428 | 2.69 |
| ENSMUSG00000112124 | Gm19056 | predicted gene, 19056 | 2.69 |
| ENSMUSG00000060550 | H2-Q7 | histocompatibility 2, Q region locus 7 | 2.68 |
| ENSMUSG00000056665 | Them6 | thioesterase superfamily member 6 | 2.68 |
| ENSMUSG00000092569 | Gm20544 | predicted gene 20544 | 2.68 |
| ENSMUSG00000020804 | Aanat | arylalkylamine N-acetyltransferase | 2.68 |
| ENSMUSG00000000957 | Mmp14 | matrix metallopeptidase 14 (membrane-inserted) | 2.67 |
| ENSMUSG00000042306 | S100a14 | S100 calcium binding protein A14 | 2.67 |
| ENSMUSG00000097494 | 4933406C10Rik | RIKEN cDNA 4933406C10 gene | 2.67 |
| ENSMUSG00000015243 | Abca1 | ATP-binding cassette, sub-family A (ABC1), member 1 | 2.66 |
| ENSMUSG00000008090 | Fgfrl1 | fibroblast growth factor receptor-like 1 | 2.66 |
| ENSMUSG00000020773 | Trim47 | tripartite motif-containing 47 | 2.66 |
| ENSMUSG00000041889 | Shisa4 | shisa family member 4 | 2.66 |
| ENSMUSG00000026211 | Obsl1 | obscurin-like 1 | 2.65 |
| ENSMUSG00000040990 | Sh3kbp1 | SH3-domain kinase binding protein 1 | 2.64 |
| ENSMUSG00000018008 | Cyth4 | cytohesin 4 | 2.64 |
| ENSMUSG00000035274 | Tpbg | trophoblast glycoprotein | 2.63 |
| ENSMUSG00000024736 | Tmem132a | transmembrane protein 132A | 2.63 |
| ENSMUSG00000038079 | Tmem237 | transmembrane protein 237 | 2.63 |
| ENSMUSG00000030222 | Rerg | RAS-like, estrogen-regulated, growth-inhibitor | 2.63 |
| ENSMUSG00000037936 | Scarb1 | scavenger receptor class B, member 1 | 2.62 |
| ENSMUSG00000049823 | Zbtb12 | zinc finger and BTB domain containing 12 | 2.61 |
| ENSMUSG00000086395 | A630014C17Rik | RIKEN cDNA A630014C17 gene | 2.61 |
| ENSMUSG00000026473 | Glul | glutamate-ammonia ligase (glutamine synthetase) | 2.60 |
| ENSMUSG00000046731 | Kctd11 | potassium channel tetramerisation domain containing 11 | 2.60 |
| ENSMUSG00000033287 | Kctd17 | potassium channel tetramerisation domain containing 17 | 2.60 |
| ENSMUSG00000022421 | Nptxr | neuronal pentraxin receptor | 2.60 |
| ENSMUSG00000002633 | Shh | sonic hedgehog | 2.60 |
| ENSMUSG00000031351 | Zfp185 | zinc finger protein 185 | 2.60 |
| ENSMUSG00000002847 | Pla1a | phospholipase A1 member A | 2.60 |
| ENSMUSG00000037902 | Sirpa | signal-regulatory protein alpha | 2.59 |
| ENSMUSG00000024427 | Spry4 | sprouty RTK signaling antagonist 4 | 2.59 |
| ENSMUSG00000073791 | Efcab7 | EF-hand calcium binding domain 7 | 2.59 |
| ENSMUSG00000031530 | Dusp4 | dual specificity phosphatase 4 | 2.58 |

| | | | |
|---|---|---|---|
| ENSMUSG00000036764 | Dnajc12 | DnaJ heat shock protein family (Hsp40) member C12 | 2.58 |
| ENSMUSG00000106219 | 5830416I19Rik | RIKEN cDNA 5830416I19 gene | 2.58 |
| ENSMUSG00000097993 | Ptprv | protein tyrosine phosphatase, receptor type, V | 2.58 |
| ENSMUSG00000021340 | Gpld1 | glycosylphosphatidylinositol specific phospholipase D1 | 2.57 |
| ENSMUSG00000009378 | Slc16a12 | solute carrier family 16 (monocarboxylic acid transporters), member 12 | 2.57 |
| ENSMUSG00000071042 | Rasgrp3 | RAS, guanyl releasing protein 3 | 2.57 |
| ENSMUSG00000104415 | Gm37069 | predicted gene, 37069 | 2.57 |
| ENSMUSG00000028766 | Alpl | alkaline phosphatase, liver/bone/kidney | 2.57 |
| ENSMUSG00000042156 | Dzip1 | DAZ interacting protein 1 | 2.57 |
| ENSMUSG00000032849 | Abcc4 | ATP-binding cassette, sub-family C (CFTR/MRP), member 4 | 2.56 |
| ENSMUSG00000041930 | Fam222a | family with sequence similarity 222, member A | 2.55 |
| ENSMUSG00000026411 | Tmem9 | transmembrane protein 9 | 2.55 |
| ENSMUSG00000024912 | Fosl1 | fos-like antigen 1 | 2.55 |
| ENSMUSG00000031925 | Maml2 | mastermind like transcriptional coactivator 2 | 2.55 |
| ENSMUSG00000029771 | Irf5 | interferon regulatory factor 5 | 2.55 |
| ENSMUSG00000049281 | Scn3b | sodium channel, voltage-gated, type III, beta | 2.55 |
| ENSMUSG00000029094 | Afap1 | actin filament associated protein 1 | 2.54 |
| ENSMUSG00000035914 | Cd276 | CD276 antigen | 2.54 |
| ENSMUSG00000032013 | Trim29 | tripartite motif-containing 29 | 2.54 |
| ENSMUSG00000033730 | Egr3 | early growth response 3 | 2.53 |
| ENSMUSG00000037341 | Slc9a7 | solute carrier family 9 (sodium/hydrogen exchanger), member 7 | 2.53 |
| ENSMUSG00000016534 | Lamp2 | lysosomal-associated membrane protein 2 | 2.52 |
| ENSMUSG00000048232 | Fbxo10 | F-box protein 10 | 2.52 |
| ENSMUSG00000102460 | Gm38197 | predicted gene, 38197 | 2.52 |
| ENSMUSG00000028152 | Tspan5 | tetraspanin 5 | 2.51 |
| ENSMUSG00000053137 | Mapk11 | mitogen-activated protein kinase 11 | 2.51 |
| ENSMUSG00000026405 | C4bp | complement component 4 binding protein | 2.50 |
| ENSMUSG00000040118 | Cacna2d1 | calcium channel, voltage-dependent, alpha2/delta subunit 1 | 2.50 |
| ENSMUSG00000055485 | Soga1 | suppressor of glucose, autophagy associated 1 | 2.49 |
| ENSMUSG00000031292 | Cdkl5 | cyclin-dependent kinase-like 5 | 2.49 |
| ENSMUSG00000060671 | Atp8b2 | ATPase, class I, type 8B, member 2 | 2.49 |
| ENSMUSG00000002603 | Tgfb1 | transforming growth factor, beta 1 | 2.49 |
| ENSMUSG00000022199 | Slc22a17 | solute carrier family 22 (organic cation transporter), member 17 | 2.49 |
| ENSMUSG00000031239 | Itm2a | integral membrane protein 2A | 2.49 |
| ENSMUSG00000025921 | Rdh10 | retinol dehydrogenase 10 (all-trans) | 2.48 |
| ENSMUSG00000013033 | Adgrl1 | adhesion G protein-coupled receptor L1 | 2.48 |
| ENSMUSG00000052852 | Reep1 | receptor accessory protein 1 | 2.48 |
| ENSMUSG00000025592 | Dach2 | dachshund family transcription factor 2 | 2.48 |
| ENSMUSG00000102278 | Gm37145 | predicted gene, 37145 | 2.48 |
| ENSMUSG00000034872 | Gipc3 | GIPC PDZ domain containing family, member 3 | 2.48 |
| ENSMUSG00000041245 | Wnk3 | WNK lysine deficient protein kinase 3 | 2.48 |
| ENSMUSG00000056758 | Hmga2 | high mobility group AT-hook 2 | 2.47 |
| ENSMUSG00000036902 | Neto2 | neuropilin (NRP) and tolloid (TLL)-like 2 | 2.47 |
| ENSMUSG00000031398 | Plxna3 | plexin A3 | 2.47 |
| ENSMUSG00000028464 | Tpm2 | tropomyosin 2, beta | 2.47 |
| ENSMUSG00000029093 | Sorcs2 | sortilin-related VPS10 domain containing receptor 2 | 2.46 |
| ENSMUSG00000039478 | Micu3 | mitochondrial calcium uptake family, member 3 | 2.46 |
| ENSMUSG00000025479 | Cyp2e1 | cytochrome P450, family 2, subfamily e, polypeptide 1 | 2.46 |
| ENSMUSG00000071724 | Smpd5 | sphingomyelin phosphodiesterase 5 | 2.46 |
| ENSMUSG00000006720 | Zfp184 | zinc finger protein 184 (Kruppel-like) | 2.45 |
| ENSMUSG00000031872 | Bean1 | brain expressed, associated with Nedd4, 1 | 2.45 |
| ENSMUSG00000037716 | Ccdc33 | coiled-coil domain containing 33 | 2.44 |
| ENSMUSG00000044716 | Dok7 | docking protein 7 | 2.43 |
| ENSMUSG00000017723 | Wfdc2 | WAP four-disulfide core domain 2 | 2.43 |
| ENSMUSG00000013921 | Clip3 | CAP-GLY domain containing linker protein 3 | 2.43 |
| ENSMUSG00000027999 | Pla2g12a | phospholipase A2, group XIIA | 2.42 |
| ENSMUSG00000044927 | H1fx | H1 histone family, member X | 2.42 |
| ENSMUSG00000040808 | S100g | S100 calcium binding protein G | 2.42 |
| ENSMUSG00000115275 | 9630050E16Rik | RIKEN cDNA 9630050E16 gene | 2.42 |
| ENSMUSG00000060591 | Ifitm2 | interferon induced transmembrane protein 2 | 2.41 |
| ENSMUSG00000022505 | Emp2 | epithelial membrane protein 2 | 2.41 |
| ENSMUSG00000041540 | Sox5 | SRY (sex determining region Y)-box 5 | 2.41 |
| ENSMUSG00000068245 | Phf11d | PHD finger protein 11D | 2.41 |
| ENSMUSG00000000486 | Sept1 | septin 1 | 2.41 |
| ENSMUSG00000032470 | Mras | muscle and microspikes RAS | 2.40 |
| ENSMUSG00000056486 | Chn1 | chimerin 1 | 2.40 |
| ENSMUSG00000028100 | Nudt17 | nudix (nucleoside diphosphate linked moiety X)-type motif 17 | 2.39 |
| ENSMUSG00000028836 | Slc30a2 | solute carrier family 30 (zinc transporter), member 2 | 2.38 |
| ENSMUSG00000045932 | Ifit2 | interferon-induced protein with tetratricopeptide repeats 2 | 2.38 |
| ENSMUSG00000031919 | Tmed6 | transmembrane p24 trafficking protein 6 | 2.38 |
| ENSMUSG00000020599 | Rgs9 | regulator of G-protein signaling 9 | 2.37 |
| ENSMUSG00000018012 | Rac3 | Rac family small GTPase 3 | 2.37 |
| ENSMUSG00000020660 | Pomc | pro-opiomelanocortin-alpha | 2.37 |
| ENSMUSG00000036667 | Tcaf1 | TRPM8 channel-associated factor 1 | 2.36 |
| ENSMUSG00000018906 | P4ha2 | procollagen-proline, 2-oxoglutarate 4-dioxygenase (proline 4-hydroxylase), alpha II polypeptide | 2.36 |
| ENSMUSG00000029762 | Akr1b8 | aldo-keto reductase family 1, member B8 | 2.36 |

| | | | |
|---|---|---|---|
| ENSMUSG00000034771 | Tle2 | transducin-like enhancer of split 2 | 2.36 |
| ENSMUSG00000049191 | Rtl5 | retrotransposon Gag like 5 | 2.36 |
| ENSMUSG00000037060 | Cavin3 | caveolae associated 3 | 2.36 |
| ENSMUSG00000031207 | Msn | moesin | 2.35 |
| ENSMUSG00000028545 | Bend5 | BEN domain containing 5 | 2.35 |
| ENSMUSG00000022696 | Sidt1 | SID1 transmembrane family, member 1 | 2.34 |
| ENSMUSG00000069255 | Dusp22 | dual specificity phosphatase 22 | 2.34 |
| ENSMUSG00000113326 | Gm47586 | predicted gene, 47586 | 2.34 |
| ENSMUSG00000030022 | Adamts9 | a disintegrin-like and metallopeptidase (reprolysin type) with thrombospondin type 1 motif, 9 | 2.34 |
| ENSMUSG00000023913 | Pla2g7 | phospholipase A2, group VII (platelet-activating factor acetylhydrolase, plasma) | 2.34 |
| ENSMUSG00000045629 | Sh3tc2 | SH3 domain and tetratricopeptide repeats 2 | 2.34 |
| ENSMUSG00000038296 | Galnt18 | polypeptide N-acetylgalactosaminyltransferase 18 | 2.34 |
| ENSMUSG00000031633 | Slc25a4 | solute carrier family 25 (mitochondrial carrier, adenine nucleotide translocator), member 4 | 2.33 |
| ENSMUSG00000048696 | Mex3d | mex3 RNA binding family member D | 2.33 |
| ENSMUSG00000031112 | Stk26 | serine/threonine kinase 26 | 2.33 |
| ENSMUSG00000021756 | Il6st | interleukin 6 signal transducer | 2.32 |
| ENSMUSG00000053141 | Ptprt | protein tyrosine phosphatase, receptor type, T | 2.32 |
| ENSMUSG00000084910 | C630043F03Rik | RIKEN cDNA C630043F03 gene | 2.32 |
| ENSMUSG00000030789 | Itgax | integrin alpha X | 2.32 |
| ENSMUSG00000024558 | Mapk4 | mitogen-activated protein kinase 4 | 2.31 |
| ENSMUSG00000031596 | Slc7a2 | solute carrier family 7 (cationic amino acid transporter, y+ system), member 2 | 2.31 |
| ENSMUSG00000110218 | Gm20219 | predicted gene, 20219 | 2.31 |
| ENSMUSG00000039328 | Rnf122 | ring finger protein 122 | 2.30 |
| ENSMUSG00000032607 | Amt | aminomethyltransferase | 2.30 |
| ENSMUSG00000085421 | 4732490B19Rik | RIKEN cDNA 4732490B19 gene | 2.30 |
| ENSMUSG00000020620 | Abca8b | ATP-binding cassette, sub-family A (ABC1), member 8b | 2.30 |
| ENSMUSG00000097709 | 2810429I04Rik | RIKEN cDNA 2810429I04 gene | 2.30 |
| ENSMUSG00000022433 | Csnk1e | casein kinase 1, epsilon | 2.29 |
| ENSMUSG00000038545 | Cul7 | cullin 7 | 2.29 |
| ENSMUSG00000040711 | Sh3pxd2b | SH3 and PX domains 2B | 2.29 |
| ENSMUSG00000028838 | Extl1 | exostoses (multiple)-like 1 | 2.29 |
| ENSMUSG00000104292 | Gm38042 | predicted gene, 38042 | 2.29 |
| ENSMUSG00000069892 | 9930111J21Rik2 | RIKEN cDNA 9930111J21 gene 2 | 2.29 |
| ENSMUSG00000032289 | Thsd4 | thrombospondin, type I, domain containing 4 | 2.29 |
| ENSMUSG00000039457 | Ppl | periplakin | 2.28 |
| ENSMUSG00000056515 | Rab31 | RAB31, member RAS oncogene family | 2.28 |
| ENSMUSG00000034593 | Myo5a | myosin VA | 2.28 |
| ENSMUSG00000039103 | Nexn | nexilin | 2.28 |
| ENSMUSG00000002059 | Rab34 | RAB34, member RAS oncogene family | 2.28 |
| ENSMUSG00000044906 | 4930503L19Rik | RIKEN cDNA 4930503L19 gene | 2.27 |
| ENSMUSG00000046971 | Pla2g4f | phospholipase A2, group IVF | 2.27 |
| ENSMUSG00000082361 | Btc | betacellulin, epidermal growth factor family member | 2.26 |
| ENSMUSG00000045287 | Rtn4rl1 | reticulon 4 receptor-like 1 | 2.26 |
| ENSMUSG00000038884 | A230050P20Rik | RIKEN cDNA A230050P20 gene | 2.26 |
| ENSMUSG00000076431 | Sox4 | SRY (sex determining region Y)-box 4 | 2.25 |
| ENSMUSG00000006777 | Krt23 | keratin 23 | 2.25 |
| ENSMUSG00000047747 | Rnf150 | ring finger protein 150 | 2.25 |
| ENSMUSG00000058317 | Ube2e2 | ubiquitin-conjugating enzyme E2E 2 | 2.25 |
| ENSMUSG00000085329 | 2810404F17Rik | RIKEN cDNA 2810404F17 gene | 2.25 |
| ENSMUSG00000078762 | Haus5 | HAUS augmin-like complex, subunit 5 | 2.25 |
| ENSMUSG00000113811 | Gm47882 | predicted gene, 47882 | 2.25 |
| ENSMUSG00000026223 | Itm2c | integral membrane protein 2C | 2.24 |
| ENSMUSG00000004032 | Gstm5 | glutathione S-transferase, mu 5 | 2.24 |
| ENSMUSG00000037568 | Vash2 | vasohibin 2 | 2.24 |
| ENSMUSG00000021835 | Bmp4 | bone morphogenetic protein 4 | 2.24 |
| ENSMUSG00000056476 | Med12l | mediator complex subunit 12-like | 2.24 |
| ENSMUSG00000020668 | Kif3c | kinesin family member 3C | 2.24 |
| ENSMUSG00000104150 | Gm44573 | predicted gene 44573 | 2.24 |
| ENSMUSG00000044468 | Tent5c | terminal nucleotidyltransferase 5C | 2.23 |
| ENSMUSG00000038740 | Mvb12b | multivesicular body subunit 12B | 2.23 |
| ENSMUSG00000058672 | Tubb2a | tubulin, beta 2A class IIA | 2.23 |
| ENSMUSG00000025854 | Fam20c | family with sequence similarity 20, member C | 2.23 |
| ENSMUSG00000111313 | Gm18942 | predicted gene, 18942 | 2.23 |
| ENSMUSG00000009418 | Nav1 | neuron navigator 1 | 2.22 |
| ENSMUSG00000033032 | Afap1l1 | actin filament associated protein 1-like 1 | 2.22 |
| ENSMUSG00000001248 | Gramd1a | GRAM domain containing 1A | 2.22 |
| ENSMUSG00000032640 | Chsy1 | chondroitin sulfate synthase 1 | 2.22 |
| ENSMUSG00000028076 | Cd1d1 | CD1d1 antigen | 2.22 |
| ENSMUSG00000020396 | Nefh | neurofilament, heavy polypeptide | 2.22 |
| ENSMUSG00000023031 | Cela1 | chymotrypsin-like elastase family, member 1 | 2.21 |
| ENSMUSG00000055409 | Nell1 | NEL-like 1 | 2.21 |
| ENSMUSG00000021506 | Pitx1 | paired-like homeodomain transcription factor 1 | 2.20 |
| ENSMUSG00000023007 | Prpf40b | pre-mRNA processing factor 40B | 2.20 |
| ENSMUSG00000028654 | Mycl | v-myc avian myelocytomatosis viral oncogene lung carcinoma derived | 2.19 |
| ENSMUSG00000021464 | Ror2 | receptor tyrosine kinase-like orphan receptor 2 | 2.19 |
| ENSMUSG00000038594 | Cep85l | centrosomal protein 85-like | 2.19 |

| | | | |
|---|---|---|---|
| ENSMUSG00000029669 | Tspan12 | tetraspanin 12 | 2.18 |
| ENSMUSG00000097715 | Gpr137b-ps | G protein-coupled receptor 137B, pseudogene | 2.18 |
| ENSMUSG00000046191 | Pcdhb20 | protocadherin beta 20 | 2.18 |
| ENSMUSG00000032514 | Ttc21a | tetratricopeptide repeat domain 21A | 2.18 |
| ENSMUSG00000027346 | Gpcpd1 | glycerophosphocholine phosphodiesterase 1 | 2.17 |
| ENSMUSG00000025017 | Pik3ap1 | phosphoinositide-3-kinase adaptor protein 1 | 2.17 |
| ENSMUSG00000027712 | Anxa5 | annexin A5 | 2.17 |
| ENSMUSG00000026944 | Abca2 | ATP-binding cassette, sub-family A (ABC1), member 2 | 2.16 |
| ENSMUSG00000033233 | Trim45 | tripartite motif-containing 45 | 2.16 |
| ENSMUSG00000050071 | Bex1 | brain expressed X-linked 1 | 2.15 |
| ENSMUSG00000034848 | Ttc21b | tetratricopeptide repeat domain 21B | 2.15 |
| ENSMUSG00000039934 | Gsap | gamma-secretase activating protein | 2.15 |
| ENSMUSG00000037833 | Sh2d4b | SH2 domain containing 4B | 2.15 |
| ENSMUSG00000060985 | Tdrd5 | tudor domain containing 5 | 2.15 |
| ENSMUSG00000105140 | Gm43127 | predicted gene 43127 | 2.15 |
| ENSMUSG00000049577 | Zfpm1 | zinc finger protein, multitype 1 | 2.14 |
| ENSMUSG00000047501 | Cldn4 | claudin 4 | 2.14 |
| ENSMUSG00000041308 | Sntb2 | syntrophin, basic 2 | 2.14 |
| ENSMUSG00000031066 | Usp11 | ubiquitin specific peptidase 11 | 2.14 |
| ENSMUSG00000109079 | Gm44755 | predicted gene 44755 | 2.14 |
| ENSMUSG00000027186 | Elf5 | E74-like factor 5 | 2.14 |
| ENSMUSG00000000142 | Axin2 | axin 2 | 2.13 |
| ENSMUSG00000028613 | Lrp8 | low density lipoprotein receptor-related protein 8, apolipoprotein e receptor | 2.13 |
| ENSMUSG00000055926 | Gm14137 | predicted gene 14137 | 2.13 |
| ENSMUSG00000111905 | Gm48045 | predicted gene, 48045 | 2.13 |
| ENSMUSG00000034177 | Rnf43 | ring finger protein 43 | 2.12 |
| ENSMUSG00000022415 | Syngr1 | synaptogyrin 1 | 2.12 |
| ENSMUSG00000047146 | Tet1 | tet methylcytosine dioxygenase 1 | 2.12 |
| ENSMUSG00000110411 | Gm45457 | predicted gene 45457 | 2.12 |
| ENSMUSG00000062937 | Mtap | methylthioadenosine phosphorylase | 2.11 |
| ENSMUSG00000017390 | Aldoc | aldolase C, fructose-bisphosphate | 2.11 |
| ENSMUSG00000032363 | Adamts7 | a disintegrin-like and metallopeptidase (reprolysin type) with thrombospondin type 1 motif, 7 | 2.11 |
| ENSMUSG00000090100 | Ttbk2 | tau tubulin kinase 2 | 2.10 |
| ENSMUSG00000049670 | Morn4 | MORN repeat containing 4 | 2.10 |
| ENSMUSG00000031907 | Zfp90 | zinc finger protein 90 | 2.10 |
| ENSMUSG00000052212 | Cd177 | CD177 antigen | 2.09 |
| ENSMUSG00000024270 | Slc39a6 | solute carrier family 39 (metal ion transporter), member 6 | 2.09 |
| ENSMUSG00000029312 | Klhl8 | kelch-like 8 | 2.08 |
| ENSMUSG00000027674 | Pex5l | peroxisomal biogenesis factor 5-like | 2.08 |
| ENSMUSG00000046589 | Lrrc8e | leucine rich repeat containing 8 family, member E | 2.08 |
| ENSMUSG00000031996 | Aplp2 | amyloid beta (A4) precursor-like protein 2 | 2.07 |
| ENSMUSG00000073987 | Ggh | gamma-glutamyl hydrolase | 2.07 |
| ENSMUSG00000032184 | Lysmd2 | LysM, putative peptidoglycan-binding, domain containing 2 | 2.07 |
| ENSMUSG00000037946 | Fgd3 | FYVE, RhoGEF and PH domain containing 3 | 2.07 |
| ENSMUSG00000001025 | S100a6 | S100 calcium binding protein A6 (calcyclin) | 2.06 |
| ENSMUSG00000026478 | Lamc1 | laminin, gamma 1 | 2.06 |
| ENSMUSG00000046329 | Slc25a23 | solute carrier family 25 (mitochondrial carrier; phosphate carrier), member 23 | 2.06 |
| ENSMUSG00000076441 | Ass1 | argininosuccinate synthetase 1 | 2.06 |
| ENSMUSG00000041293 | Adgrf1 | adhesion G protein-coupled receptor F1 | 2.06 |
| ENSMUSG00000080775 | Gm6368 | predicted gene 6368 | 2.06 |
| ENSMUSG00000025038 | Efhc2 | EF-hand domain (C-terminal) containing 2 | 2.06 |
| ENSMUSG00000087792 | Gm25968 | predicted gene, 25968 | 2.06 |
| ENSMUSG00000028184 | Adgrl2 | adhesion G protein-coupled receptor L2 | 2.05 |
| ENSMUSG00000024896 | Minpp1 | multiple inositol polyphosphate histidine phosphatase 1 | 2.05 |
| ENSMUSG00000089809 | Rasgef1b | RasGEF domain family, member 1B | 2.05 |
| ENSMUSG00000071855 | Ccdc112 | coiled-coil domain containing 112 | 2.05 |
| ENSMUSG00000109341 | Gm30873 | predicted gene, 30873 | 2.05 |
| ENSMUSG00000042524 | Sun2 | Sad1 and UNC84 domain containing 2 | 2.04 |
| ENSMUSG00000058656 | Samd12 | sterile alpha motif domain containing 12 | 2.04 |
| ENSMUSG00000056832 | Ttc26 | tetratricopeptide repeat domain 26 | 2.04 |
| ENSMUSG00000014158 | Trpv4 | transient receptor potential cation channel, subfamily V, member 4 | 2.04 |
| ENSMUSG00000044229 | Nxpe4 | neurexophilin and PC-esterase domain family, member 4 | 2.04 |
| ENSMUSG00000062184 | Hs6st2 | heparan sulfate 6-O-sulfotransferase 2 | 2.04 |
| ENSMUSG00000060261 | Gtf2i | general transcription factor II I | 2.03 |
| ENSMUSG00000057092 | Fxyd3 | FXYD domain-containing ion transport regulator 3 | 2.03 |
| ENSMUSG00000059895 | Ptp4a3 | protein tyrosine phosphatase 4a3 | 2.03 |
| ENSMUSG00000027698 | Nceh1 | neutral cholesterol ester hydrolase 1 | 2.02 |
| ENSMUSG00000021589 | Rhobtb3 | Rho-related BTB domain containing 3 | 2.02 |
| ENSMUSG00000001249 | Hpn | hepsin | 2.02 |
| ENSMUSG00000028348 | Cavin4 | caveolae associated 4 | 2.02 |
| ENSMUSG00000059741 | Myl3 | myosin, light polypeptide 3 | 2.02 |
| ENSMUSG00000053398 | Phgdh | 3-phosphoglycerate dehydrogenase | 2.01 |
| ENSMUSG00000037287 | Tbcel | tubulin folding cofactor E-like | 2.00 |
| ENSMUSG00000041025 | Iffo2 | intermediate filament family orphan 2 | 2.00 |
| ENSMUSG00000026604 | Ptpn14 | protein tyrosine phosphatase, non-receptor type 14 | 2.00 |
| ENSMUSG00000050359 | Sprr1a | small proline-rich protein 1A | 2.00 |

| | | | |
|---|---|---|---|
| ENSMUSG00000110256 | Gm45412 | predicted gene 45412 | 2.00 |
| ENSMUSG00000027907 | S100a11 | S100 calcium binding protein A11 | 1.99 |
| ENSMUSG00000009739 | Pou6f1 | POU domain, class 6, transcription factor 1 | 1.99 |
| ENSMUSG00000020546 | Stxbp4 | syntaxin binding protein 4 | 1.99 |
| ENSMUSG00000030862 | Cpxm2 | carboxypeptidase X 2 (M14 family) | 1.99 |
| ENSMUSG00000007989 | Fzd3 | frizzled class receptor 3 | 1.98 |
| ENSMUSG00000038156 | Spon1 | spondin 1, (f-spondin) extracellular matrix protein | 1.98 |
| ENSMUSG00000024990 | Rbp4 | retinol binding protein 4, plasma | 1.98 |
| ENSMUSG00000042684 | Npl | N-acetylneuraminate pyruvate lyase | 1.98 |
| ENSMUSG00000051510 | Mafg | v-maf musculoaponeurotic fibrosarcoma oncogene family, protein G (avian) | 1.97 |
| ENSMUSG00000084939 | Gm830 | predicted gene 830 | 1.97 |
| ENSMUSG00000062044 | Lmtk3 | lemur tyrosine kinase 3 | 1.97 |
| ENSMUSG00000113876 | Gm48042 | predicted gene, 48042 | 1.97 |
| ENSMUSG00000116656 | AC131339.2 | novel transcript | 1.97 |
| ENSMUSG00000107388 | Gm42788 | predicted gene 42788 | 1.97 |
| ENSMUSG00000027333 | Smox | spermine oxidase | 1.96 |
| ENSMUSG00000056648 | Hoxb8 | homeobox B8 | 1.96 |
| ENSMUSG00000021611 | Tert | telomerase reverse transcriptase | 1.96 |
| ENSMUSG00000024968 | Rcor2 | REST corepressor 2 | 1.96 |
| ENSMUSG00000038065 | Mturn | maturin, neural progenitor differentiation regulator homolog (Xenopus) | 1.96 |
| ENSMUSG00000051652 | Lrrc3 | leucine rich repeat containing 3 | 1.95 |
| ENSMUSG00000046805 | Mpeg1 | macrophage expressed gene 1 | 1.95 |
| ENSMUSG00000034573 | Ptpn13 | protein tyrosine phosphatase, non-receptor type 13 | 1.95 |
| ENSMUSG00000013846 | St3gal1 | ST3 beta-galactoside alpha-2,3-sialyltransferase 1 | 1.94 |
| ENSMUSG00000069874 | Irgm2 | immunity-related GTPase family M member 2 | 1.94 |
| ENSMUSG00000030284 | Creld1 | cysteine-rich with EGF-like domains 1 | 1.93 |
| ENSMUSG00000032194 | Kank2 | KN motif and ankyrin repeat domains 2 | 1.93 |
| ENSMUSG00000066643 | Wdr35 | WD repeat domain 35 | 1.93 |
| ENSMUSG00000071984 | Fndc1 | fibronectin type III domain containing 1 | 1.93 |
| ENSMUSG00000051359 | Ncald | neurocalcin delta | 1.92 |
| ENSMUSG00000039621 | Prex1 | phosphatidylinositol-3,4,5-trisphosphate-dependent Rac exchange factor 1 | 1.92 |
| ENSMUSG00000032578 | Cish | cytokine inducible SH2-containing protein | 1.92 |
| ENSMUSG00000073591 | Pcdhb22 | protocadherin beta 22 | 1.92 |
| ENSMUSG00000026727 | Rsu1 | Ras suppressor protein 1 | 1.91 |
| ENSMUSG00000024524 | Gnal | guanine nucleotide binding protein, alpha stimulating, olfactory type | 1.91 |
| ENSMUSG00000031617 | Tmem184c | transmembrane protein 184C | 1.91 |
| ENSMUSG00000020389 | Cdkl3 | cyclin-dependent kinase-like 3 | 1.91 |
| ENSMUSG00000099404 | Gm28172 | predicted gene 28172 | 1.91 |
| ENSMUSG00000028581 | Laptm5 | lysosomal-associated protein transmembrane 5 | 1.91 |
| ENSMUSG00000048728 | Zfp454 | zinc finger protein 454 | 1.91 |
| ENSMUSG00000047945 | Marcksl1 | MARCKS-like 1 | 1.90 |
| ENSMUSG00000006369 | Fbln1 | fibulin 1 | 1.90 |
| ENSMUSG00000003363 | Pld3 | phospholipase D family, member 3 | 1.90 |
| ENSMUSG00000025986 | Slc39a10 | solute carrier family 39 (zinc transporter), member 10 | 1.90 |
| ENSMUSG00000041096 | Tspyl2 | TSPY-like 2 | 1.90 |
| ENSMUSG00000008206 | Cers4 | ceramide synthase 4 | 1.90 |
| ENSMUSG00000003812 | Dnase2a | deoxyribonuclease II alpha | 1.90 |
| ENSMUSG00000049804 | Armcx4 | armadillo repeat containing, X-linked 4 | 1.90 |
| ENSMUSG00000111212 | Gm47087 | predicted gene, 47087 | 1.90 |
| ENSMUSG00000044062 | Plekhd1os | pleckstrin homology domain containing, family D (with coiled-coil domains) member 1, opposite stra | 1.90 |
| ENSMUSG00000023088 | Abcc1 | ATP-binding cassette, sub-family C (CFTR/MRP), member 1 | 1.89 |
| ENSMUSG00000035376 | Hacd2 | 3-hydroxyacyl-CoA dehydratase 2 | 1.88 |
| ENSMUSG00000020453 | Patz1 | POZ (BTB) and AT hook containing zinc finger 1 | 1.88 |
| ENSMUSG00000037447 | Arid5a | AT rich interactive domain 5A (MRF1-like) | 1.88 |
| ENSMUSG00000040717 | Il17rd | interleukin 17 receptor D | 1.88 |
| ENSMUSG00000033906 | Zdhhc15 | zinc finger, DHHC domain containing 15 | 1.88 |
| ENSMUSG00000022297 | Fzd6 | frizzled class receptor 6 | 1.88 |
| ENSMUSG00000035638 | Muc20 | mucin 20 | 1.88 |
| ENSMUSG00000109957 | Gm45353 | predicted gene 45353 | 1.88 |
| ENSMUSG00000037254 | Itih2 | inter-alpha trypsin inhibitor, heavy chain 2 | 1.88 |
| ENSMUSG00000027997 | Casp6 | caspase 6 | 1.87 |
| ENSMUSG00000106847 | Peg13 | paternally expressed 13 | 1.87 |
| ENSMUSG00000073565 | Prr16 | proline rich 16 | 1.87 |
| ENSMUSG00000022512 | Cldn1 | claudin 1 | 1.87 |
| ENSMUSG00000086544 | Chn1os3 | chimerin 1, opposite strand 3 | 1.87 |
| ENSMUSG00000048924 | Ccdc125 | coiled-coil domain containing 125 | 1.86 |
| ENSMUSG00000059975 | Zfp74 | zinc finger protein 74 | 1.86 |
| ENSMUSG00000029156 | Sgcb | sarcoglycan, beta (dystrophin-associated glycoprotein) | 1.86 |
| ENSMUSG00000016942 | Tmprss6 | transmembrane serine protease 6 | 1.86 |
| ENSMUSG00000109245 | Gm44860 | predicted gene 44860 | 1.86 |
| ENSMUSG00000056155 | Nanos3 | nanos C2HC-type zinc finger 3 | 1.86 |
| ENSMUSG00000019841 | Rev3l | REV3 like, DNA directed polymerase zeta catalytic subunit | 1.85 |
| ENSMUSG00000071637 | Cebpd | CCAAT/enhancer binding protein (C/EBP), delta | 1.85 |
| ENSMUSG00000016541 | Atxn10 | ataxin 10 | 1.85 |
| ENSMUSG00000042155 | Klhl23 | kelch-like 23 | 1.85 |
| ENSMUSG00000051243 | Islr2 | immunoglobulin superfamily containing leucine-rich repeat 2 | 1.85 |

| | | | |
|---|---|---|---|
| ENSMUSG00000049521 | Cdc42ep1 | CDC42 effector protein (Rho GTPase binding) 1 | 1.84 |
| ENSMUSG00000114524 | Gm48357 | predicted gene, 48357 | 1.84 |
| ENSMUSG00000026383 | Epb41l5 | erythrocyte membrane protein band 4.1 like 5 | 1.83 |
| ENSMUSG00000019256 | Ahr | aryl-hydrocarbon receptor | 1.83 |
| ENSMUSG00000025075 | Habp2 | hyaluronic acid binding protein 2 | 1.83 |
| ENSMUSG00000102758 | Naaladl2 | N-acetylated alpha-linked acidic dipeptidase-like 2 | 1.83 |
| ENSMUSG00000049939 | Lrrc4 | leucine rich repeat containing 4 | 1.83 |
| ENSMUSG00000069208 | Zfp825 | zinc finger protein 825 | 1.83 |
| ENSMUSG00000026458 | Ppfia4 | protein tyrosine phosphatase, receptor type, f polypeptide (PTPRF), interacting protein (liprin), alph | 1.83 |
| ENSMUSG00000070822 | Zscan18 | zinc finger and SCAN domain containing 18 | 1.83 |
| ENSMUSG00000023348 | Trip6 | thyroid hormone receptor interactor 6 | 1.82 |
| ENSMUSG00000054412 | Gm4793 | predicted gene 4793 | 1.82 |
| ENSMUSG00000103818 | Gm38009 | predicted gene, 38009 | 1.82 |
| ENSMUSG00000058799 | Nap1l1 | nucleosome assembly protein 1-like 1 | 1.81 |
| ENSMUSG00000064061 | Dzip3 | DAZ interacting protein 3, zinc finger | 1.81 |
| ENSMUSG00000035829 | Ppp1r26 | protein phosphatase 1, regulatory subunit 26 | 1.81 |
| ENSMUSG00000035390 | Brsk1 | BR serine/threonine kinase 1 | 1.81 |
| ENSMUSG00000031480 | Thsd1 | thrombospondin, type I, domain 1 | 1.81 |
| ENSMUSG00000078716 | Tmem8b | transmembrane protein 8B | 1.81 |
| ENSMUSG00000050821 | Fam131a | family with sequence similarity 131, member A | 1.81 |
| ENSMUSG00000031660 | Brd7 | bromodomain containing 7 | 1.80 |
| ENSMUSG00000070000 | Fcho1 | FCH domain only 1 | 1.80 |
| ENSMUSG00000050737 | Ptges | prostaglandin E synthase | 1.80 |
| ENSMUSG00000026479 | Lamc2 | laminin, gamma 2 | 1.79 |
| ENSMUSG00000030486 | Zfp108 | zinc finger protein 108 | 1.79 |
| ENSMUSG00000030161 | Gabarapl1 | gamma-aminobutyric acid (GABA) A receptor-associated protein-like 1 | 1.78 |
| ENSMUSG00000038205 | Prkab2 | protein kinase, AMP-activated, beta 2 non-catalytic subunit | 1.78 |
| ENSMUSG00000054580 | Pla2r1 | phospholipase A2 receptor 1 | 1.78 |
| ENSMUSG00000083816 | Gm13033 | predicted gene 13033 | 1.78 |
| ENSMUSG00000036867 | Smad6 | SMAD family member 6 | 1.77 |
| ENSMUSG00000032300 | 1700017B05Rik | RIKEN cDNA 1700017B05 gene | 1.76 |
| ENSMUSG00000032440 | Tgfbr2 | transforming growth factor, beta receptor II | 1.76 |
| ENSMUSG00000021262 | Evl | Ena-vasodilator stimulated phosphoprotein | 1.76 |
| ENSMUSG00000039384 | Dusp10 | dual specificity phosphatase 10 | 1.76 |
| ENSMUSG00000020650 | Bcap29 | B cell receptor associated protein 29 | 1.76 |
| ENSMUSG00000040591 | 1110051M20Rik | RIKEN cDNA 1110051M20 gene | 1.76 |
| ENSMUSG00000020253 | Ppm1m | protein phosphatase 1M | 1.76 |
| ENSMUSG00000053931 | Cnn3 | calponin 3, acidic | 1.75 |
| ENSMUSG00000028211 | Trp53inp1 | transformation related protein 53 inducible nuclear protein 1 | 1.75 |
| ENSMUSG00000008999 | Bmp7 | bone morphogenetic protein 7 | 1.75 |
| ENSMUSG00000070305 | Mpzl3 | myelin protein zero-like 3 | 1.75 |
| ENSMUSG00000034854 | Mfsd12 | major facilitator superfamily domain containing 12 | 1.75 |
| ENSMUSG00000020032 | Nuak1 | NUAK family, SNF1-like kinase, 1 | 1.75 |
| ENSMUSG00000038797 | Zscan2 | zinc finger and SCAN domain containing 2 | 1.75 |
| ENSMUSG00000103088 | Pcdhgb6 | protocadherin gamma subfamily B, 6 | 1.74 |
| ENSMUSG00000016024 | Lbp | lipopolysaccharide binding protein | 1.74 |
| ENSMUSG00000025185 | Loxl4 | lysyl oxidase-like 4 | 1.74 |
| ENSMUSG00000026278 | Bok | BCL2-related ovarian killer | 1.73 |
| ENSMUSG00000053716 | Dusp7 | dual specificity phosphatase 7 | 1.73 |
| ENSMUSG00000004665 | Cnn2 | calponin 2 | 1.73 |
| ENSMUSG00000073000 | Gm10451 | predicted gene 10451 | 1.73 |
| ENSMUSG00000020570 | Sypl | synaptophysin-like protein | 1.72 |
| ENSMUSG00000028849 | Map7d1 | MAP7 domain containing 1 | 1.72 |
| ENSMUSG00000030315 | Vgll4 | vestigial like family member 4 | 1.72 |
| ENSMUSG00000045319 | Proser2 | proline and serine rich 2 | 1.72 |
| ENSMUSG00000025265 | Fgd1 | FYVE, RhoGEF and PH domain containing 1 | 1.72 |
| ENSMUSG00000103037 | Pcdhgb1 | protocadherin gamma subfamily B, 1 | 1.72 |
| ENSMUSG00000047344 | Lancl3 | LanC lantibiotic synthetase component C-like 3 (bacterial) | 1.72 |
| ENSMUSG00000027796 | Smad9 | SMAD family member 9 | 1.71 |
| ENSMUSG00000039081 | Zfp503 | zinc finger protein 503 | 1.70 |
| ENSMUSG00000032348 | Gsta4 | glutathione S-transferase, alpha 4 | 1.70 |
| ENSMUSG00000041886 | Macc1 | metastasis associated in colon cancer 1 | 1.70 |
| ENSMUSG00000024013 | Fgd2 | FYVE, RhoGEF and PH domain containing 2 | 1.70 |
| ENSMUSG00000002107 | Celf2 | CUGBP, Elav-like family member 2 | 1.70 |
| ENSMUSG00000096544 | Gm4617 | predicted pseudogene 4617 | 1.70 |
| ENSMUSG00000000058 | Cav2 | caveolin 2 | 1.70 |
| ENSMUSG00000038872 | Zfhx3 | zinc finger homeobox 3 | 1.69 |
| ENSMUSG00000047407 | Tgif1 | TGFB-induced factor homeobox 1 | 1.69 |
| ENSMUSG00000020776 | Fbf1 | Fas (TNFRSF6) binding factor 1 | 1.69 |
| ENSMUSG00000017376 | Nlk | nemo like kinase | 1.69 |
| ENSMUSG00000020848 | Doc2b | double C2, beta | 1.69 |
| ENSMUSG00000035778 | Ggta1 | glycoprotein galactosyltransferase alpha 1, 3 | 1.69 |
| ENSMUSG00000031098 | Syt8 | synaptotagmin VIII | 1.69 |
| ENSMUSG00000020592 | Sdc1 | syndecan 1 | 1.68 |
| ENSMUSG00000009585 | Apobec3 | apolipoprotein B mRNA editing enzyme, catalytic polypeptide 3 | 1.68 |
| ENSMUSG00000038400 | Pmepa1 | prostate transmembrane protein, androgen induced 1 | 1.68 |

| | | | |
|---|---|---|---|
| ENSMUSG00000030759 | Far1 | fatty acyl CoA reductase 1 | 1.68 |
| ENSMUSG00000045045 | Lrfn4 | leucine rich repeat and fibronectin type III domain containing 4 | 1.68 |
| ENSMUSG00000028737 | Aldh4a1 | aldehyde dehydrogenase 4 family, member A1 | 1.68 |
| ENSMUSG00000028378 | Ptgr1 | prostaglandin reductase 1 | 1.67 |
| ENSMUSG00000025161 | Slc16a3 | solute carrier family 16 (monocarboxylic acid transporters), member 3 | 1.67 |
| ENSMUSG00000029246 | Ppat | phosphoribosyl pyrophosphate amidotransferase | 1.67 |
| ENSMUSG00000000555 | Itga5 | integrin alpha 5 (fibronectin receptor alpha) | 1.67 |
| ENSMUSG00000046572 | Zfp518b | zinc finger protein 518B | 1.67 |
| ENSMUSG00000028573 | Fggy | FGGY carbohydrate kinase domain containing | 1.67 |
| ENSMUSG00000038068 | Rnf144b | ring finger protein 144B | 1.67 |
| ENSMUSG00000037892 | Pcdh18 | protocadherin 18 | 1.67 |
| ENSMUSG00000040473 | Cfap69 | cilia and flagella associated protein 69 | 1.67 |
| ENSMUSG00000025534 | Gusb | glucuronidase, beta | 1.66 |
| ENSMUSG00000024070 | Prkd3 | protein kinase D3 | 1.66 |
| ENSMUSG00000004105 | Angptl2 | angiopoietin-like 2 | 1.66 |
| ENSMUSG00000027995 | Tlr2 | toll-like receptor 2 | 1.66 |
| ENSMUSG00000021373 | Cap2 | CAP, adenylate cyclase-associated protein, 2 (yeast) | 1.66 |
| ENSMUSG00000041836 | Ptpre | protein tyrosine phosphatase, receptor type, E | 1.65 |
| ENSMUSG00000053897 | Slc39a8 | solute carrier family 39 (metal ion transporter), member 8 | 1.65 |
| ENSMUSG00000037656 | Slc20a2 | solute carrier family 20, member 2 | 1.65 |
| ENSMUSG00000040466 | Blvrb | biliverdin reductase B (flavin reductase (NADPH)) | 1.65 |
| ENSMUSG00000021792 | Fam213a | family with sequence similarity 213, member A | 1.65 |
| ENSMUSG00000044636 | Csrnp2 | cysteine-serine-rich nuclear protein 2 | 1.65 |
| ENSMUSG00000037499 | Nenf | neuron derived neurotrophic factor | 1.65 |
| ENSMUSG00000020656 | Grhl1 | grainyhead like transcription factor 1 | 1.65 |
| ENSMUSG00000018822 | Sfrp5 | secreted frizzled-related sequence protein 5 | 1.65 |
| ENSMUSG00000020176 | Grb10 | growth factor receptor bound protein 10 | 1.65 |
| ENSMUSG00000091438 | Gm17088 | predicted gene 17088 | 1.65 |
| ENSMUSG00000078853 | Igtp | interferon gamma induced GTPase | 1.64 |
| ENSMUSG00000031365 | Zfp275 | zinc finger protein 275 | 1.64 |
| ENSMUSG00000081480 | Gm13608 | predicted gene 13608 | 1.64 |
| ENSMUSG00000025083 | Afap1l2 | actin filament associated protein 1-like 2 | 1.64 |
| ENSMUSG00000007891 | Ctsd | cathepsin D | 1.63 |
| ENSMUSG00000033039 | Micall1 | microtubule associated monooxygenase, calponin and LIM domain containing -like 1 | 1.63 |
| ENSMUSG00000025268 | Maged2 | melanoma antigen, family D, 2 | 1.63 |
| ENSMUSG00000047264 | Zfp358 | zinc finger protein 358 | 1.63 |
| ENSMUSG00000017724 | Etv4 | ets variant 4 | 1.63 |
| ENSMUSG00000059493 | Nhs | NHS actin remodeling regulator | 1.63 |
| ENSMUSG00000085972 | 1110028F11Rik | RIKEN cDNA 1110028F11 gene | 1.63 |
| ENSMUSG00000037185 | Krt80 | keratin 80 | 1.63 |
| ENSMUSG00000023224 | Serping1 | serine (or cysteine) peptidase inhibitor, clade G, member 1 | 1.63 |
| ENSMUSG00000027860 | Vangl1 | VANGL planar cell polarity 1 | 1.62 |
| ENSMUSG00000053110 | Yap1 | yes-associated protein 1 | 1.61 |
| ENSMUSG00000037679 | Inf2 | inverted formin, FH2 and WH2 domain containing | 1.61 |
| ENSMUSG00000063760 | Rnf217 | ring finger protein 217 | 1.61 |
| ENSMUSG00000032531 | Amotl2 | angiomotin-like 2 | 1.61 |
| ENSMUSG00000025577 | Cbx2 | chromobox 2 | 1.61 |
| ENSMUSG00000032657 | Fam189b | family with sequence similarity 189, member B | 1.61 |
| ENSMUSG00000038022 | Mindy4 | MINDY lysine 48 deubiquitinase 4 | 1.61 |
| ENSMUSG00000025145 | Lrrc45 | leucine rich repeat containing 45 | 1.60 |
| ENSMUSG00000022043 | Trim35 | tripartite motif-containing 35 | 1.60 |
| ENSMUSG00000047749 | Zc3hav1l | zinc finger CCCH-type, antiviral 1-like | 1.60 |
| ENSMUSG00000028214 | Gem | GTP binding protein (gene overexpressed in skeletal muscle) | 1.60 |
| ENSMUSG00000117231 | AC079441.1 | novel transcript, antisense to Dlgap1 | 1.60 |
| ENSMUSG00000045349 | Sh2d5 | SH2 domain containing 5 | 1.60 |
| ENSMUSG00000018378 | Cuedc1 | CUE domain containing 1 | 1.59 |
| ENSMUSG00000010080 | Epn3 | epsin 3 | 1.59 |
| ENSMUSG00000023495 | Pcbp4 | poly(rC) binding protein 4 | 1.59 |
| ENSMUSG00000043895 | S1pr2 | sphingosine-1-phosphate receptor 2 | 1.59 |
| ENSMUSG00000068617 | Efcab1 | EF-hand calcium binding domain 1 | 1.59 |
| ENSMUSG00000026335 | Pam | peptidylglycine alpha-amidating monooxygenase | 1.58 |
| ENSMUSG00000025372 | Baiap2 | brain-specific angiogenesis inhibitor 1-associated protein 2 | 1.58 |
| ENSMUSG00000024579 | Pcyox1l | prenylcysteine oxidase 1 like | 1.58 |
| ENSMUSG00000113665 | Gm47603 | predicted gene, 47603 | 1.58 |
| ENSMUSG00000032228 | Tcf12 | transcription factor 12 | 1.57 |
| ENSMUSG00000053580 | Tanc2 | tetratricopeptide repeat, ankyrin repeat and coiled-coil containing 2 | 1.57 |
| ENSMUSG00000042439 | Zfp532 | zinc finger protein 532 | 1.57 |
| ENSMUSG00000020882 | Cacnb1 | calcium channel, voltage-dependent, beta 1 subunit | 1.57 |
| ENSMUSG00000054967 | Zfp647 | zinc finger protein 647 | 1.57 |
| ENSMUSG00000030729 | Pgm2l1 | phosphoglucomutase 2-like 1 | 1.56 |
| ENSMUSG00000032320 | Rcn2 | reticulocalbin 2 | 1.56 |
| ENSMUSG00000002458 | Rgs19 | regulator of G-protein signaling 19 | 1.56 |
| ENSMUSG00000027737 | Slc7a11 | solute carrier family 7 (cationic amino acid transporter, y+ system), member 11 | 1.56 |
| ENSMUSG00000085116 | Gm2735 | predicted gene 2735 | 1.56 |
| ENSMUSG00000087201 | Gm15261 | predicted gene 15261 | 1.56 |
| ENSMUSG00000052563 | D930048N14Rik | RIKEN cDNA D930048N14 gene | 1.55 |

| | | | |
|---|---|---|---|
| ENSMUSG00000064125 | Prr36 | proline rich 36 | 1.55 |
| ENSMUSG00000008153 | Clstn3 | calsyntenin 3 | 1.55 |
| ENSMUSG00000042426 | Dhx29 | DEAH (Asp-Glu-Ala-His) box polypeptide 29 | 1.54 |
| ENSMUSG00000001053 | N4bp3 | NEDD4 binding protein 3 | 1.54 |
| ENSMUSG00000068115 | Ninl | ninein-like | 1.54 |
| ENSMUSG00000102440 | Pcdhga9 | protocadherin gamma subfamily A, 9 | 1.54 |
| ENSMUSG00000034118 | Tpst1 | protein-tyrosine sulfotransferase 1 | 1.54 |
| ENSMUSG00000020723 | Cacng4 | calcium channel, voltage-dependent, gamma subunit 4 | 1.54 |
| ENSMUSG00000049580 | Tsku | tsukushi, small leucine rich proteoglycan | 1.54 |
| ENSMUSG00000006519 | Cyba | cytochrome b-245, alpha polypeptide | 1.53 |
| ENSMUSG00000025151 | Maged1 | melanoma antigen, family D, 1 | 1.53 |
| ENSMUSG00000033319 | Fem1c | fem-1 homolog c (C.elegans) | 1.53 |
| ENSMUSG00000026425 | Srgap2 | SLIT-ROBO Rho GTPase activating protein 2 | 1.53 |
| ENSMUSG00000021720 | Rnf180 | ring finger protein 180 | 1.53 |
| ENSMUSG00000026627 | Tmem206 | transmembrane protein 206 | 1.53 |
| ENSMUSG00000024909 | Efemp2 | epidermal growth factor-containing fibulin-like extracellular matrix protein 2 | 1.53 |
| ENSMUSG00000037890 | Wdr19 | WD repeat domain 19 | 1.52 |
| ENSMUSG00000005087 | Cd44 | CD44 antigen | 1.50 |
| ENSMUSG00000032359 | Ctsh | cathepsin H | 1.50 |
| ENSMUSG00000039831 | Arhgap29 | Rho GTPase activating protein 29 | 1.50 |
| ENSMUSG00000025035 | Arl3 | ADP-ribosylation factor-like 3 | 1.50 |
| ENSMUSG00000102748 | Pcdhgb2 | protocadherin gamma subfamily B, 2 | 1.50 |
| ENSMUSG00000022895 | Ets2 | E26 avian leukemia oncogene 2, 3' domain | 1.49 |
| ENSMUSG00000025153 | Fasn | fatty acid synthase | 1.49 |
| ENSMUSG00000021482 | Aaed1 | AhpC/TSA antioxidant enzyme domain containing 1 | 1.49 |
| ENSMUSG00000028268 | Gbp3 | guanylate binding protein 3 | 1.49 |
| ENSMUSG00000074211 | Sdhaf1 | succinate dehydrogenase complex assembly factor 1 | 1.49 |
| ENSMUSG00000055639 | Dach1 | dachshund family transcription factor 1 | 1.48 |
| ENSMUSG00000024177 | Nme4 | NME/NM23 nucleoside diphosphate kinase 4 | 1.48 |
| ENSMUSG00000029999 | Tgfa | transforming growth factor alpha | 1.47 |
| ENSMUSG00000038007 | Acer2 | alkaline ceramidase 2 | 1.47 |
| ENSMUSG00000038886 | Man2a2 | mannosidase 2, alpha 2 | 1.47 |
| ENSMUSG00000068270 | Shroom4 | shroom family member 4 | 1.47 |
| ENSMUSG00000038587 | Akap12 | A kinase (PRKA) anchor protein (gravin) 12 | 1.47 |
| ENSMUSG00000021109 | Hif1a | hypoxia inducible factor 1, alpha subunit | 1.46 |
| ENSMUSG00000028173 | Wls | wntless WNT ligand secretion mediator | 1.46 |
| ENSMUSG00000036368 | Rmdn2 | regulator of microtubule dynamics 2 | 1.46 |
| ENSMUSG00000027223 | Mapk8ip1 | mitogen-activated protein kinase 8 interacting protein 1 | 1.46 |
| ENSMUSG00000034586 | Hid1 | HID1 domain containing | 1.45 |
| ENSMUSG00000020536 | Llgl1 | LLGL1 scribble cell polarity complex component | 1.45 |
| ENSMUSG00000040511 | Pvr | poliovirus receptor | 1.45 |
| ENSMUSG00000031303 | Map3k15 | mitogen-activated protein kinase kinase kinase 15 | 1.45 |
| ENSMUSG00000027784 | Ppm1l | protein phosphatase 1 (formerly 2C)-like | 1.45 |
| ENSMUSG00000072235 | Tuba1a | tubulin, alpha 1A | 1.45 |
| ENSMUSG00000048755 | Mcat | malonyl CoA:ACP acyltransferase (mitochondrial) | 1.45 |
| ENSMUSG00000041729 | Coro2b | coronin, actin binding protein, 2B | 1.45 |
| ENSMUSG00000059248 | Sept9 | septin 9 | 1.44 |
| ENSMUSG00000024098 | Twsg1 | twisted gastrulation BMP signaling modulator 1 | 1.44 |
| ENSMUSG00000046668 | Cxxc5 | CXXC finger 5 | 1.44 |
| ENSMUSG00000047090 | Tmem198b | transmembrane protein 198b | 1.44 |
| ENSMUSG00000022637 | Cblb | Casitas B-lineage lymphoma b | 1.43 |
| ENSMUSG00000040669 | Phc1 | polyhomeotic 1 | 1.43 |
| ENSMUSG00000049532 | Sall2 | spalt like transcription factor 2 | 1.43 |
| ENSMUSG00000083012 | Fam220a | family with sequence similarity 220, member A | 1.43 |
| ENSMUSG00000004837 | Grap | GRB2-related adaptor protein | 1.43 |
| ENSMUSG00000022096 | Hr | hairless | 1.42 |
| ENSMUSG00000038648 | Creb3l2 | cAMP responsive element binding protein 3-like 2 | 1.42 |
| ENSMUSG00000054256 | Msi1 | musashi RNA-binding protein 1 | 1.42 |
| ENSMUSG00000031360 | Ctps2 | cytidine 5'-triphosphate synthase 2 | 1.42 |
| ENSMUSG00000000861 | Bcl11a | B cell CLL/lymphoma 11A (zinc finger protein) | 1.42 |
| ENSMUSG00000036968 | Cnpy4 | canopy FGF signaling regulator 4 | 1.42 |
| ENSMUSG00000024818 | Slc25a45 | solute carrier family 25, member 45 | 1.42 |
| ENSMUSG00000025787 | Tgm4 | transglutaminase 4 (prostate) | 1.42 |
| ENSMUSG00000013698 | Pea15a | phosphoprotein enriched in astrocytes 15A | 1.41 |
| ENSMUSG00000033900 | Map9 | microtubule-associated protein 9 | 1.41 |
| ENSMUSG00000019986 | Ahi1 | Abelson helper integration site 1 | 1.41 |
| ENSMUSG00000051391 | Ywhag | tyrosine 3-monooxygenase/tryptophan 5-monooxygenase activation protein, gamma polypeptide | 1.40 |
| ENSMUSG00000022390 | Zc3h7b | zinc finger CCCH type containing 7B | 1.40 |
| ENSMUSG00000025223 | Ldb1 | LIM domain binding 1 | 1.40 |
| ENSMUSG00000000085 | Scmh1 | sex comb on midleg homolog 1 | 1.40 |
| ENSMUSG00000038151 | Prdm1 | PR domain containing 1, with ZNF domain | 1.40 |
| ENSMUSG00000022604 | Cep97 | centrosomal protein 97 | 1.40 |
| ENSMUSG00000041135 | Ripk2 | receptor (TNFRSF)-interacting serine-threonine kinase 2 | 1.40 |
| ENSMUSG00000052534 | Pbx1 | pre B cell leukemia homeobox 1 | 1.39 |
| ENSMUSG00000067071 | Hes6 | hairy and enhancer of split 6 | 1.39 |
| ENSMUSG00000028749 | Pla2g2f | phospholipase A2, group IIF | 1.39 |

| | | | |
|---|---|---|---|
| ENSMUSG00000069631 | Strada | STE20-related kinase adaptor alpha | 1.39 |
| ENSMUSG00000042506 | Usp22 | ubiquitin specific peptidase 22 | 1.38 |
| ENSMUSG00000029763 | Exoc4 | exocyst complex component 4 | 1.38 |
| ENSMUSG00000060510 | Zfp266 | zinc finger protein 266 | 1.38 |
| ENSMUSG00000040681 | Hmgn1 | high mobility group nucleosomal binding domain 1 | 1.38 |
| ENSMUSG00000053414 | Hunk | hormonally upregulated Neu-associated kinase | 1.38 |
| ENSMUSG00000041440 | Gk5 | glycerol kinase 5 (putative) | 1.38 |
| ENSMUSG00000032285 | Dnaja4 | DnaJ heat shock protein family (Hsp40) member A4 | 1.38 |
| ENSMUSG00000030725 | Lipt2 | lipoyl(octanoyl) transferase 2 (putative) | 1.38 |
| ENSMUSG00000070883 | Ccdc173 | coiled-coil domain containing 173 | 1.38 |
| ENSMUSG00000031626 | Sorbs2 | sorbin and SH3 domain containing 2 | 1.37 |
| ENSMUSG00000039801 | Cplane1 | ciliogenesis and planar polarity effector 1 | 1.37 |
| ENSMUSG00000024855 | Pacs1 | phosphofurin acidic cluster sorting protein 1 | 1.37 |
| ENSMUSG00000004768 | Rab23 | RAB23, member RAS oncogene family | 1.37 |
| ENSMUSG00000031342 | Gpm6b | glycoprotein m6b | 1.37 |
| ENSMUSG00000107205 | Gm42576 | predicted gene 42576 | 1.37 |
| ENSMUSG00000036097 | Slf2 | SMC5-SMC6 complex localization factor 2 | 1.36 |
| ENSMUSG00000002489 | Tiam1 | T cell lymphoma invasion and metastasis 1 | 1.36 |
| ENSMUSG00000027189 | Trim44 | tripartite motif-containing 44 | 1.36 |
| ENSMUSG00000066357 | Wdr6 | WD repeat domain 6 | 1.36 |
| ENSMUSG00000057147 | Dph6 | diphthamine biosynthesis 6 | 1.36 |
| ENSMUSG00000095098 | Ccdc85b | coiled-coil domain containing 85B | 1.36 |
| ENSMUSG00000074825 | Itpripl1 | inositol 1,4,5-triphosphate receptor interacting protein-like 1 | 1.36 |
| ENSMUSG00000035104 | Eva1a | eva-1 homolog A (C. elegans) | 1.36 |
| ENSMUSG00000039262 | Prrc2b | proline-rich coiled-coil 2B | 1.35 |
| ENSMUSG00000026305 | Lrrfip1 | leucine rich repeat (in FLII) interacting protein 1 | 1.35 |
| ENSMUSG00000037145 | 2210407C18Rik | RIKEN cDNA 2210407C18 gene | 1.35 |
| ENSMUSG00000022475 | Hdac7 | histone deacetylase 7 | 1.35 |
| ENSMUSG00000018199 | Trove2 | TROVE domain family, member 2 | 1.35 |
| ENSMUSG00000000325 | Arvcf | armadillo repeat gene deleted in velocardiofacial syndrome | 1.35 |
| ENSMUSG00000037640 | Zfp60 | zinc finger protein 60 | 1.35 |
| ENSMUSG00000030980 | Knop1 | lysine rich nucleolar protein 1 | 1.34 |
| ENSMUSG00000039976 | Tbc1d16 | TBC1 domain family, member 16 | 1.34 |
| ENSMUSG00000020654 | Adcy3 | adenylate cyclase 3 | 1.34 |
| ENSMUSG00000031861 | Lpar2 | lysophosphatidic acid receptor 2 | 1.34 |
| ENSMUSG00000039481 | Nrtn | neurturin | 1.34 |
| ENSMUSG00000054693 | Adam10 | a disintegrin and metallopeptidase domain 10 | 1.33 |
| ENSMUSG00000029822 | Osbpl3 | oxysterol binding protein-like 3 | 1.33 |
| ENSMUSG00000019461 | Plscr3 | phospholipid scramblase 3 | 1.33 |
| ENSMUSG00000031434 | Morc4 | microrchidia 4 | 1.33 |
| ENSMUSG00000024350 | Dnajc18 | DnaJ heat shock protein family (Hsp40) member C18 | 1.33 |
| ENSMUSG00000023947 | Nfkbie | nuclear factor of kappa light polypeptide gene enhancer in B cells inhibitor, epsilon | 1.33 |
| ENSMUSG00000038264 | Sema7a | sema domain, immunoglobulin domain (Ig), and GPI membrane anchor, (semaphorin) 7A | 1.32 |
| ENSMUSG00000020015 | Cdk17 | cyclin-dependent kinase 17 | 1.32 |
| ENSMUSG00000035776 | Cd99l2 | CD99 antigen-like 2 | 1.32 |
| ENSMUSG00000033773 | Rpap2 | RNA polymerase II associated protein 2 | 1.32 |
| ENSMUSG00000036052 | Dnajb5 | DnaJ heat shock protein family (Hsp40) member B5 | 1.32 |
| ENSMUSG00000025321 | Itgb8 | integrin beta 8 | 1.32 |
| ENSMUSG00000027878 | Notch2 | notch 2 | 1.31 |
| ENSMUSG00000033669 | Zfp7 | zinc finger protein 7 | 1.31 |
| ENSMUSG00000033545 | Znrf1 | zinc and ring finger 1 | 1.31 |
| ENSMUSG00000061981 | Flot2 | flotillin 2 | 1.31 |
| ENSMUSG00000040811 | Eml2 | echinoderm microtubule associated protein like 2 | 1.31 |
| ENSMUSG00000038718 | Pbx3 | pre B cell leukemia homeobox 3 | 1.31 |
| ENSMUSG00000014907 | Naf1 | nuclear assembly factor 1 ribonucleoprotein | 1.31 |
| ENSMUSG00000046223 | Plaur | plasminogen activator, urokinase receptor | 1.31 |
| ENSMUSG00000107000 | Gm43481 | predicted gene 43481 | 1.31 |
| ENSMUSG00000085433 | Gm16001 | predicted gene 16001 | 1.31 |
| ENSMUSG00000072694 | 1500011B03Rik | RIKEN cDNA 1500011B03 gene | 1.31 |
| ENSMUSG00000027684 | Mecom | MDS1 and EVI1 complex locus | 1.30 |
| ENSMUSG00000033767 | Tmem131l | transmembrane 131 like | 1.30 |
| ENSMUSG00000086040 | Wipf3 | WAS/WASL interacting protein family, member 3 | 1.30 |
| ENSMUSG00000022098 | Bmp1 | bone morphogenetic protein 1 | 1.30 |
| ENSMUSG00000025203 | Scd2 | stearoyl-Coenzyme A desaturase 2 | 1.29 |
| ENSMUSG00000036206 | Sh3bp4 | SH3-domain binding protein 4 | 1.29 |
| ENSMUSG00000025969 | Nrp2 | neuropilin 2 | 1.29 |
| ENSMUSG00000036006 | Ripor2 | RHO family interacting cell polarization regulator 2 | 1.29 |
| ENSMUSG00000021368 | Tbc1d7 | TBC1 domain family, member 7 | 1.29 |
| ENSMUSG00000007659 | Bcl2l1 | BCL2-like 1 | 1.28 |
| ENSMUSG00000041238 | Rbbp8 | retinoblastoma binding protein 8, endonuclease | 1.28 |
| ENSMUSG00000046574 | Prr12 | proline rich 12 | 1.28 |
| ENSMUSG00000020142 | Slc1a4 | solute carrier family 1 (glutamate/neutral amino acid transporter), member 4 | 1.28 |
| ENSMUSG00000040296 | Ddx58 | DEAD (Asp-Glu-Ala-Asp) box polypeptide 58 | 1.28 |
| ENSMUSG00000024501 | Dpysl3 | dihydropyrimidinase-like 3 | 1.28 |
| ENSMUSG00000028599 | Tnfrsf1b | tumor necrosis factor receptor superfamily, member 1b | 1.28 |
| ENSMUSG00000025931 | Paqr8 | progestin and adipoQ receptor family member VIII | 1.28 |

| | | | |
|---|---|---|---|
| ENSMUSG00000032883 | Acsl3 | acyl-CoA synthetase long-chain family member 3 | 1.27 |
| ENSMUSG00000021277 | Traf3 | TNF receptor-associated factor 3 | 1.27 |
| ENSMUSG00000006800 | Sulf2 | sulfatase 2 | 1.27 |
| ENSMUSG00000027122 | Arl14ep | ADP-ribosylation factor-like 14 effector protein | 1.27 |
| ENSMUSG00000030170 | Wnt5b | wingless-type MMTV integration site family, member 5B | 1.27 |
| ENSMUSG00000020607 | Fam84a | family with sequence similarity 84, member A | 1.26 |
| ENSMUSG00000029229 | Chic2 | cysteine-rich hydrophobic domain 2 | 1.26 |
| ENSMUSG00000022971 | Ifnar2 | interferon (alpha and beta) receptor 2 | 1.26 |
| ENSMUSG00000025082 | Vwa2 | von Willebrand factor A domain containing 2 | 1.26 |
| ENSMUSG00000031799 | Tpm4 | tropomyosin 4 | 1.25 |
| ENSMUSG00000035547 | Capn5 | calpain 5 | 1.25 |
| ENSMUSG00000048371 | Pdp2 | pyruvate dehyrogenase phosphatase catalytic subunit 2 | 1.25 |
| ENSMUSG00000043467 | Zbtb37 | zinc finger and BTB domain containing 37 | 1.25 |
| ENSMUSG00000019261 | Map1s | microtubule-associated protein 1S | 1.25 |
| ENSMUSG00000030323 | Ift122 | intraflagellar transport 122 | 1.25 |
| ENSMUSG00000117123 | AL805899.1 | novel transcript | 1.25 |
| ENSMUSG00000043461 | Sptssb | serine palmitoyltransferase, small subunit B | 1.25 |
| ENSMUSG00000021782 | Dlg5 | discs large MAGUK scaffold protein 5 | 1.24 |
| ENSMUSG00000036885 | Arhgef26 | Rho guanine nucleotide exchange factor (GEF) 26 | 1.24 |
| ENSMUSG00000031278 | Acsl4 | acyl-CoA synthetase long-chain family member 4 | 1.24 |
| ENSMUSG00000070803 | Cited4 | Cbp/p300-interacting transactivator, with Glu/Asp-rich carboxy-terminal domain, 4 | 1.24 |
| ENSMUSG00000079038 | D130040H23Rik | RIKEN cDNA D130040H23 gene | 1.24 |
| ENSMUSG00000031903 | Pla2g15 | phospholipase A2, group XV | 1.24 |
| ENSMUSG00000023169 | Slc38a1 | solute carrier family 38, member 1 | 1.23 |
| ENSMUSG00000022723 | Crybg3 | beta-gamma crystallin domain containing 3 | 1.23 |
| ENSMUSG00000058135 | Gstm1 | glutathione S-transferase, mu 1 | 1.23 |
| ENSMUSG00000021608 | Lpcat1 | lysophosphatidylcholine acyltransferase 1 | 1.23 |
| ENSMUSG00000068758 | Il3ra | interleukin 3 receptor, alpha chain | 1.23 |
| ENSMUSG00000019966 | Kitl | kit ligand | 1.22 |
| ENSMUSG00000050912 | Tmem123 | transmembrane protein 123 | 1.22 |
| ENSMUSG00000019433 | Gipc1 | GIPC PDZ domain containing family, member 1 | 1.22 |
| ENSMUSG00000021774 | Ube2e1 | ubiquitin-conjugating enzyme E2E 1 | 1.22 |
| ENSMUSG00000028344 | Invs | inversin | 1.22 |
| ENSMUSG00000038721 | Hoxb7 | homeobox B7 | 1.21 |
| ENSMUSG00000028601 | Echdc2 | enoyl Coenzyme A hydratase domain containing 2 | 1.21 |
| ENSMUSG00000045211 | Nudt18 | nudix (nucleoside diphosphate linked moiety X)-type motif 18 | 1.21 |
| ENSMUSG00000006356 | Crip2 | cysteine rich protein 2 | 1.21 |
| ENSMUSG00000083282 | Ctsf | cathepsin F | 1.21 |
| ENSMUSG00000035021 | Baz1a | bromodomain adjacent to zinc finger domain 1A | 1.20 |
| ENSMUSG00000039985 | Sinhcaf | SIN3-HDAC complex associated factor | 1.20 |
| ENSMUSG00000040446 | Rprd1a | regulation of nuclear pre-mRNA domain containing 1A | 1.20 |
| ENSMUSG00000025262 | Fam120c | family with sequence similarity 120, member C | 1.20 |
| ENSMUSG00000025196 | Cpn1 | carboxypeptidase N, polypeptide 1 | 1.20 |
| ENSMUSG00000051705 | Senp8 | SUMO/sentrin specific peptidase 8 | 1.20 |
| ENSMUSG00000035713 | Usp35 | ubiquitin specific peptidase 35 | 1.20 |
| ENSMUSG00000015745 | Plekho1 | pleckstrin homology domain containing, family O member 1 | 1.20 |
| ENSMUSG00000063382 | Bcl9l | B cell CLL/lymphoma 9-like | 1.19 |
| ENSMUSG00000020376 | Rnf130 | ring finger protein 130 | 1.19 |
| ENSMUSG00000048264 | Dip2c | disco interacting protein 2 homolog C | 1.19 |
| ENSMUSG00000058704 | Memo1 | mediator of cell motility 1 | 1.19 |
| ENSMUSG00000020393 | Kremen1 | kringle containing transmembrane protein 1 | 1.19 |
| ENSMUSG00000033970 | Rfc3 | replication factor C (activator 1) 3 | 1.19 |
| ENSMUSG00000038301 | Snx10 | sorting nexin 10 | 1.19 |
| ENSMUSG00000067928 | Zfp760 | zinc finger protein 760 | 1.19 |
| ENSMUSG00000043542 | Zc2hc1a | zinc finger, C2HC-type containing 1A | 1.19 |
| ENSMUSG00000074785 | Plxnc1 | plexin C1 | 1.19 |
| ENSMUSG00000070565 | Rasal2 | RAS protein activator like 2 | 1.18 |
| ENSMUSG00000042105 | Inpp5f | inositol polyphosphate-5-phosphatase F | 1.18 |
| ENSMUSG00000040502 | 39873 | membrane-associated ring finger (C3HC4) 9 | 1.18 |
| ENSMUSG00000044254 | Pcsk9 | proprotein convertase subtilisin/kexin type 9 | 1.17 |
| ENSMUSG00000040928 | S100pbp | S100P binding protein | 1.17 |
| ENSMUSG00000049439 | Cyp20a1 | cytochrome P450, family 20, subfamily a, polypeptide 1 | 1.17 |
| ENSMUSG00000038387 | Rras | related RAS viral (r-ras) oncogene | 1.17 |
| ENSMUSG00000087366 | Junos | jun proto-oncogene, opposite strand | 1.17 |
| ENSMUSG00000109324 | Prmt1 | protein arginine N-methyltransferase 1 | 1.16 |
| ENSMUSG00000041895 | Wipi1 | WD repeat domain, phosphoinositide interacting 1 | 1.16 |
| ENSMUSG00000021451 | Sema4d | sema domain, immunoglobulin domain (Ig), transmembrane domain (TM) and short cytoplasmic do | 1.16 |
| ENSMUSG00000027533 | Fabp5 | fatty acid binding protein 5, epidermal | 1.16 |
| ENSMUSG00000025887 | Casp12 | caspase 12 | 1.16 |
| ENSMUSG00000004961 | Syt5 | synaptotagmin V | 1.16 |
| ENSMUSG00000041298 | Katnal1 | katanin p60 subunit A-like 1 | 1.16 |
| ENSMUSG00000020875 | Hoxb9 | homeobox B9 | 1.15 |
| ENSMUSG00000026421 | Csrp1 | cysteine and glycine-rich protein 1 | 1.15 |
| ENSMUSG00000033454 | Zbtb1 | zinc finger and BTB domain containing 1 | 1.15 |
| ENSMUSG00000030946 | Lhpp | phospholysine phosphohistidine inorganic pyrophosphate phosphatase | 1.15 |
| ENSMUSG00000031641 | Cbr4 | carbonyl reductase 4 | 1.15 |

| | | | |
|---|---|---|---|
| ENSMUSG00000069114 | Zbtb10 | zinc finger and BTB domain containing 10 | 1.15 |
| ENSMUSG00000087177 | E130307A14Rik | RIKEN cDNA E130307A14 gene | 1.15 |
| ENSMUSG00000057133 | Chd6 | chromodomain helicase DNA binding protein 6 | 1.14 |
| ENSMUSG00000026566 | Mpzl1 | myelin protein zero-like 1 | 1.14 |
| ENSMUSG00000031150 | Ccdc120 | coiled-coil domain containing 120 | 1.14 |
| ENSMUSG00000028542 | Slc6a9 | solute carrier family 6 (neurotransmitter transporter, glycine), member 9 | 1.14 |
| ENSMUSG00000025323 | Sp4 | trans-acting transcription factor 4 | 1.14 |
| ENSMUSG00000062590 | Armc9 | armadillo repeat containing 9 | 1.14 |
| ENSMUSG00000115902 | AC113595.1 | novel transcript, antisense to Apobec3 | 1.14 |
| ENSMUSG00000022837 | Iqcb1 | IQ calmodulin-binding motif containing 1 | 1.14 |
| ENSMUSG00000115783 | Bc1 | brain cytoplasmic RNA 1 | 1.14 |
| ENSMUSG00000032216 | Nedd4 | neural precursor cell expressed, developmentally down-regulated 4 | 1.13 |
| ENSMUSG00000014601 | Strip1 | striatin interacting protein 1 | 1.13 |
| ENSMUSG00000025142 | Aspscr1 | alveolar soft part sarcoma chromosome region, candidate 1 (human) | 1.13 |
| ENSMUSG00000034981 | Parm1 | prostate androgen-regulated mucin-like protein 1 | 1.13 |
| ENSMUSG00000022822 | Abcc5 | ATP-binding cassette, sub-family C (CFTR/MRP), member 5 | 1.13 |
| ENSMUSG00000028245 | Nsmaf | neutral sphingomyelinase (N-SMase) activation associated factor | 1.13 |
| ENSMUSG00000046432 | Bex3 | brain expressed X-linked 3 | 1.13 |
| ENSMUSG00000026484 | Rnf2 | ring finger protein 2 | 1.13 |
| ENSMUSG00000032481 | Smarcc1 | SWI/SNF related, matrix associated, actin dependent regulator of chromatin, subfamily c, member 1 | 1.12 |
| ENSMUSG00000029478 | Ncor2 | nuclear receptor co-repressor 2 | 1.12 |
| ENSMUSG00000030413 | Pglyrp1 | peptidoglycan recognition protein 1 | 1.12 |
| ENSMUSG00000004846 | Plod3 | procollagen-lysine, 2-oxoglutarate 5-dioxygenase 3 | 1.12 |
| ENSMUSG00000022197 | Pdzd2 | PDZ domain containing 2 | 1.12 |
| ENSMUSG00000056148 | Rdh9 | retinol dehydrogenase 9 | 1.12 |
| ENSMUSG00000042050 | Wdr60 | WD repeat domain 60 | 1.12 |
| ENSMUSG00000074283 | Zfp109 | zinc finger protein 109 | 1.12 |
| ENSMUSG00000112964 | E430024I08Rik | RIKEN cDNA E430024I08 gene | 1.12 |
| ENSMUSG00000019838 | Slc16a10 | solute carrier family 16 (monocarboxylic acid transporters), member 10 | 1.11 |
| ENSMUSG00000028919 | Arhgef19 | Rho guanine nucleotide exchange factor (GEF) 19 | 1.11 |
| ENSMUSG00000037933 | Bicd2 | BICD cargo adaptor 2 | 1.11 |
| ENSMUSG00000028126 | Pip5k1a | phosphatidylinositol-4-phosphate 5-kinase, type 1 alpha | 1.11 |
| ENSMUSG00000067430 | Zfp763 | zinc finger protein 763 | 1.11 |
| ENSMUSG00000025171 | Ubtd1 | ubiquitin domain containing 1 | 1.11 |
| ENSMUSG00000011254 | Thg1l | tRNA-histidine guanylyltransferase 1-like (S. cerevisiae) | 1.11 |
| ENSMUSG00000016487 | Ppfibp1 | PTPRF interacting protein, binding protein 1 (liprin beta 1) | 1.10 |
| ENSMUSG00000052949 | Rnf157 | ring finger protein 157 | 1.10 |
| ENSMUSG00000024640 | Psat1 | phosphoserine aminotransferase 1 | 1.10 |
| ENSMUSG00000020167 | Tcf3 | transcription factor 3 | 1.09 |
| ENSMUSG00000036275 | 9530068E07Rik | RIKEN cDNA 9530068E07 gene | 1.09 |
| ENSMUSG00000051934 | Spats2 | spermatogenesis associated, serine-rich 2 | 1.09 |
| ENSMUSG00000026782 | Abi2 | abl-interactor 2 | 1.09 |
| ENSMUSG00000050714 | Zbtb26 | zinc finger and BTB domain containing 26 | 1.09 |
| ENSMUSG00000022003 | Slc25a30 | solute carrier family 25, member 30 | 1.09 |
| ENSMUSG00000029657 | Hsph1 | heat shock 105kDa/110kDa protein 1 | 1.08 |
| ENSMUSG00000037706 | Cd81 | CD81 antigen | 1.08 |
| ENSMUSG00000020140 | Lgr5 | leucine rich repeat containing G protein coupled receptor 5 | 1.08 |
| ENSMUSG00000048047 | Zbtb33 | zinc finger and BTB domain containing 33 | 1.08 |
| ENSMUSG00000040327 | Cul9 | cullin 9 | 1.08 |
| ENSMUSG00000049300 | Prmt6 | protein arginine N-methyltransferase 6 | 1.08 |
| ENSMUSG00000024269 | Tpgs2 | tubulin polyglutamylase complex subunit 2 | 1.08 |
| ENSMUSG00000021108 | Prkch | protein kinase C, eta | 1.08 |
| ENSMUSG00000033565 | Rbfox2 | RNA binding protein, fox-1 homolog (C. elegans) 2 | 1.07 |
| ENSMUSG00000046312 | AI464131 | expressed sequence AI464131 | 1.07 |
| ENSMUSG00000027495 | Fam210b | family with sequence similarity 210, member B | 1.07 |
| ENSMUSG00000022434 | Fam118a | family with sequence similarity 118, member A | 1.07 |
| ENSMUSG00000045071 | E130308A19Rik | RIKEN cDNA E130308A19 gene | 1.07 |
| ENSMUSG00000037907 | Ankrd13b | ankyrin repeat domain 13b | 1.07 |
| ENSMUSG00000034723 | Tmx4 | thioredoxin-related transmembrane protein 4 | 1.07 |
| ENSMUSG00000027778 | Ift80 | intraflagellar transport 80 | 1.07 |
| ENSMUSG00000108238 | Gm43984 | predicted gene, 43984 | 1.07 |
| ENSMUSG00000028953 | Abcf2 | ATP-binding cassette, sub-family F (GCN20), member 2 | 1.06 |
| ENSMUSG00000006728 | Cdk4 | cyclin-dependent kinase 4 | 1.06 |
| ENSMUSG00000059552 | Trp53 | transformation related protein 53 | 1.06 |
| ENSMUSG00000061689 | Dlgap4 | DLG associated protein 4 | 1.06 |
| ENSMUSG00000001761 | Smo | smoothened, frizzled class receptor | 1.06 |
| ENSMUSG00000045598 | Zfp553 | zinc finger protein 553 | 1.06 |
| ENSMUSG00000027551 | Zfp64 | zinc finger protein 64 | 1.06 |
| ENSMUSG00000079478 | Sssca1 | Sjogren syndrome/scleroderma autoantigen 1 | 1.06 |
| ENSMUSG00000001525 | Tubb5 | tubulin, beta 5 class I | 1.05 |
| ENSMUSG00000020458 | Rtn4 | reticulon 4 | 1.05 |
| ENSMUSG00000023039 | Krt7 | keratin 7 | 1.05 |
| ENSMUSG00000034300 | Fam53c | family with sequence similarity 53, member C | 1.05 |
| ENSMUSG00000034708 | Grn | granulin | 1.04 |
| ENSMUSG00000028484 | Psip1 | PC4 and SFRS1 interacting protein 1 | 1.04 |
| ENSMUSG00000022673 | Mcm4 | minichromosome maintenance complex component 4 | 1.04 |

| | | | |
|---|---|---|---|
| ENSMUSG00000036099 | Vezt | vezatin, adherens junctions transmembrane protein | 1.04 |
| ENSMUSG00000044968 | Napepld | N-acyl phosphatidylethanolamine phospholipase D | 1.04 |
| ENSMUSG00000024906 | Mus81 | MUS81 structure-specific endonuclease subunit | 1.04 |
| ENSMUSG00000035451 | Foxa1 | forkhead box A1 | 1.03 |
| ENSMUSG00000028832 | Stmn1 | stathmin 1 | 1.03 |
| ENSMUSG00000038034 | Igsf8 | immunoglobulin superfamily, member 8 | 1.03 |
| ENSMUSG00000060301 | 2610008E11Rik | RIKEN cDNA 2610008E11 gene | 1.03 |
| ENSMUSG00000049882 | Vcpkmt | valosin containing protein lysine (K) methyltransferase | 1.03 |
| ENSMUSG00000033060 | Lmo7 | LIM domain only 7 | 1.02 |
| ENSMUSG00000024811 | Tnks2 | tankyrase, TRF1-interacting ankyrin-related ADP-ribose polymerase 2 | 1.02 |
| ENSMUSG00000025085 | Ablim1 | actin-binding LIM protein 1 | 1.02 |
| ENSMUSG00000027087 | Itgav | integrin alpha V | 1.02 |
| ENSMUSG00000022952 | Runx1 | runt related transcription factor 1 | 1.02 |
| ENSMUSG00000001630 | Stk38l | serine/threonine kinase 38 like | 1.02 |
| ENSMUSG00000002870 | Mcm2 | minichromosome maintenance complex component 2 | 1.02 |
| ENSMUSG00000105703 | Gm43305 | predicted gene 43305 | 1.02 |
| ENSMUSG00000058318 | Phf21a | PHD finger protein 21A | 1.02 |
| ENSMUSG00000019971 | Cep290 | centrosomal protein 290 | 1.02 |
| ENSMUSG00000021068 | Nin | ninein | 1.02 |
| ENSMUSG00000028517 | Plpp3 | phospholipid phosphatase 3 | 1.02 |
| ENSMUSG00000051022 | Hs3st1 | heparan sulfate (glucosamine) 3-O-sulfotransferase 1 | 1.02 |
| ENSMUSG00000033987 | Dnah17 | dynein, axonemal, heavy chain 17 | 1.02 |
| ENSMUSG00000022773 | Ypel1 | yippee like 1 | 1.02 |
| ENSMUSG00000039671 | Zmynd8 | zinc finger, MYND-type containing 8 | 1.01 |
| ENSMUSG00000026639 | Lamb3 | laminin, beta 3 | 1.01 |
| ENSMUSG00000035671 | Zswim4 | zinc finger SWIM-type containing 4 | 1.01 |
| ENSMUSG00000022964 | Tmem50b | transmembrane protein 50B | 1.01 |
| ENSMUSG00000056612 | Ppp1r14b | protein phosphatase 1, regulatory inhibitor subunit 14B | 1.01 |
| ENSMUSG00000055897 | Ppp4r1l-ps | protein phosphatase 4, regulatory subunit 1-like, pseudogene | 1.01 |
| ENSMUSG00000018001 | Cyth3 | cytohesin 3 | 1.01 |
| ENSMUSG00000034064 | Poglut1 | protein O-glucosyltransferase 1 | 1.01 |
| ENSMUSG00000036093 | Arl5a | ADP-ribosylation factor-like 5A | 1.00 |
| ENSMUSG00000029461 | Fam168a | family with sequence similarity 168, member A | 1.00 |
| ENSMUSG00000041343 | Ankrd42 | ankyrin repeat domain 42 | 1.00 |
| ENSMUSG00000042638 | Gucy2c | guanylate cyclase 2c | -1.00 |
| ENSMUSG00000022900 | Ildr1 | immunoglobulin-like domain containing receptor 1 | -1.00 |
| ENSMUSG00000032633 | Flcn | folliculin | -1.00 |
| ENSMUSG00000062110 | Scfd2 | Sec1 family domain containing 2 | -1.00 |
| ENSMUSG00000032593 | Amigo3 | adhesion molecule with Ig like domain 3 | -1.00 |
| ENSMUSG00000049608 | Gpr55 | G protein-coupled receptor 55 | -1.00 |
| ENSMUSG00000027422 | Rrbp1 | ribosome binding protein 1 | -1.01 |
| ENSMUSG00000044042 | Fmn1 | formin 1 | -1.01 |
| ENSMUSG00000090035 | Galnt4 | polypeptide N-acetylgalactosaminyltransferase 4 | -1.01 |
| ENSMUSG00000040048 | Ndufb10 | NADH:ubiquinone oxidoreductase subunit B10 | -1.01 |
| ENSMUSG00000005802 | Slc30a4 | solute carrier family 30 (zinc transporter), member 4 | -1.01 |
| ENSMUSG00000063206 | Defa34 | defensin, alpha, 34 | -1.01 |
| ENSMUSG00000017707 | Serinc3 | serine incorporator 3 | -1.02 |
| ENSMUSG00000020456 | Ogdh | oxoglutarate (alpha-ketoglutarate) dehydrogenase (lipoamide) | -1.02 |
| ENSMUSG00000028792 | Ak2 | adenylate kinase 2 | -1.02 |
| ENSMUSG00000039031 | Arhgap18 | Rho GTPase activating protein 18 | -1.02 |
| ENSMUSG00000018427 | Ypel2 | yippee like 2 | -1.02 |
| ENSMUSG00000070284 | Gmppb | GDP-mannose pyrophosphorylase B | -1.02 |
| ENSMUSG00000021684 | Pde8b | phosphodiesterase 8B | -1.02 |
| ENSMUSG00000051335 | Gfod1 | glucose-fructose oxidoreductase domain containing 1 | -1.02 |
| ENSMUSG00000051517 | Arhgef39 | Rho guanine nucleotide exchange factor (GEF) 39 | -1.02 |
| ENSMUSG00000029136 | Rbks | ribokinase | -1.02 |
| ENSMUSG00000104443 | 4932442E05Rik | RIKEN cDNA 4932442E05 gene | -1.02 |
| ENSMUSG00000038859 | Baiap2l1 | BAI1-associated protein 2-like 1 | -1.03 |
| ENSMUSG00000018899 | Irf1 | interferon regulatory factor 1 | -1.03 |
| ENSMUSG00000009863 | Sdhb | succinate dehydrogenase complex, subunit B, iron sulfur (Ip) | -1.03 |
| ENSMUSG00000024190 | Dusp1 | dual specificity phosphatase 1 | -1.03 |
| ENSMUSG00000008540 | Mgst1 | microsomal glutathione S-transferase 1 | -1.03 |
| ENSMUSG00000032602 | Slc25a20 | solute carrier family 25 (mitochondrial carnitine/acylcarnitine translocase), member 20 | -1.03 |
| ENSMUSG00000015536 | Mocs2 | molybdenum cofactor synthesis 2 | -1.03 |
| ENSMUSG00000025006 | Sorbs1 | sorbin and SH3 domain containing 1 | -1.03 |
| ENSMUSG00000031089 | Slc6a14 | solute carrier family 6 (neurotransmitter transporter), member 14 | -1.03 |
| ENSMUSG00000022332 | Khdrbs3 | KH domain containing, RNA binding, signal transduction associated 3 | -1.03 |
| ENSMUSG00000034731 | Dgkh | diacylglycerol kinase, eta | -1.03 |
| ENSMUSG00000020334 | Slc22a4 | solute carrier family 22 (organic cation transporter), member 4 | -1.03 |
| ENSMUSG00000025545 | Clybl | citrate lyase beta like | -1.03 |
| ENSMUSG00000038702 | Dsel | dermatan sulfate epimerase-like | -1.03 |
| ENSMUSG00000028137 | Celf3 | CUGBP, Elav-like family member 3 | -1.03 |
| ENSMUSG00000050777 | Tmem37 | transmembrane protein 37 | -1.03 |
| ENSMUSG00000020220 | Vps13d | vacuolar protein sorting 13D | -1.04 |
| ENSMUSG00000041870 | Ankrd13a | ankyrin repeat domain 13a | -1.04 |
| ENSMUSG00000020741 | Cluh | clustered mitochondria (cluA/CLU1) homolog | -1.04 |

| | | | |
|---|---|---|---|
| ENSMUSG00000048707 | Tprn | taperin | -1.04 |
| ENSMUSG00000007036 | Abhd16a | abhydrolase domain containing 16A | -1.04 |
| ENSMUSG00000070730 | Rmdn3 | regulator of microtubule dynamics 3 | -1.04 |
| ENSMUSG00000033902 | Mapkbp1 | mitogen-activated protein kinase binding protein 1 | -1.04 |
| ENSMUSG00000025466 | Fuom | fucose mutarotase | -1.04 |
| ENSMUSG00000050721 | Plekho2 | pleckstrin homology domain containing, family O member 2 | -1.04 |
| ENSMUSG00000032278 | Paqr5 | progestin and adipoQ receptor family member V | -1.04 |
| ENSMUSG00000030739 | Myh14 | myosin, heavy polypeptide 14 | -1.05 |
| ENSMUSG00000039953 | Clstn1 | calsyntenin 1 | -1.05 |
| ENSMUSG00000038393 | Txnip | thioredoxin interacting protein | -1.05 |
| ENSMUSG00000034109 | Golim4 | golgi integral membrane protein 4 | -1.05 |
| ENSMUSG00000021876 | Rnase4 | ribonuclease, RNase A family 4 | -1.05 |
| ENSMUSG00000024338 | Psmb8 | proteasome (prosome, macropain) subunit, beta type 8 (large multifunctional peptidase 7) | -1.05 |
| ENSMUSG00000042677 | Zc3h12a | zinc finger CCCH type containing 12A | -1.05 |
| ENSMUSG00000052396 | Mogat2 | monoacylglycerol O-acyltransferase 2 | -1.05 |
| ENSMUSG00000033187 | BC016579 | cDNA sequence, BC016579 | -1.05 |
| ENSMUSG00000038332 | Sesn1 | sestrin 1 | -1.05 |
| ENSMUSG00000092274 | Neat1 | nuclear paraspeckle assembly transcript 1 (non-protein coding) | -1.06 |
| ENSMUSG00000019302 | Atp6v0a1 | ATPase, H+ transporting, lysosomal V0 subunit A1 | -1.06 |
| ENSMUSG00000021591 | Glrx | glutaredoxin | -1.06 |
| ENSMUSG00000021236 | Entpd5 | ectonucleoside triphosphate diphosphohydrolase 5 | -1.06 |
| ENSMUSG00000033124 | Atg9a | autophagy related 9A | -1.06 |
| ENSMUSG00000064120 | Mocs1 | molybdenum cofactor synthesis 1 | -1.06 |
| ENSMUSG00000025002 | Cyp2c55 | cytochrome P450, family 2, subfamily c, polypeptide 55 | -1.06 |
| ENSMUSG00000038987 | Cfap157 | cilia and flagella associated protein 157 | -1.06 |
| ENSMUSG00000020029 | Nudt4 | nudix (nucleoside diphosphate linked moiety X)-type motif 4 | -1.07 |
| ENSMUSG00000040479 | Dgkz | diacylglycerol kinase zeta | -1.07 |
| ENSMUSG00000036402 | Gng12 | guanine nucleotide binding protein (G protein), gamma 12 | -1.07 |
| ENSMUSG00000022575 | Gsdmd | gasdermin D | -1.07 |
| ENSMUSG00000024887 | Asah2 | N-acylsphingosine amidohydrolase 2 | -1.07 |
| ENSMUSG00000027524 | Edn3 | endothelin 3 | -1.07 |
| ENSMUSG00000026856 | Dolpp1 | dolichyl pyrophosphate phosphatase 1 | -1.07 |
| ENSMUSG00000024875 | Yif1a | Yip1 interacting factor homolog A (S. cerevisiae) | -1.07 |
| ENSMUSG00000033161 | Atp1a1 | ATPase, Na+/K+ transporting, alpha 1 polypeptide | -1.08 |
| ENSMUSG00000007038 | Neu1 | neuraminidase 1 | -1.08 |
| ENSMUSG00000033526 | Ppip5k1 | diphosphoinositol pentakisphosphate kinase 1 | -1.08 |
| ENSMUSG00000034613 | Ppm1h | protein phosphatase 1H (PP2C domain containing) | -1.08 |
| ENSMUSG00000035509 | Fbxl21 | F-box and leucine-rich repeat protein 21 | -1.08 |
| ENSMUSG00000024900 | Cpt1a | carnitine palmitoyltransferase 1a, liver | -1.09 |
| ENSMUSG00000027406 | Idh3b | isocitrate dehydrogenase 3 (NAD+) beta | -1.09 |
| ENSMUSG00000034656 | Cacna1a | calcium channel, voltage-dependent, P/Q type, alpha 1A subunit | -1.09 |
| ENSMUSG00000040618 | Pck2 | phosphoenolpyruvate carboxykinase 2 (mitochondrial) | -1.09 |
| ENSMUSG00000031327 | Chic1 | cysteine-rich hydrophobic domain 1 | -1.09 |
| ENSMUSG00000041654 | Slc39a11 | solute carrier family 39 (metal ion transporter), member 11 | -1.09 |
| ENSMUSG00000052776 | Oas1a | 2'-5' oligoadenylate synthetase 1A | -1.09 |
| ENSMUSG00000103711 | Tstd1 | thiosulfate sulfurtransferase (rhodanese)-like domain containing 1 | -1.09 |
| ENSMUSG00000035041 | Creb3l3 | cAMP responsive element binding protein 3-like 3 | -1.09 |
| ENSMUSG00000089960 | Ugt1a1 | UDP glucuronosyltransferase 1 family, polypeptide A1 | -1.09 |
| ENSMUSG00000103693 | Gm37529 | predicted gene, 37529 | -1.09 |
| ENSMUSG00000029189 | Sel1l3 | sel-1 suppressor of lin-12-like 3 (C. elegans) | -1.10 |
| ENSMUSG00000044786 | Zfp36 | zinc finger protein 36 | -1.10 |
| ENSMUSG00000020736 | Nt5c | 5',3'-nucleotidase, cytosolic | -1.10 |
| ENSMUSG00000000204 | Slfn4 | schlafen 4 | -1.10 |
| ENSMUSG00000016239 | Lonrf3 | LON peptidase N-terminal domain and ring finger 3 | -1.10 |
| ENSMUSG00000000184 | Ccnd2 | cyclin D2 | -1.11 |
| ENSMUSG00000046447 | Camk2n1 | calcium/calmodulin-dependent protein kinase II inhibitor 1 | -1.11 |
| ENSMUSG00000020577 | Tspan13 | tetraspanin 13 | -1.11 |
| ENSMUSG00000018334 | Ksr1 | kinase suppressor of ras 1 | -1.11 |
| ENSMUSG00000018509 | Cenpv | centromere protein V | -1.11 |
| ENSMUSG00000031073 | Fgf15 | fibroblast growth factor 15 | -1.11 |
| ENSMUSG00000025227 | Mfsd13a | major facilitator superfamily domain containing 13a | -1.11 |
| ENSMUSG00000023473 | Celsr3 | cadherin, EGF LAG seven-pass G-type receptor 3 | -1.11 |
| ENSMUSG00000025195 | Dnmbp | dynamin binding protein | -1.12 |
| ENSMUSG00000034485 | Uaca | uveal autoantigen with coiled-coil domains and ankyrin repeats | -1.12 |
| ENSMUSG00000020432 | Tcn2 | transcobalamin 2 | -1.12 |
| ENSMUSG00000061455 | Stx17 | syntaxin 17 | -1.12 |
| ENSMUSG00000013160 | Atp6v0d1 | ATPase, H+ transporting, lysosomal V0 subunit D1 | -1.12 |
| ENSMUSG00000049422 | Chchd10 | coiled-coil-helix-coiled-coil-helix domain containing 10 | -1.12 |
| ENSMUSG00000031168 | Ebp | phenylalkylamine Ca2+ antagonist (emopamil) binding protein | -1.12 |
| ENSMUSG00000036390 | Gadd45a | growth arrest and DNA-damage-inducible 45 alpha | -1.12 |
| ENSMUSG00000031788 | Kifc3 | kinesin family member C3 | -1.12 |
| ENSMUSG00000060147 | Serpinb6a | serine (or cysteine) peptidase inhibitor, clade B, member 6a | -1.13 |
| ENSMUSG00000019866 | Crybg1 | crystallin beta-gamma domain containing 1 | -1.13 |
| ENSMUSG00000034435 | Tmem30b | transmembrane protein 30B | -1.13 |
| ENSMUSG00000043003 | Rasef | RAS and EF hand domain containing | -1.13 |
| ENSMUSG00000042377 | Fam83g | family with sequence similarity 83, member G | -1.13 |

| | | | |
|---|---|---|---|
| ENSMUSG00000045312 | Lhfpl2 | lipoma HMGIC fusion partner-like 2 | -1.13 |
| ENSMUSG00000041625 | Ggact | gamma-glutamylamine cyclotransferase | -1.13 |
| ENSMUSG00000002329 | Mdp1 | magnesium-dependent phosphatase 1 | -1.13 |
| ENSMUSG00000045934 | Mtmr11 | myotubularin related protein 11 | -1.13 |
| ENSMUSG00000035835 | Plppr3 | phospholipid phosphatase related 3 | -1.13 |
| ENSMUSG00000069456 | Rdh16 | retinol dehydrogenase 16 | -1.13 |
| ENSMUSG00000056429 | Tgoln1 | trans-golgi network protein | -1.14 |
| ENSMUSG00000063952 | Brpf3 | bromodomain and PHD finger containing, 3 | -1.14 |
| ENSMUSG00000002250 | Ppard | peroxisome proliferator activator receptor delta | -1.14 |
| ENSMUSG00000039193 | Nlrc4 | NLR family, CARD domain containing 4 | -1.14 |
| ENSMUSG00000036138 | Acaa1a | acetyl-Coenzyme A acyltransferase 1A | -1.14 |
| ENSMUSG00000035877 | Zhx3 | zinc fingers and homeoboxes 3 | -1.14 |
| ENSMUSG00000020272 | Stk10 | serine/threonine kinase 10 | -1.14 |
| ENSMUSG00000058022 | Adtrp | androgen dependent TFPI regulating protein | -1.14 |
| ENSMUSG00000037279 | Ovol2 | ovo like zinc finger 2 | -1.14 |
| ENSMUSG00000038777 | Sema6c | sema domain, transmembrane domain (TM), and cytoplasmic domain, (semaphorin) 6C | -1.14 |
| ENSMUSG00000104114 | Gm37297 | predicted gene, 37297 | -1.14 |
| ENSMUSG00000031971 | Ccsap | centriole, cilia and spindle associated protein | -1.14 |
| ENSMUSG00000019370 | Calm3 | calmodulin 3 | -1.15 |
| ENSMUSG00000024378 | Stard4 | StAR-related lipid transfer (START) domain containing 4 | -1.15 |
| ENSMUSG00000029032 | Arhgef16 | Rho guanine nucleotide exchange factor (GEF) 16 | -1.15 |
| ENSMUSG00000028217 | Cdh17 | cadherin 17 | -1.15 |
| ENSMUSG00000021457 | Syk | spleen tyrosine kinase | -1.15 |
| ENSMUSG00000022265 | Ank | progressive ankylosis | -1.15 |
| ENSMUSG00000025648 | Pfkfb4 | 6-phosphofructo-2-kinase/fructose-2,6-biphosphatase 4 | -1.15 |
| ENSMUSG00000039395 | Mreg | melanoregulin | -1.15 |
| ENSMUSG00000028709 | Mob3c | MOB kinase activator 3C | -1.15 |
| ENSMUSG00000085333 | 1700030A11Rik | RIKEN cDNA 1700030A11 gene | -1.15 |
| ENSMUSG00000032083 | Apoa1 | apolipoprotein A-I | -1.15 |
| ENSMUSG00000056025 | Clca3a1 | chloride channel accessory 3A1 | -1.15 |
| ENSMUSG00000030769 | Slc5a11 | solute carrier family 5 (sodium/glucose cotransporter), member 11 | -1.15 |
| ENSMUSG00000025575 | Cant1 | calcium activated nucleotidase 1 | -1.16 |
| ENSMUSG00000019982 | Myb | myeloblastosis oncogene | -1.16 |
| ENSMUSG00000028088 | Fmo5 | flavin containing monooxygenase 5 | -1.16 |
| ENSMUSG00000040631 | Dok4 | docking protein 4 | -1.16 |
| ENSMUSG00000026614 | Slc30a10 | solute carrier family 30, member 10 | -1.16 |
| ENSMUSG00000037868 | Egr2 | early growth response 2 | -1.16 |
| ENSMUSG00000087691 | Gm15674 | predicted gene 15674 | -1.16 |
| ENSMUSG00000020009 | Ifngr1 | interferon gamma receptor 1 | -1.17 |
| ENSMUSG00000006651 | Aplp1 | amyloid beta (A4) precursor-like protein 1 | -1.17 |
| ENSMUSG00000063931 | Pepd | peptidase D | -1.17 |
| ENSMUSG00000042659 | Arrdc4 | arrestin domain containing 4 | -1.17 |
| ENSMUSG00000045314 | Sowahb | sosondowah ankyrin repeat domain family member B | -1.17 |
| ENSMUSG00000047409 | Ctdspl | CTD (carboxy-terminal domain, RNA polymerase II, polypeptide A) small phosphatase-like | -1.17 |
| ENSMUSG00000075420 | Smim6 | small integral membrane protein 6 | -1.17 |
| ENSMUSG00000047557 | Lxn | latexin | -1.17 |
| ENSMUSG00000057315 | Arhgap24 | Rho GTPase activating protein 24 | -1.17 |
| ENSMUSG00000040740 | Slc25a34 | solute carrier family 25, member 34 | -1.17 |
| ENSMUSG00000035852 | Misp | mitotic spindle positioning | -1.18 |
| ENSMUSG00000026090 | 2010300C02Rik | RIKEN cDNA 2010300C02 gene | -1.18 |
| ENSMUSG00000032306 | Mpi | mannose phosphate isomerase | -1.18 |
| ENSMUSG00000025190 | Got1 | glutamic-oxaloacetic transaminase 1, soluble | -1.18 |
| ENSMUSG00000033083 | Tbc1d4 | TBC1 domain family, member 4 | -1.18 |
| ENSMUSG00000017868 | Sgk2 | serum/glucocorticoid regulated kinase 2 | -1.18 |
| ENSMUSG00000001901 | Kcnh6 | potassium voltage-gated channel, subfamily H (eag-related), member 6 | -1.18 |
| ENSMUSG00000020841 | Cpd | carboxypeptidase D | -1.19 |
| ENSMUSG00000015846 | Rxra | retinoid X receptor alpha | -1.19 |
| ENSMUSG00000034248 | Slc25a37 | solute carrier family 25, member 37 | -1.19 |
| ENSMUSG00000076435 | Acsf2 | acyl-CoA synthetase family member 2 | -1.19 |
| ENSMUSG00000007833 | Aldh16a1 | aldehyde dehydrogenase 16 family, member A1 | -1.19 |
| ENSMUSG00000033917 | Gde1 | glycerophosphodiester phosphodiesterase 1 | -1.19 |
| ENSMUSG00000031378 | Abcd1 | ATP-binding cassette, sub-family D (ALD), member 1 | -1.19 |
| ENSMUSG00000043445 | Pgp | phosphoglycolate phosphatase | -1.19 |
| ENSMUSG00000067297 | Ifit1bl2 | interferon induced protein with tetratricopeptide repeats 1B like 2 | -1.19 |
| ENSMUSG00000029618 | Ocm | oncomodulin | -1.19 |
| ENSMUSG00000079174 | Gm3054 | predicted gene 3054 | -1.19 |
| ENSMUSG00000028399 | Ptprd | protein tyrosine phosphatase, receptor type, D | -1.20 |
| ENSMUSG00000027957 | Slc35a3 | solute carrier family 35 (UDP-N-acetylglucosamine (UDP-GlcNAc) transporter), member 3 | -1.20 |
| ENSMUSG00000084883 | Ccdc85c | coiled-coil domain containing 85C | -1.20 |
| ENSMUSG00000005615 | Pcyt1a | phosphate cytidylyltransferase 1, choline, alpha isoform | -1.20 |
| ENSMUSG00000025477 | Inpp5a | inositol polyphosphate-5-phosphatase A | -1.20 |
| ENSMUSG00000031156 | Slc35a2 | solute carrier family 35 (UDP-galactose transporter), member A2 | -1.20 |
| ENSMUSG00000016495 | Plgrkt | plasminogen receptor, C-terminal lysine transmembrane protein | -1.20 |
| ENSMUSG00000028488 | Sh3gl2 | SH3-domain GRB2-like 2 | -1.20 |
| ENSMUSG00000029471 | Camkk2 | calcium/calmodulin-dependent protein kinase kinase 2, beta | -1.20 |
| ENSMUSG00000056185 | Snx32 | sorting nexin 32 | -1.20 |

| | | | |
|---|---|---|---|
| ENSMUSG00000010122 | Slc47a1 | solute carrier family 47, member 1 | -1.20 |
| ENSMUSG00000025762 | Larp1b | La ribonucleoprotein domain family, member 1B | -1.21 |
| ENSMUSG00000024036 | Slc37a1 | solute carrier family 37 (glycerol-3-phosphate transporter), member 1 | -1.21 |
| ENSMUSG00000032802 | Srxn1 | sulfiredoxin 1 homolog (S. cerevisiae) | -1.21 |
| ENSMUSG00000026062 | Slc9a2 | solute carrier family 9 (sodium/hydrogen exchanger), member 2 | -1.21 |
| ENSMUSG00000033885 | Pxk | PX domain containing serine/threonine kinase | -1.21 |
| ENSMUSG00000054499 | Dedd2 | death effector domain-containing DNA binding protein 2 | -1.21 |
| ENSMUSG00000053898 | Ech1 | enoyl coenzyme A hydratase 1, peroxisomal | -1.21 |
| ENSMUSG00000026922 | Agpat2 | 1-acylglycerol-3-phosphate O-acyltransferase 2 (lysophosphatidic acid acyltransferase, beta) | -1.21 |
| ENSMUSG00000026102 | Inpp1 | inositol polyphosphate-1-phosphatase | -1.21 |
| ENSMUSG00000102953 | Gm37019 | predicted gene, 37019 | -1.21 |
| ENSMUSG00000040276 | Pacsin1 | protein kinase C and casein kinase substrate in neurons 1 | -1.21 |
| ENSMUSG00000003948 | Mmd | monocyte to macrophage differentiation-associated | -1.21 |
| ENSMUSG00000025359 | Pmel | premelanosome protein | -1.21 |
| ENSMUSG00000026879 | Gsn | gelsolin | -1.22 |
| ENSMUSG00000009828 | Ick | intestinal cell kinase | -1.22 |
| ENSMUSG00000031827 | Cotl1 | coactosin-like 1 (Dictyostelium) | -1.22 |
| ENSMUSG00000028555 | Ttc39a | tetratricopeptide repeat domain 39A | -1.22 |
| ENSMUSG00000032661 | Oas3 | 2'-5' oligoadenylate synthetase 3 | -1.22 |
| ENSMUSG00000040147 | Maob | monoamine oxidase B | -1.22 |
| ENSMUSG00000087535 | Zmiz1os1 | Zmiz1 opposite strand 1 | -1.22 |
| ENSMUSG00000023861 | Mpc1 | mitochondrial pyruvate carrier 1 | -1.22 |
| ENSMUSG00000067813 | Xkr9 | X-linked Kx blood group related 9 | -1.22 |
| ENSMUSG00000034271 | Jdp2 | Jun dimerization protein 2 | -1.22 |
| ENSMUSG00000034330 | Plcg2 | phospholipase C, gamma 2 | -1.23 |
| ENSMUSG00000000538 | Tom1l2 | target of myb1-like 2 (chicken) | -1.23 |
| ENSMUSG00000038894 | Irs2 | insulin receptor substrate 2 | -1.23 |
| ENSMUSG00000039197 | Adk | adenosine kinase | -1.23 |
| ENSMUSG00000034522 | Zfp395 | zinc finger protein 395 | -1.23 |
| ENSMUSG00000042198 | Chchd7 | coiled-coil-helix-coiled-coil-helix domain containing 7 | -1.23 |
| ENSMUSG00000042097 | Zfp239 | zinc finger protein 239 | -1.23 |
| ENSMUSG00000024610 | Cd74 | CD74 antigen (invariant polypeptide of major histocompatibility complex, class II antigen-associated | -1.24 |
| ENSMUSG00000026303 | Mlph | melanophilin | -1.24 |
| ENSMUSG00000020111 | Micu1 | mitochondrial calcium uptake 1 | -1.24 |
| ENSMUSG00000036452 | Arhgap26 | Rho GTPase activating protein 26 | -1.24 |
| ENSMUSG00000029561 | Oasl2 | 2'-5' oligoadenylate synthetase-like 2 | -1.24 |
| ENSMUSG00000059323 | Tonsl | tonsoku-like, DNA repair protein | -1.24 |
| ENSMUSG00000031337 | Mtm1 | X-linked myotubular myopathy gene 1 | -1.24 |
| ENSMUSG00000041143 | Tmco4 | transmembrane and coiled-coil domains 4 | -1.24 |
| ENSMUSG00000031021 | Tmem9b | TMEM9 domain family, member B | -1.24 |
| ENSMUSG00000035967 | Ints6l | integrator complex subunit 6 like | -1.24 |
| ENSMUSG00000064210 | Ano6 | anoctamin 6 | -1.25 |
| ENSMUSG00000038260 | Trpm4 | transient receptor potential cation channel, subfamily M, member 4 | -1.25 |
| ENSMUSG00000020099 | Unc5b | unc-5 netrin receptor B | -1.25 |
| ENSMUSG00000014418 | Hps5 | HPS5, biogenesis of lysosomal organelles complex 2 subunit 2 | -1.25 |
| ENSMUSG00000035722 | Abca7 | ATP-binding cassette, sub-family A (ABC1), member 7 | -1.25 |
| ENSMUSG00000042228 | Lyn | LYN proto-oncogene, Src family tyrosine kinase | -1.25 |
| ENSMUSG00000028293 | Slc35a1 | solute carrier family 35 (CMP-sialic acid transporter), member 1 | -1.25 |
| ENSMUSG00000063838 | Cdc42ep5 | CDC42 effector protein (Rho GTPase binding) 5 | -1.25 |
| ENSMUSG00000026259 | Ngef | neuronal guanine nucleotide exchange factor | -1.25 |
| ENSMUSG00000031570 | Plpp5 | phospholipid phosphatase 5 | -1.25 |
| ENSMUSG00000110277 | Gm45871 | predicted gene 45871 | -1.25 |
| ENSMUSG00000020377 | Ltc4s | leukotriene C4 synthase | -1.25 |
| ENSMUSG00000061601 | Pclo | piccolo (presynaptic cytomatrix protein) | -1.25 |
| ENSMUSG00000018166 | Erbb3 | erb-b2 receptor tyrosine kinase 3 | -1.26 |
| ENSMUSG00000036918 | Ttc7 | tetratricopeptide repeat domain 7 | -1.26 |
| ENSMUSG00000037852 | Cpe | carboxypeptidase E | -1.26 |
| ENSMUSG00000028229 | Rmdn1 | regulator of microtubule dynamics 1 | -1.26 |
| ENSMUSG00000103983 | Gm20045 | predicted gene, 20045 | -1.26 |
| ENSMUSG00000020733 | Slc9a3r1 | solute carrier family 9 (sodium/hydrogen exchanger), member 3 regulator 1 | -1.27 |
| ENSMUSG00000022186 | Oxct1 | 3-oxoacid CoA transferase 1 | -1.27 |
| ENSMUSG00000028064 | Sema4a | sema domain, immunoglobulin domain (Ig), transmembrane domain (TM) and short cytoplasmic do | -1.27 |
| ENSMUSG00000025764 | Jade1 | jade family PHD finger 1 | -1.27 |
| ENSMUSG00000017801 | Mlx | MAX-like protein X | -1.27 |
| ENSMUSG00000026304 | Rab17 | RAB17, member RAS oncogene family | -1.27 |
| ENSMUSG00000035186 | Ubd | ubiquitin D | -1.27 |
| ENSMUSG00000026976 | Pax8 | paired box 8 | -1.27 |
| ENSMUSG00000024975 | Pdcd4 | programmed cell death 4 | -1.28 |
| ENSMUSG00000038342 | Mlxip | MLX interacting protein | -1.28 |
| ENSMUSG00000031955 | Bcar1 | breast cancer anti-estrogen resistance 1 | -1.28 |
| ENSMUSG00000038365 | Fbxo25 | F-box protein 25 | -1.28 |
| ENSMUSG00000056313 | Tcim | transcriptional and immune response regulator | -1.28 |
| ENSMUSG00000006395 | Hyi | hydroxypyruvate isomerase (putative) | -1.28 |
| ENSMUSG00000044072 | Eml6 | echinoderm microtubule associated protein like 6 | -1.28 |
| ENSMUSG00000003585 | Sec14l2 | SEC14-like lipid binding 2 | -1.28 |
| ENSMUSG00000104940 | Gm42908 | predicted gene 42908 | -1.28 |

| | | | |
|---|---|---|---|
| ENSMUSG00000018554 | Ybx2 | Y box protein 2 | -1.28 |
| ENSMUSG00000031845 | Bco1 | beta-carotene oxygenase 1 | -1.28 |
| ENSMUSG00000100183 | Gm28512 | predicted gene 28512 | -1.28 |
| ENSMUSG00000028479 | Gne | glucosamine (UDP-N-acetyl)-2-epimerase/N-acetylmannosamine kinase | -1.29 |
| ENSMUSG00000022844 | Pdia5 | protein disulfide isomerase associated 5 | -1.29 |
| ENSMUSG00000055116 | Arntl | aryl hydrocarbon receptor nuclear translocator-like | -1.29 |
| ENSMUSG00000038473 | Nos1ap | nitric oxide synthase 1 (neuronal) adaptor protein | -1.29 |
| ENSMUSG00000048970 | C1galt1c1 | C1GALT1-specific chaperone 1 | -1.29 |
| ENSMUSG00000021665 | Hexb | hexosaminidase B | -1.29 |
| ENSMUSG00000042747 | Krtcap2 | keratinocyte associated protein 2 | -1.29 |
| ENSMUSG00000086784 | Isoc2a | isochorismatase domain containing 2a | -1.29 |
| ENSMUSG00000026204 | Ptprn | protein tyrosine phosphatase, receptor type, N | -1.29 |
| ENSMUSG00000005089 | Slc1a2 | solute carrier family 1 (glial high affinity glutamate transporter), member 2 | -1.29 |
| ENSMUSG00000032601 | Prkar2a | protein kinase, cAMP dependent regulatory, type II alpha | -1.30 |
| ENSMUSG00000017057 | Il13ra1 | interleukin 13 receptor, alpha 1 | -1.30 |
| ENSMUSG00000030987 | Stim1 | stromal interaction molecule 1 | -1.30 |
| ENSMUSG00000028862 | Map3k6 | mitogen-activated protein kinase kinase kinase 6 | -1.30 |
| ENSMUSG00000054342 | Kcnn4 | potassium intermediate/small conductance calcium-activated channel, subfamily N, member 4 | -1.30 |
| ENSMUSG00000054752 | Fsd1l | fibronectin type III and SPRY domain containing 1-like | -1.30 |
| ENSMUSG00000046949 | Nqo2 | N-ribosyldihydronicotinamide quinone reductase 2 | -1.30 |
| ENSMUSG00000106205 | C230096K16Rik | RIKEN cDNA C230096K16 gene | -1.30 |
| ENSMUSG00000001227 | Sema6b | sema domain, transmembrane domain (TM), and cytoplasmic domain, (semaphorin) 6B | -1.30 |
| ENSMUSG00000013418 | B4galnt2 | beta-1,4-N-acetyl-galactosaminyl transferase 2 | -1.31 |
| ENSMUSG00000027984 | Hadh | hydroxyacyl-Coenzyme A dehydrogenase | -1.31 |
| ENSMUSG00000049493 | Pls1 | plastin 1 (I-isoform) | -1.31 |
| ENSMUSG00000005125 | Ndrg1 | N-myc downstream regulated gene 1 | -1.31 |
| ENSMUSG00000024937 | Ehbp1l1 | EH domain binding protein 1-like 1 | -1.31 |
| ENSMUSG00000041827 | Oasl1 | 2'-5' oligoadenylate synthetase-like 1 | -1.31 |
| ENSMUSG00000034591 | Slc41a2 | solute carrier family 41, member 2 | -1.31 |
| ENSMUSG00000026880 | Stom | stomatin | -1.31 |
| ENSMUSG00000041120 | Nbl1 | neuroblastoma, suppression of tumorigenicity 1 | -1.31 |
| ENSMUSG00000039304 | Tnfsf10 | tumor necrosis factor (ligand) superfamily, member 10 | -1.31 |
| ENSMUSG00000020447 | Npc1l1 | NPC1 like intracellular cholesterol transporter 1 | -1.31 |
| ENSMUSG00000064177 | Ghrl | ghrelin | -1.31 |
| ENSMUSG00000019122 | Ccl9 | chemokine (C-C motif) ligand 9 | -1.31 |
| ENSMUSG00000033577 | Myo6 | myosin VI | -1.32 |
| ENSMUSG00000060012 | Kif13b | kinesin family member 13B | -1.32 |
| ENSMUSG00000033684 | Qsox1 | quiescin Q6 sulfhydryl oxidase 1 | -1.32 |
| ENSMUSG00000024431 | Nr3c1 | nuclear receptor subfamily 3, group C, member 1 | -1.32 |
| ENSMUSG00000061845 | Defa35 | defensin, alpha, 35 | -1.32 |
| ENSMUSG00000026749 | Nek6 | NIMA (never in mitosis gene a)-related expressed kinase 6 | -1.32 |
| ENSMUSG00000047604 | Frat2 | frequently rearranged in advanced T cell lymphomas 2 | -1.32 |
| ENSMUSG00000022683 | Pla2g10 | phospholipase A2, group X | -1.32 |
| ENSMUSG00000024395 | Lims2 | LIM and senescent cell antigen like domains 2 | -1.32 |
| ENSMUSG00000070574 | 2310016G11Rik | RIKEN cDNA 2310016G11 gene | -1.32 |
| ENSMUSG00000052738 | Suclg1 | succinate-CoA ligase, GDP-forming, alpha subunit | -1.33 |
| ENSMUSG00000005580 | Adcy9 | adenylate cyclase 9 | -1.33 |
| ENSMUSG00000031887 | Tradd | TNFRSF1A-associated via death domain | -1.33 |
| ENSMUSG00000030761 | Myo7a | myosin VIIA | -1.33 |
| ENSMUSG00000029859 | Epha1 | Eph receptor A1 | -1.33 |
| ENSMUSG00000020609 | Apob | apolipoprotein B | -1.33 |
| ENSMUSG00000093880 | Tmem181c-ps | transmembrane protein 181C, pseudogene | -1.33 |
| ENSMUSG00000035441 | Myo1d | myosin ID | -1.34 |
| ENSMUSG00000039611 | Tmem246 | transmembrane protein 246 | -1.34 |
| ENSMUSG00000042797 | Aqp11 | aquaporin 11 | -1.34 |
| ENSMUSG00000079429 | Mroh2a | maestro heat-like repeat family member 2A | -1.34 |
| ENSMUSG00000042737 | Dpm3 | dolichyl-phosphate mannosyltransferase polypeptide 3 | -1.34 |
| ENSMUSG00000097006 | 9530082P21Rik | RIKEN cDNA 9530082P21 gene | -1.34 |
| ENSMUSG00000022946 | Dopey2 | dopey family member 2 | -1.35 |
| ENSMUSG00000042808 | Gpx2 | glutathione peroxidase 2 | -1.35 |
| ENSMUSG00000039234 | Sec24d | Sec24 related gene family, member D (S. cerevisiae) | -1.35 |
| ENSMUSG00000038244 | Mical2 | microtubule associated monooxygenase, calponin and LIM domain containing 2 | -1.35 |
| ENSMUSG00000020826 | Nos2 | nitric oxide synthase 2, inducible | -1.35 |
| ENSMUSG00000025429 | Pstpip2 | proline-serine-threonine phosphatase-interacting protein 2 | -1.35 |
| ENSMUSG00000038967 | Pdk2 | pyruvate dehydrogenase kinase, isoenzyme 2 | -1.35 |
| ENSMUSG00000022750 | Klhl22 | kelch-like 22 | -1.35 |
| ENSMUSG00000037709 | Fam13a | family with sequence similarity 13, member A | -1.35 |
| ENSMUSG00000074882 | Cyp2c68 | cytochrome P450, family 2, subfamily c, polypeptide 68 | -1.35 |
| ENSMUSG00000032766 | Gng11 | guanine nucleotide binding protein (G protein), gamma 11 | -1.35 |
| ENSMUSG00000028719 | Cmpk1 | cytidine monophosphate (UMP-CMP) kinase 1 | -1.36 |
| ENSMUSG00000033318 | Gstt2 | glutathione S-transferase, theta 2 | -1.36 |
| ENSMUSG00000030935 | Acsm3 | acyl-CoA synthetase medium-chain family member 3 | -1.36 |
| ENSMUSG00000074218 | Cox7a1 | cytochrome c oxidase subunit 7A1 | -1.36 |
| ENSMUSG00000020827 | Mink1 | misshapen-like kinase 1 (zebrafish) | -1.37 |
| ENSMUSG00000009647 | Mcu | mitochondrial calcium uniporter | -1.37 |
| ENSMUSG00000028334 | Nans | N-acetylneuraminic acid synthase (sialic acid synthase) | -1.37 |

| | | | |
|---|---|---|---|
| ENSMUSG00000020752 | Recql5 | RecQ protein-like 5 | -1.37 |
| ENSMUSG00000059316 | Slc27a4 | solute carrier family 27 (fatty acid transporter), member 4 | -1.37 |
| ENSMUSG00000026896 | Ifih1 | interferon induced with helicase C domain 1 | -1.37 |
| ENSMUSG00000047496 | Rnf152 | ring finger protein 152 | -1.37 |
| ENSMUSG00000027412 | Lpin3 | lipin 3 | -1.37 |
| ENSMUSG00000004815 | Dgkq | diacylglycerol kinase, theta | -1.37 |
| ENSMUSG00000072115 | Ang | angiogenin, ribonuclease, RNase A family, 5 | -1.37 |
| ENSMUSG00000029188 | Slc34a2 | solute carrier family 34 (sodium phosphate), member 2 | -1.37 |
| ENSMUSG00000029802 | Abcg2 | ATP binding cassette subfamily G member 2 (Junior blood group) | -1.37 |
| ENSMUSG00000051339 | 2900026A02Rik | RIKEN cDNA 2900026A02 gene | -1.38 |
| ENSMUSG00000039131 | Gipc2 | GIPC PDZ domain containing family, member 2 | -1.38 |
| ENSMUSG00000033416 | Gucd1 | guanylyl cyclase domain containing 1 | -1.38 |
| ENSMUSG00000036499 | Eea1 | early endosome antigen 1 | -1.38 |
| ENSMUSG00000024867 | Pip5k1b | phosphatidylinositol-4-phosphate 5-kinase, type 1 beta | -1.38 |
| ENSMUSG00000031823 | Zdhhc7 | zinc finger, DHHC domain containing 7 | -1.38 |
| ENSMUSG00000034371 | Tkfc | triokinase, FMN cyclase | -1.38 |
| ENSMUSG00000026961 | Lrrc26 | leucine rich repeat containing 26 | -1.38 |
| ENSMUSG00000001670 | Tat | tyrosine aminotransferase | -1.38 |
| ENSMUSG00000067199 | Frat1 | frequently rearranged in advanced T cell lymphomas | -1.38 |
| ENSMUSG00000035769 | Xylb | xylulokinase homolog (H. influenzae) | -1.38 |
| ENSMUSG00000102336 | Gm37233 | predicted gene, 37233 | -1.38 |
| ENSMUSG00000106659 | Gm42161 | predicted gene, 42161 | -1.38 |
| ENSMUSG00000044231 | Nhlrc1 | NHL repeat containing 1 | -1.38 |
| ENSMUSG00000045019 | Acer1 | alkaline ceramidase 1 | -1.38 |
| ENSMUSG00000061751 | Kalrn | kalirin, RhoGEF kinase | -1.39 |
| ENSMUSG00000043079 | Synpo | synaptopodin | -1.39 |
| ENSMUSG00000050931 | Sgms2 | sphingomyelin synthase 2 | -1.39 |
| ENSMUSG00000022105 | Rb1 | RB transcriptional corepressor 1 | -1.39 |
| ENSMUSG00000029727 | Cyp3a13 | cytochrome P450, family 3, subfamily a, polypeptide 13 | -1.39 |
| ENSMUSG00000021185 | Dglucy | D-glutamate cyclase | -1.39 |
| ENSMUSG00000036863 | Syde2 | synapse defective 1, Rho GTPase, homolog 2 (C. elegans) | -1.39 |
| ENSMUSG00000029780 | Nt5c3 | 5'-nucleotidase, cytosolic III | -1.39 |
| ENSMUSG00000108291 | Gm44292 | predicted gene, 44292 | -1.39 |
| ENSMUSG00000027457 | Snph | syntaphilin | -1.39 |
| ENSMUSG00000029452 | Tmem116 | transmembrane protein 116 | -1.39 |
| ENSMUSG00000025068 | Gsto1 | glutathione S-transferase omega 1 | -1.40 |
| ENSMUSG00000003746 | Man1a | mannosidase 1, alpha | -1.40 |
| ENSMUSG00000074443 | Defa22 | defensin, alpha, 22 | -1.40 |
| ENSMUSG00000046519 | Golph3l | golgi phosphoprotein 3-like | -1.40 |
| ENSMUSG00000067219 | Nipal1 | NIPA-like domain containing 1 | -1.40 |
| ENSMUSG00000024776 | Stambpl1 | STAM binding protein like 1 | -1.40 |
| ENSMUSG00000036106 | Prr5 | proline rich 5 (renal) | -1.40 |
| ENSMUSG00000039699 | Batf2 | basic leucine zipper transcription factor, ATF-like 2 | -1.40 |
| ENSMUSG00000028518 | Prkaa2 | protein kinase, AMP-activated, alpha 2 catalytic subunit | -1.41 |
| ENSMUSG00000024074 | Crim1 | cysteine rich transmembrane BMP regulator 1 (chordin like) | -1.41 |
| ENSMUSG00000025993 | Slc40a1 | solute carrier family 40 (iron-regulated transporter), member 1 | -1.41 |
| ENSMUSG00000038240 | Pdss2 | prenyl (solanesyl) diphosphate synthase, subunit 2 | -1.41 |
| ENSMUSG00000022040 | Ephx2 | epoxide hydrolase 2, cytoplasmic | -1.41 |
| ENSMUSG00000030428 | Ttyh1 | tweety family member 1 | -1.41 |
| ENSMUSG00000032737 | Inppl1 | inositol polyphosphate phosphatase-like 1 | -1.42 |
| ENSMUSG00000024955 | Esrra | estrogen related receptor, alpha | -1.42 |
| ENSMUSG00000003134 | Tbc1d8 | TBC1 domain family, member 8 | -1.42 |
| ENSMUSG00000036136 | Fam110c | family with sequence similarity 110, member C | -1.42 |
| ENSMUSG00000058586 | Serhl | serine hydrolase-like | -1.42 |
| ENSMUSG00000026018 | Ica1l | islet cell autoantigen 1-like | -1.42 |
| ENSMUSG00000024644 | Cndp2 | CNDP dipeptidase 2 (metallopeptidase M20 family) | -1.43 |
| ENSMUSG00000047153 | Khnyn | KH and NYN domain containing | -1.43 |
| ENSMUSG00000022330 | Osr2 | odd-skipped related 2 | -1.43 |
| ENSMUSG00000028121 | Bcar3 | breast cancer anti-estrogen resistance 3 | -1.43 |
| ENSMUSG00000037921 | Ddx60 | DEAD (Asp-Glu-Ala-Asp) box polypeptide 60 | -1.43 |
| ENSMUSG00000022687 | Boc | biregional cell adhesion molecule-related/down-regulated by oncogenes (Cdon) binding protein | -1.43 |
| ENSMUSG00000040350 | Trim7 | tripartite motif-containing 7 | -1.43 |
| ENSMUSG00000104737 | Gm42937 | predicted gene 42937 | -1.43 |
| ENSMUSG00000043155 | Hpdl | 4-hydroxyphenylpyruvate dioxygenase-like | -1.43 |
| ENSMUSG00000106191 | 4930557B06Rik | RIKEN cDNA 4930557B06 gene | -1.43 |
| ENSMUSG00000068341 | Reg3d | regenerating islet-derived 3 delta | -1.43 |
| ENSMUSG00000001156 | Mxd1 | MAX dimerization protein 1 | -1.44 |
| ENSMUSG00000015533 | Itga2 | integrin alpha 2 | -1.44 |
| ENSMUSG00000034066 | Farp2 | FERM, RhoGEF and pleckstrin domain protein 2 | -1.44 |
| ENSMUSG00000026360 | Rgs2 | regulator of G-protein signaling 2 | -1.44 |
| ENSMUSG00000013539 | Tango2 | transport and golgi organization 2 | -1.44 |
| ENSMUSG00000086894 | Gm15708 | predicted gene 15708 | -1.44 |
| ENSMUSG00000046861 | Hectd3 | HECT domain E3 ubiquitin protein ligase 3 | -1.45 |
| ENSMUSG00000031090 | Nadsyn1 | NAD synthetase 1 | -1.45 |
| ENSMUSG00000021708 | Rasgrf2 | RAS protein-specific guanine nucleotide-releasing factor 2 | -1.45 |
| ENSMUSG00000067931 | Zfp948 | zinc finger protein 948 | -1.45 |

| | | | |
|---|---|---|---|
| ENSMUSG00000004610 | Etfb | electron transferring flavoprotein, beta polypeptide | -1.45 |
| ENSMUSG00000043629 | 1700019D03Rik | RIKEN cDNA 1700019D03 gene | -1.45 |
| ENSMUSG00000056035 | Cyp3a11 | cytochrome P450, family 3, subfamily a, polypeptide 11 | -1.45 |
| ENSMUSG00000085088 | 4931413K12Rik | RIKEN cDNA 4931413K12 gene | -1.45 |
| ENSMUSG00000053819 | Camk2d | calcium/calmodulin-dependent protein kinase II, delta | -1.46 |
| ENSMUSG00000021556 | Golm1 | golgi membrane protein 1 | -1.46 |
| ENSMUSG00000022949 | Clic6 | chloride intracellular channel 6 | -1.46 |
| ENSMUSG00000039637 | Coro7 | coronin 7 | -1.46 |
| ENSMUSG00000028538 | St3gal3 | ST3 beta-galactoside alpha-2,3-sialyltransferase 3 | -1.46 |
| ENSMUSG00000097059 | Fam120aos | family with sequence similarity 120A, opposite strand | -1.46 |
| ENSMUSG00000023206 | Il15ra | interleukin 15 receptor, alpha chain | -1.46 |
| ENSMUSG00000054150 | Syne3 | spectrin repeat containing, nuclear envelope family member 3 | -1.46 |
| ENSMUSG00000023057 | Fabp2 | fatty acid binding protein 2, intestinal | -1.46 |
| ENSMUSG00000030681 | Mvp | major vault protein | -1.47 |
| ENSMUSG00000003546 | Klc4 | kinesin light chain 4 | -1.47 |
| ENSMUSG00000025498 | Irf7 | interferon regulatory factor 7 | -1.47 |
| ENSMUSG00000035473 | Galm | galactose mutarotase | -1.47 |
| ENSMUSG00000028480 | Glipr2 | GLI pathogenesis-related 2 | -1.47 |
| ENSMUSG00000025198 | Erlin1 | ER lipid raft associated 1 | -1.47 |
| ENSMUSG00000014778 | Fhod1 | formin homology 2 domain containing 1 | -1.47 |
| ENSMUSG00000042333 | Tnfrsf14 | tumor necrosis factor receptor superfamily, member 14 (herpesvirus entry mediator) | -1.47 |
| ENSMUSG00000028158 | Mttp | microsomal triglyceride transfer protein | -1.47 |
| ENSMUSG00000108526 | Gm45828 | predicted gene 45828 | -1.47 |
| ENSMUSG00000025978 | Rftn2 | raftlin family member 2 | -1.47 |
| ENSMUSG00000029163 | Emilin1 | elastin microfibril interfacer 1 | -1.47 |
| ENSMUSG00000045817 | Zfp36l2 | zinc finger protein 36, C3H type-like 2 | -1.48 |
| ENSMUSG00000028943 | Espn | espin | -1.48 |
| ENSMUSG00000039166 | Akap7 | A kinase (PRKA) anchor protein 7 | -1.48 |
| ENSMUSG00000032842 | Abcc10 | ATP-binding cassette, sub-family C (CFTR/MRP), member 10 | -1.48 |
| ENSMUSG00000068874 | Selenbp1 | selenium binding protein 1 | -1.48 |
| ENSMUSG00000015405 | Ace2 | angiotensin I converting enzyme (peptidyl-dipeptidase A) 2 | -1.48 |
| ENSMUSG00000042367 | Gjb3 | gap junction protein, beta 3 | -1.48 |
| ENSMUSG00000052135 | Foxo6 | forkhead box O6 | -1.48 |
| ENSMUSG00000112346 | Gm48768 | predicted gene, 48768 | -1.48 |
| ENSMUSG00000039278 | Pcsk1n | proprotein convertase subtilisin/kexin type 1 inhibitor | -1.48 |
| ENSMUSG00000050395 | Tnfsf15 | tumor necrosis factor (ligand) superfamily, member 15 | -1.48 |
| ENSMUSG00000054733 | Msra | methionine sulfoxide reductase A | -1.49 |
| ENSMUSG00000037455 | Slc18b1 | solute carrier family 18, subfamily B, member 1 | -1.49 |
| ENSMUSG00000068417 | Pnp2 | purine-nucleoside phosphorylase 2 | -1.49 |
| ENSMUSG00000021280 | Exoc3l4 | exocyst complex component 3-like 4 | -1.49 |
| ENSMUSG00000063652 | Slc22a21 | solute carrier family 22 (organic cation transporter), member 21 | -1.49 |
| ENSMUSG00000072571 | Tmem253 | transmembrane protein 253 | -1.49 |
| ENSMUSG00000054200 | Ffar4 | free fatty acid receptor 4 | -1.49 |
| ENSMUSG00000037169 | Mycn | v-myc avian myelocytomatosis viral related oncogene, neuroblastoma derived | -1.49 |
| ENSMUSG00000087651 | 1500009L16Rik | RIKEN cDNA 1500009L16 gene | -1.49 |
| ENSMUSG00000000915 | Hip1r | huntingtin interacting protein 1 related | -1.50 |
| ENSMUSG00000029646 | Cdx2 | caudal type homeobox 2 | -1.50 |
| ENSMUSG00000055675 | Kbtbd11 | kelch repeat and BTB (POZ) domain containing 11 | -1.50 |
| ENSMUSG00000021587 | Pcsk1 | proprotein convertase subtilisin/kexin type 1 | -1.50 |
| ENSMUSG00000030545 | Pex11a | peroxisomal biogenesis factor 11 alpha | -1.50 |
| ENSMUSG00000104693 | Gm42941 | predicted gene 42941 | -1.50 |
| ENSMUSG00000017718 | Afmid | arylformamidase | -1.50 |
| ENSMUSG00000041617 | Ccdc74a | coiled-coil domain containing 74A | -1.50 |
| ENSMUSG00000086189 | Gm15462 | predicted gene 15462 | -1.50 |
| ENSMUSG00000024066 | Xdh | xanthine dehydrogenase | -1.51 |
| ENSMUSG00000024712 | Rfk | riboflavin kinase | -1.51 |
| ENSMUSG00000031482 | Slc25a15 | solute carrier family 25 (mitochondrial carrier ornithine transporter), member 15 | -1.51 |
| ENSMUSG00000041237 | Pklr | pyruvate kinase liver and red blood cell | -1.51 |
| ENSMUSG00000024727 | Trpm6 | transient receptor potential cation channel, subfamily M, member 6 | -1.51 |
| ENSMUSG00000042073 | Abhd14b | abhydrolase domain containing 14b | -1.51 |
| ENSMUSG00000098678 | Mroh6 | maestro heat-like repeat family member 6 | -1.51 |
| ENSMUSG00000039653 | Baat | bile acid-Coenzyme A: amino acid N-acyltransferase | -1.51 |
| ENSMUSG00000022836 | Mylk | myosin, light polypeptide kinase | -1.52 |
| ENSMUSG00000061666 | Gdpd1 | glycerophosphodiester phosphodiesterase domain containing 1 | -1.52 |
| ENSMUSG00000020681 | Ace | angiotensin I converting enzyme (peptidyl-dipeptidase A) 1 | -1.52 |
| ENSMUSG00000033826 | Dnah8 | dynein, axonemal, heavy chain 8 | -1.52 |
| ENSMUSG00000021286 | Zfyve21 | zinc finger, FYVE domain containing 21 | -1.52 |
| ENSMUSG00000028795 | Ccdc28b | coiled coil domain containing 28B | -1.52 |
| ENSMUSG00000032380 | Dapk2 | death-associated protein kinase 2 | -1.52 |
| ENSMUSG00000023393 | Slc17a9 | solute carrier family 17, member 9 | -1.52 |
| ENSMUSG00000038058 | Nod1 | nucleotide-binding oligomerization domain containing 1 | -1.52 |
| ENSMUSG00000082148 | Gm12266 | predicted gene 12266 | -1.52 |
| ENSMUSG00000087611 | 4930458D05Rik | RIKEN cDNA 4930458D05 gene | -1.52 |
| ENSMUSG00000029102 | Hgfac | hepatocyte growth factor activator | -1.53 |
| ENSMUSG00000020573 | Pik3cg | phosphatidylinositol-4,5-bisphosphate 3-kinase catalytic subunit gamma | -1.53 |
| ENSMUSG00000075010 | AW112010 | expressed sequence AW112010 | -1.53 |

| | | | |
|---|---|---|---|
| ENSMUSG00000073402 | Gm8909 | predicted gene 8909 | -1.53 |
| ENSMUSG00000017453 | Pipox | pipecolic acid oxidase | -1.53 |
| ENSMUSG00000029923 | Rab19 | RAB19, member RAS oncogene family | -1.53 |
| ENSMUSG00000069227 | Gprin1 | G protein-regulated inducer of neurite outgrowth 1 | -1.53 |
| ENSMUSG00000097924 | A730020E08Rik | RIKEN cDNA A730020E08 gene | -1.53 |
| ENSMUSG00000074628 | Tldc2 | TBC/LysM associated domain containing 2 | -1.53 |
| ENSMUSG00000041757 | Plekha6 | pleckstrin homology domain containing, family A member 6 | -1.54 |
| ENSMUSG00000023073 | Slc10a2 | solute carrier family 10, member 2 | -1.54 |
| ENSMUSG00000030155 | Clec2e | C-type lectin domain family 2, member e | -1.54 |
| ENSMUSG00000052102 | Gnpda1 | glucosamine-6-phosphate deaminase 1 | -1.54 |
| ENSMUSG00000029413 | Naaa | N-acylethanolamine acid amidase | -1.54 |
| ENSMUSG00000027765 | P2ry1 | purinergic receptor P2Y, G-protein coupled 1 | -1.54 |
| ENSMUSG00000052271 | Bhlha15 | basic helix-loop-helix family, member a15 | -1.54 |
| ENSMUSG00000051065 | Mb21d2 | Mab-21 domain containing 2 | -1.54 |
| ENSMUSG00000097354 | 2310001H17Rik | RIKEN cDNA 2310001H17 gene | -1.54 |
| ENSMUSG00000080715 | Gm5406 | predicted gene 5406 | -1.54 |
| ENSMUSG00000105586 | Gm43009 | predicted gene 43009 | -1.54 |
| ENSMUSG00000016206 | H2-M3 | histocompatibility 2, M region locus 3 | -1.54 |
| ENSMUSG00000032243 | Itga11 | integrin alpha 11 | -1.54 |
| ENSMUSG00000055197 | Fev | FEV (ETS oncogene family) | -1.54 |
| ENSMUSG00000034781 | Gna11 | guanine nucleotide binding protein, alpha 11 | -1.55 |
| ENSMUSG00000036813 | Entpd8 | ectonucleoside triphosphate diphosphohydrolase 8 | -1.55 |
| ENSMUSG00000104535 | Gm42686 | predicted gene 42686 | -1.55 |
| ENSMUSG00000022711 | Pmm2 | phosphomannomutase 2 | -1.56 |
| ENSMUSG00000024187 | Fam234a | family with sequence similarity 234, member A | -1.56 |
| ENSMUSG00000026687 | Aldh9a1 | aldehyde dehydrogenase 9, subfamily A1 | -1.56 |
| ENSMUSG00000028223 | Decr1 | 2,4-dienoyl CoA reductase 1, mitochondrial | -1.56 |
| ENSMUSG00000027314 | Dll4 | delta like canonical Notch ligand 4 | -1.56 |
| ENSMUSG00000039208 | Metrnl | meteorin, glial cell differentiation regulator-like | -1.56 |
| ENSMUSG00000044080 | S100a1 | S100 calcium binding protein A1 | -1.56 |
| ENSMUSG00000026189 | Pecr | peroxisomal trans-2-enoyl-CoA reductase | -1.56 |
| ENSMUSG00000074653 | Lrrc31 | leucine rich repeat containing 31 | -1.56 |
| ENSMUSG00000011034 | Slc5a1 | solute carrier family 5 (sodium/glucose cotransporter), member 1 | -1.56 |
| ENSMUSG00000025584 | Pde8a | phosphodiesterase 8A | -1.56 |
| ENSMUSG00000042258 | Isl1 | ISL1 transcription factor, LIM/homeodomain | -1.56 |
| ENSMUSG00000055670 | Zzef1 | zinc finger, ZZ-type with EF hand domain 1 | -1.57 |
| ENSMUSG00000022680 | Pdxdc1 | pyridoxal-dependent decarboxylase domain containing 1 | -1.57 |
| ENSMUSG00000116594 | AC133488.1 | novel protein | -1.57 |
| ENSMUSG00000026322 | Htr4 | 5 hydroxytryptamine (serotonin) receptor 4 | -1.57 |
| ENSMUSG00000070331 | Qrich2 | glutamine rich 2 | -1.57 |
| ENSMUSG00000108850 | Gm44914 | predicted gene 44914 | -1.57 |
| ENSMUSG00000048992 | Prss32 | protease, serine 32 | -1.58 |
| ENSMUSG00000020451 | Limk2 | LIM motif-containing protein kinase 2 | -1.58 |
| ENSMUSG00000024063 | Lbh | limb-bud and heart | -1.58 |
| ENSMUSG00000001348 | Acp5 | acid phosphatase 5, tartrate resistant | -1.58 |
| ENSMUSG00000070563 | Spaca4 | sperm acrosome associated 4 | -1.58 |
| ENSMUSG00000039145 | Camk1d | calcium/calmodulin-dependent protein kinase ID | -1.59 |
| ENSMUSG00000029053 | Prkcz | protein kinase C, zeta | -1.59 |
| ENSMUSG00000020019 | Ntn4 | netrin 4 | -1.59 |
| ENSMUSG00000041372 | B4galnt3 | beta-1,4-N-acetyl-galactosaminyl transferase 3 | -1.59 |
| ENSMUSG00000034320 | Slc26a2 | solute carrier family 26 (sulfate transporter), member 2 | -1.60 |
| ENSMUSG00000054662 | Ano9 | anoctamin 9 | -1.60 |
| ENSMUSG00000007034 | Slc44a4 | solute carrier family 44, member 4 | -1.60 |
| ENSMUSG00000041737 | Tmem45b | transmembrane protein 45b | -1.60 |
| ENSMUSG00000027938 | Creb3l4 | cAMP responsive element binding protein 3-like 4 | -1.60 |
| ENSMUSG00000055866 | Per2 | period circadian clock 2 | -1.60 |
| ENSMUSG00000043333 | Rhbdl2 | rhomboid like 2 | -1.60 |
| ENSMUSG00000071604 | Fam189a2 | family with sequence similarity 189, member A2 | -1.60 |
| ENSMUSG00000036553 | Sh3tc1 | SH3 domain and tetratricopeptide repeats 1 | -1.60 |
| ENSMUSG00000036473 | Tbc1d24 | TBC1 domain family, member 24 | -1.60 |
| ENSMUSG00000027834 | Serpini1 | serine (or cysteine) peptidase inhibitor, clade I, member 1 | -1.60 |
| ENSMUSG00000029082 | Bst1 | bone marrow stromal cell antigen 1 | -1.60 |
| ENSMUSG00000111468 | 5033425B01Rik | RIKEN cDNA 5033425B01 gene | -1.60 |
| ENSMUSG00000013653 | 1810065E05Rik | RIKEN cDNA 1810065E05 gene | -1.60 |
| ENSMUSG00000049037 | Clec4a1 | C-type lectin domain family 4, member a1 | -1.60 |
| ENSMUSG00000063730 | Hsd3b2 | hydroxy-delta-5-steroid dehydrogenase, 3 beta- and steroid delta-isomerase 2 | -1.60 |
| ENSMUSG00000029381 | Shroom3 | shroom family member 3 | -1.61 |
| ENSMUSG00000040584 | Abcb1a | ATP-binding cassette, sub-family B (MDR/TAP), member 1A | -1.61 |
| ENSMUSG00000074445 | Sprr2a3 | small proline-rich protein 2A3 | -1.61 |
| ENSMUSG00000058624 | Gda | guanine deaminase | -1.61 |
| ENSMUSG00000022555 | Dgat1 | diacylglycerol O-acyltransferase 1 | -1.61 |
| ENSMUSG00000064373 | Selenop | selenoprotein P | -1.61 |
| ENSMUSG00000037649 | H2-DMa | histocompatibility 2, class II, locus DMa | -1.61 |
| ENSMUSG00000019762 | Iyd | iodotyrosine deiodinase | -1.61 |
| ENSMUSG00000093580 | Gm20706 | predicted gene 20706 | -1.61 |
| ENSMUSG00000071506 | Tmem139 | transmembrane protein 139 | -1.61 |

| | | | |
|---|---|---|---|
| ENSMUSG00000036533 | Cdc42ep3 | CDC42 effector protein (Rho GTPase binding) 3 | -1.61 |
| ENSMUSG00000009772 | Nuak2 | NUAK family, SNF1-like kinase, 2 | -1.62 |
| ENSMUSG00000072214 | Sept5 | septin 5 | -1.62 |
| ENSMUSG00000027510 | Rbm38 | RNA binding motif protein 38 | -1.62 |
| ENSMUSG00000020774 | Aspa | aspartoacylase | -1.62 |
| ENSMUSG00000022574 | Naprt | nicotinate phosphoribosyltransferase | -1.62 |
| ENSMUSG00000054252 | Fgfr3 | fibroblast growth factor receptor 3 | -1.62 |
| ENSMUSG00000005107 | Slc2a9 | solute carrier family 2 (facilitated glucose transporter), member 9 | -1.62 |
| ENSMUSG00000039747 | Orai2 | ORAI calcium release-activated calcium modulator 2 | -1.62 |
| ENSMUSG00000029314 | Gpat3 | glycerol-3-phosphate acyltransferase 3 | -1.62 |
| ENSMUSG00000045679 | Pqlc3 | PQ loop repeat containing | -1.62 |
| ENSMUSG00000021125 | Arg2 | arginase type II | -1.62 |
| ENSMUSG00000003849 | Nqo1 | NAD(P)H dehydrogenase, quinone 1 | -1.62 |
| ENSMUSG00000078954 | Arhgap8 | Rho GTPase activating protein 8 | -1.62 |
| ENSMUSG00000050762 | Prss27 | protease, serine 27 | -1.62 |
| ENSMUSG00000040600 | Eps8l3 | EPS8-like 3 | -1.63 |
| ENSMUSG00000027695 | Pld1 | phospholipase D1 | -1.63 |
| ENSMUSG00000000958 | Slc7a7 | solute carrier family 7 (cationic amino acid transporter, y+ system), member 7 | -1.63 |
| ENSMUSG00000021509 | Slc25a48 | solute carrier family 25, member 48 | -1.63 |
| ENSMUSG00000029409 | U90926 | cDNA sequence U90926 | -1.63 |
| ENSMUSG00000021337 | Scgn | secretagogin, EF-hand calcium binding protein | -1.63 |
| ENSMUSG00000085944 | 1700003D09Rik | RIKEN cDNA 1700003D09 gene | -1.63 |
| ENSMUSG00000023909 | Paqr4 | progestin and adipoQ receptor family member IV | -1.64 |
| ENSMUSG00000064254 | Ethe1 | ethylmalonic encephalopathy 1 | -1.64 |
| ENSMUSG00000033538 | Casp4 | caspase 4, apoptosis-related cysteine peptidase | -1.64 |
| ENSMUSG00000020847 | Rph3al | rabphilin 3A-like (without C2 domains) | -1.64 |
| ENSMUSG00000038811 | Gngt2 | guanine nucleotide binding protein (G protein), gamma transducing activity polypeptide 2 | -1.64 |
| ENSMUSG00000070802 | Pnmal2 | PNMA-like 2 | -1.64 |
| ENSMUSG00000030909 | Anks4b | ankyrin repeat and sterile alpha motif domain containing 4B | -1.65 |
| ENSMUSG00000102752 | Gm7694 | predicted gene 7694 | -1.65 |
| ENSMUSG00000051379 | Flrt3 | fibronectin leucine rich transmembrane protein 3 | -1.65 |
| ENSMUSG00000029847 | Slc23a4 | solute carrier family 23 member 4 | -1.65 |
| ENSMUSG00000057069 | Ero1lb | ERO1-like beta (S. cerevisiae) | -1.65 |
| ENSMUSG00000025557 | Slc15a1 | solute carrier family 15 (oligopeptide transporter), member 1 | -1.65 |
| ENSMUSG00000104316 | Gm37909 | predicted gene, 37909 | -1.65 |
| ENSMUSG00000085196 | Gm14963 | predicted gene 14963 | -1.65 |
| ENSMUSG00000032500 | Dclk3 | doublecortin-like kinase 3 | -1.65 |
| ENSMUSG00000025357 | Dgka | diacylglycerol kinase, alpha | -1.66 |
| ENSMUSG00000021266 | Wars | tryptophanyl-tRNA synthetase | -1.66 |
| ENSMUSG00000037579 | Kcnh3 | potassium voltage-gated channel, subfamily H (eag-related), member 3 | -1.66 |
| ENSMUSG00000022219 | Cideb | cell death-inducing DNA fragmentation factor, alpha subunit-like effector B | -1.66 |
| ENSMUSG00000115062 | 4930544F09Rik | RIKEN cDNA 4930544F09 gene | -1.66 |
| ENSMUSG00000086187 | Gm12860 | predicted gene 12860 | -1.66 |
| ENSMUSG00000026175 | Vil1 | villin 1 | -1.67 |
| ENSMUSG00000060733 | Ipmk | inositol polyphosphate multikinase | -1.67 |
| ENSMUSG00000040712 | Camta2 | calmodulin binding transcription activator 2 | -1.67 |
| ENSMUSG00000020520 | Galnt10 | polypeptide N-acetylgalactosaminyltransferase 10 | -1.67 |
| ENSMUSG00000027225 | Duoxa2 | dual oxidase maturation factor 2 | -1.67 |
| ENSMUSG00000028519 | Dab1 | disabled 1 | -1.67 |
| ENSMUSG00000057596 | Trim30d | tripartite motif-containing 30D | -1.67 |
| ENSMUSG00000039813 | Tbc1d2 | TBC1 domain family, member 2 | -1.67 |
| ENSMUSG00000064262 | Gimap8 | GTPase, IMAP family member 8 | -1.67 |
| ENSMUSG00000085747 | Slc13a2os | solute carrier family 13 (sodium-dependent dicarboxylate transporter), member 2, opposite strand | -1.67 |
| ENSMUSG00000022861 | Dgkg | diacylglycerol kinase, gamma | -1.67 |
| ENSMUSG00000030499 | Kctd15 | potassium channel tetramerisation domain containing 15 | -1.67 |
| ENSMUSG00000004748 | Mtfp1 | mitochondrial fission process 1 | -1.67 |
| ENSMUSG00000078202 | Nrarp | Notch-regulated ankyrin repeat protein | -1.67 |
| ENSMUSG00000108210 | Gm35808 | predicted gene, 35808 | -1.67 |
| ENSMUSG00000048905 | 4930539E08Rik | RIKEN cDNA 4930539E08 gene | -1.68 |
| ENSMUSG00000024556 | Me2 | malic enzyme 2, NAD(+)-dependent, mitochondrial | -1.68 |
| ENSMUSG00000031402 | Mpp1 | membrane protein, palmitoylated | -1.68 |
| ENSMUSG00000050747 | Trim15 | tripartite motif-containing 15 | -1.68 |
| ENSMUSG00000030790 | Adm | adrenomedullin | -1.68 |
| ENSMUSG00000029455 | Aldh2 | aldehyde dehydrogenase 2, mitochondrial | -1.69 |
| ENSMUSG00000038497 | Tmco3 | transmembrane and coiled-coil domains 3 | -1.69 |
| ENSMUSG00000116542 | Gm17783 | predicted, 17783 | -1.69 |
| ENSMUSG00000025467 | Prap1 | proline-rich acidic protein 1 | -1.69 |
| ENSMUSG00000006567 | Atp7b | ATPase, Cu++ transporting, beta polypeptide | -1.69 |
| ENSMUSG00000042041 | 2010003K11Rik | RIKEN cDNA 2010003K11 gene | -1.69 |
| ENSMUSG00000062410 | Hsd3b3 | hydroxy-delta-5-steroid dehydrogenase, 3 beta- and steroid delta-isomerase 3 | -1.69 |
| ENSMUSG00000106206 | Gm43094 | predicted gene 43094 | -1.69 |
| ENSMUSG00000050158 | Olfr165 | olfactory receptor 165 | -1.69 |
| ENSMUSG00000023150 | Ivns1abp | influenza virus NS1A binding protein | -1.70 |
| ENSMUSG00000029772 | Ahcyl2 | S-adenosylhomocysteine hydrolase-like 2 | -1.70 |
| ENSMUSG00000034275 | Igsf9b | immunoglobulin superfamily, member 9B | -1.70 |
| ENSMUSG00000029401 | Rilpl2 | Rab interacting lysosomal protein-like 2 | -1.70 |

| | | | |
|---|---|---|---|
| ENSMUSG00000074063 | Osgin1 | oxidative stress induced growth inhibitor 1 | -1.70 |
| ENSMUSG00000039062 | Anpep | alanyl (membrane) aminopeptidase | -1.70 |
| ENSMUSG00000112592 | Gm19972 | predicted gene, 19972 | -1.70 |
| ENSMUSG00000110520 | Gm45776 | predicted gene 45776 | -1.70 |
| ENSMUSG00000062962 | Gm6378 | predicted pseudogene 6378 | -1.70 |
| ENSMUSG00000029470 | P2rx4 | purinergic receptor P2X, ligand-gated ion channel 4 | -1.71 |
| ENSMUSG00000038642 | Ctss | cathepsin S | -1.71 |
| ENSMUSG00000045038 | Prkce | protein kinase C, epsilon | -1.71 |
| ENSMUSG00000034413 | Neurl1b | neuralized E3 ubiquitin protein ligase 1B | -1.71 |
| ENSMUSG00000036594 | H2-Aa | histocompatibility 2, class II antigen A, alpha | -1.71 |
| ENSMUSG00000042312 | S100a13 | S100 calcium binding protein A13 | -1.71 |
| ENSMUSG00000037139 | Myom3 | myomesin family, member 3 | -1.71 |
| ENSMUSG00000104350 | Gm38244 | predicted gene, 38244 | -1.71 |
| ENSMUSG00000020405 | Fabp6 | fatty acid binding protein 6, ileal (gastrotropin) | -1.71 |
| ENSMUSG00000100927 | Gm28536 | predicted gene 28536 | -1.71 |
| ENSMUSG00000022445 | Cyp2d26 | cytochrome P450, family 2, subfamily d, polypeptide 26 | -1.71 |
| ENSMUSG00000057963 | Itpk1 | inositol 1,3,4-triphosphate 5/6 kinase | -1.72 |
| ENSMUSG00000041605 | Inava | innate immunity activator | -1.72 |
| ENSMUSG00000034714 | Ttyh2 | tweety family member 2 | -1.72 |
| ENSMUSG00000023915 | Tnfrsf21 | tumor necrosis factor receptor superfamily, member 21 | -1.72 |
| ENSMUSG00000019970 | Sgk1 | serum/glucocorticoid regulated kinase 1 | -1.72 |
| ENSMUSG00000022180 | Slc7a8 | solute carrier family 7 (cationic amino acid transporter, y+ system), member 8 | -1.72 |
| ENSMUSG00000037157 | Il22ra1 | interleukin 22 receptor, alpha 1 | -1.73 |
| ENSMUSG00000038331 | Satb2 | special AT-rich sequence binding protein 2 | -1.73 |
| ENSMUSG00000097099 | Gm9917 | predicted gene 9917 | -1.73 |
| ENSMUSG00000037390 | Muc3 | mucin 3, intestinal | -1.73 |
| ENSMUSG00000024386 | Proc | protein C | -1.73 |
| ENSMUSG00000106417 | Gm42628 | predicted gene 42628 | -1.73 |
| ENSMUSG00000037605 | Adgrl3 | adhesion G protein-coupled receptor L3 | -1.73 |
| ENSMUSG00000106542 | Gm43410 | predicted gene 43410 | -1.73 |
| ENSMUSG00000019558 | Slc6a8 | solute carrier family 6 (neurotransmitter transporter, creatine), member 8 | -1.74 |
| ENSMUSG00000025076 | Casp7 | caspase 7 | -1.74 |
| ENSMUSG00000039910 | Cited2 | Cbp/p300-interacting transactivator, with Glu/Asp-rich carboxy-terminal domain, 2 | -1.74 |
| ENSMUSG00000037661 | Gpr160 | G protein-coupled receptor 160 | -1.74 |
| ENSMUSG00000038843 | Gcnt1 | glucosaminyl (N-acetyl) transferase 1, core 2 | -1.74 |
| ENSMUSG00000079507 | H2-Q1 | histocompatibility 2, Q region locus 1 | -1.74 |
| ENSMUSG00000022048 | Dpysl2 | dihydropyrimidinase-like 2 | -1.75 |
| ENSMUSG00000039774 | Galnt12 | polypeptide N-acetylgalactosaminyltransferase 12 | -1.75 |
| ENSMUSG00000102802 | Mgam2-ps | maltase-glucoamylase 2, pseudogene | -1.75 |
| ENSMUSG00000061411 | Nol4l | nucleolar protein 4-like | -1.75 |
| ENSMUSG00000031897 | Psmb10 | proteasome (prosome, macropain) subunit, beta type 10 | -1.75 |
| ENSMUSG00000028194 | Ddah1 | dimethylarginine dimethylaminohydrolase 1 | -1.75 |
| ENSMUSG00000022504 | Ciita | class II transactivator | -1.75 |
| ENSMUSG00000028536 | 2610528J11Rik | RIKEN cDNA 2610528J11 gene | -1.75 |
| ENSMUSG00000009646 | Pla2g12b | phospholipase A2, group XIIB | -1.75 |
| ENSMUSG00000054474 | Thnsl2 | threonine synthase-like 2 (bacterial) | -1.75 |
| ENSMUSG00000015452 | Ager | advanced glycosylation end product-specific receptor | -1.75 |
| ENSMUSG00000032536 | Trak1 | trafficking protein, kinesin binding 1 | -1.76 |
| ENSMUSG00000032120 | C2cd2l | C2 calcium-dependent domain containing 2-like | -1.76 |
| ENSMUSG00000035642 | Aamdc | adipogenesis associated Mth938 domain containing | -1.76 |
| ENSMUSG00000025092 | Hspa12a | heat shock protein 12A | -1.76 |
| ENSMUSG00000032773 | Chrm1 | cholinergic receptor, muscarinic 1, CNS | -1.76 |
| ENSMUSG00000015468 | Notch4 | notch 4 | -1.76 |
| ENSMUSG00000104094 | Gm37314 | predicted gene, 37314 | -1.76 |
| ENSMUSG00000056133 | Unc93a2 | unc-93 homolog A2 | -1.76 |
| ENSMUSG00000068452 | Duox2 | dual oxidase 2 | -1.77 |
| ENSMUSG00000020889 | Nr1d1 | nuclear receptor subfamily 1, group D, member 1 | -1.77 |
| ENSMUSG00000020182 | Ddc | dopa decarboxylase | -1.77 |
| ENSMUSG00000046352 | Gjb2 | gap junction protein, beta 2 | -1.77 |
| ENSMUSG00000040432 | Ltb4r2 | leukotriene B4 receptor 2 | -1.77 |
| ENSMUSG00000066516 | Klk1b21 | kallikrein 1-related peptidase b21 | -1.77 |
| ENSMUSG00000071036 | Gm10309 | predicted gene 10309 | -1.77 |
| ENSMUSG00000061718 | Ppp1r1b | protein phosphatase 1, regulatory inhibitor subunit 1B | -1.78 |
| ENSMUSG00000032246 | Calml4 | calmodulin-like 4 | -1.78 |
| ENSMUSG00000026726 | Cubn | cubilin (intrinsic factor-cobalamin receptor) | -1.78 |
| ENSMUSG00000046688 | Tifa | TRAF-interacting protein with forkhead-associated domain | -1.78 |
| ENSMUSG00000034687 | Fras1 | Fraser extracellular matrix complex subunit 1 | -1.78 |
| ENSMUSG00000022799 | Arhgap31 | Rho GTPase activating protein 31 | -1.78 |
| ENSMUSG00000023262 | Acy1 | aminoacylase 1 | -1.78 |
| ENSMUSG00000034765 | Dusp5 | dual specificity phosphatase 5 | -1.78 |
| ENSMUSG00000002108 | Nr1h3 | nuclear receptor subfamily 1, group H, member 3 | -1.78 |
| ENSMUSG00000038128 | Camk4 | calcium/calmodulin-dependent protein kinase IV | -1.78 |
| ENSMUSG00000106541 | Gm43246 | predicted gene 43246 | -1.78 |
| ENSMUSG00000110062 | B430319F04Rik | RIKEN cDNA B430319F04 gene | -1.78 |
| ENSMUSG00000020017 | Hal | histidine ammonia lyase | -1.78 |
| ENSMUSG00000074444 | Defa30 | defensin, alpha, 30 | -1.79 |

| | | | |
|---|---|---|---|
| ENSMUSG00000038178 | Slc43a2 | solute carrier family 43, member 2 | -1.79 |
| ENSMUSG00000026271 | Gpr35 | G protein-coupled receptor 35 | -1.79 |
| ENSMUSG00000026692 | Fmo4 | flavin containing monooxygenase 4 | -1.79 |
| ENSMUSG00000078451 | Ppil6 | peptidylprolyl isomerase (cyclophilin)-like 6 | -1.79 |
| ENSMUSG00000024511 | Rab27b | RAB27B, member RAS oncogene family | -1.80 |
| ENSMUSG00000030281 | Il17rc | interleukin 17 receptor C | -1.80 |
| ENSMUSG00000041992 | Rapgef5 | Rap guanine nucleotide exchange factor (GEF) 5 | -1.80 |
| ENSMUSG00000032373 | Car12 | carbonic anhydrase 12 | -1.80 |
| ENSMUSG00000032105 | Pdzd3 | PDZ domain containing 3 | -1.80 |
| ENSMUSG00000026389 | Steap3 | STEAP family member 3 | -1.80 |
| ENSMUSG00000040562 | Gstm2 | glutathione S-transferase, mu 2 | -1.80 |
| ENSMUSG00000106078 | Gm43697 | predicted gene 43697 | -1.80 |
| ENSMUSG00000114705 | Gm18760 | predicted gene, 18760 | -1.80 |
| ENSMUSG00000031144 | Syp | synaptophysin | -1.80 |
| ENSMUSG00000006281 | Tep1 | telomerase associated protein 1 | -1.81 |
| ENSMUSG00000029322 | Plac8 | placenta-specific 8 | -1.81 |
| ENSMUSG00000021336 | Slc17a4 | solute carrier family 17 (sodium phosphate), member 4 | -1.81 |
| ENSMUSG00000021779 | Thrb | thyroid hormone receptor beta | -1.81 |
| ENSMUSG00000070661 | Rnf186 | ring finger protein 186 | -1.81 |
| ENSMUSG00000074261 | Erich4 | glutamate rich 4 | -1.81 |
| ENSMUSG00000055963 | Triqk | triple QxxK/R motif containing | -1.81 |
| ENSMUSG00000041323 | Ak7 | adenylate kinase 7 | -1.81 |
| ENSMUSG00000110289 | 4930412F12Rik | RIKEN cDNA 4930412F12 gene | -1.81 |
| ENSMUSG00000090608 | Gm17200 | predicted gene 17200 | -1.81 |
| ENSMUSG00000017950 | Hnf4a | hepatic nuclear factor 4, alpha | -1.82 |
| ENSMUSG00000021097 | Clmn | calmin | -1.82 |
| ENSMUSG00000031209 | Heph | hephaestin | -1.82 |
| ENSMUSG00000026480 | Ncf2 | neutrophil cytosolic factor 2 | -1.82 |
| ENSMUSG00000056602 | Fry | FRY microtubule binding protein | -1.82 |
| ENSMUSG00000027605 | Acss2 | acyl-CoA synthetase short-chain family member 2 | -1.82 |
| ENSMUSG00000049799 | Lrrc19 | leucine rich repeat containing 19 | -1.82 |
| ENSMUSG00000109438 | Gm45073 | predicted gene 45073 | -1.82 |
| ENSMUSG00000019699 | Akt3 | thymoma viral proto-oncogene 3 | -1.82 |
| ENSMUSG00000035878 | Hykk | hydroxylysine kinase 1 | -1.82 |
| ENSMUSG00000030616 | Sytl2 | synaptotagmin-like 2 | -1.83 |
| ENSMUSG00000030934 | Oat | ornithine aminotransferase | -1.83 |
| ENSMUSG00000037031 | Tspan15 | tetraspanin 15 | -1.83 |
| ENSMUSG00000055435 | Maf | avian musculoaponeurotic fibrosarcoma oncogene homolog | -1.83 |
| ENSMUSG00000031853 | Map3k21 | mitogen-activated protein kinase kinase kinase 21 | -1.83 |
| ENSMUSG00000021830 | Txndc16 | thioredoxin domain containing 16 | -1.83 |
| ENSMUSG00000029377 | Ereg | epiregulin | -1.83 |
| ENSMUSG00000054598 | 9130230L23Rik | RIKEN cDNA 9130230L23 gene | -1.83 |
| ENSMUSG00000038239 | Hrc | histidine rich calcium binding protein | -1.83 |
| ENSMUSG00000000530 | Acvrl1 | activin A receptor, type II-like 1 | -1.83 |
| ENSMUSG00000109293 | Dcst2 | DC-STAMP domain containing 2 | -1.83 |
| ENSMUSG00000022824 | Muc13 | mucin 13, epithelial transmembrane | -1.84 |
| ENSMUSG00000022938 | Fam3b | family with sequence similarity 3, member B | -1.84 |
| ENSMUSG00000022469 | Rapgef3 | Rap guanine nucleotide exchange factor (GEF) 3 | -1.84 |
| ENSMUSG00000034265 | Zdhhc14 | zinc finger, DHHC domain containing 14 | -1.84 |
| ENSMUSG00000051452 | Gm11437 | predicted gene 11437 | -1.84 |
| ENSMUSG00000112944 | Gm48885 | predicted gene, 48885 | -1.84 |
| ENSMUSG00000096546 | Smlr1 | small leucine-rich protein 1 | -1.84 |
| ENSMUSG00000033872 | Best4-ps | bestrophin 4, pseudogene | -1.84 |
| ENSMUSG00000059146 | Ntrk3 | neurotrophic tyrosine kinase, receptor, type 3 | -1.84 |
| ENSMUSG00000108622 | Gm36864 | predicted gene, 36864 | -1.84 |
| ENSMUSG00000003545 | Fosb | FBJ osteosarcoma oncogene B | -1.85 |
| ENSMUSG00000024254 | Abcg8 | ATP binding cassette subfamily G member 8 | -1.85 |
| ENSMUSG00000049690 | Nckap5 | NCK-associated protein 5 | -1.85 |
| ENSMUSG00000090214 | Gm15657 | predicted gene 15657 | -1.85 |
| ENSMUSG00000096954 | Gdap10 | ganglioside-induced differentiation-associated-protein 10 | -1.85 |
| ENSMUSG00000020600 | Slc7a15 | solute carrier family 7 (cationic amino acid transporter, y+ system), member 15 | -1.85 |
| ENSMUSG00000038188 | Scarf1 | scavenger receptor class F, member 1 | -1.85 |
| ENSMUSG00000097575 | Gm26796 | predicted gene, 26796 | -1.85 |
| ENSMUSG00000030587 | 2200002D01Rik | RIKEN cDNA 2200002D01 gene | -1.86 |
| ENSMUSG00000042428 | Mgat3 | mannoside acetylglucosaminyltransferase 3 | -1.86 |
| ENSMUSG00000038903 | Ccdc68 | coiled-coil domain containing 68 | -1.86 |
| ENSMUSG00000039405 | Prss23 | protease, serine 23 | -1.86 |
| ENSMUSG00000022853 | Ehhadh | enoyl-Coenzyme A, hydratase/3-hydroxyacyl Coenzyme A dehydrogenase | -1.86 |
| ENSMUSG00000034957 | Cebpa | CCAAT/enhancer binding protein (C/EBP), alpha | -1.86 |
| ENSMUSG00000029630 | Cyp3a25 | cytochrome P450, family 3, subfamily a, polypeptide 25 | -1.86 |
| ENSMUSG00000050106 | Tmc8 | transmembrane channel-like gene family 8 | -1.86 |
| ENSMUSG00000021223 | Papln | papilin, proteoglycan-like sulfated glycoprotein | -1.86 |
| ENSMUSG00000053675 | Tgm5 | transglutaminase 5 | -1.86 |
| ENSMUSG00000008461 | Fut1 | fucosyltransferase 1 | -1.86 |
| ENSMUSG00000054146 | Krt15 | keratin 15 | -1.86 |
| ENSMUSG00000091475 | 2810468N07Rik | RIKEN cDNA 2810468N07 gene | -1.86 |

| | | | |
|---|---|---|---|
| ENSMUSG00000100182 | 1810006J02Rik | RIKEN cDNA 1810006J02 gene | -1.86 |
| ENSMUSG00000043020 | Wdr63 | WD repeat domain 63 | -1.86 |
| ENSMUSG00000024180 | Tmem8 | transmembrane protein 8 (five membrane-spanning domains) | -1.87 |
| ENSMUSG00000056413 | Adap1 | ArfGAP with dual PH domains 1 | -1.87 |
| ENSMUSG00000053310 | Nrgn | neurogranin | -1.87 |
| ENSMUSG00000105872 | Gm9515 | predicted gene 9515 | -1.87 |
| ENSMUSG00000042846 | Lrrtm3 | leucine rich repeat transmembrane neuronal 3 | -1.87 |
| ENSMUSG00000027463 | Slc52a3 | solute carrier protein family 52, member 3 | -1.88 |
| ENSMUSG00000027358 | Bmp2 | bone morphogenetic protein 2 | -1.88 |
| ENSMUSG00000024131 | Slc3a1 | solute carrier family 3, member 1 | -1.88 |
| ENSMUSG00000045316 | Fahd1 | fumarylacetoacetate hydrolase domain containing 1 | -1.88 |
| ENSMUSG00000052085 | Dock8 | dedicator of cytokinesis 8 | -1.88 |
| ENSMUSG00000074622 | Mafb | v-maf musculoaponeurotic fibrosarcoma oncogene family, protein B (avian) | -1.88 |
| ENSMUSG00000032487 | Ptgs2 | prostaglandin-endoperoxide synthase 2 | -1.88 |
| ENSMUSG00000004151 | Etv1 | ets variant 1 | -1.88 |
| ENSMUSG00000071550 | Cfap44 | cilia and flagella associated protein 44 | -1.88 |
| ENSMUSG00000030838 | Ush1c | USH1 protein network component harmonin | -1.89 |
| ENSMUSG00000018909 | Arrb1 | arrestin, beta 1 | -1.89 |
| ENSMUSG00000030650 | Tmc5 | transmembrane channel-like gene family 5 | -1.89 |
| ENSMUSG00000025888 | Casp1 | caspase 1 | -1.89 |
| ENSMUSG00000053862 | Slc51b | solute carrier family 51, beta subunit | -1.89 |
| ENSMUSG00000026784 | Pdss1 | prenyl (solanesyl) diphosphate synthase, subunit 1 | -1.89 |
| ENSMUSG00000021798 | Ldb3 | LIM domain binding 3 | -1.89 |
| ENSMUSG00000101693 | Gm19461 | predicted gene, 19461 | -1.89 |
| ENSMUSG00000078650 | G6pc | glucose-6-phosphatase, catalytic | -1.89 |
| ENSMUSG00000108353 | Gm45205 | predicted gene 45205 | -1.89 |
| ENSMUSG00000029992 | Gfpt1 | glutamine fructose-6-phosphate transaminase 1 | -1.90 |
| ENSMUSG00000024597 | Slc12a2 | solute carrier family 12, member 2 | -1.90 |
| ENSMUSG00000027901 | Dennd2d | DENN/MADD domain containing 2D | -1.90 |
| ENSMUSG00000073421 | H2-Ab1 | histocompatibility 2, class II antigen A, beta 1 | -1.90 |
| ENSMUSG00000044626 | Liph | lipase, member H | -1.90 |
| ENSMUSG00000020151 | Ptprr | protein tyrosine phosphatase, receptor type, R | -1.90 |
| ENSMUSG00000056671 | Prelid2 | PRELI domain containing 2 | -1.90 |
| ENSMUSG00000043432 | Leng9 | leukocyte receptor cluster (LRC) member 9 | -1.91 |
| ENSMUSG00000029919 | Hpgds | hematopoietic prostaglandin D synthase | -1.91 |
| ENSMUSG00000029059 | Fam213b | family with sequence similarity 213, member B | -1.91 |
| ENSMUSG00000037440 | Vnn1 | vanin 1 | -1.91 |
| ENSMUSG00000096001 | 2610528A11Rik | RIKEN cDNA 2610528A11 gene | -1.91 |
| ENSMUSG00000104966 | Gm43273 | predicted gene 43273 | -1.91 |
| ENSMUSG00000098609 | Anxa11os | annexin A11, opposite strand | -1.91 |
| ENSMUSG00000085890 | Tnfsf13os | tumor necrosis factor (ligand) superfamily, member 13, opposite strand | -1.91 |
| ENSMUSG00000044576 | Garem2 | GRB2 associated regulator of MAPK1 subtype 2 | -1.91 |
| ENSMUSG00000109093 | Gm19950 | predicted gene, 19950 | -1.91 |
| ENSMUSG00000034570 | Inpp5j | inositol polyphosphate 5-phosphatase J | -1.92 |
| ENSMUSG00000035000 | Dpp4 | dipeptidylpeptidase 4 | -1.92 |
| ENSMUSG00000012123 | Crybg2 | crystallin beta-gamma domain containing 2 | -1.92 |
| ENSMUSG00000079440 | Alpi | alkaline phosphatase, intestinal | -1.92 |
| ENSMUSG00000054999 | Naaladl1 | N-acetylated alpha-linked acidic dipeptidase-like 1 | -1.92 |
| ENSMUSG00000029811 | Aoc1 | amine oxidase, copper-containing 1 | -1.92 |
| ENSMUSG00000069814 | Ccdc92b | coiled-coil domain containing 92B | -1.92 |
| ENSMUSG00000033491 | Prss35 | protease, serine 35 | -1.92 |
| ENSMUSG00000112129 | Pbld1 | phenazine biosynthesis-like protein domain containing 1 | -1.92 |
| ENSMUSG00000116738 | CT009627.2 | TEC | -1.92 |
| ENSMUSG00000086012 | Gm15902 | predicted gene 15902 | -1.92 |
| ENSMUSG00000113935 | Gm35732 | predicted gene, 35732 | -1.92 |
| ENSMUSG00000034127 | Tspan8 | tetraspanin 8 | -1.93 |
| ENSMUSG00000026670 | Uap1 | UDP-N-acetylglucosamine pyrophosphorylase 1 | -1.93 |
| ENSMUSG00000026399 | Cd55 | CD55 molecule, decay accelerating factor for complement | -1.93 |
| ENSMUSG00000051111 | Sv2c | synaptic vesicle glycoprotein 2c | -1.93 |
| ENSMUSG00000109585 | Gm45358 | predicted gene 45358 | -1.93 |
| ENSMUSG00000047414 | Flrt2 | fibronectin leucine rich transmembrane protein 2 | -1.93 |
| ENSMUSG00000109711 | Gm45445 | predicted gene 45445 | -1.93 |
| ENSMUSG00000071185 | Olfr1357 | olfactory receptor 1357 | -1.93 |
| ENSMUSG00000029054 | Gabrd | gamma-aminobutyric acid (GABA) A receptor, subunit delta | -1.93 |
| ENSMUSG00000029821 | Gsdme | gasdermin E | -1.94 |
| ENSMUSG00000041268 | Dmxl2 | Dmx-like 2 | -1.94 |
| ENSMUSG00000001095 | Slc13a2 | solute carrier family 13 (sodium-dependent dicarboxylate transporter), member 2 | -1.94 |
| ENSMUSG00000053279 | Aldh1a1 | aldehyde dehydrogenase family 1, subfamily A1 | -1.94 |
| ENSMUSG00000021913 | Ogdhl | oxoglutarate dehydrogenase-like | -1.94 |
| ENSMUSG00000110060 | Gm9860 | predicted gene 9860 | -1.94 |
| ENSMUSG00000020839 | Tmigd1 | transmembrane and immunoglobulin domain containing 1 | -1.94 |
| ENSMUSG00000112513 | 4930422I22Rik | RIKEN cDNA 4930422I22 gene | -1.94 |
| ENSMUSG00000087047 | 1700110K17Rik | RIKEN cDNA 1700110K17 gene | -1.94 |
| ENSMUSG00000040528 | Milr1 | mast cell immunoglobulin like receptor 1 | -1.94 |
| ENSMUSG00000044734 | Serpinb1a | serine (or cysteine) peptidase inhibitor, clade B, member 1a | -1.95 |
| ENSMUSG00000074272 | Ceacam1 | carcinoembryonic antigen-related cell adhesion molecule 1 | -1.95 |

| | | | |
|---|---|---|---|
| ENSMUSG00000052595 | A1cf | APOBEC1 complementation factor | -1.95 |
| ENSMUSG00000025059 | Gk | glycerol kinase | -1.95 |
| ENSMUSG00000067206 | Lrrc66 | leucine rich repeat containing 66 | -1.95 |
| ENSMUSG00000027377 | Mall | mal, T cell differentiation protein-like | -1.95 |
| ENSMUSG00000033213 | AA467197 | expressed sequence AA467197 | -1.95 |
| ENSMUSG00000106475 | Gm43011 | predicted gene 43011 | -1.95 |
| ENSMUSG00000010830 | Kdelr3 | KDEL (Lys-Asp-Glu-Leu) endoplasmic reticulum protein retention receptor 3 | -1.95 |
| ENSMUSG00000050944 | Efcab5 | EF-hand calcium binding domain 5 | -1.95 |
| ENSMUSG00000004038 | Gstm3 | glutathione S-transferase, mu 3 | -1.95 |
| ENSMUSG00000086479 | Gm16014 | predicted gene 16014 | -1.95 |
| ENSMUSG00000111379 | Gm48346 | predicted gene, 48346 | -1.95 |
| ENSMUSG00000024960 | Plcb3 | phospholipase C, beta 3 | -1.96 |
| ENSMUSG00000031391 | L1cam | L1 cell adhesion molecule | -1.96 |
| ENSMUSG00000024105 | Themis3 | thymocyte selection associated family member 3 | -1.96 |
| ENSMUSG00000006345 | Ggt1 | gamma-glutamyltransferase 1 | -1.96 |
| ENSMUSG00000110622 | Iqcn | IQ motif containing N | -1.96 |
| ENSMUSG00000069804 | Gm10277 | predicted gene 10277 | -1.96 |
| ENSMUSG00000047728 | BC025446 | cDNA sequence BC025446 | -1.96 |
| ENSMUSG00000038195 | Rilp | Rab interacting lysosomal protein | -1.96 |
| ENSMUSG00000087230 | Mroh3 | maestro heat-like repeat family member 3 | -1.96 |
| ENSMUSG00000003032 | Klf4 | Kruppel-like factor 4 (gut) | -1.97 |
| ENSMUSG00000030865 | Chp2 | calcineurin-like EF hand protein 2 | -1.97 |
| ENSMUSG00000104571 | Gm43010 | predicted gene 43010 | -1.97 |
| ENSMUSG00000019278 | Dpep1 | dipeptidase 1 (renal) | -1.97 |
| ENSMUSG00000020709 | Adap2 | ArfGAP with dual PH domains 2 | -1.97 |
| ENSMUSG00000029352 | Crybb3 | crystallin, beta B3 | -1.97 |
| ENSMUSG00000099025 | Gm27162 | predicted gene 27162 | -1.97 |
| ENSMUSG00000097257 | Gm26768 | predicted gene, 26768 | -1.97 |
| ENSMUSG00000040249 | Lrp1 | low density lipoprotein receptor-related protein 1 | -1.98 |
| ENSMUSG00000074447 | Defa21 | defensin, alpha, 21 | -1.98 |
| ENSMUSG00000041515 | Irf8 | interferon regulatory factor 8 | -1.98 |
| ENSMUSG00000005686 | Ampd3 | adenosine monophosphate deaminase 3 | -1.98 |
| ENSMUSG00000024222 | Fkbp5 | FK506 binding protein 5 | -1.98 |
| ENSMUSG00000047250 | Ptgs1 | prostaglandin-endoperoxide synthase 1 | -1.98 |
| ENSMUSG00000041565 | L3mbtl4 | L3MBTL4 histone methyl-lysine binding protein | -1.98 |
| ENSMUSG00000038372 | Gmds | GDP-mannose 4, 6-dehydratase | -1.99 |
| ENSMUSG00000053886 | Sh2d4a | SH2 domain containing 4A | -1.99 |
| ENSMUSG00000032047 | Acat1 | acetyl-Coenzyme A acetyltransferase 1 | -1.99 |
| ENSMUSG00000037818 | Abhd18 | abhydrolase domain containing 18 | -1.99 |
| ENSMUSG00000046159 | Chrm3 | cholinergic receptor, muscarinic 3, cardiac | -1.99 |
| ENSMUSG00000019232 | Etnppl | ethanolamine phosphate phospholyase | -1.99 |
| ENSMUSG00000038209 | Itln1 | intelectin 1 (galactofuranose binding) | -2.00 |
| ENSMUSG00000021196 | Pfkp | phosphofructokinase, platelet | -2.00 |
| ENSMUSG00000003420 | Fcgrt | Fc receptor, IgG, alpha chain transporter | -2.00 |
| ENSMUSG00000029700 | Slc13a1 | solute carrier family 13 (sodium/sulfate symporters), member 1 | -2.00 |
| ENSMUSG00000086825 | Gm15675 | predicted gene 15675 | -2.00 |
| ENSMUSG00000024503 | Spink1 | serine peptidase inhibitor, Kazal type 1 | -2.00 |
| ENSMUSG00000024885 | Aldh3b1 | aldehyde dehydrogenase 3 family, member B1 | -2.00 |
| ENSMUSG00000021379 | Id4 | inhibitor of DNA binding 4 | -2.00 |
| ENSMUSG00000079559 | Colca2 | COLCA2 homolog | -2.00 |
| ENSMUSG00000106802 | Gm42885 | predicted gene 42885 | -2.00 |
| ENSMUSG00000015970 | Chdh | choline dehydrogenase | -2.01 |
| ENSMUSG00000022419 | Deptor | DEP domain containing MTOR-interacting protein | -2.01 |
| ENSMUSG00000091705 | H2-Q2 | histocompatibility 2, Q region locus 2 | -2.01 |
| ENSMUSG00000038020 | Rapgefl1 | Rap guanine nucleotide exchange factor (GEF)-like 1 | -2.01 |
| ENSMUSG00000061825 | Ces2c | carboxylesterase 2C | -2.01 |
| ENSMUSG00000057182 | Scn3a | sodium channel, voltage-gated, type III, alpha | -2.01 |
| ENSMUSG00000028337 | Coro2a | coronin, actin binding protein 2A | -2.02 |
| ENSMUSG00000038807 | Rap1gap2 | RAP1 GTPase activating protein 2 | -2.02 |
| ENSMUSG00000028699 | Tspan1 | tetraspanin 1 | -2.02 |
| ENSMUSG00000006360 | Crip1 | cysteine-rich protein 1 (intestinal) | -2.02 |
| ENSMUSG00000024642 | Tle4 | transducin-like enhancer of split 4 | -2.02 |
| ENSMUSG00000034919 | Ttc22 | tetratricopeptide repeat domain 22 | -2.02 |
| ENSMUSG00000074115 | Saa1 | serum amyloid A 1 | -2.02 |
| ENSMUSG00000034845 | Plvap | plasmalemma vesicle associated protein | -2.02 |
| ENSMUSG00000032899 | Styk1 | serine/threonine/tyrosine kinase 1 | -2.03 |
| ENSMUSG00000026688 | Mgst3 | microsomal glutathione S-transferase 3 | -2.03 |
| ENSMUSG00000039899 | Fgl2 | fibrinogen-like protein 2 | -2.03 |
| ENSMUSG00000023045 | Soat2 | sterol O-acyltransferase 2 | -2.03 |
| ENSMUSG00000034947 | Tmem106a | transmembrane protein 106A | -2.03 |
| ENSMUSG00000028544 | Slc5a9 | solute carrier family 5 (sodium/glucose cotransporter), member 9 | -2.03 |
| ENSMUSG00000001665 | Gstt3 | glutathione S-transferase, theta 3 | -2.03 |
| ENSMUSG00000115970 | 8430426J06Rik | RIKEN cDNA 8430426J06 gene | -2.03 |
| ENSMUSG00000004864 | Mapk13 | mitogen-activated protein kinase 13 | -2.04 |
| ENSMUSG00000066800 | Rnasel | ribonuclease L (2', 5'-oligoisoadenylate synthetase-dependent) | -2.04 |
| ENSMUSG00000024313 | Mep1b | meprin 1 beta | -2.04 |

| | | | |
|---|---|---|---|
| ENSMUSG00000074604 | Mgst2 | microsomal glutathione S-transferase 2 | -2.04 |
| ENSMUSG00000024124 | Prss30 | protease, serine 30 | -2.04 |
| ENSMUSG00000022546 | Gpt | glutamic pyruvic transaminase, soluble | -2.04 |
| ENSMUSG00000028976 | Slc2a5 | solute carrier family 2 (facilitated glucose transporter), member 5 | -2.04 |
| ENSMUSG00000074196 | Clca4c-ps | chloride channel accessory 4C, pseudogene | -2.04 |
| ENSMUSG00000108206 | Gm44427 | predicted gene, 44427 | -2.04 |
| ENSMUSG00000085152 | Gm11496 | predicted gene 11496 | -2.04 |
| ENSMUSG00000021250 | Fos | FBJ osteosarcoma oncogene | -2.05 |
| ENSMUSG00000028307 | Aldob | aldolase B, fructose-bisphosphate | -2.05 |
| ENSMUSG00000023914 | Mep1a | meprin 1 alpha | -2.05 |
| ENSMUSG00000023243 | Kcnk5 | potassium channel, subfamily K, member 5 | -2.05 |
| ENSMUSG00000032978 | Guca2b | guanylate cyclase activator 2b (retina) | -2.05 |
| ENSMUSG00000030214 | Plbd1 | phospholipase B domain containing 1 | -2.05 |
| ENSMUSG00000084948 | 1700061H18Rik | RIKEN cDNA 1700061H18 gene | -2.05 |
| ENSMUSG00000040505 | Abcg5 | ATP binding cassette subfamily G member 5 | -2.06 |
| ENSMUSG00000010601 | Apol7a | apolipoprotein L 7a | -2.06 |
| ENSMUSG00000038534 | Osbpl7 | oxysterol binding protein-like 7 | -2.06 |
| ENSMUSG00000030492 | Slc7a9 | solute carrier family 7 (cationic amino acid transporter, y+ system), member 9 | -2.06 |
| ENSMUSG00000044165 | Bcl2l15 | BCLl2-like 15 | -2.06 |
| ENSMUSG00000026928 | Card9 | caspase recruitment domain family, member 9 | -2.06 |
| ENSMUSG00000057074 | Ces1g | carboxylesterase 1G | -2.06 |
| ENSMUSG00000054161 | Fam83e | family with sequence similarity 83, member E | -2.07 |
| ENSMUSG00000041642 | Kif21b | kinesin family member 21B | -2.07 |
| ENSMUSG00000035429 | Ptprh | protein tyrosine phosphatase, receptor type, H | -2.07 |
| ENSMUSG00000103696 | Gm37531 | predicted gene, 37531 | -2.07 |
| ENSMUSG00000087326 | Gm12503 | predicted gene 12503 | -2.07 |
| ENSMUSG00000087530 | Gm15533 | predicted gene 15533 | -2.07 |
| ENSMUSG00000050866 | Clrn3 | clarin 3 | -2.08 |
| ENSMUSG00000054469 | Lclat1 | lysocardiolipin acyltransferase 1 | -2.08 |
| ENSMUSG00000032548 | Slco2a1 | solute carrier organic anion transporter family, member 2a1 | -2.08 |
| ENSMUSG00000020062 | Slc5a8 | solute carrier family 5 (iodide transporter), member 8 | -2.08 |
| ENSMUSG00000028150 | Rorc | RAR-related orphan receptor gamma | -2.08 |
| ENSMUSG00000074217 | 2210011C24Rik | RIKEN cDNA 2210011C24 gene | -2.08 |
| ENSMUSG00000115388 | Eppk1 | epiplakin 1 | -2.09 |
| ENSMUSG00000035493 | Tgfbi | transforming growth factor, beta induced | -2.09 |
| ENSMUSG00000017639 | Rab11fip4 | RAB11 family interacting protein 4 (class II) | -2.09 |
| ENSMUSG00000032776 | Mctp2 | multiple C2 domains, transmembrane 2 | -2.09 |
| ENSMUSG00000103527 | Gm37261 | predicted gene, 37261 | -2.09 |
| ENSMUSG00000025189 | Cnnm1 | cyclin M1 | -2.09 |
| ENSMUSG00000072244 | Trim6 | tripartite motif-containing 6 | -2.09 |
| ENSMUSG00000024112 | Cacna1h | calcium channel, voltage-dependent, T type, alpha 1H subunit | -2.10 |
| ENSMUSG00000019478 | Rab4a | RAB4A, member RAS oncogene family | -2.10 |
| ENSMUSG00000031453 | Rasa3 | RAS p21 protein activator 3 | -2.10 |
| ENSMUSG00000094651 | Gal3st2 | galactose-3-O-sulfotransferase 2 | -2.10 |
| ENSMUSG00000028024 | Enpep | glutamyl aminopeptidase | -2.10 |
| ENSMUSG00000034918 | Cdhr2 | cadherin-related family member 2 | -2.10 |
| ENSMUSG00000074345 | Tnfaip8l3 | tumor necrosis factor, alpha-induced protein 8-like 3 | -2.10 |
| ENSMUSG00000030834 | Abcc6 | ATP-binding cassette, sub-family C (CFTR/MRP), member 6 | -2.10 |
| ENSMUSG00000035775 | Krt20 | keratin 20 | -2.11 |
| ENSMUSG00000070777 | Ceacam20 | carcinoembryonic antigen-related cell adhesion molecule 20 | -2.11 |
| ENSMUSG00000108476 | Gm44974 | predicted gene 44974 | -2.11 |
| ENSMUSG00000024292 | Cyp4f14 | cytochrome P450, family 4, subfamily f, polypeptide 14 | -2.12 |
| ENSMUSG00000019888 | Mgat4c | MGAT4 family, member C | -2.12 |
| ENSMUSG00000085683 | Tmem238l | transmembrane protein 238 like | -2.12 |
| ENSMUSG00000027508 | Pag1 | phosphoprotein associated with glycosphingolipid microdomains 1 | -2.12 |
| ENSMUSG00000058163 | Gm5431 | predicted gene 5431 | -2.12 |
| ENSMUSG00000040441 | Slc26a10 | solute carrier family 26, member 10 | -2.12 |
| ENSMUSG00000072618 | Gm10384 | predicted gene 10384 | -2.12 |
| ENSMUSG00000108961 | Gm32540 | predicted gene, 32540 | -2.12 |
| ENSMUSG00000114945 | Gm48265 | predicted gene, 48265 | -2.12 |
| ENSMUSG00000004266 | Ptpn6 | protein tyrosine phosphatase, non-receptor type 6 | -2.13 |
| ENSMUSG00000073802 | Cdkn2b | cyclin dependent kinase inhibitor 2B | -2.13 |
| ENSMUSG00000019872 | Smpdl3a | sphingomyelin phosphodiesterase, acid-like 3A | -2.13 |
| ENSMUSG00000068587 | Mgam | maltase-glucoamylase | -2.14 |
| ENSMUSG00000060586 | H2-Eb1 | histocompatibility 2, class II antigen E beta | -2.14 |
| ENSMUSG00000018983 | E2f2 | E2F transcription factor 2 | -2.14 |
| ENSMUSG00000026222 | Sp100 | nuclear antigen Sp100 | -2.14 |
| ENSMUSG00000031980 | Agt | angiotensinogen (serpin peptidase inhibitor, clade A, member 8) | -2.14 |
| ENSMUSG00000024846 | Cst6 | cystatin E/M | -2.14 |
| ENSMUSG00000044361 | BC024139 | cDNA sequence BC024139 | -2.14 |
| ENSMUSG00000026546 | Cfap45 | cilia and flagella associated protein 45 | -2.14 |
| ENSMUSG00000043953 | Ccrl2 | chemokine (C-C motif) receptor-like 2 | -2.14 |
| ENSMUSG00000024734 | Zp1 | zona pellucida glycoprotein 1 | -2.14 |
| ENSMUSG00000078945 | Naip2 | NLR family, apoptosis inhibitory protein 2 | -2.15 |
| ENSMUSG00000024713 | Pcsk5 | proprotein convertase subtilisin/kexin type 5 | -2.15 |
| ENSMUSG00000072812 | Ahnak2 | AHNAK nucleoprotein 2 | -2.15 |

| | | | |
|---|---|---|---|
| ENSMUSG00000090115 | Usp49 | ubiquitin specific peptidase 49 | -2.15 |
| ENSMUSG00000000805 | Car4 | carbonic anhydrase 4 | -2.15 |
| ENSMUSG00000033615 | Cplx1 | complexin 1 | -2.15 |
| ENSMUSG00000091443 | Gm17023 | predicted gene 17023 | -2.15 |
| ENSMUSG00000038167 | Plekhg6 | pleckstrin homology domain containing, family G (with RhoGef domain) member 6 | -2.16 |
| ENSMUSG00000031844 | Hsd17b2 | hydroxysteroid (17-beta) dehydrogenase 2 | -2.16 |
| ENSMUSG00000048572 | Tmem252 | transmembrane protein 252 | -2.16 |
| ENSMUSG00000056162 | Cndp1 | carnosine dipeptidase 1 (metallopeptidase M20 family) | -2.16 |
| ENSMUSG00000030621 | Me3 | malic enzyme 3, NADP(+)-dependent, mitochondrial | -2.16 |
| ENSMUSG00000110597 | Gm8798 | predicted gene 8798 | -2.16 |
| ENSMUSG00000021062 | Rab15 | RAB15, member RAS oncogene family | -2.17 |
| ENSMUSG00000024747 | Aldh1a7 | aldehyde dehydrogenase family 1, subfamily A7 | -2.17 |
| ENSMUSG00000097789 | Gm2115 | predicted gene 2115 | -2.17 |
| ENSMUSG00000034116 | Vav1 | vav 1 oncogene | -2.18 |
| ENSMUSG00000020614 | Fam20a | family with sequence similarity 20, member A | -2.18 |
| ENSMUSG00000024247 | Pkdcc | protein kinase domain containing, cytoplasmic | -2.18 |
| ENSMUSG00000061728 | Btnl7-ps | butyrophilin-like 7, pseudogene | -2.18 |
| ENSMUSG00000091956 | C2cd4b | C2 calcium-dependent domain containing 4B | -2.18 |
| ENSMUSG00000026077 | Npas2 | neuronal PAS domain protein 2 | -2.19 |
| ENSMUSG00000028635 | Edn2 | endothelin 2 | -2.19 |
| ENSMUSG00000024697 | Gna14 | guanine nucleotide binding protein, alpha 14 | -2.19 |
| ENSMUSG00000114662 | Gm31683 | predicted gene, 31683 | -2.19 |
| ENSMUSG00000104917 | Gm43289 | predicted gene 43289 | -2.19 |
| ENSMUSG00000031442 | Mcf2l | mcf.2 transforming sequence-like | -2.20 |
| ENSMUSG00000027452 | Acss1 | acyl-CoA synthetase short-chain family member 1 | -2.20 |
| ENSMUSG00000046697 | Enpp7 | ectonucleotide pyrophosphatase/phosphodiesterase 7 | -2.20 |
| ENSMUSG00000052131 | Akr1b7 | aldo-keto reductase family 1, member B7 | -2.20 |
| ENSMUSG00000106969 | Gm42882 | predicted gene 42882 | -2.20 |
| ENSMUSG00000027230 | Creb3l1 | cAMP responsive element binding protein 3-like 1 | -2.21 |
| ENSMUSG00000045441 | Gprin3 | GPRIN family member 3 | -2.21 |
| ENSMUSG00000038305 | Spats2l | spermatogenesis associated, serine-rich 2-like | -2.21 |
| ENSMUSG00000023259 | Slc26a6 | solute carrier family 26, member 6 | -2.21 |
| ENSMUSG00000103761 | Gm37859 | predicted gene, 37859 | -2.21 |
| ENSMUSG00000033268 | Duox1 | dual oxidase 1 | -2.21 |
| ENSMUSG00000019359 | Gdpd2 | glycerophosphodiester phosphodiesterase domain containing 2 | -2.21 |
| ENSMUSG00000100241 | Slc18a3 | solute carrier family 18 (vesicular monoamine), member 3 | -2.21 |
| ENSMUSG00000041119 | Pde9a | phosphodiesterase 9A | -2.22 |
| ENSMUSG00000027200 | Sema6d | sema domain, transmembrane domain (TM), and cytoplasmic domain, (semaphorin) 6D | -2.22 |
| ENSMUSG00000031255 | Sytl4 | synaptotagmin-like 4 | -2.22 |
| ENSMUSG00000067231 | Cyp2c65 | cytochrome P450, family 2, subfamily c, polypeptide 65 | -2.22 |
| ENSMUSG00000035969 | Rusc2 | RUN and SH3 domain containing 2 | -2.22 |
| ENSMUSG00000046840 | Hnf4aos | hepatic nuclear factor 4 alpha, opposite strand | -2.22 |
| ENSMUSG00000030364 | Clec2h | C-type lectin domain family 2, member h | -2.23 |
| ENSMUSG00000031838 | Ifi30 | interferon gamma inducible protein 30 | -2.23 |
| ENSMUSG00000062638 | Btnl1 | butyrophilin-like 1 | -2.23 |
| ENSMUSG00000022809 | Nr1i2 | nuclear receptor subfamily 1, group I, member 2 | -2.23 |
| ENSMUSG00000010660 | Plcd1 | phospholipase C, delta 1 | -2.23 |
| ENSMUSG00000038963 | Slco4a1 | solute carrier organic anion transporter family, member 4a1 | -2.23 |
| ENSMUSG00000052229 | Gpr17 | G protein-coupled receptor 17 | -2.23 |
| ENSMUSG00000021456 | Fbp2 | fructose bisphosphatase 2 | -2.24 |
| ENSMUSG00000031803 | B3gnt3 | UDP-GlcNAc:betaGal beta-1,3-N-acetylglucosaminyltransferase 3 | -2.24 |
| ENSMUSG00000030157 | Clec2d | C-type lectin domain family 2, member d | -2.24 |
| ENSMUSG00000036151 | Tm6sf2 | transmembrane 6 superfamily member 2 | -2.24 |
| ENSMUSG00000025790 | Slco3a1 | solute carrier organic anion transporter family, member 3a1 | -2.24 |
| ENSMUSG00000025528 | 2010106E10Rik | RIKEN cDNA 2010106E10 gene | -2.24 |
| ENSMUSG00000079547 | H2-DMb1 | histocompatibility 2, class II, locus Mb1 | -2.24 |
| ENSMUSG00000031608 | Galnt7 | polypeptide N-acetylgalactosaminyltransferase 7 | -2.25 |
| ENSMUSG00000023959 | Clic5 | chloride intracellular channel 5 | -2.25 |
| ENSMUSG00000021565 | Slc6a19 | solute carrier family 6 (neurotransmitter transporter), member 19 | -2.25 |
| ENSMUSG00000044017 | Adgrd1 | adhesion G protein-coupled receptor D1 | -2.25 |
| ENSMUSG00000033721 | Vav3 | vav 3 oncogene | -2.25 |
| ENSMUSG00000090293 | Gm17034 | predicted gene 17034 | -2.25 |
| ENSMUSG00000045912 | C2cd4c | C2 calcium-dependent domain containing 4C | -2.25 |
| ENSMUSG00000104519 | Gm37161 | predicted gene, 37161 | -2.25 |
| ENSMUSG00000090145 | Ugt1a6b | UDP glucuronosyltransferase 1 family, polypeptide A6B | -2.25 |
| ENSMUSG00000021676 | Iqgap2 | IQ motif containing GTPase activating protein 2 | -2.26 |
| ENSMUSG00000027227 | Sord | sorbitol dehydrogenase | -2.26 |
| ENSMUSG00000034785 | Dio1 | deiodinase, iodothyronine, type I | -2.26 |
| ENSMUSG00000031497 | Tnfsf13b | tumor necrosis factor (ligand) superfamily, member 13b | -2.26 |
| ENSMUSG00000025497 | Cdhr5 | cadherin-related family member 5 | -2.27 |
| ENSMUSG00000031886 | Ces2e | carboxylesterase 2E | -2.27 |
| ENSMUSG00000024140 | Epas1 | endothelial PAS domain protein 1 | -2.27 |
| ENSMUSG00000021553 | Slc28a3 | solute carrier family 28 (sodium-coupled nucleoside transporter), member 3 | -2.27 |
| ENSMUSG00000023827 | Agpat4 | 1-acylglycerol-3-phosphate O-acyltransferase 4 (lysophosphatidic acid acyltransferase, delta) | -2.27 |
| ENSMUSG00000029275 | Gfi1 | growth factor independent 1 | -2.27 |
| ENSMUSG00000105456 | Gm43745 | predicted gene 43745 | -2.27 |

| | | | |
|---|---|---|---|
| ENSMUSG00000097316 | Gm10516 | predicted gene 10516 | -2.27 |
| ENSMUSG00000109372 | Gm19410 | predicted gene, 19410 | -2.27 |
| ENSMUSG00000017688 | Hnf4g | hepatocyte nuclear factor 4, gamma | -2.28 |
| ENSMUSG00000024277 | Mapre2 | microtubule-associated protein, RP/EB family, member 2 | -2.28 |
| ENSMUSG00000040537 | Adam22 | a disintegrin and metallopeptidase domain 22 | -2.28 |
| ENSMUSG00000105986 | Gm43065 | predicted gene 43065 | -2.28 |
| ENSMUSG00000037408 | Cnnm4 | cyclin M4 | -2.29 |
| ENSMUSG00000024340 | Btnl2 | butyrophilin-like 2 | -2.29 |
| ENSMUSG00000025401 | Myo1a | myosin IA | -2.30 |
| ENSMUSG00000034449 | Dhrs11 | dehydrogenase/reductase (SDR family) member 11 | -2.30 |
| ENSMUSG00000085289 | Gm15337 | predicted gene 15337 | -2.30 |
| ENSMUSG00000037033 | Clca3b | chloride channel accessory 3B | -2.31 |
| ENSMUSG00000035699 | Slc51a | solute carrier family 51, alpha subunit | -2.31 |
| ENSMUSG00000028940 | Hes2 | hes family bHLH transcription factor 2 | -2.31 |
| ENSMUSG00000089635 | Gm16559 | predicted gene 16559 | -2.31 |
| ENSMUSG00000031618 | Nr3c2 | nuclear receptor subfamily 3, group C, member 2 | -2.32 |
| ENSMUSG00000096299 | Gm21814 | predicted gene, 21814 | -2.32 |
| ENSMUSG00000110557 | Gm5159 | predicted gene 5159 | -2.32 |
| ENSMUSG00000082677 | Gm15371 | predicted gene 15371 | -2.32 |
| ENSMUSG00000043501 | Lgals2 | lectin, galactose-binding, soluble 2 | -2.33 |
| ENSMUSG00000027790 | Sis | sucrase isomaltase (alpha-glucosidase) | -2.33 |
| ENSMUSG00000049858 | Suox | sulfite oxidase | -2.33 |
| ENSMUSG00000027317 | Ppp1r14d | protein phosphatase 1, regulatory inhibitor subunit 14D | -2.33 |
| ENSMUSG00000055730 | Ces2a | carboxylesterase 2A | -2.33 |
| ENSMUSG00000110266 | Gm32742 | predicted gene, 32742 | -2.33 |
| ENSMUSG00000000308 | Ckmt1 | creatine kinase, mitochondrial 1, ubiquitous | -2.34 |
| ENSMUSG00000021640 | Naip1 | NLR family, apoptosis inhibitory protein 1 | -2.34 |
| ENSMUSG00000038751 | Ptk6 | PTK6 protein tyrosine kinase 6 | -2.34 |
| ENSMUSG00000073403 | Gm10499 | predicted gene 10499 | -2.34 |
| ENSMUSG00000023328 | Ache | acetylcholinesterase | -2.34 |
| ENSMUSG00000059213 | Ddn | dendrin | -2.34 |
| ENSMUSG00000043822 | Adamtsl5 | ADAMTS-like 5 | -2.35 |
| ENSMUSG00000027173 | Depdc7 | DEP domain containing 7 | -2.35 |
| ENSMUSG00000113680 | Gm47405 | predicted gene, 47405 | -2.35 |
| ENSMUSG00000032726 | Bmp8a | bone morphogenetic protein 8a | -2.35 |
| ENSMUSG00000018126 | Baiap2l2 | BAI1-associated protein 2-like 2 | -2.36 |
| ENSMUSG00000092618 | Btnl6 | butyrophilin-like 6 | -2.36 |
| ENSMUSG00000005373 | Mlxipl | MLX interacting protein-like | -2.36 |
| ENSMUSG00000033308 | Dpyd | dihydropyrimidine dehydrogenase | -2.36 |
| ENSMUSG00000040767 | Snrnp25 | small nuclear ribonucleoprotein 25 (U11/U12) | -2.36 |
| ENSMUSG00000049307 | Fut4 | fucosyltransferase 4 | -2.36 |
| ENSMUSG00000050097 | Ces2b | carboxyesterase 2B | -2.36 |
| ENSMUSG00000087615 | Pnpla1os | patatin-like phospholipase domain containing 1, opposite strand | -2.36 |
| ENSMUSG00000052415 | Tchh | trichohyalin | -2.37 |
| ENSMUSG00000034427 | Myo15b | myosin XVB | -2.38 |
| ENSMUSG00000078439 | Smim24 | small integral membrane protein 24 | -2.38 |
| ENSMUSG00000037003 | Tns2 | tensin 2 | -2.38 |
| ENSMUSG00000032690 | Oas2 | 2'-5' oligoadenylate synthetase 2 | -2.38 |
| ENSMUSG00000050592 | Fam78a | family with sequence similarity 78, member A | -2.38 |
| ENSMUSG00000025726 | Slc28a1 | solute carrier family 28 (sodium-coupled nucleoside transporter), member 1 | -2.38 |
| ENSMUSG00000100738 | 2010106C02Rik | RIKEN cDNA 2010106C02 gene | -2.38 |
| ENSMUSG00000030366 | Ceacam12 | carcinoembryonic antigen-related cell adhesion molecule 12 | -2.38 |
| ENSMUSG00000029269 | Sult1b1 | sulfotransferase family 1B, member 1 | -2.39 |
| ENSMUSG00000036377 | C530008M17Rik | RIKEN cDNA C530008M17 gene | -2.39 |
| ENSMUSG00000043592 | Unc5cl | unc-5 family C-terminal like | -2.39 |
| ENSMUSG00000033871 | Ppargc1b | peroxisome proliferative activated receptor, gamma, coactivator 1 beta | -2.39 |
| ENSMUSG00000030683 | Sez6l2 | seizure related 6 homolog like 2 | -2.39 |
| ENSMUSG00000024619 | Cdx1 | caudal type homeobox 1 | -2.40 |
| ENSMUSG00000037280 | Galnt6 | polypeptide N-acetylgalactosaminyltransferase 6 | -2.40 |
| ENSMUSG00000021416 | Eci3 | enoyl-Coenzyme A delta isomerase 3 | -2.40 |
| ENSMUSG00000092517 | Art2a-ps | ADP-ribosyltransferase 2a, pseudogene | -2.40 |
| ENSMUSG00000047298 | Kcnv2 | potassium channel, subfamily V, member 2 | -2.40 |
| ENSMUSG00000030017 | Reg3g | regenerating islet-derived 3 gamma | -2.41 |
| ENSMUSG00000034528 | Hsd17b13 | hydroxysteroid (17-beta) dehydrogenase 13 | -2.41 |
| ENSMUSG00000039339 | Mfsd4b2 | major facilitator superfamily domain containing 4B2 | -2.41 |
| ENSMUSG00000014786 | Slc9a5 | solute carrier family 9 (sodium/hydrogen exchanger), member 5 | -2.41 |
| ENSMUSG00000049999 | Ppp1r3d | protein phosphatase 1, regulatory subunit 3D | -2.41 |
| ENSMUSG00000071265 | 1700086L19Rik | RIKEN cDNA 1700086L19 gene | -2.41 |
| ENSMUSG00000035561 | Aldh1b1 | aldehyde dehydrogenase 1 family, member B1 | -2.42 |
| ENSMUSG00000064213 | Defa24 | defensin, alpha, 24 | -2.42 |
| ENSMUSG00000005045 | Chd5 | chromodomain helicase DNA binding protein 5 | -2.42 |
| ENSMUSG00000028874 | Fgr | FGR proto-oncogene, Src family tyrosine kinase | -2.42 |
| ENSMUSG00000095328 | Defa-ps6 | defensin, alpha, pseudogene 6 | -2.43 |
| ENSMUSG00000042828 | Trim72 | tripartite motif-containing 72 | -2.43 |
| ENSMUSG00000081245 | Gm6587 | predicted gene 6587 | -2.43 |
| ENSMUSG00000105822 | Gm42969 | predicted gene 42969 | -2.43 |

| | | | |
|---|---|---|---|
| ENSMUSG00000050965 | Prkca | protein kinase C, alpha | -2.44 |
| ENSMUSG00000050982 | Apol10a | apolipoprotein L 10A | -2.44 |
| ENSMUSG00000006642 | Tcf23 | transcription factor 23 | -2.44 |
| ENSMUSG00000071356 | Reg3b | regenerating islet-derived 3 beta | -2.45 |
| ENSMUSG00000017978 | Cadps2 | Ca2+-dependent activator protein for secretion 2 | -2.45 |
| ENSMUSG00000054453 | Sytl5 | synaptotagmin-like 5 | -2.45 |
| ENSMUSG00000093458 | Gm20688 | predicted gene 20688 | -2.45 |
| ENSMUSG00000094091 | Gm21885 | predicted gene, 21885 | -2.46 |
| ENSMUSG00000057614 | Gnai1 | guanine nucleotide binding protein (G protein), alpha inhibiting 1 | -2.46 |
| ENSMUSG00000024957 | Kcnk4 | potassium channel, subfamily K, member 4 | -2.46 |
| ENSMUSG00000024548 | Setbp1 | SET binding protein 1 | -2.47 |
| ENSMUSG00000021278 | Amn | amnionless | -2.47 |
| ENSMUSG00000057880 | Abat | 4-aminobutyrate aminotransferase | -2.48 |
| ENSMUSG00000039981 | Zc3h12d | zinc finger CCCH type containing 12D | -2.48 |
| ENSMUSG00000021221 | Dpf3 | D4, zinc and double PHD fingers, family 3 | -2.48 |
| ENSMUSG00000107310 | Gm7452 | predicted pseudogene 7452 | -2.48 |
| ENSMUSG00000049598 | Vsig8 | V-set and immunoglobulin domain containing 8 | -2.48 |
| ENSMUSG00000026825 | Dnm1 | dynamin 1 | -2.49 |
| ENSMUSG00000039578 | Ccser1 | coiled-coil serine rich 1 | -2.49 |
| ENSMUSG00000079445 | B3gnt7 | UDP-GlcNAc:betaGal beta-1,3-N-acetylglucosaminyltransferase 7 | -2.49 |
| ENSMUSG00000031937 | Vstm5 | V-set and transmembrane domain containing 5 | -2.49 |
| ENSMUSG00000025876 | Unc5a | unc-5 netrin receptor A | -2.49 |
| ENSMUSG00000060208 | Defa17 | defensin, alpha, 17 | -2.49 |
| ENSMUSG00000040770 | Il25 | interleukin 25 | -2.49 |
| ENSMUSG00000024743 | Syt7 | synaptotagmin VII | -2.50 |
| ENSMUSG00000073399 | Trim40 | tripartite motif-containing 40 | -2.50 |
| ENSMUSG00000024731 | Ms4a10 | membrane-spanning 4-domains, subfamily A, member 10 | -2.50 |
| ENSMUSG00000071551 | Akr1c19 | aldo-keto reductase family 1, member C19 | -2.50 |
| ENSMUSG00000047528 | Als2cr12 | amyotrophic lateral sclerosis 2 chromosome region 12 | -2.50 |
| ENSMUSG00000112489 | 9230116L04Rik | RIKEN cDNA 9230116L04 gene | -2.50 |
| ENSMUSG00000012777 | Acp4 | acid phosphatase 4 | -2.50 |
| ENSMUSG00000066178 | 6030445D17Rik | RIKEN cDNA 6030445D17 gene | -2.50 |
| ENSMUSG00000063142 | Kcnma1 | potassium large conductance calcium-activated channel, subfamily M, alpha member 1 | -2.51 |
| ENSMUSG00000054640 | Slc8a1 | solute carrier family 8 (sodium/calcium exchanger), member 1 | -2.51 |
| ENSMUSG00000031883 | Car7 | carbonic anhydrase 7 | -2.51 |
| ENSMUSG00000026417 | Pigr | polymeric immunoglobulin receptor | -2.52 |
| ENSMUSG00000038515 | Grtp1 | GH regulated TBC protein 1 | -2.52 |
| ENSMUSG00000030472 | Ceacam18 | carcinoembryonic antigen-related cell adhesion molecule 18 | -2.52 |
| ENSMUSG00000108688 | Gm44985 | predicted gene 44985 | -2.52 |
| ENSMUSG00000047878 | A4galt | alpha 1,4-galactosyltransferase | -2.52 |
| ENSMUSG00000060703 | Cd302 | CD302 antigen | -2.52 |
| ENSMUSG00000019906 | Lin7a | lin-7 homolog A (C. elegans) | -2.52 |
| ENSMUSG00000031621 | Isx | intestine specific homeobox | -2.53 |
| ENSMUSG00000025347 | Mettl7b | methyltransferase like 7B | -2.53 |
| ENSMUSG00000053004 | Hrh1 | histamine receptor H1 | -2.53 |
| ENSMUSG00000005360 | Slc1a3 | solute carrier family 1 (glial high affinity glutamate transporter), member 3 | -2.53 |
| ENSMUSG00000032034 | Kcnj5 | potassium inwardly-rectifying channel, subfamily J, member 5 | -2.53 |
| ENSMUSG00000101814 | Gm17807 | predicted gene, 17807 | -2.54 |
| ENSMUSG00000025037 | Maoa | monoamine oxidase A | -2.55 |
| ENSMUSG00000036814 | Slc6a20a | solute carrier family 6 (neurotransmitter transporter), member 20A | -2.55 |
| ENSMUSG00000040471 | Ggt6 | gamma-glutamyltransferase 6 | -2.55 |
| ENSMUSG00000110573 | Gm5485 | predicted gene 5485 | -2.55 |
| ENSMUSG00000102140 | Gm38058 | predicted gene, 38058 | -2.55 |
| ENSMUSG00000025207 | Sema4g | sema domain, immunoglobulin domain (Ig), transmembrane domain (TM) and short cytoplasmic do | -2.56 |
| ENSMUSG00000005672 | Kit | KIT proto-oncogene receptor tyrosine kinase | -2.56 |
| ENSMUSG00000060002 | Chpt1 | choline phosphotransferase 1 | -2.56 |
| ENSMUSG00000057897 | Camk2b | calcium/calmodulin-dependent protein kinase II, beta | -2.56 |
| ENSMUSG00000028262 | Clca3a2 | chloride channel accessory 3A2 | -2.56 |
| ENSMUSG00000032098 | Treh | trehalase (brush-border membrane glycoprotein) | -2.56 |
| ENSMUSG00000109644 | 0610005C13Rik | RIKEN cDNA 0610005C13 gene | -2.56 |
| ENSMUSG00000061762 | Tac1 | tachykinin 1 | -2.56 |
| ENSMUSG00000110238 | Gm19269 | predicted gene, 19269 | -2.56 |
| ENSMUSG00000020072 | Pbld2 | phenazine biosynthesis-like protein domain containing 2 | -2.57 |
| ENSMUSG00000031445 | Proz | protein Z, vitamin K-dependent plasma glycoprotein | -2.57 |
| ENSMUSG00000075611 | Gm11545 | predicted gene 11545 | -2.57 |
| ENSMUSG00000022747 | St3gal6 | ST3 beta-galactoside alpha-2,3-sialyltransferase 6 | -2.58 |
| ENSMUSG00000032181 | Scg3 | secretogranin III | -2.58 |
| ENSMUSG00000004113 | Cacna1b | calcium channel, voltage-dependent, N type, alpha 1B subunit | -2.58 |
| ENSMUSG00000053964 | Lgals4 | lectin, galactose binding, soluble 4 | -2.59 |
| ENSMUSG00000037005 | Xpnpep2 | X-prolyl aminopeptidase (aminopeptidase P) 2, membrane-bound | -2.59 |
| ENSMUSG00000029162 | Khk | ketohexokinase | -2.59 |
| ENSMUSG00000084978 | Gm11655 | predicted gene 11655 | -2.59 |
| ENSMUSG00000075517 | Cyp2d37-ps | cytochrome P450, family 2, subfamily d, polypeptide 37, pseudogene | -2.59 |
| ENSMUSG00000032377 | Plscr4 | phospholipid scramblase 4 | -2.59 |
| ENSMUSG00000055567 | Unc80 | unc-80, NALCN activator | -2.59 |
| ENSMUSG00000108155 | Gm44443 | predicted gene, 44443 | -2.59 |

| | | | |
|---|---|---|---|
| ENSMUSG00000054128 | H2-T3 | histocompatibility 2, T region locus 3 | -2.60 |
| ENSMUSG00000058435 | Btnl4 | butyrophilin-like 4 | -2.60 |
| ENSMUSG00000071203 | Naip5 | NLR family, apoptosis inhibitory protein 5 | -2.61 |
| ENSMUSG00000029084 | Cd38 | CD38 antigen | -2.61 |
| ENSMUSG00000028179 | Cth | cystathionase (cystathionine gamma-lyase) | -2.61 |
| ENSMUSG00000040653 | Ppp1r14c | protein phosphatase 1, regulatory inhibitor subunit 14C | -2.61 |
| ENSMUSG00000086067 | Gm16183 | predicted gene 16183 | -2.61 |
| ENSMUSG00000050296 | Abca12 | ATP-binding cassette, sub-family A (ABC1), member 12 | -2.62 |
| ENSMUSG00000072941 | Sod3 | superoxide dismutase 3, extracellular | -2.62 |
| ENSMUSG00000022947 | Cbr3 | carbonyl reductase 3 | -2.62 |
| ENSMUSG00000013643 | Lypd8 | LY6/PLAUR domain containing 8 | -2.63 |
| ENSMUSG00000055114 | Anxa13 | annexin A13 | -2.63 |
| ENSMUSG00000030866 | Ern2 | endoplasmic reticulum (ER) to nucleus signalling 2 | -2.63 |
| ENSMUSG00000038910 | Plcl2 | phospholipase C-like 2 | -2.63 |
| ENSMUSG00000027876 | Reg4 | regenerating islet-derived family, member 4 | -2.63 |
| ENSMUSG00000014773 | Dll1 | delta like canonical Notch ligand 1 | -2.63 |
| ENSMUSG00000107524 | Gm44136 | predicted gene, 44136 | -2.63 |
| ENSMUSG00000033847 | Pla2g4c | phospholipase A2, group IVC (cytosolic, calcium-independent) | -2.64 |
| ENSMUSG00000013523 | Bcas1 | breast carcinoma amplified sequence 1 | -2.64 |
| ENSMUSG00000020080 | Hkdc1 | hexokinase domain containing 1 | -2.64 |
| ENSMUSG00000058921 | Slc10a5 | solute carrier family 10 (sodium/bile acid cotransporter family), member 5 | -2.64 |
| ENSMUSG00000027555 | Car13 | carbonic anhydrase 13 | -2.64 |
| ENSMUSG00000027297 | Ltk | leukocyte tyrosine kinase | -2.64 |
| ENSMUSG00000032038 | St3gal4 | ST3 beta-galactoside alpha-2,3-sialyltransferase 4 | -2.65 |
| ENSMUSG00000026126 | Ptpn18 | protein tyrosine phosphatase, non-receptor type 18 | -2.65 |
| ENSMUSG00000021208 | Ifi27l2b | interferon, alpha-inducible protein 27 like 2B | -2.65 |
| ENSMUSG00000073420 | Btnl5-ps | butyrophilin-like 5, pseudogene | -2.65 |
| ENSMUSG00000047638 | Nr1h4 | nuclear receptor subfamily 1, group H, member 4 | -2.65 |
| ENSMUSG00000023235 | Ccl25 | chemokine (C-C motif) ligand 25 | -2.66 |
| ENSMUSG00000060508 | Nlrp9b | NLR family, pyrin domain containing 9B | -2.66 |
| ENSMUSG00000074300 | BC030870 | cDNA sequence BC030870 | -2.66 |
| ENSMUSG00000116929 | AC164088.2 | TEC | -2.66 |
| ENSMUSG00000070315 | 4930581F22Rik | RIKEN cDNA 4930581F22 gene | -2.67 |
| ENSMUSG00000022340 | Sybu | syntabulin (syntaxin-interacting) | -2.67 |
| ENSMUSG00000037451 | Slc22a20 | solute carrier family 22 (organic anion transporter), member 20 | -2.67 |
| ENSMUSG00000091387 | Gcnt4 | glucosaminyl (N-acetyl) transferase 4, core 2 (beta-1,6-N-acetylglucosaminyltransferase) | -2.68 |
| ENSMUSG00000014361 | Mertk | c-mer proto-oncogene tyrosine kinase | -2.68 |
| ENSMUSG00000085042 | Abhd11os | abhydrolase domain containing 11, opposite strand | -2.68 |
| ENSMUSG00000026489 | Coq8a | coenzyme Q8A | -2.68 |
| ENSMUSG00000070034 | Sp110 | Sp110 nuclear body protein | -2.68 |
| ENSMUSG00000096914 | Galntl6 | UDP-N-acetyl-alpha-D-galactosamine:polypeptide N-acetylgalactosaminyltransferase-like 6 | -2.68 |
| ENSMUSG00000032297 | Celf6 | CUGBP, Elav-like family member 6 | -2.68 |
| ENSMUSG00000075702 | Selenom | selenoprotein M | -2.69 |
| ENSMUSG00000024225 | Clps | colipase, pancreatic | -2.69 |
| ENSMUSG00000074067 | Gm10619 | predicted gene 10619 | -2.69 |
| ENSMUSG00000046410 | Kcnk6 | potassium inwardly-rectifying channel, subfamily K, member 6 | -2.70 |
| ENSMUSG00000059654 | Reg1 | regenerating islet-derived 1 | -2.70 |
| ENSMUSG00000021947 | Cryl1 | crystallin, lambda 1 | -2.71 |
| ENSMUSG00000031163 | Glod5 | glyoxalase domain containing 5 | -2.71 |
| ENSMUSG00000071335 | Mfsd4b3 | major facilitator superfamily domain containing 4B3 | -2.71 |
| ENSMUSG00000027107 | Chrna1 | cholinergic receptor, nicotinic, alpha polypeptide 1 (muscle) | -2.71 |
| ENSMUSG00000046908 | Ltb4r1 | leukotriene B4 receptor 1 | -2.71 |
| ENSMUSG00000040703 | Cyp2s1 | cytochrome P450, family 2, subfamily s, polypeptide 1 | -2.72 |
| ENSMUSG00000091575 | 2010016I18Rik | RIKEN cDNA 2010016I18 gene | -2.72 |
| ENSMUSG00000031613 | Hpgd | hydroxyprostaglandin dehydrogenase 15 (NAD) | -2.73 |
| ENSMUSG00000020838 | Slc6a4 | solute carrier family 6 (neurotransmitter transporter, serotonin), member 4 | -2.73 |
| ENSMUSG00000032357 | Tinag | tubulointerstitial nephritis antigen | -2.73 |
| ENSMUSG00000027792 | Bche | butyrylcholinesterase | -2.74 |
| ENSMUSG00000057719 | Sh3rf2 | SH3 domain containing ring finger 2 | -2.74 |
| ENSMUSG00000028427 | Aqp7 | aquaporin 7 | -2.74 |
| ENSMUSG00000116010 | Gm36026 | predicted gene, 36026 | -2.74 |
| ENSMUSG00000063296 | Tmem117 | transmembrane protein 117 | -2.75 |
| ENSMUSG00000044352 | Sowaha | sosondowah ankyrin repeat domain family member A | -2.75 |
| ENSMUSG00000067377 | Tspan6 | tetraspanin 6 | -2.75 |
| ENSMUSG00000038745 | Nlrp6 | NLR family, pyrin domain containing 6 | -2.76 |
| ENSMUSG00000006731 | B4galnt1 | beta-1,4-N-acetyl-galactosaminyl transferase 1 | -2.76 |
| ENSMUSG00000061531 | Tmem236 | transmembrane protein 236 | -2.76 |
| ENSMUSG00000022235 | Cmbl | carboxymethylenebutenolidase-like (Pseudomonas) | -2.76 |
| ENSMUSG00000043705 | Capn13 | calpain 13 | -2.77 |
| ENSMUSG00000041741 | Pde3a | phosphodiesterase 3A, cGMP inhibited | -2.77 |
| ENSMUSG00000031364 | Grpr | gastrin releasing peptide receptor | -2.77 |
| ENSMUSG00000032528 | Vipr1 | vasoactive intestinal peptide receptor 1 | -2.78 |
| ENSMUSG00000020105 | Lrig3 | leucine-rich repeats and immunoglobulin-like domains 3 | -2.78 |
| ENSMUSG00000045775 | Slc16a5 | solute carrier family 16 (monocarboxylic acid transporters), member 5 | -2.78 |
| ENSMUSG00000060176 | Kif27 | kinesin family member 27 | -2.78 |
| ENSMUSG00000024730 | Ms4a8a | membrane-spanning 4-domains, subfamily A, member 8A | -2.79 |

| | | | |
|---|---|---|---|
| ENSMUSG00000032226 | Gcnt3 | glucosaminyl (N-acetyl) transferase 3, mucin type | -2.80 |
| ENSMUSG00000029330 | Cds1 | CDP-diacylglycerol synthase 1 | -2.80 |
| ENSMUSG00000024411 | Aqp4 | aquaporin 4 | -2.80 |
| ENSMUSG00000036298 | Slc2a13 | solute carrier family 2 (facilitated glucose transporter), member 13 | -2.80 |
| ENSMUSG00000033174 | Mgll | monoglyceride lipase | -2.80 |
| ENSMUSG00000021396 | Nxnl2 | nucleoredoxin-like 2 | -2.80 |
| ENSMUSG00000004655 | Aqp1 | aquaporin 1 | -2.81 |
| ENSMUSG00000022755 | Adgrg7 | adhesion G protein-coupled receptor G7 | -2.81 |
| ENSMUSG00000081303 | Gm16011 | predicted gene 16011 | -2.81 |
| ENSMUSG00000026870 | Cutal | cutA divalent cation tolerance homolog-like | -2.81 |
| ENSMUSG00000029420 | Rimbp2 | RIMS binding protein 2 | -2.81 |
| ENSMUSG00000057914 | Cacnb2 | calcium channel, voltage-dependent, beta 2 subunit | -2.81 |
| ENSMUSG00000090700 | Cyp4f40 | cytochrome P450, family 4, subfamily f, polypeptide 40 | -2.83 |
| ENSMUSG00000073413 | Ly6g6d | lymphocyte antigen 6 complex, locus G6D | -2.83 |
| ENSMUSG00000041644 | Slc5a12 | solute carrier family 5 (sodium/glucose cotransporter), member 12 | -2.83 |
| ENSMUSG00000027420 | Bfsp1 | beaded filament structural protein 1, in lens-CP94 | -2.84 |
| ENSMUSG00000028278 | Rragd | Ras-related GTP binding D | -2.84 |
| ENSMUSG00000027224 | Duoxa1 | dual oxidase maturation factor 1 | -2.84 |
| ENSMUSG00000097445 | Gm26631 | predicted gene, 26631 | -2.84 |
| ENSMUSG00000021216 | Tubal3 | tubulin, alpha-like 3 | -2.85 |
| ENSMUSG00000047115 | Fam221a | family with sequence similarity 221, member A | -2.85 |
| ENSMUSG00000039419 | Cntnap2 | contactin associated protein-like 2 | -2.85 |
| ENSMUSG00000045005 | Fzd5 | frizzled class receptor 5 | -2.86 |
| ENSMUSG00000053182 | Gm609 | predicted gene 609 | -2.86 |
| ENSMUSG00000006362 | Cbfa2t3 | core-binding factor, runt domain, alpha subunit 2, translocated to, 3 (human) | -2.86 |
| ENSMUSG00000051726 | Kcnf1 | potassium voltage-gated channel, subfamily F, member 1 | -2.86 |
| ENSMUSG00000071633 | Gm4952 | predicted gene 4952 | -2.86 |
| ENSMUSG00000041301 | Cftr | cystic fibrosis transmembrane conductance regulator | -2.87 |
| ENSMUSG00000075520 | Malrd1 | MAM and LDL receptor class A domain containing 1 | -2.87 |
| ENSMUSG00000019989 | Enpp3 | ectonucleotide pyrophosphatase/phosphodiesterase 3 | -2.87 |
| ENSMUSG00000087430 | Gm13405 | predicted gene 13405 | -2.87 |
| ENSMUSG00000026380 | Tfcp2l1 | transcription factor CP2-like 1 | -2.88 |
| ENSMUSG00000040412 | 5330417C22Rik | RIKEN cDNA 5330417C22 gene | -2.88 |
| ENSMUSG00000032033 | Barx2 | BarH-like homeobox 2 | -2.88 |
| ENSMUSG00000036912 | Piwil4 | piwi-like RNA-mediated gene silencing 4 | -2.88 |
| ENSMUSG00000046804 | Phgr1 | proline/histidine/glycine-rich 1 | -2.89 |
| ENSMUSG00000037686 | Aspg | asparaginase | -2.89 |
| ENSMUSG00000108388 | Gm44673 | predicted gene 44673 | -2.89 |
| ENSMUSG00000109446 | Gm9195 | predicted gene 9195 | -2.89 |
| ENSMUSG00000029161 | Cgref1 | cell growth regulator with EF hand domain 1 | -2.90 |
| ENSMUSG00000054385 | Ceacam2 | carcinoembryonic antigen-related cell adhesion molecule 2 | -2.90 |
| ENSMUSG00000027843 | Ptpn22 | protein tyrosine phosphatase, non-receptor type 22 (lymphoid) | -2.90 |
| ENSMUSG00000035849 | Krt222 | keratin 222 | -2.90 |
| ENSMUSG00000027669 | Gnb4 | guanine nucleotide binding protein (G protein), beta 4 | -2.90 |
| ENSMUSG00000058063 | Trim31 | tripartite motif-containing 31 | -2.91 |
| ENSMUSG00000116220 | A430088P11Rik | RIKEN cDNA A430088P11 gene | -2.91 |
| ENSMUSG00000036422 | Pcdh8 | protocadherin 8 | -2.91 |
| ENSMUSG00000021194 | Chga | chromogranin A | -2.91 |
| ENSMUSG00000041261 | Car8 | carbonic anhydrase 8 | -2.92 |
| ENSMUSG00000024620 | Pdgfrb | platelet derived growth factor receptor, beta polypeptide | -2.92 |
| ENSMUSG00000087389 | Gm15592 | predicted gene 15592 | -2.92 |
| ENSMUSG00000094840 | Muc3a | mucin 3A, cell surface associated | -2.93 |
| ENSMUSG00000005268 | Prlr | prolactin receptor | -2.93 |
| ENSMUSG00000049134 | Nrap | nebulin-related anchoring protein | -2.93 |
| ENSMUSG00000103579 | Gm37113 | predicted gene, 37113 | -2.93 |
| ENSMUSG00000090291 | Lrrc10b | leucine rich repeat containing 10B | -2.93 |
| ENSMUSG00000078670 | Fam174b | family with sequence similarity 174, member B | -2.94 |
| ENSMUSG00000029134 | Plb1 | phospholipase B1 | -2.94 |
| ENSMUSG00000046613 | Vwa5b2 | von Willebrand factor A domain containing 5B2 | -2.94 |
| ENSMUSG00000073608 | Gal3st2c | galactose-3-O-sulfotransferase 2C | -2.94 |
| ENSMUSG00000018919 | Tm4sf5 | transmembrane 4 superfamily member 5 | -2.95 |
| ENSMUSG00000041696 | Rasl12 | RAS-like, family 12 | -2.95 |
| ENSMUSG00000015224 | Cyp2j9 | cytochrome P450, family 2, subfamily j, polypeptide 9 | -2.95 |
| ENSMUSG00000001739 | Cldn15 | claudin 15 | -2.96 |
| ENSMUSG00000010066 | Cacna2d2 | calcium channel, voltage-dependent, alpha 2/delta subunit 2 | -2.96 |
| ENSMUSG00000028327 | Stra6l | STRA6-like | -2.97 |
| ENSMUSG00000031451 | Gas6 | growth arrest specific 6 | -2.98 |
| ENSMUSG00000026398 | Nr5a2 | nuclear receptor subfamily 5, group A, member 2 | -2.98 |
| ENSMUSG00000030340 | Scnn1a | sodium channel, nonvoltage-gated 1 alpha | -2.98 |
| ENSMUSG00000021850 | ccdc198 | coiled-coil domain containing 198 | -2.98 |
| ENSMUSG00000091189 | Ear-ps3 | eosinophil-associated, ribonuclease A family, pseudogene 3 | -2.98 |
| ENSMUSG00000042284 | Itga1 | integrin alpha 1 | -2.99 |
| ENSMUSG00000001166 | Oas1c | 2'-5' oligoadenylate synthetase 1C | -2.99 |
| ENSMUSG00000015401 | Cltrn | collectrin, amino acid transport regulator | -2.99 |
| ENSMUSG00000030793 | Pycard | PYD and CARD domain containing | -3.00 |
| ENSMUSG00000094219 | Calhm3 | calcium homeostasis modulator 3 | -3.00 |

| | | | |
|---|---|---|---|
| ENSMUSG00000027870 | Hao2 | hydroxyacid oxidase 2 | -3.00 |
| ENSMUSG00000028251 | Tstd3 | thiosulfate sulfurtransferase (rhodanese)-like domain containing 3 | -3.01 |
| ENSMUSG00000036960 | Clca2 | chloride channel accessory 2 | -3.01 |
| ENSMUSG00000024379 | Tslp | thymic stromal lymphopoietin | -3.01 |
| ENSMUSG00000093867 | Gsdmcl2 | gasdermin C-like 2 | -3.02 |
| ENSMUSG00000022661 | Cd200 | CD200 antigen | -3.02 |
| ENSMUSG00000085723 | Gm15915 | predicted gene 15915 | -3.02 |
| ENSMUSG00000022479 | Vdr | vitamin D (1,25-dihydroxyvitamin D3) receptor | -3.03 |
| ENSMUSG00000022408 | Fam83f | family with sequence similarity 83, member F | -3.03 |
| ENSMUSG00000023032 | Slc4a8 | solute carrier family 4 (anion exchanger), member 8 | -3.03 |
| ENSMUSG00000022262 | Dnah5 | dynein, axonemal, heavy chain 5 | -3.03 |
| ENSMUSG00000056752 | Dnah9 | dynein, axonemal, heavy chain 9 | -3.04 |
| ENSMUSG00000027513 | Pck1 | phosphoenolpyruvate carboxykinase 1, cytosolic | -3.05 |
| ENSMUSG00000038917 | 3930402G23Rik | RIKEN cDNA 3930402G23 gene | -3.05 |
| ENSMUSG00000001225 | Slc26a3 | solute carrier family 26, member 3 | -3.06 |
| ENSMUSG00000090124 | Ugt1a7c | UDP glucuronosyltransferase 1 family, polypeptide A7C | -3.06 |
| ENSMUSG00000048022 | Tmem229a | transmembrane protein 229A | -3.06 |
| ENSMUSG00000059003 | Grin2a | glutamate receptor, ionotropic, NMDA2A (epsilon 1) | -3.06 |
| ENSMUSG00000028003 | Lrat | lecithin-retinol acyltransferase (phosphatidylcholine-retinol-O-acyltransferase) | -3.07 |
| ENSMUSG00000020865 | Abcc3 | ATP-binding cassette, sub-family C (CFTR/MRP), member 3 | -3.08 |
| ENSMUSG00000024935 | Slc1a1 | solute carrier family 1 (neuronal/epithelial high affinity glutamate transporter, system Xag), membe | -3.08 |
| ENSMUSG00000029490 | Mfsd7a | major facilitator superfamily domain containing 7A | -3.08 |
| ENSMUSG00000079355 | Ackr4 | atypical chemokine receptor 4 | -3.09 |
| ENSMUSG00000028525 | Pde4b | phosphodiesterase 4B, cAMP specific | -3.09 |
| ENSMUSG00000027350 | Chgb | chromogranin B | -3.09 |
| ENSMUSG00000025010 | Ccnj | cyclin J | -3.10 |
| ENSMUSG00000035506 | Slc12a8 | solute carrier family 12 (potassium/chloride transporters), member 8 | -3.10 |
| ENSMUSG00000034112 | Atp2c2 | ATPase, Ca++ transporting, type 2C, member 2 | -3.10 |
| ENSMUSG00000096751 | Gm28373 | predicted gene 28373 | -3.10 |
| ENSMUSG00000044748 | Defb1 | defensin beta 1 | -3.10 |
| ENSMUSG00000021749 | Oit1 | oncoprotein induced transcript 1 | -3.11 |
| ENSMUSG00000074207 | Adh1 | alcohol dehydrogenase 1 (class I) | -3.12 |
| ENSMUSG00000028457 | Atp8b5 | ATPase, class I, type 8B, member 5 | -3.12 |
| ENSMUSG00000055027 | Smyd1 | SET and MYND domain containing 1 | -3.13 |
| ENSMUSG00000024371 | C2 | complement component 2 (within H-2S) | -3.14 |
| ENSMUSG00000085941 | Gm11201 | predicted gene 11201 | -3.14 |
| ENSMUSG00000116288 | Gm41349 | predicted gene, 41349 | -3.14 |
| ENSMUSG00000060615 | Ang4 | angiogenin, ribonuclease A family, member 4 | -3.15 |
| ENSMUSG00000098557 | Kctd12 | potassium channel tetramerisation domain containing 12 | -3.15 |
| ENSMUSG00000073406 | H2-Bl | histocompatibility 2, blastocyst | -3.15 |
| ENSMUSG00000033740 | St18 | suppression of tumorigenicity 18 | -3.16 |
| ENSMUSG00000040896 | Kcnd3 | potassium voltage-gated channel, Shal-related family, member 3 | -3.16 |
| ENSMUSG00000070424 | Art5 | ADP-ribosyltransferase 5 | -3.16 |
| ENSMUSG00000103356 | Gm37595 | predicted gene, 37595 | -3.17 |
| ENSMUSG00000108260 | Gm3793 | predicted gene 3793 | -3.17 |
| ENSMUSG00000027254 | Map1a | microtubule-associated protein 1 A | -3.18 |
| ENSMUSG00000039878 | Slc39a5 | solute carrier family 39 (metal ion transporter), member 5 | -3.18 |
| ENSMUSG00000094584 | Ms4a18 | membrane-spanning 4-domains, subfamily A, member 18 | -3.18 |
| ENSMUSG00000061414 | Cracr2a | calcium release activated channel regulator 2A | -3.19 |
| ENSMUSG00000034057 | Myrfl | myelin regulatory factor-like | -3.20 |
| ENSMUSG00000090125 | Pou3f1 | POU domain, class 3, transcription factor 1 | -3.20 |
| ENSMUSG00000042770 | Hebp1 | heme binding protein 1 | -3.21 |
| ENSMUSG00000038580 | Sct | secretin | -3.22 |
| ENSMUSG00000072623 | Zfp9 | zinc finger protein 9 | -3.22 |
| ENSMUSG00000027359 | Slc27a2 | solute carrier family 27 (fatty acid transporter), member 2 | -3.23 |
| ENSMUSG00000078942 | Naip6 | NLR family, apoptosis inhibitory protein 6 | -3.24 |
| ENSMUSG00000033715 | Akr1c14 | aldo-keto reductase family 1, member C14 | -3.25 |
| ENSMUSG00000049154 | Fam183b | family with sequence similarity 183, member B | -3.25 |
| ENSMUSG00000001604 | Tcea3 | transcription elongation factor A (SII), 3 | -3.26 |
| ENSMUSG00000029273 | Sult1d1 | sulfotransferase family 1D, member 1 | -3.26 |
| ENSMUSG00000022383 | Ppara | peroxisome proliferator activated receptor alpha | -3.27 |
| ENSMUSG00000030786 | Itgam | integrin alpha M | -3.27 |
| ENSMUSG00000072723 | Gm10044 | predicted gene 10044 | -3.28 |
| ENSMUSG00000030303 | Far2 | fatty acyl CoA reductase 2 | -3.29 |
| ENSMUSG00000027474 | Ccm2l | cerebral cavernous malformation 2-like | -3.29 |
| ENSMUSG00000023247 | Guca2a | guanylate cyclase activator 2a (guanylin) | -3.30 |
| ENSMUSG00000030560 | Ctsc | cathepsin C | -3.30 |
| ENSMUSG00000029095 | Ablim2 | actin-binding LIM protein 2 | -3.30 |
| ENSMUSG00000049892 | Rasd1 | RAS, dexamethasone-induced 1 | -3.30 |
| ENSMUSG00000021806 | Nid2 | nidogen 2 | -3.30 |
| ENSMUSG00000005640 | Insrr | insulin receptor-related receptor | -3.30 |
| ENSMUSG00000021675 | F2rl2 | coagulation factor II (thrombin) receptor-like 2 | -3.31 |
| ENSMUSG00000108818 | Gm44869 | predicted gene 44869 | -3.32 |
| ENSMUSG00000052026 | Slc6a7 | solute carrier family 6 (neurotransmitter transporter, L-proline), member 7 | -3.33 |
| ENSMUSG00000033453 | Adamts15 | a disintegrin-like and metallopeptidase (reprolysin type) with thrombospondin type 1 motif, 15 | -3.33 |
| ENSMUSG00000114576 | Naip3-ps1 | NLR family, apoptosis inhibitory protein 3, pseudogene 1 | -3.33 |

| | | | |
|---|---|---|---|
| ENSMUSG00000042388 | Dlgap3 | DLG associated protein 3 | -3.33 |
| ENSMUSG00000050556 | Kcnb1 | potassium voltage gated channel, Shab-related subfamily, member 1 | -3.33 |
| ENSMUSG00000096852 | Cyp2d12 | cytochrome P450, family 2, subfamily d, polypeptide 12 | -3.33 |
| ENSMUSG00000071424 | Grid2 | glutamate receptor, ionotropic, delta 2 | -3.33 |
| ENSMUSG00000031551 | Ido1 | indoleamine 2,3-dioxygenase 1 | -3.34 |
| ENSMUSG00000048373 | Fgfbp1 | fibroblast growth factor binding protein 1 | -3.34 |
| ENSMUSG00000061808 | Ttr | transthyretin | -3.34 |
| ENSMUSG00000048217 | Nags | N-acetylglutamate synthase | -3.35 |
| ENSMUSG00000020120 | Plek | pleckstrin | -3.35 |
| ENSMUSG00000044827 | Tlr1 | toll-like receptor 1 | -3.35 |
| ENSMUSG00000056973 | Ces1d | carboxylesterase 1D | -3.35 |
| ENSMUSG00000031444 | F10 | coagulation factor X | -3.35 |
| ENSMUSG00000011008 | Mcoln2 | mucolipin 2 | -3.36 |
| ENSMUSG00000031725 | Ces1f | carboxylesterase 1F | -3.36 |
| ENSMUSG00000035200 | Chrnb4 | cholinergic receptor, nicotinic, beta polypeptide 4 | -3.37 |
| ENSMUSG00000074441 | Gm15292 | predicted gene 15292 | -3.39 |
| ENSMUSG00000068323 | Slc4a5 | solute carrier family 4, sodium bicarbonate cotransporter, member 5 | -3.39 |
| ENSMUSG00000020805 | Slc13a5 | solute carrier family 13 (sodium-dependent citrate transporter), member 5 | -3.40 |
| ENSMUSG00000027801 | Tm4sf4 | transmembrane 4 superfamily member 4 | -3.41 |
| ENSMUSG00000027661 | Slc2a10 | solute carrier family 2 (facilitated glucose transporter), member 10 | -3.41 |
| ENSMUSG00000108169 | Gm43958 | predicted gene, 43958 | -3.41 |
| ENSMUSG00000110397 | Gm45540 | predicted gene 45540 | -3.42 |
| ENSMUSG00000027983 | Cyp2u1 | cytochrome P450, family 2, subfamily u, polypeptide 1 | -3.43 |
| ENSMUSG00000020589 | Fam49a | family with sequence similarity 49, member A | -3.43 |
| ENSMUSG00000074437 | Defa29 | defensin, alpha, 29 | -3.44 |
| ENSMUSG00000114443 | Gm19241 | predicted gene, 19241 | -3.44 |
| ENSMUSG00000032015 | Pou2f3 | POU domain, class 2, transcription factor 3 | -3.45 |
| ENSMUSG00000041794 | Myrip | myosin VIIA and Rab interacting protein | -3.46 |
| ENSMUSG00000023092 | Fhl1 | four and a half LIM domains 1 | -3.46 |
| ENSMUSG00000104841 | Gm17743 | predicted gene, 17743 | -3.46 |
| ENSMUSG00000042351 | Grap2 | GRB2-related adaptor protein 2 | -3.46 |
| ENSMUSG00000026610 | Esrrg | estrogen-related receptor gamma | -3.47 |
| ENSMUSG00000097254 | C430042M11Rik | RIKEN cDNA C430042M11 gene | -3.47 |
| ENSMUSG00000015966 | Il17rb | interleukin 17 receptor B | -3.48 |
| ENSMUSG00000020732 | Rab37 | RAB37, member RAS oncogene family | -3.48 |
| ENSMUSG00000026811 | St6galnac6 | ST6 (alpha-N-acetyl-neuraminyl-2,3-beta-galactosyl-1,3)-N-acetylgalactosaminide alpha-2,6-sialyltra | -3.49 |
| ENSMUSG00000052861 | Dnah6 | dynein, axonemal, heavy chain 6 | -3.49 |
| ENSMUSG00000107425 | Gm44216 | predicted gene, 44216 | -3.49 |
| ENSMUSG00000038665 | Dgki | diacylglycerol kinase, iota | -3.50 |
| ENSMUSG00000024215 | Spdef | SAM pointed domain containing ets transcription factor | -3.50 |
| ENSMUSG00000048473 | Sult6b2 | sulfotransferase family 6B, member 2 | -3.51 |
| ENSMUSG00000044860 | Gm1123 | predicted gene 1123 | -3.52 |
| ENSMUSG00000002565 | Scin | scinderin | -3.52 |
| ENSMUSG00000022840 | Adcy5 | adenylate cyclase 5 | -3.52 |
| ENSMUSG00000026834 | Acvr1c | activin A receptor, type IC | -3.52 |
| ENSMUSG00000064036 | Mro | maestro | -3.52 |
| ENSMUSG00000020434 | 4921536K21Rik | RIKEN cDNA 4921536K21 gene | -3.52 |
| ENSMUSG00000074715 | Ccl28 | chemokine (C-C motif) ligand 28 | -3.53 |
| ENSMUSG00000031410 | Nxf7 | nuclear RNA export factor 7 | -3.53 |
| ENSMUSG00000021730 | Hcn1 | hyperpolarization-activated, cyclic nucleotide-gated K+ 1 | -3.53 |
| ENSMUSG00000082485 | Gm15693 | predicted gene 15693 | -3.53 |
| ENSMUSG00000105975 | Gm42609 | predicted gene 42609 | -3.54 |
| ENSMUSG00000032202 | Rab27a | RAB27A, member RAS oncogene family | -3.55 |
| ENSMUSG00000116383 | Apol10c-ps | apolipoprotein L 10C, pseudogene | -3.55 |
| ENSMUSG00000095457 | Gm8989 | predicted gene 8989 | -3.55 |
| ENSMUSG00000008193 | Spib | Spi-B transcription factor (Spi-1/PU.1 related) | -3.56 |
| ENSMUSG00000036492 | Rnf39 | ring finger protein 39 | -3.56 |
| ENSMUSG00000017311 | Pyy | peptide YY | -3.56 |
| ENSMUSG00000045930 | Clec14a | C-type lectin domain family 14, member a | -3.56 |
| ENSMUSG00000021624 | Cd180 | CD180 antigen | -3.57 |
| ENSMUSG00000105105 | Gm31363 | predicted gene, 31363 | -3.57 |
| ENSMUSG00000030302 | Atp2b2 | ATPase, Ca++ transporting, plasma membrane 2 | -3.59 |
| ENSMUSG00000042115 | Klhdc8a | kelch domain containing 8A | -3.59 |
| ENSMUSG00000103320 | Gm37403 | predicted gene, 37403 | -3.59 |
| ENSMUSG00000056293 | Gsdmc2 | gasdermin C2 | -3.60 |
| ENSMUSG00000078776 | 9530053A07Rik | RIKEN cDNA 9530053A07 gene | -3.60 |
| ENSMUSG00000072849 | Serpina1e | serine (or cysteine) peptidase inhibitor, clade A, member 1E | -3.60 |
| ENSMUSG00000066687 | Zbtb16 | zinc finger and BTB domain containing 16 | -3.61 |
| ENSMUSG00000038599 | Capn8 | calpain 8 | -3.62 |
| ENSMUSG00000060063 | Alox5ap | arachidonate 5-lipoxygenase activating protein | -3.62 |
| ENSMUSG00000030228 | Pik3c2g | phosphatidylinositol-4-phosphate 3-kinase catalytic subunit type 2 gamma | -3.62 |
| ENSMUSG00000021700 | Rab3c | RAB3C, member RAS oncogene family | -3.63 |
| ENSMUSG00000050100 | Hmx2 | H6 homeobox 2 | -3.64 |
| ENSMUSG00000068154 | Insm1 | insulinoma-associated 1 | -3.64 |
| ENSMUSG00000025738 | Fbxl16 | F-box and leucine-rich repeat protein 16 | -3.64 |
| ENSMUSG00000061958 | Gm14851 | predicted gene 14851 | -3.65 |

| | | | |
|---|---|---|---|
| ENSMUSG00000106651 | Gm42608 | predicted gene 42608 | -3.65 |
| ENSMUSG00000107296 | Gm43500 | predicted gene 43500 | -3.65 |
| ENSMUSG00000027797 | Dclk1 | doublecortin-like kinase 1 | -3.66 |
| ENSMUSG00000090062 | Galnt6os | polypeptide N-acetylgalactosaminyltransferase 6, opposite strand | -3.66 |
| ENSMUSG00000085184 | 4933439K11Rik | RIKEN cDNA 4933439K11 gene | -3.66 |
| ENSMUSG00000020901 | Pik3r5 | phosphoinositide-3-kinase regulatory subunit 5 | -3.67 |
| ENSMUSG00000060224 | Pyroxd2 | pyridine nucleotide-disulphide oxidoreductase domain 2 | -3.67 |
| ENSMUSG00000020279 | Il9r | interleukin 9 receptor | -3.67 |
| ENSMUSG00000001985 | Grik3 | glutamate receptor, ionotropic, kainate 3 | -3.67 |
| ENSMUSG00000029445 | Hpd | 4-hydroxyphenylpyruvic acid dioxygenase | -3.68 |
| ENSMUSG00000033200 | Tpsg1 | tryptase gamma 1 | -3.69 |
| ENSMUSG00000056220 | Pla2g4a | phospholipase A2, group IVA (cytosolic, calcium-dependent) | -3.69 |
| ENSMUSG00000101581 | C430002N11Rik | RIKEN cDNA C430002N11 gene | -3.69 |
| ENSMUSG00000085451 | Gm40614 | predicted gene, 40614 | -3.69 |
| ENSMUSG00000070332 | Trim80 | tripartite motif-containing 80 | -3.69 |
| ENSMUSG00000030737 | Slco2b1 | solute carrier organic anion transporter family, member 2b1 | -3.70 |
| ENSMUSG00000113800 | 2210039B01Rik | RIKEN cDNA 2210039B01 gene | -3.70 |
| ENSMUSG00000019890 | Nts | neurotensin | -3.71 |
| ENSMUSG00000107456 | Gm10400 | predicted gene 10400 | -3.71 |
| ENSMUSG00000060070 | Defa26 | defensin, alpha, 26 | -3.72 |
| ENSMUSG00000058216 | Gstp3 | glutathione S-transferase pi 3 | -3.73 |
| ENSMUSG00000026815 | Gfi1b | growth factor independent 1B | -3.74 |
| ENSMUSG00000058740 | Kcnt1 | potassium channel, subfamily T, member 1 | -3.75 |
| ENSMUSG00000095026 | Gm3336 | predicted gene 3336 | -3.76 |
| ENSMUSG00000089948 | Far2os1 | fatty acyl CoA reductase 2, opposite strand 1 | -3.76 |
| ENSMUSG00000055748 | Gsdmc4 | gasdermin C4 | -3.77 |
| ENSMUSG00000079180 | Mptx2 | mucosal pentraxin 2 | -3.77 |
| ENSMUSG00000098426 | Gm27149 | predicted gene 27149 | -3.77 |
| ENSMUSG00000022148 | Fyb | FYN binding protein | -3.78 |
| ENSMUSG00000028415 | Spink4 | serine peptidase inhibitor, Kazal type 4 | -3.78 |
| ENSMUSG00000026544 | Dusp23 | dual specificity phosphatase 23 | -3.79 |
| ENSMUSG00000034107 | Ano7 | anoctamin 7 | -3.80 |
| ENSMUSG00000036242 | Armh4 | armadillo-like helical domain containing 4 | -3.80 |
| ENSMUSG00000055827 | Gsdmc3 | gasdermin C3 | -3.81 |
| ENSMUSG00000082956 | Naip3 | NLR family, apoptosis inhibitory protein 3 | -3.82 |
| ENSMUSG00000047517 | Dmbt1 | deleted in malignant brain tumors 1 | -3.84 |
| ENSMUSG00000085651 | Gm11695 | predicted gene 11695 | -3.84 |
| ENSMUSG00000031891 | Hsd11b2 | hydroxysteroid 11-beta dehydrogenase 2 | -3.85 |
| ENSMUSG00000037953 | A4gnt | alpha-1,4-N-acetylglucosaminyltransferase | -3.85 |
| ENSMUSG00000110101 | Gm6249 | predicted gene 6249 | -3.85 |
| ENSMUSG00000073043 | Atoh1 | atonal bHLH transcription factor 1 | -3.86 |
| ENSMUSG00000026288 | Inpp5d | inositol polyphosphate-5-phosphatase D | -3.87 |
| ENSMUSG00000040121 | Rep15 | RAB15 effector protein | -3.88 |
| ENSMUSG00000104291 | A130071D04Rik | RIKEN cDNA A130071D04 gene | -3.88 |
| ENSMUSG00000084174 | Sycn | syncollin | -3.88 |
| ENSMUSG00000102615 | Gm37844 | predicted gene, 37844 | -3.89 |
| ENSMUSG00000039264 | Gimap3 | GTPase, IMAP family member 3 | -3.90 |
| ENSMUSG00000111377 | Gm40634 | predicted gene, 40634 | -3.90 |
| ENSMUSG00000046460 | Sh2d7 | SH2 domain containing 7 | -3.91 |
| ENSMUSG00000050321 | Neto1 | neuropilin (NRP) and tolloid (TLL)-like 1 | -3.91 |
| ENSMUSG00000041476 | Smpx | small muscle protein, X-linked | -3.91 |
| ENSMUSG00000009246 | Trpm5 | transient receptor potential cation channel, subfamily M, member 5 | -3.92 |
| ENSMUSG00000004933 | Matk | megakaryocyte-associated tyrosine kinase | -3.92 |
| ENSMUSG00000111340 | Gm47171 | predicted gene, 47171 | -3.92 |
| ENSMUSG00000026452 | Syt2 | synaptotagmin II | -3.92 |
| ENSMUSG00000027762 | Sucnr1 | succinate receptor 1 | -3.93 |
| ENSMUSG00000056553 | Ptprn2 | protein tyrosine phosphatase, receptor type, N polypeptide 2 | -3.94 |
| ENSMUSG00000063903 | Klk1 | kallikrein 1 | -3.94 |
| ENSMUSG00000110405 | Gm45534 | predicted gene 45534 | -3.95 |
| ENSMUSG00000020581 | Agr2 | anterior gradient 2 | -3.96 |
| ENSMUSG00000036027 | 1810046K07Rik | RIKEN cDNA 1810046K07 gene | -3.96 |
| ENSMUSG00000020788 | Atp2a3 | ATPase, Ca++ transporting, ubiquitous | -3.97 |
| ENSMUSG00000026832 | Cytip | cytohesin 1 interacting protein | -4.00 |
| ENSMUSG00000040478 | Prdm13 | PR domain containing 13 | -4.00 |
| ENSMUSG00000034472 | Rasd2 | RASD family, member 2 | -4.02 |
| ENSMUSG00000022686 | B3gnt5 | UDP-GlcNAc:betaGal beta-1,3-N-acetylglucosaminyltransferase 5 | -4.02 |
| ENSMUSG00000058260 | Serpina9 | serine (or cysteine) peptidase inhibitor, clade A (alpha-1 antiproteinase, antitrypsin), member 9 | -4.04 |
| ENSMUSG00000034701 | Neurod1 | neurogenic differentiation 1 | -4.05 |
| ENSMUSG00000024553 | Galr1 | galanin receptor 1 | -4.05 |
| ENSMUSG00000037106 | Fer1l6 | fer-1-like 6 (C. elegans) | -4.06 |
| ENSMUSG00000040061 | Plcb2 | phospholipase C, beta 2 | -4.06 |
| ENSMUSG00000054422 | Fabp1 | fatty acid binding protein 1, liver | -4.07 |
| ENSMUSG00000096146 | Kcnj11 | potassium inwardly rectifying channel, subfamily J, member 11 | -4.07 |
| ENSMUSG00000031377 | Bmx | BMX non-receptor tyrosine kinase | -4.08 |
| ENSMUSG00000030263 | Lrmp | lymphoid-restricted membrane protein | -4.09 |
| ENSMUSG00000039372 | 38047 | membrane-associated ring finger (C3HC4) 4 | -4.09 |

| | | | |
|---|---|---|---|
| ENSMUSG00000034353 | Ramp1 | receptor (calcitonin) activity modifying protein 1 | -4.10 |
| ENSMUSG00000044156 | Hepacam2 | HEPACAM family member 2 | -4.15 |
| ENSMUSG00000084325 | Gm15550 | predicted gene 15550 | -4.16 |
| ENSMUSG00000047730 | Fcgbp | Fc fragment of IgG binding protein | -4.17 |
| ENSMUSG00000001827 | Folr1 | folate receptor 1 (adult) | -4.17 |
| ENSMUSG00000006586 | Runx1t1 | runt-related transcription factor 1; translocated to, 1 (cyclin D-related) | -4.17 |
| ENSMUSG00000087801 | Gm25980 | predicted gene, 25980 | -4.19 |
| ENSMUSG00000023122 | Sult1c2 | sulfotransferase family, cytosolic, 1C, member 2 | -4.19 |
| ENSMUSG00000039629 | Strip2 | striatin interacting protein 2 | -4.20 |
| ENSMUSG00000032561 | Acpp | acid phosphatase, prostate | -4.20 |
| ENSMUSG00000061171 | Slc38a11 | solute carrier family 38, member 11 | -4.20 |
| ENSMUSG00000004952 | Rasa4 | RAS p21 protein activator 4 | -4.21 |
| ENSMUSG00000029335 | Bmp3 | bone morphogenetic protein 3 | -4.21 |
| ENSMUSG00000030207 | Fam234b | family with sequence similarity 234, member B | -4.21 |
| ENSMUSG00000042096 | Dao | D-amino acid oxidase | -4.21 |
| ENSMUSG00000028186 | Uox | urate oxidase | -4.21 |
| ENSMUSG00000078964 | Ces1b | carboxylesterase 1B | -4.21 |
| ENSMUSG00000026828 | Galnt5 | polypeptide N-acetylgalactosaminyltransferase 5 | -4.22 |
| ENSMUSG00000087518 | Gm13561 | predicted gene 13561 | -4.22 |
| ENSMUSG00000047564 | Krtap3-1 | keratin associated protein 3-1 | -4.23 |
| ENSMUSG00000043673 | Kcns3 | potassium voltage-gated channel, delayed-rectifier, subfamily S, member 3 | -4.23 |
| ENSMUSG00000051079 | Rgs13 | regulator of G-protein signaling 13 | -4.24 |
| ENSMUSG00000021069 | Pygl | liver glycogen phosphorylase | -4.24 |
| ENSMUSG00000058618 | AY761184 | cDNA sequence AY761184 | -4.24 |
| ENSMUSG00000017007 | Rbpjl | recombination signal binding protein for immunoglobulin kappa J region-like | -4.24 |
| ENSMUSG00000038528 | Mfsd4b5 | major facilitator superfamily domain containing 4B5 | -4.24 |
| ENSMUSG00000064272 | Gpbar1 | G protein-coupled bile acid receptor 1 | -4.25 |
| ENSMUSG00000049350 | Zg16 | zymogen granule protein 16 | -4.27 |
| ENSMUSG00000024029 | Tff3 | trefoil factor 3, intestinal | -4.28 |
| ENSMUSG00000061959 | Ces1e | carboxylesterase 1E | -4.28 |
| ENSMUSG00000026395 | Ptprc | protein tyrosine phosphatase, receptor type, C | -4.29 |
| ENSMUSG00000056032 | BC018473 | cDNA sequence BC018473 | -4.29 |
| ENSMUSG00000018776 | Slc35g3 | solute carrier family 35, member G3 | -4.29 |
| ENSMUSG00000032036 | Kirrel3 | kirre like nephrin family adhesion molecule 3 | -4.31 |
| ENSMUSG00000051606 | 2010001K21Rik | RIKEN cDNA 2010001K21 gene | -4.32 |
| ENSMUSG00000044453 | Ffar1 | free fatty acid receptor 1 | -4.32 |
| ENSMUSG00000079242 | C730034F03Rik | RIKEN cDNA C730034F03 gene | -4.32 |
| ENSMUSG00000071489 | Ptgdr | prostaglandin D receptor | -4.32 |
| ENSMUSG00000035930 | Chst4 | carbohydrate (chondroitin 6/keratan) sulfotransferase 4 | -4.33 |
| ENSMUSG00000006313 | Upk1a | uroplakin 1A | -4.33 |
| ENSMUSG00000082211 | Defa27 | defensin, alpha, 27 | -4.34 |
| ENSMUSG00000045826 | Ptprcap | protein tyrosine phosphatase, receptor type, C polypeptide-associated protein | -4.34 |
| ENSMUSG00000090174 | Gm10612 | predicted gene 10612 | -4.34 |
| ENSMUSG00000031173 | Otc | ornithine transcarbamylase | -4.35 |
| ENSMUSG00000046314 | Stxbp6 | syntaxin binding protein 6 (amisyn) | -4.35 |
| ENSMUSG00000105302 | Gm19817 | predicted gene, 19817 | -4.37 |
| ENSMUSG00000078137 | Ankrd63 | ankyrin repeat domain 63 | -4.37 |
| ENSMUSG00000033220 | Rac2 | Rac family small GTPase 2 | -4.38 |
| ENSMUSG00000025515 | Muc2 | mucin 2 | -4.40 |
| ENSMUSG00000052631 | Sh2d6 | SH2 domain containing 6 | -4.41 |
| ENSMUSG00000084384 | Gm12251 | predicted gene 12251 | -4.41 |
| ENSMUSG00000049036 | Tmem121 | transmembrane protein 121 | -4.46 |
| ENSMUSG00000056468 | 5730596B20Rik | RIKEN cDNA 5730596B20 gene | -4.47 |
| ENSMUSG00000095649 | Gm8979 | predicted gene 8979 | -4.48 |
| ENSMUSG00000109244 | Gm44751 | predicted gene 44751 | -4.48 |
| ENSMUSG00000000049 | Apoh | apolipoprotein H | -4.48 |
| ENSMUSG00000101031 | Ms4a12 | membrane-spanning 4-domains, subfamily A, member 12 | -4.49 |
| ENSMUSG00000019429 | Ffar3 | free fatty acid receptor 3 | -4.51 |
| ENSMUSG00000025991 | Cps1 | carbamoyl-phosphate synthetase 1 | -4.53 |
| ENSMUSG00000037973 | Ccdc129 | coiled-coil domain containing 129 | -4.54 |
| ENSMUSG00000029865 | Sval1 | seminal vesicle antigen-like 1 | -4.54 |
| ENSMUSG00000032087 | Dscaml1 | DS cell adhesion molecule like 1 | -4.54 |
| ENSMUSG00000003283 | Hck | hemopoietic cell kinase | -4.55 |
| ENSMUSG00000064140 | Trim38 | tripartite motif-containing 38 | -4.55 |
| ENSMUSG00000028255 | Clca1 | chloride channel accessory 1 | -4.56 |
| ENSMUSG00000047798 | Cd300lf | CD300 molecule like family member F | -4.57 |
| ENSMUSG00000100022 | Gm29590 | predicted gene 29590 | -4.58 |
| ENSMUSG00000044359 | P2ry4 | pyrimidinergic receptor P2Y, G-protein coupled, 4 | -4.60 |
| ENSMUSG00000079588 | Tmem182 | transmembrane protein 182 | -4.61 |
| ENSMUSG00000049122 | Frmd3 | FERM domain containing 3 | -4.62 |
| ENSMUSG00000068231 | Vmn1r43 | vomeronasal 1 receptor 43 | -4.62 |
| ENSMUSG00000025701 | Alox5 | arachidonate 5-lipoxygenase | -4.63 |
| ENSMUSG00000040148 | Hmx3 | H6 homeobox 3 | -4.64 |
| ENSMUSG00000040136 | Abcc8 | ATP-binding cassette, sub-family C (CFTR/MRP), member 8 | -4.64 |
| ENSMUSG00000044505 | Lingo4 | leucine rich repeat and Ig domain containing 4 | -4.64 |
| ENSMUSG00000018927 | Ccl6 | chemokine (C-C motif) ligand 6 | -4.65 |

| | | | |
|---|---|---|---|
| ENSMUSG00000034645 | Zyg11a | zyg-11 family member A, cell cycle regulator | -4.67 |
| ENSMUSG00000026163 | Sphkap | SPHK1 interactor, AKAP domain containing | -4.68 |
| ENSMUSG00000040298 | Btbd16 | BTB (POZ) domain containing 16 | -4.68 |
| ENSMUSG00000027220 | Syt13 | synaptotagmin XIII | -4.71 |
| ENSMUSG00000089669 | Tnfsf13 | tumor necrosis factor (ligand) superfamily, member 13 | -4.71 |
| ENSMUSG00000027296 | Itpka | inositol 1,4,5-trisphosphate 3-kinase A | -4.72 |
| ENSMUSG00000052955 | Cpvl | carboxypeptidase, vitellogenic-like | -4.75 |
| ENSMUSG00000071178 | Serpina1b | serine (or cysteine) preptidase inhibitor, clade A, member 1B | -4.77 |
| ENSMUSG00000028339 | Col15a1 | collagen, type XV, alpha 1 | -4.78 |
| ENSMUSG00000031981 | Capn9 | calpain 9 | -4.79 |
| ENSMUSG00000066176 | Gm12511 | predicted gene 12511 | -4.79 |
| ENSMUSG00000027168 | Pax6 | paired box 6 | -4.83 |
| ENSMUSG00000039013 | Siglecf | sialic acid binding Ig-like lectin F | -4.84 |
| ENSMUSG00000022489 | Pde1b | phosphodiesterase 1B, Ca2+-calmodulin dependent | -4.85 |
| ENSMUSG00000097266 | Gm26802 | predicted gene, 26802 | -4.86 |
| ENSMUSG00000090066 | 1110002E22Rik | RIKEN cDNA 1110002E22 gene | -4.92 |
| ENSMUSG00000086513 | 9130208D14Rik | RIKEN cDNA 9130208D14 gene | -4.94 |
| ENSMUSG00000070704 | Ugt2b36 | UDP glucuronosyltransferase 2 family, polypeptide B36 | -4.95 |
| ENSMUSG00000085095 | Gm15635 | predicted gene 15635 | -4.95 |
| ENSMUSG00000108950 | 9130015G15Rik | RIKEN cDNA 9130015G15 gene | -4.96 |
| ENSMUSG00000029797 | Sspo | SCO-spondin | -4.97 |
| ENSMUSG00000028777 | Gnat3 | guanine nucleotide binding protein, alpha transducing 3 | -5.00 |
| ENSMUSG00000040345 | Arhgap9 | Rho GTPase activating protein 9 | -5.02 |
| ENSMUSG00000042010 | Acacb | acetyl-Coenzyme A carboxylase beta | -5.04 |
| ENSMUSG00000022026 | Olfm4 | olfactomedin 4 | -5.08 |
| ENSMUSG00000037627 | Rgs22 | regulator of G-protein signalling 22 | -5.12 |
| ENSMUSG00000110195 | Pde2a | phosphodiesterase 2A, cGMP-stimulated | -5.14 |
| ENSMUSG00000040035 | Disp2 | dispatched RND tramsporter family member 2 | -5.19 |
| ENSMUSG00000109685 | Gm45912 | predicted gene 45912 | -5.20 |
| ENSMUSG00000050711 | Scg2 | secretogranin II | -5.24 |
| ENSMUSG00000034762 | Glis1 | GLIS family zinc finger 1 | -5.34 |
| ENSMUSG00000034923 | Ly6g6f | lymphocyte antigen 6 complex, locus G6F | -5.37 |
| ENSMUSG00000034115 | Scn11a | sodium channel, voltage-gated, type XI, alpha | -5.39 |
| ENSMUSG00000032826 | Ank2 | ankyrin 2, brain | -5.43 |
| ENSMUSG00000057400 | Ces1c | carboxylesterase 1C | -5.49 |
| ENSMUSG00000040680 | Kremen2 | kringle containing transmembrane protein 2 | -5.61 |
| ENSMUSG00000055235 | Wdr86 | WD repeat domain 86 | -5.61 |
| ENSMUSG00000051497 | Kcnj16 | potassium inwardly-rectifying channel, subfamily J, member 16 | -5.62 |
| ENSMUSG00000096630 | Vmn2r26 | vomeronasal 2, receptor 26 | -5.63 |
| ENSMUSG00000021919 | Chat | choline acetyltransferase | -5.68 |
| ENSMUSG00000107624 | Gm44005 | predicted gene, 44005 | -5.68 |
| ENSMUSG00000090254 | Gm1965 | predicted gene 1965 | -5.70 |
| ENSMUSG00000021070 | Bdkrb2 | bradykinin receptor, beta 2 | -5.70 |
| ENSMUSG00000035277 | Arx | aristaless related homeobox | -5.75 |
| ENSMUSG00000028996 | Rbp7 | retinol binding protein 7, cellular | -5.83 |
| ENSMUSG00000012819 | Cdh23 | cadherin 23 (otocadherin) | -5.88 |
| ENSMUSG00000106354 | Gm42607 | predicted gene 42607 | -5.89 |
| ENSMUSG00000020787 | P2rx1 | purinergic receptor P2X, ligand-gated ion channel, 1 | -5.98 |
| ENSMUSG00000002100 | Mybpc3 | myosin binding protein C, cardiac | -5.99 |
| ENSMUSG00000022650 | Retnlb | resistin like beta | -6.04 |
| ENSMUSG00000029878 | Dbpht2 | DNA binding protein with his-thr domain | -6.04 |
| ENSMUSG00000033569 | Adgrb3 | adhesion G protein-coupled receptor B3 | -6.12 |
| ENSMUSG00000046318 | Ccbe1 | collagen and calcium binding EGF domains 1 | -6.22 |
| ENSMUSG00000046215 | Rprml | reprimo-like | -6.27 |
| ENSMUSG00000040380 | Cbln3 | cerebellin 3 precursor protein | -6.31 |
| ENSMUSG00000050824 | Sstr5 | somatostatin receptor 5 | -6.44 |
| ENSMUSG00000115816 | Gm34589 | predicted gene, 34589 | -6.45 |
| ENSMUSG00000102759 | Gm10463 | predicted gene 10463 | -6.62 |
| ENSMUSG00000109237 | 9130214F15Rik | RIKEN cDNA 9130214F15 gene | -6.99 |
| ENSMUSG00000026531 | Mptx1 | mucosal pentraxin 1 | -7.01 |
| ENSMUSG00000024366 | Gfra3 | glial cell line derived neurotrophic factor family receptor alpha 3 | -7.18 |
| ENSMUSG00000032532 | Cck | cholecystokinin | -7.26 |
| ENSMUSG00000021536 | Adcy2 | adenylate cyclase 2 | -7.91 |
| ENSMUSG00000053852 | Adgrg4 | adhesion G protein-coupled receptor G4 | -8.21 |
| ENSMUSG00000026938 | Fcna | ficolin A | -8.30 |
| ENSMUSG00000047497 | Adamts12 | a disintegrin-like and metallopeptidase (reprolysin type) with thrombospondin type 1 motif, 12 | -8.61 |
| ENSMUSG00000000394 | Gcg | glucagon | -8.91 |
| ENSMUSG00000014351 | Gip | gastric inhibitory polypeptide | -9.22 |
| ENSMUSG00000026532 | Spta1 | spectrin alpha, erythrocytic 1 | -9.57 |

**Tom+Trop2- vs. differentiated cells**

| Ensemble gene code | Symbol | Gene name | logFC |
|---|---|---|---|
| ENSMUSG00000031841 | Cdh13 | cadherin 13 | 8.43 |
| ENSMUSG00000035615 | Frmpd1 | FERM and PDZ domain containing 1 | 7.13 |
| ENSMUSG00000020388 | Pdlim4 | PDZ and LIM domain 4 | 6.99 |
| ENSMUSG00000013415 | Igf2bp1 | insulin-like growth factor 2 mRNA binding protein 1 | 6.96 |
| ENSMUSG00000037035 | Inhbb | inhibin beta-B | 6.67 |
| ENSMUSG00000042268 | Slc26a9 | solute carrier family 26, member 9 | 6.58 |
| ENSMUSG00000040852 | Plekhh2 | pleckstrin homology domain containing, family H (with MyTH4 domain) member 2 | 6.55 |
| ENSMUSG00000079330 | Lemd1 | LEM domain containing 1 | 6.42 |
| ENSMUSG00000029368 | Alb | albumin | 6.38 |
| ENSMUSG00000031871 | Cdh5 | cadherin 5 | 6.23 |
| ENSMUSG00000025330 | Padi4 | peptidyl arginine deiminase, type IV | 6.19 |
| ENSMUSG00000028031 | Dkk2 | dickkopf WNT signaling pathway inhibitor 2 | 6.14 |
| ENSMUSG00000055322 | Tns1 | tensin 1 | 6.13 |
| ENSMUSG00000017491 | Rarb | retinoic acid receptor, beta | 6.12 |
| ENSMUSG00000053613 | Notumos | notum palmitoleoyl-protein carboxylesterase, opposite strand | 6.11 |
| ENSMUSG00000029371 | Cxcl5 | chemokine (C-X-C motif) ligand 5 | 6.04 |
| ENSMUSG00000023885 | Thbs2 | thrombospondin 2 | 6.02 |
| ENSMUSG00000029838 | Ptn | pleiotrophin | 6.02 |
| ENSMUSG00000031503 | Col4a2 | collagen, type IV, alpha 2 | 6.02 |
| ENSMUSG00000032502 | Stac | src homology three (SH3) and cysteine rich domain | 5.95 |
| ENSMUSG00000097651 | 4930461G14Rik | RIKEN cDNA 4930461G14 gene | 5.82 |
| ENSMUSG00000012428 | Steap4 | STEAP family member 4 | 5.74 |
| ENSMUSG00000022883 | Robo1 | roundabout guidance receptor 1 | 5.70 |
| ENSMUSG00000029223 | Uchl1 | ubiquitin carboxy-terminal hydrolase L1 | 5.68 |
| ENSMUSG00000022754 | Tmem45a | transmembrane protein 45a | 5.55 |
| ENSMUSG00000097767 | Miat | myocardial infarction associated transcript (non-protein coding) | 5.48 |
| ENSMUSG00000068699 | Flnc | filamin C, gamma | 5.35 |
| ENSMUSG00000054263 | Lifr | leukemia inhibitory factor receptor | 5.34 |
| ENSMUSG00000024268 | Celf4 | CUGBP, Elav-like family member 4 | 5.27 |
| ENSMUSG00000036480 | Prss56 | protease, serine 56 | 5.25 |
| ENSMUSG00000028360 | Slc44a5 | solute carrier family 44, member 5 | 5.22 |
| ENSMUSG00000059898 | Dsc3 | desmocollin 3 | 5.16 |
| ENSMUSG00000026167 | Wnt10a | wingless-type MMTV integration site family, member 10A | 5.14 |
| ENSMUSG00000110290 | Gm45336 | predicted gene 45336 | 5.06 |
| ENSMUSG00000021091 | Serpina3n | serine (or cysteine) peptidase inhibitor, clade A, member 3N | 4.94 |
| ENSMUSG00000029869 | Ephb6 | Eph receptor B6 | 4.92 |
| ENSMUSG00000033377 | Palmd | palmdelphin | 4.84 |
| ENSMUSG00000110187 | Gm45496 | predicted gene 45496 | 4.74 |
| ENSMUSG00000051159 | Cited1 | Cbp/p300-interacting transactivator with Glu/Asp-rich carboxy-terminal domain 1 | 4.72 |
| ENSMUSG00000022464 | Slc38a4 | solute carrier family 38, member 4 | 4.69 |
| ENSMUSG00000052516 | Robo2 | roundabout guidance receptor 2 | 4.64 |
| ENSMUSG00000038415 | Foxq1 | forkhead box Q1 | 4.59 |
| ENSMUSG00000032965 | Ift57 | intraflagellar transport 57 | 4.44 |
| ENSMUSG00000026117 | Zap70 | zeta-chain (TCR) associated protein kinase | 4.41 |
| ENSMUSG00000060962 | Dmkn | dermokine | 4.32 |
| ENSMUSG00000016763 | Scube1 | signal peptide, CUB domain, EGF-like 1 | 4.28 |
| ENSMUSG00000005503 | Evx1 | even-skipped homeobox 1 | 4.22 |
| ENSMUSG00000042988 | Notum | notum palmitoleoyl-protein carboxylesterase | 4.16 |
| ENSMUSG00000032643 | Fhl3 | four and a half LIM domains 3 | 4.16 |
| ENSMUSG00000024810 | Il33 | interleukin 33 | 4.15 |
| ENSMUSG00000021950 | Anxa8 | annexin A8 | 4.12 |
| ENSMUSG00000108897 | Gm44861 | predicted gene 44861 | 4.11 |
| ENSMUSG00000039232 | Stx11 | syntaxin 11 | 4.04 |
| ENSMUSG00000031661 | Nkd1 | naked cuticle 1 | 4.03 |
| ENSMUSG00000054555 | Adam12 | a disintegrin and metallopeptidase domain 12 (meltrin alpha) | 4.03 |
| ENSMUSG00000004552 | Ctse | cathepsin E | 3.97 |
| ENSMUSG00000030772 | Dkk3 | dickkopf WNT signaling pathway inhibitor 3 | 3.96 |
| ENSMUSG00000071531 | Gprin2 | G protein regulated inducer of neurite outgrowth 2 | 3.96 |
| ENSMUSG00000067818 | Myl9 | myosin, light polypeptide 9, regulatory | 3.93 |
| ENSMUSG00000037712 | Fermt2 | fermitin family member 2 | 3.89 |
| ENSMUSG00000026950 | Neb | nebulin | 3.87 |
| ENSMUSG00000040488 | Ltbp4 | latent transforming growth factor beta binding protein 4 | 3.83 |
| ENSMUSG00000066607 | Insyn1 | inhibitory synaptic factor 1 | 3.79 |
| ENSMUSG00000051236 | Msrb3 | methionine sulfoxide reductase B3 | 3.78 |
| ENSMUSG00000032492 | Pth1r | parathyroid hormone 1 receptor | 3.73 |
| ENSMUSG00000045991 | Onecut2 | one cut domain, family member 2 | 3.72 |
| ENSMUSG00000000031 | H19 | H19, imprinted maternally expressed transcript | 3.71 |
| ENSMUSG00000035164 | Zc3h12c | zinc finger CCCH type containing 12C | 3.69 |

| | | | |
|---|---|---|---|
| ENSMUSG00000030911 | Zp2 | zona pellucida glycoprotein 2 | 3.69 |
| ENSMUSG00000048078 | Tenm4 | teneurin transmembrane protein 4 | 3.67 |
| ENSMUSG00000041046 | Ramp3 | receptor (calcitonin) activity modifying protein 3 | 3.61 |
| ENSMUSG00000022176 | Rem2 | rad and gem related GTP binding protein 2 | 3.57 |
| ENSMUSG00000038860 | Garnl3 | GTPase activating RANGAP domain-like 3 | 3.57 |
| ENSMUSG00000004668 | Abca13 | ATP-binding cassette, sub-family A (ABC1), member 13 | 3.54 |
| ENSMUSG00000034460 | Six4 | sine oculis-related homeobox 4 | 3.53 |
| ENSMUSG00000022037 | Clu | clusterin | 3.48 |
| ENSMUSG00000049866 | Arl4c | ADP-ribosylation factor-like 4C | 3.48 |
| ENSMUSG00000040836 | Gpr161 | G protein-coupled receptor 161 | 3.44 |
| ENSMUSG00000063531 | Sema3e | sema domain, immunoglobulin domain (Ig), short basic domain, secreted, (semaphorin) 3E | 3.41 |
| ENSMUSG00000013584 | Aldh1a2 | aldehyde dehydrogenase family 1, subfamily A2 | 3.38 |
| ENSMUSG00000062327 | T | brachyury, T-box transcription factor T | 3.36 |
| ENSMUSG00000037016 | Frem2 | Fras1 related extracellular matrix protein 2 | 3.33 |
| ENSMUSG00000034684 | Sema3f | sema domain, immunoglobulin domain (Ig), short basic domain, secreted, (semaphorin) 3F | 3.26 |
| ENSMUSG00000030048 | Gkn3 | gastrokine 3 | 3.26 |
| ENSMUSG00000038146 | Notch3 | notch 3 | 3.25 |
| ENSMUSG00000078235 | Fam43b | family with sequence similarity 43, member B | 3.25 |
| ENSMUSG00000048450 | Msx1 | msh homeobox 1 | 3.16 |
| ENSMUSG00000026579 | F5 | coagulation factor V | 3.13 |
| ENSMUSG00000029769 | Ccdc136 | coiled-coil domain containing 136 | 3.12 |
| ENSMUSG00000006205 | Htra1 | HtrA serine peptidase 1 | 3.11 |
| ENSMUSG00000062591 | Tubb4a | tubulin, beta 4A class IVA | 3.09 |
| ENSMUSG00000027570 | Col9a3 | collagen, type IX, alpha 3 | 3.09 |
| ENSMUSG00000039239 | Tgfb2 | transforming growth factor, beta 2 | 3.08 |
| ENSMUSG00000027985 | Lef1 | lymphoid enhancer binding factor 1 | 3.06 |
| ENSMUSG00000043903 | Zfp469 | zinc finger protein 469 | 3.02 |
| ENSMUSG00000052957 | Gas1 | growth arrest specific 1 | 3.02 |
| ENSMUSG00000058070 | Eml1 | echinoderm microtubule associated protein like 1 | 2.98 |
| ENSMUSG00000031595 | Pdgfrl | platelet-derived growth factor receptor-like | 2.96 |
| ENSMUSG00000036036 | Zfp57 | zinc finger protein 57 | 2.95 |
| ENSMUSG00000028497 | Hacd4 | 3-hydroxyacyl-CoA dehydratase 4 | 2.94 |
| ENSMUSG00000061048 | Cdh3 | cadherin 3 | 2.94 |
| ENSMUSG00000086484 | Nron | non-protein coding RNA, repressor of NFAT | 2.91 |
| ENSMUSG00000020218 | Wif1 | Wnt inhibitory factor 1 | 2.89 |
| ENSMUSG00000025352 | Gdf11 | growth differentiation factor 11 | 2.87 |
| ENSMUSG00000000983 | Wfdc18 | WAP four-disulfide core domain 18 | 2.85 |
| ENSMUSG00000048960 | Prex2 | phosphatidylinositol-3,4,5-trisphosphate-dependent Rac exchange factor 2 | 2.81 |
| ENSMUSG00000031074 | Fgf3 | fibroblast growth factor 3 | 2.80 |
| ENSMUSG00000038775 | Vill | villin-like | 2.78 |
| ENSMUSG00000032717 | Mdfi | MyoD family inhibitor | 2.78 |
| ENSMUSG00000019997 | Ctgf | connective tissue growth factor | 2.77 |
| ENSMUSG00000059854 | Hydin | HYDIN, axonemal central pair apparatus protein | 2.75 |
| ENSMUSG00000059991 | Nptx2 | neuronal pentraxin 2 | 2.74 |
| ENSMUSG00000043924 | Ncmap | noncompact myelin associated protein | 2.72 |
| ENSMUSG00000026193 | Fn1 | fibronectin 1 | 2.70 |
| ENSMUSG00000055653 | Gpc3 | glypican 3 | 2.70 |
| ENSMUSG00000032186 | Tmod2 | tropomodulin 2 | 2.68 |
| ENSMUSG00000031502 | Col4a1 | collagen, type IV, alpha 1 | 2.67 |
| ENSMUSG00000023008 | Fmnl3 | formin-like 3 | 2.66 |
| ENSMUSG00000022371 | Col14a1 | collagen, type XIV, alpha 1 | 2.64 |
| ENSMUSG00000063605 | Ccdc102a | coiled-coil domain containing 102A | 2.64 |
| ENSMUSG00000017897 | Eya2 | EYA transcriptional coactivator and phosphatase 2 | 2.63 |
| ENSMUSG00000074813 | Gm14005 | predicted gene 14005 | 2.63 |
| ENSMUSG00000074796 | Slc4a11 | solute carrier family 4, sodium bicarbonate transporter-like, member 11 | 2.62 |
| ENSMUSG00000021539 | Lect2 | leukocyte cell-derived chemotaxin 2 | 2.61 |
| ENSMUSG00000062184 | Hs6st2 | heparan sulfate 6-O-sulfotransferase 2 | 2.56 |
| ENSMUSG00000045932 | Ifit2 | interferon-induced protein with tetratricopeptide repeats 2 | 2.51 |
| ENSMUSG00000026822 | Lcn2 | lipocalin 2 | 2.50 |
| ENSMUSG00000029674 | Limk1 | LIM-domain containing, protein kinase | 2.50 |
| ENSMUSG00000049848 | Ceacam19 | carcinoembryonic antigen-related cell adhesion molecule 19 | 2.50 |
| ENSMUSG00000032625 | Thsd7a | thrombospondin, type I, domain containing 7A | 2.49 |
| ENSMUSG00000088609 | Gm24187 | predicted gene, 24187 | 2.47 |
| ENSMUSG00000060548 | Tnfrsf19 | tumor necrosis factor receptor superfamily, member 19 | 2.46 |
| ENSMUSG00000070867 | Trabd2b | TraB domain containing 2B | 2.46 |
| ENSMUSG00000020844 | Nxn | nucleoredoxin | 2.44 |
| ENSMUSG00000038295 | Atg9b | autophagy related 9B | 2.43 |
| ENSMUSG00000063727 | Tnfrsf11b | tumor necrosis factor receptor superfamily, member 11b (osteoprotegerin) | 2.42 |
| ENSMUSG00000025938 | Slco5a1 | solute carrier organic anion transporter family, member 5A1 | 2.42 |
| ENSMUSG00000026981 | Il1rn | interleukin 1 receptor antagonist | 2.39 |
| ENSMUSG00000068859 | Sp9 | trans-acting transcription factor 9 | 2.39 |

| | | | |
|---|---|---|---|
| ENSMUSG00000052105 | Mtcl1 | microtubule crosslinking factor 1 | 2.38 |
| ENSMUSG00000055945 | Prr18 | proline rich 18 | 2.38 |
| ENSMUSG00000058297 | Spock2 | sparc/osteonectin, cwcv and kazal-like domains proteoglycan 2 | 2.37 |
| ENSMUSG00000040998 | Npnt | nephronectin | 2.33 |
| ENSMUSG00000109245 | Gm44860 | predicted gene 44860 | 2.33 |
| ENSMUSG00000059336 | Slc14a1 | solute carrier family 14 (urea transporter), member 1 | 2.31 |
| ENSMUSG00000058952 | Cfi | complement component factor i | 2.28 |
| ENSMUSG00000040663 | Clcf1 | cardiotrophin-like cytokine factor 1 | 2.23 |
| ENSMUSG00000020937 | Plcd3 | phospholipase C, delta 3 | 2.19 |
| ENSMUSG00000040415 | Dtx3 | deltex 3, E3 ubiquitin ligase | 2.19 |
| ENSMUSG00000024659 | Anxa1 | annexin A1 | 2.15 |
| ENSMUSG00000027068 | Dhrs9 | dehydrogenase/reductase (SDR family) member 9 | 2.13 |
| ENSMUSG00000046402 | Rbp1 | retinol binding protein 1, cellular | 2.12 |
| ENSMUSG00000028364 | Tnc | tenascin C | 2.11 |
| ENSMUSG00000025492 | Ifitm3 | interferon induced transmembrane protein 3 | 2.10 |
| ENSMUSG00000085903 | Gm15340 | predicted gene 15340 | 2.10 |
| ENSMUSG00000032875 | Arhgef17 | Rho guanine nucleotide exchange factor (GEF) 17 | 2.09 |
| ENSMUSG00000030796 | Tead2 | TEA domain family member 2 | 2.08 |
| ENSMUSG00000040808 | S100g | S100 calcium binding protein G | 2.08 |
| ENSMUSG00000016942 | Tmprss6 | transmembrane serine protease 6 | 2.07 |
| ENSMUSG00000078161 | Erich3 | glutamate rich 3 | 2.06 |
| ENSMUSG00000015647 | Lama5 | laminin, alpha 5 | 2.02 |
| ENSMUSG00000048191 | Muc6 | mucin 6, gastric | 2.01 |
| ENSMUSG00000030223 | Ptpro | protein tyrosine phosphatase, receptor type, O | 2.01 |
| ENSMUSG00000040430 | Pitpnc1 | phosphatidylinositol transfer protein, cytoplasmic 1 | 2.01 |
| ENSMUSG00000020086 | H2afy2 | H2A histone family, member Y2 | 1.98 |
| ENSMUSG00000025491 | Ifitm1 | interferon induced transmembrane protein 1 | 1.97 |
| ENSMUSG00000032076 | Cadm1 | cell adhesion molecule 1 | 1.95 |
| ENSMUSG00000050989 | Selenon | selenoprotein N | 1.94 |
| ENSMUSG00000036123 | Slc9a3 | solute carrier family 9 (sodium/hydrogen exchanger), member 3 | 1.93 |
| ENSMUSG00000038578 | Susd1 | sushi domain containing 1 | 1.92 |
| ENSMUSG00000010175 | Prox1 | prospero homeobox 1 | 1.90 |
| ENSMUSG00000032068 | Plet1 | placenta expressed transcript 1 | 1.90 |
| ENSMUSG00000039607 | Rbms3 | RNA binding motif, single stranded interacting protein | 1.89 |
| ENSMUSG00000026888 | Grb14 | growth factor receptor bound protein 14 | 1.89 |
| ENSMUSG00000036902 | Neto2 | neuropilin (NRP) and tolloid (TLL)-like 2 | 1.87 |
| ENSMUSG00000051343 | Rab11fip5 | RAB11 family interacting protein 5 (class I) | 1.87 |
| ENSMUSG00000014599 | Csf1 | colony stimulating factor 1 (macrophage) | 1.86 |
| ENSMUSG00000060181 | Slc35e3 | solute carrier family 35, member E3 | 1.85 |
| ENSMUSG00000040118 | Cacna2d1 | calcium channel, voltage-dependent, alpha2/delta subunit 1 | 1.81 |
| ENSMUSG00000020646 | Mboat2 | membrane bound O-acyltransferase domain containing 2 | 1.79 |
| ENSMUSG00000025089 | Gfra1 | glial cell line derived neurotrophic factor family receptor alpha 1 | 1.79 |
| ENSMUSG00000026728 | Vim | vimentin | 1.78 |
| ENSMUSG00000068923 | Syt11 | synaptotagmin XI | 1.78 |
| ENSMUSG00000008398 | Elk3 | ELK3, member of ETS oncogene family | 1.76 |
| ENSMUSG00000026676 | Ccdc3 | coiled-coil domain containing 3 | 1.76 |
| ENSMUSG00000071847 | Apcdd1 | adenomatosis polyposis coli down-regulated 1 | 1.75 |
| ENSMUSG00000041801 | Phlda3 | pleckstrin homology like domain, family A, member 3 | 1.75 |
| ENSMUSG00000040187 | Arntl2 | aryl hydrocarbon receptor nuclear translocator-like 2 | 1.74 |
| ENSMUSG00000060429 | Sntb1 | syntrophin, basic 1 | 1.73 |
| ENSMUSG00000013089 | Etv5 | ets variant 5 | 1.71 |
| ENSMUSG00000017466 | Timp2 | tissue inhibitor of metalloproteinase 2 | 1.70 |
| ENSMUSG00000017493 | Igfbp4 | insulin-like growth factor binding protein 4 | 1.69 |
| ENSMUSG00000026509 | Capn2 | calpain 2 | 1.69 |
| ENSMUSG00000049823 | Zbtb12 | zinc finger and BTB domain containing 12 | 1.69 |
| ENSMUSG00000043439 | Epop | elongin BC and polycomb repressive complex 2 associated protein | 1.67 |
| ENSMUSG00000062661 | Ncs1 | neuronal calcium sensor 1 | 1.67 |
| ENSMUSG00000021822 | Plau | plasminogen activator, urokinase | 1.67 |
| ENSMUSG00000037071 | Scd1 | stearoyl-Coenzyme A desaturase 1 | 1.65 |
| ENSMUSG00000072437 | Nanos1 | nanos C2HC-type zinc finger 1 | 1.65 |
| ENSMUSG00000049807 | Arhgap23 | Rho GTPase activating protein 23 | 1.64 |
| ENSMUSG00000081016 | Olfr1397-ps1 | olfactory receptor 1397, pseudogene 1 | 1.63 |
| ENSMUSG00000014592 | Camta1 | calmodulin binding transcription activator 1 | 1.62 |
| ENSMUSG00000034220 | Gpc1 | glypican 1 | 1.60 |
| ENSMUSG00000022146 | Osmr | oncostatin M receptor | 1.60 |
| ENSMUSG00000037379 | Spon2 | spondin 2, extracellular matrix protein | 1.58 |
| ENSMUSG00000036667 | Tcaf1 | TRPM8 channel-associated factor 1 | 1.57 |
| ENSMUSG00000037664 | Cdkn1c | cyclin-dependent kinase inhibitor 1C (P57) | 1.57 |
| ENSMUSG00000001521 | Tulp3 | tubby-like protein 3 | 1.57 |
| ENSMUSG00000025479 | Cyp2e1 | cytochrome P450, family 2, subfamily e, polypeptide 1 | 1.57 |
| ENSMUSG00000081683 | Fzd10 | frizzled class receptor 10 | 1.56 |

| | | | |
|---|---|---|---|
| ENSMUSG00000063522 | 2010109I03Rik | RIKEN cDNA 2010109I03 gene | 1.54 |
| ENSMUSG00000067889 | Sptbn2 | spectrin beta, non-erythrocytic 2 | 1.52 |
| ENSMUSG00000028654 | Mycl | v-myc avian myelocytomatosis viral oncogene lung carcinoma derived | 1.52 |
| ENSMUSG00000055333 | Fat2 | FAT atypical cadherin 2 | 1.52 |
| ENSMUSG00000039109 | F13a1 | coagulation factor XIII, A1 subunit | 1.51 |
| ENSMUSG00000027999 | Pla2g12a | phospholipase A2, group XIIA | 1.50 |
| ENSMUSG00000030409 | Dmpk | dystrophia myotonica-protein kinase | 1.49 |
| ENSMUSG00000078949 | R3hdml | R3H domain containing-like | 1.47 |
| ENSMUSG00000031328 | Flna | filamin, alpha | 1.46 |
| ENSMUSG00000085329 | 2810404F17Rik | RIKEN cDNA 2810404F17 gene | 1.46 |
| ENSMUSG00000058908 | Pla2g2a | phospholipase A2, group IIA (platelets, synovial fluid) | 1.45 |
| ENSMUSG00000037095 | Lrg1 | leucine-rich alpha-2-glycoprotein 1 | 1.45 |
| ENSMUSG00000029359 | Tesc | tescalcin | 1.43 |
| ENSMUSG00000000782 | Tcf7 | transcription factor 7, T cell specific | 1.41 |
| ENSMUSG00000013846 | St3gal1 | ST3 beta-galactoside alpha-2,3-sialyltransferase 1 | 1.41 |
| ENSMUSG00000033149 | Phldb2 | pleckstrin homology like domain, family B, member 2 | 1.40 |
| ENSMUSG00000032698 | Lmo2 | LIM domain only 2 | 1.40 |
| ENSMUSG00000027698 | Nceh1 | neutral cholesterol ester hydrolase 1 | 1.39 |
| ENSMUSG00000025902 | Sox17 | SRY (sex determining region Y)-box 17 | 1.39 |
| ENSMUSG00000048776 | Pthlh | parathyroid hormone-like peptide | 1.37 |
| ENSMUSG00000048058 | Ldlrad3 | low density lipoprotein receptor class A domain containing 3 | 1.34 |
| ENSMUSG00000026620 | Mark1 | MAP/microtubule affinity regulating kinase 1 | 1.34 |
| ENSMUSG00000028613 | Lrp8 | low density lipoprotein receptor-related protein 8, apolipoprotein e receptor | 1.33 |
| ENSMUSG00000039621 | Prex1 | phosphatidylinositol-3,4,5-trisphosphate-dependent Rac exchange factor 1 | 1.33 |
| ENSMUSG00000038151 | Prdm1 | PR domain containing 1, with ZNF domain | 1.32 |
| ENSMUSG00000032232 | Cgnl1 | cingulin-like 1 | 1.28 |
| ENSMUSG00000029832 | Nfe2l3 | nuclear factor, erythroid derived 2, like 3 | 1.28 |
| ENSMUSG00000038400 | Pmepa1 | prostate transmembrane protein, androgen induced 1 | 1.24 |
| ENSMUSG00000044033 | Ccdc141 | coiled-coil domain containing 141 | 1.24 |
| ENSMUSG00000017204 | Gsdma | gasdermin A | 1.23 |
| ENSMUSG00000008090 | Fgfrl1 | fibroblast growth factor receptor-like 1 | 1.20 |
| ENSMUSG00000060487 | Samd5 | sterile alpha motif domain containing 5 | 1.20 |
| ENSMUSG00000013275 | Slc41a1 | solute carrier family 41, member 1 | 1.19 |
| ENSMUSG00000014602 | Kif1a | kinesin family member 1A | 1.19 |
| ENSMUSG00000027257 | Pacsin3 | protein kinase C and casein kinase substrate in neurons 3 | 1.18 |
| ENSMUSG00000068547 | Clca4a | chloride channel accessory 4A | 1.17 |
| ENSMUSG00000025608 | Podxl | podocalyxin-like | 1.17 |
| ENSMUSG00000025075 | Habp2 | hyaluronic acid binding protein 2 | 1.16 |
| ENSMUSG00000038740 | Mvb12b | multivesicular body subunit 12B | 1.16 |
| ENSMUSG00000001473 | Tubb6 | tubulin, beta 6 class V | 1.16 |
| ENSMUSG00000035678 | Tnfsf9 | tumor necrosis factor (ligand) superfamily, member 9 | 1.15 |
| ENSMUSG00000021493 | Pdlim7 | PDZ and LIM domain 7 | 1.14 |
| ENSMUSG00000028268 | Gbp3 | guanylate binding protein 3 | 1.13 |
| ENSMUSG00000028836 | Slc30a2 | solute carrier family 30 (zinc transporter), member 2 | 1.10 |
| ENSMUSG00000045136 | Tubb2b | tubulin, beta 2B class IIB | 1.10 |
| ENSMUSG00000097494 | 4933406C10Rik | RIKEN cDNA 4933406C10 gene | 1.10 |
| ENSMUSG00000030352 | Tspan9 | tetraspanin 9 | 1.07 |
| ENSMUSG00000054793 | Cadm4 | cell adhesion molecule 4 | 1.03 |
| ENSMUSG00000050295 | Foxc1 | forkhead box C1 | 1.03 |
| ENSMUSG00000047945 | Marcksl1 | MARCKS-like 1 | 1.01 |
| ENSMUSG00000020186 | Csrp2 | cysteine and glycine-rich protein 2 | -1.06 |
| ENSMUSG00000095325 | Zfp870 | zinc finger protein 870 | -1.09 |
| ENSMUSG00000095649 | Gm8979 | predicted gene 8979 | -1.11 |
| ENSMUSG00000086513 | 9130208D14Rik | RIKEN cDNA 9130208D14 gene | -1.23 |
| ENSMUSG00000061958 | Gm14851 | predicted gene 14851 | -1.24 |
| ENSMUSG00000053846 | Lipg | lipase, endothelial | -1.48 |
| ENSMUSG00000066438 | Plekhd1 | pleckstrin homology domain containing, family D (with coiled-coil domains) member 1 | -1.51 |
| ENSMUSG00000109685 | Gm45912 | predicted gene 45912 | -1.68 |
| ENSMUSG00000026531 | Mptx1 | mucosal pentraxin 1 | -2.13 |
| ENSMUSG00000104918 | Gm42944 | predicted gene 42944 | -2.19 |
| ENSMUSG00000033209 | Ttc28 | tetratricopeptide repeat domain 28 | -2.83 |

#### Supplementary Table S7

Overlap of differentially expressed genes in $Trop2^{+}$ $RFP/tdTomato^{+}$ vs $Trop2^{-}$ $RFP/tdTomato^{+}$ neoplastic cells obtained in hyperplastic epithelium and adenomas 7 days and 6 weeks after Apc inactivation, respectively. Genes with adjusted p-value < 0.05 and $|\log_2 FC| \geq 1$ experiments were used for comparison.

**Upregulated genes**

| Ensemble gene code | Symbol | Gene name | logFC hyperplasia | logFC microadenoma |
|---|---|---|---|---|
| ENSMUSG00000051397 | Tacstd2 | tumor-associated calcium signal transducer 2 | 6.26 | 6.34 |
| ENSMUSG00000004951 | Hspb1 | heat shock protein 1 | 2.09 | 5.49 |
| ENSMUSG00000108218 | Olfr1372-ps1 | olfactory receptor 1372, pseudogene 1 | 2.38 | 5.08 |
| ENSMUSG00000053007 | Creb5 | cAMP responsive element binding protein 5 | 2.11 | 4.6 |
| ENSMUSG00000028031 | Dkk2 | dickkopf WNT signaling pathway inhibitor 2 | 2.02 | 4.56 |
| ENSMUSG00000029648 | Flt1 | FMS-like tyrosine kinase 1 | 3.69 | 4.52 |
| ENSMUSG00000062591 | Tubb4a | tubulin, beta 4A class IVA | 2.43 | 4.43 |
| ENSMUSG00000034220 | Gpc1 | glypican 1 | 1.95 | 4.43 |
| ENSMUSG00000025473 | Adam8 | a disintegrin and metallopeptidase domain 8 | 2.45 | 4.39 |
| ENSMUSG00000026167 | Wnt10a | wingless-type MMTV integration site family, member 10A | 1.85 | 4.34 |
| ENSMUSG00000022037 | Clu | clusterin | 4.57 | 4.32 |
| ENSMUSG00000046623 | Gjb4 | gap junction protein, beta 4 | 2.49 | 4.31 |
| ENSMUSG00000034463 | Scara3 | scavenger receptor class A, member 3 | 3.71 | 4.2 |
| ENSMUSG00000037188 | Grhl3 | grainyhead like transcription factor 3 | 3.54 | 4.12 |
| ENSMUSG00000018340 | Anxa6 | annexin A6 | 2.19 | 4.11 |
| ENSMUSG00000085412 | Halr1 | Hoxa adjacent long noncoding RNA 1 | 2.06 | 4.1 |
| ENSMUSG00000009097 | Tbx1 | T-box 1 | 1.73 | 4.09 |
| ENSMUSG00000027210 | Meis2 | Meis homeobox 2 | 1.71 | 4.05 |
| ENSMUSG00000031538 | Plat | plasminogen activator, tissue | 1.94 | 3.97 |
| ENSMUSG00000039232 | Stx11 | syntaxin 11 | 1.83 | 3.88 |
| ENSMUSG00000049807 | Arhgap23 | Rho GTPase activating protein 23 | 1.12 | 3.84 |
| ENSMUSG00000039153 | Runx2 | runt related transcription factor 2 | 2.93 | 3.79 |
| ENSMUSG00000043430 | Psapl1 | prosaposin-like 1 | 2.01 | 3.79 |
| ENSMUSG00000032496 | Ltf | lactotransferrin | 1.5 | 3.77 |
| ENSMUSG00000025330 | Padi4 | peptidyl arginine deiminase, type IV | 1.79 | 3.75 |
| ENSMUSG00000048960 | Prex2 | phosphatidylinositol-3,4,5-trisphosphate-dependent Rac exchange factor 2 | 1.58 | 3.73 |
| ENSMUSG00000022665 | Ccdc80 | coiled-coil domain containing 80 | 1.19 | 3.73 |
| ENSMUSG00000021765 | Fst | follistatin | 3.64 | 3.7 |
| ENSMUSG00000063727 | Tnfrsf11b | tumor necrosis factor receptor superfamily, member 11b (osteoprotegerin) | 2.48 | 3.61 |
| ENSMUSG00000041608 | Entpd3 | ectonucleoside triphosphate diphosphohydrolase 3 | 1.43 | 3.54 |
| ENSMUSG00000009628 | Tex15 | testis expressed gene 15 | 1.21 | 3.53 |
| ENSMUSG00000020651 | Slc26a4 | solute carrier family 26, member 4 | 2.35 | 3.47 |
| ENSMUSG00000029761 | Cald1 | caldesmon 1 | 1.72 | 3.46 |
| ENSMUSG00000041189 | Chrnb1 | cholinergic receptor, nicotinic, beta polypeptide 1 (muscle) | 1.17 | 3.46 |
| ENSMUSG00000034634 | Ly6d | lymphocyte antigen 6 complex, locus D | 2.18 | 3.43 |
| ENSMUSG00000031925 | Maml2 | mastermind like transcriptional coactivator 2 | 1.29 | 3.41 |
| ENSMUSG00000040998 | Npnt | nephronectin | 3.5 | 3.4 |
| ENSMUSG00000025938 | Slco5a1 | solute carrier organic anion transporter family, member 5A1 | 2.46 | 3.39 |
| ENSMUSG00000032186 | Tmod2 | tropomodulin 2 | 1.68 | 3.39 |
| ENSMUSG00000053137 | Mapk11 | mitogen-activated protein kinase 11 | 2.06 | 3.32 |
| ENSMUSG00000024912 | Fosl1 | fos-like antigen 1 | 1.21 | 3.27 |
| ENSMUSG00000004552 | Ctse | cathepsin E | 2.95 | 3.26 |
| ENSMUSG00000050640 | Tmem150c | transmembrane protein 150C | 2.7 | 3.26 |
| ENSMUSG00000013338 | Fer1l4 | fer-1-like 4 (C. elegans) | 2.77 | 3.23 |
| ENSMUSG00000055333 | Fat2 | FAT atypical cadherin 2 | 2.75 | 3.23 |
| ENSMUSG00000026620 | Mark1 | MAP/microtubule affinity regulating kinase 1 | 2.73 | 3.2 |
| ENSMUSG00000080316 | Spaca6 | sperm acrosome associated 6 | 2.15 | 3.18 |
| ENSMUSG00000031841 | Cdh13 | cadherin 13 | 4.27 | 3.16 |
| ENSMUSG00000038587 | Akap12 | A kinase (PRKA) anchor protein (gravin) 12 | 2.21 | 3.14 |
| ENSMUSG00000004044 | Cavin1 | caveolae associated 1 | 1.82 | 3.13 |
| ENSMUSG00000040152 | Thbs1 | thrombospondin 1 | 2.1 | 3.12 |
| ENSMUSG00000000983 | Wfdc18 | WAP four-disulfide core domain 18 | 1.95 | 3.12 |
| ENSMUSG00000020674 | Pxdn | peroxidasin | 1.08 | 3.11 |
| ENSMUSG00000027376 | Prom2 | prominin 2 | 3.33 | 3.09 |
| ENSMUSG00000034295 | Fhod3 | formin homology 2 domain containing 3 | 1.1 | 3.06 |
| ENSMUSG00000046449 | Nexmif | neurite extension and migration factor | 2.5 | 3.04 |
| ENSMUSG00000048078 | Tenm4 | teneurin transmembrane protein 4 | 1.51 | 3.04 |
| ENSMUSG00000049866 | Arl4c | ADP-ribosylation factor-like 4C | 1.01 | 3.03 |
| ENSMUSG00000031328 | Flna | filamin, alpha | 1.65 | 3.02 |
| ENSMUSG00000013076 | Amotl1 | angiomotin-like 1 | 1.26 | 2.98 |
| ENSMUSG00000026193 | Fn1 | fibronectin 1 | 2.46 | 2.95 |
| ENSMUSG00000046731 | Kctd11 | potassium channel tetramerisation domain containing 11 | 1.19 | 2.95 |
| ENSMUSG00000024659 | Anxa1 | annexin A1 | 4.59 | 2.93 |
| ENSMUSG00000037833 | Sh2d4b | SH2 domain containing 4B | 3.16 | 2.91 |
| ENSMUSG00000024940 | Ltbp3 | latent transforming growth factor beta binding protein 3 | 1.12 | 2.91 |
| ENSMUSG00000033227 | Wnt6 | wingless-type MMTV integration site family, member 6 | 2.59 | 2.9 |
| ENSMUSG00000023008 | Fmnl3 | formin-like 3 | 1.63 | 2.89 |
| ENSMUSG00000020181 | Nav3 | neuron navigator 3 | 1.27 | 2.89 |
| ENSMUSG00000029869 | Ephb6 | Eph receptor B6 | 2.67 | 2.88 |
| ENSMUSG00000032717 | Mdfi | MyoD family inhibitor | 1.35 | 2.88 |
| ENSMUSG00000034573 | Ptpn13 | protein tyrosine phosphatase, non-receptor type 13 | 3.29 | 2.86 |
| ENSMUSG00000067889 | Sptbn2 | spectrin beta, non-erythrocytic 2 | 1.98 | 2.85 |
| ENSMUSG00000031871 | Cdh5 | cadherin 5 | 1.94 | 2.83 |

| | | | | |
|---|---|---|---|---|
| ENSMUSG00000002980 | Bcam | basal cell adhesion molecule | 1.59 | 2.83 |
| ENSMUSG00000001288 | Rarg | retinoic acid receptor, gamma | 2.72 | 2.82 |
| ENSMUSG00000022176 | Rem2 | rad and gem related GTP binding protein 2 | 1.23 | 2.82 |
| ENSMUSG00000041390 | Mdfic | MyoD family inhibitor domain containing | 1.25 | 2.8 |
| ENSMUSG00000057193 | Slc44a2 | solute carrier family 44, member 2 | 1.01 | 2.8 |
| ENSMUSG00000035678 | Tnfsf9 | tumor necrosis factor (ligand) superfamily, member 9 | 1.47 | 2.78 |
| ENSMUSG00000040430 | Pitpnc1 | phosphatidylinositol transfer protein, cytoplasmic 1 | 1.91 | 2.77 |
| ENSMUSG00000069255 | Dusp22 | dual specificity phosphatase 22 | 1.16 | 2.73 |
| ENSMUSG00000038415 | Foxq1 | forkhead box Q1 | 1.54 | 2.7 |
| ENSMUSG00000032327 | Stra6 | stimulated by retinoic acid gene 6 | 3.35 | 2.68 |
| ENSMUSG00000020427 | Igfbp3 | insulin-like growth factor binding protein 3 | 1.57 | 2.68 |
| ENSMUSG00000032875 | Arhgef17 | Rho guanine nucleotide exchange factor (GEF) 17 | 1.36 | 2.67 |
| ENSMUSG00000033377 | Palmd | palmdelphin | 4.44 | 2.65 |
| ENSMUSG00000040488 | Ltbp4 | latent transforming growth factor beta binding protein 4 | 2.58 | 2.65 |
| ENSMUSG00000015134 | Aldh1a3 | aldehyde dehydrogenase family 1, subfamily A3 | 3.3 | 2.62 |
| ENSMUSG00000040836 | Gpr161 | G protein-coupled receptor 161 | 1.91 | 2.61 |
| ENSMUSG00000021493 | Pdlim7 | PDZ and LIM domain 7 | 1.72 | 2.6 |
| ENSMUSG00000061517 | Sox21 | SRY (sex determining region Y)-box 21 | 3.2 | 2.57 |
| ENSMUSG00000090394 | 4930523C07Rik | RIKEN cDNA 4930523C07 gene | 1.43 | 2.56 |
| ENSMUSG00000006411 | Nectin4 | nectin cell adhesion molecule 4 | 1.87 | 2.54 |
| ENSMUSG00000027257 | Pacsin3 | protein kinase C and casein kinase substrate in neurons 3 | 1.2 | 2.44 |
| ENSMUSG00000059668 | Krt4 | keratin 4 | 4.76 | 2.42 |
| ENSMUSG00000009687 | Fxyd5 | FXYD domain-containing ion transport regulator 5 | 2.59 | 2.42 |
| ENSMUSG00000008398 | Elk3 | ELK3, member of ETS oncogene family | 2.27 | 2.4 |
| ENSMUSG00000024401 | Tnf | tumor necrosis factor | 1.72 | 2.4 |
| ENSMUSG00000019853 | Hebp2 | heme binding protein 2 | 1.16 | 2.36 |
| ENSMUSG00000022371 | Col14a1 | collagen, type XIV, alpha 1 | 1.46 | 2.33 |
| ENSMUSG00000024810 | Il33 | interleukin 33 | 1.28 | 2.32 |
| ENSMUSG00000052105 | Mtcl1 | microtubule crosslinking factor 1 | 1.65 | 2.31 |
| ENSMUSG00000041570 | Camsap2 | calmodulin regulated spectrin-associated protein family, member 2 | 1.4 | 2.26 |
| ENSMUSG00000029832 | Nfe2l3 | nuclear factor, erythroid derived 2, like 3 | 1.08 | 2.26 |
| ENSMUSG00000082361 | Btc | betacellulin, epidermal growth factor family member | 1.28 | 2.2 |
| ENSMUSG00000001435 | Col18a1 | collagen, type XVIII, alpha 1 | 2.34 | 2.17 |
| ENSMUSG00000042306 | S100a14 | S100 calcium binding protein A14 | 1.89 | 2.15 |
| ENSMUSG00000031441 | Atp11a | ATPase, class VI, type 11A | 1.05 | 2.14 |
| ENSMUSG00000039457 | Ppl | periplakin | 1.7 | 2.13 |
| ENSMUSG00000048058 | Ldlrad3 | low density lipoprotein receptor class A domain containing 3 | 1.27 | 2.12 |
| ENSMUSG00000001473 | Tubb6 | tubulin, beta 6 class V | 1.1 | 2.12 |
| ENSMUSG00000032013 | Trim29 | tripartite motif-containing 29 | 1.96 | 2.11 |
| ENSMUSG00000032643 | Fhl3 | four and a half LIM domains 3 | 1.16 | 2.09 |
| ENSMUSG00000022754 | Tmem45a | transmembrane protein 45a | 2.56 | 2.08 |
| ENSMUSG00000023885 | Thbs2 | thrombospondin 2 | 1.59 | 2.08 |
| ENSMUSG00000017723 | Wfdc2 | WAP four-disulfide core domain 2 | 1.7 | 2.07 |
| ENSMUSG00000026405 | C4bp | complement component 4 binding protein | 2.24 | 2.05 |
| ENSMUSG00000110218 | Gm20219 | predicted gene, 20219 | 1.23 | 2.03 |
| ENSMUSG00000037946 | Fgd3 | FYVE, RhoGEF and PH domain containing 3 | 3.28 | 2.02 |
| ENSMUSG00000003541 | Ier3 | immediate early response 3 | 1.41 | 2.01 |
| ENSMUSG00000037060 | Cavin3 | caveolae associated 3 | 2.29 | 1.98 |
| ENSMUSG00000021477 | Ctsl | cathepsin L | 1.34 | 1.98 |
| ENSMUSG00000027381 | Bcl2l11 | BCL2-like 11 (apoptosis facilitator) | 1.2 | 1.97 |
| ENSMUSG00000038775 | Vill | villin-like | 1.89 | 1.94 |
| ENSMUSG00000030352 | Tspan9 | tetraspanin 9 | 1.23 | 1.94 |
| ENSMUSG00000040990 | Sh3kbp1 | SH3-domain kinase binding protein 1 | 2.67 | 1.92 |
| ENSMUSG00000007039 | Ddah2 | dimethylarginine dimethylaminohydrolase 2 | 2.05 | 1.92 |
| ENSMUSG00000022178 | Ajuba | ajuba LIM protein | 1.7 | 1.87 |
| ENSMUSG00000026478 | Lamc1 | laminin, gamma 1 | 1.01 | 1.87 |
| ENSMUSG00000025921 | Rdh10 | retinol dehydrogenase 10 (all-trans) | 1.83 | 1.83 |
| ENSMUSG00000062661 | Ncs1 | neuronal calcium sensor 1 | 1.63 | 1.82 |
| ENSMUSG00000026065 | Slc9a4 | solute carrier family 9 (sodium/hydrogen exchanger), member 4 | 3.37 | 1.79 |
| ENSMUSG00000027712 | Anxa5 | annexin A5 | 2.12 | 1.79 |
| ENSMUSG00000046589 | Lrrc8e | leucine rich repeat containing 8 family, member E | 1.54 | 1.79 |
| ENSMUSG00000027858 | Tspan2 | tetraspanin 2 | 1.76 | 1.78 |
| ENSMUSG00000027860 | Vangl1 | VANGL planar cell polarity 1 | 1.07 | 1.77 |
| ENSMUSG00000025372 | Baiap2 | brain-specific angiogenesis inhibitor 1-associated protein 2 | 1.34 | 1.76 |
| ENSMUSG00000047501 | Cldn4 | claudin 4 | 2.13 | 1.71 |
| ENSMUSG00000028464 | Tpm2 | tropomyosin 2, beta | 1.82 | 1.71 |
| ENSMUSG00000031430 | Vsig1 | V-set and immunoglobulin domain containing 1 | 2.76 | 1.7 |
| ENSMUSG00000022505 | Emp2 | epithelial membrane protein 2 | 1.97 | 1.69 |
| ENSMUSG00000027737 | Slc7a11 | solute carrier family 7 (cationic amino acid transporter, y+ system), member 11 | 1.61 | 1.68 |
| ENSMUSG00000038295 | Atg9b | autophagy related 9B | 1.06 | 1.66 |
| ENSMUSG00000055407 | Map6 | microtubule-associated protein 6 | 2.93 | 1.64 |
| ENSMUSG00000031586 | Rbpms | RNA binding protein gene with multiple splicing | 2.02 | 1.63 |
| ENSMUSG00000006777 | Krt23 | keratin 23 | 1.63 | 1.62 |
| ENSMUSG00000079330 | Lemd1 | LEM domain containing 1 | 2.11 | 1.6 |
| ENSMUSG00000027485 | Bpifb1 | BPI fold containing family B, member 1 | 6.53 | 1.57 |
| ENSMUSG00000006435 | Neurl1a | neuralized E3 ubiquitin protein ligase 1A | 1.09 | 1.55 |

| | | | | |
|---|---|---|---|---|
| ENSMUSG00000001025 | S100a6 | S100 calcium binding protein A6 (calcyclin) | 2.11 | 1.52 |
| ENSMUSG00000017897 | Eya2 | EYA transcriptional coactivator and phosphatase 2 | 1.52 | 1.51 |
| ENSMUSG00000069114 | Zbtb10 | zinc finger and BTB domain containing 10 | 1.02 | 1.49 |
| ENSMUSG00000027907 | S100a11 | S100 calcium binding protein A11 | 1.44 | 1.47 |
| ENSMUSG00000020773 | Trim47 | tripartite motif-containing 47 | 1.38 | 1.47 |
| ENSMUSG00000021108 | Prkch | protein kinase C, eta | 1.23 | 1.46 |
| ENSMUSG00000039384 | Dusp10 | dual specificity phosphatase 10 | 2.15 | 1.44 |
| ENSMUSG00000030759 | Far1 | fatty acyl CoA reductase 1 | 1.38 | 1.44 |
| ENSMUSG00000022055 | Nefl | neurofilament, light polypeptide | 2.01 | 1.39 |
| ENSMUSG00000041886 | Macc1 | metastasis associated in colon cancer 1 | 1.83 | 1.39 |
| ENSMUSG00000021792 | Fam213a | family with sequence similarity 213, member A | 2.72 | 1.38 |
| ENSMUSG00000032348 | Gsta4 | glutathione S-transferase, alpha 4 | 2.55 | 1.3 |
| ENSMUSG00000026335 | Pam | peptidylglycine alpha-amidating monooxygenase | 1.31 | 1.26 |
| ENSMUSG00000035547 | Capn5 | calpain 5 | 1.07 | 1.26 |
| ENSMUSG00000038463 | Olfml2b | olfactomedin-like 2B | 1.98 | 1.23 |
| ENSMUSG00000028378 | Ptgr1 | prostaglandin reductase 1 | 1.25 | 1.22 |
| ENSMUSG00000039831 | Arhgap29 | Rho GTPase activating protein 29 | 1.33 | 1.17 |
| ENSMUSG00000040711 | Sh3pxd2b | SH3 and PX domains 2B | 1.26 | 1.17 |
| ENSMUSG00000040543 | Pitpnm3 | PITPNM family member 3 | 1.45 | 1.16 |
| ENSMUSG00000038387 | Rras | related RAS viral (r-ras) oncogene | 1.16 | 1.12 |
| ENSMUSG00000089809 | Rasgef1b | RasGEF domain family, member 1B | 1.31 | 1.09 |
| ENSMUSG00000032068 | Plet1 | placenta expressed transcript 1 | 2.88 | 1.08 |
| ENSMUSG00000026796 | Fam129b | family with sequence similarity 129, member B | 1.01 | 1.06 |
| ENSMUSG00000026509 | Capn2 | calpain 2 | 2.17 | 1.05 |
| ENSMUSG00000024501 | Dpysl3 | dihydropyrimidinase-like 3 | 1.84 | 1.04 |
| ENSMUSG00000030342 | Cd9 | CD9 antigen | 1.03 | 1.04 |

**Downregulated genes**

| Ensemble gene code | Symbol | Gene name | logFC hyperplasia | logFC microadenoma |
|---|---|---|---|---|
| ENSMUSG00000022650 | Retnlb | resistin like beta | -1.33 | -5.1 |
| ENSMUSG00000022026 | Olfm4 | olfactomedin 4 | -2.76 | -4.12 |
| ENSMUSG00000051079 | Rgs13 | regulator of G-protein signaling 13 | -1.04 | -4.05 |
| ENSMUSG00000032561 | Acpp | acid phosphatase, prostate | -1.44 | -3.64 |
| ENSMUSG00000086513 | 9130208D14Rik | RIKEN cDNA 9130208D14 gene | -1.65 | -3.63 |
| ENSMUSG00000056553 | Ptprn2 | protein tyrosine phosphatase, receptor type, N polypeptide 2 | -1.25 | -3.52 |
| ENSMUSG00000025991 | Cps1 | carbamoyl-phosphate synthetase 1 | -1.6 | -3.48 |
| ENSMUSG00000109685 | Gm45912 | predicted gene 45912 | -1.85 | -3.37 |
| ENSMUSG00000031173 | Otc | ornithine transcarbamylase | -1.29 | -3.34 |
| ENSMUSG00000033740 | St18 | suppression of tumorigenicity 18 | -1.25 | -3.34 |
| ENSMUSG00000058618 | AY761184 | cDNA sequence AY761184 | -1.33 | -3.33 |
| ENSMUSG00000028415 | Spink4 | serine peptidase inhibitor, Kazal type 4 | -1.01 | -3.3 |
| ENSMUSG00000095649 | Gm8979 | predicted gene 8979 | -1.75 | -3.29 |
| ENSMUSG00000029445 | Hpd | 4-hydroxyphenylpyruvic acid dioxygenase | -1.76 | -3.2 |
| ENSMUSG00000031891 | Hsd11b2 | hydroxysteroid 11-beta dehydrogenase 2 | -1.04 | -3.17 |
| ENSMUSG00000030207 | Fam234b | family with sequence similarity 234, member B | -1.77 | -3.14 |
| ENSMUSG00000034472 | Rasd2 | RASD family, member 2 | -2.09 | -3.07 |
| ENSMUSG00000079180 | Mptx2 | mucosal pentraxin 2 | -1.27 | -3.04 |
| ENSMUSG00000037953 | A4gnt | alpha-1,4-N-acetylglucosaminyltransferase | -1.81 | -2.94 |
| ENSMUSG00000074715 | Ccl28 | chemokine (C-C motif) ligand 28 | -1.19 | -2.79 |
| ENSMUSG00000005268 | Prlr | prolactin receptor | -1.39 | -2.75 |
| ENSMUSG00000026610 | Esrrg | estrogen-related receptor gamma | -1.13 | -2.73 |
| ENSMUSG00000074437 | Defa29 | defensin, alpha, 29 | -1.39 | -2.68 |
| ENSMUSG00000033715 | Akr1c14 | aldo-keto reductase family 1, member C14 | -1.15 | -2.67 |
| ENSMUSG00000046613 | Vwa5b2 | von Willebrand factor A domain containing 5B2 | -1.06 | -2.67 |
| ENSMUSG00000052026 | Slc6a7 | solute carrier family 6 (neurotransmitter transporter, L-proline), member 7 | -1.16 | -2.63 |
| ENSMUSG00000036422 | Pcdh8 | protocadherin 8 | -2.6 | -2.57 |
| ENSMUSG00000027359 | Slc27a2 | solute carrier family 27 (fatty acid transporter), member 2 | -1.66 | -2.54 |
| ENSMUSG00000028327 | Stra6l | STRA6-like | -1.3 | -2.5 |
| ENSMUSG00000026398 | Nr5a2 | nuclear receptor subfamily 5, group A, member 2 | -1.21 | -2.44 |
| ENSMUSG00000038745 | Nlrp6 | NLR family, pyrin domain containing 6 | -1.16 | -2.38 |
| ENSMUSG00000031451 | Gas6 | growth arrest specific 6 | -1.16 | -2.38 |
| ENSMUSG00000040412 | 5330417C22Rik | RIKEN cDNA 5330417C22 gene | -1.25 | -2.35 |
| ENSMUSG00000079559 | Colca2 | COLCA2 homolog | -1.38 | -2.32 |
| ENSMUSG00000026786 | Apbb1ip | amyloid beta (A4) precursor protein-binding, family B, member 1 interacting protein | -1.56 | -2.31 |
| ENSMUSG00000109936 | Gm45889 | predicted gene 45889 | -1.25 | -2.31 |
| ENSMUSG00000025790 | Slco3a1 | solute carrier organic anion transporter family, member 3a1 | -1.32 | -2.3 |
| ENSMUSG00000024411 | Aqp4 | aquaporin 4 | -2 | -2.25 |
| ENSMUSG00000004655 | Aqp1 | aquaporin 1 | -1.36 | -2.24 |
| ENSMUSG00000025738 | Fbxl16 | F-box and leucine-rich repeat protein 16 | -1.02 | -2.24 |
| ENSMUSG00000066687 | Zbtb16 | zinc finger and BTB domain containing 16 | -1.89 | -2.19 |
| ENSMUSG00000060586 | H2-Eb1 | histocompatibility 2, class II antigen E beta | -1.04 | -2.16 |
| ENSMUSG00000026489 | Coq8a | coenzyme Q8A | -1.61 | -2.08 |
| ENSMUSG00000027452 | Acss1 | acyl-CoA synthetase short-chain family member 1 | -1.01 | -2.04 |
| ENSMUSG00000079547 | H2-DMb1 | histocompatibility 2, class II, locus Mb1 | -1 | -2.01 |
| ENSMUSG00000052271 | Bhlha15 | basic helix-loop-helix family, member a15 | -1.38 | -1.96 |
| ENSMUSG00000022018 | Rgcc | regulator of cell cycle | -1.32 | -1.89 |
| ENSMUSG00000024225 | Clps | colipase, pancreatic | -1.18 | -1.88 |
| ENSMUSG00000110266 | Gm32742 | predicted gene, 32742 | -2.31 | -1.85 |
| ENSMUSG00000035561 | Aldh1b1 | aldehyde dehydrogenase 1 family, member B1 | -1.61 | -1.83 |
| ENSMUSG00000060208 | Defa17 | defensin, alpha, 17 | -1.19 | -1.79 |
| ENSMUSG00000063142 | Kcnma1 | potassium large conductance calcium-activated channel, subfamily M, alpha member 1 | -1.08 | -1.78 |
| ENSMUSG00000022504 | Ciita | class II transactivator | -1.43 | -1.77 |
| ENSMUSG00000005089 | Slc1a2 | solute carrier family 1 (glial high affinity glutamate transporter), member 2 | -1.26 | -1.77 |
| ENSMUSG00000021830 | Txndc16 | thioredoxin domain containing 16 | -1.09 | -1.77 |
| ENSMUSG00000058740 | Kcnt1 | potassium channel, subfamily T, member 1 | -1.16 | -1.71 |
| ENSMUSG00000005360 | Slc1a3 | solute carrier family 1 (glial high affinity glutamate transporter), member 3 | -1.48 | -1.68 |
| ENSMUSG00000040528 | Milr1 | mast cell immunoglobulin like receptor 1 | -1.6 | -1.55 |
| ENSMUSG00000030934 | Oat | ornithine aminotransferase | -1.1 | -1.41 |
| ENSMUSG00000033721 | Vav3 | vav 3 oncogene | -1.34 | -1.38 |
| ENSMUSG00000031453 | Rasa3 | RAS p21 protein activator 3 | -1.03 | -1.31 |
| ENSMUSG00000003545 | Fosb | FBJ osteosarcoma oncogene B | -1.07 | -1.07 |
| ENSMUSG00000002346 | Slc25a42 | solute carrier family 25, member 42 | -1.21 | -1.03 |
| ENSMUSG00000074653 | Lrrc31 | leucine rich repeat containing 31 | -1.03 | -1.03 |
| ENSMUSG00000041577 | Prelp | proline arginine-rich end leucine-rich repeat | -2.09 | -1 |
